# Transcriptome profiling of human colonic cells exposed to the gut pathobiont *Streptococcus gallolyticus* subsp. *gallolyticus*

**DOI:** 10.1101/2023.05.16.540927

**Authors:** Pasquereau-Kotula Ewa, Laurence du Merle, Odile Sismeiro, Natalia Pietrosemoli, Hugo Varet, Rachel Legendre, Patrick Trieu-Cuot, Shaynoor Dramsi

## Abstract

*Streptococcus gallolyticus sp. gallolyticus (SGG)* is a gut pathobiont involved in the development of colorectal cancer (CRC). To decipher the contribution of *SGG* in tumor initiation and/or acceleration respectively, a global transcriptome was performed in normal colonic cells (FHC) and in tumoral colonic cells (HT29). To identify *SGG*-specific alterations, we chose the phylogenetically closest relative, *Streptococcus gallolyticus* subsp. *macedonicus* (*SGM)* as the control bacterium. We show that *SGM,* a bacterium generally considered as safe, did not induce any transcriptional changes on the two human colonic cells. The transcriptional reprogramming induced by *SGG* was significantly different in FHC and HT29 cells, with most of the up- and down-regulated genes associated with cancer disease. Top up-regulated genes related to cancer were: (i) *IL-20, CLK1, SORBS2, ERG1, PIM1, SNORD3A* for normal FHC cells and (ii) *TSLP, BHLHA15, LAMP3, ZNF27B, KRT17, ATF3* for cancerous HT29 cells. *SGG* induces much stronger transcriptional changes in cancerous than in normal colonic cells (2,090 *vs* 128 genes being affected, respectively). Gene set enrichment analysis reveals that *SGG*-induced strong ER- (endoplasmic reticulum) stress and UPR- (unfolded protein response) activation in colonic epithelial cells. Our results suggest that *SGG* induces a pro-tumoral shift in human colonic cells, particularly in transformed cells potentially accelerating tumor development in the colon.

## Introduction

Colorectal cancer (CRC) is the third leading malignancy worldwide and the second most common cause of cancer mortality, regardless of gender [1]. The majority of CRC cases are sporadic, occurring in people with no family history or genetic predisposition [2]. Environmental risk factors such as lifestyle, diet, smoking and gut microbiota contribute significantly to the development of sporadic CRCs [3]. *Streptococcus gallolyticus* was one of the first bacteria to be strongly associated with CRC by epidemiological studies [4–8]. In line with the literature, we showed that *SGG* prevalence in the French population (patients with normal colonoscopies) is about 32.5%, and is increased to 50% in the stools of patients with adenocarcinomas [9].

*Streptococcus gallolyticus* subsp. *gallolyticus* (*SGG)* (formerly known as *S. bovis* type I) is an extracellular opportunistic pathogen responsible for septicemia and endocarditis in the elderly. In the last decade, several groups have contributed in deciphering the mechanisms underlying *SGG* association with CRC [10–14].

*S. gallolyticus* has been subdivided into three subspecies, subsp. *gallolyticus* (*SGG*), subsp. *pasteurianus (SGP)* and subsp. *macedonicus (SGM)*. Among them, only *SGG* is associated with CRC, suggesting that *SGG*-specific attributes contribute to this association. *SGP* causes bacteremia, endocarditis, and urinary tract infection in elderly and immunodeficient people, septicemia and meningitis in newborns and intrauterine infections in pregnant woman [15–18]*. SGM*, the genetically closest *SGG* relative is generally considered as safe and is a non-pathogenic species [19,20]. *SGM* is a homofermentative lactic acid bacterium which was first isolated from a typical Greek cheese obtained by natural fermentation [21]. A couple of studies have reported that some *SGM* strains possess probiotic properties [22,23].

A recent study reported the transcription profiling of *in vitro* cultured HT29 cells infected with *SGG* for 4 h in which 44 genes were significantly up- (21 genes) or down-regulated (23 genes) [24]. Most up-regulated genes were involved in detoxification or bio-activation of toxic compounds [24], which might alter intestinal susceptibility to DNA damaging events and on long-term contribute to carcinogenesis.

We aimed to discover pathways specifically induced by *SGG* in the host colon which could contribute to tumor initiation or/and progression. To identify cancer-related genes or pathways activated *in vitro* upon infection with *SGG*, a global transcriptome analysis was performed in normal human colonic cells (FHC) and in transformed (HT29) cells after 24 h of co-culture. We also decided to compare gene expression profiles of normal vs cancerous colonic cell in response to *SGG* vs *SGM* to infer more robust conclusions. We reasoned that if *SGG* is an oncogenic bacterium involved in tumor initiation, we should see a specific transcriptomic signature on normal, non-transformed cells such as the FHC cell line (CRL-1831, ATCC) [25], which is an epithelial cell line isolated from the large intestine of a 13-week-old human embryo. The second possibility being that *SGG* acts as an accelerator of tumor development and thus will have a much stronger impact on pre-transformed cells such as the HT29 cell line (HTB-38, ATCC) [26] which is a human colorectal adenocarcinoma cell line with epithelial morphology.

We present here a comprehensive analysis of the transcriptome alterations induced by *SGG* infection in normal colonic cells and in tumoral colonic cells. The total of 2,090 genes were differentially altered in tumoral HT29 cells as compared to 128 genes in normal FHC cells, suggesting that *SGG* is rather a tumor accelerator than an initiator which fits well with our previous *in vivo* data [14].

## Materials and methods

### Bacterial strains and culture conditions

*SGG* UCN34 [27] and *SGM* strain CIP105683T [28] were grown at 37°C in Todd Hewitt Yeast (THY) broth in standing filled flasks or on THY agar plates (Difco Laboratories).

### Cell culture

The normal human normal colon epithelial cell lines, FHC (ATCC: CRL-1831 [25]) was cultured in DMEM/F12 medium (Gibco, France) supplemented with 20 % heat-inactivated calf serum and additional factors (25 mM HEPES; 10 ng/mL cholera toxin; 0.005 mg/mL insulin; 0.005 mg/mL transferrin; 100 ng/mL hydrocortisone; EFG 20 ng/mL; 10% SVF) to sustain their growth and could be passed 5-10 times only. The human cancerous cell line HT-29 (ATCC: HTB-38 [26]) was cultivated in DMEM with 10% heat-inactivated calf serum and supplemented with 25 mM HEPES. Cells were cultured in ventilated T75 flasks at 37 °C and 5 % CO_2_.

### Infection of epithelial cells with bacteria

FHC cells were seeded into 6-well plates at a density of 2 x 10^5^ cells per well and HT29 cells at 4 x 10^5^ cells per well and incubated for 16-20 hours. Stationary phase bacteria were scraped from fresh THY plates (overnight culture) and washed with sterile phosphate buffered saline, pH 7.4 (PBS) at a final OD_600_ of 1 (corresponding to 6.5 x 10^8^ CFU). Bacteria were diluted in DMEM (Gibco, Ref. 12320032, low glucose, pyruvate, HEPES) to obtain the following desired concentrations (i) *SGM* of 6.5 x 10^5^ CFU/ml and (ii) *SGG* UCN34 of 6.5 x 10^4^ CFU/ml. Cells were washed once with DMEM and then infected with the medium containing the bacteria by adding 2 ml of bacterial suspension per well in the 6-well plate. For the non-treated (NT) condition, only 2 ml of sterile fresh media was added. For each condition, a complete 6-well plate was used and then pooled together to obtain enough cellular material for further RNA extraction. Trimethoprim (Sigma, Ref. T7883), a bacteriostatic antibiotic was added at 50 μg.ml^-1^ final concentration after 6 h of co-culture to prevent media acidification due to bacterial growth. The total co-culture incubation time was 24 h. Total RNA was extracted from cell monolayers with the RNeasy Plus Mini kit (Qiagen, USA), according to the manufacturer’s instructions. Three independent replicates for each condition were performed and analyzed. RNAs were conserved at -80°C and used in transcriptome Illumina assay and quantitative RT-PCR.

### Transcriptome Illumina analysis

Total RNA from three independent replicates of each experimental condition: FHC NT (n=3), FHC *SGM* (n=3), FHC *SGG* UCN34 (n=3), HT29 NT (n=3), HT29 *SGM* (n=3), HT29 *SGG* UCN34 (n=3) were checked on RNA 6000 Nano chips (Bioanalyzer, Agilent) for quality and integrity. Directional libraries were prepared using the TruSeq Stranded mRNA Sample preparation kit following the manufacturer’s instructions (Illumina). Libraries were checked for quality on DNA 1000 chips (Bioanalyzer, Agilent). Quantification was performed with the fluorescent-based quantitation Qubit dsDNA HS Assay Kit (Thermo Fisher Scientific). Sequencing was performed as a Single Read Multiplexed run for 65 bp sequences on HiSeq 2500 sequencer (Illumina). The multiplexing level was 6 samples per lane. Reads were cleaned of adapter sequences and low-quality sequences using cutadapt version 1.11 [29]. Only sequences at least 25 nt in length were considered for further analysis. STAR version 2.5.0a [30], with default parameters, was used for alignment on the reference genome (Human genome hg38 from Ensembl). Genes were counted using featureCounts version 1.4.6-p3 [31] from Subreads package (parameters: -t exon -g gene_id -s 1). Count data were analyzed using R version 3.5.1 [32] and the Bioconductor package DESeq2 version 1.20.0 [33]. The normalization and dispersion estimation were performed with DESeq2 using the default parameters and statistical test for differential expression were performed applying the independent filtering algorithm. A generalized linear model including the replicate effect, the cell line, the infection status as well as the cell line - infection interaction was set up to test for the differential expression between the conditions. For each pairwise comparison, raw p-values were adjusted for multiple testing according to the Benjamini and Hochberg (BH) procedure [34] and genes with an adjusted p-value lower than 0.05 were considered differentially expressed.

Gene Set Enrichment analysis (GSEA) was performed using the Camera (competitive gene set test accounting for inter-gene correlation) [35] method from the limma R package (version 3.34.9). Functional annotation of the genes was obtained using the Hallmark gene sets, the KEGG and Reactome biological pathway and the Gene Ontology collections from the MSigDB database (GSEA, UC San Diego, CA, USA) [36]. GSEA was performed on the complete count matrix and gene sets with p-value ≤ 0.05 were considered statistically significant. Ingenuity Pathway Analysis (IPA) was performed using genes differentially expressed between *SGG* UCN34 and *SGM* to specifically identify diseases associated to those genes.

Each principal component analysis (PCA) is based on the variance-stabilized transformed count matrix that has been adjusted for the replicate effect using the removeBatchEffect function or the limma R package (version 3.52.4).

### Gene expression analysis

Total RNA was extracted from cell monolayers with the RNeasy mini kit (Qiagen, USA), according to the manufacturer’s instructions. First-strand cDNA synthesis was performed using the iScript cDNA synthesis kit (Biorad). To do so, a 20 μL solution was prepared with 1 μg of RNA, 4 μL of the 5X iScript reaction mix, 1 μL of the iScript Reverse Transcriptase, and completed with RNase free water. The reverse transcription program was as follows: 5 min at 25°C, 30 min at 42°C, 5 min at 85°C and 15 min at 15°C. cDNAs were diluted to the hundredth and stored at -20°C. Quantitative real-time PCR was performed using SsoFast EvaGreen Supermix (Bio-Rad) with EvaGreen as fluorescent dye according to the manufacturer’s protocol. Primers used for amplification of cDNA are specified in **Supp Table S1**. GAPDH quantification was used as internal control for normalization. Fold difference of mRNA levels were calculated using the ΔΔCt method. All PCR reactions were performed in duplicates and repeated three independent times.

### Statistical Analysis

Mann-Whitney nonparametric test was used to test the statistical significance of differences between the different group parameters. *p* values of less than 0.05 were considered statistically significant.

## Results

### Comparison of host transcriptomic responses in normal and tumoral colonic cells exposed to *SGG*

To unravel the effects of *SGG* on gene expression of human colonic cells, a global transcriptome was performed in normal FHC and tumoral HT29 cells following 24 h of co-culture with *SGG* UCN34 vs *SGM*. The experimental protocol is schematized in **Fig. 1A** and the differential analysis results of the transcriptomic profiles in **Fig. 1B**. Both infected conditions were compared to the control non-treated (NT) cells. The initial inoculum for *SGG* and *SGM* was calculated experimentally to have the same number of bacteria at the end of the experiment at 24 h post infection and thus *SGM* cell count was about 10-fold higher than that of *SGG* UCN34 (**Supp Fig. S1A**). As shown in **Supp Fig. S1B** and **C,** *SGG* adheres very well to normal FHC colonic cells and makes bacterial aggregates (white arrow). Of note, a very different adhesion pattern was observed in HT29 cells with most of the bacteria adhering to the cell-free surface covered with extracellular matrix secreted by the tumoral cells. A few *SGG* were found adhering and making aggregates partly on cells and partly on the cell culture surface (white arrow) (**Supp Fig. S1B, C**). *SGM* adheres well to both cell types (white arrow) but no bacterial aggregates were observed (**Supp Fig. S1B**). To ensure proper growth and health of normal colonic FHC cells, transcriptomics experiments were performed under sub-confluent conditions (50-60% of confluence). To properly compare adhesion of *SGG* and *SGM* to FHC and HT29 cells, assays were performed on 100% confluent monolayers showing that *SGM* adheres as well as *SGG* on both cell lines after 24 h of co-culture (**Supp Fig. S1D).**

**Figure 1.**
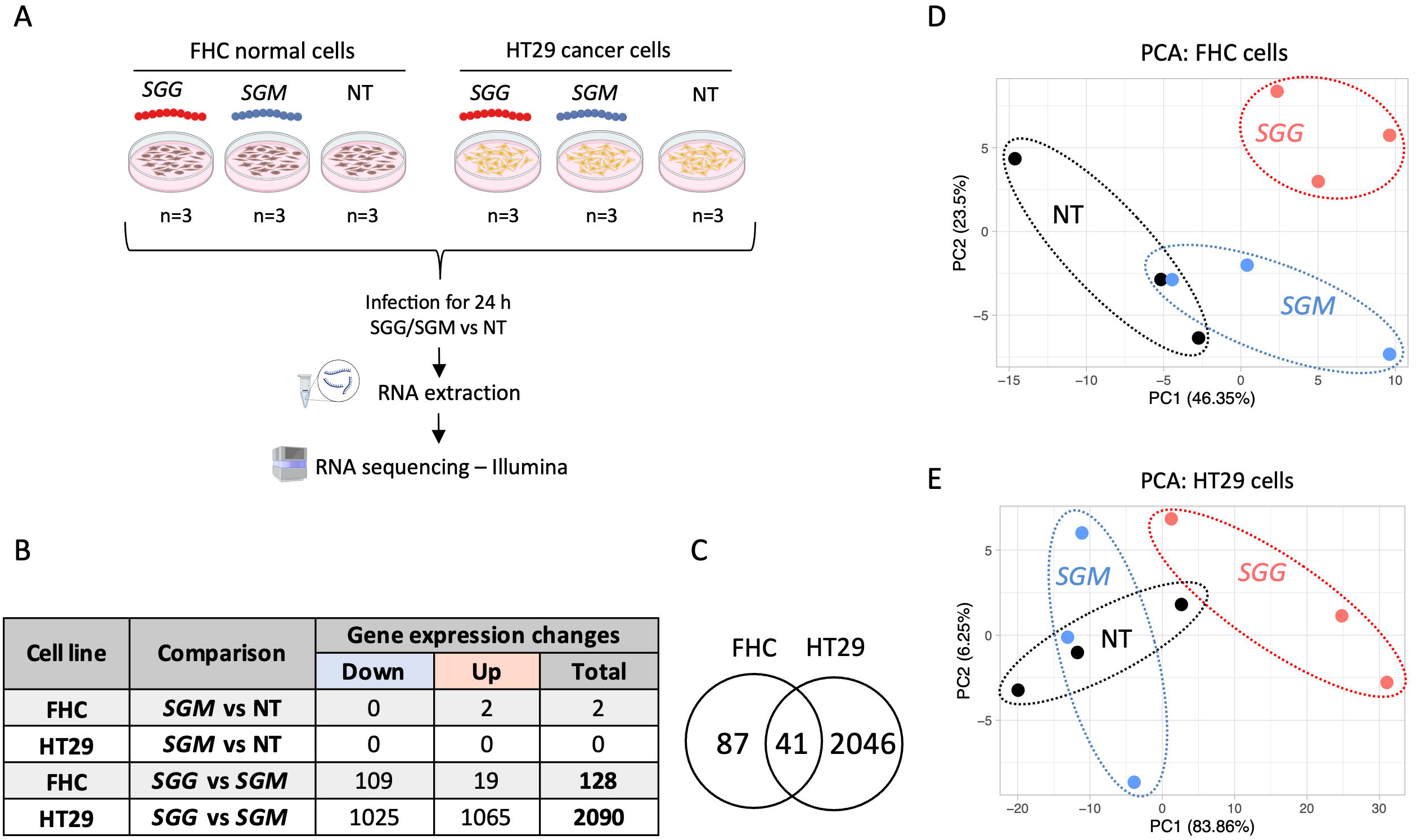
Comparison of host transcriptomic responses in normal FHC and tumoral HT29 colonic cells exposed to *SGG* vs *SGM.* **A.** Experimental design of cell-bacteria co-culture used for transcriptomic study (created with BioRender.com). Cells were seeded onto 6-wells plate (FHC: 2 x 10^5^ cells/well and HT29: 4 x 10^5^ cells/well) and incubated for 16-20 hours. Cells were infected with bacteria: *SGG* UCN34 (6.5 x 10^4^ CFU/ml) and *SGM* (6.5 x 10^5^ CFU/ml). Trimethoprim (50 μg /ml) was added to the wells after 6 hours of co-culture to prevent bacterial over-growth. The total infection time was 24 h. Total RNA was prepared, processed and sequenced using Illumina technology. This experiment was performed in triplicate (n=3). **B**. Table recapitulating the transcriptional changes in FHC and HT29 induced by *SGM* vs NT and *SGG* UCN34 vs *SGM* conditions. Criteria for significant gene changes were as follows: adjusted p < 0.05 and log2 fold change (FC) > 0.5 or < -0.5). **C**. Venn diagram showing that 41 genes regulated upon *SGG* UCN34 infection are common to FHC and HT29 cells. **D**. PCA of transcriptome samples in FHC cells: *SGG* (n=3); *SGM* (n=3) and NT (n=3). **E**. PCA of transcriptome samples in HT29 cells: *SGG* (n=3); *SGM* (n=3) and NT (n=3). D and E. Each PCA is based on the variance-stabilized transformed count matrix that has been adjusted for the replicate effect using the remove BatchEffect function or the limma R package (version 3.52.4).

Analysis of differentially expressed genes (adjusted p-value < 0.05 and log_2_ fold change (FC) > 0.5 or < -0.5) between cells co-cultured with *SGM* bacteria vs control NT condition revealed that *SGM* did not induce any transcriptional changes on HT29 and FHC cells. Only two genes were found dysregulated in FHC cells (**Fig. 1B**). These results indicate that *SGM* is an appropriate control bacterium to decipher the specific pathogenic traits of *SGG*. Next, we focused on comparing *SGG* vs *SGM* conditions in more details. We identified 128 differentially expressed genes (adjusted p-value < 0.05 and log_2_ fold change (FC) > 0.5 or < 0.5) in FHC cells with most of them being down-regulated (109 genes) and only 19 being up-regulated (**Fig. 1B; Supp. Table S2**). Interestingly, we identified 2,090 differentially expressed genes (1,025 down- and 1,065 up-regulated) in HT29 cells (**Fig. 1B; Supp. Table S3**). Only 41 genes were found to be common to both cell lines (2 up- and 39 down-regulated; **Fig. 1C**; **Supp. Table S4)**. It is worth noting that the 2 up-regulated genes were: (i) CDC like kinase 1 (*CLK1*) and (ii) DNA polymerase beta (*POLB*). *CLK1* plays a crucial role in the regulation of proliferation, invasion and migration in gastric cancer and was described as a novel therapeutic target [37]. *POLB* performs base excision repair (BER) in case of DNA damage and its levels are often unbalanced in CRC [38].

Principal component analysis (PCA) shows a clear separation of *SGG* from *SGM* and NT and less good separation in between *SGM* and NT groups in FHC (**Fig. 1D**) and HT29 cells (**Fig. 1E**), confirming the specific *SGG*-induced gene changes.

In general, our results show that *SGG* induces a specific transcriptomic signature dependent on the cell type (normal or cancerous) suggesting different consequences of *SGG* presence in healthy colon vs colon with neoplastic or pre-neoplastic lesions.

### In depth analysis of the transcriptomic responses of normal colonic cells

*SGG* altered the expression of 128 genes in normal colonic human FHC cells. The heat map shown in **Fig. 2A** displaying the six most up-regulated genes and two down-regulated genes clearly shows the existence of 2 groups: *SGG* on one hand and *SGM* and NT on the other hand. The 6 up-regulated genes (*IL-20, CLK1, SORBS2, ERG1, PIM1, SNORD3A*) are involved in cancer development [39–44]. Interleukin 20 (*IL-20*) is a cytokine assigned to the interleukin 10 family described as tumor promoting [45]. Sorbin and SH3 domain containing 2 (*SORBS2*) promotes colon cancer migration though activation of the Notch pathway [44]. Small nucleolar RNA, C/D box 3A (*SNORD3A*) was shown to be involved in doxorubicin resistance in human osteosarcoma cells by modulating multiple genes promoting proliferation, ribosome biogenesis, DNA damaging sensing, and DNA repair [46]. Early growth response 1 (*ERG1*) is implicated in the regulation of cell growth, proliferation, differentiation and apoptosis in CRC [47,43]. Proto-oncogene serine/threonine-protein kinase (*Pim-1*) promotes CRC growth and metastasis [48]. We confirmed the up-regulation of these 6 genes using q RT-PCR (**Fig. 2B**). Both techniques gave similar fold-changes between *SGG* and *SGM* (R= 0.97; **Fig. 2C**).

**Figure 2.**
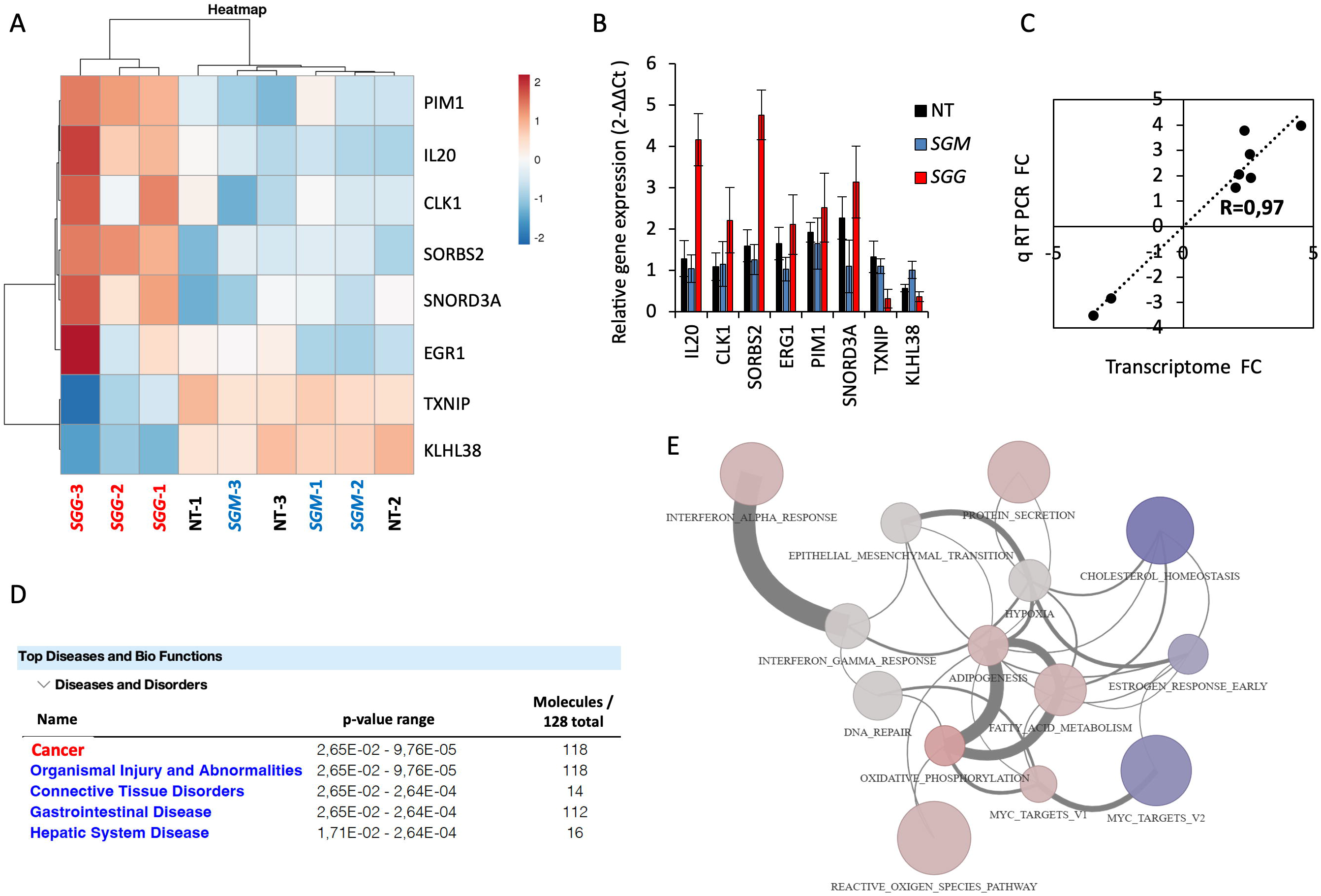
Transcriptional responses of human normal FHC cells upon *SGG* infection. **A.** Heat map of transcriptomic data displaying the top six up-regulated and two down-regulated genes. **B**. Relative quantification (2^-ΔΔCt^) of mRNA levels at 24 h of infection with *SGG*, *SGM* or non-treated (NT). **C**. Correlation of fold changes between transcriptome and quantitative real time PCR for the 8 genes shown in B. **D**. Top Disease and Bio Functions using Ingenuity Pathway Analysis (IPA) using genes differentially expressed between *SGG* UCN34 and *SGM*. Molecules indicate the number of genes associated with indicated diseases and disorders. A right-tailed Fisher’s Exact Test was used to calculate a p-value determining the probability that each biological function and/or disease assigned to that data set is due to chance alone. **E**. Network representation of the 14 significantly enriched hallmark gene sets upon *SGG* infection on FHC cells. Each node (circle) represents a gene set. Node size is proportional to the number of genes present in the dataset and belonging to the gene set. Node colors reflect the average expression directionality (based on the average of the_log fold expression for each gene belonging to the gene set) (red = over expressed; blue = down regulated; grey=no statistically significant difference in expression). The color intensity is proportional to the strength of the average expression. Each edge connecting two nodes means that two gene sets share the same genes. The edge width is proportional to the number of genes shared among the two gene sets. Edges are shown only if at least two genes are shared among the two gene sets (30 edges).

These results were analyzed using the Ingenuity Pathway Analysis (IPA, Qiagen) showing that most of the differentially expressed genes were involved in cancer disease (118 out of 128; **Fig. 2D**). Gene Set Enrichment Analysis (GSEA) with Camera showed 14 gene sets from the HALLMARK collection significantly changed: (i) oxidative phosphorylation; (ii) adipogenesis; (iii) reactive oxygen species; (iv) cholesterol homeostasis; (v) interferon alpha response; (vi) myc targets v1; (vii) protein secretion; (viii) fatty acid metabolism; (ix) epithelial to mesenchymal transition; (x) interferon gamma response; (xi) myc targets v2; (xii) hypoxia; (xiii) estrogen response early and (xiv) DNA repair (**Fig. 2E, Supp Table S7**). Most of these pathways (11) were predicted to be activated (in pink) and 3 to be down-regulated (blue) (**Fig. 2E**).

### In depth analysis of the transcriptomic responses of tumoral colonic cells

*SGG* altered the expression of as many as 2,090 genes in tumoral colonic human HT29 cells. The heat map shown in **Fig. 3A** displays the 10 most up-regulated genes and one down-regulated gene. Of note, the ratio changes in up- and down-regulated genes in tumoral HT29 cells was much greater than for normal FHC cells (≈10-fold). The most 10 up-regulated genes are: *TSLP, SPX, BHLHA15, GPR1, NGFR, ZNF27B, KLHDC7B, LAMP3, KRT17* and *ATF3.* The up-regulation of these 10 genes was confirmed independently using q RT-PCR (**Fig. 3B**). Both techniques gave similar fold changes between *SGG* and *SGM* (R= 0.77; **Fig. 3C**). Importantly, 6 of the 10 most up-regulated genes are involved in CRC development. Indeed, the high expression of thymic stromal lymphopoietin (*TSLP*) in cancer cells correlates with a poor prognosis for CRC patients [49]. Basic helix-loop-helix family member a15 (*BHLHA15*) is involved in CRC initiation through YAP/Wnt pathway activation [50]. Lysosomal associated membrane protein 3 (*LAMP3*) and zinc finger protein 274 (*ZNF27B*) are both up-regulated in CRC patients [51,52]. Keratin 17 (*KRT17*) is an oncogene in several types of cancer and is used as a biomarker of CRC [53]. Activating transcription factor 3 (*ATF3*) promotes tumor growth in CRC [54].

**Figure 3.**
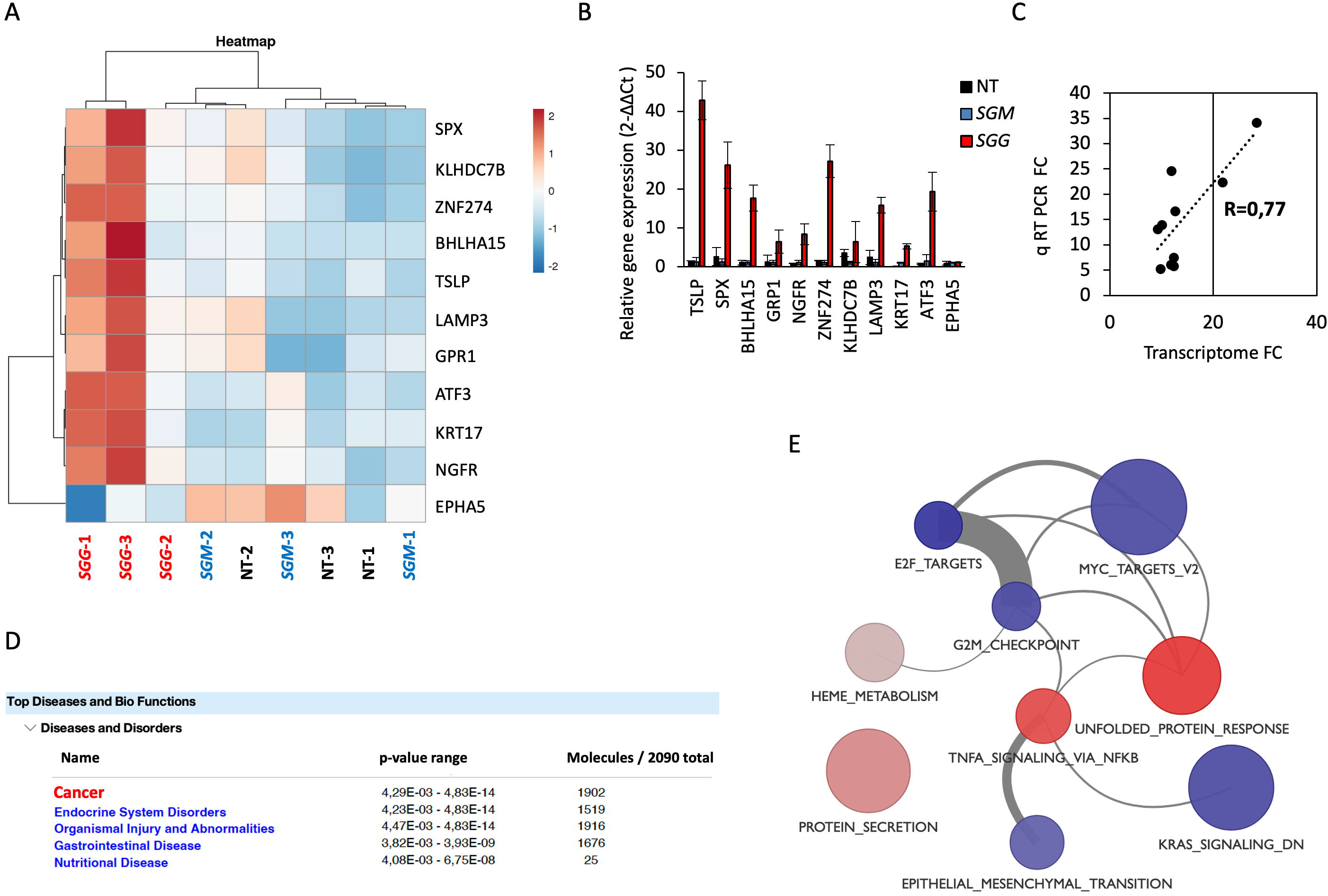
Transcriptional respo0nses of human tumoral HT29 cells upon *SGG* infection. **A.** Heat map of transcriptomic data displaying the top ten up-regulated and one down-regulated genes. **B**. Relative quantification (2^-ΔΔCt^) of mRNA levels at 24 h of infection with *SGG*, *SGM* and non-treated (NT). **C**. Correlation of fold changes between transcriptome and quantitative real time PCR for the 11 genes shown in B. **D**. Top Disease and Bio Functions using Ingenuity Pathway Analysis (IPA) using genes differentially expressed between *SGG* UCN34 and *SGM*. Molecules indicate the number of genes associated with indicated diseases and disorders. A right-tailed Fisher’s Exact Test was used to calculate a p-value determining the probability that each biological function and/or disease assigned to that data set is due to chance alone. **E**. Network of representation of the 9 significantly enriched hallmark gene sets upon *SGG* infection on HT29 cells. Each node (circle) represents a gene set. Node size is proportional to the number of genes present in the dataset and belonging to the gene set. Node colors reflect the average expression directionality (based on the average of the log fold expression for each gene belonging to the gene set) (red = over expressed; blue = down regulated; grey=no statistically significant difference in expression). The color intensity is proportional to the strength of the average expression. Each edge connecting two nodes means that two gene sets share the same genes. The edge width is proportional to the number of genes shared among the two gene sets. Edges are shown only if at least two genes are shared among the two gene sets (11 edges).

IPA analysis revealed that the most of the differentially expressed genes were involved in cancer disease (1,902 out of 2,090; **Fig. 3D**). In HT29 cells. 9 HALLMARK gene sets were significantly enriched, with four predicted to be activated: (i) unfolded protein response (UPR); (ii) protein secretion; (iii) TNFα signaling via NF-kB; (iv) Heme metabolism and five gene sets were predicted to be down-regulated: (v) E2F targets; (vi) G2M checkpoint; (vii) KRAS signaling; (viii) myc targets v2 and (ix) epithelial to mesenchymal transition (**Fig. 3E, Supp Table S8**). KEGG Pathway Analysis reveals a clear up-regulation of “Protein processing in endoplasmic reticulum (ER)” (48/169 total (41 up/7 down) **Supp. Table S5**). IPA tool revealed that the top canonical pathways were UPR and ER stress (**Supp. Fig S2**) but no directionality (activation or down-regulation) was attributed. The visualization of UPR in presented in **Fig. 4** and ER stress in **Supp. Fig S3**.

**Figure 4.**
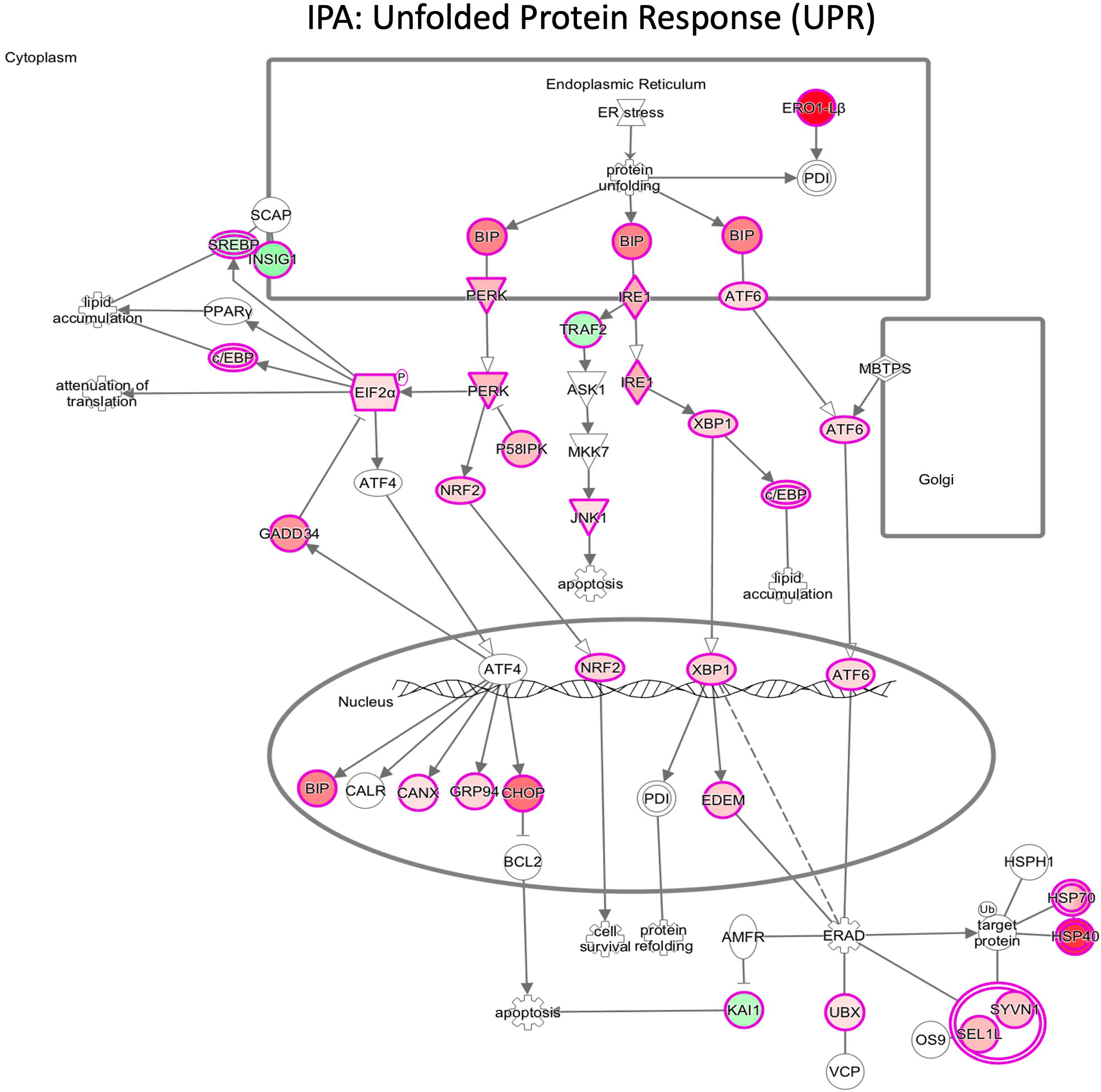
Activation of Unfolded Protein Response (UPR) signaling pathway in HT29 cells upon *SGG* infection. Differentially expressed genes in HT29 cells after 24 h of infection with *SGG* UCN34 vs *SGM*. Nodes represent molecules in a pathway, while the biological relationship between nodes is represented by a line (edge). Edges are supported by at least one reference in the Ingenuity Knowledge Base. The color intensity in a node indicates the degree of up- (red) or down- (green) regulation. Nodes that are red and green represent the increased and decreased measurements respectively. Nodes in orange represents predicted (hypothetical) activation and nodes in blue predicted (hypothetical) inhibition. Nodes are displayed using shapes that represent the functional class of a gene product (Circle = Other, Nested Circle = Group or Complex, Rhombus = Peptidase, Square = Cytokine, Triangle = Kinase, Vertical ellipse = Transmembrane receptor). Edges are marked with symbols to represent the relationship between nodes (Line only = Binding only, Flat line = inhibits, Solid arrow = Acts on, Solid arrow with flat line = inhibits and acts on, Open circle = leads to, Open arrow = translocates to). An orange line indicates predicted upregulation, whereas a blue line indicates predicted downregulation. A yellow line indicates expression being contradictory to the prediction. Gray line indicates that direction of change is not predicted. Solid or broken edges indicate direct or indirect relationships, respectively.

Taken together, transcriptomic data in HT29 cells shows a massive pro-tumoral shift with strong induction of ER stress and UPR activation by *SGG*.

## Discussion

We aimed to identify transcriptomic alterations in human colonic cells cultivated *in vitro* during interaction with the CRC-associated gut pathobiont *SGG*. As a control condition for comparison, we have chosen host cells infected with the genetically closely related but non-pathogenic bacterium *SGM*. As expected for a commensal bacterium, *SGM* did not induce any significant transcriptional changes on human colonic cells. In contrast, *SGG* altered a higher number of genes (2,090) in tumoral HT29 cells as compared to normal FHC cells (128 genes). Most of these genes were associated with cancer disease. This specific pro-tumoral transcriptional shift on colonic cells suggests that *SGG* is rather a weak tumor initiator and potentially a strong tumor accelerator. This result fits well with our previous *in vivo* data showing that *SGG* contributes to tumor development in mice pre-treated with a chemical mutagen such as azoxymethane (AOM) [14].

*SGG* displayed a weak (few genes and with quite a low fold-change) but specific pro-tumoral transcriptional signature on normal colonic cells. All the most up-regulated genes (*IL-20, CLK1, SORBS2, ERG1, PIM1, SNORD3A*) are involved in cancer development [37,39,40,42,44]. It is likely that simultaneous activation of these genes will in the long term contribute to pre-cancerous transformation of colonic cells. Only two genes, *clk1* and *polB,* were found up-regulated in both FHC and HT29 cells, suggesting a role in cell proliferation and/or DNA damage. We were not able to show a significant effect of *SGG* UCN34 on host cell proliferation (HT-29 and HCT-116) by direct counting of epithelial colonic cells [14]. To experimentally test whether *SGG* can induce DNA damage of colonic cells, DNA damage quantification using confocal microscopy (γH2AX foci detection [55]) in human colonic Caco-2 cells was performed showing a slight but not significant increase of DNA breaks induced by *SGG* as compared to *SGM* [14]. As a positive control, we used the *Escherichia coli* cells expressing the genotoxic toxin colibactin. Of note, Taddese *et al.* also evaluated the DNA damaging effect of *SGG* UCN34 using the DNA comet assay and showed that *SGG* alone had no effect while this bacterium increased the DNA damaging effect induced by the mutagen polycyclic aromatic hydrocarbon 3-methylcholanthrene [24]. This result also fits well with an acceleration role for *SGG*.

Taddese *et al* [24] also performed a global transcriptome on HT29 cells exposed to *SGG* UCN34 for 4 h and found 44 genes differentially regulated (21 genes up-regulated and 23 downregulated). The most up-regulated gene was *CYP1A1* (3.01-fold change) encoding cytochrome P450 suggesting that *SGG* can modify the capacity of intestinal epithelial or pre-cancerous cells to de-toxify dietary components. *CYPA1* (2.69-fold change) was also found up-regulated in our experimental set up (**Supp. Table S2**.). No other gene was found in common in HT29 after 24 h of infection with *SGG* UCN34. Our approach with 24 h of co-culture vs 4h from Taddese *et al* [24] revealed much more changes (2,090 vs 44 genes) and specific pro-tumoral shift. Thus, longer infection at low-bacterial dose appears physiologically relevant and informative.

A growing number of researchers have demonstrated that the Unfolded Protein Response (UPR) is closely involved in CRC development [56]. Our transcriptomic data shows clear induction of UPR by *SGG* in colonic epithelial cells (**Fig 5**; **Supp Fig S4)**. The expression of most of the proteins from these pathways were affected by *SGG* and almost all were up-regulated (red color) indicating their activation. UPR operates as a metabolic shift increasing cancer cell survival and adaptation to cope with major intrinsic and environmental challenges, promoting metastasis and angiogenesis, modulating inflammatory/immune responses [57]. ER stress could be a consequence of: (i) bacterial adhesion factors; (ii) bacterial secreted toxins/ effectors; or (iii) nutritional stress (glucose deprivation) [58],[59]. Group A *Streptococcus*, an extracellular human pathogen, was shown to induce ER stress to retrieve host nutrients such as asparagine and to increase host cell viability [60]. It is worth mentioning that *SGG* efficiently utilizes glucose and glycolysis metabolites for its own multiplication in HT29 culture spent medium [61]. Thus, we cannot completely rule out that it is the glucose consumption by *SGG* during proliferation in the cell culture medium that induced the strong UPR response detected at 24 h of co-culture. ER stress-mediated activation of the UPR is a double-edge sword in cancer development. During ER stress, cells either survive by inducing adaptation mechanisms or suicide by apoptosis. PERK is the most dominant branch of UPR to accelerate metastasis, as it can promote angiogenesis through increased expression of vascular endothelial growth factor (VEGF). Both PERK (2.82-fold change) and VEGF (2.38-fold change) expression were increased in our transcriptomic data. Of note, ER stress activates the TOR pathway through ATF6 [62]. We thus hypothesize that activation of ATF6 by *SGG* (**Fig. 4**) may lead to TOR pathway activation, a major signaling pathway identified recently by our proteomics and phospoproteomics analysis [14].

GSEA analysis also suggests also that *SGG* may trigger up-regulation of TNFα signaling via NFκB. The transcriptional factor NF-κB plays a crucial role in the host response to microbial infection by orchestrating innate and adaptive immune functions [63]. Its activity is linked to gastrointestinal cancer initiation and development through induction of chronic inflammation, cellular transformation and proliferation [64]. The modulation of the NF-κB signaling pathway was shown for several pathogens (e.g. *Helicobacter pylori, Fusobacterium nucleatum, Peptostreptococcus anaerobius, E. coli pks+, B. fragilis bft+*) and directly linked to their oncogenic potential (reviewed in [64]).

In conclusion, our data indicate that *SGG,* an opportunistic gut pathobiont associated with colorectal cancer, can alter host cell transcription in human colonic cells whereas the non-pathogenic commensal bacterium *SGM* does not. Comparing normal and tumoral responses, our results suggest that *SGG* accelerates colon tumor development rather than initiates this process and the strong activation of the UPR response by *SGG* requires further investigation.

## Data Availability

All data generated or analyzed during this study are included in this published article and its supplementary information files. In addition, the raw datasets of the transcriptomic study were deposited in the GEO database under the accession number GSE232211.

### Acknowledgements

This work was supported by the Institut National contre le Cancer (INCA, grant PLBIO16-025) and from the French Government’s Investissement d’Avenir program, Laboratoire d’Excellence Integrative Biology of Emerging Infectious Diseases (grant no. ANR-10-LABX-62-IBEID). E.P-K received a 2-year Roux-Cantarini post-doctoral fellowship. We thank Tarek Msadek for the careful editing of this manuscript.

## Author’s contributions

P-K.E. and D.S. conceived the study; P-K.E. and L.D. conducted the experiments; P-K.E. analyzed the data; L.D. and D.F. performed the transcriptomic analyses; N.P. performed bioinformatic analyses. H.V. analyzed the transcriptomic data. E.P-K. and S.D., wrote the paper. All authors edited and reviewed the manuscript.

## Supplemental Figure legends

**Figure S1. Cellular model of *SGG* UCN34 and *SGM* 24 h co-culture with human colonic normal FHC and cancerous HT29 cells. A.** Graph representing the number of bacteria present in cell culture wells (one well of a 6-well plate) after 24 h of co-culture. ns; Mann-Whitney test (p=0,33; p=0,66). **B**. Colored confocal pictures show DAPI nuclei labeling (blue), Phalloidin (red) and *SGG* UCN34 or *SGM* (green). Scale bar: 50 μm. **C**. 3D reconstruction of z-stack obtained by confocal microscopy. DAPI (blue), Phalloidin (red) and *SGG* UCN34 or *SGM* (green). **D**. Graph representing the number of adherent bacteria after 24 h of infection. Cells were seeded and grown until the full confluence in 6-well plate then inoculated with 6×10^5^ of *SGM* and 6×10^4^ of *SGG*. After the first 6 h of co-culture the trimethoprim (50μg/ml) was added to the cell culture media and infection was followed for 24 h.

**Figure S2. Top 5 canonical pathways altered by *SGG* UCN34 in HT29 cells visualized using IPA software.** The data set used in this analysis was genes differentially expressed between *SGG* UCN34 and *SGM* detected in HT29 after 24 h of infection. Canonical pathways that were most significant to the data set were identified from the QIAGEN Ingenuity Pathway Analysis library of canonical pathways. Canonical pathways with p-values < 0.05 (Fischer’s exact test) were statistically significant. The activation Z-score was calculated to predict activation or inhibition of transcriptional regulators based on published findings accessible through the Ingenuity knowledge base. Regulators with Z-score greater than 2 (positive Z-score) or less than −2 (negative Z-score) were significantly activated (orange) or inhibited (blue). Regulators with Z-score of 0 are represented in white and those for which the Z-score couldn’t be calculated are shown in grey.

**Figure S3. IPA identified the Endoplasmic reticulum (ER) stress pathway as enriched in HT29 cells after *SGG* UCN34 infection**. The data set used in this analysis was differentially expressed genes in HT29 cells after 24 h of infection with *SGG* UCN34 vs *SGM*. Nodes represent molecules in a pathway, while the biological relationship between nodes is represented by a line (edge). Edges are supported by at least one reference in the Ingenuity Knowledge Base. The color intensity in a node indicates the degree of up- (red) or down- (green) regulation. Nodes that are red and green represent the increased and decreased measurements respectively. Nodes in orange represents predicted (hypothetical) activation and nodes in blue predicted (hypothetical) inhibition. Nodes are displayed using shapes that represent the functional class of a gene product (Circle = Other, Nested Circle = Group or Complex, Rhombus = Peptidase, Square = Cytokine, Triangle = Kinase, Vertical ellipse = Transmembrane receptor). Edges are marked with symbols to represent the relationship between nodes (Line only = Binding only, Flat line = inhibits, Solid arrow = Acts on, Solid arrow with flat line = inhibits and acts on, Open circle = leads to, Open arrow = translocates to). An orange line indicates predicted upregulation, whereas a blue line indicates predicted downregulation. A yellow line indicates expression being contradictory to the prediction. Gray line indicates that direction of change is not predicted. Solid or broken edges indicate direct or indirect relationships, respectively.

## Supplemental Tables

**Table S1.** List of primers used for q RT-PCR.

**Table S2.** List of the 128 genes differentially expressed in FHC cells infected with *SGG* UCN34 vs *SGM* for 24 h.

**Table S3**. List of the 2,090 genes differentially expressed in HT29 cells infected with *SGG* UCN34 vs *SGM* for 24 h.

**Table S4.** List of the 41 genes differentially expressed in both HT29 and in FHC cells upon infection with *SGG* UCN34.

**Table S5.** List of 54 significantly up- and down-regulated pathways found by KEGG pathway analysis in HT29 cells comparing *SGG* UCN34 vs *SGM*.

**Table S6.** List of the 88 up- and down-regulated pathways found using Reactome pathway analysis in HT29 cells upon *SGG* infection.

**Table S7.** List of the 14 hallmark gene sets found with GSEA in FHC cells upon *SGG* infection.

**Table S8.** List of 9 hallmark gene sets found with GSEA in HT29 cells upon *SGG* infection.

