## Supplementary figures and images for "Transcriptome profiling of human colonic cells exposed to the gut pathobiont *Streptococcus gallolyticus* subsp. *gallolyticus*"

### Supplemental Fig 1

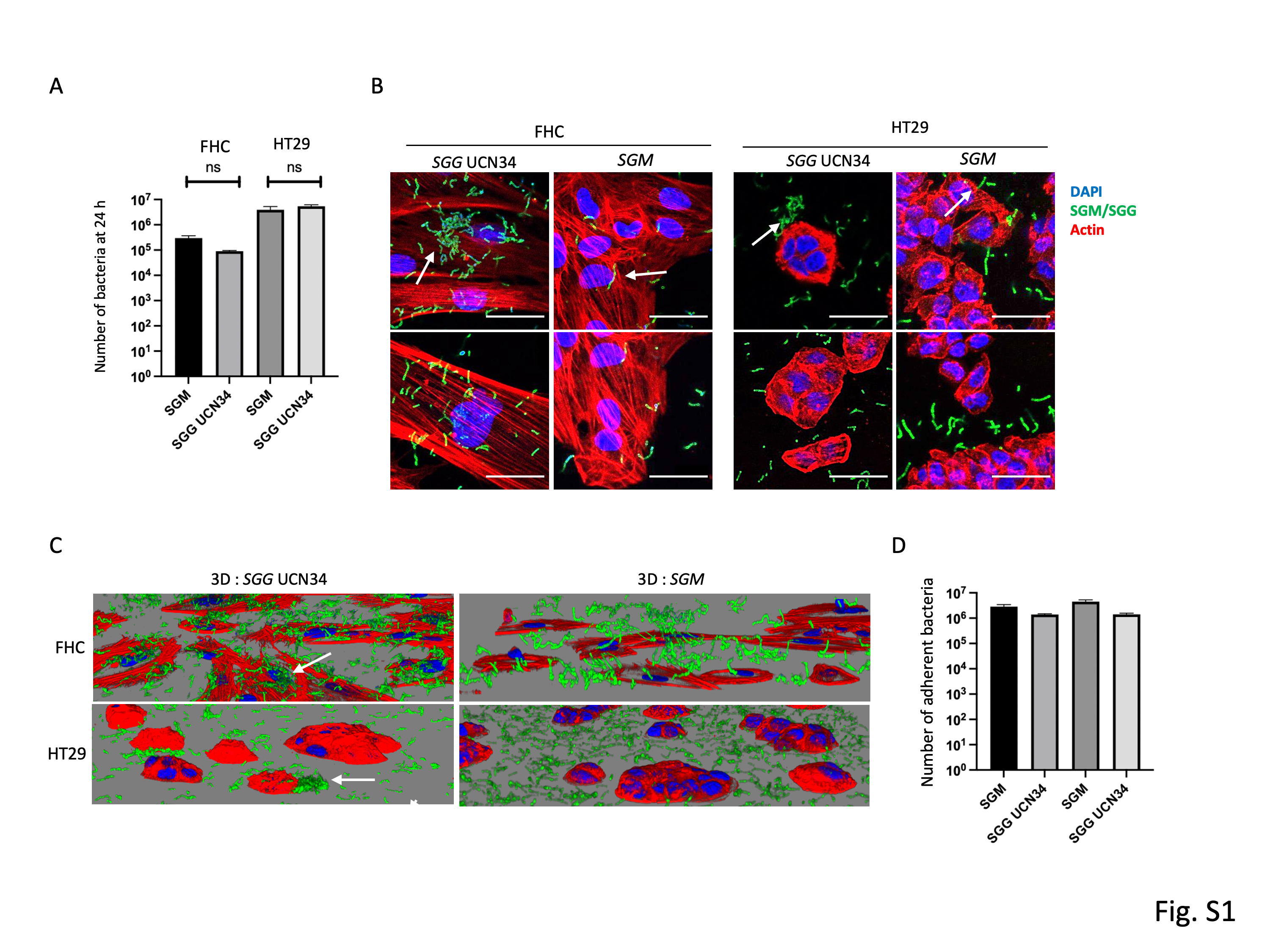

### Supplemental Fig 2

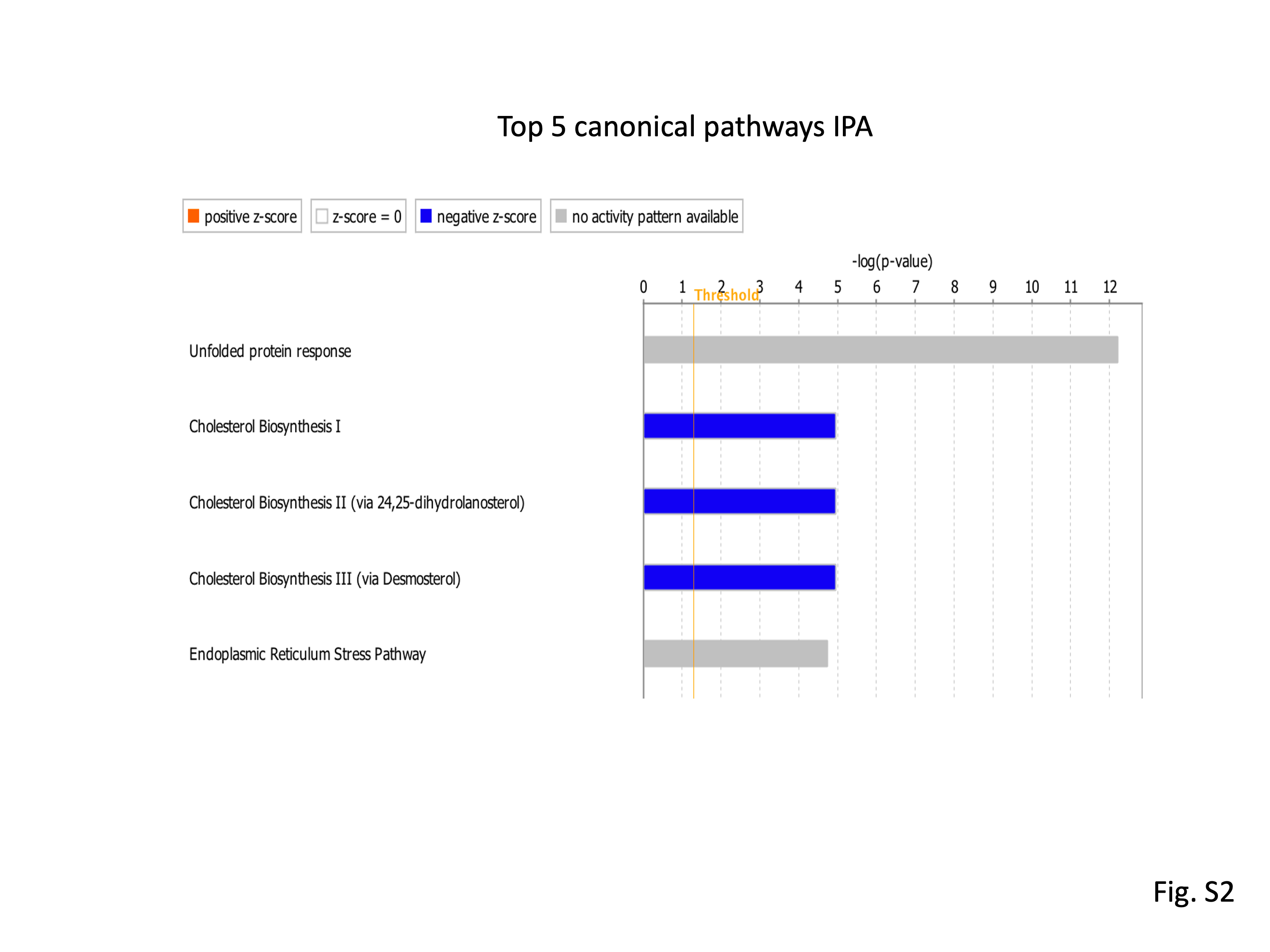

### Supplemental Fig 3

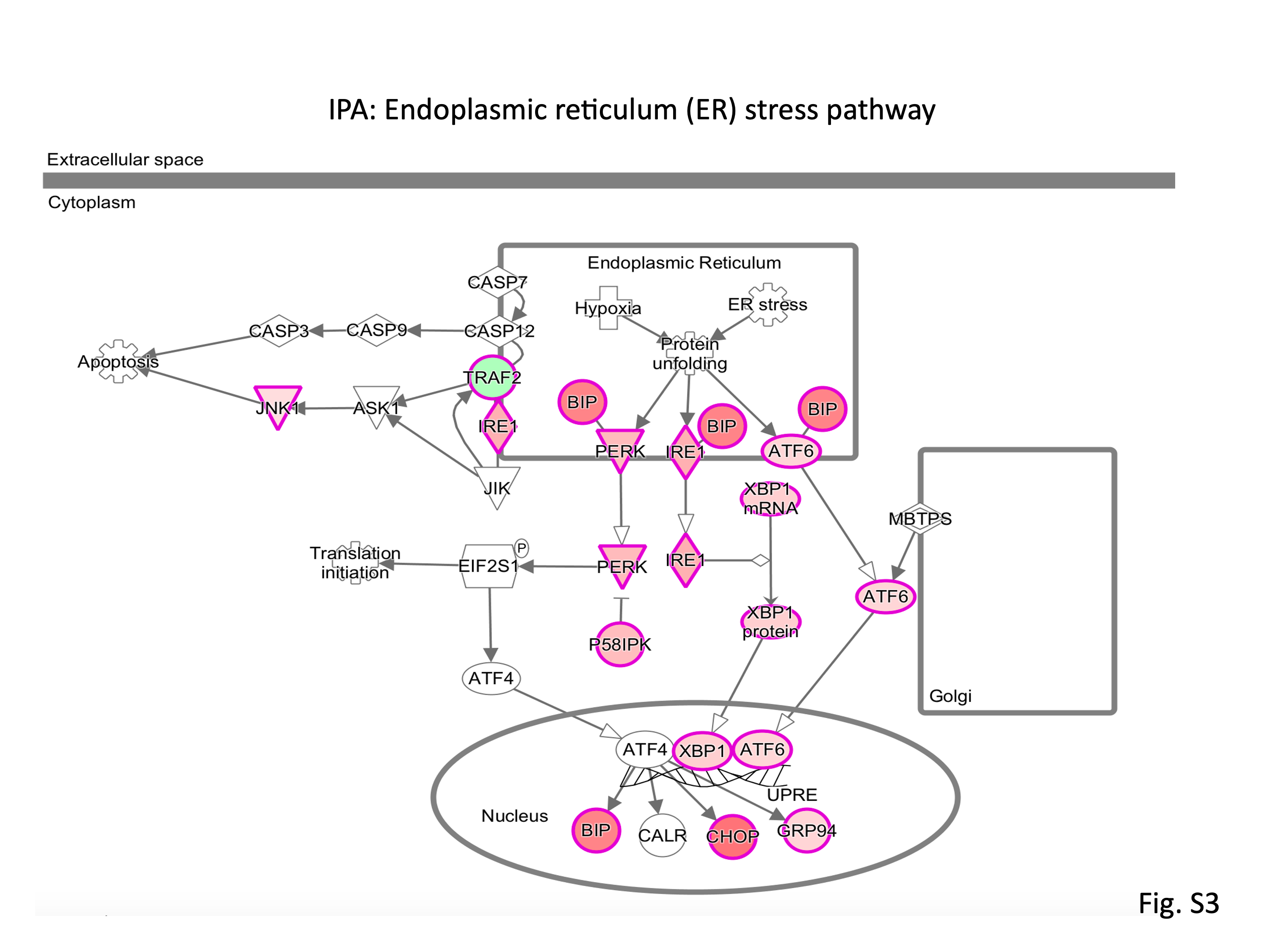
