## Supplemental Table 1 for "Transcriptome profiling of human colonic cells exposed to the gut pathobiont *Streptococcus gallolyticus* subsp. *gallolyticus*"

**Supplementary Table S1:** List of primers with sequences for quantitative real-time PCR.

| For HT29 cells | | PRIMERS | |
|---|---|---|---|
| Gene | Organism | Forward 5'- 3' | Reverse 5'- 3' |
| TSLP | human | TATCTGGTGCCCAGGCTATTCG | TGAAGCGACGCCACAATCCTTG |
| SPX | human | TCGCCGCTTCATCTCCGACCA | GACGCCAGTAAGATGGTTGCTG |
| BHLHA15 | human | GCGGACAAGAAGCTCTCCAAGA | TGGTAGTGCTGGTAGAGCTTGG |
| GPR1 | human | CCTTTGGCATCTGGCTGTGCAA | GCCGATGAGATAAGACAGGATGG |
| NGFR | human | CCTCATCCCTGTCTATTGCTCC | GTTGGCTCCTTGCTTGTTCTGC |
| ZNF274 | human | AGTACCGCGATGTGATGCTGGA | GGCTGGTTCTTCCTCAGGAATG |
| KLHDC7B | human | GCACAACTACCTGTTTCTGGCG | TGGCTCCAGATGTTGGTCAGAG |
| LAMP3 | human | TGGGAGCCTATTTGACCGTCTC | GCTGACAACTGGAGGCTCTGTT |
| KRT17 | human | GACTCAAGCTCACACAGCCCTT | ACACTCAGCCTGTGCTTGCCAA |
| ATF3 | human | CGCTGGAATCAGTCACTGTCAG | CTTGTTTCGGCACTTTGCAGCTG |
| EPHA5 | human | CAGCAGGCTATGGTGTCTTCAG | ACGCCGATAACCACTGCCAACA |
| GAPDH | human | GTCTCCTCTGACTTCAACAGCG | ACCACCCTGTTGCTGTAGCCAA |

| For FHC cells | | PRIMERS | |
|---|---|---|---|
| Gene | Organism | Forward 5'- 3' | Reverse 5'- 3' |
| IL20 | human | AGATCAGCAGCCTCGCCAATTC | CAAAGTGACTCAGAATCTGGCTG |
| CLK1 | human | CACACGATAGTAAGGAGCATTTAG | GGCAGAACTGTGTTCATCCCAG |
| SNORD3A | human | TAGAGCACCGAAAACCACGA | CCTCTCACTCCCCAATACGG |
| SORBS2 | human | GGAGATACTGTCTACATCCTCAG | AGGCTGTGCTTTCTCAGGAGGT |
| EGR1 | human | AGCAGCACCTTCAACCCTCAGG | GAGTGGTTTGGCTGGGGTAACT |
| PIM1 | human | TCTACTCAGGCATCCGCGTCTC | CTTCAGCAGGACCACTTCCATG |
| TXNIP | human | CAGCAGTGCAAACAGACTTCGG | CTGAGGAAGCTCAAAGCCGAAC |
| KLHL38 | human | ATGCAGAACCCTGTGCGCCTTA | CCTCCCACAATGACAATCCGCT |
| GAPDH | human | GTCTCCTCTGACTTCAACAGCG | ACCACCCTGTTGCTGTAGCCAA |
