## Supplemental Table 2 for "Transcriptome profiling of human colonic cells exposed to the gut pathobiont *Streptococcus gallolyticus* subsp. *gallolyticus*"

**Supplemental Table S2. Transcriptome:** List of 128 genes differentially expressed in FHC cells infected with *SGG* UCN34 vs *SGM* during 24 h

| EnsEMBL ID | Gene Symbol | Entrez Gene ID | Gene Name | Regulation | Fold-Change | Adjusted P-Value |
|---|---|---|---|---|---|---|
| **ENSG00000162891** | **IL20** | **50604** | **interleukin 20** | **up** | **4,51** | **3,87E-03** |
| **ENSG00000013441** | **CLK1** | **1195** | **CDC like kinase 1** | **up** | **2,58** | **3,17E-02** |
| **ENSG00000263934** | **SNORD3A** | **780851** | **small nucleolar RNA, C/D box 3A** | **up** | **2,54** | **1,02E-02** |
| **ENSG00000154556** | **SORBS2** | **8470** | **sorbin and SH3 domain containing** | **up** | **2,33** | **1,62E-04** |
| **ENSG00000120738** | **EGR1** | **1958** | **early growth response 1** | **up** | **2,13** | **4,98E-02** |
| **ENSG00000137193** | **PIM1** | **5292** | **Pim-1 proto-oncogene, serine/thre** | **up** | **1,99** | **2,97E-03** |
| ENSG00000184635 | ZNF93 | 81931 | zinc finger protein 93 | up | 1,70 | 2,43E-02 |
| ENSG00000105982 | RNF32 | 140545 | ring finger protein 32 | up | 1,63 | 1,95E-02 |
| ENSG00000235652 | LOC100507557 | 100507557 | uncharacterized LOC100507557 | up | 1,48 | 3,96E-02 |
| ENSG00000140876 | NUDT7 | 283927 | nudix hydrolase 7 | up | 1,45 | 7,67E-03 |
| ENSG00000205544 | TMEM256 | 254863 | transmembrane protein 256 | up | 1,43 | 1,78E-02 |
| ENSG00000234608 | MAPKAPK5-AS1 | 51275 | MAPKAPK5 antisense RNA 1 | up | 1,39 | 3,58E-02 |
| ENSG00000138175 | ARL3 | 403 | ADP ribosylation factor like GTPase | up | 1,36 | 2,53E-03 |
| ENSG00000198270 | TMEM116 | 89894 | transmembrane protein 116 | up | 1,35 | 3,99E-02 |
| ENSG00000070501 | POLB | 5423 | DNA polymerase beta | up | 1,33 | 3,69E-02 |
| ENSG00000106804 | C5 | 727 | complement C5 | up | 1,32 | 3,96E-02 |
| ENSG00000124596 | OARD1 | 221443 | O-acyl-ADP-ribose deacylase 1 | up | 1,28 | 3,70E-02 |
| ENSG00000168803 | ADAL | 161823 | adenosine deaminase like | up | 1,26 | 3,07E-03 |
| ENSG00000103479 | RBL2 | 5934 | RB transcriptional corepressor like 2 | up | 1,24 | 7,73E-03 |
| **ENSG00000265972** | **TXNIP** | **10628** | **thioredoxin interacting protein** | **down** | **3,48** | **5,66E-03** |
| **ENSG00000175946** | **KLHL38** | **340359** | **kelch like family member 38** | **down** | **2,79** | **7,27E-03** |
| ENSG00000127528 | KLF2 | 10365 | Kruppel like factor 2 | down | 2,76 | 4,29E-03 |
| ENSG00000164056 | SPRY1 | 10252 | sprouty RTK signaling antagonist 1 | down | 2,68 | 1,51E-03 |
| ENSG00000140450 | ARRDC4 | 91947 | arrestin domain containing 4 | down | 2,58 | 6,16E-04 |
| ENSG00000168906 | MAT2A | 4144 | methionine adenosyltransferase 2A | down | 2,29 | 6,64E-03 |
| ENSG00000031003 | FAM13B | 51306 | family with sequence similarity 13 me | down | 2,25 | 5,66E-03 |
| ENSG00000101188 | NTSR1 | 4923 | neurotensin receptor 1 | down | 2,10 | 3,92E-02 |
| ENSG00000282164 | PEG13 | 359809 | paternally expressed 13 | down | 2,07 | 2,69E-02 |
| ENSG00000101670 | LIPG | 9388 | lipase G, endothelial type | down | 2,02 | 2,62E-05 |
| ENSG00000043591 | ADRB1 | 153 | adrenoceptor beta 1 | down | 2,02 | 9,90E-03 |
| ENSG00000162650 | ATXN7L2 | 127002 | ataxin 7 like 2 | down | 1,87 | 6,36E-03 |
| ENSG00000258429 | PDF | 64146 | peptide deformylase, mitochondrial | down | 1,83 | 1,30E-02 |
| ENSG00000196843 | ARID5A | 10865 | AT-rich interaction domain 5A | down | 1,81 | 3,58E-02 |
| ENSG00000125347 | IRF1 | 3659 | interferon regulatory factor 1 | down | 1,75 | 3,58E-02 |
| ENSG00000247626 | MARS2 | 92935 | methionyl-tRNA synthetase 2, mitoch | down | 1,71 | 1,25E-06 |
| ENSG00000140691 | ARMC5 | 79798 | armadillo repeat containing 5 | down | 1,67 | 1,14E-03 |
| ENSG00000167173 | C15orf39 | 56905 | chromosome 15 open reading frame | down | 1,66 | 6,77E-05 |
| ENSG00000254470 | AP5B1 | 91056 | adaptor related protein complex 5 be | down | 1,66 | 8,23E-04 |
| ENSG00000139438 | FAM222A | 84915 | family with sequence similarity 222 m | down | 1,65 | 1,71E-02 |
| ENSG00000034152 | MAP2K3 | 5606 | mitogen-activated protein kinase kina | down | 1,62 | 1,18E-02 |
| ENSG00000113645 | WWC1 | 23286 | WW and C2 domain containing 1 | down | 1,61 | 9,51E-05 |
| ENSG00000145990 | GFOD1 | 54438 | glucose-fructose oxidoreductase dor | down | 1,61 | 3,58E-02 |
| ENSG00000135083 | CCNJL | 79616 | cyclin J like | down | 1,60 | 2,95E-02 |
| ENSG00000130810 | PPAN | 56342 | peter pan homolog (Drosophila) | down | 1,58 | 7,73E-03 |
| ENSG00000140987 | ZSCAN32 | 54925 | zinc finger and SCAN domain contai | down | 1,56 | 1,55E-02 |
| ENSG00000167394 | ZNF668 | 79759 | zinc finger protein 668 | down | 1,56 | 6,77E-05 |
| ENSG00000126391 | FRMD8 | 83786 | FERM domain containing 8 | down | 1,53 | 1,02E-02 |
| ENSG00000165684 | SNAPC4 | 6621 | small nuclear RNA activating comple | down | 1,51 | 1,34E-03 |
| ENSG00000108840 | HDAC5 | 10014 | histone deacetylase 5 | down | 1,51 | 7,67E-03 |
| ENSG00000177427 | MIEF2 | 125170 | mitochondrial elongation factor 2 | down | 1,50 | 1,51E-03 |
| ENSG00000068137 | PLEKHH3 | 79990 | pleckstrin homology, MyTH4 and FE | down | 1,49 | 2,52E-02 |
| ENSG00000135763 | URB2 | 9816 | URB2 ribosome biogenesis 2 homolo | down | 1,49 | 1,41E-02 |
| ENSG00000105738 | SIPA1L3 | 23094 | signal induced proliferation associate | down | 1,49 | 5,55E-03 |
| ENSG00000167716 | WDR81 | 124997 | WD repeat domain 81 | down | 1,49 | 3,52E-03 |
| ENSG00000007376 | RPUSD1 | 113000 | RNA pseudouridylate synthase doma | down | 1,47 | 5,55E-03 |
| ENSG00000131759 | RARA | 5914 | retinoic acid receptor alpha | down | 1,47 | 3,90E-04 |
| ENSG00000185085 | INTS5 | 80789 | integrator complex subunit 5 | down | 1,47 | 6,77E-05 |
| ENSG00000179041 | RRS1 | 23212 | ribosome biogenesis regulator homo | down | 1,47 | 5,55E-03 |
| ENSG00000160193 | WDR4 | 10785 | WD repeat domain 4 | down | 1,47 | 7,73E-03 |
| ENSG00000100038 | TOP3B | 8940 | DNA topoisomerase III beta | down | 1,47 | 4,12E-03 |
| ENSG00000266962 | LOC108783654 | 108783654 | uncharacterized LOC108783654 | down | 1,46 | 4,58E-02 |
| ENSG00000149930 | TAOK2 | 9344 | TAO kinase 2 | down | 1,45 | 1,99E-02 |
| ENSG00000099625 | CBARP | 255057 | CACN beta subunit associated regu | down | 1,44 | 4,11E-02 |
| ENSG00000150990 | DHX37 | 57647 | DEAH-box helicase 37 | down | 1,44 | 4,36E-03 |
| ENSG00000127334 | DYRK2 | 8445 | dual specificity tyrosine phosphoryla | down | 1,44 | 2,96E-05 |
| ENSG00000099364 | FBXL19 | 54620 | F-box and leucine rich repeat protein | down | 1,44 | 2,65E-02 |
| ENSG00000177548 | RABEP2 | 79874 | rabaptin, RAB GTPase binding effe | down | 1,43 | 3,29E-02 |
| ENSG00000165804 | ZNF219 | 51222 | zinc finger protein 219 | down | 1,43 | 4,82E-03 |
| ENSG00000185989 | RASA3 | 22821 | RAS p21 protein activator 3 | down | 1,43 | 3,99E-02 |
| ENSG00000047056 | WDR37 | 22884 | WD repeat domain 37 | down | 1,43 | 4,71E-02 |
| ENSG00000129911 | KLF16 | 83855 | Kruppel like factor 16 | down | 1,42 | 3,69E-02 |
| ENSG00000161847 | RAVER1 | 125950 | ribonucleoprotein, PTB binding 1 | down | 1,42 | 2,01E-02 |
| ENSG00000134815 | DHX34 | 9704 | DExH-box helicase 34 | down | 1,42 | 9,51E-03 |
| ENSG00000180921 | FAM83H | 286077 | family with sequence similarity 83 me | down | 1,41 | 4,82E-02 |
| ENSG00000119574 | ZBTB45 | 84878 | zinc finger and BTB domain contain | down | 1,40 | 7,67E-03 |
| ENSG00000183963 | SMTN | 6525 | smoothelin | down | 1,40 | 3,75E-05 |
| ENSG00000148840 | PPRC1 | 23082 | peroxisome proliferator-activated rec | down | 1,40 | 5,56E-03 |
| ENSG00000172534 | HCFC1 | 3054 | host cell factor C1 | down | 1,40 | 3,40E-05 |
| ENSG00000197119 | SLC25A29 | 123096 | solute carrier family 25 member 29 | down | 1,39 | 4,74E-02 |
| ENSG00000104142 | VPS18 | 57617 | VPS18, CORVET/HOPS core subun | down | 1,38 | 2,94E-02 |
| ENSG00000177303 | CASKIN2 | 57513 | CASK interacting protein 2 | down | 1,38 | 3,22E-04 |
| ENSG00000180035 | ZNF48 | 197407 | zinc finger protein 48 | down | 1,37 | 3,92E-02 |
| ENSG00000099991 | CABIN1 | 23523 | calcineurin binding protein 1 | down | 1,37 | 3,17E-02 |
| ENSG00000124160 | NCOA5 | 57727 | nuclear receptor coactivator 5 | down | 1,36 | 4,00E-02 |
| ENSG00000105662 | CRTC1 | 23373 | CREB regulated transcription coactiv | down | 1,36 | 2,58E-02 |
| ENSG00000126003 | PLAGL2 | 5326 | PLAG1 like zinc finger 2 | down | 1,36 | 4,82E-02 |
| ENSG00000130254 | SAFB2 | 9667 | scaffold attachment factor B2 | down | 1,35 | 7,73E-03 |
| ENSG00000173327 | MAP3K11 | 4296 | mitogen-activated protein kinase kin | down | 1,35 | 2,62E-02 |
| ENSG00000181830 | SLC35C1 | 55343 | solute carrier family 35 member C1 | down | 1,35 | 1,68E-02 |
| ENSG00000070047 | PHRF1 | 57661 | PHD and ring finger domains 1 | down | 1,34 | 2,14E-04 |
| ENSG00000138756 | BMP2K | 55589 | BMP2 inducible kinase | down | 1,34 | 4,58E-02 |
| ENSG00000197312 | DDI2 | 84301 | DNA damage inducible 1 homolog 2 | down | 1,34 | 1,89E-03 |
| ENSG00000105127 | AKAP8 | 10270 | A-kinase anchoring protein 8 | down | 1,34 | 6,45E-03 |
| ENSG00000103653 | CSK | 1445 | C-terminal Src kinase | down | 1,33 | 6,44E-04 |
| ENSG00000168040 | FADD | 8772 | Fas associated via death domain | down | 1,32 | 3,52E-02 |
| ENSG00000085872 | CHERP | 10523 | calcium homeostasis endoplasmic re | down | 1,32 | 1,14E-03 |
| ENSG00000241878 | PISD | 23761 | phosphatidylserine decarboxylase | down | 1,32 | 1,88E-03 |
| ENSG00000103042 | SLC38A7 | 55238 | solute carrier family 38 member 7 | down | 1,31 | 1,92E-02 |
| ENSG00000110104 | CCDC86 | 79080 | coiled-coil domain containing 86 | down | 1,31 | 5,55E-03 |
| ENSG00000173599 | PC | 5091 | pyruvate carboxylase | down | 1,31 | 4,71E-02 |
| ENSG00000124217 | MOCS3 | 27304 | molybdenum cofactor synthesis 3 | down | 1,31 | 5,55E-03 |
| ENSG00000185504 | FAAP100 | 80233 | Fanconi anemia core complex assoc | down | 1,31 | 5,55E-03 |
| ENSG00000177302 | TOP3A | 7156 | DNA topoisomerase III alpha | down | 1,31 | 1,78E-02 |
| ENSG00000238227 | TMEM250 | 90120 | transmembrane protein 250 | down | 1,31 | 2,69E-02 |
| ENSG00000198911 | SREBF2 | 6721 | sterol regulatory element binding tra | down | 1,30 | 1,29E-02 |
| ENSG00000197136 | PCNX3 | 399909 | pecanex homolog 3 | down | 1,30 | 2,43E-02 |
| ENSG00000108963 | DPH1 | 1801 | diphthamide biosynthesis 1 | down | 1,30 | 3,55E-02 |
| ENSG00000140320 | BAHD1 | 22893 | bromo adjacent homology domain co | down | 1,29 | 5,01E-03 |
| ENSG00000157933 | SKI | 6497 | SKI proto-oncogene | down | 1,29 | 4,14E-02 |
| ENSG00000125485 | DDX31 | 64794 | DEAD-box helicase 31 | down | 1,28 | 3,65E-02 |
| ENSG00000132694 | ARHGEF11 | 9826 | Rho guanine nucleotide exchange f | down | 1,28 | 2,43E-02 |
| ENSG00000185024 | BRF1 | 2972 | BRF1, RNA polymerase III transcript | down | 1,28 | 4,22E-02 |
| ENSG00000164077 | MON1A | 84315 | MON1 homolog A, secretory traffickin | down | 1,27 | 1,14E-03 |
| ENSG00000104228 | TRIM35 | 23087 | tripartite motif containing 35 | down | 1,27 | 3,58E-02 |
| ENSG00000198863 | RUNDC1 | 146923 | RUN domain containing 1 | down | 1,26 | 5,55E-03 |
| ENSG00000154832 | CXXC1 | 30827 | CXXC finger protein 1 | down | 1,26 | 3,96E-02 |
| ENSG00000148296 | SURF6 | 6838 | surfeit 6 | down | 1,26 | 3,52E-03 |
| ENSG00000169718 | DUS1L | 64118 | dihydrouridine synthase 1 like | down | 1,26 | 1,42E-02 |
| ENSG00000106344 | RBM28 | 55131 | RNA binding motif protein 28 | down | 1,25 | 5,55E-03 |
| ENSG00000186575 | NF2 | 4771 | neurofibromin 2 | down | 1,25 | 2,31E-03 |
| ENSG00000181523 | SGSH | 6448 | N-sulfoglucosamine sulfohydrolase | down | 1,23 | 4,22E-02 |
| ENSG00000141564 | RPTOR | 57521 | regulatory associated protein of MTC | down | 1,22 | 2,44E-02 |
| ENSG00000143157 | POGK | 57645 | pogo transposable element derived | down | 1,21 | 3,96E-02 |
| ENSG00000166170 | BAG5 | 9529 | BCL2 associated athanogene 5 | down | 1,21 | 3,16E-02 |
| ENSG00000072518 | MARK2 | 2011 | microtubule affinity regulating kinase | down | 1,20 | 3,25E-02 |
| ENSG00000100029 | PES1 | 23481 | pescadillo ribosomal biogenesis fact | down | 1,20 | 2,73E-02 |
| ENSG00000259956 | RBM15B | 29890 | RNA binding motif protein 15B | down | 1,19 | 4,82E-02 |
| ENSG00000140521 | POLG | 5428 | DNA polymerase gamma, catalytic su | down | 1,17 | 4,16E-02 |
