## Supplemental Table 3 for "Transcriptome profiling of human colonic cells exposed to the gut pathobiont *Streptococcus gallolyticus* subsp. *gallolyticus*"

**Supplemental Table S3. Transcriptome:** List of 2090 genes differentially expressed in HT29 cells infected with *SGG* UCN34 vs *SGM* during 24 h.

| EnsEMBL ID | Gene Symbol | Entrez Gene ID | Gene Name | Regulation | Fold-Change | Adjusted P-Value |
|---|---|---|---|---|---|---|
| **ENSG00000145777** | **TSLP** | **85480** | **thymic stromal lymphopoietin** | **up** | **28,36** | **1,78E-04** |
| **ENSG00000134548** | **SPX** | **80763** | **spexin hormone** | **up** | **21,75** | **6,80E-04** |
| **ENSG00000180535** | **BHLHA15** | **168620** | **basic helix-loop-helix family member a15** | **up** | **12,57** | **2,59E-02** |
| **ENSG00000183671** | **GPR1** | **2825** | **G protein-coupled receptor 1** | **up** | **12,36** | **2,50E-02** |
| **ENSG00000064300** | **NGFR** | **4804** | **nerve growth factor receptor** | **up** | **12,35** | **2,75E-03** |
| **ENSG00000171606** | **ZNF274** | **10782** | **zinc finger protein 274** | **up** | **11,88** | **2,26E-03** |
| **ENSG00000130487** | **KLHDC7B** | **113730** | **kelch domain containing 7B** | **up** | **11,82** | **1,01E-02** |
| **ENSG00000078081** | **LAMP3** | **27074** | **lysosomal associated membrane protein 3** | **up** | **10,09** | **5,27E-03** |
| **ENSG00000128422** | **KRT17** | **3872** | **keratin 17** | **up** | **9,79** | **8,14E-03** |
| **ENSG00000162772** | **ATF3** | **467** | **activating transcription factor 3** | **up** | **9,22** | **4,75E-04** |
| ENSG00000086619 | ERO1B | 56605 | endoplasmic reticulum oxidoreductase 1 beta | up | 8,94 | 1,02E-02 |
| ENSG00000115008 | IL1A | 3552 | interleukin 1 alpha | up | 8,56 | 1,06E-02 |
| ENSG00000128590 | DNAJB9 | 4189 | DnaJ heat shock protein family (Hsp40) member B9 | up | 7,76 | 1,31E-04 |
| ENSG00000187122 | SLIT1 | 6585 | slit guidance ligand 1 | up | 7,59 | 2,14E-03 |
| ENSG00000135406 | PRPH | 5630 | peripherin | up | 7,22 | 8,91E-03 |
| ENSG00000081181 | ARG2 | 384 | arginase 2 | up | 7,09 | 3,92E-04 |
| ENSG00000187800 | PEAR1 | 375033 | platelet endothelial aggregation receptor 1 | up | 6,01 | 1,12E-03 |
| ENSG00000153714 | LURAP1L | 286343 | leucine rich adaptor protein 1 like | up | 5,90 | 1,32E-03 |
| ENSG00000166750 | SLFN5 | 162394 | schlafen family member 5 | up | 5,78 | 1,60E-02 |
| ENSG00000175197 | DDIT3 | 1649 | DNA damage inducible transcript 3 | up | 5,65 | 3,67E-03 |
| ENSG00000073712 | FERMT2 | 10979 | fermitin family member 2 | up | 5,34 | 2,65E-03 |
| ENSG00000263006 | ROCK1P1 | 727758 | Rho associated coiled-coil containing protein kinase 1 pseudogene 1 | up | 5,31 | 4,04E-03 |
| ENSG00000235899 | LINC01564 | 101927171 | long intergenic non-protein coding RNA 1564 | up | 5,18 | 4,43E-03 |
| ENSG00000044574 | HSPA5 | 3309 | heat shock protein family A (Hsp70) member 5 | up | 5,02 | 6,99E-04 |
| ENSG00000172602 | RND1 | 27289 | Rho family GTPase 1 | up | 4,86 | 3,18E-03 |
| ENSG00000198855 | FICD | 11153 | FIC domain containing | up | 4,68 | 9,56E-04 |
| ENSG00000170909 | OSCAR | 126014 | osteoclast associated, immunoglobulin-like receptor | up | 4,60 | 2,11E-02 |
| ENSG00000115295 | CLIP4 | 79745 | CAP-Gly domain containing linker protein family member 4 | up | 4,59 | 2,57E-03 |
| ENSG00000051108 | HERPUD1 | 9709 | homocysteine inducible ER protein with ubiquitin like domain 1 | up | 4,55 | 1,12E-03 |
| ENSG00000150961 | SEC24D | 9871 | SEC24 homolog D, COPII coat complex component | up | 4,45 | 6,80E-04 |
| ENSG00000183696 | UPP1 | 7378 | uridine phosphorylase 1 | up | 4,41 | 1,63E-03 |
| ENSG00000107731 | UNC5B | 219699 | unc-5 netrin receptor B | up | 4,39 | 5,27E-03 |
| ENSG00000171658 | NMRAL2P | 344887 | NmrA like redox sensor 2, pseudogene | up | 4,38 | 1,20E-02 |
| ENSG00000092295 | TGM1 | 7051 | transglutaminase 1 | up | 4,37 | 2,22E-03 |
| ENSG00000087074 | PPP1R15A | 23645 | protein phosphatase 1 regulatory subunit 15A | up | 4,33 | 1,06E-03 |
| ENSG00000186594 | MIR22HG | 84981 | MIR22 host gene | up | 4,23 | 1,69E-03 |
| ENSG00000165125 | TRPV6 | 55503 | transient receptor potential cation channel subfamily V member 6 | up | 4,22 | 2,17E-02 |
| ENSG00000163393 | SLC22A15 | 55356 | solute carrier family 22 member 15 | up | 4,05 | 6,80E-04 |
| ENSG00000178381 | ZFAND2A | 90637 | zinc finger AN1-type containing 2A | up | 3,95 | 5,64E-04 |
| ENSG00000132837 | DMGDH | 29958 | dimethylglycine dehydrogenase | up | 3,92 | 1,01E-02 |
| ENSG00000118985 | ELL2 | 22936 | elongation factor for RNA polymerase II 2 | up | 3,83 | 1,27E-03 |
| ENSG00000128165 | ADM2 | 79924 | adrenomedullin 2 | up | 3,79 | 7,17E-03 |
| ENSG00000133639 | BTG1 | 694 | BTG anti-proliferation factor 1 | up | 3,75 | 1,43E-03 |
| ENSG00000144452 | ABCA12 | 26154 | ATP binding cassette subfamily A member 12 | up | 3,73 | 1,00E-02 |
| ENSG00000174564 | IL20RB | 53833 | interleukin 20 receptor subunit beta | up | 3,70 | 6,96E-03 |
| ENSG00000155304 | HSPA13 | 6782 | heat shock protein family A (Hsp70) member 13 | up | 3,69 | 7,84E-04 |
| ENSG00000113070 | HBEGF | 1839 | heparin binding EGF like growth factor | up | 3,67 | 5,68E-03 |
| ENSG00000135842 | FAM129A | 116496 | family with sequence similarity 129 member A | up | 3,66 | 2,70E-03 |
| ENSG00000138166 | DUSP5 | 1847 | dual specificity phosphatase 5 | up | 3,63 | 9,77E-03 |
| ENSG00000124762 | CDKN1A | 1026 | cyclin dependent kinase inhibitor 1A | up | 3,62 | 5,43E-03 |
| ENSG00000197860 | SGTB | 54557 | small glutamine rich tetratricopeptide repeat containing beta | up | 3,56 | 1,35E-03 |
| ENSG00000140961 | OSGIN1 | 29948 | oxidative stress induced growth inhibitor 1 | up | 3,55 | 8,36E-03 |
| ENSG00000114248 | LRRC31 | 79782 | leucine rich repeat containing 31 | up | 3,49 | 2,18E-02 |
| ENSG00000148677 | ANKRD1 | 27063 | ankyrin repeat domain 1 | up | 3,48 | 3,01E-02 |
| ENSG00000146278 | PNRC1 | 10957 | proline rich nuclear receptor coactivator 1 | up | 3,45 | 1,50E-03 |
| ENSG00000196352 | CD55 | 1604 | CD55 molecule (Cromer blood group) | up | 3,43 | 2,24E-03 |
| ENSG00000204758 | LOC100268168 | 100268168 | uncharacterized LOC100268168 | up | 3,41 | 4,03E-03 |
| ENSG00000116761 | CTH | 1491 | cystathionine gamma-lyase | up | 3,41 | 4,95E-03 |
| ENSG00000107242 | PIP5K1B | 8395 | phosphatidylinositol-4-phosphate 5-kinase type 1 beta | up | 3,40 | 1,85E-02 |
| ENSG00000070540 | WIPI1 | 55062 | WD repeat domain, phosphoinositide interacting 1 | up | 3,38 | 2,87E-03 |
| ENSG00000172432 | GTPBP2 | 54676 | GTP binding protein 2 | up | 3,38 | 2,11E-03 |
| ENSG00000156510 | HKDC1 | 80201 | hexokinase domain containing 1 | up | 3,37 | 4,73E-03 |
| ENSG00000104419 | NDRG1 | 10397 | N-myc downstream regulated 1 | up | 3,34 | 3,54E-02 |
| ENSG00000159885 | ZNF222 | 7673 | zinc finger protein 222 | up | 3,33 | 1,41E-03 |
| ENSG00000109321 | AREG | 374 | amphiregulin | up | 3,33 | 2,37E-02 |
| ENSG00000164683 | HEY1 | 23462 | hes related family bHLH transcription factor with YRPW motif 1 | up | 3,28 | 1,40E-02 |
| ENSG00000178607 | ERN1 | 2081 | endoplasmic reticulum to nucleus signaling 1 | up | 3,27 | 7,85E-03 |
| ENSG00000106080 | FKBP14 | 55033 | FK506 binding protein 14 | up | 3,26 | 1,30E-03 |
| ENSG00000176438 | SYNE3 | 161176 | spectrin repeat containing nuclear envelope family member 3 | up | 3,24 | 3,41E-03 |
| ENSG00000146592 | CREB5 | 9586 | cAMP responsive element binding protein 5 | up | 3,24 | 2,64E-03 |
| ENSG00000154153 | RETREG1 | 54463 | reticulophagy regulator 1 | up | 3,23 | 1,95E-02 |
| ENSG00000099860 | GADD45B | 4616 | growth arrest and DNA damage inducible beta | up | 3,21 | 3,92E-04 |
| ENSG00000101255 | TRIB3 | 57761 | tribbles pseudokinase 3 | up | 3,17 | 8,07E-03 |
| ENSG00000080031 | PTPRH | 5794 | protein tyrosine phosphatase, receptor type H | up | 3,16 | 2,17E-02 |
| ENSG00000125089 | SH3TC1 | 54436 | SH3 domain and tetratricopeptide repeats 1 | up | 3,14 | 3,06E-03 |
| ENSG00000102401 | ARMCX3 | 51566 | armadillo repeat containing, X-linked 3 | up | 3,13 | 1,19E-03 |
| ENSG00000078018 | MAP2 | 4133 | microtubule associated protein 2 | up | 3,05 | 3,16E-02 |
| ENSG00000116717 | GADD45A | 1647 | growth arrest and DNA damage inducible alpha | up | 3,05 | 4,75E-04 |
| ENSG00000131018 | SYNE1 | 23345 | spectrin repeat containing nuclear envelope protein 1 | up | 3,01 | 1,12E-03 |
| ENSG00000118292 | C1orf54 | 79630 | chromosome 1 open reading frame 54 | up | 2,99 | 3,98E-03 |
| ENSG00000260916 | CCPG1 | 9236 | cell cycle progression 1 | up | 2,96 | 4,42E-03 |
| ENSG00000149428 | HYOU1 | 10525 | hypoxia up-regulated 1 | up | 2,96 | 5,95E-03 |
| ENSG00000134278 | SPIRE1 | 56907 | spire type actin nucleation factor 1 | up | 2,93 | 2,14E-03 |
| ENSG00000198380 | GFPT1 | 2673 | glutamine--fructose-6-phosphate transaminase 1 | up | 2,92 | 2,14E-03 |
| ENSG00000140941 | MAP1LC3B | 81631 | microtubule associated protein 1 light chain 3 beta | up | 2,92 | 2,24E-03 |
| ENSG00000088826 | SMOX | 54498 | spermine oxidase | up | 2,90 | 6,10E-03 |
| ENSG00000130766 | SESN2 | 83667 | sestrin 2 | up | 2,90 | 1,62E-02 |
| ENSG00000139364 | TMEM132B | 114795 | transmembrane protein 132B | up | 2,90 | 1,81E-02 |
| ENSG00000163827 | LRRC2 | 79442 | leucine rich repeat containing 2 | up | 2,87 | 2,44E-02 |
| ENSG00000261040 | WFDC21P | 645638 | WAP four-disulfide core domain 21, pseudogene | up | 2,87 | 2,51E-02 |
| ENSG00000108176 | DNAJC12 | 56521 | DnaJ heat shock protein family (Hsp40) member C12 | up | 2,86 | 2,84E-02 |
| ENSG00000070081 | NUCB2 | 4925 | nucleobindin 2 | up | 2,86 | 3,18E-03 |
| ENSG00000204334 | ERICH2 | 285141 | glutamate rich 2 | up | 2,86 | 3,06E-02 |
| ENSG00000074935 | TUBE1 | 51175 | tubulin epsilon 1 | up | 2,86 | 1,41E-03 |
| ENSG00000166173 | LARP6 | 55323 | La ribonucleoprotein domain family member 6 | up | 2,84 | 5,01E-03 |
| ENSG00000163121 | NEURL3 | 93082 | neuralized E3 ubiquitin protein ligase 3 | up | 2,82 | 4,57E-02 |
| ENSG00000172071 | EIF2AK3 | 9451 | eukaryotic translation initiation factor 2 alpha kinase 3 | up | 2,82 | 1,12E-03 |
| ENSG00000176018 | LYSMD3 | 116068 | LysM domain containing 3 | up | 2,79 | 1,50E-03 |
| ENSG00000185022 | MAFF | 23764 | MAF bZIP transcription factor F | up | 2,78 | 1,03E-02 |
| ENSG00000120889 | TNFRSF10B | 8795 | TNF receptor superfamily member 10b | up | 2,77 | 1,99E-03 |
| ENSG00000145390 | USP53 | 54532 | ubiquitin specific peptidase 53 | up | 2,76 | 4,78E-03 |
| ENSG00000071537 | SEL1L | 6400 | SEL1L ERAD E3 ligase adaptor subunit | up | 2,76 | 4,04E-03 |
| ENSG00000242247 | ARFGAP3 | 26286 | ADP ribosylation factor GTPase activating protein 3 | up | 2,76 | 2,05E-03 |
| ENSG00000102580 | DNAJC3 | 5611 | DnaJ heat shock protein family (Hsp40) member C3 | up | 2,75 | 2,24E-03 |
| ENSG00000137094 | DNAJB5 | 25822 | DnaJ heat shock protein family (Hsp40) member B5 | up | 2,75 | 1,07E-02 |
| ENSG00000258102 | MAP1LC3B2 | 643246 | microtubule associated protein 1 light chain 3 beta 2 | up | 2,75 | 6,96E-03 |
| ENSG00000125772 | GPCPD1 | 56261 | glycerophosphocholine phosphodiesterase 1 | up | 2,75 | 2,24E-03 |
| ENSG00000196878 | LAMB3 | 3914 | laminin subunit beta 3 | up | 2,74 | 1,16E-02 |
| ENSG00000166401 | SERPINB8 | 5271 | serpin family B member 8 | up | 2,74 | 1,06E-02 |
| ENSG00000115977 | AAK1 | 22848 | AP2 associated kinase 1 | up | 2,72 | 2,09E-03 |
| ENSG00000116285 | ERRFI1 | 54206 | ERBB receptor feedback inhibitor 1 | up | 2,72 | 5,35E-03 |
| ENSG00000176142 | TMEM39A | 55254 | transmembrane protein 39A | up | 2,71 | 1,30E-03 |
| ENSG00000113742 | CPEB4 | 80315 | cytoplasmic polyadenylation element binding protein 4 | up | 2,70 | 6,55E-03 |
| ENSG00000141582 | CBX4 | 8535 | chromobox 4 | up | 2,70 | 4,01E-03 |
| ENSG00000059728 | MXD1 | 4084 | MAX dimerization protein 1 | up | 2,70 | 2,75E-03 |
| ENSG00000143333 | RGS16 | 6004 | regulator of G protein signaling 16 | up | 2,69 | 4,58E-03 |
| ENSG00000140465 | CYP1A1 | 1543 | cytochrome P450 family 1 subfamily A member 1 | up | 2,69 | 1,95E-02 |
| ENSG00000048052 | HDAC9 | 9734 | histone deacetylase 9 | up | 2,67 | 1,19E-02 |
| ENSG00000183508 | FAM46C | 54855 | family with sequence similarity 46 member C | up | 2,67 | 2,85E-02 |
| ENSG00000042832 | TG | 7038 | thyroglobulin | up | 2,66 | 1,33E-02 |
| ENSG00000197978 | GOLGA6L9 | 440295 | golgin A6 family-like 9 | up | 2,65 | 2,48E-02 |
| ENSG00000143479 | DYRK3 | 8444 | dual specificity tyrosine phosphorylation regulated kinase 3 | up | 2,65 | 1,81E-02 |
| ENSG00000165891 | E2F7 | 144455 | E2F transcription factor 7 | up | 2,65 | 1,22E-02 |
| ENSG00000134668 | SPOCD1 | 90853 | SPOC domain containing 1 | up | 2,64 | 2,24E-03 |
| ENSG00000166046 | TCP11L2 | 255394 | t-complex 11 like 2 | up | 2,64 | 6,03E-03 |
| ENSG00000139112 | GABARAPL1 | 23710 | GABA type A receptor associated protein like 1 | up | 2,62 | 4,13E-03 |
| ENSG00000116514 | RNF19B | 127544 | ring finger protein 19B | up | 2,61 | 4,56E-03 |
| ENSG00000152689 | RASGRP3 | 25780 | RAS guanyl releasing protein 3 | up | 2,59 | 1,79E-02 |
| ENSG00000151694 | ADAM17 | 6868 | ADAM metallopeptidase domain 17 | up | 2,59 | 1,61E-03 |
| ENSG00000152527 | PLEKHH2 | 130271 | pleckstrin homology, MyTH4 and FERM domain containing H2 | up | 2,56 | 2,41E-03 |
| ENSG00000204983 | PRSS1 | 5644 | protease, serine 1 | up | 2,54 | 4,99E-02 |
| ENSG00000020181 | ADGRA2 | 25960 | adhesion G protein-coupled receptor A2 | up | 2,51 | 1,87E-02 |
| ENSG00000259429 | UBE2Q2P2 | 100134869 | ubiquitin conjugating enzyme E2 Q2 pseudogene 2 | up | 2,50 | 1,58E-02 |
| ENSG00000118523 | CTGF | 1490 | connective tissue growth factor | up | 2,49 | 1,32E-02 |
| ENSG00000144655 | CSRNP1 | 64651 | cysteine and serine rich nuclear protein 1 | up | 2,49 | 2,24E-02 |
| ENSG00000160712 | IL6R | 3570 | interleukin 6 receptor | up | 2,48 | 3,34E-02 |
| ENSG00000135241 | PNPLA8 | 50640 | patatin like phospholipase domain containing 8 | up | 2,48 | 3,37E-03 |
| ENSG00000267757 | EML2-AS1 | 100287177 | EML2 antisense RNA 1 | up | 2,47 | 3,19E-02 |
| ENSG00000070669 | ASNS | 440 | asparagine synthetase (glutamine-hydrolyzing) | up | 2,47 | 2,91E-02 |
| ENSG00000236830 | CBR3-AS1 | 100506428 | CBR3 antisense RNA 1 | up | 2,46 | 1,13E-02 |
| ENSG00000138623 | SEMA7A | 8482 | semaphorin 7A (John Milton Hagen blood group) | up | 2,44 | 8,67E-03 |
| ENSG00000113811 | SELENOK | 58515 | selenoprotein K | up | 2,44 | 3,33E-03 |
| ENSG00000113369 | ARRDC3 | 57561 | arrestin domain containing 3 | up | 2,43 | 1,80E-03 |
| ENSG00000173846 | PLK3 | 1263 | polo like kinase 3 | up | 2,42 | 3,61E-03 |
| ENSG00000184545 | DUSP8 | 1850 | dual specificity phosphatase 8 | up | 2,42 | 1,40E-02 |
| ENSG00000145780 | FEM1C | 56929 | fem-1 homolog C | up | 2,42 | 2,14E-03 |
| ENSG00000175106 | TVP23C | 201158 | trans-golgi network vesicle protein 23 homolog C | up | 2,41 | 2,19E-02 |
| ENSG00000168003 | SLC3A2 | 6520 | solute carrier family 3 member 2 | up | 2,41 | 7,95E-03 |
| ENSG00000070190 | DAPP1 | 27071 | dual adaptor of phosphotyrosine and 3-phosphoinositides 1 | up | 2,40 | 3,92E-02 |
| ENSG00000147883 | CDKN2B | 1030 | cyclin dependent kinase inhibitor 2B | up | 2,40 | 3,90E-02 |
| ENSG00000167995 | BEST1 | 7439 | bestrophin 1 | up | 2,40 | 5,91E-03 |
| ENSG00000145088 | EAF2 | 55840 | ELL associated factor 2 | up | 2,39 | 6,68E-03 |
| ENSG00000280670 | CCDC163 | 126661 | coiled-coil domain containing 163 | up | 2,39 | 1,25E-02 |
| ENSG00000112715 | VEGFA | 7422 | vascular endothelial growth factor A | up | 2,38 | 1,68E-02 |
| ENSG00000013392 | RWDD2A | 112611 | RWD domain containing 2A | up | 2,37 | 4,21E-03 |
| ENSG00000165030 | NFIL3 | 4783 | nuclear factor, interleukin 3 regulated | up | 2,37 | 1,12E-03 |
| ENSG00000179094 | PER1 | 5187 | period circadian regulator 1 | up | 2,36 | 1,39E-02 |
| ENSG00000173275 | ZNF449 | 203523 | zinc finger protein 449 | up | 2,35 | 1,36E-03 |
| ENSG00000115844 | DLX2 | 1746 | distal-less homeobox 2 | up | 2,35 | 2,46E-02 |
| ENSG00000138709 | LARP1B | 55132 | La ribonucleoprotein domain family member 1B | up | 2,35 | 3,46E-03 |
| ENSG00000143507 | DUSP10 | 11221 | dual specificity phosphatase 10 | up | 2,34 | 2,20E-03 |
| ENSG00000105829 | BET1 | 10282 | Bet1 golgi vesicular membrane trafficking protein | up | 2,34 | 3,79E-03 |
| ENSG00000149201 | CCDC81 | 60494 | coiled-coil domain containing 81 | up | 2,34 | 2,43E-02 |
| ENSG00000150527 | MIA2 | 4253 | melanoma inhibitory activity 2 | up | 2,33 | 4,65E-03 |
| ENSG00000173166 | RAPH1 | 65059 | Ras association (RalGDS/AF-6) and pleckstrin homology domains 1 | up | 2,33 | 3,37E-03 |
| ENSG00000162298 | SYVN1 | 84447 | synoviolin 1 | up | 2,33 | 1,93E-02 |
| ENSG00000165195 | PIGA | 5277 | phosphatidylinositol glycan anchor biosynthesis class A | up | 2,32 | 6,64E-03 |
| ENSG00000182158 | CREB3L2 | 64764 | cAMP responsive element binding protein 3 like 2 | up | 2,31 | 8,54E-03 |
| ENSG00000130066 | SAT1 | 6303 | spermidine/spermine N1-acetyltransferase 1 | up | 2,30 | 3,56E-02 |
| ENSG00000151135 | TMEM263 | 90488 | transmembrane protein 263 | up | 2,30 | 1,31E-02 |
| ENSG00000120053 | GOT1 | 2805 | glutamic-oxaloacetic transaminase 1 | up | 2,30 | 5,68E-03 |
| ENSG00000163536 | SERPINI1 | 5274 | serpin family I member 1 | up | 2,29 | 9,29E-03 |
| ENSG00000090020 | SLC9A1 | 6548 | solute carrier family 9 member A1 | up | 2,29 | 9,81E-03 |
| ENSG00000179918 | SEPHS2 | 22928 | selenophosphate synthetase 2 | up | 2,29 | 7,53E-03 |
| ENSG00000154781 | CCDC174 | 51244 | coiled-coil domain containing 174 | up | 2,29 | 2,51E-03 |
| ENSG00000152503 | TRIM36 | 55521 | tripartite motif containing 36 | up | 2,29 | 6,64E-03 |
| ENSG00000135269 | TES | 26136 | testin LIM domain protein | up | 2,28 | 5,68E-03 |
| ENSG00000198205 | ZXDA | 7789 | zinc finger, X-linked, duplicated A | up | 2,28 | 8,42E-03 |
| ENSG00000152137 | HSPB8 | 26353 | heat shock protein family B (small) member 8 | up | 2,28 | 4,99E-03 |
| ENSG00000137962 | ARHGAP29 | 9411 | Rho GTPase activating protein 29 | up | 2,27 | 7,27E-03 |
| ENSG00000145050 | MANF | 7873 | mesencephalic astrocyte derived neurotrophic factor | up | 2,27 | 6,96E-03 |
| ENSG00000261824 | LINC00662 | 148189 | long intergenic non-protein coding RNA 662 | up | 2,25 | 6,61E-03 |
| ENSG00000197976 | AKAP17A | 8227 | A-kinase anchoring protein 17A | up | 2,25 | 5,76E-03 |
| ENSG00000186472 | PCLO | 27445 | piccolo presynaptic cytomatrix protein | up | 2,24 | 2,31E-02 |
| ENSG00000184205 | TSPYL2 | 64061 | TSPY like 2 | up | 2,24 | 3,19E-02 |
| ENSG00000185561 | TLCD2 | 727910 | TLC domain containing 2 | up | 2,23 | 5,52E-03 |
| ENSG00000170222 | ADPRM | 56985 | ADP-ribose/CDP-alcohol diphosphatase, manganese dependent | up | 2,23 | 2,73E-03 |
| ENSG00000134970 | TMED7 | 51014 | transmembrane p24 trafficking protein 7 | up | 2,23 | 4,98E-02 |
| ENSG00000135046 | ANXA1 | 301 | annexin A1 | up | 2,22 | 9,89E-03 |
| ENSG00000176428 | VPS37D | 155382 | VPS37D, ESCRT-I subunit | up | 2,22 | 5,84E-03 |
| ENSG00000074527 | NTN4 | 59277 | netrin 4 | up | 2,22 | 5,94E-03 |
| ENSG00000013375 | PGM3 | 5238 | phosphoglucomutase 3 | up | 2,22 | 4,78E-03 |
| ENSG00000198961 | PJA2 | 9867 | praja ring finger ubiquitin ligase 2 | up | 2,22 | 3,10E-03 |
| ENSG00000169826 | CSGALNACT2 | 55454 | chondroitin sulfate N-acetylgalactosaminyltransferase 2 | up | 2,21 | 1,16E-02 |
| ENSG00000185950 | IRS2 | 8660 | insulin receptor substrate 2 | up | 2,21 | 2,23E-02 |
| ENSG00000163449 | TMEM169 | 92691 | transmembrane protein 169 | up | 2,20 | 3,37E-03 |
| ENSG00000143498 | TAF1A | 9015 | TATA-box binding protein associated factor, RNA polymerase I subunit A | up | 2,20 | 2,24E-03 |
| ENSG00000148572 | NRBF2 | 29982 | nuclear receptor binding factor 2 | up | 2,19 | 2,71E-03 |
| ENSG00000214293 | APTR | 100505854 | Alu-mediated CDKN1A/p21 transcriptional regulator (non-protein coding) | up | 2,18 | 1,21E-02 |
| ENSG00000274605 | LOC105370333 | 105370333 | uncharacterized LOC105370333 | up | 2,18 | 8,98E-03 |
| ENSG00000163703 | CRELD1 | 78987 | cysteine rich with EGF like domains 1 | up | 2,17 | 3,26E-02 |
| ENSG00000101577 | LPIN2 | 9663 | lipin 2 | up | 2,16 | 6,88E-03 |
| ENSG00000078246 | TULP3 | 7289 | tubby like protein 3 | up | 2,16 | 7,18E-03 |
| ENSG00000132510 | KDM6B | 23135 | lysine demethylase 6B | up | 2,16 | 2,06E-02 |
| ENSG00000163644 | PPM1K | 152926 | protein phosphatase, Mg2+/Mn2+ dependent 1K | up | 2,16 | 3,87E-02 |
| ENSG00000134109 | EDEM1 | 9695 | ER degradation enhancing alpha-mannosidase like protein 1 | up | 2,16 | 1,32E-02 |
| ENSG00000164543 | STK17A | 9263 | serine/threonine kinase 17a | up | 2,16 | 2,71E-03 |
| ENSG00000181722 | ZBTB20 | 26137 | zinc finger and BTB domain containing 20 | up | 2,15 | 1,57E-02 |
| ENSG00000248932 | LOC100507291 | 100507291 | uncharacterized LOC100507291 | up | 2,15 | 2,19E-02 |
| ENSG00000111981 | ULBP1 | 80329 | UL16 binding protein 1 | up | 2,14 | 2,20E-02 |
| ENSG00000059769 | DNAJC25 | 548645 | DnaJ heat shock protein family (Hsp40) member C25 | up | 2,14 | 3,19E-03 |
| ENSG00000110925 | CSRNP2 | 81566 | cysteine and serine rich nuclear protein 2 | up | 2,14 | 3,62E-03 |
| ENSG00000128228 | SDF2L1 | 23753 | stromal cell derived factor 2 like 1 | up | 2,13 | 1,86E-02 |

| | | | | | | |
|---|---|---|---|---|---|---|
| ENSG00000153879 | CEBPG | 1054 | CCAAT/enhancer binding protein gamma | up | 2,13 | 5,68E-03 |
| ENSG00000163347 | CLDN1 | 9076 | claudin 1 | up | 2,13 | 6,96E-03 |
| ENSG00000165997 | ARL5B | 221079 | ADP ribosylation factor like GTPase 5B | up | 2,13 | 4,51E-03 |
| ENSG00000142871 | CYR61 | 3491 | cysteine rich angiogenic inducer 61 | up | 2,13 | 4,03E-03 |
| ENSG00000168994 | PXDC1 | 221749 | PX domain containing 1 | up | 2,12 | 1,53E-02 |
| ENSG00000172216 | CEBPB | 1051 | CCAAT/enhancer binding protein beta | up | 2,12 | 2,85E-02 |
| ENSG00000100784 | RPS6KA5 | 9252 | ribosomal protein S6 kinase A5 | up | 2,11 | 5,95E-03 |
| ENSG00000132950 | ZMYM5 | 9205 | zinc finger MYM-type containing 5 | up | 2,11 | 3,37E-03 |
| ENSG00000115641 | FHL2 | 2274 | four and a half LIM domains 2 | up | 2,10 | 1,13E-02 |
| ENSG00000001084 | GCLC | 2729 | glutamate-cysteine ligase catalytic subunit | up | 2,09 | 2,28E-02 |
| ENSG00000196083 | IL1RAP | 3556 | interleukin 1 receptor accessory protein | up | 2,09 | 9,73E-03 |
| ENSG00000148339 | SLC25A25 | 114789 | solute carrier family 25 member 25 | up | 2,09 | 5,76E-03 |
| ENSG00000105327 | BBC3 | 27113 | BCL2 binding component 3 | up | 2,09 | 3,18E-02 |
| ENSG00000013441 | CLK1 | 1195 | CDC like kinase 1 | up | 2,09 | 3,38E-02 |
| ENSG00000065911 | MTHFD2 | 10797 | methylenetetrahydrofolate dehydrogenase (NADP+ dependent) 2, metheny | up | 2,09 | 9,37E-03 |
| ENSG00000116584 | ARHGEF2 | 9181 | Rho/Rac guanine nucleotide exchange factor 2 | up | 2,09 | 1,11E-02 |
| ENSG00000090520 | DNAJB11 | 51726 | DnaJ heat shock protein family (Hsp40) member B11 | up | 2,08 | 1,18E-02 |
| ENSG00000106948 | AKNA | 80709 | AT-hook transcription factor | up | 2,07 | 1,00E-02 |
| ENSG00000197279 | ZNF165 | 7718 | zinc finger protein 165 | up | 2,07 | 2,14E-02 |
| ENSG00000134954 | ETS1 | 2113 | ETS proto-oncogene 1, transcription factor | up | 2,07 | 1,23E-02 |
| ENSG00000142867 | BCL10 | 8915 | B-cell CLL/lymphoma 10 | up | 2,06 | 5,26E-04 |
| ENSG00000146859 | TMEM140 | 55281 | transmembrane protein 140 | up | 2,05 | 2,11E-02 |
| ENSG00000102908 | NFAT5 | 10725 | nuclear factor of activated T-cells 5 | up | 2,05 | 1,13E-02 |
| ENSG00000184988 | TMEM106A | 113277 | transmembrane protein 106A | up | 2,04 | 2,36E-02 |
| ENSG00000118197 | DDX59 | 83479 | DEAD-box helicase 59 | up | 2,04 | 5,95E-03 |
| ENSG00000069956 | MAPK6 | 5597 | mitogen-activated protein kinase 6 | up | 2,04 | 5,55E-03 |
| ENSG00000171522 | PTGER4 | 5734 | prostaglandin E receptor 4 | up | 2,03 | 2,73E-03 |
| ENSG00000162616 | DNAJB4 | 11080 | DnaJ heat shock protein family (Hsp40) member B4 | up | 2,03 | 3,41E-03 |
| ENSG00000120278 | PLEKHG1 | 57480 | pleckstrin homology and RhoGEF domain containing G1 | up | 2,03 | 3,58E-02 |
| ENSG00000127528 | KLF2 | 10365 | Kruppel like factor 2 | up | 2,02 | 4,17E-02 |
| ENSG00000131871 | SELENOS | 55829 | selenoprotein S | up | 2,02 | 1,16E-02 |
| ENSG00000174738 | NR1D2 | 9975 | nuclear receptor subfamily 1 group D member 2 | up | 2,02 | 4,50E-03 |
| ENSG00000132002 | DNAJB1 | 3337 | DnaJ heat shock protein family (Hsp40) member B1 | up | 2,02 | 1,42E-02 |
| ENSG00000111912 | NCOA7 | 135112 | nuclear receptor coactivator 7 | up | 2,02 | 9,51E-03 |
| ENSG00000214050 | FBXO16 | 157574 | F-box protein 16 | up | 2,01 | 4,57E-02 |
| ENSG00000008311 | AASS | 10157 | aminoadipate-semialdehyde synthase | up | 2,01 | 4,96E-02 |
| ENSG00000141682 | PMAIP1 | 5366 | phorbol-12-myristate-13-acetate-induced protein 1 | up | 2,01 | 4,53E-02 |
| ENSG00000249345 | LINC02405 | 101927592 | long intergenic non-protein coding RNA 2405 | up | 2,01 | 2,19E-02 |
| ENSG00000075420 | FNDC3B | 64778 | fibronectin type III domain containing 3B | up | 2,00 | 2,16E-03 |
| ENSG00000144366 | GULP1 | 51454 | GULP, engulfment adaptor PTB domain containing 1 | up | 2,00 | 2,31E-02 |
| ENSG00000205189 | ZBTB10 | 65986 | zinc finger and BTB domain containing 10 | up | 2,00 | 2,07E-02 |
| ENSG00000100219 | XBP1 | 7494 | X-box binding protein 1 | up | 2,00 | 5,68E-03 |
| ENSG00000198018 | ENTPD7 | 57089 | ectonucleoside triphosphate diphosphohydrolase 7 | up | 2,00 | 2,10E-02 |
| ENSG00000151743 | AMN1 | 196394 | antagonist of mitotic exit network 1 homolog | up | 2,00 | 2,24E-03 |
| ENSG00000139289 | PHLDA1 | 22822 | pleckstrin homology like domain family A member 1 | up | 2,00 | 2,22E-02 |
| ENSG00000197608 | ZNF841 | 284371 | zinc finger protein 841 | up | 1,99 | 1,22E-02 |
| ENSG00000132823 | OSER1 | 51526 | oxidative stress responsive serine rich 1 | up | 1,99 | 5,44E-03 |
| ENSG00000213963 | LOC100130691 | 100130691 | uncharacterized LOC100130691 | up | 1,99 | 1,23E-02 |
| ENSG00000069869 | NEDD4 | 4734 | neural precursor cell expressed, developmentally down-regulated 4, E3 ubiq | up | 1,99 | 3,69E-03 |
| ENSG00000120129 | DUSP1 | 1843 | dual specificity phosphatase 1 | up | 1,99 | 2,51E-02 |
| ENSG00000144674 | GOLGA4 | 2803 | golgin A4 | up | 1,99 | 1,50E-03 |
| ENSG00000173812 | EIF1 | 10209 | eukaryotic translation initiation factor 1 | up | 1,98 | 9,37E-03 |
| ENSG00000232160 | RAP2C-AS1 | 101928578 | RAP2C antisense RNA 1 | up | 1,98 | 8,36E-03 |
| ENSG00000104231 | ZFAND1 | 79752 | zinc finger AN1-type containing 1 | up | 1,97 | 3,25E-03 |
| ENSG00000166579 | NDEL1 | 81565 | nudE neurodevelopment protein 1 like 1 | up | 1,97 | 6,96E-03 |
| ENSG00000068912 | ERLEC1 | 27248 | endoplasmic reticulum lectin 1 | up | 1,96 | 7,72E-03 |
| ENSG00000136826 | KLF4 | 9314 | Kruppel like factor 4 | up | 1,96 | 3,95E-02 |
| ENSG00000137331 | IER3 | 8870 | immediate early response 3 | up | 1,96 | 3,99E-02 |
| ENSG00000120742 | SERP1 | 27230 | stress associated endoplasmic reticulum protein 1 | up | 1,95 | 1,28E-02 |
| ENSG00000008086 | CDKL5 | 6792 | cyclin dependent kinase like 5 | up | 1,95 | 3,05E-02 |
| ENSG00000102531 | FNDC3A | 22862 | fibronectin type III domain containing 3A | up | 1,95 | 2,73E-03 |
| ENSG00000186376 | ZNF75D | 7626 | zinc finger protein 75D | up | 1,95 | 7,65E-03 |
| ENSG00000240303 | ACAD11 | 84129 | acyl-CoA dehydrogenase family member 11 | up | 1,95 | 3,58E-02 |
| ENSG00000243335 | KCTD7 | 154881 | potassium channel tetramerization domain containing 7 | up | 1,95 | 1,85E-03 |
| ENSG00000067082 | KLF6 | 1316 | Kruppel like factor 6 | up | 1,94 | 7,14E-03 |
| ENSG00000005700 | IBTK | 25998 | inhibitor of Bruton tyrosine kinase | up | 1,94 | 4,13E-03 |
| ENSG00000154305 | MIA3 | 375056 | MIA family member 3, ER export factor | up | 1,93 | 2,26E-03 |
| ENSG00000166471 | TMEM41B | 440026 | transmembrane protein 41B | up | 1,92 | 1,57E-02 |
| ENSG00000196182 | STK40 | 83931 | serine/threonine kinase 40 | up | 1,92 | 2,37E-02 |
| ENSG00000251192 | ZNF674 | 641339 | zinc finger protein 674 | up | 1,92 | 1,01E-02 |
| ENSG00000177410 | ZFAS1 | 441951 | ZNFX1 antisense RNA 1 | up | 1,92 | 1,42E-02 |
| ENSG00000137502 | RAB30 | 27314 | RAB30, member RAS oncogene family | up | 1,92 | 5,34E-03 |
| ENSG00000152409 | JMY | 133746 | junction mediating and regulatory protein, p53 cofactor | up | 1,91 | 2,54E-02 |
| ENSG00000164463 | CREBRF | 153222 | CREB3 regulatory factor | up | 1,91 | 8,94E-03 |
| ENSG00000130150 | MOSPD2 | 158747 | motile sperm domain containing 2 | up | 1,90 | 1,53E-03 |
| ENSG00000128016 | ZFP36 | 7538 | ZFP36 ring finger protein | up | 1,90 | 4,85E-02 |
| ENSG00000116604 | MEF2D | 4209 | myocyte enhancer factor 2D | up | 1,90 | 4,04E-02 |
| ENSG00000006459 | KDM7A | 80853 | lysine demethylase 7A | up | 1,89 | 3,98E-02 |
| ENSG00000115520 | COQ10B | 80219 | coenzyme Q10B | up | 1,89 | 7,17E-03 |
| ENSG00000156273 | BACH1 | 571 | BTB domain and CNC homolog 1 | up | 1,89 | 4,11E-03 |
| ENSG00000156273 | GRIK1-AS2 | 100379661 | GRIK1 antisense RNA 2 | up | 1,89 | 4,11E-03 |
| ENSG00000164951 | PDP1 | 54704 | pyruvate dehyrogenase phosphatase catalytic subunit 1 | up | 1,89 | 4,13E-03 |
| ENSG00000185015 | CA13 | 377677 | carbonic anhydrase 13 | up | 1,89 | 7,38E-03 |
| ENSG00000142627 | EPHA2 | 1969 | EPH receptor A2 | up | 1,89 | 2,51E-02 |
| ENSG00000140105 | WARS | 7453 | tryptophanyl-tRNA synthetase | up | 1,88 | 3,69E-02 |
| ENSG00000116044 | NFE2L2 | 4780 | nuclear factor, erythroid 2 like 2 | up | 1,88 | 2,41E-03 |
| ENSG00000161011 | SQSTM1 | 8878 | sequestosome 1 | up | 1,88 | 2,42E-02 |
| ENSG00000196110 | ZNF699 | 374879 | zinc finger protein 699 | up | 1,88 | 2,17E-02 |
| ENSG00000116675 | DNAJC6 | 9829 | DnaJ heat shock protein family (Hsp40) member C6 | up | 1,87 | 1,57E-02 |
| ENSG00000173540 | GMPPB | 29925 | GDP-mannose pyrophosphorylase B | up | 1,87 | 4,53E-02 |
| ENSG00000197019 | SERTAD1 | 29950 | SERTA domain containing 1 | up | 1,87 | 1,89E-02 |
| ENSG00000171988 | JMJD1C | 221037 | jumonji domain containing 1C | up | 1,87 | 6,39E-03 |
| ENSG00000212719 | C17orf51 | 339263 | chromosome 17 open reading frame 51 | up | 1,87 | 4,97E-02 |
| ENSG00000166012 | TAF1D | 79101 | TATA-box binding protein associated factor, RNA polymerase I subunit D | up | 1,87 | 1,01E-02 |
| ENSG00000136868 | SLC31A1 | 1317 | solute carrier family 31 member 1 | up | 1,86 | 1,00E-02 |
| ENSG00000066455 | GOLGA5 | 9950 | golgin A5 | up | 1,86 | 2,74E-03 |
| ENSG00000251136 | LOC101929709 | 101929709 | uncharacterized LOC101929709 | up | 1,86 | 3,30E-02 |
| ENSG00000173575 | CHD2 | 1106 | chromodomain helicase DNA binding protein 2 | up | 1,86 | 2,28E-02 |
| ENSG00000198833 | UBE2J1 | 51465 | ubiquitin conjugating enzyme E2 J1 | up | 1,86 | 8,98E-03 |
| ENSG00000106070 | GRB10 | 2887 | growth factor receptor bound protein 10 | up | 1,86 | 2,08E-02 |
| ENSG00000147535 | PLPP5 | 84513 | phospholipid phosphatase 5 | up | 1,85 | 4,16E-02 |
| ENSG00000167110 | GOLGA2 | 2801 | golgin A2 | up | 1,85 | 1,28E-02 |
| ENSG00000234741 | GAS5 | 60674 | growth arrest specific 5 (non-protein coding) | up | 1,85 | 2,75E-03 |
| ENSG00000173276 | ZBTB21 | 49854 | zinc finger and BTB domain containing 21 | up | 1,85 | 2,14E-03 |
| ENSG00000204498 | NFKBIL1 | 4795 | NFKB inhibitor like 1 | up | 1,85 | 2,93E-02 |
| ENSG00000109572 | CLCN3 | 1182 | chloride voltage-gated channel 3 | up | 1,85 | 9,81E-03 |
| ENSG00000148926 | ADM | 133 | adrenomedullin | up | 1,84 | 4,75E-04 |
| ENSG00000150403 | TMCO3 | 55002 | transmembrane and coiled-coil domains 3 | up | 1,84 | 1,70E-02 |
| ENSG00000101384 | JAG1 | 182 | jagged 1 | up | 1,84 | 3,28E-02 |
| ENSG00000143457 | GOLPH3L | 55204 | golgi phosphoprotein 3 like | up | 1,84 | 3,00E-03 |
| ENSG00000120686 | UFM1 | 51569 | ubiquitin fold modifier 1 | up | 1,84 | 1,26E-02 |
| ENSG00000131791 | PRKAB2 | 5565 | protein kinase AMP-activated non-catalytic subunit beta 2 | up | 1,83 | 1,28E-02 |
| ENSG00000143367 | TUFT1 | 7286 | tuftelin 1 | up | 1,83 | 9,81E-03 |
| ENSG00000114423 | CBLB | 868 | Cbl proto-oncogene B | up | 1,83 | 4,36E-02 |
| ENSG00000175395 | ZNF25 | 219749 | zinc finger protein 25 | up | 1,82 | 5,70E-04 |
| ENSG00000149260 | CAPN5 | 726 | calpain 5 | up | 1,82 | 1,33E-02 |
| ENSG00000136152 | COG3 | 83548 | component of oligomeric golgi complex 3 | up | 1,82 | 2,64E-03 |
| ENSG00000143067 | ZNF697 | 90874 | zinc finger protein 697 | up | 1,82 | 6,33E-03 |
| ENSG00000100485 | SOS2 | 6655 | SOS Ras/Rho guanine nucleotide exchange factor 2 | up | 1,82 | 1,41E-02 |
| ENSG00000031698 | SARS | 6301 | seryl-tRNA synthetase | up | 1,82 | 1,32E-02 |
| ENSG00000090861 | AARS | 16 | alanyl-tRNA synthetase | up | 1,82 | 1,85E-02 |
| ENSG00000196428 | TSC22D2 | 9819 | TSC22 domain family member 2 | up | 1,81 | 6,25E-03 |
| ENSG00000065802 | ASB1 | 51665 | ankyrin repeat and SOCS box containing 1 | up | 1,81 | 3,60E-02 |
| ENSG00000134107 | BHLHE40 | 8553 | basic helix-loop-helix family member e40 | up | 1,80 | 3,57E-02 |
| ENSG00000204387 | C6orf48 | 50854 | chromosome 6 open reading frame 48 | up | 1,80 | 1,39E-02 |
| ENSG00000160961 | ZNF333 | 84449 | zinc finger protein 333 | up | 1,80 | 2,51E-03 |
| ENSG00000107864 | CPEB3 | 22849 | cytoplasmic polyadenylation element binding protein 3 | up | 1,80 | 2,54E-02 |
| ENSG00000155158 | TTC39B | 158219 | tetratricopeptide repeat domain 39B | up | 1,80 | 2,24E-03 |
| ENSG00000269893 | SNHG8 | 100093630 | small nucleolar RNA host gene 8 | up | 1,80 | 7,61E-03 |
| ENSG00000151014 | NOCT | 25819 | nocturnin | up | 1,79 | 6,10E-03 |
| ENSG00000129472 | RAB2B | 84932 | RAB2B, member RAS oncogene family | up | 1,79 | 7,65E-03 |
| ENSG00000166348 | USP54 | 159195 | ubiquitin specific peptidase 54 | up | 1,79 | 3,77E-02 |
| ENSG00000283050 | GTF2IP12 | 101926918 | general transcription factor IIi pseudogene 12 | up | 1,79 | 4,28E-02 |
| ENSG00000139725 | RHOF | 54509 | ras homolog family member F, filopodia associated | up | 1,79 | 3,66E-02 |
| ENSG00000135709 | KIAA0513 | 9764 | KIAA0513 | up | 1,79 | 2,03E-02 |
| ENSG00000180747 | SMG1P3 | 100271836 | SMG1 pseudogene 3 | up | 1,79 | 4,28E-02 |
| ENSG00000115966 | ATF2 | 1386 | activating transcription factor 2 | up | 1,79 | 1,01E-02 |
| ENSG00000154447 | SH3RF1 | 57630 | SH3 domain containing ring finger 1 | up | 1,79 | 1,12E-02 |
| ENSG00000081087 | OSTM1 | 28962 | osteopetrosis associated transmembrane protein 1 | up | 1,78 | 9,61E-03 |
| ENSG00000116954 | RRAGC | 64121 | Ras related GTP binding C | up | 1,78 | 6,10E-03 |
| ENSG00000131016 | AKAP12 | 9590 | A-kinase anchoring protein 12 | up | 1,78 | 2,20E-02 |
| ENSG00000143622 | RIT1 | 6016 | Ras like without CAAX 1 | up | 1,78 | 1,44E-02 |
| ENSG00000167842 | MIS12 | 79003 | MIS12, kinetochore complex component | up | 1,78 | 1,58E-02 |
| ENSG00000197063 | MAFG | 4097 | MAF bZIP transcription factor G | up | 1,78 | 3,51E-02 |
| ENSG00000144566 | RAB5A | 5868 | RAB5A, member RAS oncogene family | up | 1,78 | 3,11E-03 |
| ENSG00000138835 | RGS3 | 5998 | regulator of G protein signaling 3 | up | 1,78 | 1,19E-02 |
| ENSG00000123091 | RNF11 | 26994 | ring finger protein 11 | up | 1,77 | 1,57E-02 |
| ENSG00000138678 | GPAT3 | 84803 | glycerol-3-phosphate acyltransferase 3 | up | 1,77 | 5,00E-02 |
| ENSG00000072364 | AFF4 | 27125 | AF4/FMR2 family member 4 | up | 1,77 | 8,70E-03 |
| ENSG00000147894 | C9orf72 | 203228 | chromosome 9 open reading frame 72 | up | 1,77 | 5,09E-03 |
| ENSG00000138768 | USO1 | 8615 | USO1 vesicle transport factor | up | 1,77 | 6,10E-03 |
| ENSG00000186300 | ZNF555 | 148254 | zinc finger protein 555 | up | 1,77 | 5,21E-03 |
| ENSG00000221883 | ARIH2OS | 646450 | ariadne RBR E3 ubiquitin protein ligase 2 opposite strand | up | 1,76 | 4,30E-02 |
| ENSG00000115425 | PECR | 55825 | peroxisomal trans-2-enoyl-CoA reductase | up | 1,76 | 4,64E-02 |
| ENSG00000203875 | SNHG5 | 387066 | small nucleolar RNA host gene 5 | up | 1,76 | 1,72E-02 |
| ENSG00000106610 | STAG3L4 | 64940 | stromal antigen 3-like 4 (pseudogene) | up | 1,76 | 3,18E-02 |
| ENSG00000179476 | C14orf28 | 122525 | chromosome 14 open reading frame 28 | up | 1,76 | 4,55E-02 |
| ENSG00000163697 | APBB2 | 323 | amyloid beta precursor protein binding family B member 2 | up | 1,76 | 8,58E-03 |
| ENSG00000145817 | YIPF5 | 81555 | Yip1 domain family member 5 | up | 1,76 | 1,33E-02 |
| ENSG00000138073 | PREB | 10113 | prolactin regulatory element binding | up | 1,76 | 3,49E-02 |
| ENSG00000135164 | DMTF1 | 9988 | cyclin D binding myb like transcription factor 1 | up | 1,76 | 4,26E-02 |
| ENSG00000115170 | ACVR1 | 90 | activin A receptor type 1 | up | 1,76 | 1,12E-02 |
| ENSG00000187735 | TCEA1 | 6917 | transcription elongation factor A1 | up | 1,76 | 1,82E-02 |
| ENSG00000221926 | TRIM16 | 10626 | tripartite motif containing 16 | up | 1,76 | 3,40E-02 |
| ENSG00000221926 | TRIM16L | 147166 | tripartite motif containing 16 like | up | 1,76 | 3,40E-02 |
| ENSG00000067167 | TRAM1 | 23471 | translocation associated membrane protein 1 | up | 1,76 | 4,37E-02 |
| ENSG00000115289 | PCGF1 | 84759 | polycomb group ring finger 1 | up | 1,76 | 9,57E-03 |
| ENSG00000111011 | RSRC2 | 65117 | arginine and serine rich coiled-coil 2 | up | 1,76 | 9,74E-03 |
| ENSG00000181982 | CCDC149 | 91050 | coiled-coil domain containing 149 | up | 1,75 | 4,01E-02 |
| ENSG00000115548 | KDM3A | 55818 | lysine demethylase 3A | up | 1,75 | 2,45E-02 |
| ENSG00000104497 | SNX16 | 64089 | sorting nexin 16 | up | 1,75 | 5,01E-03 |
| ENSG00000165813 | CCDC186 | 55088 | coiled-coil domain containing 186 | up | 1,75 | 8,08E-03 |
| ENSG00000196470 | SIAH1 | 6477 | siah E3 ubiquitin protein ligase 1 | up | 1,74 | 1,66E-02 |
| ENSG00000213047 | DENND1B | 163486 | DENN domain containing 1B | up | 1,74 | 1,66E-02 |
| ENSG00000169239 | CA5B | 11238 | carbonic anhydrase 5B | up | 1,74 | 2,94E-02 |
| ENSG00000173482 | PTPRM | 5797 | protein tyrosine phosphatase, receptor type M | up | 1,74 | 7,91E-03 |
| ENSG00000047346 | FAM214A | 56204 | family with sequence similarity 214 member A | up | 1,74 | 3,23E-02 |
| ENSG00000115963 | RND3 | 390 | Rho family GTPase 3 | up | 1,73 | 3,10E-03 |
| ENSG00000107290 | SETX | 23064 | senataxin | up | 1,73 | 1,47E-02 |
| ENSG00000124783 | SSR1 | 6745 | signal sequence receptor subunit 1 | up | 1,73 | 2,06E-02 |
| ENSG00000163512 | AZI2 | 64343 | 5-azacytidine induced 2 | up | 1,73 | 9,03E-03 |
| ENSG00000104957 | CCDC130 | 81576 | coiled-coil domain containing 130 | up | 1,73 | 2,43E-02 |
| ENSG00000120519 | SLC10A7 | 84068 | solute carrier family 10 member 7 | up | 1,73 | 3,01E-02 |
| ENSG00000101745 | ANKRD12 | 23253 | ankyrin repeat domain 12 | up | 1,73 | 1,36E-02 |
| ENSG00000177888 | ZBTB41 | 360023 | zinc finger and BTB domain containing 41 | up | 1,72 | 2,11E-02 |
| ENSG00000215126 | CBWD2 | 150472 | COBW domain containing 2 | up | 1,72 | 4,61E-02 |
| ENSG00000215126 | CBWD6 | 644019 | COBW domain containing 6 | up | 1,72 | 4,61E-02 |
| ENSG00000163867 | ZMYM6 | 9204 | zinc finger MYM-type containing 6 | up | 1,72 | 1,64E-02 |
| ENSG00000172007 | RAB33B | 83452 | RAB33B, member RAS oncogene family | up | 1,72 | 1,79E-02 |
| ENSG00000204152 | TIMM23B | 100652748 | translocase of inner mitochondrial membrane 23 homolog B | up | 1,72 | 2,21E-02 |
| ENSG00000104312 | RIPK2 | 8767 | receptor interacting serine/threonine kinase 2 | up | 1,72 | 8,32E-03 |
| ENSG00000131503 | ANKHD1 | 54882 | ankyrin repeat and KH domain containing 1 | up | 1,72 | 1,81E-02 |
| ENSG00000088451 | TGDS | 23483 | TDP-glucose 4,6-dehydratase | up | 1,72 | 2,21E-02 |
| ENSG00000114120 | SLC25A36 | 55186 | solute carrier family 25 member 36 | up | 1,72 | 1,13E-02 |
| ENSG00000169359 | SLC33A1 | 9197 | solute carrier family 33 member 1 | up | 1,72 | 2,24E-02 |
| ENSG00000130779 | CLIP1 | 6249 | CAP-Gly domain containing linker protein 1 | up | 1,72 | 1,43E-03 |
| ENSG00000179361 | ARID3B | 10620 | AT-rich interaction domain 3B | up | 1,72 | 4,67E-02 |
| ENSG00000166598 | HSP90B1 | 7184 | heat shock protein 90 beta family member 1 | up | 1,71 | 2,12E-02 |
| ENSG00000101310 | SEC23B | 10483 | Sec23 homolog B, coat complex II component | up | 1,71 | 1,74E-02 |
| ENSG00000144747 | TMF1 | 7110 | TATA element modulatory factor 1 | up | 1,71 | 3,05E-03 |
| ENSG00000113615 | SEC24A | 10802 | SEC24 homolog A, COPII coat complex component | up | 1,71 | 3,96E-03 |
| ENSG00000156639 | ZFAND3 | 60685 | zinc finger AN1-type containing 3 | up | 1,71 | 2,58E-02 |
| ENSG00000147421 | HMBOX1 | 79618 | homeobox containing 1 | up | 1,71 | 2,25E-02 |
| ENSG00000111727 | HCFC2 | 29915 | host cell factor C2 | up | 1,71 | 9,52E-03 |
| ENSG00000177683 | THAP5 | 168451 | THAP domain containing 5 | up | 1,71 | 2,61E-02 |
| ENSG00000143669 | LYST | 1130 | lysosomal trafficking regulator | up | 1,70 | 2,05E-02 |
| ENSG00000075643 | MOCOS | 55034 | molybdenum cofactor sulfurase | up | 1,70 | 2,55E-02 |
| ENSG00000182827 | ACBD3 | 64746 | acyl-CoA binding domain containing 3 | up | 1,70 | 3,10E-03 |
| ENSG00000139793 | MBNL2 | 10150 | muscleblind like splicing regulator 2 | up | 1,70 | 4,92E-03 |
| ENSG00000136603 | SKIL | 6498 | SKI like proto-oncogene | up | 1,70 | 1,92E-02 |
| ENSG00000139517 | LNX2 | 222484 | ligand of numb-protein X 2 | up | 1,70 | 1,86E-02 |
| ENSG00000182400 | TRAPPC6B | 122553 | trafficking protein particle complex 6B | up | 1,70 | 3,07E-02 |
| ENSG00000010818 | HIVEP2 | 3097 | human immunodeficiency virus type I enhancer binding protein 2 | up | 1,69 | 1,13E-02 |
| ENSG00000147905 | ZCCHC7 | 84186 | zinc finger CCHC-type containing 7 | up | 1,69 | 8,32E-03 |
| ENSG00000197647 | ZNF433 | 163059 | zinc finger protein 433 | up | 1,69 | 5,68E-03 |
| ENSG00000168374 | ARF4 | 378 | ADP ribosylation factor 4 | up | 1,69 | 2,56E-02 |
| ENSG00000066651 | TRMT11 | 60487 | tRNA methyltransferase 11 homolog | up | 1,68 | 6,36E-03 |

| | | | | | | |
|---|---|---|---|---|---|---|
| ENSG00000137955 | RABGGTB | 5876 | Rab geranylgeranyltransferase beta subunit | up | 1,68 | 4,78E-03 |
| ENSG00000072849 | DERL2 | 51009 | derlin 2 | up | 1,68 | 4,88E-02 |
| ENSG00000118217 | ATF6 | 22926 | activating transcription factor 6 | up | 1,68 | 1,14E-02 |
| ENSG00000107854 | TNKS2 | 80351 | tankyrase 2 | up | 1,68 | 3,03E-03 |
| ENSG00000171928 | TVP23B | 51030 | trans-golgi network vesicle protein 23 homolog B | up | 1,68 | 1,21E-02 |
| ENSG00000153037 | SRP19 | 6728 | signal recognition particle 19 | up | 1,68 | 4,13E-02 |
| ENSG00000156976 | EIF4A2 | 1974 | eukaryotic translation initiation factor 4A2 | up | 1,68 | 4,70E-03 |
| ENSG00000187792 | ZNF70 | 7621 | zinc finger protein 70 | up | 1,68 | 4,63E-02 |
| ENSG00000198496 | NBR2 | 10230 | neighbor of BRCA1 gene 2 (non-protein coding) | up | 1,68 | 1,63E-02 |
| ENSG00000120063 | GNA13 | 10672 | G protein subunit alpha 13 | up | 1,68 | 2,41E-03 |
| ENSG00000119771 | KLHL29 | 114818 | kelch like family member 29 | up | 1,67 | 4,47E-02 |
| ENSG00000071243 | ING3 | 54556 | inhibitor of growth family member 3 | up | 1,67 | 2,14E-02 |
| ENSG00000073008 | PVR | 5817 | poliovirus receptor | up | 1,67 | 4,18E-02 |
| ENSG00000133794 | ARNTL | 406 | aryl hydrocarbon receptor nuclear translocator like | up | 1,67 | 5,95E-03 |
| ENSG00000138380 | CARF | 79800 | calcium responsive transcription factor | up | 1,67 | 8,22E-03 |
| ENSG00000164983 | TMEM65 | 157378 | transmembrane protein 65 | up | 1,67 | 4,07E-02 |
| ENSG00000165312 | OTUD1 | 220213 | OTU deubiquitinase 1 | up | 1,67 | 3,89E-02 |
| ENSG00000162236 | STX5 | 6811 | syntaxin 5 | up | 1,66 | 2,40E-02 |
| ENSG00000172164 | SNTB1 | 6641 | syntrophin beta 1 | up | 1,66 | 8,86E-03 |
| ENSG00000143256 | PFDN2 | 5202 | prefoldin subunit 2 | up | 1,66 | 1,44E-02 |
| ENSG00000181264 | TMEM136 | 219902 | transmembrane protein 136 | up | 1,66 | 1,61E-02 |
| ENSG00000125812 | GZF1 | 64412 | GDNF inducible zinc finger protein 1 | up | 1,66 | 1,31E-02 |
| ENSG00000187866 | FAM122A | 116224 | family with sequence similarity 122A | up | 1,66 | 1,82E-02 |
| ENSG00000023318 | ERP44 | 23071 | endoplasmic reticulum protein 44 | up | 1,66 | 2,10E-02 |
| ENSG00000223547 | ZNF844 | 284391 | zinc finger protein 844 | up | 1,65 | 2,03E-02 |
| ENSG00000106105 | GARS | 2617 | glycyl-tRNA synthetase | up | 1,65 | 1,80E-02 |
| ENSG00000143878 | RHOB | 388 | ras homolog family member B | up | 1,65 | 2,44E-02 |
| ENSG00000119048 | UBE2B | 7320 | ubiquitin conjugating enzyme E2 B | up | 1,65 | 4,38E-02 |
| ENSG00000101782 | RIOK3 | 8780 | RIO kinase 3 | up | 1,65 | 1,28E-02 |
| ENSG00000188994 | ZNF292 | 23036 | zinc finger protein 292 | up | 1,64 | 9,49E-03 |
| ENSG00000144468 | RHBDD1 | 84236 | rhomboid domain containing 1 | up | 1,64 | 9,81E-03 |
| ENSG00000168386 | FILIP1L | 11259 | filamin A interacting protein 1 like | up | 1,64 | 4,88E-02 |
| ENSG00000155313 | USP25 | 29761 | ubiquitin specific peptidase 25 | up | 1,64 | 2,64E-03 |
| ENSG00000167766 | ZNF83 | 55769 | zinc finger protein 83 | up | 1,64 | 2,73E-03 |
| ENSG00000089177 | KIF16B | 55614 | kinesin family member 16B | up | 1,64 | 5,27E-03 |
| ENSG00000066855 | MTFR1 | 9650 | mitochondrial fission regulator 1 | up | 1,63 | 2,19E-02 |
| ENSG00000162994 | CLHC1 | 130162 | clathrin heavy chain linker domain containing 1 | up | 1,63 | 4,18E-02 |
| ENSG00000138772 | ANXA3 | 306 | annexin A3 | up | 1,63 | 1,58E-02 |
| ENSG00000198853 | RUSC2 | 9853 | RUN and SH3 domain containing 2 | up | 1,63 | 3,73E-02 |
| ENSG00000086544 | ITPKC | 80271 | inositol-trisphosphate 3-kinase C | up | 1,63 | 3,33E-02 |
| ENSG00000130449 | ZSWIM6 | 57688 | zinc finger SWIM-type containing 6 | up | 1,63 | 2,36E-02 |
| ENSG00000144182 | LIPT1 | 51601 | lipoyltransferase 1 | up | 1,63 | 1,74E-02 |
| ENSG00000179454 | KLHL28 | 54813 | kelch like family member 28 | up | 1,63 | 3,99E-02 |
| ENSG00000198707 | CEP290 | 80184 | centrosomal protein 290 | up | 1,63 | 1,81E-02 |
| ENSG00000137574 | TGS1 | 96764 | trimethylguanosine synthase 1 | up | 1,63 | 9,37E-03 |
| ENSG00000167528 | ZNF641 | 121274 | zinc finger protein 641 | up | 1,63 | 1,31E-02 |
| ENSG00000119004 | CYP20A1 | 57404 | cytochrome P450 family 20 subfamily A member 1 | up | 1,62 | 2,88E-02 |
| ENSG00000008294 | SPAG9 | 9043 | sperm associated antigen 9 | up | 1,62 | 6,50E-03 |
| ENSG00000134294 | SLC38A2 | 54407 | solute carrier family 38 member 2 | up | 1,62 | 4,57E-02 |
| ENSG00000114331 | ACAP2 | 23527 | ArfGAP with coiled-coil, ankyrin repeat and PH domains 2 | up | 1,62 | 6,27E-03 |
| ENSG00000121964 | GTDC1 | 79712 | glycosyltransferase like domain containing 1 | up | 1,62 | 1,50E-02 |
| ENSG00000148426 | PROSER2 | 254427 | proline and serine rich 2 | up | 1,62 | 2,60E-02 |
| ENSG00000080822 | CLDND1 | 56650 | claudin domain containing 1 | up | 1,62 | 8,55E-03 |
| ENSG00000173230 | GOLGB1 | 2804 | golgin B1 | up | 1,62 | 4,77E-04 |
| ENSG00000114784 | EIF1B | 10289 | eukaryotic translation initiation factor 1B | up | 1,61 | 1,63E-02 |
| ENSG00000104375 | STK3 | 6788 | serine/threonine kinase 3 | up | 1,61 | 2,09E-03 |
| ENSG00000119523 | ALG2 | 85365 | ALG2, alpha-1,3/1,6-mannosyltransferase | up | 1,61 | 2,75E-02 |
| ENSG00000121350 | PYROXD1 | 79912 | pyridine nucleotide-disulphide oxidoreductase domain 1 | up | 1,61 | 1,61E-02 |
| ENSG00000139496 | NUP58 | 9818 | nucleoporin 58 | up | 1,61 | 1,29E-02 |
| ENSG00000111897 | SERINC1 | 57515 | serine incorporator 1 | up | 1,61 | 1,07E-02 |
| ENSG00000119801 | YPEL5 | 51646 | yippee like 5 | up | 1,61 | 3,95E-02 |
| ENSG00000170448 | NFXL1 | 152518 | nuclear transcription factor, X-box binding like 1 | up | 1,61 | 7,69E-03 |
| ENSG00000174373 | RALGAPA1 | 253959 | Ral GTPase activating protein catalytic alpha subunit 1 | up | 1,61 | 7,17E-03 |
| ENSG00000186432 | KPNA4 | 3840 | karyopherin subunit alpha 4 | up | 1,61 | 4,03E-03 |
| ENSG00000020577 | SAMD4A | 23034 | sterile alpha motif domain containing 4A | up | 1,61 | 1,95E-02 |
| ENSG00000112763 | BTN2A1 | 11120 | butyrophilin subfamily 2 member A1 | up | 1,61 | 1,62E-02 |
| ENSG00000196911 | KPNA5 | 3841 | karyopherin subunit alpha 5 | up | 1,61 | 1,31E-02 |
| ENSG00000119711 | ALDH6A1 | 4329 | aldehyde dehydrogenase 6 family member A1 | up | 1,60 | 3,51E-02 |
| ENSG00000133816 | MICAL2 | 9645 | microtubule associated monooxygenase, calponin and LIM domain containi | up | 1,60 | 3,30E-02 |
| ENSG00000219626 | FAM228B | 375190 | family with sequence similarity 228 member B | up | 1,60 | 1,66E-02 |
| ENSG00000139132 | FGD4 | 121512 | FYVE, RhoGEF and PH domain containing 4 | up | 1,60 | 1,35E-02 |
| ENSG00000274211 | SOCS7 | 30837 | suppressor of cytokine signaling 7 | up | 1,60 | 3,58E-02 |
| ENSG00000025796 | SEC63 | 11231 | SEC63 homolog, protein translocation regulator | up | 1,60 | 8,90E-03 |
| ENSG00000096717 | SIRT1 | 23411 | sirtuin 1 | up | 1,60 | 1,00E-02 |
| ENSG00000113013 | HSPA9 | 3313 | heat shock protein family A (Hsp70) member 9 | up | 1,60 | 5,69E-03 |
| ENSG00000181852 | RNF41 | 10193 | ring finger protein 41 | up | 1,59 | 2,87E-02 |
| ENSG00000226864 | ATE1-AS1 | 100130887 | ATE1 antisense RNA 1 | up | 1,59 | 4,84E-02 |
| ENSG00000179912 | R3HDM2 | 22864 | R3H domain containing 2 | up | 1,59 | 5,78E-03 |
| ENSG00000105856 | HBP1 | 26959 | HMG-box transcription factor 1 | up | 1,59 | 3,18E-02 |
| ENSG00000244405 | ETV5 | 2119 | ETS variant 5 | up | 1,59 | 2,20E-02 |
| ENSG00000168939 | SPRY3 | 10251 | sprouty RTK signaling antagonist 3 | up | 1,59 | 3,58E-02 |
| ENSG00000076053 | RBM7 | 10179 | RNA binding motif protein 7 | up | 1,58 | 7,14E-03 |
| ENSG00000184575 | XPOT | 11260 | exportin for tRNA | up | 1,58 | 1,21E-02 |
| ENSG00000184009 | ACTG1 | 71 | actin gamma 1 | up | 1,58 | 1,66E-02 |
| ENSG00000140743 | LOC101060399 | 101060399 | cerebellar degeneration-related protein 2 | up | 1,58 | 1,01E-02 |
| ENSG00000117616 | RSRP1 | 57035 | arginine and serine rich protein 1 | up | 1,58 | 4,00E-02 |
| ENSG00000197044 | ZNF441 | 126068 | zinc finger protein 441 | up | 1,58 | 3,64E-02 |
| ENSG00000140743 | CDR2 | 1039 | cerebellar degeneration related protein 2 | up | 1,58 | 1,01E-02 |
| ENSG00000137996 | RTCA | 8634 | RNA 3'-terminal phosphate cyclase | up | 1,57 | 1,00E-02 |
| ENSG00000196365 | LONP1 | 9361 | lon peptidase 1, mitochondrial | up | 1,57 | 4,38E-02 |
| ENSG00000176597 | B3GNT5 | 84002 | UDP-GlcNAc:betaGal beta-1,3-N-acetylglucosaminyltransferase 5 | up | 1,57 | 1,70E-02 |
| ENSG00000186952 | TMEM232 | 642987 | transmembrane protein 232 | up | 1,57 | 4,47E-02 |
| ENSG00000164975 | SNAPC3 | 6619 | small nuclear RNA activating complex polypeptide 3 | up | 1,57 | 2,08E-03 |
| ENSG00000204256 | BRD2 | 6046 | bromodomain containing 2 | up | 1,57 | 3,67E-02 |
| ENSG00000174796 | THAP6 | 152815 | THAP domain containing 6 | up | 1,57 | 1,11E-02 |
| ENSG00000187801 | ZFP69B | 65243 | ZFP69 zinc finger protein B | up | 1,57 | 2,18E-02 |
| ENSG00000084764 | MAPRE3 | 22924 | microtubule associated protein RP/EB family member 3 | up | 1,57 | 2,61E-02 |
| ENSG00000121073 | SLC35B1 | 10237 | solute carrier family 35 member B1 | up | 1,57 | 4,81E-02 |
| ENSG00000116752 | BCAS2 | 10286 | BCAS2, pre-mRNA processing factor | up | 1,57 | 2,14E-02 |
| ENSG00000166037 | CEP57 | 9702 | centrosomal protein 57 | up | 1,57 | 1,19E-02 |
| ENSG00000171365 | CLCN5 | 1184 | chloride voltage-gated channel 5 | up | 1,57 | 9,66E-03 |
| ENSG00000141441 | GAREM1 | 64762 | GRB2 associated regulator of MAPK1 subtype 1 | up | 1,57 | 4,54E-02 |
| ENSG00000100592 | DAAM1 | 23002 | dishevelled associated activator of morphogenesis 1 | up | 1,56 | 1,64E-02 |
| ENSG00000107771 | CCSER2 | 54462 | coiled-coil serine rich protein 2 | up | 1,56 | 1,20E-02 |
| ENSG00000034677 | RNF19A | 25897 | ring finger protein 19A, RBR E3 ubiquitin protein ligase | up | 1,56 | 8,54E-03 |
| ENSG00000109046 | WSB1 | 26118 | WD repeat and SOCS box containing 1 | up | 1,56 | 2,11E-02 |
| ENSG00000198342 | ZNF442 | 79973 | zinc finger protein 442 | up | 1,56 | 2,53E-02 |
| ENSG00000224531 | SMIM13 | 221710 | small integral membrane protein 13 | up | 1,56 | 6,88E-03 |
| ENSG00000130522 | JUND | 3727 | JunD proto-oncogene, AP-1 transcription factor subunit | up | 1,56 | 4,42E-02 |
| ENSG00000120370 | GORAB | 92344 | golgin, RAB6 interacting | up | 1,56 | 2,85E-02 |
| ENSG00000132256 | TRIM5 | 85363 | tripartite motif containing 5 | up | 1,56 | 4,95E-02 |
| ENSG00000108433 | GOSR2 | 9570 | golgi SNAP receptor complex member 2 | up | 1,55 | 2,21E-02 |
| ENSG00000167767 | KRT80 | 144501 | keratin 80 | up | 1,55 | 4,17E-02 |
| ENSG00000068305 | MEF2A | 4205 | myocyte enhancer factor 2A | up | 1,55 | 1,41E-02 |
| ENSG00000124788 | ATXN1 | 6310 | ataxin 1 | up | 1,55 | 2,07E-02 |
| ENSG00000131931 | THAP1 | 55145 | THAP domain containing 1 | up | 1,55 | 2,21E-02 |
| ENSG00000023287 | RB1CC1 | 9821 | RB1 inducible coiled-coil 1 | up | 1,55 | 2,96E-02 |
| ENSG00000110619 | CARS | 833 | cysteinyl-tRNA synthetase | up | 1,55 | 4,39E-02 |
| ENSG00000162695 | SLC30A7 | 148867 | solute carrier family 30 member 7 | up | 1,55 | 2,39E-02 |
| ENSG00000108061 | SHOC2 | 8036 | SHOC2, leucine rich repeat scaffold protein | up | 1,54 | 9,37E-03 |
| ENSG00000177425 | PAWR | 5074 | pro-apoptotic WT1 regulator | up | 1,54 | 9,76E-03 |
| ENSG00000164244 | PRRC1 | 133619 | proline rich coiled-coil 1 | up | 1,54 | 1,00E-02 |
| ENSG00000175691 | ZNF77 | 58492 | zinc finger protein 77 | up | 1,54 | 2,97E-02 |
| ENSG00000116874 | WARS2 | 10352 | tryptophanyl tRNA synthetase 2, mitochondrial | up | 1,54 | 9,55E-03 |
| ENSG00000148690 | FRA10AC1 | 118924 | FRA10A associated CGG repeat 1 | up | 1,54 | 9,89E-03 |
| ENSG00000153989 | NUS1 | 116150 | NUS1 dehydrodolichyl diphosphate synthase subunit | up | 1,54 | 4,28E-02 |
| ENSG00000167565 | SERTAD3 | 29946 | SERTA domain containing 3 | up | 1,54 | 5,95E-03 |
| ENSG00000178397 | FAM220A | 84792 | family with sequence similarity 220 member A | up | 1,54 | 1,11E-02 |
| ENSG00000074695 | LMAN1 | 3998 | lectin, mannose binding 1 | up | 1,54 | 4,15E-02 |
| ENSG00000102158 | MAGT1 | 84061 | magnesium transporter 1 | up | 1,54 | 4,98E-02 |
| ENSG00000182831 | C16orf72 | 29035 | chromosome 16 open reading frame 72 | up | 1,54 | 1,70E-02 |
| ENSG00000092108 | SCFD1 | 23256 | sec1 family domain containing 1 | up | 1,54 | 1,06E-02 |
| ENSG00000107372 | ZFAND5 | 7763 | zinc finger AN1-type containing 5 | up | 1,54 | 2,26E-02 |
| ENSG00000138032 | PPM1B | 5495 | protein phosphatase, Mg2+/Mn2+ dependent 1B | up | 1,53 | 2,84E-02 |
| ENSG00000158615 | PPP1R15B | 84919 | protein phosphatase 1 regulatory subunit 15B | up | 1,53 | 2,64E-02 |
| ENSG00000198839 | ZNF277 | 11179 | zinc finger protein 277 | up | 1,53 | 1,28E-02 |
| ENSG00000037637 | FBXO42 | 54455 | F-box protein 42 | up | 1,53 | 2,76E-02 |
| ENSG00000119328 | FAM206A | 54942 | family with sequence similarity 206 member A | up | 1,53 | 1,23E-02 |
| ENSG00000124151 | NCOA3 | 8202 | nuclear receptor coactivator 3 | up | 1,53 | 4,14E-02 |
| ENSG00000138674 | SEC31A | 22872 | SEC31 homolog A, COPII coat complex component | up | 1,53 | 3,03E-02 |
| ENSG00000144597 | EAF1 | 85403 | ELL associated factor 1 | up | 1,53 | 2,92E-02 |
| ENSG00000158079 | PTPDC1 | 138639 | protein tyrosine phosphatase domain containing 1 | up | 1,53 | 1,40E-02 |
| ENSG00000162434 | JAK1 | 3716 | Janus kinase 1 | up | 1,53 | 2,13E-02 |
| ENSG00000106993 | CDC37L1 | 55664 | cell division cycle 37 like 1 | up | 1,53 | 1,70E-02 |
| ENSG00000121749 | TBC1D15 | 64786 | TBC1 domain family member 15 | up | 1,53 | 2,01E-02 |
| ENSG00000185917 | SETD4 | 54093 | SET domain containing 4 | up | 1,53 | 3,81E-02 |
| ENSG00000109814 | UGDH | 7358 | UDP-glucose 6-dehydrogenase | up | 1,53 | 3,50E-02 |
| ENSG00000110048 | OSBP | 5007 | oxysterol binding protein | up | 1,53 | 1,86E-02 |
| ENSG00000136021 | SCYL2 | 55681 | SCY1 like pseudokinase 2 | up | 1,53 | 9,48E-03 |
| ENSG00000169249 | ZRSR2 | 8233 | zinc finger CCCH-type, RNA binding motif and serine/arginine rich 2 | up | 1,53 | 1,68E-02 |
| ENSG00000138078 | PREPL | 9581 | prolyl endopeptidase-like | up | 1,53 | 2,61E-02 |
| ENSG00000120910 | PPP3CC | 5533 | protein phosphatase 3 catalytic subunit gamma | up | 1,53 | 1,79E-02 |
| ENSG00000160570 | DEDD2 | 162989 | death effector domain containing 2 | up | 1,53 | 2,03E-02 |
| ENSG00000248049 | UBA6-AS1 | 550112 | UBA6 antisense RNA 1 (head to head) | up | 1,53 | 1,27E-02 |
| ENSG00000187240 | DYNC2H1 | 79659 | dynein cytoplasmic 2 heavy chain 1 | up | 1,52 | 9,52E-03 |
| ENSG00000085433 | WDR47 | 22911 | WD repeat domain 47 | up | 1,52 | 1,28E-03 |
| ENSG00000134352 | IL6ST | 3572 | interleukin 6 signal transducer | up | 1,52 | 1,96E-02 |
| ENSG00000162971 | TYW5 | 129450 | tRNA-yW synthesizing protein 5 | up | 1,52 | 3,54E-02 |
| ENSG00000196792 | STRN3 | 29966 | striatin 3 | up | 1,52 | 3,66E-03 |
| ENSG00000013561 | RNF14 | 9604 | ring finger protein 14 | up | 1,52 | 2,07E-02 |
| ENSG00000141446 | ESCO1 | 114799 | establishment of sister chromatid cohesion N-acetyltransferase 1 | up | 1,52 | 2,97E-02 |
| ENSG00000168917 | SLC35G2 | 80723 | solute carrier family 35 member G2 | up | 1,52 | 3,69E-03 |
| ENSG00000175471 | MCTP1 | 79772 | multiple C2 and transmembrane domain containing 1 | up | 1,52 | 4,42E-02 |
| ENSG00000073331 | ALPK1 | 80216 | alpha kinase 1 | up | 1,52 | 1,00E-02 |
| ENSG00000164823 | OSGIN2 | 734 | oxidative stress induced growth inhibitor family member 2 | up | 1,52 | 1,79E-02 |
| ENSG00000111696 | NT5DC3 | 51559 | 5'-nucleotidase domain containing 3 | up | 1,52 | 4,75E-02 |
| ENSG00000168495 | POLR3D | 661 | RNA polymerase III subunit D | up | 1,52 | 2,72E-02 |
| ENSG00000151929 | BAG3 | 9531 | BCL2 associated athanogene 3 | up | 1,52 | 4,07E-02 |
| ENSG00000157764 | BRAF | 673 | B-Raf proto-oncogene, serine/threonine kinase | up | 1,52 | 4,53E-02 |
| ENSG00000145685 | LHFPL2 | 10184 | LHFPL tetraspan subfamily member 2 | up | 1,52 | 1,21E-02 |
| ENSG00000113734 | BNIP1 | 662 | BCL2 interacting protein 1 | up | 1,52 | 2,19E-02 |
| ENSG00000110851 | PRDM4 | 11108 | PR/SET domain 4 | up | 1,52 | 2,92E-02 |
| ENSG00000066427 | ATXN3 | 4287 | ataxin 3 | up | 1,51 | 4,51E-03 |
| ENSG00000143751 | SDE2 | 163859 | SDE2 telomere maintenance homolog | up | 1,51 | 1,00E-02 |
| ENSG00000163602 | RYBP | 23429 | RING1 and YY1 binding protein | up | 1,51 | 1,40E-02 |
| ENSG00000014123 | UFL1 | 23376 | UFM1 specific ligase 1 | up | 1,51 | 1,67E-02 |
| ENSG00000132912 | DCTN4 | 51164 | dynactin subunit 4 | up | 1,51 | 1,31E-02 |
| ENSG00000131013 | PPIL4 | 85313 | peptidylprolyl isomerase like 4 | up | 1,51 | 4,67E-03 |
| ENSG00000107249 | GLIS3 | 169792 | GLIS family zinc finger 3 | up | 1,51 | 4,38E-02 |
| ENSG00000118260 | CREB1 | 1385 | cAMP responsive element binding protein 1 | up | 1,51 | 7,18E-03 |
| ENSG00000142459 | EVI5L | 115704 | ecotropic viral integration site 5 like | up | 1,51 | 4,73E-02 |
| ENSG00000115993 | TRAK2 | 66008 | trafficking kinesin protein 2 | up | 1,51 | 2,96E-02 |
| ENSG00000211455 | STK38L | 23012 | serine/threonine kinase 38 like | up | 1,51 | 1,00E-02 |
| ENSG00000141428 | C18orf21 | 83608 | chromosome 18 open reading frame 21 | up | 1,51 | 1,53E-02 |
| ENSG00000078747 | ITCH | 83737 | itchy E3 ubiquitin protein ligase | up | 1,51 | 6,64E-03 |
| ENSG00000116668 | SWT1 | 54823 | SWT1, RNA endoribonuclease homolog | up | 1,51 | 4,63E-02 |
| ENSG00000163659 | TIPARP | 25976 | TCDD inducible poly(ADP-ribose) polymerase | up | 1,51 | 3,69E-02 |
| ENSG00000169379 | ARL13B | 200894 | ADP ribosylation factor like GTPase 13B | up | 1,51 | 1,10E-02 |
| ENSG00000196639 | HRH1 | 3269 | histamine receptor H1 | up | 1,51 | 2,04E-02 |
| ENSG00000198900 | TOP1 | 7150 | DNA topoisomerase I | up | 1,51 | 1,95E-02 |
| ENSG00000106392 | C1GALT1 | 56913 | core 1 synthase, glycoprotein-N-acetylgalactosamine 3-beta-galactosyltransf | up | 1,51 | 1,70E-02 |
| ENSG00000089818 | NECAP1 | 25977 | NECAP endocytosis associated 1 | up | 1,50 | 1,94E-02 |
| ENSG00000220205 | VAMP2 | 6844 | vesicle associated membrane protein 2 | up | 1,50 | 4,83E-02 |
| ENSG00000181789 | COPG1 | 22820 | coatomer protein complex subunit gamma 1 | up | 1,50 | 4,75E-02 |
| ENSG00000091527 | CDV3 | 55573 | CDV3 homolog | up | 1,50 | 9,77E-03 |
| ENSG00000028203 | VEZT | 55591 | vezatin, adherens junctions transmembrane protein | up | 1,50 | 5,68E-03 |
| ENSG00000107560 | RAB11FIP2 | 22841 | RAB11 family interacting protein 2 | up | 1,50 | 4,24E-02 |
| ENSG00000176407 | KCMF1 | 56888 | potassium channel modulatory factor 1 | up | 1,50 | 1,57E-02 |
| ENSG00000204344 | STK19 | 8859 | serine/threonine kinase 19 | up | 1,50 | 3,24E-02 |
| ENSG00000105821 | DNAJC2 | 27000 | DnaJ heat shock protein family (Hsp40) member C2 | up | 1,50 | 1,50E-02 |
| ENSG00000113595 | TRIM23 | 373 | tripartite motif containing 23 | up | 1,50 | 1,76E-02 |
| ENSG00000129083 | COPB1 | 1315 | coatomer protein complex subunit beta 1 | up | 1,50 | 1,68E-02 |
| ENSG00000137944 | KYAT3 | 56267 | kynurenine aminotransferase 3 | up | 1,50 | 3,91E-02 |
| ENSG00000095564 | BTAF1 | 9044 | B-TFIID TATA-box binding protein associated factor 1 | up | 1,50 | 1,71E-02 |
| ENSG00000180917 | CMTR2 | 55783 | cap methyltransferase 2 | up | 1,50 | 2,63E-02 |
| ENSG00000215717 | TMEM167B | 56900 | transmembrane protein 167B | up | 1,50 | 4,58E-03 |
| ENSG00000169087 | HSPBAP1 | 79663 | HSPB1 associated protein 1 | up | 1,49 | 4,53E-02 |
| ENSG00000114354 | TFG | 10342 | TRK-fused gene | up | 1,49 | 1,96E-02 |
| ENSG00000122643 | NT5C3A | 51251 | 5'-nucleotidase, cytosolic IIIA | up | 1,49 | 1,52E-02 |
| ENSG00000033867 | SLC4A7 | 9497 | solute carrier family 4 member 7 | up | 1,49 | 3,78E-03 |
| ENSG00000159256 | MORC3 | 23515 | MORC family CW-type zinc finger 3 | up | 1,49 | 1,67E-02 |
| ENSG00000105835 | NAMPT | 10135 | nicotinamide phosphoribosyltransferase | up | 1,49 | 4,91E-02 |
| ENSG00000143384 | MCL1 | 4170 | MCL1, BCL2 family apoptosis regulator | up | 1,49 | 1,70E-02 |
| ENSG00000061987 | MON2 | 23041 | MON2 homolog, regulator of endosome-to-Golgi trafficking | up | 1,49 | 5,95E-03 |
| ENSG00000108854 | SMURF2 | 64750 | SMAD specific E3 ubiquitin protein ligase 2 | up | 1,49 | 4,17E-02 |
| ENSG00000112658 | SRF | 6722 | serum response factor | up | 1,49 | 3,97E-02 |
| ENSG00000158092 | NCK1 | 4690 | NCK adaptor protein 1 | up | 1,49 | 1,42E-02 |

| | | | | | | |
|---|---|---|---|---|---|---|
| ENSG00000205323 | SARNP | 84324 | SAP domain containing ribonucleoprotein | up | 1,48 | 4,36E-02 |
| ENSG00000097033 | SH3GLB1 | 51100 | SH3 domain containing GRB2 like, endophilin B1 | up | 1,48 | 1,94E-02 |
| ENSG00000117500 | TMED5 | 50999 | transmembrane p24 trafficking protein 5 | up | 1,48 | 4,72E-02 |
| ENSG00000144895 | EIF2A | 83939 | eukaryotic translation initiation factor 2A | up | 1,48 | 9,76E-03 |
| ENSG00000155099 | PIP4P2 | 55529 | phosphatidylinositol-4,5-bisphosphate 4-phosphatase 2 | up | 1,48 | 2,19E-02 |
| ENSG00000141452 | WDR98 | 29919 | WD repeat domain 98 | up | 1,48 | 1,22E-02 |
| ENSG00000155975 | VPS37A | 137492 | VPS37A, ESCRT-I subunit | up | 1,48 | 1,14E-02 |
| ENSG00000146833 | TRIM4 | 89122 | tripartite motif containing 4 | up | 1,48 | 3,49E-02 |
| ENSG00000180769 | WDFY3-AS2 | 404201 | WDFY3 antisense RNA 2 | up | 1,48 | 3,15E-02 |
| ENSG00000129292 | PHF20L1 | 51105 | PHD finger protein 20 like 1 | up | 1,48 | 1,50E-02 |
| ENSG00000151414 | NEK7 | 140609 | NIMA related kinase 7 | up | 1,48 | 2,18E-02 |
| ENSG00000036054 | TBC1D23 | 55773 | TBC1 domain family member 23 | up | 1,48 | 1,79E-02 |
| ENSG00000123240 | OPTN | 10133 | optineurin | up | 1,48 | 2,60E-02 |
| ENSG00000135749 | PCNX2 | 80003 | pecanex homolog 2 | up | 1,47 | 1,00E-02 |
| ENSG00000095787 | WAC | 51322 | WW domain containing adaptor with coiled-coil | up | 1,47 | 1,00E-02 |
| ENSG00000065613 | SLK | 9748 | STE20 like kinase | up | 1,47 | 2,14E-03 |
| ENSG00000103047 | TANGO6 | 79613 | transport and golgi organization 6 homolog | up | 1,47 | 1,32E-02 |
| ENSG00000152104 | PTPN14 | 5784 | protein tyrosine phosphatase, non-receptor type 14 | up | 1,47 | 2,82E-02 |
| ENSG00000166783 | MARF1 | 9665 | meiosis regulator and mRNA stability factor 1 | up | 1,47 | 2,81E-02 |
| ENSG00000135297 | MTO1 | 25821 | mitochondrial tRNA translation optimization 1 | up | 1,47 | 2,12E-02 |
| ENSG00000270362 | HMGN3-AS1 | 100288198 | HMGN3 antisense RNA 1 | up | 1,47 | 1,91E-02 |
| ENSG00000217128 | FNIP1 | 96459 | folliculin interacting protein 1 | up | 1,47 | 1,66E-02 |
| ENSG00000106733 | NMRK1 | 54981 | nicotinamide riboside kinase 1 | up | 1,47 | 1,76E-02 |
| ENSG00000100814 | CCNB1IP1 | 57820 | cyclin B1 interacting protein 1 | up | 1,47 | 4,80E-02 |
| ENSG00000117151 | CTBS | 1486 | chitobiase | up | 1,47 | 8,32E-03 |
| ENSG00000127914 | AKAP9 | 10142 | A-kinase anchoring protein 9 | up | 1,47 | 1,81E-02 |
| ENSG00000159200 | RCAN1 | 1827 | regulator of calcineurin 1 | up | 1,47 | 4,96E-02 |
| ENSG00000110274 | CEP164 | 22897 | centrosomal protein 164 | up | 1,46 | 2,91E-02 |
| ENSG00000104067 | TJP1 | 7082 | tight junction protein 1 | up | 1,46 | 8,91E-03 |
| ENSG00000169155 | ZBTB43 | 23099 | zinc finger and BTB domain containing 43 | up | 1,46 | 1,07E-02 |
| ENSG00000179119 | SPTY2D1 | 144108 | SPT2 chromatin protein domain containing 1 | up | 1,46 | 1,81E-02 |
| ENSG00000100934 | SEC23A | 10484 | Sec23 homolog A, coat complex II component | up | 1,46 | 2,47E-02 |
| ENSG00000124209 | RAB22A | 57403 | RAB22A, member RAS oncogene family | up | 1,46 | 3,17E-02 |
| ENSG00000123595 | RAB9A | 9367 | RAB9A, member RAS oncogene family | up | 1,46 | 1,66E-02 |
| ENSG00000106665 | CLIP2 | 7461 | CAP-Gly domain containing linker protein 2 | up | 1,46 | 3,48E-02 |
| ENSG00000136051 | WASHC4 | 23325 | WASH complex subunit 4 | up | 1,46 | 1,05E-02 |
| ENSG00000066697 | MSANTD3 | 91283 | Myb/SANT DNA binding domain containing 3 | up | 1,46 | 1,35E-02 |
| ENSG00000173451 | THAP2 | 83591 | THAP domain containing 2 | up | 1,46 | 4,82E-02 |
| ENSG00000198060 | MARCH5 | 54708 | membrane associated ring-CH-type finger 5 | up | 1,46 | 1,95E-02 |
| ENSG00000151327 | FAM177A1 | 283635 | family with sequence similarity 177 member A1 | up | 1,45 | 2,44E-02 |
| ENSG00000151327 | LOC101927178 | 101927178 | uncharacterized LOC101927178 | up | 1,45 | 2,44E-02 |
| ENSG00000050426 | LETMD1 | 25875 | LETM1 domain containing 1 | up | 1,45 | 7,28E-03 |
| ENSG00000115109 | EPB41L5 | 57669 | erythrocyte membrane protein band 4.1 like 5 | up | 1,45 | 6,92E-03 |
| ENSG00000113407 | TARS | 6897 | threonyl-tRNA synthetase | up | 1,45 | 1,96E-02 |
| ENSG00000179335 | CLK3 | 1198 | CDC like kinase 3 | up | 1,45 | 2,44E-02 |
| ENSG00000127870 | RNF6 | 6049 | ring finger protein 6 | up | 1,45 | 1,98E-02 |
| ENSG00000166398 | KIAA0355 | 9710 | KIAA0355 | up | 1,45 | 3,73E-02 |
| ENSG00000095139 | ARCN1 | 372 | archain 1 | up | 1,45 | 2,54E-02 |
| ENSG00000144445 | KANSL1L | 151050 | KAT8 regulatory NSL complex subunit 1 like | up | 1,45 | 4,79E-02 |
| ENSG00000106052 | TAX1BP1 | 8887 | Tax1 binding protein 1 | up | 1,45 | 1,33E-03 |
| ENSG00000155621 | C9orf85 | 138241 | chromosome 9 open reading frame 85 | up | 1,45 | 2,00E-02 |
| ENSG00000176853 | FAM91A1 | 157769 | family with sequence similarity 91 member A1 | up | 1,45 | 1,60E-02 |
| ENSG00000256060 | TRAPPC2B | 10597 | trafficking protein particle complex 2B | up | 1,45 | 3,77E-02 |
| ENSG00000107625 | DDX50 | 79009 | DExD-box helicase 50 | up | 1,45 | 5,81E-03 |
| ENSG00000138468 | SENP7 | 57337 | SUMO1/sentrin specific peptidase 7 | up | 1,45 | 3,58E-02 |
| ENSG00000164221 | CCDC112 | 153733 | coiled-coil domain containing 112 | up | 1,45 | 1,19E-02 |
| ENSG00000169762 | TAPT1 | 202018 | transmembrane anterior posterior transformation 1 | up | 1,44 | 1,41E-02 |
| ENSG00000064652 | SNX24 | 28966 | sorting nexin 24 | up | 1,44 | 2,85E-02 |
| ENSG00000036549 | ZZZ3 | 26009 | zinc finger ZZ-type containing 3 | up | 1,44 | 3,37E-03 |
| ENSG00000112851 | ERBIN | 55914 | erbb2 interacting protein | up | 1,44 | 6,50E-03 |
| ENSG00000156650 | KAT6B | 23522 | lysine acetyltransferase 6B | up | 1,44 | 4,06E-02 |
| ENSG00000169251 | NMD3 | 51068 | NMD3 ribosome export adaptor | up | 1,44 | 3,25E-03 |
| ENSG00000123106 | CCDC91 | 55297 | coiled-coil domain containing 91 | up | 1,44 | 3,38E-02 |
| ENSG00000137575 | SDCBP | 6386 | syndecan binding protein | up | 1,44 | 8,08E-03 |
| ENSG00000163818 | LZTFL1 | 54585 | leucine zipper transcription factor like 1 | up | 1,44 | 4,05E-02 |
| ENSG00000115839 | RAB3GAP1 | 22930 | RAB3 GTPase activating protein catalytic subunit 1 | up | 1,44 | 1,49E-02 |
| ENSG00000181704 | YIPF6 | 286451 | Yip1 domain family member 6 | up | 1,44 | 2,30E-02 |
| ENSG00000117475 | BLZF1 | 8548 | basic leucine zipper nuclear factor 1 | up | 1,43 | 2,31E-02 |
| ENSG00000174780 | SRP72 | 6731 | signal recognition particle 72 | up | 1,43 | 9,37E-03 |
| ENSG00000010017 | RANBP9 | 10048 | RAN binding protein 9 | up | 1,43 | 1,66E-02 |
| ENSG00000186141 | POLR3C | 10623 | RNA polymerase III subunit C | up | 1,43 | 4,98E-02 |
| ENSG00000122741 | DCAF10 | 79269 | DDB1 and CUL4 associated factor 10 | up | 1,43 | 3,90E-02 |
| ENSG00000114850 | SSR3 | 6747 | signal sequence receptor subunit 3 | up | 1,43 | 4,56E-02 |
| ENSG00000092148 | HECTD1 | 25831 | HECT domain E3 ubiquitin protein ligase 1 | up | 1,43 | 1,58E-02 |
| ENSG00000092978 | GPATCH2 | 55105 | G-patch domain containing 2 | up | 1,43 | 6,92E-03 |
| ENSG00000110888 | CAPRIN2 | 65981 | caprin family member 2 | up | 1,42 | 2,53E-02 |
| ENSG00000182923 | CEP63 | 80254 | centrosomal protein 63 | up | 1,42 | 2,18E-02 |
| ENSG00000075223 | SEMA3C | 10512 | semaphorin 3C | up | 1,42 | 3,05E-02 |
| ENSG00000111911 | HINT3 | 135114 | histidine triad nucleotide binding protein 3 | up | 1,42 | 4,73E-02 |
| ENSG00000155744 | FAM126B | 285172 | family with sequence similarity 126 member B | up | 1,42 | 3,30E-02 |
| ENSG00000186908 | ZDHHC17 | 23390 | zinc finger DHHC-type containing 17 | up | 1,42 | 4,56E-02 |
| ENSG00000076685 | NT5C2 | 22978 | 5'-nucleotidase, cytosolic II | up | 1,42 | 1,00E-02 |
| ENSG00000188786 | MTF1 | 4520 | metal regulatory transcription factor 1 | up | 1,42 | 3,66E-02 |
| ENSG00000151320 | AKAP6 | 9472 | A-kinase anchoring protein 6 | up | 1,42 | 4,80E-02 |
| ENSG00000156256 | USP16 | 10600 | ubiquitin specific peptidase 16 | up | 1,42 | 1,31E-02 |
| ENSG00000118816 | CCNI | 10983 | cyclin I | up | 1,42 | 2,19E-02 |
| ENSG00000132952 | USPL1 | 10208 | ubiquitin specific peptidase like 1 | up | 1,42 | 2,68E-02 |
| ENSG00000130340 | SNX9 | 51429 | sorting nexin 9 | up | 1,42 | 4,22E-02 |
| ENSG00000184903 | IMMP2L | 83943 | inner mitochondrial membrane peptidase subunit 2 | up | 1,42 | 2,85E-02 |
| ENSG00000188554 | NBR1 | 4077 | NBR1, autophagy cargo receptor | up | 1,42 | 2,18E-02 |
| ENSG00000136628 | EPRS | 2058 | glutamyl-prolyl-tRNA synthetase | up | 1,42 | 1,95E-02 |
| ENSG00000204524 | ZNF805 | 390980 | zinc finger protein 805 | up | 1,42 | 3,69E-02 |
| ENSG00000118873 | RAB3GAP2 | 25782 | RAB3 GTPase activating non-catalytic protein subunit 2 | up | 1,42 | 3,40E-02 |
| ENSG00000143155 | TIPRL | 261726 | TOR signaling pathway regulator | up | 1,42 | 2,93E-02 |
| ENSG00000105865 | DUS4L | 11062 | dihydrouridine synthase 4 like | up | 1,42 | 2,35E-02 |
| ENSG00000179833 | SERTAD2 | 9792 | SERTA domain containing 2 | up | 1,42 | 1,21E-02 |
| ENSG00000137831 | UACA | 55075 | uveal autoantigen with coiled-coil domains and ankyrin repeats | up | 1,42 | 3,62E-03 |
| ENSG00000173273 | TNKS | 8658 | tankyrase | up | 1,42 | 8,32E-03 |
| ENSG00000115306 | SPTBN1 | 6711 | spectrin beta, non-erythrocytic 1 | up | 1,42 | 1,66E-02 |
| ENSG00000166848 | LOC105371348 | 105371348 | uncharacterized LOC105371348 | up | 1,41 | 1,58E-02 |
| ENSG00000166848 | TERF2IP | 54386 | TERF2 interacting protein | up | 1,41 | 1,58E-02 |
| ENSG00000168944 | CEP120 | 153241 | centrosomal protein 120 | up | 1,41 | 3,66E-02 |
| ENSG00000173744 | AGFG1 | 3267 | ArfGAP with FG repeats 1 | up | 1,41 | 1,40E-02 |
| ENSG00000167257 | RNF214 | 257160 | ring finger protein 214 | up | 1,41 | 4,25E-02 |
| ENSG00000173281 | PPP1R3B | 79660 | protein phosphatase 1 regulatory subunit 3B | up | 1,41 | 4,67E-02 |
| ENSG00000070778 | PTPN21 | 11099 | protein tyrosine phosphatase, non-receptor type 21 | up | 1,41 | 3,58E-02 |
| ENSG00000122912 | SLC25A16 | 8034 | solute carrier family 25 member 16 | up | 1,41 | 4,87E-02 |
| ENSG00000156671 | SAMD8 | 142891 | sterile alpha motif domain containing 8 | up | 1,41 | 2,54E-02 |
| ENSG00000163510 | CWC22 | 57703 | CWC22 spliceosome associated protein homolog | up | 1,41 | 8,63E-03 |
| ENSG00000048544 | MRPS10 | 55173 | mitochondrial ribosomal protein S10 | up | 1,41 | 1,63E-02 |
| ENSG00000115145 | STAM2 | 10254 | signal transducing adaptor molecule 2 | up | 1,41 | 7,00E-03 |
| ENSG00000136436 | CALCOCO2 | 10241 | calcium binding and coiled-coil domain 2 | up | 1,41 | 2,11E-02 |
| ENSG00000085721 | RRN3 | 54700 | RRN3 homolog, RNA polymerase I transcription factor | up | 1,41 | 4,25E-02 |
| ENSG00000135968 | GCC2 | 9648 | GRIP and coiled-coil domain containing 2 | up | 1,41 | 1,40E-02 |
| ENSG00000058063 | ATP11B | 23200 | ATPase phospholipid transporting 11B (putative) | up | 1,40 | 9,63E-03 |
| ENSG00000151422 | FER | 2241 | FER tyrosine kinase | up | 1,40 | 6,10E-03 |
| ENSG00000175105 | ZNF654 | 55279 | zinc finger protein 654 | up | 1,40 | 4,36E-02 |
| ENSG00000138081 | FBXO11 | 80204 | F-box protein 11 | up | 1,40 | 2,20E-02 |
| ENSG00000075413 | MARK3 | 4140 | microtubule affinity regulating kinase 3 | up | 1,40 | 1,07E-02 |
| ENSG00000120616 | EPC1 | 80314 | enhancer of polycomb homolog 1 | up | 1,40 | 2,88E-02 |
| ENSG00000138036 | DYNC2LI1 | 51626 | dynein cytoplasmic 2 light intermediate chain 1 | up | 1,40 | 3,10E-02 |
| ENSG00000204406 | MBD5 | 55777 | methyl-CpG binding domain protein 5 | up | 1,40 | 2,85E-02 |
| ENSG00000141447 | OSBPL1A | 114876 | oxysterol binding protein like 1A | up | 1,40 | 2,14E-03 |
| ENSG00000125977 | EIF2S2 | 8894 | eukaryotic translation initiation factor 2 subunit beta | up | 1,40 | 2,40E-02 |
| ENSG00000170759 | KIF5B | 3799 | kinesin family member 5B | up | 1,40 | 7,18E-03 |
| ENSG00000167881 | SRP68 | 6730 | signal recognition particle 68 | up | 1,40 | 3,03E-02 |
| ENSG00000038219 | BOD1L1 | 259282 | biorientation of chromosomes in cell division 1 like 1 | up | 1,40 | 3,33E-02 |
| ENSG00000123636 | BAZ2B | 29994 | bromodomain adjacent to zinc finger domain 2B | up | 1,40 | 3,86E-02 |
| ENSG00000164597 | COG5 | 10466 | component of oligomeric golgi complex 5 | up | 1,40 | 3,37E-03 |
| ENSG00000157625 | TAB3 | 257397 | TGF-beta activated kinase 1 and MAP3K7 binding protein 3 | up | 1,40 | 4,94E-02 |
| ENSG00000135049 | AGTPBP1 | 23287 | ATP/GTP binding protein 1 | up | 1,40 | 1,30E-02 |
| ENSG00000179152 | TCAIM | 285343 | T-cell activation inhibitor, mitochondrial | up | 1,40 | 1,31E-02 |
| ENSG00000175354 | PTPN2 | 5771 | protein tyrosine phosphatase, non-receptor type 2 | up | 1,40 | 1,73E-02 |
| ENSG00000015153 | YAF2 | 10138 | YY1 associated factor 2 | up | 1,40 | 2,35E-02 |
| ENSG00000125962 | ARMCX5 | 64860 | armadillo repeat containing, X-linked 5 | up | 1,40 | 3,10E-02 |
| ENSG00000019995 | ZRANB1 | 54764 | zinc finger RANBP2-type containing 1 | up | 1,39 | 1,57E-02 |
| ENSG00000132824 | SERINC3 | 10955 | serine incorporator 3 | up | 1,39 | 4,64E-02 |
| ENSG00000151553 | FAM160B1 | 57700 | family with sequence similarity 160 member B1 | up | 1,39 | 2,29E-02 |
| ENSG00000154222 | CC2D1B | 200014 | coiled-coil and C2 domain containing 1B | up | 1,39 | 4,35E-02 |
| ENSG00000136003 | ISCU | 23479 | iron-sulfur cluster assembly enzyme | up | 1,39 | 3,98E-02 |
| ENSG00000171843 | MLLT3 | 4300 | MLLT3, super elongation complex subunit | up | 1,39 | 2,58E-02 |
| ENSG00000163001 | CFAP36 | 112942 | cilia and flagella associated protein 36 | up | 1,39 | 2,03E-02 |
| ENSG00000164023 | SGMS2 | 166929 | sphingomyelin synthase 2 | up | 1,39 | 4,00E-02 |
| ENSG00000183621 | ZNF438 | 220929 | zinc finger protein 438 | up | 1,39 | 4,73E-02 |
| ENSG00000111790 | FGFR1OP2 | 26127 | FGFR1 oncogene partner 2 | up | 1,39 | 2,10E-02 |
| ENSG00000123836 | PFKFB2 | 5208 | 6-phosphofructo-2-kinase/fructose-2,6-biphosphatase 2 | up | 1,39 | 4,33E-02 |
| ENSG00000118997 | DNAH7 | 56171 | dynein axonemal heavy chain 7 | up | 1,39 | 4,33E-02 |
| ENSG00000086102 | NFX1 | 4799 | nuclear transcription factor, X-box binding 1 | up | 1,39 | 2,52E-02 |
| ENSG00000108587 | GOSR1 | 9527 | golgi SNAP receptor complex member 1 | up | 1,39 | 3,34E-02 |
| ENSG00000164329 | PAPD4 | 167153 | poly(A) RNA polymerase D4, non-canonical | up | 1,39 | 1,94E-02 |
| ENSG00000148925 | BTBD10 | 84280 | BTB domain containing 10 | up | 1,38 | 4,52E-02 |
| ENSG00000178996 | SNX18 | 112574 | sorting nexin 18 | up | 1,38 | 4,81E-02 |
| ENSG00000058729 | RIOK2 | 55781 | RIO kinase 2 | up | 1,38 | 3,10E-03 |
| ENSG00000103978 | TMEM87A | 25963 | transmembrane protein 87A | up | 1,38 | 2,31E-02 |
| ENSG00000204842 | ATXN2 | 6311 | ataxin 2 | up | 1,38 | 2,82E-02 |
| ENSG00000147854 | UHRF2 | 115426 | ubiquitin like with PHD and ring finger domains 2 | up | 1,38 | 2,49E-02 |
| ENSG00000119820 | YIPF4 | 84272 | Yip1 domain family member 4 | up | 1,38 | 3,19E-02 |
| ENSG00000157224 | CLDN12 | 9069 | claudin 12 | up | 1,38 | 3,08E-02 |
| ENSG00000156232 | WHAMM | 123720 | WAS protein homolog associated with actin, golgi membranes and microtubu | up | 1,38 | 2,44E-02 |
| ENSG00000151092 | NGLY1 | 55768 | N-glycanase 1 | up | 1,38 | 1,28E-02 |
| ENSG00000110330 | BIRC2 | 329 | baculoviral IAP repeat containing 2 | up | 1,38 | 1,86E-02 |
| ENSG00000196505 | GDAP2 | 54834 | ganglioside induced differentiation associated protein 2 | up | 1,38 | 2,16E-02 |
| ENSG00000196284 | SUPT3H | 8464 | SPT3 homolog, SAGA and STAGA complex component | up | 1,38 | 1,01E-02 |
| ENSG00000107643 | MAPK8 | 5599 | mitogen-activated protein kinase 8 | up | 1,38 | 3,05E-02 |
| ENSG00000138138 | ATAD1 | 84896 | ATPase family, AAA domain containing 1 | up | 1,38 | 1,16E-02 |
| ENSG00000166685 | COG1 | 9382 | component of oligomeric golgi complex 1 | up | 1,38 | 4,80E-02 |
| ENSG00000042286 | AIFM2 | 84883 | apoptosis inducing factor, mitochondria associated 2 | up | 1,38 | 4,97E-02 |
| ENSG00000104412 | EMC2 | 9694 | ER membrane protein complex subunit 2 | up | 1,38 | 1,03E-02 |
| ENSG00000165630 | PRPF18 | 8559 | pre-mRNA processing factor 18 | up | 1,37 | 1,40E-02 |
| ENSG00000119446 | RBM18 | 92400 | RNA binding motif protein 18 | up | 1,37 | 4,06E-02 |
| ENSG00000130363 | RSPH3 | 83861 | radial spoke head 3 homolog | up | 1,37 | 3,38E-02 |
| ENSG00000136770 | DNAJC1 | 64215 | DnaJ heat shock protein family (Hsp40) member C1 | up | 1,37 | 4,59E-02 |
| ENSG00000163681 | SLMAP | 7871 | sarcolemma associated protein | up | 1,37 | 9,55E-03 |
| ENSG00000169764 | UGP2 | 7360 | UDP-glucose pyrophosphorylase 2 | up | 1,37 | 3,48E-02 |
| ENSG00000214174 | AMZ2P1 | 201283 | archaelysin family metallopeptidase 2 pseudogene 1 | up | 1,37 | 1,16E-02 |
| ENSG00000102786 | INTS6 | 26512 | integrator complex subunit 6 | up | 1,37 | 9,77E-03 |
| ENSG00000157181 | ODR4 | 54953 | odr-4 GPCR localization factor homolog | up | 1,37 | 1,40E-02 |
| ENSG00000126524 | SBDS | 51119 | SBDS, ribosome maturation factor | up | 1,37 | 4,62E-02 |
| ENSG00000134371 | CDC73 | 79577 | cell division cycle 73 | up | 1,37 | 4,42E-02 |
| ENSG00000175198 | PCCA | 5095 | propionyl-CoA carboxylase alpha subunit | up | 1,37 | 2,03E-02 |
| ENSG00000070495 | JMJD6 | 23210 | arginine demethylase and lysine hydroxylase | up | 1,37 | 4,72E-02 |
| ENSG00000134058 | CDK7 | 1022 | cyclin dependent kinase 7 | up | 1,37 | 4,38E-02 |
| ENSG00000179818 | PCBP1-AS1 | 400960 | PCBP1 antisense RNA 1 | up | 1,37 | 3,61E-02 |
| ENSG00000198648 | STK39 | 27347 | serine/threonine kinase 39 | up | 1,36 | 8,70E-03 |
| ENSG00000253797 | UTP14C | 9724 | UTP14C, small subunit processome component | up | 1,36 | 4,71E-02 |
| ENSG00000136273 | HUS1 | 3364 | HUS1 checkpoint clamp component | up | 1,36 | 4,97E-02 |
| ENSG00000137821 | LRRC49 | 54839 | leucine rich repeat containing 49 | up | 1,36 | 3,58E-02 |
| ENSG00000166233 | ARIH1 | 25820 | ariadne RBR E3 ubiquitin protein ligase 1 | up | 1,36 | 8,38E-03 |
| ENSG00000140830 | TXNL4B | 54957 | thioredoxin like 4B | up | 1,36 | 4,14E-02 |
| ENSG00000141337 | ARSG | 22901 | arylsulfatase G | up | 1,36 | 3,94E-02 |
| ENSG00000197498 | RPF2 | 84154 | ribosome production factor 2 homolog | up | 1,36 | 3,39E-02 |
| ENSG00000128908 | INO80 | 54617 | INO80 complex subunit | up | 1,36 | 1,66E-02 |
| ENSG00000126070 | AGO3 | 192669 | argonaute 3, RISC catalytic component | up | 1,36 | 1,80E-02 |
| ENSG00000120254 | MTHFD1L | 25902 | methylenetetrahydrofolate dehydrogenase (NADP+ dependent) 1 like | up | 1,36 | 4,52E-02 |
| ENSG00000127022 | CANX | 821 | calnexin | up | 1,36 | 3,99E-02 |
| ENSG00000141198 | TOM1L1 | 10040 | target of myb1 like 1 membrane trafficking protein | up | 1,36 | 1,19E-02 |
| ENSG00000170242 | USP47 | 55031 | ubiquitin specific peptidase 47 | up | 1,36 | 3,59E-03 |
| ENSG00000188215 | DCUN1D3 | 123879 | defective in cullin neddylation 1 domain containing 3 | up | 1,36 | 4,53E-02 |
| ENSG00000281649 | EBLN3P | 100506710 | endogenous Bornavirus-like nucleoprotein 3, pseudogene | up | 1,36 | 4,86E-02 |
| ENSG00000135541 | AHI1 | 54806 | Abelson helper integration site 1 | up | 1,35 | 2,73E-03 |
| ENSG00000112701 | SENP6 | 26054 | SUMO1/sentrin specific peptidase 6 | up | 1,35 | 1,16E-02 |
| ENSG00000113712 | CSNK1A1 | 1452 | casein kinase 1 alpha 1 | up | 1,35 | 2,85E-02 |
| ENSG00000101882 | NKAP | 79576 | NFKB activating protein | up | 1,35 | 9,48E-03 |
| ENSG00000122299 | ZC3H7A | 29066 | zinc finger CCCH-type containing 7A | up | 1,35 | 3,89E-02 |
| ENSG00000126777 | KTN1 | 3895 | kinectin 1 | up | 1,35 | 2,24E-03 |
| ENSG00000138433 | CIR1 | 9541 | corepressor interacting with RBPJ, 1 | up | 1,35 | 3,15E-02 |
| ENSG00000145782 | ATG12 | 9140 | autophagy related 12 | up | 1,35 | 4,11E-02 |
| ENSG00000152242 | C18orf25 | 147339 | chromosome 18 open reading frame 25 | up | 1,35 | 1,04E-02 |
| ENSG00000001167 | NFYA | 4800 | nuclear transcription factor Y subunit alpha | up | 1,35 | 3,30E-02 |
| ENSG00000104756 | KCTD9 | 54793 | potassium channel tetramerization domain containing 9 | up | 1,35 | 4,42E-02 |
| ENSG00000070961 | ATP2B1 | 490 | ATPase plasma membrane Ca2+ transporting 1 | up | 1,35 | 4,18E-02 |
| ENSG00000056277 | ZNF280C | 55609 | zinc finger protein 280C | up | 1,35 | 5,95E-03 |
| ENSG00000168288 | MMADHC | 27249 | methylmalonic aciduria and homocystinuria, cblD type | up | 1,35 | 3,58E-02 |
| ENSG00000044090 | CUL7 | 9820 | cullin 7 | up | 1,35 | 4,05E-02 |
| ENSG00000129315 | CCNT1 | 904 | cyclin T1 | up | 1,34 | 3,05E-02 |
| ENSG00000203705 | TATDN3 | 128387 | TatD DNase domain containing 3 | up | 1,34 | 1,67E-02 |
| ENSG00000140262 | TCF12 | 6938 | transcription factor 12 | up | 1,34 | 4,99E-02 |

| | | | | | | |
|---|---|---|---|---|---|---|
| ENSG00000152684 | PELO | 53918 | pelota mRNA surveillance and ribosome rescue factor | up | 1,34 | 4,50E-02 |
| ENSG00000115808 | STRN | 6801 | striatin | up | 1,34 | 2,37E-02 |
| ENSG00000073921 | PICALM | 8301 | phosphatidylinositol binding clathrin assembly protein | up | 1,34 | 3,22E-02 |
| ENSG00000089234 | BRAP | 8315 | BRCA1 associated protein | up | 1,34 | 1,15E-02 |
| ENSG00000173875 | ZNF791 | 163049 | zinc finger protein 791 | up | 1,34 | 2,88E-02 |
| ENSG00000131263 | RLIM | 51132 | ring finger protein, LIM domain interacting | up | 1,34 | 9,04E-03 |
| ENSG00000169018 | FEM1B | 10116 | fem-1 homolog B | up | 1,34 | 2,19E-02 |
| ENSG00000083799 | CYLD | 1540 | CYLD lysine 63 deubiquitinase | up | 1,34 | 4,86E-02 |
| ENSG00000134313 | KIDINS220 | 57498 | kinase D interacting substrate 220 | up | 1,34 | 3,95E-02 |
| ENSG00000173226 | IQCB1 | 9657 | IQ motif containing B1 | up | 1,34 | 1,42E-02 |
| ENSG00000140386 | SCAPER | 49855 | S-phase cyclin A associated protein in the ER | up | 1,34 | 4,13E-02 |
| ENSG00000136813 | KIAA0368 | 23392 | KIAA0368 | up | 1,34 | 1,86E-02 |
| ENSG00000110768 | GTF2H1 | 2965 | general transcription factor IIH subunit 1 | up | 1,34 | 1,86E-02 |
| ENSG00000112305 | SMAP1 | 60682 | small ArfGAP 1 | up | 1,34 | 2,17E-02 |
| ENSG00000112414 | ADGRG6 | 57211 | adhesion G protein-coupled receptor G6 | up | 1,34 | 2,88E-02 |
| ENSG00000172965 | CYTOR | 112597 | cytoskeleton regulator RNA | up | 1,33 | 3,58E-02 |
| ENSG00000172965 | MIR4435-2HG | 541471 | MIR4435-2 host gene | up | 1,33 | 3,58E-02 |
| ENSG00000275052 | PPP4R3B | 57223 | protein phosphatase 4 regulatory subunit 3B | up | 1,33 | 4,11E-02 |
| ENSG00000121879 | PIK3CA | 5290 | phosphatidylinositol-4,5-bisphosphate 3-kinase catalytic subunit alpha | up | 1,33 | 3,18E-02 |
| ENSG00000100815 | TRIP11 | 9321 | thyroid hormone receptor interactor 11 | up | 1,33 | 2,24E-03 |
| ENSG00000168522 | FNTA | 2339 | farnesyltransferase, CAAX box, alpha | up | 1,33 | 3,74E-03 |
| ENSG00000214413 | BBIP1 | 92482 | BBSome interacting protein 1 | up | 1,33 | 3,49E-02 |
| ENSG00000128607 | KLHDC10 | 23008 | kelch domain containing 10 | up | 1,33 | 4,80E-02 |
| ENSG00000196151 | WDSUB1 | 151525 | WD repeat, sterile alpha motif and U-box domain containing 1 | up | 1,33 | 4,98E-02 |
| ENSG00000272886 | DCP1A | 55802 | decapping mRNA 1A | up | 1,33 | 4,54E-02 |
| ENSG00000075651 | PLD1 | 5337 | phospholipase D1 | up | 1,33 | 1,61E-02 |
| ENSG00000139505 | MTMR6 | 9107 | myotubularin related protein 6 | up | 1,33 | 2,92E-02 |
| ENSG00000134265 | NAPG | 8774 | NSF attachment protein gamma | up | 1,33 | 1,70E-02 |
| ENSG00000093167 | LRRFIP2 | 9209 | LRR binding FLII interacting protein 2 | up | 1,33 | 1,28E-02 |
| ENSG00000055147 | FAM114A2 | 10827 | family with sequence similarity 114 member A2 | up | 1,33 | 2,43E-02 |
| ENSG00000116678 | LEPR | 3953 | leptin receptor | up | 1,33 | 4,95E-03 |
| ENSG00000109466 | KLHL2 | 11275 | kelch like family member 2 | up | 1,33 | 3,06E-02 |
| ENSG00000144451 | SPAG16 | 79582 | sperm associated antigen 16 | up | 1,33 | 2,95E-02 |
| ENSG00000171862 | PTEN | 5728 | phosphatase and tensin homolog | up | 1,33 | 4,73E-02 |
| ENSG00000119778 | ATAD2B | 54454 | ATPase family, AAA domain containing 2B | up | 1,33 | 3,73E-02 |
| ENSG00000171466 | ZNF562 | 54811 | zinc finger protein 562 | up | 1,33 | 5,21E-03 |
| ENSG00000110841 | PPFIBP1 | 8496 | PPFIA binding protein 1 | up | 1,33 | 4,01E-02 |
| ENSG00000157796 | WDR19 | 57728 | WD repeat domain 19 | up | 1,33 | 4,96E-02 |
| ENSG00000114416 | FXR1 | 8087 | FMR1 autosomal homolog 1 | up | 1,32 | 6,51E-03 |
| ENSG00000164010 | ERMAP | 114625 | erythroblast membrane associated protein (Scianna blood group) | up | 1,32 | 3,38E-02 |
| ENSG00000177200 | CHD9 | 80205 | chromodomain helicase DNA binding protein 9 | up | 1,32 | 1,42E-03 |
| ENSG00000126653 | NSRP1 | 84081 | nuclear speckle splicing regulatory protein 1 | up | 1,32 | 1,31E-02 |
| ENSG00000086065 | CHMP5 | 51510 | charged multivesicular body protein 5 | up | 1,32 | 3,38E-02 |
| ENSG00000122882 | ECD | 11319 | ecdysoneless cell cycle regulator | up | 1,32 | 3,91E-02 |
| ENSG00000140455 | USP3 | 9960 | ubiquitin specific peptidase 3 | up | 1,32 | 2,23E-02 |
| ENSG00000131374 | TBC1D5 | 9779 | TBC1 domain family member 5 | up | 1,31 | 4,12E-03 |
| ENSG00000143493 | INTS7 | 25896 | integrator complex subunit 7 | up | 1,31 | 1,74E-02 |
| ENSG00000144224 | UBXN4 | 23190 | UBX domain protein 4 | up | 1,31 | 8,48E-03 |
| ENSG00000155906 | RMND1 | 55005 | required for meiotic nuclear division 1 homolog | up | 1,31 | 2,36E-02 |
| ENSG00000100575 | TIMM9 | 26520 | translocase of inner mitochondrial membrane 9 | up | 1,31 | 2,36E-02 |
| ENSG00000140299 | BNIP2 | 663 | BCL2 interacting protein 2 | up | 1,31 | 3,45E-02 |
| ENSG00000062725 | APPBP2 | 10513 | amyloid beta precursor protein binding protein 2 | up | 1,31 | 3,37E-03 |
| ENSG00000117000 | RLF | 6018 | rearranged L-myc fusion | up | 1,31 | 4,49E-02 |
| ENSG00000138592 | USP8 | 9101 | ubiquitin specific peptidase 8 | up | 1,31 | 7,82E-03 |
| ENSG00000180357 | ZNF609 | 23060 | zinc finger protein 609 | up | 1,31 | 1,86E-02 |
| ENSG00000067066 | SP100 | 6672 | SP100 nuclear antigen | up | 1,31 | 2,72E-02 |
| ENSG00000104133 | SPG11 | 80208 | SPG11, spatacsin vesicle trafficking associated | up | 1,31 | 2,11E-02 |
| ENSG00000152256 | PDK1 | 5163 | pyruvate dehydrogenase kinase 1 | up | 1,31 | 2,69E-03 |
| ENSG00000104064 | GABPB1 | 2553 | GA binding protein transcription factor beta subunit 1 | up | 1,31 | 1,57E-02 |
| ENSG00000104064 | GABPB1-IT1 | 55056 | GABPB1 intronic transcript | up | 1,31 | 1,57E-02 |
| ENSG00000117153 | KLHL12 | 59349 | kelch like family member 12 | up | 1,31 | 4,38E-02 |
| ENSG00000146085 | MUT | 4594 | methylmalonyl-CoA mutase | up | 1,31 | 4,58E-03 |
| ENSG00000112200 | ZNF451 | 26036 | zinc finger protein 451 | up | 1,31 | 4,42E-02 |
| ENSG00000138095 | LRPPRC | 10128 | leucine rich pentatricopeptide repeat containing | up | 1,31 | 4,03E-03 |
| ENSG00000204138 | PHACTR4 | 65979 | phosphatase and actin regulator 4 | up | 1,31 | 4,68E-02 |
| ENSG00000119760 | SUPT7L | 9913 | SPT7 like, STAGA complex gamma subunit | up | 1,30 | 3,95E-02 |
| ENSG00000011405 | PIK3C2A | 5286 | phosphatidylinositol-4-phosphate 3-kinase catalytic subunit type 2 alpha | up | 1,30 | 1,80E-02 |
| ENSG00000135655 | USP15 | 9958 | ubiquitin specific peptidase 15 | up | 1,30 | 2,19E-02 |
| ENSG00000164190 | NIPBL | 25836 | NIPBL, cohesin loading factor | up | 1,30 | 4,13E-02 |
| ENSG00000136758 | YME1L1 | 10730 | YME1 like 1 ATPase | up | 1,30 | 2,92E-02 |
| ENSG00000104695 | PPP2CB | 5516 | protein phosphatase 2 catalytic subunit beta | up | 1,30 | 4,79E-02 |
| ENSG00000152404 | CWF19L2 | 143884 | CWF19 like 2, cell cycle control (S. pombe) | up | 1,30 | 3,86E-02 |
| ENSG00000176105 | YES1 | 7525 | YES proto-oncogene 1, Src family tyrosine kinase | up | 1,30 | 2,09E-02 |
| ENSG00000085832 | EPS15 | 2060 | epidermal growth factor receptor pathway substrate 15 | up | 1,30 | 7,73E-03 |
| ENSG00000164615 | CAMLG | 819 | calcium modulating ligand | up | 1,30 | 4,05E-02 |
| ENSG00000114127 | XRN1 | 54464 | 5'-3' exoribonuclease 1 | up | 1,30 | 1,96E-02 |
| ENSG00000131626 | PPFIA1 | 8500 | PTPRF interacting protein alpha 1 | up | 1,30 | 3,69E-02 |
| ENSG00000187953 | PMS2CL | 441194 | PMS2 C-terminal like pseudogene | up | 1,30 | 3,99E-02 |
| ENSG00000064313 | TAF2 | 6873 | TATA-box binding protein associated factor 2 | up | 1,30 | 5,95E-03 |
| ENSG00000089289 | IGBP1 | 3476 | immunoglobulin binding protein 1 | up | 1,29 | 1,46E-02 |
| ENSG00000113272 | THG1L | 54974 | tRNA-histidine guanylyltransferase 1 like | up | 1,29 | 4,59E-02 |
| ENSG00000079950 | STX7 | 8417 | syntaxin 7 | up | 1,29 | 3,62E-02 |
| ENSG00000120690 | ELF1 | 1997 | E74 like ETS transcription factor 1 | up | 1,29 | 2,90E-02 |
| ENSG00000175387 | SMAD2 | 4087 | SMAD family member 2 | up | 1,29 | 2,19E-02 |
| ENSG00000101558 | VAPA | 9218 | VAMP associated protein A | up | 1,29 | 4,33E-02 |
| ENSG00000162869 | PPP1R21 | 129285 | protein phosphatase 1 regulatory subunit 21 | up | 1,29 | 2,42E-02 |
| ENSG00000107938 | EDRF1 | 26098 | erythroid differentiation regulatory factor 1 | up | 1,29 | 2,68E-02 |
| ENSG00000145349 | CAMK2D | 817 | calcium/calmodulin dependent protein kinase II delta | up | 1,29 | 1,49E-02 |
| ENSG00000163788 | SNRK | 54861 | SNF related kinase | up | 1,29 | 2,44E-02 |
| ENSG00000102710 | SUPT20H | 55578 | SPT20 homolog, SAGA complex component | up | 1,29 | 4,66E-02 |
| ENSG00000134440 | NARS | 4677 | asparaginyl-tRNA synthetase | up | 1,29 | 2,58E-02 |
| ENSG00000174718 | KIAA1551 | 55196 | KIAA1551 | up | 1,29 | 3,30E-02 |
| ENSG00000028116 | VRK2 | 7444 | vaccinia related kinase 2 | up | 1,29 | 9,55E-03 |
| ENSG00000078070 | MCCC1 | 56922 | methylcrotonoyl-CoA carboxylase 1 | up | 1,29 | 4,04E-02 |
| ENSG00000118518 | RNF146 | 81847 | ring finger protein 146 | up | 1,29 | 3,58E-02 |
| ENSG00000197603 | C5orf42 | 65250 | chromosome 5 open reading frame 42 | up | 1,29 | 4,63E-02 |
| ENSG00000119041 | GTF3C3 | 9330 | general transcription factor IIIC subunit 3 | up | 1,28 | 2,82E-03 |
| ENSG00000083312 | TNPO1 | 3842 | transportin 1 | up | 1,28 | 4,94E-02 |
| ENSG00000143756 | FBXO28 | 23219 | F-box protein 28 | up | 1,28 | 4,53E-02 |
| ENSG00000101166 | PRELID3B | 51012 | PRELI domain containing 3B | up | 1,28 | 4,61E-02 |
| ENSG00000124831 | LRRFIP1 | 9208 | LRR binding FLII interacting protein 1 | up | 1,28 | 2,23E-02 |
| ENSG00000164219 | PGGT1B | 5229 | protein geranylgeranyltransferase type I subunit beta | up | 1,28 | 4,67E-02 |
| ENSG00000171566 | PLRG1 | 5356 | pleiotropic regulator 1 | up | 1,28 | 3,19E-02 |
| ENSG00000106617 | PRKAG2 | 51422 | protein kinase AMP-activated non-catalytic subunit gamma 2 | up | 1,28 | 4,57E-02 |
| ENSG00000096654 | ZNF184 | 7738 | zinc finger protein 184 | up | 1,28 | 2,31E-02 |
| ENSG00000138286 | FAM149B1 | 317662 | family with sequence similarity 149 member B1 | up | 1,28 | 4,91E-02 |
| ENSG00000164610 | RP9 | 6100 | RP9, pre-mRNA splicing factor | up | 1,28 | 3,45E-02 |
| ENSG00000139746 | RBM26 | 64062 | RNA binding motif protein 26 | up | 1,28 | 2,69E-02 |
| ENSG00000156026 | MCU | 90550 | mitochondrial calcium uniporter | up | 1,27 | 2,20E-02 |
| ENSG00000181004 | BBS12 | 166379 | Bardet-Biedl syndrome 12 | up | 1,27 | 4,94E-02 |
| ENSG00000198677 | TTC37 | 9652 | tetratricopeptide repeat domain 37 | up | 1,27 | 2,64E-03 |
| ENSG00000109171 | SLAIN2 | 57606 | SLAIN motif family member 2 | up | 1,27 | 1,21E-02 |
| ENSG00000112343 | TRIM38 | 10475 | tripartite motif containing 38 | up | 1,27 | 3,50E-02 |
| ENSG00000165678 | GHITM | 27069 | growth hormone inducible transmembrane protein | up | 1,27 | 4,79E-02 |
| ENSG00000205339 | IPO7 | 10527 | importin 7 | up | 1,27 | 1,16E-02 |
| ENSG00000264522 | OTUD7B | 56957 | OTU deubiquitinase 7B | up | 1,27 | 3,81E-02 |
| ENSG00000163608 | NEPRO | 25871 | nucleolus and neural progenitor protein | up | 1,27 | 3,08E-02 |
| ENSG00000117614 | SYF2 | 25949 | SYF2 pre-mRNA splicing factor | up | 1,27 | 4,77E-02 |
| ENSG00000169967 | MAP3K2 | 10746 | mitogen-activated protein kinase kinase kinase 2 | up | 1,27 | 8,97E-03 |
| ENSG00000141627 | DYM | 54808 | dymeclin | up | 1,27 | 4,26E-02 |
| ENSG00000172765 | TMCC1 | 23023 | transmembrane and coiled-coil domain family 1 | up | 1,27 | 1,80E-02 |
| ENSG00000153317 | ASAP1 | 50807 | ArfGAP with SH3 domain, ankyrin repeat and PH domain 1 | up | 1,27 | 4,96E-02 |
| ENSG00000105793 | GTPBP10 | 85865 | GTP binding protein 10 | up | 1,26 | 4,63E-02 |
| ENSG00000105793 | LOC101927446 | 101927446 | uncharacterized LOC101927446 | up | 1,26 | 4,63E-02 |
| ENSG00000134759 | ELP2 | 55250 | elongator acetyltransferase complex subunit 2 | up | 1,26 | 4,37E-02 |
| ENSG00000134146 | DPH6 | 89978 | diphthamine biosynthesis 6 | up | 1,26 | 3,03E-02 |
| ENSG00000163798 | SLC4A1AP | 22950 | solute carrier family 4 member 1 adaptor protein | up | 1,26 | 1,23E-02 |
| ENSG00000221914 | PPP2R2A | 5520 | protein phosphatase 2 regulatory subunit Balpha | up | 1,26 | 3,62E-02 |
| ENSG00000085760 | MTIF2 | 4528 | mitochondrial translational initiation factor 2 | up | 1,26 | 1,83E-02 |
| ENSG00000198087 | CD2AP | 23607 | CD2 associated protein | up | 1,26 | 4,33E-02 |
| ENSG00000048649 | RSF1 | 51773 | remodeling and spacing factor 1 | up | 1,26 | 3,94E-02 |
| ENSG00000101972 | STAG2 | 10735 | stromal antigen 2 | up | 1,26 | 3,70E-02 |
| ENSG00000137055 | PLAA | 9373 | phospholipase A2 activating protein | up | 1,26 | 1,64E-02 |
| ENSG00000155096 | AZIN1 | 51582 | antizyme inhibitor 1 | up | 1,26 | 4,56E-02 |
| ENSG00000213020 | ZNF611 | 81856 | zinc finger protein 611 | up | 1,26 | 2,68E-02 |
| ENSG00000147316 | MCPH1 | 79648 | microcephalin 1 | up | 1,26 | 4,63E-02 |
| ENSG00000134900 | TPP2 | 7174 | tripeptidyl peptidase 2 | up | 1,25 | 4,04E-02 |
| ENSG00000136935 | GOLGA1 | 2800 | golgin A1 | up | 1,25 | 3,77E-02 |
| ENSG00000147202 | DIAPH2 | 1730 | diaphanous related formin 2 | up | 1,25 | 2,61E-02 |
| ENSG00000101557 | USP14 | 9097 | ubiquitin specific peptidase 14 | up | 1,25 | 2,47E-02 |
| ENSG00000104320 | NBN | 4683 | nibrin | up | 1,25 | 3,16E-02 |
| ENSG00000070501 | POLB | 5423 | DNA polymerase beta | up | 1,25 | 3,76E-02 |
| ENSG00000132664 | POLR3F | 10621 | RNA polymerase III subunit F | up | 1,25 | 3,24E-02 |
| ENSG00000149100 | EIF3M | 10480 | eukaryotic translation initiation factor 3 subunit M | up | 1,25 | 2,52E-02 |
| ENSG00000082213 | C5orf22 | 55322 | chromosome 5 open reading frame 22 | up | 1,25 | 4,00E-02 |
| ENSG00000058272 | PPP1R12A | 4659 | protein phosphatase 1 regulatory subunit 12A | up | 1,25 | 1,03E-02 |
| ENSG00000187742 | SECISBP2 | 79048 | SECIS binding protein 2 | up | 1,25 | 3,40E-02 |
| ENSG00000131375 | CAPN7 | 23473 | calpain 7 | up | 1,25 | 4,72E-02 |
| ENSG00000185728 | YTHDF3 | 253943 | YTH N6-methyladenosine RNA binding protein 3 | up | 1,25 | 4,67E-02 |
| ENSG00000116580 | GON4L | 54856 | gon-4 like | up | 1,25 | 3,69E-02 |
| ENSG00000089048 | ESF1 | 51575 | ESF1 nucleolar pre-rRNA processing protein homolog | up | 1,24 | 1,54E-02 |
| ENSG00000164609 | SLU7 | 10569 | SLU7 homolog, splicing factor | up | 1,24 | 3,64E-02 |
| ENSG00000033178 | UBA6 | 55236 | ubiquitin like modifier activating enzyme 6 | up | 1,24 | 1,11E-02 |
| ENSG00000154174 | TOMM70 | 9868 | translocase of outer mitochondrial membrane 70 | up | 1,24 | 3,74E-02 |
| ENSG00000130956 | HABP4 | 22927 | hyaluronan binding protein 4 | up | 1,24 | 3,62E-02 |
| ENSG00000170234 | PWWP2A | 114825 | PWWP domain containing 2A | up | 1,24 | 2,10E-02 |
| ENSG00000172262 | ZNF131 | 7690 | zinc finger protein 131 | up | 1,24 | 4,59E-02 |
| ENSG00000122958 | VPS26A | 9559 | VPS26, retromer complex component A | up | 1,24 | 4,07E-02 |
| ENSG00000114742 | WDR48 | 57599 | WD repeat domain 48 | up | 1,24 | 3,95E-02 |
| ENSG00000119318 | RAD23B | 5887 | RAD23 homolog B, nucleotide excision repair protein | up | 1,24 | 3,35E-02 |
| ENSG00000129566 | TEP1 | 7011 | telomerase associated protein 1 | up | 1,23 | 6,61E-03 |
| ENSG00000078674 | PCM1 | 5108 | pericentriolar material 1 | up | 1,23 | 5,27E-03 |
| ENSG00000165660 | ABRAXAS2 | 23172 | abraxas 2, BRISC complex subunit | up | 1,23 | 8,54E-03 |
| ENSG00000136738 | STAM | 8027 | signal transducing adaptor molecule | up | 1,23 | 3,53E-02 |
| ENSG00000143748 | NVL | 4931 | nuclear VCP-like | up | 1,23 | 2,64E-02 |
| ENSG00000164654 | MIOS | 54468 | meiosis regulator for oocyte development | up | 1,23 | 9,66E-03 |
| ENSG00000087053 | MTMR2 | 8898 | myotubularin related protein 2 | up | 1,23 | 2,62E-02 |
| ENSG00000159592 | GPBP1L1 | 60313 | GC-rich promoter binding protein 1 like 1 | up | 1,23 | 4,14E-02 |
| ENSG00000011566 | MAP4K3 | 8491 | mitogen-activated protein kinase kinase kinase kinase 3 | up | 1,23 | 4,83E-02 |
| ENSG00000100644 | HIF1A | 3091 | hypoxia inducible factor 1 alpha subunit | up | 1,23 | 4,67E-02 |
| ENSG00000197062 | ZSCAN26 | 7741 | zinc finger and SCAN domain containing 26 | up | 1,22 | 4,98E-02 |
| ENSG00000170852 | KBTBD2 | 25948 | kelch repeat and BTB domain containing 2 | up | 1,22 | 3,47E-02 |
| ENSG00000147677 | EIF3H | 8667 | eukaryotic translation initiation factor 3 subunit H | up | 1,22 | 3,84E-02 |
| ENSG00000151923 | TIAL1 | 7073 | TIA1 cytotoxic granule associated RNA binding protein like 1 | up | 1,22 | 3,57E-02 |
| ENSG00000153113 | CAST | 831 | calpastatin | up | 1,22 | 4,61E-02 |
| ENSG00000082212 | ME2 | 4200 | malic enzyme 2 | up | 1,22 | 4,89E-02 |
| ENSG00000125124 | BBS2 | 583 | Bardet-Biedl syndrome 2 | up | 1,21 | 3,77E-02 |
| ENSG00000035687 | ADSS | 159 | adenylosuccinate synthase | up | 1,21 | 2,47E-02 |
| ENSG00000135720 | DYNC1LI2 | 1783 | dynein cytoplasmic 1 light intermediate chain 2 | up | 1,21 | 2,48E-02 |
| ENSG00000011454 | GPR21 | 2844 | G protein-coupled receptor 21 | up | 1,21 | 2,65E-02 |
| ENSG00000011454 | RABGAP1 | 23637 | RAB GTPase activating protein 1 | up | 1,21 | 2,65E-02 |
| ENSG00000168538 | TRAPPC11 | 60684 | trafficking protein particle complex 11 | up | 1,20 | 1,57E-02 |
| ENSG00000115816 | CEBPZ | 10153 | CCAAT/enhancer binding protein zeta | up | 1,20 | 5,00E-02 |
| ENSG00000171490 | RSL1D1 | 26156 | ribosomal L1 domain containing 1 | up | 1,20 | 4,73E-02 |
| ENSG00000138029 | HADHB | 3032 | hydroxyacyl-CoA dehydrogenase trifunctional multienzyme complex subunit | up | 1,20 | 4,67E-02 |
| ENSG00000165288 | BRWD3 | 254065 | bromodomain and WD repeat domain containing 3 | up | 1,20 | 2,37E-02 |
| ENSG00000075151 | EIF4G3 | 8672 | eukaryotic translation initiation factor 4 gamma 3 | up | 1,19 | 4,60E-02 |
| ENSG00000130741 | EIF2S3 | 1968 | eukaryotic translation initiation factor 2 subunit gamma | up | 1,19 | 2,93E-02 |
| ENSG00000111596 | CNOT2 | 4848 | CCR4-NOT transcription complex subunit 2 | up | 1,19 | 4,40E-02 |
| ENSG00000159459 | UBR1 | 197131 | ubiquitin protein ligase E3 component n-recognin 1 | up | 1,19 | 2,30E-02 |
| ENSG00000116198 | CEP104 | 9731 | centrosomal protein 104 | up | 1,19 | 4,76E-02 |
| ENSG00000046604 | DSG2 | 1829 | desmoglein 2 | up | 1,19 | 2,35E-02 |
| ENSG00000135018 | UBQLN1 | 29979 | ubiquilin 1 | up | 1,19 | 4,56E-03 |
| ENSG00000005020 | SKAP2 | 8935 | src kinase associated phosphoprotein 2 | up | 1,19 | 3,74E-02 |
| ENSG00000156931 | VPS8 | 23355 | VPS8, CORVET complex subunit | up | 1,18 | 4,37E-02 |
| ENSG00000196504 | PRPF40A | 55660 | pre-mRNA processing factor 40 homolog A | up | 1,18 | 3,23E-02 |
| ENSG00000101596 | SMCHD1 | 23347 | structural maintenance of chromosomes flexible hinge domain containing 1 | up | 1,18 | 1,97E-02 |
| ENSG00000032742 | IFT88 | 8100 | intraflagellar transport 88 | up | 1,17 | 4,84E-02 |
| ENSG00000091140 | DLD | 1738 | dihydrolipoamide dehydrogenase | up | 1,16 | 4,53E-02 |
| ENSG00000106346 | USP42 | 84132 | ubiquitin specific peptidase 42 | up | 1,16 | 4,56E-02 |
| ENSG00000035928 | RFC1 | 5981 | replication factor C subunit 1 | up | 1,15 | 2,18E-02 |
| ENSG00000038532 | MFSD7 | 84179 | major facilitator superfamily domain containing 7 | down | 21,28 | 1,20E-02 |
| ENSG00000101444 | EPHA5 | 2044 | EPH receptor A5 | down | 18,87 | 1,97E-02 |
| ENSG00000144043 | CRISPLD1 | 83690 | cysteine rich secretory protein LCCL domain containing 1 | down | 17,24 | 2,24E-02 |
| ENSG00000011304 | TXLNGY | 246126 | taxilin gamma pseudogene, Y-linked | down | 16,95 | 2,92E-02 |
| ENSG00000159692 | SCN5A | 6331 | sodium voltage-gated channel alpha subunit 5 | down | 15,38 | 3,10E-02 |
| ENSG00000100403 | MEG3 | 55384 | maternally expressed 3 (non-protein coding) | down | 13,89 | 8,01E-03 |
| ENSG00000125970 | FAM155A | 728215 | family with sequence similarity 155 member A | down | 8,55 | 4,47E-02 |
| ENSG00000105323 | PLAT | 5327 | plasminogen activator, tissue type | down | 6,67 | 2,47E-02 |
| ENSG00000104964 | GSDME | 1687 | gasdermin E | down | 5,71 | 1,57E-02 |
| ENSG00000125149 | RPS4Y1 | 6192 | ribosomal protein S4, Y-linked 1 | down | 5,59 | 7,82E-03 |
| ENSG00000186187 | SUSD5 | 26032 | sushi domain containing 5 | down | 5,38 | 3,88E-02 |
| ENSG00000177370 | FGF2 | 2247 | fibroblast growth factor 2 | down | 5,35 | 3,10E-02 |
| ENSG00000108469 | DACT1 | 51339 | dishevelled binding antagonist of beta catenin 1 | down | 5,29 | 2,24E-02 |
| ENSG00000088256 | GYPC | 2995 | glycophorin C (Gerbich blood group) | down | 3,94 | 2,34E-02 |
| ENSG00000070047 | ADRA2A | 150 | adrenoceptor alpha 2A | down | 3,92 | 2,46E-02 |
| ENSG00000169592 | APLN | 8862 | apelin | down | 3,89 | 3,03E-02 |

| | | | | | | |
|---|---|---|---|---|---|---|
| ENSG00000123992 | BICC1 | 80114 | BicC family RNA binding protein 1 | down | 3,77 | 8,32E-03 |
| ENSG00000177963 | VAT1L | 57687 | vesicle amine transport 1 like | down | 3,75 | 1,12E-03 |
| ENSG00000163812 | EFEMP1 | 2202 | EGF containing fibulin extracellular matrix protein 1 | down | 3,55 | 4,54E-02 |
| ENSG00000167535 | HTRA1 | 5654 | HtrA serine peptidase 1 | down | 3,48 | 2,24E-02 |
| ENSG00000130764 | MAMLD1 | 10046 | mastermind like domain containing 1 | down | 3,36 | 4,56E-02 |
| ENSG00000140983 | FENDRR | 400550 | FOXF1 adjacent non-coding developmental regulatory RNA | down | 3,31 | 4,63E-02 |
| ENSG00000148300 | DAAM2 | 23500 | dishevelled associated activator of morphogenesis 2 | down | 3,21 | 3,09E-02 |
| ENSG00000085872 | SLC16A2 | 6567 | solute carrier family 16 member 2 | down | 3,17 | 2,47E-02 |
| ENSG00000141564 | TMPO-AS1 | 100128191 | TMPO antisense RNA 1 | down | 2,99 | 5,95E-03 |
| ENSG00000129250 | BDNF | 627 | brain derived neurotrophic factor | down | 2,98 | 1,16E-02 |
| ENSG00000125656 | GATA3 | 2625 | GATA binding protein 3 | down | 2,92 | 3,46E-02 |
| ENSG00000166228 | MGAT3 | 4248 | mannosyl (beta-1,4-)-glycoprotein beta-1,4-N-acetylglucosaminyltransferase | down | 2,92 | 7,17E-03 |
| ENSG00000241878 | HPDL | 84842 | 4-hydroxyphenylpyruvate dioxygenase like | down | 2,92 | 3,85E-03 |
| ENSG00000002834 | RIMBP3 | 85376 | RIMS binding protein 3 | down | 2,88 | 2,83E-02 |
| ENSG00000133313 | TP73 | 7161 | tumor protein p73 | down | 2,86 | 3,40E-02 |
| ENSG00000105993 | L1CAM | 3897 | L1 cell adhesion molecule | down | 2,86 | 3,10E-02 |
| ENSG00000101849 | HOXB8 | 3218 | homeobox B8 | down | 2,86 | 1,77E-02 |
| ENSG00000108518 | UHRF1 | 29128 | ubiquitin like with PHD and ring finger domains 1 | down | 2,85 | 1,97E-02 |
| ENSG00000163161 | RASL11A | 387496 | RAS like family 11 member A | down | 2,82 | 3,58E-02 |
| ENSG00000137409 | PDE4B | 5142 | phosphodiesterase 4B | down | 2,82 | 4,91E-02 |
| ENSG00000105618 | NAT8L | 339983 | N-acetyltransferase 8 like | down | 2,79 | 1,70E-02 |
| ENSG00000204673 | AXIN2 | 8313 | axin 2 | down | 2,79 | 3,67E-03 |
| ENSG00000090316 | VASH1 | 22846 | vasohibin 1 | down | 2,76 | 6,96E-03 |
| ENSG00000135632 | HOXA13 | 3209 | homeobox A13 | down | 2,75 | 2,80E-02 |
| ENSG00000274523 | NSG1 | 27065 | neuronal vesicle trafficking associated 1 | down | 2,74 | 4,17E-02 |
| ENSG00000154832 | VSNL1 | 7447 | visinin like 1 | down | 2,70 | 3,86E-02 |
| ENSG00000172663 | CBX2 | 84733 | chromobox 2 | down | 2,70 | 8,10E-03 |
| ENSG00000135587 | RIBC2 | 26150 | RIB43A domain with coiled-coils 2 | down | 2,70 | 3,33E-02 |
| ENSG00000087088 | LGR5 | 8549 | leucine rich repeat containing G protein-coupled receptor 5 | down | 2,70 | 3,09E-02 |
| ENSG00000136856 | SOWAHD | 347454 | sosondowah ankyrin repeat domain family member D | down | 2,67 | 3,34E-02 |
| ENSG00000119401 | VWA1 | 64856 | von Willebrand factor A domain containing 1 | down | 2,67 | 4,01E-03 |
| ENSG00000114353 | SP5 | 389058 | Sp5 transcription factor | down | 2,65 | 1,04E-02 |
| ENSG00000100034 | CNIH2 | 254263 | cornichon family AMPA receptor auxiliary protein 2 | down | 2,64 | 1,06E-02 |
| ENSG00000103042 | OLFML2A | 169611 | olfactomedin like 2A | down | 2,62 | 1,89E-02 |
| ENSG00000162585 | ARL10 | 285598 | ADP ribosylation factor like GTPase 10 | down | 2,61 | 2,93E-02 |
| ENSG00000175634 | BMP4 | 652 | bone morphogenetic protein 4 | down | 2,60 | 1,26E-02 |
| ENSG00000140474 | RHPN1-AS1 | 78998 | RHPN1 antisense RNA 1 (head to head) | down | 2,54 | 1,01E-02 |
| ENSG00000181472 | PAQR4 | 124222 | progestin and adipoQ receptor family member 4 | down | 2,52 | 1,01E-02 |
| ENSG00000196976 | E2F8 | 79733 | E2F transcription factor 8 | down | 2,48 | 3,90E-02 |
| ENSG00000160209 | TRIB2 | 28951 | tribbles pseudokinase 2 | down | 2,43 | 3,49E-02 |
| ENSG00000145912 | GPR161 | 23432 | G protein-coupled receptor 161 | down | 2,43 | 1,50E-03 |
| ENSG00000160209 | B4GAT1 | 11041 | beta-1,4-glucuronyltransferase 1 | down | 2,42 | 1,50E-02 |
| ENSG00000080189 | LHX4 | 89884 | LIM homeobox 4 | down | 2,42 | 1,29E-02 |
| ENSG00000160959 | ZNF792 | 126375 | zinc finger protein 792 | down | 2,40 | 1,00E-02 |
| ENSG00000158545 | FGFBP1 | 9982 | fibroblast growth factor binding protein 1 | down | 2,40 | 2,73E-02 |
| ENSG00000047315 | B3GNT7 | 93010 | UDP-GlcNAc:betaGal beta-1,3-N-acetylglucosaminyltransferase 7 | down | 2,40 | 3,48E-02 |
| ENSG00000104973 | DLX1 | 1745 | distal-less homeobox 1 | down | 2,38 | 1,88E-02 |
| ENSG00000066117 | RASSF5 | 83593 | Ras association domain family member 5 | down | 2,36 | 2,59E-02 |
| ENSG00000100243 | HOXD13 | 3239 | homeobox D13 | down | 2,36 | 1,81E-02 |
| ENSG00000227057 | LINC01124 | 440925 | long intergenic non-protein coding RNA 1124 | down | 2,36 | 3,58E-02 |
| ENSG00000115317 | DDN-AS1 | 105369758 | DDN and PRKAG1 antisense RNA 1 | down | 2,35 | 1,68E-02 |
| ENSG00000158941 | PHGDH | 26227 | phosphoglycerate dehydrogenase | down | 2,34 | 2,54E-02 |
| ENSG00000169692 | ISL2 | 64843 | ISL LIM homeobox 2 | down | 2,33 | 1,37E-02 |
| ENSG00000072958 | GPRC5B | 51704 | G protein-coupled receptor class C group 5 member B | down | 2,32 | 4,64E-02 |
| ENSG00000148343 | CDX2 | 1045 | caudal type homeobox 2 | down | 2,31 | 4,63E-02 |
| ENSG00000177303 | HOXA-AS3 | 100133311 | HOXA cluster antisense RNA 3 | down | 2,30 | 6,10E-03 |
| ENSG00000186635 | H2AFY2 | 55506 | H2A histone family member Y2 | down | 2,30 | 4,64E-02 |
| ENSG00000103266 | RHOBTB2 | 23221 | Rho related BTB domain containing 2 | down | 2,29 | 7,15E-03 |
| ENSG00000124535 | DFFB | 1677 | DNA fragmentation factor subunit beta | down | 2,29 | 3,17E-02 |
| ENSG00000144021 | C2orf72 | 257407 | chromosome 2 open reading frame 72 | down | 2,29 | 4,50E-02 |
| ENSG00000105220 | ENHO | 375704 | energy homeostasis associated | down | 2,28 | 2,41E-03 |
| ENSG00000130726 | PCOTH | 542767 | Pro-X-Gly collagen triple helix like repeat containing | down | 2,28 | 6,61E-03 |
| ENSG00000184900 | LFNG | 3955 | LFNG O-fucosylpeptide 3-beta-N-acetylglucosaminyltransferase | down | 2,26 | 1,16E-02 |
| ENSG00000108883 | XKRX | 402415 | XK related, X-linked | down | 2,26 | 1,57E-02 |
| ENSG00000154122 | NNMT | 4837 | nicotinamide N-methyltransferase | down | 2,26 | 3,06E-02 |
| ENSG00000158669 | SLITRK6 | 84189 | SLIT and NTRK like family member 6 | down | 2,25 | 3,71E-02 |
| ENSG00000171453 | ASF1B | 55723 | anti-silencing function 1B histone chaperone | down | 2,25 | 1,42E-02 |
| ENSG00000167112 | AQP1 | 358 | aquaporin 1 (Colton blood group) | down | 2,24 | 2,61E-02 |
| ENSG00000110107 | LARGE2 | 120071 | LARGE xylosyl- and glucuronyltransferase 2 | down | 2,23 | 3,90E-02 |
| ENSG00000139998 | SERTAD4 | 56256 | SERTA domain containing 4 | down | 2,23 | 2,42E-02 |
| ENSG00000103653 | CSPG4 | 1464 | chondroitin sulfate proteoglycan 4 | down | 2,22 | 4,00E-02 |
| ENSG00000141698 | ZNF793 | 390927 | zinc finger protein 793 | down | 2,21 | 4,12E-02 |
| ENSG00000172531 | SKP2 | 6502 | S-phase kinase associated protein 2 | down | 2,21 | 3,69E-02 |
| ENSG00000148308 | PCDHGA4 | 56111 | protocadherin gamma subfamily A, 4 | down | 2,21 | 6,25E-03 |
| ENSG00000102977 | FAM78A | 286336 | family with sequence similarity 78 member A | down | 2,20 | 1,01E-02 |
| ENSG00000118046 | ZNF618 | 114991 | zinc finger protein 618 | down | 2,19 | 4,39E-02 |
| ENSG00000130826 | BEND3 | 57673 | BEN domain containing 3 | down | 2,19 | 1,25E-02 |
| ENSG00000131899 | DHCR24 | 1718 | 24-dehydrocholesterol reductase | down | 2,17 | 4,89E-02 |
| ENSG00000165802 | FOXD2-AS1 | 84793 | FOXD2 adjacent opposite strand RNA 1 | down | 2,17 | 1,28E-02 |
| ENSG00000227500 | GINS1 | 9837 | GINS complex subunit 1 | down | 2,17 | 3,10E-02 |
| ENSG00000105676 | FZD4 | 8322 | frizzled class receptor 4 | down | 2,17 | 3,82E-02 |
| ENSG00000198931 | SIRPA | 140885 | signal regulatory protein alpha | down | 2,17 | 1,00E-02 |
| ENSG00000169718 | SDC1 | 6382 | syndecan 1 | down | 2,17 | 1,62E-02 |
| ENSG00000178252 | NES | 10763 | nestin | down | 2,17 | 2,68E-02 |
| ENSG00000189306 | ZNF367 | 195828 | zinc finger protein 367 | down | 2,17 | 4,13E-02 |
| ENSG00000181830 | NPFFR1 | 64106 | neuropeptide FF receptor 1 | down | 2,16 | 3,35E-02 |
| ENSG00000130165 | CCNF | 899 | cyclin F | down | 2,16 | 6,61E-03 |
| ENSG00000149761 | GATA2 | 2624 | GATA binding protein 2 | down | 2,16 | 2,07E-02 |
| ENSG00000174939 | NYNRIN | 57523 | NYN domain and retroviral integrase containing | down | 2,15 | 3,38E-02 |
| ENSG00000168936 | PCNA | 5111 | proliferating cell nuclear antigen | down | 2,14 | 4,63E-02 |
| ENSG00000169188 | BOLA3-AS1 | 100507171 | BOLA3 antisense RNA 1 (head to head) | down | 2,14 | 3,53E-02 |
| ENSG00000142207 | UNG | 7374 | uracil DNA glycosylase | down | 2,13 | 1,95E-02 |
| ENSG00000148730 | C21orf58 | 54058 | chromosome 21 open reading frame 58 | down | 2,13 | 2,18E-02 |
| ENSG00000177663 | LMNB1 | 4001 | lamin B1 | down | 2,12 | 3,86E-02 |
| ENSG00000232677 | DEPP1 | 11067 | DEPP1, autophagy regulator | down | 2,11 | 4,04E-02 |
| ENSG00000105355 | H2AFX | 3014 | H2A histone family member X | down | 2,11 | 1,30E-03 |
| ENSG00000144579 | FBXL16 | 146330 | F-box and leucine rich repeat protein 16 | down | 2,11 | 4,03E-03 |
| ENSG00000176248 | ZNF540 | 163255 | zinc finger protein 540 | down | 2,09 | 2,24E-03 |
| ENSG00000126062 | MYORG | 57462 | myogenesis regulating glycosidase (putative) | down | 2,09 | 1,26E-02 |
| ENSG00000126003 | KCNC4 | 3749 | potassium voltage-gated channel subfamily C member 4 | down | 2,09 | 3,27E-02 |
| ENSG00000075240 | ARID5B | 84159 | AT-rich interaction domain 5B | down | 2,09 | 3,59E-02 |
| ENSG00000165271 | TICRR | 90381 | TOPBP1 interacting checkpoint and replication regulator | down | 2,08 | 1,18E-02 |
| ENSG00000165684 | ADAM11 | 4185 | ADAM metallopeptidase domain 11 | down | 2,07 | 1,87E-02 |
| ENSG00000165688 | ADM5 | 199800 | adrenomedullin 5 (putative) | down | 2,07 | 2,20E-02 |
| ENSG00000130254 | THEM6 | 51337 | thioesterase superfamily member 6 | down | 2,07 | 5,68E-03 |
| ENSG00000167815 | SPC24 | 147841 | SPC24, NDC80 kinetochore complex component | down | 2,07 | 3,62E-02 |
| ENSG00000164366 | MCM3 | 4172 | minichromosome maintenance complex component 3 | down | 2,07 | 4,13E-02 |
| ENSG00000167394 | SH2B2 | 10603 | SH2B adaptor protein 2 | down | 2,07 | 1,95E-02 |
| ENSG00000134780 | LRRC20 | 55222 | leucine rich repeat containing 20 | down | 2,06 | 2,51E-02 |
| ENSG00000096746 | TNFAIP8L3 | 388121 | TNF alpha induced protein 8 like 3 | down | 2,06 | 1,40E-02 |
| ENSG00000198131 | SQLE | 6713 | squalene epoxidase | down | 2,05 | 1,00E-02 |
| ENSG00000142082 | PIK3R1 | 5295 | phosphoinositide-3-kinase regulatory subunit 1 | down | 2,04 | 8,41E-03 |
| ENSG00000014216 | PKDCC | 91461 | protein kinase domain containing, cytoplasmic | down | 2,04 | 2,29E-02 |
| ENSG00000063244 | FAM171A2 | 284069 | family with sequence similarity 171 member A2 | down | 2,04 | 1,20E-02 |
| ENSG00000103365 | FEN1 | 2237 | flap structure-specific endonuclease 1 | down | 2,04 | 1,61E-02 |
| ENSG00000106263 | ERCC6L | 54821 | ERCC excision repair 6 like, spindle assembly checkpoint helicase | down | 2,03 | 2,61E-02 |
| ENSG00000149480 | ZNF488 | 118738 | zinc finger protein 488 | down | 2,03 | 3,48E-02 |
| ENSG00000174903 | AGAP2 | 116986 | ArfGAP with GTPase domain, ankyrin repeat and PH domain 2 | down | 2,02 | 3,75E-02 |
| ENSG00000030110 | EMILIN1 | 11117 | elastin microfibril interfacer 1 | down | 2,02 | 1,50E-02 |
| ENSG00000149547 | MCM2 | 4171 | minichromosome maintenance complex component 2 | down | 2,01 | 1,43E-02 |
| ENSG00000005007 | SAC3D1 | 29901 | SAC3 domain containing 1 | down | 2,01 | 3,22E-03 |
| ENSG00000157014 | FHOD1 | 29109 | formin homology 2 domain containing 1 | down | 2,00 | 1,54E-02 |
| ENSG00000177830 | EFNA3 | 1944 | ephrin A3 | down | 2,00 | 4,89E-02 |
| ENSG00000110104 | CAMK2N2 | 94032 | calcium/calmodulin dependent protein kinase II inhibitor 2 | down | 2,00 | 1,47E-02 |
| ENSG00000150990 | FAM84A | 151354 | family with sequence similarity 84 member A | down | 2,00 | 3,93E-02 |
| ENSG00000169230 | DHCR7 | 1717 | 7-dehydrocholesterol reductase | down | 2,00 | 2,14E-03 |
| ENSG00000196363 | TREX2 | 11219 | three prime repair exonuclease 2 | down | 2,00 | 3,47E-02 |
| ENSG00000129347 | KHK | 3795 | ketohexokinase | down | 2,00 | 2,31E-02 |
| ENSG00000103495 | SPHK1 | 8877 | sphingosine kinase 1 | down | 2,00 | 2,35E-02 |
| ENSG00000164081 | LOC100287042 | 100287042 | uncharacterized LOC100287042 | down | 2,00 | 1,12E-03 |
| ENSG00000153406 | SYDE1 | 85360 | synapse defective Rho GTPase homolog 1 | down | 1,99 | 3,41E-03 |
| ENSG00000114554 | MVK | 4598 | mevalonate kinase | down | 1,99 | 3,37E-03 |
| ENSG00000159335 | WWTR1 | 25937 | WW domain containing transcription regulator 1 | down | 1,99 | 1,00E-02 |
| ENSG00000111669 | FAM213B | 127281 | family with sequence similarity 213 member B | down | 1,99 | 1,18E-02 |
| ENSG00000090565 | BAIAP2-AS1 | 440465 | BAIAP2 antisense RNA 1 (head to head) | down | 1,98 | 1,47E-02 |
| ENSG00000105221 | XYLT2 | 64132 | xylosyltransferase 2 | down | 1,98 | 2,04E-02 |
| ENSG00000167925 | SBK1 | 388228 | SH3 domain binding kinase 1 | down | 1,98 | 3,92E-02 |
| ENSG00000068650 | ADAMTS9 | 56999 | ADAM metallopeptidase with thrombospondin type 1 motif 9 | down | 1,98 | 4,91E-02 |
| ENSG00000104884 | TMEM109 | 79073 | transmembrane protein 109 | down | 1,97 | 1,42E-02 |
| ENSG00000185085 | HOXB7 | 3217 | homeobox B7 | down | 1,96 | 3,18E-03 |
| ENSG00000238227 | ARAP3 | 64411 | ArfGAP with RhoGAP domain, ankyrin repeat and PH domain 3 | down | 1,96 | 6,61E-03 |
| ENSG00000013306 | FASN | 2194 | fatty acid synthase | down | 1,96 | 8,32E-03 |
| ENSG00000130560 | ARHGEF39 | 84904 | Rho guanine nucleotide exchange factor 39 | down | 1,96 | 3,08E-02 |
| ENSG00000135916 | MARVELD1 | 83742 | MARVEL domain containing 1 | down | 1,95 | 1,38E-02 |
| ENSG00000167085 | KIFC1 | 3833 | kinesin family member C1 | down | 1,95 | 2,60E-02 |
| ENSG00000146828 | GINS3 | 64785 | GINS complex subunit 3 | down | 1,95 | 1,67E-02 |
| ENSG00000160767 | KIF18B | 146909 | kinesin family member 18B | down | 1,94 | 3,01E-02 |
| ENSG00000144455 | TMEM38A | 79041 | transmembrane protein 38A | down | 1,94 | 1,99E-02 |
| ENSG00000172757 | ENC1 | 8507 | ectodermal-neural cortex 1 | down | 1,94 | 4,67E-02 |
| ENSG00000128626 | INSIG1 | 3638 | insulin induced gene 1 | down | 1,94 | 1,61E-02 |
| ENSG00000181396 | P3H4 | 10609 | prolyl 3-hydroxylase family member 4 (non-enzymatic) | down | 1,93 | 1,10E-02 |
| ENSG00000247626 | SLC19A1 | 6573 | solute carrier family 19 member 1 | down | 1,93 | 4,76E-03 |
| ENSG00000130204 | CDCA4 | 55038 | cell division cycle associated 4 | down | 1,93 | 4,12E-03 |
| ENSG00000132507 | MPP2 | 4355 | membrane palmitoylated protein 2 | down | 1,93 | 3,82E-02 |
| ENSG00000176973 | CHST14 | 113189 | carbohydrate sulfotransferase 14 | down | 1,92 | 4,01E-03 |
| ENSG00000103415 | ZNF503-AS2 | 100131213 | ZNF503 antisense RNA 2 | down | 1,92 | 6,94E-03 |
| ENSG00000011485 | CDC25A | 993 | cell division cycle 25A | down | 1,92 | 1,49E-02 |
| ENSG00000100324 | CTXN1 | 404217 | cortexin 1 | down | 1,92 | 3,96E-03 |
| ENSG00000144580 | ALDH1B1 | 219 | aldehyde dehydrogenase 1 family member B1 | down | 1,92 | 9,37E-03 |
| ENSG00000160216 | TFAP4 | 7023 | transcription factor AP-4 | down | 1,91 | 1,28E-02 |
| ENSG00000100395 | S1PR2 | 9294 | sphingosine-1-phosphate receptor 2 | down | 1,91 | 1,42E-02 |
| ENSG00000007520 | RGCC | 28984 | regulator of cell cycle | down | 1,90 | 2,18E-02 |
| ENSG00000155189 | HOXB5 | 3215 | homeobox B5 | down | 1,90 | 2,91E-02 |
| ENSG00000169972 | PSRC1 | 84722 | proline and serine rich coiled-coil 1 | down | 1,90 | 3,19E-02 |
| ENSG00000134287 | IRS1 | 3667 | insulin receptor substrate 1 | down | 1,89 | 6,25E-03 |
| ENSG00000135966 | GEMIN4 | 50628 | gem nuclear organelle associated protein 4 | down | 1,89 | 5,68E-03 |
| ENSG00000068654 | LOC101927751 | 101927751 | uncharacterized LOC101927751 | down | 1,89 | 3,30E-02 |
| ENSG00000148803 | MANEAL | 149175 | mannosidase endo-alpha like | down | 1,89 | 4,58E-03 |
| ENSG00000006744 | TTC30B | 150737 | tetratricopeptide repeat domain 30B | down | 1,89 | 3,05E-02 |
| ENSG00000160271 | ESPL1 | 9700 | extra spindle pole bodies like 1, separase | down | 1,88 | 3,18E-02 |
| ENSG00000174915 | SOX12 | 6666 | SRY-box 12 | down | 1,88 | 1,81E-02 |
| ENSG00000141570 | SCD | 6319 | stearoyl-CoA desaturase | down | 1,88 | 3,59E-02 |
| ENSG00000106628 | PRR19 | 284338 | proline rich 19 | down | 1,88 | 1,24E-02 |
| ENSG00000115207 | CYP2U1 | 113612 | cytochrome P450 family 2 subfamily U member 1 | down | 1,88 | 2,20E-02 |
| ENSG00000093010 | GDF11 | 10220 | growth differentiation factor 11 | down | 1,88 | 4,03E-03 |
| ENSG00000099817 | SPRY1 | 10252 | sprouty RTK signaling antagonist 1 | down | 1,88 | 2,51E-02 |
| ENSG00000110651 | ABHD15 | 116236 | abhydrolase domain containing 15 | down | 1,87 | 9,48E-03 |
| ENSG00000168040 | ASB13 | 79754 | ankyrin repeat and SOCS box containing 13 | down | 1,87 | 4,56E-03 |
| ENSG00000105447 | TRIM47 | 91107 | tripartite motif containing 47 | down | 1,86 | 4,36E-02 |
| ENSG00000156471 | TEDC2 | 80178 | tubulin epsilon and delta complex 2 | down | 1,86 | 1,59E-02 |
| ENSG00000168395 | CNPY4 | 245812 | canopy FGF signaling regulator 4 | down | 1,86 | 3,50E-02 |
| ENSG00000178927 | IGFBP7 | 3490 | insulin like growth factor binding protein 7 | down | 1,86 | 2,80E-02 |
| ENSG00000178188 | DUBR | 344595 | DPPA2 upstream binding RNA | down | 1,86 | 3,40E-02 |
| ENSG00000188735 | RASL10B | 91608 | RAS like family 10 member B | down | 1,85 | 9,37E-03 |
| ENSG00000119185 | LRP8 | 7804 | LDL receptor related protein 8 | down | 1,85 | 4,38E-02 |
| ENSG00000165752 | NRGN | 4900 | neurogranin | down | 1,85 | 9,47E-03 |
| ENSG00000119574 | DGCR8 | 54487 | DGCR8, microprocessor complex subunit | down | 1,84 | 1,01E-02 |
| ENSG00000184508 | STMN3 | 50861 | stathmin 3 | down | 1,84 | 1,29E-02 |
| ENSG00000247077 | OBSCN-AS1 | 574407 | OBSCN antisense RNA 1 | down | 1,84 | 1,41E-02 |
| ENSG00000183751 | THAP11 | 57215 | THAP domain containing 11 | down | 1,83 | 2,24E-03 |
| ENSG00000183963 | TMEM102 | 284114 | transmembrane protein 102 | down | 1,83 | 7,17E-03 |
| ENSG00000130749 | CAMKK1 | 84254 | calcium/calmodulin dependent protein kinase kinase 1 | down | 1,83 | 2,57E-02 |
| ENSG00000162729 | SLC25A10 | 1468 | solute carrier family 25 member 10 | down | 1,83 | 5,12E-03 |
| ENSG00000170043 | MCM5 | 4174 | minichromosome maintenance complex component 5 | down | 1,83 | 2,23E-02 |
| ENSG00000174775 | TCF19 | 6941 | transcription factor 19 | down | 1,82 | 2,14E-02 |
| ENSG00000198917 | KANK2 | 25959 | KN motif and ankyrin repeat domains 2 | down | 1,82 | 2,00E-02 |
| ENSG00000110442 | LYPD6 | 130574 | LY6/PLAUR domain containing 6 | down | 1,82 | 3,33E-02 |
| ENSG00000100364 | ERVMER34-1 | 100288413 | endogenous retrovirus group MER34 member 1, envelope | down | 1,82 | 4,42E-02 |
| ENSG00000140326 | NR2F1 | 7025 | nuclear receptor subfamily 2 group F member 1 | down | 1,82 | 1,74E-02 |
| ENSG00000197696 | NHLRC1 | 378884 | NHL repeat containing E3 ubiquitin protein ligase 1 | down | 1,81 | 1,85E-03 |
| ENSG00000111667 | SLC2A6 | 11182 | solute carrier family 2 member 6 | down | 1,81 | 1,03E-02 |
| ENSG00000103502 | LOC101928728 | 101928728 | uncharacterized LOC101928728 | down | 1,81 | 4,41E-02 |
| ENSG00000164050 | HAPLN3 | 145864 | hyaluronan and proteoglycan link protein 3 | down | 1,81 | 1,37E-02 |
| ENSG00000152990 | OGFRP1 | 388906 | opioid growth factor receptor pseudogene 1 | down | 1,81 | 2,05E-02 |
| ENSG00000167173 | C19orf57 | 79173 | chromosome 19 open reading frame 57 | down | 1,81 | 4,01E-02 |
| ENSG00000104915 | C1QTNF6 | 114904 | C1q and TNF related 6 | down | 1,81 | 2,35E-02 |
| ENSG00000174243 | CALHM2 | 51063 | calcium homeostasis modulator family member 2 | down | 1,81 | 3,18E-02 |
| ENSG00000104812 | TCF21 | 6943 | transcription factor 21 | down | 1,80 | 3,05E-02 |
| ENSG00000167515 | JADE2 | 23338 | jade family PHD finger 2 | down | 1,80 | 8,91E-03 |
| ENSG00000173914 | SLC40A1 | 30061 | solute carrier family 40 member 1 | down | 1,80 | 2,31E-02 |
| ENSG00000189319 | TMEM201 | 199953 | transmembrane protein 201 | down | 1,80 | 4,24E-03 |
| ENSG00000148362 | POLR3H | 171568 | RNA polymerase III subunit H | down | 1,80 | 7,08E-04 |
| ENSG00000224877 | RAB3A | 5864 | RAB3A, member RAS oncogene family | down | 1,80 | 4,63E-02 |
| ENSG00000179041 | PRSS35 | 167681 | protease, serine 35 | down | 1,79 | 3,40E-02 |
| ENSG00000172830 | ZNF551 | 90233 | zinc finger protein 551 | down | 1,79 | 1,19E-02 |
| ENSG00000214706 | ARRB1 | 408 | arrestin beta 1 | down | 1,79 | 2,43E-02 |
| ENSG00000269556 | CARHSP1 | 23589 | calcium regulated heat stable protein 1 | down | 1,79 | 7,72E-03 |

| | | | | | | |
|---|---|---|---|---|---|---|
| ENSG00000126456 | ANKRD9 | 122416 | ankyrin repeat domain 9 | down | 1,79 | 5,07E-03 |
| ENSG00000160072 | TMEM147-AS1 | 100506469 | TMEM147 antisense RNA 1 | down | 1,79 | 1,38E-02 |
| ENSG00000172534 | MAT2A | 4144 | methionine adenosyltransferase 2A | down | 1,79 | 2,93E-02 |
| ENSG00000084090 | EMP2 | 2013 | epithelial membrane protein 2 | down | 1,79 | 3,46E-02 |
| ENSG00000160208 | TUBA1B | 10376 | tubulin alpha 1b | down | 1,78 | 2,31E-02 |
| ENSG00000188636 | LRRC45 | 201255 | leucine rich repeat containing 45 | down | 1,78 | 2,96E-02 |
| ENSG00000171307 | AMER1 | 139285 | APC membrane recruitment protein 1 | down | 1,78 | 4,16E-02 |
| ENSG00000226287 | CELSR2 | 1952 | cadherin EGF LAG seven-pass G-type receptor 2 | down | 1,78 | 2,11E-02 |
| ENSG00000147804 | INCENP | 3619 | inner centromere protein | down | 1,78 | 2,09E-02 |
| ENSG00000105364 | VANGL1 | 81839 | VANGL planar cell polarity protein 1 | down | 1,78 | 1,08E-02 |
| ENSG00000204514 | LPCAT4 | 254531 | lysophosphatidylcholine acyltransferase 4 | down | 1,78 | 2,53E-02 |
| ENSG00000008838 | TACC3 | 10460 | transforming acidic coiled-coil containing protein 3 | down | 1,78 | 4,69E-02 |
| ENSG00000132768 | CMTM1 | 113540 | CKLF like MARVEL transmembrane domain containing 1 | down | 1,77 | 1,97E-02 |
| ENSG00000128245 | TOMM40L | 84134 | translocase of outer mitochondrial membrane 40 like | down | 1,77 | 2,73E-03 |
| ENSG00000172009 | PYCR3 | 65263 | pyrroline-5-carboxylate reductase 3 | down | 1,77 | 1,12E-03 |
| ENSG00000213965 | MCAM | 4162 | melanoma cell adhesion molecule | down | 1,77 | 3,27E-02 |
| ENSG00000214160 | TFDP1 | 7027 | transcription factor Dp-1 | down | 1,77 | 1,08E-02 |
| ENSG00000162302 | DPYSL2 | 1808 | dihydropyrimidinase like 2 | down | 1,77 | 2,20E-02 |
| ENSG00000051523 | TGIF2 | 60436 | TGFB induced factor homeobox 2 | down | 1,76 | 2,54E-02 |
| ENSG00000182240 | GAS2L1 | 10634 | growth arrest specific 2 like 1 | down | 1,76 | 1,42E-03 |
| ENSG00000131165 | ATF5 | 22809 | activating transcription factor 5 | down | 1,76 | 1,31E-04 |
| ENSG00000172366 | ATP2A1-AS1 | 100289092 | ATP2A1 antisense RNA 1 | down | 1,76 | 3,44E-02 |
| ENSG00000172366 | CBX7 | 23492 | chromobox 7 | down | 1,76 | 5,68E-03 |
| ENSG00000107815 | LPCAT1 | 79888 | lysophosphatidylcholine acyltransferase 1 | down | 1,76 | 2,61E-02 |
| ENSG00000164077 | AK4 | 205 | adenylate kinase 4 | down | 1,76 | 4,87E-02 |
| ENSG00000186350 | RPUSD2 | 27079 | RNA pseudouridylate synthase domain containing 2 | down | 1,76 | 1,30E-03 |
| ENSG00000224051 | GPRIN1 | 114787 | G protein regulated inducer of neurite outgrowth 1 | down | 1,76 | 2,39E-02 |
| ENSG00000119383 | PRADC1 | 84279 | protease associated domain containing 1 | down | 1,76 | 2,82E-03 |
| ENSG00000180035 | WNT10B | 7480 | Wnt family member 10B | down | 1,76 | 4,79E-02 |
| ENSG00000053372 | C1GALT1C1L | 728819 | C1GALT1-specific chaperone 1 like | down | 1,76 | 2,95E-02 |
| ENSG00000124217 | NUDT8 | 254552 | nudix hydrolase 8 | down | 1,75 | 2,24E-03 |
| ENSG00000104299 | NCS1 | 23413 | neuronal calcium sensor 1 | down | 1,75 | 1,02E-02 |
| ENSG00000163513 | TTYH3 | 80727 | tweety family member 3 | down | 1,75 | 6,96E-03 |
| ENSG00000114491 | SCAMP5 | 192683 | secretory carrier membrane protein 5 | down | 1,75 | 4,80E-02 |
| ENSG00000132382 | FZD1 | 8321 | frizzled class receptor 1 | down | 1,75 | 2,85E-02 |
| ENSG00000133275 | NDRG2 | 57447 | NDRG family member 2 | down | 1,75 | 4,63E-02 |
| ENSG00000105552 | HOXA11-AS | 221883 | HOXA11 antisense RNA | down | 1,75 | 1,87E-02 |
| ENSG00000160972 | KCTD14 | 65987 | potassium channel tetramerization domain containing 14 | down | 1,74 | 4,33E-02 |
| ENSG00000106077 | SAMD1 | 90378 | sterile alpha motif domain containing 1 | down | 1,74 | 1,57E-02 |
| ENSG00000172057 | GPR153 | 387509 | G protein-coupled receptor 153 | down | 1,74 | 1,86E-02 |
| ENSG00000197586 | MFSD13A | 79847 | major facilitator superfamily domain containing 13A | down | 1,73 | 2,25E-02 |
| ENSG00000103150 | LRFN3 | 79414 | leucine rich repeat and fibronectin type III domain containing 3 | down | 1,73 | 3,23E-02 |
| ENSG00000175602 | DOLK | 22845 | dolichol kinase | down | 1,73 | 2,75E-03 |
| ENSG00000067225 | NT5DC2 | 64943 | 5'-nucleotidase domain containing 2 | down | 1,73 | 1,12E-03 |
| ENSG00000138074 | DHRS13 | 147015 | dehydrogenase/reductase 13 | down | 1,73 | 3,95E-02 |
| ENSG00000103260 | ZBED1 | 9189 | zinc finger BED-type containing 1 | down | 1,72 | 6,80E-04 |
| ENSG00000164818 | LINC01963 | 150967 | long intergenic non-protein coding RNA 1963 | down | 1,72 | 4,06E-02 |
| ENSG00000033011 | FAM86JP | 100125556 | family with sequence similarity 86, member A pseudogene | down | 1,72 | 1,58E-02 |
| ENSG00000143793 | SEMA4C | 54910 | semaphorin 4C | down | 1,72 | 8,32E-03 |
| ENSG00000181588 | COQ8A | 56997 | coenzyme Q8A | down | 1,72 | 1,12E-02 |
| ENSG00000116649 | POLA2 | 23649 | DNA polymerase alpha 2, accessory subunit | down | 1,72 | 2,47E-02 |
| ENSG00000109062 | GATS | 352954 | GATS, stromal antigen 3 opposite strand | down | 1,72 | 2,51E-02 |
| ENSG00000114383 | ZNF530 | 348327 | zinc finger protein 530 | down | 1,72 | 1,40E-02 |
| ENSG00000215012 | EBP | 10682 | emopamil binding protein (sterol isomerase) | down | 1,71 | 2,00E-02 |
| ENSG00000147383 | FRAT2 | 23401 | FRAT2, WNT signaling pathway regulator | down | 1,71 | 2,53E-04 |
| ENSG00000139546 | RPP25 | 54913 | ribonuclease P and MRP subunit p25 | down | 1,71 | 4,35E-03 |
| ENSG00000153551 | ITPKB | 3707 | inositol-trisphosphate 3-kinase B | down | 1,71 | 1,97E-02 |
| ENSG00000180822 | ZNF552 | 79818 | zinc finger protein 552 | down | 1,71 | 1,42E-02 |
| ENSG00000254087 | HYAL2 | 8692 | hyaluronoglucosaminidase 2 | down | 1,71 | 7,17E-03 |
| ENSG00000132394 | BAHCC1 | 57597 | BAH domain and coiled-coil containing 1 | down | 1,71 | 3,01E-02 |
| ENSG00000085117 | FGFR3 | 2261 | fibroblast growth factor receptor 3 | down | 1,70 | 4,61E-02 |
| ENSG00000142453 | NOV | 4856 | nephroblastoma overexpressed | down | 1,70 | 3,73E-02 |
| ENSG00000169925 | DNMT3B | 1789 | DNA methyltransferase 3 beta | down | 1,70 | 3,62E-02 |
| ENSG00000177674 | TOR3A | 64222 | torsin family 3 member A | down | 1,70 | 1,21E-02 |
| ENSG00000132793 | UBL4A | 8266 | ubiquitin like 4A | down | 1,70 | 2,24E-03 |
| ENSG00000076662 | OSR2 | 116039 | odd-skipped related transciption factor 2 | down | 1,70 | 4,85E-02 |
| ENSG00000104853 | HOXB6 | 3216 | homeobox B6 | down | 1,70 | 5,84E-03 |
| ENSG00000161847 | DCLRE1B | 64858 | DNA cross-link repair 1B | down | 1,70 | 1,16E-02 |
| ENSG00000104886 | RBM38 | 55544 | RNA binding motif protein 38 | down | 1,69 | 1,30E-03 |
| ENSG00000122490 | COLGALT1 | 79709 | collagen beta(1-O)galactosyltransferase 1 | down | 1,69 | 2,64E-02 |
| ENSG00000125648 | THAP7-AS1 | 439931 | THAP7 antisense RNA 1 | down | 1,69 | 1,43E-02 |
| ENSG00000116685 | ACAT2 | 39 | acetyl-CoA acetyltransferase 2 | down | 1,69 | 2,53E-02 |
| ENSG00000069998 | DCTPP1 | 79077 | dCTP pyrophosphatase 1 | down | 1,69 | 1,60E-02 |
| ENSG00000116685 | VPS9D1-AS1 | 100128881 | VPS9D1 antisense RNA 1 | down | 1,69 | 1,57E-02 |
| ENSG00000177469 | IQCC | 55721 | IQ motif containing C | down | 1,69 | 9,76E-03 |
| ENSG00000119669 | MAFG-AS1 | 92659 | MAFG antisense RNA 1 (head to head) | down | 1,69 | 7,75E-03 |
| ENSG00000102870 | SRSF2 | 6427 | serine and arginine rich splicing factor 2 | down | 1,69 | 1,80E-02 |
| ENSG00000142208 | HILPDA | 29923 | hypoxia inducible lipid droplet associated | down | 1,68 | 5,68E-03 |
| ENSG00000158555 | HTR7P1 | 93164 | 5-hydroxytryptamine receptor 7 pseudogene 1 | down | 1,68 | 3,32E-03 |
| ENSG00000101191 | SDC3 | 9672 | syndecan 3 | down | 1,68 | 1,98E-02 |
| ENSG00000112146 | TLCD1 | 116238 | TLC domain containing 1 | down | 1,68 | 1,27E-02 |
| ENSG00000141560 | C11orf95 | 65998 | chromosome 11 open reading frame 95 | down | 1,68 | 2,21E-02 |
| ENSG00000111328 | MAP3K6 | 9064 | mitogen-activated protein kinase kinase kinase 6 | down | 1,68 | 3,58E-02 |
| ENSG00000176087 | SLC25A19 | 60386 | solute carrier family 25 member 19 | down | 1,67 | 6,88E-03 |
| ENSG00000247596 | IKBKE | 9641 | inhibitor of nuclear factor kappa B kinase subunit epsilon | down | 1,67 | 1,00E-02 |
| ENSG00000188130 | ZBTB42 | 100128927 | zinc finger and BTB domain containing 42 | down | 1,67 | 2,71E-03 |
| ENSG00000159788 | MT1A | 4489 | metallothionein 1A | down | 1,67 | 4,76E-02 |
| ENSG00000150977 | MT2A | 4502 | metallothionein 2A | down | 1,67 | 7,14E-03 |
| ENSG00000161091 | TMEM177 | 80775 | transmembrane protein 177 | down | 1,67 | 8,32E-03 |
| ENSG00000163348 | MCM7 | 4176 | minichromosome maintenance complex component 7 | down | 1,67 | 3,36E-02 |
| ENSG00000158234 | FAM109A | 144717 | family with sequence similarity 109 member A | down | 1,67 | 1,00E-02 |
| ENSG00000068024 | ZNF324B | 388569 | zinc finger protein 324B | down | 1,67 | 2,93E-02 |
| ENSG00000121057 | ZBTB47 | 92999 | zinc finger and BTB domain containing 47 | down | 1,67 | 3,74E-02 |
| ENSG00000108819 | SLC46A1 | 113235 | solute carrier family 46 member 1 | down | 1,66 | 4,59E-02 |
| ENSG00000160703 | EPHB2 | 2048 | EPH receptor B2 | down | 1,66 | 2,53E-02 |
| ENSG00000138092 | PPIF | 10105 | peptidylprolyl isomerase F | down | 1,66 | 7,17E-03 |
| ENSG00000174307 | ZMYM3 | 9203 | zinc finger MYM-type containing 3 | down | 1,66 | 3,73E-02 |
| ENSG00000099385 | ST3GAL2 | 6483 | ST3 beta-galactoside alpha-2,3-sialyltransferase 2 | down | 1,66 | 3,85E-03 |
| ENSG00000167280 | SPATA33 | 124045 | spermatogenesis associated 33 | down | 1,66 | 6,88E-03 |
| ENSG00000182810 | UBTF | 7343 | upstream binding transcription factor, RNA polymerase I | down | 1,66 | 1,21E-02 |
| ENSG00000184209 | C1QTNF1 | 114897 | C1q and TNF related 1 | down | 1,66 | 4,93E-02 |
| ENSG00000108639 | CNNM3 | 26505 | cyclin and CBS domain divalent metal cation transport mediator 3 | down | 1,66 | 3,37E-03 |
| ENSG00000168569 | HMGB3 | 3149 | high mobility group box 3 | down | 1,66 | 2,44E-02 |
| ENSG00000168806 | LZTS2 | 84445 | leucine zipper tumor suppressor 2 | down | 1,66 | 6,92E-03 |
| ENSG00000176953 | SOGA1 | 140710 | suppressor of glucose, autophagy associated 1 | down | 1,66 | 2,85E-02 |
| ENSG00000228544 | TGM2 | 7052 | transglutaminase 2 | down | 1,65 | 4,56E-02 |
| ENSG00000234616 | ZBTB12 | 221527 | zinc finger and BTB domain containing 12 | down | 1,65 | 7,17E-03 |
| ENSG00000133612 | FLVCR2 | 55640 | feline leukemia virus subgroup C cellular receptor family member 2 | down | 1,65 | 4,27E-02 |
| ENSG00000065054 | DAG1 | 1605 | dystroglycan 1 | down | 1,65 | 1,05E-02 |
| ENSG00000157881 | MTURN | 222166 | maturin, neural progenitor differentiation regulator homolog | down | 1,65 | 4,21E-02 |
| ENSG00000100294 | ALKBH4 | 54784 | alkB homolog 4, lysine demethylase | down | 1,65 | 1,94E-03 |
| ENSG00000114993 | LIPG | 9388 | lipase G, endothelial type | down | 1,65 | 1,50E-03 |
| ENSG00000186111 | MAMSTR | 284358 | MEF2 activating motif and SAP domain containing transcriptional regulator | down | 1,65 | 4,13E-02 |
| ENSG00000100023 | POP1 | 10940 | POP1 homolog, ribonuclease P/MRP subunit | down | 1,65 | 3,99E-02 |
| ENSG00000108773 | ABCB10 | 23456 | ATP binding cassette subfamily B member 10 | down | 1,65 | 4,38E-02 |
| ENSG00000143315 | GMPR | 2766 | guanosine monophosphate reductase | down | 1,65 | 3,57E-02 |
| ENSG00000243449 | TSHZ1 | 10194 | teashirt zinc finger homeobox 1 | down | 1,65 | 3,58E-02 |
| ENSG00000135763 | FAM57A | 79850 | family with sequence similarity 57 member A | down | 1,64 | 9,66E-03 |
| ENSG00000165724 | SH3BP1 | 23616 | SH3 domain binding protein 1 | down | 1,64 | 4,67E-02 |
| ENSG00000160953 | E2F1 | 1869 | E2F transcription factor 1 | down | 1,64 | 3,25E-02 |
| ENSG00000204237 | GAREM2 | 150946 | GRB2 associated regulator of MAPK1 subtype 2 | down | 1,64 | 7,72E-03 |
| ENSG00000136699 | SIX5 | 147912 | SIX homeobox 5 | down | 1,64 | 1,12E-03 |
| ENSG00000142252 | HSD3B7 | 80270 | hydroxy-delta-5-steroid dehydrogenase, 3 beta- and steroid delta-isomerase | down | 1,64 | 3,73E-02 |
| ENSG00000162144 | ZNF385A | 25946 | zinc finger protein 385A | down | 1,64 | 3,03E-02 |
| ENSG00000275023 | MARCH9 | 92979 | membrane associated ring-CH-type finger 9 | down | 1,64 | 3,10E-03 |
| ENSG00000278540 | CAMKK2 | 10645 | calcium/calmodulin dependent protein kinase kinase 2 | down | 1,63 | 2,19E-02 |
| ENSG00000134461 | ICMT | 23463 | isoprenylcysteine carboxyl methyltransferase | down | 1,63 | 2,40E-02 |
| ENSG00000101361 | MARCKSL1 | 65108 | MARCKS like 1 | down | 1,63 | 1,62E-02 |
| ENSG00000132205 | SAMD10 | 140700 | sterile alpha motif domain containing 10 | down | 1,63 | 1,61E-02 |
| ENSG00000103707 | ZNF180 | 7733 | zinc finger protein 180 | down | 1,63 | 1,85E-02 |
| ENSG00000172922 | HOXA10 | 3206 | homeobox A10 | down | 1,63 | 1,87E-02 |
| ENSG00000172409 | TUBA4A | 7277 | tubulin alpha 4a | down | 1,63 | 2,98E-02 |
| ENSG00000130520 | SLC35E4 | 339665 | solute carrier family 35 member E4 | down | 1,63 | 4,12E-03 |
| ENSG00000167775 | PACSIN3 | 29763 | protein kinase C and casein kinase substrate in neurons 3 | down | 1,63 | 3,59E-03 |
| ENSG00000015475 | NAGS | 162417 | N-acetylglutamate synthase | down | 1,62 | 4,90E-02 |
| ENSG00000060491 | SAMHD1 | 25939 | SAM and HD domain containing deoxynucleoside triphosphate triphosphohy | down | 1,62 | 4,03E-03 |
| ENSG00000141873 | CHAF1A | 10036 | chromatin assembly factor 1 subunit A | down | 1,62 | 3,80E-02 |
| ENSG00000144647 | NOTCH1 | 4851 | notch 1 | down | 1,62 | 2,47E-02 |
| ENSG00000197912 | DAB2IP | 153090 | DAB2 interacting protein | down | 1,62 | 1,58E-02 |
| ENSG00000057252 | SAMD14 | 201191 | sterile alpha motif domain containing 14 | down | 1,62 | 3,18E-02 |
| ENSG00000099364 | PGAM1 | 5223 | phosphoglycerate mutase 1 | down | 1,62 | 5,68E-03 |
| ENSG00000160803 | ZNRF3 | 84133 | zinc and ring finger 3 | down | 1,62 | 2,51E-02 |
| ENSG00000020256 | ATP2A3 | 489 | ATPase sarcoplasmic/endoplasmic reticulum Ca2+ transporting 3 | down | 1,62 | 4,34E-02 |
| ENSG00000125841 | CASTOR2 | 729438 | cytosolic arginine sensor for mTORC1 subunit 2 | down | 1,61 | 1,95E-02 |
| ENSG00000131323 | DDX11 | 1663 | DEAD/H-box helicase 11 | down | 1,61 | 1,79E-02 |
| ENSG00000131473 | ATP6V0E2 | 155066 | ATPase H+ transporting V0 subunit e2 | down | 1,61 | 2,71E-03 |
| ENSG00000158716 | AHDC1 | 27245 | AT-hook DNA binding motif containing 1 | down | 1,61 | 1,95E-02 |
| ENSG00000125170 | SLC29A1 | 2030 | solute carrier family 29 member 1 (Augustine blood group) | down | 1,61 | 8,47E-03 |
| ENSG00000198728 | C1orf159 | 54991 | chromosome 1 open reading frame 159 | down | 1,61 | 3,67E-03 |
| ENSG00000273820 | ANO7 | 50636 | anoctamin 7 | down | 1,61 | 5,95E-03 |
| ENSG00000197451 | RGS14 | 10636 | regulator of G protein signaling 14 | down | 1,61 | 4,73E-02 |
| ENSG00000124160 | PATZ1 | 23598 | POZ/BTB and AT hook containing zinc finger 1 | down | 1,60 | 7,17E-03 |
| ENSG00000214063 | CCDC134 | 79879 | coiled-coil domain containing 134 | down | 1,60 | 4,70E-03 |
| ENSG00000115255 | FJX1 | 24147 | four jointed box 1 | down | 1,60 | 2,96E-02 |
| ENSG00000132383 | CCND3 | 896 | cyclin D3 | down | 1,60 | 4,18E-02 |
| ENSG00000160932 | TRIM65 | 201292 | tripartite motif containing 65 | down | 1,59 | 2,18E-02 |
| ENSG00000184787 | POLD1 | 5424 | DNA polymerase delta 1, catalytic subunit | down | 1,59 | 4,63E-02 |
| ENSG00000186654 | NDST1 | 3340 | N-deacetylase and N-sulfotransferase 1 | down | 1,59 | 2,40E-03 |
| ENSG00000100142 | LRFN4 | 78999 | leucine rich repeat and fibronectin type III domain containing 4 | down | 1,59 | 2,11E-03 |
| ENSG00000275464 | DBP | 1628 | D-box binding PAR bZIP transcription factor | down | 1,59 | 2,85E-02 |
| ENSG00000173457 | SLC27A4 | 10999 | solute carrier family 27 member 4 | down | 1,59 | 7,26E-03 |
| ENSG00000107882 | ZNF250 | 58500 | zinc finger protein 250 | down | 1,59 | 2,29E-02 |
| ENSG00000185024 | TSEN54 | 283989 | tRNA splicing endonuclease subunit 54 | down | 1,59 | 9,52E-03 |
| ENSG00000007376 | SNX21 | 90203 | sorting nexin family member 21 | down | 1,59 | 2,24E-03 |
| ENSG00000072163 | SLC16A13 | 201232 | solute carrier family 16 member 13 | down | 1,58 | 1,40E-02 |
| ENSG00000164086 | COA3 | 28958 | cytochrome c oxidase assembly factor 3 | down | 1,58 | 3,24E-02 |
| ENSG00000168802 | LOC171391 | 171391 | uncharacterized LOC171391 | down | 1,58 | 1,14E-02 |
| ENSG00000259479 | ZBTB14 | 7541 | zinc finger and BTB domain containing 14 | down | 1,58 | 1,93E-02 |
| ENSG00000182108 | A4GALT | 53947 | alpha 1,4-galactosyltransferase (P blood group) | down | 1,58 | 1,61E-02 |
| ENSG00000005882 | CCM2 | 83605 | CCM2 scaffolding protein | down | 1,58 | 6,37E-03 |
| ENSG00000072310 | KIF3C | 3797 | kinesin family member 3C | down | 1,58 | 4,57E-02 |
| ENSG00000110583 | CCDC106 | 29903 | coiled-coil domain containing 106 | down | 1,58 | 8,11E-04 |
| ENSG00000141562 | FBXO5 | 26271 | F-box protein 5 | down | 1,57 | 4,57E-02 |
| ENSG00000243364 | DISP2 | 85455 | dispatched RND transporter family member 2 | down | 1,57 | 3,64E-02 |
| ENSG00000139405 | VASH1-AS1 | 100506603 | VASH1 antisense RNA 1 | down | 1,57 | 3,50E-02 |
| ENSG00000172458 | SLC35B4 | 84912 | solute carrier family 35 member B4 | down | 1,57 | 3,99E-02 |
| ENSG00000240889 | HGH1 | 51236 | HGH1 homolog | down | 1,57 | 8,47E-03 |
| ENSG00000198668 | CD3EAP | 10849 | CD3e molecule associated protein | down | 1,57 | 3,13E-03 |
| ENSG00000198668 | IMP3 | 55272 | IMP3, U3 small nucleolar ribonucleoprotein | down | 1,57 | 1,42E-02 |
| ENSG00000198668 | ZNF174 | 7727 | zinc finger protein 174 | down | 1,57 | 1,01E-02 |
| ENSG00000182584 | AK1 | 203 | adenylate kinase 1 | down | 1,57 | 9,91E-03 |
| ENSG00000084774 | SLC35B2 | 347734 | solute carrier family 35 member B2 | down | 1,57 | 6,39E-03 |
| ENSG00000123810 | FUT10 | 84750 | fucosyltransferase 10 | down | 1,56 | 2,88E-02 |
| ENSG00000160446 | SNN | 8303 | stannin | down | 1,56 | 5,45E-03 |
| ENSG00000162639 | TIMM21 | 29090 | translocase of inner mitochondrial membrane 21 | down | 1,56 | 1,69E-02 |
| ENSG00000196371 | HES6 | 55502 | hes family bHLH transcription factor 6 | down | 1,56 | 1,92E-02 |
| ENSG00000003249 | TUBB | 203068 | tubulin beta class I | down | 1,56 | 2,67E-02 |
| ENSG00000139531 | UQCC3 | 790955 | ubiquinol-cytochrome c reductase complex assembly factor 3 | down | 1,56 | 5,91E-03 |
| ENSG00000168778 | NAT6 | 24142 | N-acetyltransferase 6 | down | 1,56 | 2,71E-03 |
| ENSG00000171017 | PELI3 | 246330 | pellino E3 ubiquitin protein ligase family member 3 | down | 1,56 | 1,45E-02 |
| ENSG00000069011 | LOC105374836 | 105374836 | uncharacterized LOC105374836 | down | 1,56 | 2,11E-02 |
| ENSG00000132600 | CEP19 | 84984 | centrosomal protein 19 | down | 1,56 | 8,75E-03 |
| ENSG00000106683 | TRABD2A | 129293 | TraB domain containing 2A | down | 1,56 | 2,11E-02 |
| ENSG00000111642 | IMPDH1 | 3614 | inosine monophosphate dehydrogenase 1 | down | 1,56 | 1,50E-03 |
| ENSG00000144711 | SLC25A22 | 79751 | solute carrier family 25 member 22 | down | 1,55 | 4,36E-02 |
| ENSG00000167840 | CCDC51 | 79714 | coiled-coil domain containing 51 | down | 1,55 | 2,45E-02 |
| ENSG00000122687 | HOXD9 | 3235 | homeobox D9 | down | 1,55 | 4,57E-02 |
| ENSG00000127399 | EFHD2 | 79180 | EF-hand domain family member D2 | down | 1,55 | 7,65E-03 |
| ENSG00000204315 | ARHGEF19 | 128272 | Rho guanine nucleotide exchange factor 19 | down | 1,55 | 1,82E-02 |
| ENSG00000070413 | DHRS4-AS1 | 55449 | DHRS4 antisense RNA 1 | down | 1,55 | 3,95E-02 |
| ENSG00000127838 | MYD88 | 4615 | myeloid differentiation primary response 88 | down | 1,55 | 1,86E-02 |
| ENSG00000168411 | PHF13 | 148479 | PHD finger protein 13 | down | 1,55 | 4,04E-02 |
| ENSG00000173548 | SUV39H1 | 6839 | suppressor of variegation 3-9 homolog 1 | down | 1,55 | 1,63E-02 |
| ENSG00000125817 | CALM2 | 805 | calmodulin 2 | down | 1,55 | 8,32E-03 |
| ENSG00000126351 | CALM3 | 808 | calmodulin 3 | down | 1,55 | 8,32E-03 |
| ENSG00000132589 | POLE | 5426 | DNA polymerase epsilon, catalytic subunit | down | 1,55 | 2,68E-02 |
| ENSG00000138031 | FADS1 | 3992 | fatty acid desaturase 1 | down | 1,54 | 8,39E-03 |
| ENSG00000143179 | ACSL1 | 2180 | acyl-CoA synthetase long chain family member 1 | down | 1,54 | 3,01E-02 |
| ENSG00000048140 | ANKS6 | 203286 | ankyrin repeat and sterile alpha motif domain containing 6 | down | 1,54 | 3,37E-03 |
| ENSG00000160753 | CISD3 | 284106 | CDGSH iron sulfur domain 3 | down | 1,54 | 1,51E-03 |
| ENSG00000073060 | WDR92 | 116143 | WD repeat domain 92 | down | 1,54 | 4,86E-02 |
| ENSG00000101811 | ISYNA1 | 51477 | inositol-3-phosphate synthase 1 | down | 1,54 | 7,65E-03 |
| ENSG00000170846 | CENPV | 201161 | centromere protein V | down | 1,54 | 1,40E-02 |
| ENSG00000204934 | ACTR5 | 79913 | ARP5 actin related protein 5 homolog | down | 1,54 | 9,20E-03 |
| ENSG00000010165 | SPR | 6697 | sepiapterin reductase | down | 1,54 | 1,91E-02 |
| ENSG00000171700 | NLGN2 | 57555 | neuroligin 2 | down | 1,54 | 2,37E-02 |
| ENSG00000182841 | P2RY11 | 5032 | purinergic receptor P2Y11 | down | 1,54 | 2,90E-02 |

| | | | | | | |
|---|---|---|---|---|---|---|
| ENSG00000184990 | TRMU | 55687 | tRNA 5-methylaminomethyl-2-thiouridylate methyltransferase | down | 1,54 | 1,57E-02 |
| ENSG00000117411 | SNRNP25 | 79622 | small nuclear ribonucleoprotein U11/U12 subunit 25 | down | 1,54 | 4,80E-02 |
| ENSG00000177613 | F2R | 2149 | coagulation factor II thrombin receptor | down | 1,54 | 7,69E-03 |
| ENSG00000157978 | TIGD5 | 84948 | tigger transposable element derived 5 | down | 1,53 | 1,21E-02 |
| ENSG00000187626 | CDHR3 | 222256 | cadherin related family member 3 | down | 1,53 | 4,57E-02 |
| ENSG00000017483 | DOK1 | 1796 | docking protein 1 | down | 1,53 | 2,54E-03 |
| ENSG00000182544 | MSRB1 | 51734 | methionine sulfoxide reductase B1 | down | 1,53 | 2,24E-03 |
| ENSG00000101220 | PIP4K2B | 8396 | phosphatidylinositol-5-phosphate 4-kinase type 2 beta | down | 1,53 | 1,02E-02 |
| ENSG00000177096 | SH2D4A | 63898 | SH2 domain containing 4A | down | 1,53 | 4,42E-02 |
| ENSG00000167716 | NFIC | 4782 | nuclear factor I C | down | 1,53 | 1,49E-02 |
| ENSG00000185551 | ZC3HAV1 | 56829 | zinc finger CCCH-type containing, antiviral 1 | down | 1,53 | 4,26E-02 |
| ENSG00000160877 | KDM2B | 84678 | lysine demethylase 2B | down | 1,53 | 2,57E-02 |
| ENSG00000261801 | FAM173B | 134145 | family with sequence similarity 173 member B | down | 1,53 | 4,89E-02 |
| ENSG00000102981 | PFAS | 5198 | phosphoribosylformylglycinamidine synthase | down | 1,53 | 1,70E-02 |
| ENSG00000160094 | BOLA1 | 51027 | bolA family member 1 | down | 1,53 | 1,23E-02 |
| ENSG00000160404 | PIGW | 284098 | phosphatidylinositol glycan anchor biosynthesis class W | down | 1,53 | 1,79E-02 |
| ENSG00000143995 | NEK10 | 152110 | NIMA related kinase 10 | down | 1,52 | 4,97E-02 |
| ENSG00000152518 | F12 | 2161 | coagulation factor XII | down | 1,52 | 3,17E-02 |
| ENSG00000158715 | BRD3OS | 266655 | BRD3 opposite strand | down | 1,52 | 9,20E-03 |
| ENSG00000170468 | RNF26 | 79102 | ring finger protein 26 | down | 1,52 | 1,31E-02 |
| ENSG00000081059 | GYG2 | 8908 | glycogenin 2 | down | 1,52 | 4,53E-02 |
| ENSG00000126464 | MBD3 | 53615 | methyl-CpG binding domain protein 3 | down | 1,52 | 4,26E-04 |
| ENSG00000183337 | PODXL2 | 50512 | podocalyxin like 2 | down | 1,52 | 1,81E-02 |
| ENSG00000125505 | SIGMAR1 | 10280 | sigma non-opioid intracellular receptor 1 | down | 1,52 | 1,79E-02 |
| ENSG00000127191 | POLR2L | 5441 | RNA polymerase II subunit L | down | 1,52 | 3,59E-03 |
| ENSG00000141736 | NCLN | 56926 | nicalin | down | 1,52 | 1,12E-03 |
| ENSG00000272142 | DTX4 | 23220 | deltex E3 ubiquitin ligase 4 | down | 1,52 | 4,63E-02 |
| ENSG00000103202 | WRAP53 | 55135 | WD repeat containing antisense to TP53 | down | 1,52 | 1,70E-02 |
| ENSG00000176720 | RTN4RL2 | 349667 | reticulon 4 receptor like 2 | down | 1,52 | 1,13E-02 |
| ENSG00000187713 | NUP62 | 23636 | nucleoporin 62 | down | 1,52 | 4,40E-03 |
| ENSG00000100304 | VOPP1 | 81552 | VOPP1, WBP1/VOPP1 family member | down | 1,52 | 4,69E-02 |
| ENSG00000103174 | GPR39 | 2863 | G protein-coupled receptor 39 | down | 1,52 | 4,50E-03 |
| ENSG00000135821 | AGAP2-AS1 | 100130776 | AGAP2 antisense RNA 1 | down | 1,52 | 5,16E-03 |
| ENSG00000100726 | CDC42EP4 | 23580 | CDC42 effector protein 4 | down | 1,52 | 1,16E-02 |
| ENSG00000143815 | PPT2 | 9374 | palmitoyl-protein thioesterase 2 | down | 1,51 | 1,22E-02 |
| ENSG00000148411 | CPNE2 | 221184 | copine 2 | down | 1,51 | 1,49E-02 |
| ENSG00000165879 | C1orf74 | 148304 | chromosome 1 open reading frame 74 | down | 1,51 | 3,06E-02 |
| ENSG00000106009 | PCDHGC3 | 5098 | protocadherin gamma subfamily C, 3 | down | 1,51 | 3,99E-02 |
| ENSG00000110400 | CHST12 | 55501 | carbohydrate sulfotransferase 12 | down | 1,51 | 1,31E-04 |
| ENSG00000189339 | EXOSC4 | 54512 | exosome component 4 | down | 1,51 | 2,77E-03 |
| ENSG00000183779 | PLCG1 | 5335 | phospholipase C gamma 1 | down | 1,51 | 2,86E-02 |
| ENSG00000162065 | IGFBP4 | 3487 | insulin like growth factor binding protein 4 | down | 1,51 | 1,12E-02 |
| ENSG00000185298 | RCC1 | 1104 | regulator of chromosome condensation 1 | down | 1,51 | 6,96E-03 |
| ENSG00000160113 | EXOC3-AS1 | 116349 | EXOC3 antisense RNA 1 | down | 1,51 | 3,10E-03 |
| ENSG00000141295 | KAZALD1 | 81621 | Kazal type serine peptidase inhibitor domain 1 | down | 1,51 | 4,39E-02 |
| ENSG00000130299 | RBM14 | 10432 | RNA binding motif protein 14 | down | 1,51 | 1,38E-02 |
| ENSG00000100075 | MRPS34 | 65993 | mitochondrial ribosomal protein S34 | down | 1,51 | 1,07E-02 |
| ENSG00000100401 | LOXL1 | 4016 | lysyl oxidase like 1 | down | 1,51 | 3,17E-02 |
| ENSG00000197119 | TM7SF2 | 7108 | transmembrane 7 superfamily member 2 | down | 1,51 | 4,01E-02 |
| ENSG00000278619 | PARP1 | 142 | poly(ADP-ribose) polymerase 1 | down | 1,50 | 3,92E-02 |
| ENSG00000011009 | EXOSC6 | 118460 | exosome component 6 | down | 1,50 | 1,04E-02 |
| ENSG00000103064 | ZNF689 | 115509 | zinc finger protein 689 | down | 1,50 | 9,77E-03 |
| ENSG00000125901 | TUBG1 | 7283 | tubulin gamma 1 | down | 1,50 | 1,42E-02 |
| ENSG00000158156 | C1orf216 | 127703 | chromosome 1 open reading frame 216 | down | 1,50 | 2,42E-02 |
| ENSG00000110171 | PGP | 283871 | phosphoglycolate phosphatase | down | 1,50 | 8,04E-03 |
| ENSG00000132361 | CENPX | 201254 | centromere protein X | down | 1,50 | 2,53E-02 |
| ENSG00000166189 | PRKAR1B | 5575 | protein kinase cAMP-dependent type I regulatory subunit beta | down | 1,50 | 6,80E-04 |
| ENSG00000173786 | LY6G5C | 80741 | lymphocyte antigen 6 family member G5C | down | 1,50 | 4,00E-02 |
| ENSG00000133895 | LSS | 4047 | lanosterol synthase | down | 1,49 | 1,66E-03 |
| ENSG00000182796 | ABHD6 | 57406 | abhydrolase domain containing 6 | down | 1,49 | 2,16E-02 |
| ENSG00000123933 | MMP15 | 4324 | matrix metallopeptidase 15 | down | 1,49 | 4,36E-02 |
| ENSG00000124702 | TYSND1 | 219743 | trypsin domain containing 1 | down | 1,49 | 3,10E-02 |
| ENSG00000157557 | MXD4 | 10608 | MAX dimerization protein 4 | down | 1,49 | 4,76E-03 |
| ENSG00000160285 | KLHDC3 | 116138 | kelch domain containing 3 | down | 1,49 | 2,19E-03 |
| ENSG00000163686 | ETS2 | 2114 | ETS proto-oncogene 2, transcription factor | down | 1,49 | 4,28E-02 |
| ENSG00000102996 | MEN1 | 4221 | menin 1 | down | 1,49 | 4,75E-04 |
| ENSG00000156521 | TMEM198B | 440104 | transmembrane protein 198B (pseudogene) | down | 1,49 | 3,20E-02 |
| ENSG00000184207 | SLC7A6 | 9057 | solute carrier family 7 member 6 | down | 1,49 | 1,38E-02 |
| ENSG00000169689 | MRPS26 | 64949 | mitochondrial ribosomal protein S26 | down | 1,49 | 2,65E-02 |
| ENSG00000188191 | XKR8 | 55113 | XK related 8 | down | 1,49 | 4,03E-03 |
| ENSG00000204428 | TRIM3 | 10612 | tripartite motif containing 3 | down | 1,49 | 2,77E-02 |
| ENSG00000131462 | CLUH | 23277 | clustered mitochondria homolog | down | 1,49 | 7,72E-03 |
| ENSG00000142686 | HPS6 | 79803 | HPS6, biogenesis of lysosomal organelles complex 2 subunit 3 | down | 1,49 | 3,48E-03 |
| ENSG00000223496 | CNP | 1267 | 2',3'-cyclic nucleotide 3' phosphodiesterase | down | 1,49 | 9,41E-03 |
| ENSG00000156853 | LYPLA2 | 11313 | lysophospholipase II | down | 1,49 | 2,46E-02 |
| ENSG00000143799 | GTPBP3 | 84705 | GTP binding protein 3, mitochondrial | down | 1,48 | 4,01E-02 |
| ENSG00000149809 | SLC25A1 | 6576 | solute carrier family 25 member 1 | down | 1,48 | 5,55E-04 |
| ENSG00000180198 | RANGAP1 | 5905 | Ran GTPase activating protein 1 | down | 1,48 | 1,41E-02 |
| ENSG00000221990 | SLC25A29 | 123096 | solute carrier family 25 member 29 | down | 1,48 | 2,78E-03 |
| ENSG00000107821 | MRM1 | 79922 | mitochondrial rRNA methyltransferase 1 | down | 1,48 | 2,19E-02 |
| ENSG00000239306 | TBC1D24 | 57465 | TBC1 domain family member 24 | down | 1,48 | 3,58E-02 |
| ENSG00000074071 | CCDC137 | 339230 | coiled-coil domain containing 137 | down | 1,48 | 1,00E-02 |
| ENSG00000129038 | NR2F6 | 2063 | nuclear receptor subfamily 2 group F member 6 | down | 1,48 | 1,28E-02 |
| ENSG00000124181 | SCRN2 | 90507 | secernin 2 | down | 1,48 | 2,66E-02 |
| ENSG00000141753 | BRAT1 | 221927 | BRCA1 associated ATM activator 1 | down | 1,48 | 1,14E-02 |
| ENSG00000221988 | NECTIN1 | 5818 | nectin cell adhesion molecule 1 | down | 1,48 | 7,72E-03 |
| ENSG00000140848 | SLC35E2B | 728661 | solute carrier family 35 member E2B | down | 1,48 | 4,40E-03 |
| ENSG00000162757 | ZNF703 | 80139 | zinc finger protein 703 | down | 1,48 | 4,67E-02 |
| ENSG00000240184 | LBR | 3930 | lamin B receptor | down | 1,48 | 4,92E-02 |
| ENSG00000136213 | NACC2 | 138151 | NACC family member 2 | down | 1,48 | 2,64E-03 |
| ENSG00000178896 | FRAT1 | 10023 | FRAT1, WNT signaling pathway regulator | down | 1,48 | 3,15E-02 |
| ENSG00000179604 | TTLL12 | 23170 | tubulin tyrosine ligase like 12 | down | 1,47 | 1,81E-02 |
| ENSG00000110042 | NAGPA | 51172 | N-acetylglucosamine-1-phosphodiester alpha-N-acetylglucosaminidase | down | 1,47 | 1,63E-03 |
| ENSG00000141499 | GLUL | 2752 | glutamate-ammonia ligase | down | 1,47 | 6,92E-03 |
| ENSG00000186907 | TELO2 | 9894 | telomere maintenance 2 | down | 1,47 | 2,97E-02 |
| ENSG00000213024 | NME4 | 4833 | NME/NM23 nucleoside diphosphate kinase 4 | down | 1,47 | 3,86E-02 |
| ENSG00000154978 | BOK | 666 | BOK, BCL2 family apoptosis regulator | down | 1,47 | 4,10E-04 |
| ENSG00000183840 | TMEM203 | 94107 | transmembrane protein 203 | down | 1,47 | 1,01E-02 |
| ENSG00000255737 | TRAF2 | 7186 | TNF receptor associated factor 2 | down | 1,47 | 2,66E-02 |
| ENSG00000114631 | ERBB2 | 2064 | erb-b2 receptor tyrosine kinase 2 | down | 1,47 | 1,94E-02 |
| ENSG00000147955 | LYRM4-AS1 | 100129461 | LYRM4 antisense RNA 1 | down | 1,47 | 4,12E-02 |
| ENSG00000177700 | TCF7 | 6932 | transcription factor 7 | down | 1,47 | 1,07E-02 |
| ENSG00000173456 | PRR12 | 57479 | proline rich 12 | down | 1,47 | 3,48E-03 |
| ENSG00000125912 | BCOR | 54880 | BCL6 corepressor | down | 1,47 | 2,58E-02 |
| ENSG00000056998 | MBOAT7 | 79143 | membrane bound O-acyltransferase domain containing 7 | down | 1,47 | 9,37E-03 |
| ENSG00000071655 | TOR2A | 27433 | torsin family 2 member A | down | 1,47 | 1,55E-02 |
| ENSG00000163491 | MEIS1 | 4211 | Meis homeobox 1 | down | 1,47 | 3,50E-02 |
| ENSG00000131187 | ZFP36L2 | 678 | ZFP36 ring finger protein like 2 | down | 1,47 | 2,47E-02 |
| ENSG00000235106 | SLC45A3 | 85414 | solute carrier family 45 member 3 | down | 1,47 | 1,40E-02 |
| ENSG00000178096 | RIOX1 | 79697 | ribosomal oxygenase 1 | down | 1,47 | 6,88E-03 |
| ENSG00000277161 | PARD6A | 50855 | par-6 family cell polarity regulator alpha | down | 1,46 | 8,78E-03 |
| ENSG00000089094 | ZNF362 | 149076 | zinc finger protein 362 | down | 1,46 | 4,50E-02 |
| ENSG00000150756 | NR2F2 | 7026 | nuclear receptor subfamily 2 group F member 2 | down | 1,46 | 8,32E-03 |
| ENSG00000178921 | NACC1 | 112939 | nucleus accumbens associated 1 | down | 1,46 | 1,78E-04 |
| ENSG00000104611 | LOXL1-AS1 | 100287616 | LOXL1 antisense RNA 1 | down | 1,46 | 3,57E-02 |
| ENSG00000141905 | FAM109B | 150368 | family with sequence similarity 109 member B | down | 1,46 | 2,20E-02 |
| ENSG00000105939 | WDR81 | 124997 | WD repeat domain 81 | down | 1,46 | 2,32E-03 |
| ENSG00000179886 | LDLRAP1 | 26119 | low density lipoprotein receptor adaptor protein 1 | down | 1,46 | 2,21E-02 |
| ENSG00000128536 | ZKSCAN4 | 387032 | zinc finger with KRAB and SCAN domains 4 | down | 1,46 | 3,86E-02 |
| ENSG00000115325 | SLC38A5 | 92745 | solute carrier family 38 member 5 | down | 1,46 | 4,03E-02 |
| ENSG00000198736 | MFSD5 | 84975 | major facilitator superfamily domain containing 5 | down | 1,46 | 2,58E-02 |
| ENSG00000276293 | C20orf27 | 54976 | chromosome 20 open reading frame 27 | down | 1,46 | 5,70E-04 |
| ENSG00000244165 | METTL13 | 51603 | methyltransferase like 13 | down | 1,45 | 1,12E-03 |
| ENSG00000100416 | RGS19 | 10287 | regulator of G protein signaling 19 | down | 1,45 | 1,44E-03 |
| ENSG00000161981 | RRP7BP | 91695 | ribosomal RNA processing 7 homolog B, pseudogene | down | 1,45 | 1,68E-02 |
| ENSG00000181104 | SIVA1 | 10572 | SIVA1 apoptosis inducing factor | down | 1,45 | 1,26E-02 |
| ENSG00000101442 | B4GALT2 | 8704 | beta-1,4-galactosyltransferase 2 | down | 1,45 | 2,14E-03 |
| ENSG00000116096 | CSTF2T | 23283 | cleavage stimulation factor subunit 2 tau variant | down | 1,45 | 1,70E-02 |
| ENSG00000169992 | ATP6V0E2-AS1 | 401431 | ATP6V0E2 antisense RNA 1 | down | 1,45 | 3,38E-02 |
| ENSG00000243667 | TSPAN17 | 26262 | tetraspanin 17 | down | 1,45 | 2,69E-03 |
| ENSG00000105655 | RUSC1 | 23623 | RUN and SH3 domain containing 1 | down | 1,45 | 8,90E-03 |
| ENSG00000166582 | SCARB1 | 949 | scavenger receptor class B member 1 | down | 1,45 | 4,27E-02 |
| ENSG00000149485 | CSTF2 | 1478 | cleavage stimulation factor subunit 2 | down | 1,45 | 4,39E-02 |
| ENSG00000151726 | LOC93622 | 93622 | Morf4 family associated protein 1 like 1 pseudogene | down | 1,45 | 4,35E-02 |
| ENSG00000165138 | THRA | 7067 | thyroid hormone receptor, alpha | down | 1,45 | 2,64E-02 |
| ENSG00000277972 | FLOT2 | 2319 | flotillin 2 | down | 1,45 | 3,40E-02 |
| ENSG00000116273 | ADCY3 | 109 | adenylate cyclase 3 | down | 1,45 | 1,04E-02 |
| ENSG00000101945 | UCK2 | 7371 | uridine-cytidine kinase 2 | down | 1,45 | 2,72E-02 |
| ENSG00000160014 | SNX33 | 257364 | sorting nexin 33 | down | 1,44 | 2,11E-02 |
| ENSG00000160014 | CENPB | 1059 | centromere protein B | down | 1,44 | 3,32E-03 |
| ENSG00000177084 | MRM2 | 29960 | mitochondrial rRNA methyltransferase 2 | down | 1,44 | 1,90E-02 |
| ENSG00000215256 | LRRC61 | 65999 | leucine rich repeat containing 61 | down | 1,44 | 3,37E-03 |
| ENSG00000172936 | FKBPL | 63943 | FK506 binding protein like | down | 1,44 | 1,67E-02 |
| ENSG00000128709 | DGCR2 | 9993 | DiGeorge syndrome critical region gene 2 | down | 1,44 | 3,13E-03 |
| ENSG00000142634 | PNKD | 25953 | paroxysmal nonkinesigenic dyskinesia | down | 1,44 | 2,59E-02 |
| ENSG00000142632 | RFWD3 | 55159 | ring finger and WD repeat domain 3 | down | 1,44 | 4,11E-02 |
| ENSG00000177542 | PITX1 | 5307 | paired like homeodomain 1 | down | 1,44 | 2,76E-02 |
| ENSG00000164051 | PRMT7 | 54496 | protein arginine methyltransferase 7 | down | 1,44 | 1,88E-02 |
| ENSG00000106348 | LIMK1 | 3984 | LIM domain kinase 1 | down | 1,44 | 6,50E-03 |
| ENSG00000186854 | CHD4 | 1108 | chromodomain helicase DNA binding protein 4 | down | 1,44 | 9,37E-03 |
| ENSG00000174007 | IQSEC1 | 9922 | IQ motif and Sec7 domain 1 | down | 1,44 | 3,57E-02 |
| ENSG00000186854 | ZNF232 | 7775 | zinc finger protein 232 | down | 1,44 | 3,58E-02 |
| ENSG00000174516 | FUT4 | 2526 | fucosyltransferase 4 | down | 1,44 | 1,70E-03 |
| ENSG00000075336 | DBNDD1 | 79007 | dysbindin domain containing 1 | down | 1,44 | 1,43E-02 |
| ENSG00000144485 | SUOX | 6821 | sulfite oxidase | down | 1,44 | 3,36E-02 |
| ENSG00000196230 | TCTN2 | 79867 | tectonic family member 2 | down | 1,44 | 2,64E-02 |
| ENSG00000204922 | LRRC8E | 80131 | leucine rich repeat containing 8 VRAC subunit E | down | 1,44 | 9,61E-03 |
| ENSG00000243477 | CAD | 790 | carbamoyl-phosphate synthetase 2, aspartate transcarbamylase, and dihyd | down | 1,43 | 7,14E-03 |
| ENSG00000172728 | B9D2 | 80776 | B9 domain containing 2 | down | 1,43 | 1,57E-02 |
| ENSG00000184602 | ZDHHC12 | 84885 | zinc finger DHHC-type containing 12 | down | 1,43 | 2,00E-03 |
| ENSG00000103343 | HENMT1 | 113802 | HEN methyltransferase 1 | down | 1,43 | 3,77E-02 |
| ENSG00000106992 | ACTL10 | 170487 | actin like 10 | down | 1,43 | 6,88E-03 |
| ENSG00000157593 | IL17D | 53342 | interleukin 17D | down | 1,43 | 8,17E-03 |
| ENSG00000117877 | NDUFB2-AS1 | 100134713 | NDUFB2 antisense RNA 1 | down | 1,43 | 1,60E-02 |
| ENSG00000177971 | CALM1 | 801 | calmodulin 1 | down | 1,43 | 2,24E-02 |
| ENSG00000235173 | CALM2 | 805 | calmodulin 2 | down | 1,43 | 2,24E-02 |
| ENSG00000112029 | CALM3 | 808 | calmodulin 3 | down | 1,43 | 2,24E-02 |
| ENSG00000140323 | RITA1 | 84934 | RBPJ interacting and tubulin associated 1 | down | 1,43 | 3,85E-03 |
| ENSG00000258301 | NAA40 | 79829 | N(alpha)-acetyltransferase 40, NatD catalytic subunit | down | 1,43 | 2,47E-02 |
| ENSG00000205060 | NARF | 26502 | nuclear prelamin A recognition factor | down | 1,43 | 8,90E-03 |
| ENSG00000136280 | EFNA4 | 1945 | ephrin A4 | down | 1,43 | 4,93E-02 |
| ENSG00000084731 | PDK2 | 5164 | pyruvate dehydrogenase kinase 2 | down | 1,42 | 1,03E-02 |
| ENSG00000173581 | SREBF1 | 6720 | sterol regulatory element binding transcription factor 1 | down | 1,42 | 3,68E-02 |
| ENSG00000198081 | DEXI | 28955 | Dexi homolog | down | 1,42 | 2,10E-03 |
| ENSG00000128274 | DUSP7 | 1849 | dual specificity phosphatase 7 | down | 1,42 | 1,79E-02 |
| ENSG00000174327 | CHTF8 | 54921 | chromosome transmission fidelity factor 8 | down | 1,42 | 1,16E-02 |
| ENSG00000183978 | SORD2P | 653381 | sorbitol dehydrogenase 2, pseudogene | down | 1,42 | 4,63E-02 |
| ENSG00000255284 | RPUSD1 | 113000 | RNA pseudouridylate synthase domain containing 1 | down | 1,42 | 4,13E-03 |
| ENSG00000124104 | LIMS2 | 55679 | LIM zinc finger domain containing 2 | down | 1,42 | 1,12E-02 |
| ENSG00000105516 | BRF1 | 2972 | BRF1, RNA polymerase III transcription initiation factor subunit | down | 1,42 | 3,92E-04 |
| ENSG00000167114 | POLR2F | 5435 | RNA polymerase II subunit F | down | 1,41 | 3,23E-02 |
| ENSG00000196150 | LOC102724159 | 102724159 | periodic tryptophan protein 2 homolog | down | 1,41 | 1,43E-02 |
| ENSG00000182173 | PPP1R14B | 26472 | protein phosphatase 1 regulatory inhibitor subunit 14B | down | 1,41 | 7,05E-04 |
| ENSG00000141569 | SUFU | 51684 | SUFU negative regulator of hedgehog signaling | down | 1,41 | 3,38E-02 |
| ENSG00000062822 | REEP6 | 92840 | receptor accessory protein 6 | down | 1,41 | 9,21E-03 |
| ENSG00000070614 | RPA1 | 6117 | replication protein A1 | down | 1,41 | 4,94E-02 |
| ENSG00000173621 | LY6E | 4061 | lymphocyte antigen 6 family member E | down | 1,41 | 4,10E-04 |
| ENSG00000179431 | UBE2G2 | 7327 | ubiquitin conjugating enzyme E2 G2 | down | 1,41 | 4,13E-03 |
| ENSG00000112576 | PRR5 | 55615 | proline rich 5 | down | 1,41 | 1,80E-03 |
| ENSG00000100105 | NCOA5 | 57727 | nuclear receptor coactivator 5 | down | 1,41 | 3,22E-03 |
| ENSG00000100147 | TSPAN4 | 7106 | tetraspanin 4 | down | 1,41 | 4,57E-02 |
| ENSG00000146205 | ACLY | 47 | ATP citrate lyase | down | 1,41 | 3,10E-02 |
| ENSG00000169220 | DUSP23 | 54935 | dual specificity phosphatase 23 | down | 1,41 | 3,49E-02 |
| ENSG00000126705 | DOK4 | 55715 | docking protein 4 | down | 1,41 | 2,06E-02 |
| ENSG00000112759 | LDB1 | 8861 | LIM domain binding 1 | down | 1,41 | 1,96E-02 |
| ENSG00000131591 | USP27X | 389856 | ubiquitin specific peptidase 27, X-linked | down | 1,41 | 3,52E-02 |
| ENSG00000274070 | HNRNPAB | 3182 | heterogeneous nuclear ribonucleoprotein A/B | down | 1,41 | 2,01E-02 |
| ENSG00000013573 | SOAT1 | 6646 | sterol O-acyltransferase 1 | down | 1,40 | 1,14E-02 |
| ENSG00000171130 | FBXL19 | 54620 | F-box and leucine rich repeat protein 19 | down | 1,40 | 9,52E-03 |
| ENSG00000171314 | UBQLN4 | 56893 | ubiquilin 4 | down | 1,40 | 4,39E-02 |
| ENSG00000183579 | ZFP64 | 55734 | ZFP64 zinc finger protein | down | 1,40 | 2,48E-02 |
| ENSG00000074370 | NRSN2 | 80023 | neurensin 2 | down | 1,40 | 2,54E-02 |
| ENSG00000136848 | TRAF3 | 7187 | TNF receptor associated factor 3 | down | 1,40 | 1,36E-02 |
| ENSG00000167100 | OGFR | 11054 | opioid growth factor receptor | down | 1,40 | 1,12E-02 |
| ENSG00000101347 | SLC39A3 | 29985 | solute carrier family 39 member 3 | down | 1,40 | 3,92E-04 |
| ENSG00000167670 | POMGNT2 | 84892 | protein O-linked mannose N-acetylglucosaminyltransferase 2 (beta 1,4-) | down | 1,40 | 2,18E-02 |
| ENSG00000148400 | SPG7 | 6687 | SPG7, paraplegin matrix AAA peptidase subunit | down | 1,40 | 7,74E-03 |
| ENSG00000161653 | CLP1 | 10978 | cleavage and polyadenylation factor I subunit 1 | down | 1,40 | 1,05E-02 |
| ENSG00000127824 | LSM4 | 25804 | LSM4 homolog, U6 small nuclear RNA and mRNA degradation associated | down | 1,40 | 2,18E-02 |
| ENSG00000100036 | CD320 | 51293 | CD320 molecule | down | 1,40 | 1,01E-02 |
| ENSG00000165912 | BID | 637 | BH3 interacting domain death agonist | down | 1,40 | 2,43E-02 |
| ENSG00000130590 | ANKRD16 | 54522 | ankyrin repeat domain 16 | down | 1,40 | 4,20E-03 |
| ENSG00000167384 | NOP56 | 10528 | NOP56 ribonucleoprotein | down | 1,40 | 2,85E-02 |
| ENSG00000253293 | EMILIN2 | 84034 | elastin microfibril interfacer 2 | down | 1,40 | 4,61E-02 |
| ENSG00000110931 | MTFMT | 123263 | mitochondrial methionyl-tRNA formyltransferase | down | 1,40 | 4,59E-02 |
| ENSG00000116237 | RNASEH2C | 84153 | ribonuclease H2 subunit C | down | 1,40 | 6,10E-03 |
| ENSG00000175130 | SMPD4 | 55627 | sphingomyelin phosphodiesterase 4 | down | 1,40 | 2,24E-03 |

| | | | | | | |
|---|---|---|---|---|---|---|
| ENSG00000139266 | GEMIN7 | 79760 | gem nuclear organelle associated protein 7 | down | 1,40 | 2,72E-03 |
| ENSG00000161642 | CYB561A3 | 220002 | cytochrome b561 family member A3 | down | 1,40 | 2,73E-03 |
| ENSG00000101412 | MLLT6 | 4302 | MLLT6, PHD finger containing | down | 1,40 | 2,01E-02 |
| ENSG00000157833 | ACACA | 31 | acetyl-CoA carboxylase alpha | down | 1,40 | 2,54E-02 |
| ENSG00000177045 | ZMYND19 | 116225 | zinc finger MYND-type containing 19 | down | 1,39 | 1,22E-02 |
| ENSG00000099377 | MUM1 | 84939 | melanoma associated antigen (mutated) 1 | down | 1,39 | 1,50E-02 |
| ENSG00000167695 | OXLD1 | 339229 | oxidoreductase like domain containing 1 | down | 1,39 | 2,94E-02 |
| ENSG00000100092 | URB2 | 9816 | URB2 ribosome biogenesis 2 homolog (S. cerevisiae) | down | 1,39 | 1,40E-02 |
| ENSG00000135776 | KAT2A | 2648 | lysine acetyltransferase 2A | down | 1,39 | 1,21E-02 |
| ENSG00000137198 | PIGM | 93183 | phosphatidylinositol glycan anchor biosynthesis class M | down | 1,39 | 4,67E-02 |
| ENSG00000179981 | C4orf48 | 401115 | chromosome 4 open reading frame 48 | down | 1,39 | 1,37E-02 |
| ENSG00000101670 | MCAT | 27349 | malonyl-CoA-acyl carrier protein transacylase | down | 1,39 | 3,37E-03 |
| ENSG00000176909 | RTKN | 6242 | rhotekin | down | 1,39 | 3,83E-02 |
| ENSG00000104356 | PIP5K1C | 23396 | phosphatidylinositol-4-phosphate 5-kinase type 1 gamma | down | 1,39 | 1,50E-03 |
| ENSG00000198959 | PPIL2 | 23759 | peptidylprolyl isomerase like 2 | down | 1,39 | 6,53E-03 |
| ENSG00000204366 | SLC9A3R2 | 9351 | SLC9A3 regulator 2 | down | 1,39 | 2,36E-02 |
| ENSG00000119686 | PANK4 | 55229 | pantothenate kinase 4 | down | 1,39 | 4,37E-02 |
| ENSG00000173402 | CCDC183-AS1 | 100131193 | CCDC183 antisense RNA 1 | down | 1,39 | 4,68E-02 |
| ENSG00000180354 | JRK | 8629 | Jrk helix-turn-helix protein | down | 1,39 | 4,18E-02 |
| ENSG00000160993 | AGAP3 | 116988 | ArfGAP with GTPase domain, ankyrin repeat and PH domain 3 | down | 1,39 | 4,98E-02 |
| ENSG00000029993 | LCMT2 | 9836 | leucine carboxyl methyltransferase 2 | down | 1,38 | 1,67E-02 |
| ENSG00000107816 | NFATC2IP | 84901 | nuclear factor of activated T-cells 2 interacting protein | down | 1,38 | 2,19E-02 |
| ENSG00000149639 | SNRNP35 | 11066 | small nuclear ribonucleoprotein U11/U12 subunit 35 | down | 1,38 | 4,95E-02 |
| ENSG00000108312 | SYNGR2 | 9144 | synaptogyrin 2 | down | 1,38 | 4,75E-04 |
| ENSG00000173918 | TMEM223 | 79064 | transmembrane protein 223 | down | 1,38 | 2,44E-02 |
| ENSG00000168763 | NLRX1 | 79671 | NLR family member X1 | down | 1,38 | 1,23E-02 |
| ENSG00000108179 | CENPO | 79172 | centromere protein O | down | 1,38 | 4,87E-02 |
| ENSG00000147130 | PHLDA3 | 23612 | pleckstrin homology like domain family A member 3 | down | 1,38 | 1,77E-02 |
| ENSG00000157350 | BCL7C | 9274 | BCL tumor suppressor 7C | down | 1,38 | 3,60E-04 |
| ENSG00000167523 | ENGASE | 64772 | endo-beta-N-acetylglucosaminidase | down | 1,38 | 3,04E-02 |
| ENSG00000076351 | DDX28 | 55794 | DEAD-box helicase 28 | down | 1,38 | 1,42E-02 |
| ENSG00000133216 | FAIM | 55179 | Fas apoptotic inhibitory molecule | down | 1,38 | 3,24E-02 |
| ENSG00000144120 | HDAC4 | 9759 | histone deacetylase 4 | down | 1,38 | 1,08E-02 |
| ENSG00000166508 | AKAP1 | 8165 | A-kinase anchoring protein 1 | down | 1,38 | 4,54E-02 |
| ENSG00000198324 | PPP1R9B | 84687 | protein phosphatase 1 regulatory subunit 9B | down | 1,38 | 4,33E-02 |
| ENSG00000249471 | RGS12 | 6002 | regulator of G protein signaling 12 | down | 1,37 | 2,75E-02 |
| ENSG00000114853 | RILPL2 | 196383 | Rab interacting lysosomal protein like 2 | down | 1,37 | 2,24E-02 |
| ENSG00000263528 | MFSD12 | 126321 | major facilitator superfamily domain containing 12 | down | 1,37 | 5,21E-03 |
| ENSG00000179627 | PYGO2 | 90780 | pygopus family PHD finger 2 | down | 1,37 | 1,37E-02 |
| ENSG00000205362 | CDK2AP1 | 8099 | cyclin dependent kinase 2 associated protein 1 | down | 1,37 | 2,11E-02 |
| ENSG00000125148 | SLC35A4 | 113829 | solute carrier family 35 member A4 | down | 1,37 | 5,67E-03 |
| ENSG00000125454 | TWF2 | 11344 | twinfilin actin binding protein 2 | down | 1,37 | 7,95E-03 |
| ENSG00000160606 | MAPK12 | 6300 | mitogen-activated protein kinase 12 | down | 1,37 | 3,32E-03 |
| ENSG00000188070 | DIDO1 | 11083 | death inducer-obliterator 1 | down | 1,37 | 4,03E-03 |
| ENSG00000142733 | FBXO9 | 26268 | F-box protein 9 | down | 1,37 | 4,01E-02 |
| ENSG00000135245 | FN3KRP | 79672 | fructosamine 3 kinase related protein | down | 1,37 | 1,70E-02 |
| ENSG00000183935 | CAVIN1 | 284119 | caveolae associated protein 1 | down | 1,37 | 3,95E-02 |
| ENSG00000162512 | IRF2BPL | 64207 | interferon regulatory factor 2 binding protein like | down | 1,37 | 2,58E-02 |
| ENSG00000160051 | ZNF629 | 23361 | zinc finger protein 629 | down | 1,37 | 2,48E-02 |
| ENSG00000265688 | AKT1 | 207 | AKT serine/threonine kinase 1 | down | 1,37 | 1,43E-03 |
| ENSG00000161547 | GDPD5 | 81544 | glycerophosphodiester phosphodiesterase domain containing 5 | down | 1,37 | 1,27E-02 |
| ENSG00000261373 | RAVER1 | 125950 | ribonucleoprotein, PTB binding 1 | down | 1,37 | 1,10E-02 |
| ENSG00000179958 | PLEKHJ1 | 55111 | pleckstrin homology domain containing J1 | down | 1,37 | 2,14E-02 |
| ENSG00000132819 | PQLC1 | 80148 | PQ loop repeat containing 1 | down | 1,37 | 2,89E-03 |
| ENSG00000130309 | SLC25A23 | 79085 | solute carrier family 25 member 23 | down | 1,37 | 1,88E-02 |
| ENSG00000230513 | LOC102724984 | 102724984 | uncharacterized protein KIAA2013 | down | 1,37 | 1,03E-02 |
| ENSG00000120437 | HDHD5 | 27440 | haloacid dehalogenase like hydrolase domain containing 5 | down | 1,37 | 1,14E-02 |
| ENSG00000164920 | KIAA2013 | 90231 | KIAA2013 | down | 1,37 | 1,03E-02 |
| ENSG00000108511 | LPIN3 | 64900 | lipin 3 | down | 1,36 | 4,77E-02 |
| ENSG00000118655 | ICAM3 | 3385 | intercellular adhesion molecule 3 | down | 1,36 | 1,59E-02 |
| ENSG00000102178 | CLPTM1 | 1209 | CLPTM1, transmembrane protein | down | 1,36 | 2,35E-02 |
| ENSG00000068078 | AGTRAP | 57085 | angiotensin II receptor associated protein | down | 1,36 | 2,20E-02 |
| ENSG00000136999 | CARM1 | 10498 | coactivator associated arginine methyltransferase 1 | down | 1,36 | 1,61E-02 |
| ENSG00000088305 | BRD3 | 8019 | bromodomain containing 3 | down | 1,36 | 3,35E-02 |
| ENSG00000186283 | LYN | 4067 | LYN proto-oncogene, Src family tyrosine kinase | down | 1,36 | 2,35E-02 |
| ENSG00000178718 | EEFSEC | 60678 | eukaryotic elongation factor, selenocysteine-tRNA specific | down | 1,36 | 4,96E-02 |
| ENSG00000068001 | CD82 | 3732 | CD82 molecule | down | 1,36 | 1,80E-02 |
| ENSG00000266074 | PSMG4 | 389362 | proteasome assembly chaperone 4 | down | 1,36 | 4,37E-02 |
| ENSG00000143772 | NSDHL | 50814 | NAD(P) dependent steroid dehydrogenase-like | down | 1,36 | 2,00E-02 |
| ENSG00000178935 | TARBP2 | 6895 | TARBP2, RISC loading complex RNA binding subunit | down | 1,36 | 3,80E-03 |
| ENSG00000147155 | CMTM7 | 112616 | CKLF like MARVEL transmembrane domain containing 7 | down | 1,36 | 1,80E-02 |
| ENSG00000181274 | SRM | 6723 | spermidine synthase | down | 1,35 | 8,70E-03 |
| ENSG00000183647 | SLC9A3R1 | 9368 | SLC9A3 regulator 1 | down | 1,35 | 4,57E-02 |
| ENSG00000239521 | TUSC2 | 11334 | tumor suppressor candidate 2 | down | 1,35 | 2,24E-03 |
| ENSG00000168758 | RTL10 | 79680 | retrotransposon Gag like 10 | down | 1,35 | 4,79E-02 |
| ENSG00000163050 | METRN | 79006 | meteorin, glial cell differentiation regulator | down | 1,35 | 3,70E-02 |
| ENSG00000014138 | DNAAF5 | 54919 | dynein axonemal assembly factor 5 | down | 1,35 | 3,68E-02 |
| ENSG00000214717 | ALG1 | 56052 | ALG1, chitobiosyldiphosphodolichol beta-mannosyltransferase | down | 1,35 | 3,86E-02 |
| ENSG00000260804 | C1orf35 | 79169 | chromosome 1 open reading frame 35 | down | 1,35 | 1,04E-02 |
| ENSG00000171084 | MEX3D | 399664 | mex-3 RNA binding family member D | down | 1,35 | 2,00E-02 |
| ENSG00000167536 | PKM | 5315 | pyruvate kinase M1/2 | down | 1,35 | 4,17E-02 |
| ENSG00000126243 | SLC5A6 | 8884 | solute carrier family 5 member 6 | down | 1,35 | 8,35E-03 |
| ENSG00000175283 | MLYCD | 23417 | malonyl-CoA decarboxylase | down | 1,35 | 2,54E-02 |
| ENSG00000168268 | CCDC85B | 11007 | coiled-coil domain containing 85B | down | 1,35 | 4,66E-02 |
| ENSG00000138111 | ABHD11 | 83451 | abhydrolase domain containing 11 | down | 1,35 | 2,73E-03 |
| ENSG00000158292 | ORMDL3 | 94103 | ORMDL sphingolipid biosynthesis regulator 3 | down | 1,35 | 1,52E-02 |
| ENSG00000151364 | ENTPD6 | 955 | ectonucleoside triphosphate diphosphohydrolase 6 (putative) | down | 1,35 | 4,75E-04 |
| ENSG00000141858 | UMPS | 7372 | uridine monophosphate synthetase | down | 1,34 | 4,41E-02 |
| ENSG00000165795 | MYBBP1A | 10514 | MYB binding protein 1a | down | 1,34 | 6,88E-03 |
| ENSG00000240990 | CSNK1G2 | 1455 | casein kinase 1 gamma 2 | down | 1,34 | 2,24E-03 |
| ENSG00000157240 | BCAT2 | 587 | branched chain amino acid transaminase 2 | down | 1,34 | 1,97E-02 |
| ENSG00000198794 | PPP1R16A | 84988 | protein phosphatase 1 regulatory subunit 16A | down | 1,34 | 3,18E-02 |
| ENSG00000167799 | INTS9 | 55756 | integrator complex subunit 9 | down | 1,34 | 4,01E-02 |
| ENSG00000107130 | TGFBR2 | 7048 | transforming growth factor beta receptor 2 | down | 1,34 | 2,75E-02 |
| ENSG00000136295 | PTPA | 5524 | protein phosphatase 2 phosphatase activator | down | 1,34 | 2,58E-02 |
| ENSG00000169884 | ZNF48 | 197407 | zinc finger protein 48 | down | 1,34 | 1,26E-02 |
| ENSG00000223658 | MRTO4 | 51154 | MRT4 homolog, ribosome maturation factor | down | 1,34 | 4,38E-02 |
| ENSG00000100307 | MOCS3 | 27304 | molybdenum cofactor synthesis 3 | down | 1,34 | 6,80E-04 |
| ENSG00000153395 | RXRA | 6256 | retinoid X receptor alpha | down | 1,34 | 3,68E-03 |
| ENSG00000162433 | CPTP | 80772 | ceramide-1-phosphate transfer protein | down | 1,34 | 1,22E-02 |
| ENSG00000166133 | MCRIP2 | 84331 | MAPK regulated corepressor interacting protein 2 | down | 1,34 | 1,99E-02 |
| ENSG00000169258 | LOC100287175 | 100287175 | uncharacterized LOC100287175 | down | 1,34 | 1,99E-02 |
| ENSG00000135617 | TWNK | 56652 | twinkle mtDNA helicase | down | 1,34 | 4,19E-02 |
| ENSG00000118707 | MON1A | 84315 | MON1 homolog A, secretory trafficking associated | down | 1,34 | 1,31E-04 |
| ENSG00000185340 | CYBA | 1535 | cytochrome b-245 alpha chain | down | 1,34 | 1,10E-02 |
| ENSG00000169136 | BACE2 | 25825 | beta-site APP-cleaving enzyme 2 | down | 1,34 | 1,69E-02 |
| ENSG00000260442 | CHMP1A | 5119 | charged multivesicular body protein 1A | down | 1,34 | 1,28E-05 |
| ENSG00000198176 | YWHAH | 7533 | tyrosine 3-monooxygenase/tryptophan 5-monooxygenase activation protein | down | 1,33 | 4,80E-02 |
| ENSG00000092964 | THOP1 | 7064 | thimet oligopeptidase 1 | down | 1,33 | 4,33E-02 |
| ENSG00000158882 | NUDT19 | 390916 | nudix hydrolase 19 | down | 1,33 | 1,68E-02 |
| ENSG00000104524 | ALG3 | 10195 | ALG3, alpha-1,3- mannosyltransferase | down | 1,33 | 3,99E-02 |
| ENSG00000076706 | RPS6KA4 | 8986 | ribosomal protein S6 kinase A4 | down | 1,33 | 5,68E-03 |
| ENSG00000089505 | MED24 | 9862 | mediator complex subunit 24 | down | 1,33 | 2,93E-02 |
| ENSG00000176454 | DPH2 | 1802 | DPH2 homolog | down | 1,33 | 6,99E-04 |
| ENSG00000013810 | SLC39A4 | 55630 | solute carrier family 39 member 4 | down | 1,33 | 4,56E-03 |
| ENSG00000143126 | MRPL4 | 51073 | mitochondrial ribosomal protein L4 | down | 1,33 | 1,12E-03 |
| ENSG00000149503 | ZNF814 | 730051 | zinc finger protein 814 | down | 1,33 | 4,31E-02 |
| ENSG00000173218 | STARD7 | 56910 | StAR related lipid transfer domain containing 7 | down | 1,33 | 1,21E-02 |
| ENSG00000123416 | RRP1B | 23076 | ribosomal RNA processing 1B | down | 1,33 | 4,94E-02 |
| ENSG00000169683 | RTL6 | 84247 | retrotransposon Gag like 6 | down | 1,33 | 1,42E-02 |
| ENSG00000184675 | ZDHHC16 | 84287 | zinc finger DHHC-type containing 16 | down | 1,33 | 7,02E-03 |
| ENSG00000168906 | TMEM191A | 84222 | transmembrane protein 191A (pseudogene) | down | 1,33 | 4,66E-02 |
| ENSG00000213853 | SSH3 | 54961 | slingshot protein phosphatase 3 | down | 1,33 | 3,12E-02 |
| ENSG00000204519 | IFRD2 | 7866 | interferon related developmental regulator 2 | down | 1,33 | 2,14E-03 |
| ENSG00000137486 | TMEM185A | 84548 | transmembrane protein 185A | down | 1,33 | 2,96E-02 |
| ENSG00000153048 | IRF3 | 3661 | interferon regulatory factor 3 | down | 1,33 | 2,59E-03 |
| ENSG00000156381 | ATAD3B | 83858 | ATPase family, AAA domain containing 3B | down | 1,33 | 3,64E-02 |
| ENSG00000236144 | HCFC1 | 3054 | host cell factor C1 | down | 1,33 | 4,82E-04 |
| ENSG00000146250 | RRS1 | 23212 | ribosome biogenesis regulator homolog | down | 1,32 | 2,10E-02 |
| ENSG00000188807 | TRAPPC2L | 51693 | trafficking protein particle complex 2 like | down | 1,32 | 2,18E-02 |
| ENSG00000100413 | RBM4B | 83759 | RNA binding motif protein 4B | down | 1,32 | 3,23E-02 |
| ENSG00000105649 | FAM53B | 9679 | family with sequence similarity 53 member B | down | 1,32 | 4,66E-02 |
| ENSG00000043143 | PAXX | 286257 | PAXX, non-homologous end joining factor | down | 1,32 | 2,48E-02 |
| ENSG00000138449 | NDUFAF8 | 284184 | NADH:ubiquinone oxidoreductase complex assembly factor 8 | down | 1,32 | 4,13E-02 |
| ENSG00000118526 | DDX23 | 9416 | DEAD-box helicase 23 | down | 1,32 | 1,26E-02 |
| ENSG00000132016 | GYS1 | 2997 | glycogen synthase 1 | down | 1,32 | 2,64E-02 |
| ENSG00000133466 | STX10 | 8677 | syntaxin 10 | down | 1,32 | 4,33E-02 |
| ENSG00000138172 | ADGRA3 | 166647 | adhesion G protein-coupled receptor A3 | down | 1,32 | 4,88E-02 |
| ENSG00000140511 | C15orf39 | 56905 | chromosome 15 open reading frame 39 | down | 1,32 | 2,75E-02 |
| ENSG00000182057 | USP5 | 8078 | ubiquitin specific peptidase 5 | down | 1,32 | 1,02E-02 |
| ENSG00000187566 | CDIPT | 10423 | CDP-diacylglycerol--inositol 3-phosphatidyltransferase | down | 1,32 | 3,13E-03 |
| ENSG00000160326 | PLXNB1 | 5364 | plexin B1 | down | 1,32 | 3,64E-02 |
| ENSG00000260063 | COMMD9 | 29099 | COMM domain containing 9 | down | 1,31 | 4,53E-02 |
| ENSG00000226887 | KIAA0930 | 23313 | KIAA0930 | down | 1,31 | 4,87E-02 |
| ENSG00000175745 | CDAN1 | 146059 | codanin 1 | down | 1,31 | 2,78E-02 |
| ENSG00000137310 | NMB | 4828 | neuromedin B | down | 1,31 | 2,57E-02 |
| ENSG00000197256 | ZC3H4 | 23211 | zinc finger CCCH-type containing 4 | down | 1,31 | 2,06E-02 |
| ENSG00000187123 | IGSF8 | 93185 | immunoglobulin superfamily member 8 | down | 1,31 | 9,76E-03 |
| ENSG00000100297 | TRAPPC1 | 58485 | trafficking protein particle complex 1 | down | 1,31 | 2,31E-02 |
| ENSG00000168286 | HRAS | 3265 | HRas proto-oncogene, GTPase | down | 1,31 | 2,19E-03 |
| ENSG00000181284 | SPOUT1 | 51490 | SPOUT domain containing methyltransferase 1 | down | 1,31 | 4,10E-04 |
| ENSG00000004660 | HDDC3 | 374659 | HD domain containing 3 | down | 1,31 | 6,37E-03 |
| ENSG00000183048 | PGAM5 | 192111 | PGAM family member 5, mitochondrial serine/threonine protein phosphatase | down | 1,31 | 3,69E-02 |
| ENSG00000197457 | TBL3 | 10607 | transducin beta like 3 | down | 1,31 | 6,92E-03 |
| ENSG00000162913 | SMTN | 6525 | smoothelin | down | 1,31 | 1,50E-03 |
| ENSG00000128191 | ITGB1BP1 | 9270 | integrin subunit beta 1 binding protein 1 | down | 1,31 | 2,96E-02 |
| ENSG00000154146 | STK32C | 282974 | serine/threonine kinase 32C | down | 1,31 | 1,81E-02 |
| ENSG00000157193 | ZBTB45 | 84878 | zinc finger and BTB domain containing 45 | down | 1,31 | 1,67E-02 |
| ENSG00000270885 | GRWD1 | 83743 | glutamate rich WD repeat containing 1 | down | 1,31 | 3,08E-03 |
| ENSG00000243701 | PTDSS1 | 9791 | phosphatidylserine synthase 1 | down | 1,31 | 3,90E-02 |
| ENSG00000162062 | ING5 | 84289 | inhibitor of growth family member 5 | down | 1,31 | 4,39E-02 |
| ENSG00000166997 | C17orf62 | 79415 | chromosome 17 open reading frame 62 | down | 1,31 | 1,97E-03 |
| ENSG00000163453 | SH2B1 | 25970 | SH2B adaptor protein 1 | down | 1,31 | 1,82E-02 |
| ENSG00000132481 | TMEM120B | 144404 | transmembrane protein 120B | down | 1,31 | 3,95E-02 |
| ENSG00000196372 | POLR2E | 5434 | RNA polymerase II subunit E | down | 1,31 | 3,18E-03 |
| ENSG00000168792 | CD81 | 975 | CD81 molecule | down | 1,31 | 1,16E-02 |
| ENSG00000135414 | FADD | 8772 | Fas associated via death domain | down | 1,31 | 8,90E-03 |
| ENSG00000164056 | POLD2 | 5425 | DNA polymerase delta 2, accessory subunit | down | 1,30 | 1,93E-02 |
| ENSG00000177732 | GTF3C2 | 2976 | general transcription factor IIIC subunit 2 | down | 1,30 | 1,10E-02 |
| ENSG00000099194 | COMT | 1312 | catechol-O-methyltransferase | down | 1,30 | 1,09E-02 |
| ENSG00000188368 | RALGDS | 5900 | ral guanine nucleotide dissociation stimulator | down | 1,30 | 3,50E-02 |
| ENSG00000155016 | PTDSS2 | 81490 | phosphatidylserine synthase 2 | down | 1,30 | 1,47E-02 |
| ENSG00000135476 | CBX8 | 57332 | chromobox 8 | down | 1,30 | 2,19E-02 |
| ENSG00000259583 | POLR1A | 25885 | RNA polymerase I subunit A | down | 1,30 | 4,33E-02 |
| ENSG00000185090 | FUOM | 282969 | fucose mutarotase | down | 1,30 | 1,85E-02 |
| ENSG00000196659 | ELAC2 | 60528 | elaC ribonuclease Z 2 | down | 1,30 | 2,77E-02 |
| ENSG00000169047 | TSR3 | 115939 | TSR3, acp transferase ribosome maturation factor | down | 1,30 | 2,73E-03 |
| ENSG00000179409 | AGPAT5 | 55326 | 1-acylglycerol-3-phosphate O-acyltransferase 5 | down | 1,30 | 4,71E-02 |
| ENSG00000134222 | PUSL1 | 126789 | pseudouridylate synthase-like 1 | down | 1,30 | 2,85E-02 |
| ENSG00000102760 | ARF3 | 377 | ADP ribosylation factor 3 | down | 1,30 | 4,57E-02 |
| ENSG00000120075 | TGFBRAP1 | 9392 | transforming growth factor beta receptor associated protein 1 | down | 1,30 | 1,40E-02 |
| ENSG00000090447 | L3MBTL2 | 83746 | L3MBTL2, polycomb repressive complex 1 subunit | down | 1,30 | 1,42E-02 |
| ENSG00000267534 | CNOT9 | 9125 | CCR4-NOT transcription complex subunit 9 | down | 1,30 | 4,91E-02 |
| ENSG00000164045 | AGPAT3 | 56894 | 1-acylglycerol-3-phosphate O-acyltransferase 3 | down | 1,30 | 9,80E-03 |
| ENSG00000178531 | HMOX2 | 3163 | heme oxygenase 2 | down | 1,29 | 4,87E-02 |
| ENSG00000137124 | PPP5C | 5536 | protein phosphatase 5 catalytic subunit | down | 1,29 | 4,15E-02 |
| ENSG00000169105 | TAB1 | 10454 | TGF-beta activated kinase 1 (MAP3K7) binding protein 1 | down | 1,29 | 1,94E-02 |
| ENSG00000237149 | TOMM40 | 10452 | translocase of outer mitochondrial membrane 40 | down | 1,29 | 4,10E-04 |
| ENSG00000108852 | EIF5A | 1984 | eukaryotic translation initiation factor 5A | down | 1,29 | 1,18E-02 |
| ENSG00000170779 | FAM89B | 23625 | family with sequence similarity 89 member B | down | 1,29 | 9,48E-03 |
| ENSG00000141696 | MRPS12 | 6183 | mitochondrial ribosomal protein S12 | down | 1,29 | 9,89E-03 |
| ENSG00000173638 | OGFOD3 | 79701 | 2-oxoglutarate and iron dependent oxygenase domain containing 3 | down | 1,29 | 3,76E-03 |
| ENSG00000186480 | mars-02 | 92935 | methionyl-tRNA synthetase 2, mitochondrial | down | 1,29 | 2,14E-02 |
| ENSG00000186185 | FAM189B | 10712 | family with sequence similarity 189 member B | down | 1,29 | 2,61E-02 |
| ENSG00000072954 | SUMF1 | 285362 | sulfatase modifying factor 1 | down | 1,29 | 3,58E-02 |
| ENSG00000171617 | CFL1 | 1072 | cofilin 1 | down | 1,29 | 3,03E-02 |
| ENSG00000181938 | ITM2C | 81618 | integral membrane protein 2C | down | 1,29 | 2,28E-02 |
| ENSG00000155254 | PHB | 5245 | prohibitin | down | 1,29 | 3,99E-02 |
| ENSG00000237649 | SLC12A9 | 56996 | solute carrier family 12 member 9 | down | 1,29 | 4,51E-03 |
| ENSG00000120318 | ERCC2 | 2068 | ERCC excision repair 2, TFIIH core complex helicase subunit | down | 1,29 | 2,21E-02 |
| ENSG00000169710 | INTS5 | 80789 | integrator complex subunit 5 | down | 1,29 | 8,32E-03 |
| ENSG00000137135 | TMEM250 | 90120 | transmembrane protein 250 | down | 1,29 | 9,37E-03 |
| ENSG00000260027 | SLC25A39 | 51629 | solute carrier family 25 member 39 | down | 1,29 | 1,05E-02 |
| ENSG00000110108 | UBAC1 | 10422 | UBA domain containing 1 | down | 1,29 | 2,18E-02 |
| ENSG00000163638 | GHDC | 84514 | GH3 domain containing | down | 1,28 | 7,18E-03 |
| ENSG00000015532 | ATP11A | 23250 | ATPase phospholipid transporting 11A | down | 1,28 | 4,20E-02 |
| ENSG00000188322 | RAB11FIP3 | 9727 | RAB11 family interacting protein 3 | down | 1,28 | 2,24E-02 |
| ENSG00000226137 | AKT2 | 208 | AKT serine/threonine kinase 2 | down | 1,28 | 6,33E-03 |
| ENSG00000157870 | TPI1 | 7167 | triosephosphate isomerase 1 | down | 1,28 | 3,59E-02 |
| ENSG00000105137 | NMRAL1 | 57407 | NmrA like redox sensor 1 | down | 1,28 | 3,99E-02 |
| ENSG00000110921 | PLXNA1 | 5361 | plexin A1 | down | 1,28 | 9,70E-04 |
| ENSG00000018408 | PTMS | 5763 | parathymosin | down | 1,28 | 1,26E-02 |
| ENSG00000138030 | MAZ | 4150 | MYC associated zinc finger protein | down | 1,28 | 4,88E-02 |
| ENSG00000176170 | TEX264 | 51368 | testis expressed 264 | down | 1,28 | 3,33E-02 |
| ENSG00000263843 | WDR5 | 11091 | WD repeat domain 5 | down | 1,28 | 3,82E-03 |
| ENSG00000163888 | KRI1 | 65095 | KRI1 homolog | down | 1,28 | 3,87E-02 |
| ENSG00000162981 | UPF1 | 5976 | UPF1, RNA helicase and ATPase | down | 1,27 | 2,47E-02 |

| | | | | | | |
|---|---|---|---|---|---|---|
| ENSG00000172893 | TATDN2 | 9797 | TatD DNase domain containing 2 | down | 1,27 | 4,27E-02 |
| ENSG00000183479 | CHID1 | 66005 | chitinase domain containing 1 | down | 1,27 | 2,24E-03 |
| ENSG00000135723 | CCDC86 | 79080 | coiled-coil domain containing 86 | down | 1,27 | 4,83E-03 |
| ENSG00000143590 | DHX37 | 57647 | DEAH-box helicase 37 | down | 1,27 | 3,18E-02 |
| ENSG00000168061 | PRELID1 | 27166 | PRELI domain containing 1 | down | 1,27 | 8,79E-03 |
| ENSG00000073111 | GGA2 | 23062 | golgi associated, gamma adaptin ear containing, ARF binding protein 2 | down | 1,27 | 3,87E-02 |
| ENSG00000135439 | EIF3B | 8662 | eukaryotic translation initiation factor 3 subunit B | down | 1,27 | 1,77E-02 |
| ENSG00000138080 | MTA2 | 9219 | metastasis associated 1 family member 2 | down | 1,27 | 2,46E-02 |
| ENSG00000186871 | RAB1B | 81876 | RAB1B, member RAS oncogene family | down | 1,27 | 4,10E-04 |
| ENSG00000265763 | BAK1 | 578 | BCL2 antagonist/killer 1 | down | 1,27 | 1,42E-02 |
| ENSG00000161682 | EI24 | 9538 | EI24, autophagy associated transmembrane protein | down | 1,27 | 2,32E-02 |
| ENSG00000168496 | U2AF2 | 11338 | U2 small nuclear RNA auxiliary factor 2 | down | 1,27 | 8,21E-03 |
| ENSG00000145675 | SIRT3 | 23410 | sirtuin 3 | down | 1,27 | 4,50E-02 |
| ENSG00000162878 | CAPN1 | 823 | calpain 1 | down | 1,27 | 2,65E-02 |
| ENSG00000104549 | HNRNPH3 | 3189 | heterogeneous nuclear ribonucleoprotein H3 | down | 1,27 | 4,80E-02 |
| ENSG00000172731 | ZNF544 | 27300 | zinc finger protein 544 | down | 1,27 | 4,86E-02 |
| ENSG00000183578 | ZNF668 | 79759 | zinc finger protein 668 | down | 1,27 | 3,50E-02 |
| ENSG00000130193 | DAGLA | 747 | diacylglycerol lipase alpha | down | 1,27 | 4,47E-02 |
| ENSG00000161888 | SAFB2 | 9667 | scaffold attachment factor B2 | down | 1,26 | 1,67E-02 |
| ENSG00000112118 | PRDX2 | 7001 | peroxiredoxin 2 | down | 1,26 | 3,15E-02 |
| ENSG00000160999 | CCDC127 | 133957 | coiled-coil domain containing 127 | down | 1,26 | 4,68E-02 |
| ENSG00000224420 | SNAPC4 | 6621 | small nuclear RNA activating complex polypeptide 4 | down | 1,26 | 4,39E-02 |
| ENSG00000073670 | PMPCA | 23203 | peptidase, mitochondrial processing alpha subunit | down | 1,26 | 2,78E-02 |
| ENSG00000140534 | NOL6 | 65083 | nucleolar protein 6 | down | 1,26 | 3,93E-02 |
| ENSG00000116396 | PLAGL2 | 5326 | PLAG1 like zinc finger 2 | down | 1,26 | 4,81E-02 |
| ENSG00000150347 | GRAMD4 | 23151 | GRAM domain containing 4 | down | 1,26 | 3,91E-02 |
| ENSG00000171817 | ANAPC2 | 29882 | anaphase promoting complex subunit 2 | down | 1,26 | 2,00E-02 |
| ENSG00000164976 | TMEM115 | 11070 | transmembrane protein 115 | down | 1,26 | 8,32E-03 |
| ENSG00000127585 | PLIN3 | 10226 | perilipin 3 | down | 1,26 | 4,54E-02 |
| ENSG00000165507 | CTDSP1 | 58190 | CTD small phosphatase 1 | down | 1,26 | 4,35E-03 |
| ENSG00000188486 | TMEM129 | 92305 | transmembrane protein 129 | down | 1,25 | 2,68E-02 |
| ENSG00000113368 | APEX2 | 27301 | apurinic/apyrimidinic endodeoxyribonuclease 2 | down | 1,25 | 2,35E-02 |
| ENSG00000160298 | URB1 | 9875 | URB1 ribosome biogenesis 1 homolog (S. cerevisiae) | down | 1,25 | 2,44E-02 |
| ENSG00000076248 | EIF4EBP2 | 1979 | eukaryotic translation initiation factor 4E binding protein 2 | down | 1,25 | 2,07E-02 |
| ENSG00000225439 | IL17RA | 23765 | interleukin 17 receptor A | down | 1,25 | 4,96E-02 |
| ENSG00000132646 | LINC00665 | 100506930 | long intergenic non-protein coding RNA 665 | down | 1,25 | 4,62E-02 |
| ENSG00000205978 | ELOF1 | 84337 | elongation factor 1 homolog | down | 1,25 | 2,51E-02 |
| ENSG00000179348 | NUDT22 | 84304 | nudix hydrolase 22 | down | 1,25 | 3,17E-02 |
| ENSG00000148734 | ASPHD1 | 253982 | aspartate beta-hydroxylase domain containing 1 | down | 1,25 | 2,98E-02 |
| ENSG00000162063 | DUS1L | 64118 | dihydrouridine synthase 1 like | down | 1,25 | 3,37E-03 |
| ENSG00000165244 | WDR6 | 11180 | WD repeat domain 6 | down | 1,25 | 9,81E-03 |
| ENSG00000116133 | RRP7A | 27341 | ribosomal RNA processing 7 homolog A | down | 1,25 | 6,61E-03 |
| ENSG00000237424 | SLC35C1 | 55343 | solute carrier family 35 member C1 | down | 1,25 | 2,93E-02 |
| ENSG00000101003 | ARMC6 | 93436 | armadillo repeat containing 6 | down | 1,25 | 3,95E-02 |
| ENSG00000174804 | APRT | 353 | adenine phosphoribosyltransferase | down | 1,25 | 1,64E-03 |
| ENSG00000198053 | SCAMP4 | 113178 | secretory carrier membrane protein 4 | down | 1,25 | 1,79E-02 |
| ENSG00000115884 | STK11 | 6794 | serine/threonine kinase 11 | down | 1,25 | 3,06E-02 |
| ENSG00000132688 | DKC1 | 1736 | dyskerin pseudouridine synthase 1 | down | 1,25 | 4,47E-02 |
| ENSG00000178409 | LLGL1 | 3996 | LLGL1, scribble cell polarity complex component | down | 1,25 | 4,38E-02 |
| ENSG00000157657 | NSMF | 26012 | NMDA receptor synaptonuclear signaling and neuronal migration factor | down | 1,25 | 2,66E-02 |
| ENSG00000126882 | ACD | 65057 | ACD, shelterin complex subunit and telomerase recruitment factor | down | 1,25 | 2,08E-02 |
| ENSG00000145604 | CSK | 1445 | C-terminal Src kinase | down | 1,24 | 5,68E-03 |
| ENSG00000262576 | NT5C3B | 115024 | 5'-nucleotidase, cytosolic IIIB | down | 1,24 | 3,10E-03 |
| ENSG00000188227 | PPP1CA | 5499 | protein phosphatase 1 catalytic subunit alpha | down | 1,24 | 9,77E-03 |
| ENSG00000173546 | GTF3C5 | 9328 | general transcription factor IIIC subunit 5 | down | 1,24 | 3,90E-02 |
| ENSG00000082497 | TRUB2 | 26995 | TruB pseudouridine synthase family member 2 | down | 1,24 | 1,55E-02 |
| ENSG00000165905 | PRPF19 | 27339 | pre-mRNA processing factor 19 | down | 1,24 | 4,42E-02 |
| ENSG00000240583 | RAB15 | 376267 | RAB15, member RAS oncogene family | down | 1,24 | 1,05E-02 |
| ENSG00000184564 | EFTUD2 | 9343 | elongation factor Tu GTP binding domain containing 2 | down | 1,24 | 4,00E-02 |
| ENSG00000105011 | ANKH | 56172 | ANKH inorganic pyrophosphate transport regulator | down | 1,24 | 4,15E-02 |
| ENSG00000166741 | GPAT4 | 137964 | glycerol-3-phosphate acyltransferase 4 | down | 1,24 | 4,76E-02 |
| ENSG00000106003 | POLR1C | 9533 | RNA polymerase I subunit C | down | 1,24 | 4,72E-02 |
| ENSG00000182489 | STUB1 | 10273 | STIP1 homology and U-box containing protein 1 | down | 1,24 | 4,83E-03 |
| ENSG00000205861 | WRNIP1 | 56897 | Werner helicase interacting protein 1 | down | 1,24 | 3,85E-02 |
| ENSG00000168913 | CIAO1 | 9391 | cytosolic iron-sulfur assembly component 1 | down | 1,24 | 1,26E-02 |
| ENSG00000008853 | GPI | 2821 | glucose-6-phosphate isomerase | down | 1,24 | 1,55E-02 |
| ENSG00000169598 | TRIM28 | 10155 | tripartite motif containing 28 | down | 1,24 | 4,18E-02 |
| ENSG00000204128 | SUMO3 | 6612 | small ubiquitin-like modifier 3 | down | 1,24 | 4,67E-02 |
| ENSG00000254369 | ARAP1 | 116985 | ArfGAP with RhoGAP domain, ankyrin repeat and PH domain 1 | down | 1,24 | 4,91E-02 |
| ENSG00000099284 | AP1M1 | 8907 | adaptor related protein complex 1 mu 1 subunit | down | 1,24 | 1,31E-02 |
| ENSG00000165556 | MIGA2 | 84895 | mitoguardin 2 | down | 1,24 | 1,70E-02 |
| ENSG00000167191 | CASKIN2 | 57513 | CASK interacting protein 2 | down | 1,24 | 1,33E-02 |
| ENSG00000159556 | WDR46 | 9277 | WD repeat domain 46 | down | 1,23 | 1,10E-02 |
| ENSG00000092621 | HTRA2 | 27429 | HtrA serine peptidase 2 | down | 1,23 | 2,64E-02 |
| ENSG00000257913 | CCAR2 | 57805 | cell cycle and apoptosis regulator 2 | down | 1,23 | 2,13E-02 |
| ENSG00000266094 | AGPAT2 | 10555 | 1-acylglycerol-3-phosphate O-acyltransferase 2 | down | 1,23 | 2,89E-03 |
| ENSG00000128714 | SMARCD1 | 6602 | SWI/SNF related, matrix associated, actin dependent regulator of chromatin | down | 1,23 | 2,53E-02 |
| ENSG00000222033 | CYB5R3 | 1727 | cytochrome b5 reductase 3 | down | 1,23 | 4,75E-02 |
| ENSG00000144355 | POLR2B | 5431 | RNA polymerase II subunit B | down | 1,23 | 3,81E-02 |
| ENSG00000137440 | MED25 | 81857 | mediator complex subunit 25 | down | 1,23 | 4,80E-02 |
| ENSG00000156966 | LRRC14 | 9684 | leucine rich repeat containing 14 | down | 1,23 | 4,06E-02 |
| ENSG00000180884 | ZC3H18 | 124245 | zinc finger CCCH-type containing 18 | down | 1,23 | 1,01E-02 |
| ENSG00000174684 | LOC105372824 | 105372824 | uncharacterized LOC105372824 | down | 1,23 | 2,36E-02 |
| ENSG00000121454 | PDXK | 8566 | pyridoxal kinase | down | 1,23 | 2,36E-02 |
| ENSG00000143147 | SLC35C2 | 51006 | solute carrier family 35 member C2 | down | 1,23 | 2,36E-02 |
| ENSG00000071575 | LAGE3 | 8270 | L antigen family member 3 | down | 1,23 | 3,98E-02 |
| ENSG00000129173 | NHP2 | 55651 | NHP2 ribonucleoprotein | down | 1,23 | 3,16E-02 |
| ENSG00000162073 | RPS6KB2 | 6199 | ribosomal protein S6 kinase B2 | down | 1,22 | 1,01E-02 |
| ENSG00000254389 | ULK3 | 25989 | unc-51 like kinase 3 | down | 1,22 | 4,14E-02 |
| ENSG00000125378 | ZBTB2 | 57621 | zinc finger and BTB domain containing 2 | down | 1,22 | 4,96E-02 |
| ENSG00000175414 | SLC38A7 | 55238 | solute carrier family 38 member 7 | down | 1,22 | 3,97E-02 |
| ENSG00000185585 | FAAP20 | 199990 | Fanconi anemia core complex associated protein 20 | down | 1,22 | 1,49E-02 |
| ENSG00000174871 | GNAI2 | 2771 | G protein subunit alpha i2 | down | 1,22 | 3,66E-02 |
| ENSG00000204335 | PPM1F | 9647 | protein phosphatase, Mg2+/Mn2+ dependent 1F | down | 1,22 | 2,37E-02 |
| ENSG00000187808 | TRIM32 | 22954 | tripartite motif containing 32 | down | 1,22 | 3,96E-02 |
| ENSG00000179403 | SMPD2 | 6610 | sphingomyelin phosphodiesterase 2 | down | 1,22 | 4,96E-02 |
| ENSG00000163032 | BAX | 581 | BCL2 associated X, apoptosis regulator | down | 1,22 | 2,62E-02 |
| ENSG00000173894 | SLC2A8 | 29988 | solute carrier family 2 member 8 | down | 1,22 | 3,59E-03 |
| ENSG00000128408 | TMEM134 | 80194 | transmembrane protein 134 | down | 1,21 | 6,92E-03 |
| ENSG00000139292 | CXXC1 | 30827 | CXXC finger protein 1 | down | 1,21 | 2,87E-02 |
| ENSG00000168824 | MAEA | 10296 | macrophage erythroblast attacher | down | 1,21 | 7,72E-03 |
| ENSG00000106031 | SMYD5 | 10322 | SMYD family member 5 | down | 1,21 | 2,45E-02 |
| ENSG00000071246 | RCC1L | 81554 | RCC1 like | down | 1,21 | 9,63E-03 |
| ENSG00000185818 | MTCH1 | 23787 | mitochondrial carrier 1 | down | 1,21 | 5,52E-04 |
| ENSG00000168646 | PRPF31 | 26121 | pre-mRNA processing factor 31 | down | 1,21 | 2,75E-02 |
| ENSG00000184588 | AKT1S1 | 84335 | AKT1 substrate 1 | down | 1,21 | 4,33E-02 |
| ENSG00000122035 | DNAJB6 | 10049 | DnaJ heat shock protein family (Hsp40) member B6 | down | 1,20 | 4,90E-02 |
| ENSG00000276043 | TBL1X | 6907 | transducin beta like 1 X-linked | down | 1,20 | 2,85E-02 |
| ENSG00000078900 | PFN1 | 5216 | profilin 1 | down | 1,20 | 3,99E-02 |
| ENSG00000198910 | ERCC3 | 2071 | ERCC excision repair 3, TFIIH core complex helicase subunit | down | 1,20 | 2,37E-02 |
| ENSG00000120068 | LASP1 | 3927 | LIM and SH3 protein 1 | down | 1,20 | 4,26E-02 |
| ENSG00000275793 | CNDP2 | 55748 | carnosine dipeptidase 2 | down | 1,20 | 4,79E-02 |
| ENSG00000186603 | CLPP | 8192 | caseinolytic mitochondrial matrix peptidase proteolytic subunit | down | 1,20 | 3,82E-02 |
| ENSG00000107485 | PCBD1 | 5092 | pterin-4 alpha-carbinolamine dehydratase 1 | down | 1,20 | 1,58E-02 |
| ENSG00000128268 | PISD | 23761 | phosphatidylserine decarboxylase | down | 1,20 | 2,48E-02 |
| ENSG00000176697 | CHERP | 10523 | calcium homeostasis endoplasmic reticulum protein | down | 1,20 | 2,06E-02 |
| ENSG00000257167 | RPTOR | 57521 | regulatory associated protein of MTOR complex 1 | down | 1,20 | 1,16E-02 |
| ENSG00000147100 | KIF1C | 10749 | kinesin family member 1C | down | 1,20 | 2,46E-02 |
| ENSG00000146122 | REXO4 | 57109 | REX4 homolog, 3'-5' exonuclease | down | 1,19 | 2,24E-02 |
| ENSG00000268388 | RHOT2 | 89941 | ras homolog family member T2 | down | 1,19 | 2,93E-02 |
| ENSG00000013619 | LRRC47 | 57470 | leucine rich repeat containing 47 | down | 1,19 | 3,59E-02 |
| ENSG00000166033 | ZDHHC3 | 51304 | zinc finger DHHC-type containing 3 | down | 1,19 | 4,45E-02 |
| ENSG00000115380 | CACNB3 | 784 | calcium voltage-gated channel auxiliary subunit beta 3 | down | 1,19 | 3,30E-02 |
| ENSG00000171724 | DNPEP | 23549 | aspartyl aminopeptidase | down | 1,19 | 2,68E-02 |
| ENSG00000122870 | RIC8A | 60626 | RIC8 guanine nucleotide exchange factor A | down | 1,19 | 2,13E-02 |
| ENSG00000171388 | PHRF1 | 57661 | PHD and ring finger domains 1 | down | 1,19 | 2,37E-02 |
| ENSG00000150594 | INO80E | 283899 | INO80 complex subunit E | down | 1,19 | 2,68E-02 |
| ENSG00000136732 | RECQL5 | 9400 | RecQ like helicase 5 | down | 1,18 | 4,33E-02 |
| ENSG00000165617 | GNA11 | 2767 | G protein subunit alpha 11 | down | 1,18 | 4,37E-02 |
| ENSG00000138685 | TIMM22 | 29928 | translocase of inner mitochondrial membrane 22 | down | 1,18 | 2,55E-02 |
| ENSG00000173705 | C16orf70 | 80262 | chromosome 16 open reading frame 70 | down | 1,18 | 4,56E-02 |
| ENSG00000129824 | ZNRF1 | 84937 | zinc and ring finger 1 | down | 1,18 | 1,73E-02 |
| ENSG00000105928 | AES | 166 | amino-terminal enhancer of split | down | 1,18 | 2,85E-02 |
| ENSG00000104368 | HNRNPUL1 | 11100 | heterogeneous nuclear ribonucleoprotein U like 1 | down | 1,17 | 2,87E-02 |
| ENSG00000204442 | RALY | 22913 | RALY heterogeneous nuclear ribonucleoprotein | down | 1,17 | 4,88E-02 |
| ENSG00000214548 | ZC3H7B | 23264 | zinc finger CCCH-type containing 7B | down | 1,16 | 2,53E-02 |
| ENSG00000183873 | PTBP1 | 5725 | polypyrimidine tract binding protein 1 | down | 1,16 | 4,21E-02 |
| ENSG00000131002 | CTBP1 | 1487 | C-terminal binding protein 1 | down | 1,16 | 3,77E-02 |
| ENSG00000121005 | TEX261 | 113419 | testis expressed 261 | down | 1,16 | 2,92E-02 |
| ENSG00000145242 | AHCY | 191 | adenosylhomocysteinase | down | 1,14 | 2,15E-02 |
| ENSG00000169026 | CLEC16A | 23274 | C-type lectin domain containing 16A | down | 1,13 | 4,19E-02 |
