## Supplemental Table 4 for "Transcriptome profiling of human colonic cells exposed to the gut pathobiont *Streptococcus gallolyticus* subsp. *gallolyticus*"

**Supplemental Table S4. Transcriptome:** List of 41 genes differentially expressed in HT29 and in FHC cells infected with *SGG* UCN34 vs *SGM* during 24 h.

| EnsEMBL ID | Gene Symbol | Entrez Gene ID | Gene Name | Regulation | Fold-Change | Adjusted P-Value |
|---|---|---|---|---|---|---|
| ENSG00000013441 | CLK1 | 1195 | CDC like kinase 1 | up | 2,58 | 3,17E-02 |
| ENSG00000070501 | POLB | 5423 | DNA polymerase beta | up | 1,33 | 3,69E-02 |
| ENSG00000141564 | RPTOR | 57521 | regulatory associated protein of MTC | down | 1,22 | 2,44E-02 |
| ENSG00000169718 | DUS1L | 64118 | dihydrouridine synthase 1 like | down | 1,26 | 1,42E-02 |
| ENSG00000154832 | CXXC1 | 30827 | CXXC finger protein 1 | down | 1,26 | 3,96E-02 |
| ENSG00000164077 | MON1A | 84315 | MON1 homolog A, secretory traffickir | down | 1,27 | 1,14E-03 |
| ENSG00000185024 | BRF1 | 2972 | BRF1, RNA polymerase III transcripti | down | 1,28 | 4,22E-02 |
| ENSG00000238227 | TMEM250 | 90120 | transmembrane protein 250 | down | 1,31 | 2,69E-02 |
| ENSG00000124217 | MOCS3 | 27304 | molybdenum cofactor synthesis 3 | down | 1,31 | 5,55E-03 |
| ENSG00000110104 | CCDC86 | 79080 | solute carrier family 38 member 7 | down | 1,31 | 5,55E-03 |
| ENSG00000103042 | SLC38A7 | 55238 | solute carrier family 38 member 7 | down | 1,31 | 1,92E-02 |
| ENSG00000241878 | PISD | 23761 | phosphatidylserine decarboxylase | down | 1,32 | 1,88E-03 |
| ENSG00000085872 | CHERP | 10523 | calcium homeostasis endoplasmic re | down | 1,32 | 1,14E-03 |
| ENSG00000168040 | FADD | 8772 | Fas associated via death domain | down | 1,32 | 3,52E-02 |
| ENSG00000103653 | CSK | 1445 | C-terminal Src kinase | down | 1,33 | 6,44E-04 |
| ENSG00000070047 | PHRF1 | 57661 | PHD and ring finger domains 1 | down | 1,34 | 2,14E-04 |
| ENSG00000181830 | SLC35C1 | 55343 | solute carrier family 35 member C1 | down | 1,35 | 1,68E-02 |
| ENSG00000130254 | SAFB2 | 9667 | scaffold attachment factor B2 | down | 1,35 | 7,73E-03 |
| ENSG00000126003 | PLAGL2 | 5326 | PLAG1 like zinc finger 2 | down | 1,36 | 4,82E-02 |
| ENSG00000124160 | NCOA5 | 57727 | nuclear receptor coactivator 5 | down | 1,36 | 4,00E-02 |
| ENSG00000180035 | ZNF48 | 197407 | zinc finger protein 48 | down | 1,37 | 3,92E-02 |
| ENSG00000177303 | CASKIN2 | 57513 | CASK interacting protein 2 | down | 1,38 | 3,22E-04 |
| ENSG00000197119 | SLC25A29 | 123096 | solute carrier family 25 member 29 | down | 1,39 | 4,74E-02 |
| ENSG00000172534 | HCFC1 | 3054 | host cell factor C1 | down | 1,40 | 3,40E-05 |
| ENSG00000119574 | ZBTB45 | 84878 | zinc finger and BTB domain containi | down | 1,40 | 7,67E-03 |
| ENSG00000183963 | SMTN | 6525 | smoothelin | down | 1,40 | 3,75E-05 |
| ENSG00000161847 | RAVER1 | 125950 | ribonucleoprotein, PTB binding 1 | down | 1,42 | 2,01E-02 |
| ENSG00000099364 | FBXL19 | 54620 | F-box and leucine rich repeat proteir | down | 1,44 | 2,65E-02 |
| ENSG00000150990 | DHX37 | 57647 | DEAH-box helicase 37 | down | 1,44 | 4,36E-03 |
| ENSG00000179041 | RRS1 | 23212 | ribosome biogenesis regulator homo | down | 1,47 | 5,55E-03 |
| ENSG00000185085 | INTS5 | 80789 | integrator complex subunit 5 | down | 1,47 | 6,77E-05 |
| ENSG00000007376 | RPUSD1 | 113000 | RNA pseudouridylate synthase doma | down | 1,47 | 5,55E-03 |
| ENSG00000167716 | WDR81 | 124997 | WD repeat domain 81 | down | 1,49 | 3,52E-03 |
| ENSG00000135763 | URB2 | 9816 | URB2 ribosome biogenesis 2 homolo | down | 1,49 | 1,41E-02 |
| ENSG00000165684 | SNAPC4 | 6621 | small nuclear RNA activating comple | down | 1,51 | 1,34E-03 |
| ENSG00000167394 | ZNF668 | 79759 | zinc finger protein 668 | down | 1,56 | 6,77E-05 |
| ENSG00000167173 | C15orf39 | 56905 | chromosome 15 open reading frame | down | 1,66 | 6,77E-05 |
| ENSG00000101670 | LIPG | 9388 | lipase G, endothelial type | down | 2,02 | 2,62E-05 |
| ENSG00000168906 | MAT2A | 4144 | methionine adenosyltransferase 2A | down | 2,29 | 6,64E-03 |
| ENSG00000164056 | SPRY1 | 10252 | sprouty RTK signaling antagonist 1 | down | 2,68 | 1,51E-03 |
| ENSG00000127528 | KLF2 | 10365 | Kruppel like factor 2 | down | 2,76 | 4,29E-03 |
