## Supplemental Table 5 for "Transcriptome profiling of human colonic cells exposed to the gut pathobiont *Streptococcus gallolyticus* subsp. *gallolyticus*"

**Supplemental Table S5.** KEGG: List of 54 significantly up- and down-regulated pathways found by KEGG pathway analysis in HT29 cells after *SGG* UCN34 vs *SGM* during 24 h.

| Link to KEGG Pathway | Pathway Description (KEGG) | Nb Genes in Pathway | Nb Regulated Genes (Up / Down) | P-Value (All) | P-Value (Up) | P-Value (Down) | Min P-Value | Min BH P-Value |
|---|---|---|---|---|---|---|---|---|
| hsa04141 | **Protein processing in endoplasmic reticulum** | 169 | 48 (41/7) | 1,06E-10 | 4,55E-17 | NA | 4,55E-17 | 1,10E-14 |
| hsa03030 | DNA replication | 36 | 13 (1/12) | 1,69E-04 | NA | 1,42E-06 | 1,42E-06 | 3,69E-04 |
| hsa01100 | Metabolic pathways | 1219 | 149 (51/98) | 1,60E-02 | NA | 8,54E-06 | 8,54E-06 | 1,11E-03 |
| hsa00240 | Pyrimidine metabolism | 101 | 24 (6/18) | 2,05E-04 | NA | 1,84E-05 | 1,84E-05 | 1,59E-03 |
| hsa00100 | Steroid biosynthesis | 20 | 8 (0/8) | 2,90E-03 | NA | 4,66E-05 | 4,66E-05 | 3,02E-03 |
| hsa00230 | Purine metabolism | 176 | 31 (7/24) | 4,11E-03 | NA | 4,77E-05 | 4,77E-05 | 2,47E-03 |
| hsa04152 | **AMPK signaling pathway** | 123 | 28 (12/16) | 1,15E-04 | 4,71E-02 | 2,00E-03 | 1,15E-04 | 1,58E-02 |
| hsa04144 | Endocytosis | 241 | 41 (27/14) | 1,68E-03 | 1,99E-04 | NA | 1,99E-04 | 2,37E-02 |
| hsa04068 | FoxO signaling pathway | 134 | 28 (18/10) | 5,04E-04 | 4,03E-04 | NA | 4,03E-04 | 2,30E-02 |
| hsa04722 | Neurotrophin signaling pathway | 120 | 26 (11/15) | 4,78E-04 | NA | 4,21E-03 | 4,78E-04 | 2,61E-02 |
| hsa04910 | Insulin signaling pathway | 138 | 28 (10/18) | 8,14E-04 | NA | 9,18E-04 | 8,14E-04 | 3,17E-02 |
| hsa03410 | Base excision repair | 33 | 9 (1/8) | 1,71E-02 | NA | 1,41E-03 | 1,41E-03 | 5,12E-02 |
| hsa04962 | Vasopressin-regulated water reabsorption | 44 | 10 (9/1) | 3,40E-02 | 1,44E-03 | NA | 1,44E-03 | 8,35E-02 |
| hsa04668 | **TNF signaling pathway** | 107 | 22 (12/10) | 2,87E-03 | 1,92E-02 | NA | 2,87E-03 | 8,47E-02 |
| hsa04931 | Insulin resistance | 108 | 22 (12/10) | 3,22E-03 | 2,05E-02 | NA | 3,22E-03 | 7,81E-02 |
| hsa04015 | Rap1 signaling pathway | 210 | 22 (0/22) | NA | NA | 3,53E-03 | 3,53E-03 | 9,72E-02 |
| hsa01130 | Biosynthesis of antibiotics | 212 | 35 (13/22) | 6,31E-03 | NA | 3,95E-03 | 3,95E-03 | 9,79E-02 |
| hsa00564 | Glycerophospholipid metabolism | 95 | 18 (5/13) | 1,72E-02 | NA | 4,11E-03 | 4,11E-03 | 9,28E-02 |
| hsa03020 | RNA polymerase | 32 | 10 (3/7) | 4,14E-03 | NA | 5,91E-03 | 4,14E-03 | 8,46E-02 |
| hsa03060 | Protein export | 23 | 6 (6/0) | NA | 5,15E-03 | NA | 5,15E-03 | 2,21E-01 |
| hsa05214 | Glioma | 65 | 15 (6/9) | 5,92E-03 | NA | 2,05E-02 | 5,92E-03 | 1,10E-01 |
| hsa04130 | SNARE interactions in vesicular transport | 34 | 7 (7/0) | NA | 6,56E-03 | NA | 6,56E-03 | 2,33E-01 |
| hsa05223 | Non-small cell lung cancer | 56 | 12 (3/9) | 2,68E-02 | NA | 8,69E-03 | 8,69E-03 | 1,49E-01 |
| **hsa04150** | **mTOR signaling pathway** | **58** | **13 (4/9)** | **1,43E-02** | **NA** | **1,07E-02** | **1,07E-02** | **1,70E-01** |
| hsa04360 | Axon guidance | 127 | 23 (10/13) | 1,07E-02 | NA | 3,59E-02 | 1,07E-02 | 1,70E-01 |
| hsa01212 | Fatty acid metabolism | 48 | 8 (0/8) | NA | NA | 1,23E-02 | 1,23E-02 | 1,83E-01 |
| hsa04922 | Glucagon signaling pathway | 99 | 19 (10/9) | 1,23E-02 | NA | NA | 1,23E-02 | 1,83E-01 |
| hsa04666 | Fc gamma R-mediated phagocytosis | 84 | 15 (4/11) | 4,93E-02 | NA | 1,27E-02 | 1,27E-02 | 1,77E-01 |
| hsa05230 | Central carbon metabolism in cancer | 64 | 14 (5/9) | 1,29E-02 | NA | 1,88E-02 | 1,29E-02 | 1,81E-01 |
| hsa00562 | Inositol phosphate metabolism | 71 | 15 (8/7) | 1,30E-02 | NA | NA | 1,30E-02 | 1,74E-01 |
| hsa04210 | Apoptosis | 62 | 9 (0/9) | NA | NA | 1,57E-02 | 1,57E-02 | 2,04E-01 |
| hsa04012 | ErbB signaling pathway | 87 | 17 (10/7) | 1,59E-02 | 3,15E-02 | NA | 1,59E-02 | 1,91E-01 |
| hsa00970 | Aminoacyl-tRNA biosynthesis | 66 | 9 (9/0) | NA | 1,77E-02 | NA | 1,77E-02 | 4,62E-01 |
| hsa04710 | Circadian rhythm | 31 | 6 (6/0) | NA | 1,86E-02 | NA | 1,86E-02 | 4,34E-01 |
| **hsa04115** | **p53 signaling pathway** | **67** | **14 (8/6)** | **1,86E-02** | **NA** | **NA** | **1,86E-02** | **1,88E-01** |
| hsa03420 | Nucleotide excision repair | 47 | 11 (4/7) | 2,02E-02 | NA | 3,61E-02 | 2,02E-02 | 1,95E-01 |
| hsa04920 | Adipocytokine signaling pathway | 70 | 14 (5/9) | 2,62E-02 | NA | 3,05E-02 | 2,62E-02 | 2,38E-01 |
| hsa05200 | Pathways in cancer | 393 | 31 (0/31) | NA | NA | 2,67E-02 | 2,67E-02 | 2,84E-01 |
| hsa00061 | Fatty acid biosynthesis | 13 | 4 (0/4) | NA | NA | 2,82E-02 | 2,82E-02 | 2,87E-01 |
| **hsa04010** | **MAPK signaling pathway** | **253** | **37 (21/16)** | **3,00E-02** | **3,17E-02** | **NA** | **3,00E-02** | **2,52E-01** |
| hsa04110 | Cell cycle | 124 | 13 (0/13) | NA | NA | 3,05E-02 | 3,05E-02 | 2,85E-01 |
| hsa05160 | Hepatitis C | 133 | 22 (10/12) | 3,26E-02 | NA | NA | 3,26E-02 | 2,63E-01 |
| hsa05212 | Pancreatic cancer | 65 | 13 (6/7) | 3,33E-02 | NA | NA | 3,33E-02 | 2,61E-01 |
| hsa01230 | Biosynthesis of amino acids | 72 | 9 (0/9) | NA | NA | 3,54E-02 | 3,54E-02 | 3,12E-01 |
| hsa04022 | cGMP-PKG signaling pathway | 158 | 25 (12/13) | 3,54E-02 | NA | NA | 3,54E-02 | 2,61E-01 |
| hsa05215 | Prostate cancer | 88 | 16 (9/7) | 3,60E-02 | NA | NA | 3,60E-02 | 2,58E-01 |
| hsa05213 | Endometrial cancer | 52 | 11 (4/7) | 3,87E-02 | NA | NA | 3,87E-02 | 2,68E-01 |
| hsa04071 | Sphingolipid signaling pathway | 120 | 20 (8/12) | 3,97E-02 | NA | NA | 3,97E-02 | 2,67E-01 |
| hsa04919 | Thyroid hormone signaling pathway | 115 | 19 (7/12) | 4,89E-02 | NA | 4,02E-02 | 4,02E-02 | 3,00E-01 |
| hsa00533 | Glycosaminoglycan biosynthesis - keratan sulfate | 15 | 4 (0/4) | NA | NA | 4,16E-02 | 4,16E-02 | 3,16E-01 |
| hsa04070 | Phosphatidylinositol signaling system | 98 | 17 (8/9) | 4,40E-02 | NA | NA | 4,40E-02 | 2,86E-01 |
| hsa05210 | Colorectal cancer | 62 | 8 (0/8) | NA | NA | 4,41E-02 | 4,41E-02 | 2,97E-01 |
| hsa04320 | Dorso-ventral axis formation | 27 | 5 (5/0) | NA | 4,54E-02 | NA | 4,54E-02 | 2,97E-01 |
| hsa04915 | Estrogen signaling pathway | 99 | 17 (8/9) | 4,77E-02 | NA | NA | 4,77E-02 | 2,99E-01 |

***Regulated Genes in Pathways***

| Link to KEGG Pathway | Pathway Description (KEGG) | Gene Symbol | Gene Name | Link to Entrez Gene | Regulation | Fold-Change | Adjusted P-Value |
|---|---|---|---|---|---|---|---|
| hsa04141 | Protein processing in endoplasmic reticulum | EIF2AK3 | eukaryotic translation initiation factor 2 alpha kinase 3 | 9451 | up | 2,82 | 1,11E-03 |
| hsa04141 | Protein processing in endoplasmic reticulum | PREB | prolactin regulatory element binding | 10113 | up | 1,76 | 3,49E-02 |
| hsa04141 | Protein processing in endoplasmic reticulum | DNAJC3 | DnaJ heat shock protein family (Hsp40) member C3 | 5611 | up | 2,75 | 2,24E-03 |
| hsa04141 | Protein processing in endoplasmic reticulum | SYVN1 | synoviolin 1 | 84447 | up | 2,33 | 1,93E-02 |
| hsa04141 | Protein processing in endoplasmic reticulum | SEC24A | SEC24 homolog A, COPII coat complex component | 10802 | up | 1,71 | 3,95E-03 |
| hsa04141 | Protein processing in endoplasmic reticulum | HSP90B1 | heat shock protein 90 beta family member 1 | 7184 | up | 1,71 | 2,12E-02 |
| hsa04141 | Protein processing in endoplasmic reticulum | DNAJC1 | DnaJ heat shock protein family (Hsp40) member C1 | 64215 | up | 1,37 | 4,59E-02 |
| hsa04141 | Protein processing in endoplasmic reticulum | ERO1B | endoplasmic reticulum oxidoreductase 1 beta | 56605 | up | 8,94 | 1,01E-02 |
| hsa04141 | Protein processing in endoplasmic reticulum | HERPUD1 | homocysteine inducible ER protein with ubiquitin like do | 9709 | up | 4,55 | 1,11E-03 |
| hsa04141 | Protein processing in endoplasmic reticulum | HSPA5 | heat shock protein family A (Hsp70) member 5 | 3309 | up | 5,02 | 6,98E-04 |
| hsa04141 | Protein processing in endoplasmic reticulum | DNAJB11 | DnaJ heat shock protein family (Hsp40) member B11 | 51726 | up | 2,08 | 1,17E-02 |
| hsa04141 | Protein processing in endoplasmic reticulum | BAK1 | BCL2 antagonist/killer 1 | 578 | down | 1,27 | 1,41E-02 |
| hsa04141 | Protein processing in endoplasmic reticulum | UBE2G2 | ubiquitin conjugating enzyme E2 G2 | 7327 | down | 1,41 | 4,12E-03 |
| hsa04141 | Protein processing in endoplasmic reticulum | SEC24D | SEC24 homolog D, COPII coat complex component | 9871 | up | 4,45 | 6,80E-04 |
| hsa04141 | Protein processing in endoplasmic reticulum | MAPK8 | mitogen-activated protein kinase 8 | 5599 | up | 1,38 | 3,05E-02 |
| hsa04141 | Protein processing in endoplasmic reticulum | SSR1 | signal sequence receptor subunit 1 | 6745 | up | 1,73 | 2,06E-02 |
| hsa04141 | Protein processing in endoplasmic reticulum | UBE2J1 | ubiquitin conjugating enzyme E2 J1 | 51465 | up | 1,86 | 8,97E-03 |
| hsa04141 | Protein processing in endoplasmic reticulum | SELENOS | selenoprotein S | 55829 | up | 2,02 | 1,16E-02 |
| hsa04141 | Protein processing in endoplasmic reticulum | ATXN3 | ataxin 3 | 4287 | up | 1,51 | 4,51E-03 |
| hsa04141 | Protein processing in endoplasmic reticulum | RAD23B | RAD23 homolog B, nucleotide excision repair protein | 5887 | up | 1,23 | 3,34E-02 |
| hsa04141 | Protein processing in endoplasmic reticulum | UBQLN1 | ubiquilin 1 | 29979 | up | 1,19 | 4,56E-03 |
| hsa04141 | Protein processing in endoplasmic reticulum | PPP1R15A | protein phosphatase 1 regulatory subunit 15A | 23645 | up | 4,33 | 1,06E-03 |
| hsa04141 | Protein processing in endoplasmic reticulum | TRAF2 | TNF receptor associated factor 2 | 7186 | down | 1,47 | 2,65E-02 |
| hsa04141 | Protein processing in endoplasmic reticulum | UBQLN4 | ubiquilin 4 | 56893 | down | 1,40 | 4,38E-02 |
| hsa04141 | Protein processing in endoplasmic reticulum | NFE2L2 | nuclear factor, erythroid 2 like 2 | 4780 | up | 1,88 | 2,41E-03 |
| hsa04141 | Protein processing in endoplasmic reticulum | SEL1L | SEL1L ERAD E3 ligase adaptor subunit | 6400 | up | 2,76 | 4,03E-03 |
| hsa04141 | Protein processing in endoplasmic reticulum | ERN1 | endoplasmic reticulum to nucleus signaling 1 | 2081 | up | 3,27 | 7,84E-03 |
| hsa04141 | Protein processing in endoplasmic reticulum | LMAN1 | lectin, mannose binding 1 | 3998 | up | 1,54 | 4,14E-02 |
| hsa04141 | Protein processing in endoplasmic reticulum | SSR3 | signal sequence receptor subunit 3 | 6747 | up | 1,43 | 4,56E-02 |
| hsa04141 | Protein processing in endoplasmic reticulum | DERL2 | derlin 2 | 51009 | up | 1,68 | 4,88E-02 |
| hsa04141 | Protein processing in endoplasmic reticulum | ATF6 | activating transcription factor 6 | 22926 | up | 1,68 | 1,14E-02 |
| hsa04141 | Protein processing in endoplasmic reticulum | CAPN1 | calpain 1 | 823 | down | 1,27 | 2,65E-02 |
| hsa04141 | Protein processing in endoplasmic reticulum | PLAA | phospholipase A2 activating protein | 9373 | up | 1,26 | 1,64E-02 |
| hsa04141 | Protein processing in endoplasmic reticulum | NGLY1 | N-glycanase 1 | 55768 | up | 1,38 | 1,27E-02 |
| hsa04141 | Protein processing in endoplasmic reticulum | BAX | BCL2 associated X, apoptosis regulator | 581 | down | 1,22 | 2,61E-02 |
| hsa04141 | Protein processing in endoplasmic reticulum | EDEM1 | ER degradation enhancing alpha-mannosidase like pro | 9695 | up | 2,16 | 1,32E-02 |
| hsa04141 | Protein processing in endoplasmic reticulum | SEC23B | Sec23 homolog B, coat complex II component | 10483 | up | 1,71 | 1,73E-02 |
| hsa04141 | Protein processing in endoplasmic reticulum | SEC23A | Sec23 homolog A, coat complex II component | 10484 | up | 1,46 | 2,47E-02 |
| hsa04141 | Protein processing in endoplasmic reticulum | CANX | calnexin | 821 | up | 1,36 | 3,98E-02 |
| hsa04141 | Protein processing in endoplasmic reticulum | HYOU1 | hypoxia up-regulated 1 | 10525 | up | 2,96 | 5,94E-03 |
| hsa04141 | Protein processing in endoplasmic reticulum | DNAJB1 | DnaJ heat shock protein family (Hsp40) member B1 | 3337 | up | 2,02 | 1,41E-02 |
| hsa04141 | Protein processing in endoplasmic reticulum | XBP1 | X-box binding protein 1 | 7494 | up | 2,00 | 5,67E-03 |
| hsa04141 | Protein processing in endoplasmic reticulum | STUB1 | STIP1 homology and U-box containing protein 1 | 10273 | down | 1,24 | 4,82E-03 |
| hsa04141 | Protein processing in endoplasmic reticulum | DDIT3 | DNA damage inducible transcript 3 | 1649 | up | 5,65 | 3,67E-03 |
| hsa04141 | Protein processing in endoplasmic reticulum | TRAM1 | translocation associated membrane protein 1 | 23471 | up | 1,76 | 4,37E-02 |
| hsa04141 | Protein processing in endoplasmic reticulum | ERLEC1 | endoplasmic reticulum lectin 1 | 27248 | up | 1,96 | 7,71E-03 |
| hsa04141 | Protein processing in endoplasmic reticulum | SEC31A | SEC31 homolog A, COPII coat complex component | 22872 | up | 1,53 | 3,02E-02 |
| hsa04141 | Protein processing in endoplasmic reticulum | SEC63 | SEC63 homolog, protein translocation regulator | 11231 | up | 1,60 | 8,90E-03 |
| hsa03030 | DNA replication | MCM5 | minichromosome maintenance complex component 5 | 4174 | down | 1,83 | 2,22E-02 |
| hsa03030 | DNA replication | POLA2 | DNA polymerase alpha 2, accessory subunit | 23649 | down | 1,72 | 2,46E-02 |
| hsa03030 | DNA replication | RNASEH2C | ribonuclease H2 subunit C | 84153 | down | 1,40 | 6,10E-03 |
| hsa03030 | DNA replication | RFC1 | replication factor C subunit 1 | 5981 | up | 1,15 | 2,18E-02 |
| hsa03030 | DNA replication | MCM3 | minichromosome maintenance complex component 3 | 4172 | down | 2,07 | 4,13E-02 |
| hsa03030 | DNA replication | RPA1 | replication protein A1 | 6117 | down | 1,41 | 4,93E-02 |
| hsa03030 | DNA replication | POLD2 | DNA polymerase delta 2, accessory subunit | 5425 | down | 1,30 | 1,92E-02 |
| hsa03030 | DNA replication | POLD1 | DNA polymerase delta 1, catalytic subunit | 5424 | down | 1,59 | 4,62E-02 |
| hsa03030 | DNA replication | PCNA | proliferating cell nuclear antigen | 5111 | down | 2,14 | 4,63E-02 |
| hsa03030 | DNA replication | MCM2 | minichromosome maintenance complex component 2 | 4171 | down | 2,01 | 1,42E-02 |
| hsa03030 | DNA replication | MCM7 | minichromosome maintenance complex component 7 | 4176 | down | 1,67 | 3,35E-02 |
| hsa03030 | DNA replication | POLE | DNA polymerase epsilon, catalytic subunit | 5426 | down | 1,55 | 2,67E-02 |
| hsa03030 | DNA replication | FEN1 | flap structure-specific endonuclease 1 | 2237 | down | 2,04 | 1,60E-02 |
| hsa01100 | Metabolic pathways | NT5C2 | 5'-nucleotidase, cytosolic II | 22978 | up | 1,42 | 1,00E-02 |
| hsa01100 | Metabolic pathways | A4GALT | alpha 1,4-galactosyltransferase (P blood group) | 53947 | down | 1,58 | 1,60E-02 |
| hsa01100 | Metabolic pathways | TPI1 | triosephosphate isomerase 1 | 7167 | down | 1,28 | 3,58E-02 |
| hsa01100 | Metabolic pathways | C1GALT1 | core 1 synthase, glycoprotein-N-acetylgalactosamine 3 | 56913 | up | 1,51 | 1,69E-02 |
| hsa01100 | Metabolic pathways | ALDH1B1 | aldehyde dehydrogenase 1 family member B1 | 219 | down | 1,92 | 9,37E-03 |
| hsa01100 | Metabolic pathways | GMPPB | GDP-mannose pyrophosphorylase B | 29925 | up | 1,87 | 4,52E-02 |
| hsa01100 | Metabolic pathways | CAD | carbamoyl-phosphate synthetase 2, aspartate transcar | 790 | down | 1,43 | 7,14E-03 |
| hsa01100 | Metabolic pathways | SLC33A1 | solute carrier family 33 member 1 | 9197 | up | 1,72 | 2,23E-02 |
| hsa01100 | Metabolic pathways | PIP5K1B | phosphatidylinositol-4-phosphate 5-kinase type 1 beta | 8395 | up | 3,40 | 1,84E-02 |
| hsa01100 | Metabolic pathways | PANK4 | pantothenate kinase 4 | 55229 | down | 1,39 | 4,37E-02 |
| hsa01100 | Metabolic pathways | SMPD2 | sphingomyelin phosphodiesterase 2 | 6610 | down | 1,22 | 4,95E-02 |
| hsa01100 | Metabolic pathways | MGAT3 | mannosyl (beta-1,4-)-glycoprotein beta-1,4-N-acetylgluc | 4248 | down | 2,92 | 7,16E-03 |
| hsa01100 | Metabolic pathways | POLR3F | RNA polymerase III subunit F | 10621 | up | 1,25 | 3,24E-02 |
| hsa01100 | Metabolic pathways | MTMR6 | myotubularin related protein 6 | 9107 | up | 1,33 | 2,92E-02 |
| hsa01100 | Metabolic pathways | CNDP2 | carnosine dipeptidase 2 | 55748 | down | 1,20 | 4,78E-02 |
| hsa01100 | Metabolic pathways | HSD3B7 | hydroxy-delta-5-steroid dehydrogenase, 3 beta- and st | 80270 | down | 1,64 | 3,72E-02 |
| hsa01100 | Metabolic pathways | MVK | mevalonate kinase | 4598 | down | 1,99 | 3,36E-03 |
| hsa01100 | Metabolic pathways | LPIN3 | lipin 3 | 64900 | down | 1,36 | 4,76E-02 |
| hsa01100 | Metabolic pathways | PTDSS2 | phosphatidylserine synthase 2 | 81490 | down | 1,30 | 1,46E-02 |
| hsa01100 | Metabolic pathways | PIGA | phosphatidylinositol glycan anchor biosynthesis class A | 5277 | up | 2,32 | 6,64E-03 |
| hsa01100 | Metabolic pathways | PTDSS1 | phosphatidylserine synthase 1 | 9791 | down | 1,31 | 3,90E-02 |
| hsa01100 | Metabolic pathways | GLUL | glutamate-ammonia ligase | 2752 | down | 1,47 | 6,91E-03 |
| hsa01100 | Metabolic pathways | MUT | methylmalonyl-CoA mutase | 4594 | up | 1,31 | 4,57E-03 |
| hsa01100 | Metabolic pathways | AGPAT2 | 1-acylglycerol-3-phosphate O-acyltransferase 2 | 10555 | down | 1,23 | 2,89E-03 |
| hsa01100 | Metabolic pathways | ITPKC | inositol-trisphosphate 3-kinase C | 80271 | up | 1,63 | 3,32E-02 |

| | | | | | | | |
|---|---|---|---|---|---|---|---|
| hsa01100 | Metabolic pathways | NNMT | nicotinamide N-methyltransferase | 4837 | down | 2,26 | 3,05E-02 |
| hsa01100 | Metabolic pathways | KHK | ketohexokinase | 3795 | down | 2,00 | 2,30E-02 |
| hsa01100 | Metabolic pathways | ITPKB | inositol-trisphosphate 3-kinase B | 3707 | down | 1,71 | 1,97E-02 |
| hsa01100 | Metabolic pathways | POLR1A | RNA polymerase I subunit A | 25885 | down | 1,30 | 4,32E-02 |
| hsa01100 | Metabolic pathways | MCCC1 | methylcrotonoyl-CoA carboxylase 1 | 56922 | up | 1,28 | 4,04E-02 |
| hsa01100 | Metabolic pathways | AHCY | adenosylhomocysteinase | 191 | down | 1,14 | 2,15E-02 |
| hsa01100 | Metabolic pathways | ARG2 | arginase 2 | 384 | up | 7,09 | 3,92E-04 |
| hsa01100 | Metabolic pathways | MAT2A | methionine adenosyltransferase 2A | 4144 | down | 1,79 | 2,93E-02 |
| hsa01100 | Metabolic pathways | POLA2 | DNA polymerase alpha 2, accessory subunit | 23649 | down | 1,72 | 2,46E-02 |
| hsa01100 | Metabolic pathways | ACLY | ATP citrate lyase | 47 | down | 1,41 | 3,09E-02 |
| hsa01100 | Metabolic pathways | LPCAT4 | lysophosphatidylcholine acyltransferase 4 | 254531 | down | 1,78 | 2,52E-02 |
| hsa01100 | Metabolic pathways | LIPT1 | lipoyltransferase 1 | 51601 | up | 1,63 | 1,73E-02 |
| hsa01100 | Metabolic pathways | CYP2U1 | cytochrome P450 family 2 subfamily U member 1 | 113612 | down | 1,88 | 2,19E-02 |
| hsa01100 | Metabolic pathways | POLR3C | RNA polymerase III subunit C | 10623 | up | 1,43 | 4,98E-02 |
| hsa01100 | Metabolic pathways | ALG3 | ALG3, alpha-1,3- mannosyltransferase | 10195 | down | 1,33 | 3,98E-02 |
| hsa01100 | Metabolic pathways | AK1 | adenylate kinase 1 | 203 | down | 1,57 | 9,91E-03 |
| hsa01100 | Metabolic pathways | LPCAT1 | lysophosphatidylcholine acyltransferase 1 | 79888 | down | 1,76 | 2,60E-02 |
| hsa01100 | Metabolic pathways | POLR3D | RNA polymerase III subunit D | 661 | up | 1,52 | 2,72E-02 |
| hsa01100 | Metabolic pathways | B3GNT5 | UDP-GlcNAc:betaGal beta-1,3-N-acetylglucosaminyltra | 84002 | up | 1,57 | 1,69E-02 |
| hsa01100 | Metabolic pathways | BCAT2 | branched chain amino acid transaminase 2 | 587 | down | 1,34 | 1,96E-02 |
| hsa01100 | Metabolic pathways | UGDH | UDP-glucose 6-dehydrogenase | 7358 | up | 1,53 | 3,49E-02 |
| hsa01100 | Metabolic pathways | SRM | spermidine synthase | 6723 | down | 1,35 | 8,70E-03 |
| hsa01100 | Metabolic pathways | UCK2 | uridine-cytidine kinase 2 | 7371 | down | 1,45 | 2,72E-02 |
| hsa01100 | Metabolic pathways | CSGALNACT2 | chondroitin sulfate N-acetylgalactosaminyltransferase 2 | 55454 | up | 2,21 | 1,16E-02 |
| hsa01100 | Metabolic pathways | PIP5K1C | phosphatidylinositol-4-phosphate 5-kinase type 1 gamm | 23396 | down | 1,39 | 1,49E-03 |
| hsa01100 | Metabolic pathways | PLD1 | phospholipase D1 | 5337 | up | 1,33 | 1,60E-02 |
| hsa01100 | Metabolic pathways | KYAT3 | kynurenine aminotransferase 3 | 56267 | up | 1,50 | 3,91E-02 |
| hsa01100 | Metabolic pathways | PLCG1 | phospholipase C gamma 1 | 5335 | down | 1,51 | 2,85E-02 |
| hsa01100 | Metabolic pathways | EPRS | glutamyl-prolyl-tRNA synthetase | 2058 | up | 1,42 | 1,95E-02 |
| hsa01100 | Metabolic pathways | SGMS2 | sphingomyelin synthase 2 | 166929 | up | 1,39 | 4,00E-02 |
| hsa01100 | Metabolic pathways | CDIPT | CDP-diacylglycerol--inositol 3-phosphatidyltransferase | 10423 | down | 1,32 | 3,12E-03 |
| hsa01100 | Metabolic pathways | PDXK | pyridoxal kinase | 8566 | down | 1,23 | 2,36E-02 |
| hsa01100 | Metabolic pathways | GPAT3 | glycerol-3-phosphate acyltransferase 3 | 84803 | up | 1,77 | 4,99E-02 |
| hsa01100 | Metabolic pathways | LIPG | lipase G, endothelial type | 9388 | down | 1,65 | 1,49E-03 |
| hsa01100 | Metabolic pathways | SMPD4 | sphingomyelin phosphodiesterase 4 | 55627 | down | 1,40 | 2,24E-03 |
| hsa01100 | Metabolic pathways | LSS | lanosterol synthase | 4047 | down | 1,49 | 1,65E-03 |
| hsa01100 | Metabolic pathways | EBP | emopamil binding protein (sterol isomerase) | 10682 | down | 1,71 | 1,99E-02 |
| hsa01100 | Metabolic pathways | PISD | phosphatidylserine decarboxylase | 23761 | down | 1,20 | 2,48E-02 |
| hsa01100 | Metabolic pathways | ALG2 | ALG2, alpha-1,3/1,6-mannosyltransferase | 85365 | up | 1,61 | 2,74E-02 |
| hsa01100 | Metabolic pathways | ADSS | adenylosuccinate synthase | 159 | up | 1,21 | 2,47E-02 |
| hsa01100 | Metabolic pathways | PIGW | phosphatidylinositol glycan anchor biosynthesis class W | 284098 | down | 1,53 | 1,78E-02 |
| hsa01100 | Metabolic pathways | SEPHS2 | selenophosphate synthetase 2 | 22928 | up | 2,29 | 7,52E-03 |
| hsa01100 | Metabolic pathways | DHCR7 | 7-dehydrocholesterol reductase | 1717 | down | 2,00 | 2,13E-03 |
| hsa01100 | Metabolic pathways | PKM | pyruvate kinase M1/2 | 5315 | down | 1,35 | 4,16E-02 |
| hsa01100 | Metabolic pathways | NSDHL | NAD(P) dependent steroid dehydrogenase-like | 50814 | down | 1,36 | 1,99E-02 |
| hsa01100 | Metabolic pathways | TRAK2 | trafficking kinesin protein 2 | 66008 | up | 1,51 | 2,95E-02 |
| hsa01100 | Metabolic pathways | GPAT4 | glycerol-3-phosphate acyltransferase 4 | 137964 | down | 1,24 | 4,75E-02 |
| hsa01100 | Metabolic pathways | MCAT | malonyl-CoA-acyl carrier protein transacylase | 27349 | down | 1,39 | 3,36E-03 |
| hsa01100 | Metabolic pathways | NDST1 | N-deacetylase and N-sulfotransferase 1 | 3340 | down | 1,59 | 2,40E-03 |
| hsa01100 | Metabolic pathways | HKDC1 | hexokinase domain containing 1 | 80201 | up | 3,37 | 4,73E-03 |
| hsa01100 | Metabolic pathways | CYP1A1 | cytochrome P450 family 1 subfamily A member 1 | 1543 | up | 2,69 | 1,95E-02 |
| hsa01100 | Metabolic pathways | ACACA | acetyl-CoA carboxylase alpha | 31 | down | 1,40 | 2,54E-02 |
| hsa01100 | Metabolic pathways | LPIN2 | lipin 2 | 9663 | up | 2,16 | 6,88E-03 |
| hsa01100 | Metabolic pathways | POLD2 | DNA polymerase delta 2, accessory subunit | 5425 | down | 1,30 | 1,92E-02 |
| hsa01100 | Metabolic pathways | AK4 | adenylate kinase 4 | 205 | down | 1,76 | 4,87E-02 |
| hsa01100 | Metabolic pathways | ALDH6A1 | aldehyde dehydrogenase 6 family member A1 | 4329 | up | 1,60 | 3,50E-02 |
| hsa01100 | Metabolic pathways | AASS | aminoadipate-semialdehyde synthase | 10157 | up | 2,01 | 4,95E-02 |
| hsa01100 | Metabolic pathways | POLR3H | RNA polymerase III subunit H | 171568 | down | 1,80 | 7,07E-04 |
| hsa01100 | Metabolic pathways | SPR | sepiapterin reductase | 6697 | down | 1,54 | 1,91E-02 |
| hsa01100 | Metabolic pathways | UMPS | uridine monophosphate synthetase | 7372 | down | 1,34 | 4,41E-02 |
| hsa01100 | Metabolic pathways | DCTPP1 | dCTP pyrophosphatase 1 | 79077 | down | 1,69 | 1,60E-02 |
| hsa01100 | Metabolic pathways | FUT4 | fucosyltransferase 4 | 2526 | down | 1,44 | 1,70E-03 |
| hsa01100 | Metabolic pathways | MTHFD2 | methylenetetrahydrofolate dehydrogenase (NADP+ de | 10797 | up | 2,09 | 9,37E-03 |
| hsa01100 | Metabolic pathways | ASNS | asparagine synthetase (glutamine-hydrolyzing) | 440 | up | 2,47 | 2,91E-02 |
| hsa01100 | Metabolic pathways | DOLK | dolichol kinase | 22845 | down | 1,73 | 2,74E-03 |
| hsa01100 | Metabolic pathways | FASN | fatty acid synthase | 2194 | down | 1,96 | 8,31E-03 |
| hsa01100 | Metabolic pathways | APRT | adenine phosphoribosyltransferase | 353 | down | 1,25 | 1,64E-03 |
| hsa01100 | Metabolic pathways | FAM213B | family with sequence similarity 213 member B | 127281 | down | 1,99 | 1,17E-02 |
| hsa01100 | Metabolic pathways | B4GALT2 | beta-1,4-galactosyltransferase 2 | 8704 | down | 1,45 | 2,13E-03 |
| hsa01100 | Metabolic pathways | CTH | cystathionine gamma-lyase | 1491 | up | 3,41 | 4,94E-03 |
| hsa01100 | Metabolic pathways | XYLT2 | xylosyltransferase 2 | 64132 | down | 1,98 | 2,03E-02 |
| hsa01100 | Metabolic pathways | HADHB | hydroxyacyl-CoA dehydrogenase trifunctional multienzy | 3032 | up | 1,20 | 4,66E-02 |
| hsa01100 | Metabolic pathways | COMT | catechol-O-methyltransferase | 1312 | down | 1,30 | 1,09E-02 |
| hsa01100 | Metabolic pathways | NAMPT | nicotinamide phosphoribosyltransferase | 10135 | up | 1,49 | 4,90E-02 |
| hsa01100 | Metabolic pathways | ACAT2 | acetyl-CoA acetyltransferase 2 | 39 | down | 1,69 | 2,52E-02 |
| hsa01100 | Metabolic pathways | MTHFD1L | methylenetetrahydrofolate dehydrogenase (NADP+ de | 25902 | up | 1,36 | 4,51E-02 |
| hsa01100 | Metabolic pathways | DMGDH | dimethylglycine dehydrogenase | 29958 | up | 3,92 | 1,01E-02 |
| hsa01100 | Metabolic pathways | MTMR2 | myotubularin related protein 2 | 8898 | up | 1,23 | 2,61E-02 |
| hsa01100 | Metabolic pathways | PYCR3 | pyrroline-5-carboxylate reductase 3 | 65263 | down | 1,77 | 1,11E-03 |
| hsa01100 | Metabolic pathways | NAGS | N-acetylglutamate synthase | 162417 | down | 1,62 | 4,90E-02 |
| hsa01100 | Metabolic pathways | POLR1C | RNA polymerase I subunit C | 9533 | down | 1,24 | 4,71E-02 |
| hsa01100 | Metabolic pathways | UGP2 | UDP-glucose pyrophosphorylase 2 | 7360 | up | 1,37 | 3,47E-02 |
| hsa01100 | Metabolic pathways | PGP | phosphoglycolate phosphatase | 283871 | down | 1,50 | 8,04E-03 |
| hsa01100 | Metabolic pathways | B4GAT1 | beta-1,4-glucuronyltransferase 1 | 11041 | down | 2,42 | 1,49E-02 |
| hsa01100 | Metabolic pathways | TM7SF2 | transmembrane 7 superfamily member 2 | 7108 | down | 1,51 | 4,00E-02 |
| hsa01100 | Metabolic pathways | POLR2F | RNA polymerase II subunit F | 5435 | down | 1,41 | 3,22E-02 |
| hsa01100 | Metabolic pathways | UPP1 | uridine phosphorylase 1 | 7378 | up | 4,41 | 1,62E-03 |
| hsa01100 | Metabolic pathways | PGAM1 | phosphoglycerate mutase 1 | 5223 | down | 1,62 | 5,67E-03 |
| hsa01100 | Metabolic pathways | GPI | glucose-6-phosphate isomerase | 2821 | down | 1,24 | 1,55E-02 |
| hsa01100 | Metabolic pathways | AGPAT3 | 1-acylglycerol-3-phosphate O-acyltransferase 3 | 56894 | down | 1,30 | 9,80E-03 |
| hsa01100 | Metabolic pathways | POLR2L | RNA polymerase II subunit L | 5441 | down | 1,52 | 3,58E-03 |
| hsa01100 | Metabolic pathways | ACSL1 | acyl-CoA synthetase long chain family member 1 | 2180 | down | 1,54 | 3,01E-02 |
| hsa01100 | Metabolic pathways | GCLC | glutamate-cysteine ligase catalytic subunit | 2729 | up | 2,09 | 2,27E-02 |
| hsa01100 | Metabolic pathways | PHGDH | phosphoglycerate dehydrogenase | 26227 | down | 2,34 | 2,54E-02 |
| hsa01100 | Metabolic pathways | NT5C3A | 5'-nucleotidase, cytosolic IIIA | 51251 | up | 1,49 | 1,52E-02 |
| hsa01100 | Metabolic pathways | NAT8L | N-acetyltransferase 8 like | 339983 | down | 2,79 | 1,70E-02 |
| hsa01100 | Metabolic pathways | NME4 | NME/NM23 nucleoside diphosphate kinase 4 | 4833 | down | 1,47 | 3,85E-02 |
| hsa01100 | Metabolic pathways | POLR2E | RNA polymerase II subunit E | 5434 | down | 1,31 | 3,18E-03 |
| hsa01100 | Metabolic pathways | SPHK1 | sphingosine kinase 1 | 8877 | down | 2,00 | 2,34E-02 |
| hsa01100 | Metabolic pathways | PCCA | propionyl-CoA carboxylase alpha subunit | 5095 | up | 1,37 | 2,03E-02 |
| hsa01100 | Metabolic pathways | DLD | dihydrolipoamide dehydrogenase | 1738 | up | 1,16 | 4,53E-02 |
| hsa01100 | Metabolic pathways | SQLE | squalene epoxidase | 6713 | down | 2,05 | 1,00E-02 |
| hsa01100 | Metabolic pathways | GOT1 | glutamic-oxaloacetic transaminase 1 | 2805 | up | 2,30 | 5,67E-03 |
| hsa01100 | Metabolic pathways | PFAS | phosphoribosylformylglycinamidine synthase | 5198 | down | 1,53 | 1,69E-02 |
| hsa01100 | Metabolic pathways | NT5C3B | 5'-nucleotidase, cytosolic IIIB | 115024 | down | 1,24 | 3,10E-03 |
| hsa01100 | Metabolic pathways | IMPDH1 | inosine monophosphate dehydrogenase 1 | 3614 | down | 1,56 | 1,49E-03 |
| hsa01100 | Metabolic pathways | SAT1 | spermidine/spermine N1-acetyltransferase 1 | 6303 | up | 2,30 | 3,55E-02 |
| hsa01100 | Metabolic pathways | HYAL2 | hyaluronoglucosaminidase 2 | 8692 | down | 1,71 | 7,16E-03 |
| hsa01100 | Metabolic pathways | MLYCD | malonyl-CoA decarboxylase | 23417 | down | 1,35 | 2,54E-02 |
| hsa01100 | Metabolic pathways | DNMT3B | DNA methyltransferase 3 beta | 1789 | down | 1,70 | 3,62E-02 |
| hsa01100 | Metabolic pathways | POLD1 | DNA polymerase delta 1, catalytic subunit | 5424 | down | 1,59 | 4,62E-02 |
| hsa01100 | Metabolic pathways | POLE | DNA polymerase epsilon, catalytic subunit | 5426 | down | 1,55 | 2,67E-02 |
| hsa01100 | Metabolic pathways | GFPT1 | glutamine--fructose-6-phosphate transaminase 1 | 2673 | up | 2,92 | 2,14E-03 |
| hsa01100 | Metabolic pathways | ATP6V0E2 | ATPase H+ transporting V0 subunit e2 | 155066 | down | 1,61 | 2,71E-03 |
| hsa01100 | Metabolic pathways | AGPAT5 | 1-acylglycerol-3-phosphate O-acyltransferase 5 | 55326 | down | 1,30 | 4,70E-02 |
| hsa01100 | Metabolic pathways | ALG1 | ALG1, chitobiosyldiphosphodolichol beta-mannosyltran | 56052 | down | 1,35 | 3,85E-02 |
| hsa01100 | Metabolic pathways | DHCR24 | 24-dehydrocholesterol reductase | 1718 | down | 2,17 | 4,89E-02 |
| hsa01100 | Metabolic pathways | ST3GAL2 | ST3 beta-galactoside alpha-2,3-sialyltransferase 2 | 6483 | down | 1,66 | 3,84E-03 |
| hsa01100 | Metabolic pathways | POLR2B | RNA polymerase II subunit B | 5431 | down | 1,23 | 3,81E-02 |
| hsa01100 | Metabolic pathways | ISYNA1 | inositol-3-phosphate synthase 1 | 51477 | down | 1,54 | 7,64E-03 |
| hsa01100 | Metabolic pathways | PIGM | phosphatidylinositol glycan anchor biosynthesis class M | 93183 | down | 1,39 | 4,66E-02 |
| hsa01100 | Metabolic pathways | PPT2 | palmitoyl-protein thioesterase 2 | 9374 | down | 1,51 | 1,22E-02 |
| hsa01100 | Metabolic pathways | NMRK1 | nicotinamide riboside kinase 1 | 54981 | up | 1,47 | 1,75E-02 |
| hsa01100 | Metabolic pathways | PIK3C2A | phosphatidylinositol-4-phosphate 3-kinase catalytic sub | 5286 | up | 1,30 | 1,79E-02 |
| hsa00240 | Pyrimidine metabolism | POLR3D | RNA polymerase III subunit D | 661 | up | 1,52 | 2,72E-02 |
| hsa00240 | Pyrimidine metabolism | POLE | DNA polymerase epsilon, catalytic subunit | 5426 | down | 1,55 | 2,67E-02 |
| hsa00240 | Pyrimidine metabolism | CAD | carbamoyl-phosphate synthetase 2, aspartate transcar | 790 | down | 1,43 | 7,14E-03 |
| hsa00240 | Pyrimidine metabolism | POLD1 | DNA polymerase delta 1, catalytic subunit | 5424 | down | 1,59 | 4,62E-02 |
| hsa00240 | Pyrimidine metabolism | POLR2L | RNA polymerase II subunit L | 5441 | down | 1,52 | 3,58E-03 |
| hsa00240 | Pyrimidine metabolism | NT5C3B | 5'-nucleotidase, cytosolic IIIB | 115024 | down | 1,24 | 3,10E-03 |
| hsa00240 | Pyrimidine metabolism | POLR3C | RNA polymerase III subunit C | 10623 | up | 1,43 | 4,98E-02 |
| hsa00240 | Pyrimidine metabolism | POLA2 | DNA polymerase alpha 2, accessory subunit | 23649 | down | 1,72 | 2,46E-02 |
| hsa00240 | Pyrimidine metabolism | NT5C2 | 5'-nucleotidase, cytosolic II | 22978 | up | 1,42 | 1,00E-02 |
| hsa00240 | Pyrimidine metabolism | UMPS | uridine monophosphate synthetase | 7372 | down | 1,34 | 4,41E-02 |
| hsa00240 | Pyrimidine metabolism | DCTPP1 | dCTP pyrophosphatase 1 | 79077 | down | 1,69 | 1,60E-02 |
| hsa00240 | Pyrimidine metabolism | UPP1 | uridine phosphorylase 1 | 7378 | up | 4,41 | 1,62E-03 |
| hsa00240 | Pyrimidine metabolism | POLR1A | RNA polymerase I subunit A | 25885 | down | 1,30 | 4,32E-02 |
| hsa00240 | Pyrimidine metabolism | POLR3H | RNA polymerase III subunit H | 171568 | down | 1,80 | 7,07E-04 |
| hsa00240 | Pyrimidine metabolism | ENTPD6 | ectonucleoside triphosphate diphosphohydrolase 6 (pu | 955 | down | 1,35 | 4,75E-04 |
| hsa00240 | Pyrimidine metabolism | POLR2E | RNA polymerase II subunit E | 5434 | down | 1,31 | 3,18E-03 |
| hsa00240 | Pyrimidine metabolism | POLR2B | RNA polymerase II subunit B | 5431 | down | 1,23 | 3,81E-02 |
| hsa00240 | Pyrimidine metabolism | UCK2 | uridine-cytidine kinase 2 | 7371 | down | 1,45 | 2,72E-02 |
| hsa00240 | Pyrimidine metabolism | POLR2F | RNA polymerase II subunit F | 5435 | down | 1,41 | 3,22E-02 |
| hsa00240 | Pyrimidine metabolism | POLR3F | RNA polymerase III subunit F | 10621 | up | 1,25 | 3,24E-02 |
| hsa00240 | Pyrimidine metabolism | POLD2 | DNA polymerase delta 2, accessory subunit | 5425 | down | 1,30 | 1,92E-02 |
| hsa00240 | Pyrimidine metabolism | POLR1C | RNA polymerase I subunit C | 9533 | down | 1,24 | 4,71E-02 |
| hsa00240 | Pyrimidine metabolism | NME4 | NME/NM23 nucleoside diphosphate kinase 4 | 4833 | down | 1,47 | 3,85E-02 |
| hsa00240 | Pyrimidine metabolism | NT5C3A | 5'-nucleotidase, cytosolic IIIA | 51251 | up | 1,49 | 1,52E-02 |

| | | | | | | | |
|---|---|---|---|---|---|---|---|
| hsa00100 | Steroid biosynthesis | DHCR24 | 24-dehydrocholesterol reductase | 1718 | down | 2,17 | 4,89E-02 |
| hsa00100 | Steroid biosynthesis | SQLE | squalene epoxidase | 6713 | down | 2,05 | 1,00E-02 |
| hsa00100 | Steroid biosynthesis | SOAT1 | sterol O-acyltransferase 1 | 6646 | down | 1,40 | 1,13E-02 |
| hsa00100 | Steroid biosynthesis | LSS | lanosterol synthase | 4047 | down | 1,49 | 1,65E-03 |
| hsa00100 | Steroid biosynthesis | EBP | emopamil binding protein (sterol isomerase) | 10682 | down | 1,71 | 1,99E-02 |
| hsa00100 | Steroid biosynthesis | NSDHL | NAD(P) dependent steroid dehydrogenase-like | 50814 | down | 1,36 | 1,99E-02 |
| hsa00100 | Steroid biosynthesis | DHCR7 | 7-dehydrocholesterol reductase | 1717 | down | 2,00 | 2,13E-03 |
| hsa00100 | Steroid biosynthesis | TM7SF2 | transmembrane 7 superfamily member 2 | 7108 | down | 1,51 | 4,00E-02 |
| hsa00230 | Purine metabolism | AK4 | adenylate kinase 4 | 205 | down | 1,76 | 4,87E-02 |
| hsa00230 | Purine metabolism | POLD2 | DNA polymerase delta 2, accessory subunit | 5425 | down | 1,30 | 1,92E-02 |
| hsa00230 | Purine metabolism | ADSS | adenylosuccinate synthase | 159 | up | 1,21 | 2,47E-02 |
| hsa00230 | Purine metabolism | POLR1C | RNA polymerase I subunit C | 9533 | down | 1,24 | 4,71E-02 |
| hsa00230 | Purine metabolism | NME4 | NME/NM23 nucleoside diphosphate kinase 4 | 4833 | down | 1,47 | 3,85E-02 |
| hsa00230 | Purine metabolism | ADCY3 | adenylate cyclase 3 | 109 | down | 1,45 | 1,03E-02 |
| hsa00230 | Purine metabolism | NT5C3A | 5'-nucleotidase, cytosolic IIIA | 51251 | up | 1,49 | 1,52E-02 |
| hsa00230 | Purine metabolism | HDDC3 | HD domain containing 3 | 374659 | down | 1,31 | 6,37E-03 |
| hsa00230 | Purine metabolism | POLR1A | RNA polymerase I subunit A | 25885 | down | 1,30 | 4,32E-02 |
| hsa00230 | Purine metabolism | ADPRM | ADP-ribose/CDP-alcohol diphosphatase, manganese d | 56985 | up | 2,23 | 2,72E-03 |
| hsa00230 | Purine metabolism | PDE4B | phosphodiesterase 4B | 5142 | down | 2,82 | 4,90E-02 |
| hsa00230 | Purine metabolism | POLR3H | RNA polymerase III subunit H | 171568 | down | 1,80 | 7,07E-04 |
| hsa00230 | Purine metabolism | ENTPD6 | ectonucleoside triphosphate diphosphohydrolase 6 (pu | 955 | down | 1,35 | 4,75E-04 |
| hsa00230 | Purine metabolism | POLR2E | RNA polymerase II subunit E | 5434 | down | 1,31 | 3,18E-03 |
| hsa00230 | Purine metabolism | POLR2B | RNA polymerase II subunit B | 5431 | down | 1,23 | 3,81E-02 |
| hsa00230 | Purine metabolism | PKM | pyruvate kinase M1/2 | 5315 | down | 1,35 | 4,16E-02 |
| hsa00230 | Purine metabolism | POLR2F | RNA polymerase II subunit F | 5435 | down | 1,41 | 3,22E-02 |
| hsa00230 | Purine metabolism | POLR3F | RNA polymerase III subunit F | 10621 | up | 1,25 | 3,24E-02 |
| hsa00230 | Purine metabolism | NT5C3B | 5'-nucleotidase, cytosolic IIIB | 115024 | down | 1,24 | 3,10E-03 |
| hsa00230 | Purine metabolism | PFAS | phosphoribosylformylglycinamidine synthase | 5198 | down | 1,53 | 1,69E-02 |
| hsa00230 | Purine metabolism | IMPDH1 | inosine monophosphate dehydrogenase 1 | 3614 | down | 1,56 | 1,49E-03 |
| hsa00230 | Purine metabolism | POLR3C | RNA polymerase III subunit C | 10623 | up | 1,43 | 4,98E-02 |
| hsa00230 | Purine metabolism | POLA2 | DNA polymerase alpha 2, accessory subunit | 23649 | down | 1,72 | 2,46E-02 |
| hsa00230 | Purine metabolism | NT5C2 | 5'-nucleotidase, cytosolic II | 22978 | up | 1,42 | 1,00E-02 |
| hsa00230 | Purine metabolism | POLE | DNA polymerase epsilon, catalytic subunit | 5426 | down | 1,55 | 2,67E-02 |
| hsa00230 | Purine metabolism | POLR3D | RNA polymerase III subunit D | 661 | up | 1,52 | 2,72E-02 |
| hsa00230 | Purine metabolism | GMPR | guanosine monophosphate reductase | 2766 | down | 1,65 | 3,56E-02 |
| hsa00230 | Purine metabolism | APRT | adenine phosphoribosyltransferase | 353 | down | 1,25 | 1,64E-03 |
| hsa00230 | Purine metabolism | AK1 | adenylate kinase 1 | 203 | down | 1,57 | 9,91E-03 |
| hsa00230 | Purine metabolism | POLD1 | DNA polymerase delta 1, catalytic subunit | 5424 | down | 1,59 | 4,62E-02 |
| hsa00230 | Purine metabolism | POLR2L | RNA polymerase II subunit L | 5441 | down | 1,52 | 3,58E-03 |
| hsa04152 | AMPK signaling pathway | SREBF1 | sterol regulatory element binding transcription factor 1 | 6720 | down | 1,42 | 3,67E-02 |
| hsa04152 | AMPK signaling pathway | FASN | fatty acid synthase | 2194 | down | 1,96 | 8,31E-03 |
| hsa04152 | AMPK signaling pathway | SCD | stearoyl-CoA desaturase | 6319 | down | 1,88 | 3,59E-02 |
| hsa04152 | AMPK signaling pathway | PIK3R1 | phosphoinositide-3-kinase regulatory subunit 1 | 5295 | down | 2,04 | 8,40E-03 |
| hsa04152 | AMPK signaling pathway | RPTOR | regulatory associated protein of MTOR complex 1 | 57521 | down | 1,20 | 1,16E-02 |
| hsa04152 | AMPK signaling pathway | PRKAB2 | protein kinase AMP-activated non-catalytic subunit beta | 5565 | up | 1,83 | 1,27E-02 |
| hsa04152 | AMPK signaling pathway | PPP2R2A | protein phosphatase 2 regulatory subunit Balpha | 5520 | up | 1,26 | 3,61E-02 |
| hsa04152 | AMPK signaling pathway | RPS6KB2 | ribosomal protein S6 kinase B2 | 6199 | down | 1,22 | 1,01E-02 |
| hsa04152 | AMPK signaling pathway | MLYCD | malonyl-CoA decarboxylase | 23417 | down | 1,35 | 2,54E-02 |
| hsa04152 | AMPK signaling pathway | CAMKK1 | calcium/calmodulin dependent protein kinase kinase 1 | 84254 | down | 1,83 | 2,57E-02 |
| hsa04152 | AMPK signaling pathway | STK11 | serine/threonine kinase 11 | 6794 | down | 1,25 | 3,05E-02 |
| hsa04152 | AMPK signaling pathway | PIK3CA | phosphatidylinositol-4,5-bisphosphate 3-kinase catalyti | 5290 | up | 1,33 | 3,17E-02 |
| hsa04152 | AMPK signaling pathway | LEPR | leptin receptor | 3953 | up | 1,33 | 4,94E-03 |
| hsa04152 | AMPK signaling pathway | AKT2 | AKT serine/threonine kinase 2 | 208 | down | 1,28 | 6,33E-03 |
| hsa04152 | AMPK signaling pathway | PFKFB2 | 6-phosphofructo-2-kinase/fructose-2,6-biphosphatase 2 | 5208 | up | 1,39 | 4,32E-02 |
| hsa04152 | AMPK signaling pathway | CREB1 | cAMP responsive element binding protein 1 | 1385 | up | 1,51 | 7,18E-03 |
| hsa04152 | AMPK signaling pathway | GYS1 | glycogen synthase 1 | 2997 | down | 1,32 | 2,63E-02 |
| hsa04152 | AMPK signaling pathway | IRS2 | insulin receptor substrate 2 | 8660 | up | 2,21 | 2,22E-02 |
| hsa04152 | AMPK signaling pathway | CREB5 | cAMP responsive element binding protein 5 | 9586 | up | 3,23 | 2,63E-03 |
| hsa04152 | AMPK signaling pathway | PRKAG2 | protein kinase AMP-activated non-catalytic subunit gan | 51422 | up | 1,28 | 4,57E-02 |
| hsa04152 | AMPK signaling pathway | AKT1 | AKT serine/threonine kinase 1 | 207 | down | 1,37 | 1,42E-03 |
| hsa04152 | AMPK signaling pathway | CAMKK2 | calcium/calmodulin dependent protein kinase kinase 2 | 10645 | down | 1,63 | 2,18E-02 |
| hsa04152 | AMPK signaling pathway | AKT1S1 | AKT1 substrate 1 | 84335 | down | 1,21 | 4,33E-02 |
| hsa04152 | AMPK signaling pathway | PPP2CB | protein phosphatase 2 catalytic subunit beta | 5516 | up | 1,30 | 4,79E-02 |
| hsa04152 | AMPK signaling pathway | IRS1 | insulin receptor substrate 1 | 3667 | down | 1,89 | 6,25E-03 |
| hsa04152 | AMPK signaling pathway | SIRT1 | sirtuin 1 | 23411 | up | 1,60 | 1,00E-02 |
| hsa04152 | AMPK signaling pathway | ACACA | acetyl-CoA carboxylase alpha | 31 | down | 1,40 | 2,54E-02 |
| hsa04152 | AMPK signaling pathway | CREB3L2 | cAMP responsive element binding protein 3 like 2 | 64764 | up | 2,31 | 8,54E-03 |
| hsa04144 | Endocytosis | ARF3 | ADP ribosylation factor 3 | 377 | down | 1,30 | 4,57E-02 |
| hsa04144 | Endocytosis | CHMP5 | charged multivesicular body protein 5 | 51510 | up | 1,32 | 3,37E-02 |
| hsa04144 | Endocytosis | TGFBR2 | transforming growth factor beta receptor 2 | 7048 | down | 1,34 | 2,74E-02 |
| hsa04144 | Endocytosis | STAM | signal transducing adaptor molecule | 8027 | up | 1,23 | 3,52E-02 |
| hsa04144 | Endocytosis | EPS15 | epidermal growth factor receptor pathway substrate 15 | 2060 | up | 1,30 | 7,72E-03 |
| hsa04144 | Endocytosis | VPS26A | VPS26, retromer complex component A | 9559 | up | 1,24 | 4,07E-02 |
| hsa04144 | Endocytosis | RAB22A | RAB22A, member RAS oncogene family | 57403 | up | 1,46 | 3,17E-02 |
| hsa04144 | Endocytosis | FGFR3 | fibroblast growth factor receptor 3 | 2261 | down | 1,70 | 4,60E-02 |
| hsa04144 | Endocytosis | PLD1 | phospholipase D1 | 5337 | up | 1,33 | 1,60E-02 |
| hsa04144 | Endocytosis | PIP5K1C | phosphatidylinositol-4-phosphate 5-kinase type 1 gamm | 23396 | down | 1,39 | 1,49E-03 |
| hsa04144 | Endocytosis | ARRB1 | arrestin beta 1 | 408 | down | 1,79 | 2,42E-02 |
| hsa04144 | Endocytosis | RNF41 | ring finger protein 41 | 10193 | up | 1,59 | 2,87E-02 |
| hsa04144 | Endocytosis | SMURF2 | SMAD specific E3 ubiquitin protein ligase 2 | 64750 | up | 1,49 | 4,16E-02 |
| hsa04144 | Endocytosis | ARAP1 | ArfGAP with RhoGAP domain, ankyrin repeat and PH d | 116985 | down | 1,24 | 4,90E-02 |
| hsa04144 | Endocytosis | VPS37A | VPS37A, ESCRT-I subunit | 137492 | up | 1,48 | 1,14E-02 |
| hsa04144 | Endocytosis | KIF5B | kinesin family member 5B | 3799 | up | 1,40 | 7,18E-03 |
| hsa04144 | Endocytosis | LDLRAP1 | low density lipoprotein receptor adaptor protein 1 | 26119 | down | 1,46 | 2,21E-02 |
| hsa04144 | Endocytosis | ASAP1 | ArfGAP with SH3 domain, ankyrin repeat and PH doma | 50807 | up | 1,27 | 4,95E-02 |
| hsa04144 | Endocytosis | STAM2 | signal transducing adaptor molecule 2 | 10254 | up | 1,41 | 6,99E-03 |
| hsa04144 | Endocytosis | ITCH | itchy E3 ubiquitin protein ligase | 83737 | up | 1,51 | 6,64E-03 |
| hsa04144 | Endocytosis | RAB5A | RAB5A, member RAS oncogene family | 5868 | up | 1,78 | 3,10E-03 |
| hsa04144 | Endocytosis | WASHC4 | WASH complex subunit 4 | 23325 | up | 1,46 | 1,05E-02 |
| hsa04144 | Endocytosis | SH3GLB1 | SH3 domain containing GRB2 like, endophilin B1 | 51100 | up | 1,48 | 1,93E-02 |
| hsa04144 | Endocytosis | SMAD2 | SMAD family member 2 | 4087 | up | 1,29 | 2,19E-02 |
| hsa04144 | Endocytosis | DNAJC6 | DnaJ heat shock protein family (Hsp40) member C6 | 9829 | up | 1,87 | 1,56E-02 |
| hsa04144 | Endocytosis | RAB11FIP3 | RAB11 family interacting protein 3 | 9727 | down | 1,28 | 2,24E-02 |
| hsa04144 | Endocytosis | PARD6A | par-6 family cell polarity regulator alpha | 50855 | down | 1,46 | 8,78E-03 |
| hsa04144 | Endocytosis | AGAP2 | ArfGAP with GTPase domain, ankyrin repeat and PH d | 116986 | down | 2,02 | 3,75E-02 |
| hsa04144 | Endocytosis | HRAS | HRas proto-oncogene, GTPase | 3265 | down | 1,31 | 2,19E-03 |
| hsa04144 | Endocytosis | IQSEC1 | IQ motif and Sec7 domain 1 | 9922 | down | 1,44 | 3,57E-02 |
| hsa04144 | Endocytosis | ARFGAP3 | ADP ribosylation factor GTPase activating protein 3 | 26286 | up | 2,76 | 2,05E-03 |
| hsa04144 | Endocytosis | VPS37D | VPS37D, ESCRT-I subunit | 155382 | up | 2,22 | 5,83E-03 |
| hsa04144 | Endocytosis | USP8 | ubiquitin specific peptidase 8 | 9101 | up | 1,31 | 7,82E-03 |
| hsa04144 | Endocytosis | AGAP3 | ArfGAP with GTPase domain, ankyrin repeat and PH d | 116988 | down | 1,39 | 4,98E-02 |
| hsa04144 | Endocytosis | ARAP3 | ArfGAP with RhoGAP domain, ankyrin repeat and PH d | 64411 | down | 1,96 | 6,61E-03 |
| hsa04144 | Endocytosis | RAB11FIP2 | RAB11 family interacting protein 2 | 22841 | up | 1,50 | 4,23E-02 |
| hsa04144 | Endocytosis | CBLB | Cbl proto-oncogene B | 868 | up | 1,83 | 4,35E-02 |
| hsa04144 | Endocytosis | SMAP1 | small ArfGAP 1 | 60682 | up | 1,33 | 2,16E-02 |
| hsa04144 | Endocytosis | ACAP2 | ArfGAP with coiled-coil, ankyrin repeat and PH domains | 23527 | up | 1,62 | 6,27E-03 |
| hsa04144 | Endocytosis | PIP5K1B | phosphatidylinositol-4-phosphate 5-kinase type 1 beta | 8395 | up | 3,40 | 1,84E-02 |
| hsa04144 | Endocytosis | NEDD4 | neural precursor cell expressed, developmentally down | 4734 | up | 1,99 | 3,68E-03 |
| hsa04068 | FoxO signaling pathway | PIK3R1 | phosphoinositide-3-kinase regulatory subunit 1 | 5295 | down | 2,04 | 8,40E-03 |
| hsa04068 | FoxO signaling pathway | HRAS | HRas proto-oncogene, GTPase | 3265 | down | 1,31 | 2,19E-03 |
| hsa04068 | FoxO signaling pathway | AGAP2 | ArfGAP with GTPase domain, ankyrin repeat and PH d | 116986 | down | 2,02 | 3,75E-02 |
| hsa04068 | FoxO signaling pathway | SOS2 | SOS Ras/Rho guanine nucleotide exchange factor 2 | 6655 | up | 1,82 | 1,41E-02 |
| hsa04068 | FoxO signaling pathway | TGFBR2 | transforming growth factor beta receptor 2 | 7048 | down | 1,34 | 2,74E-02 |
| hsa04068 | FoxO signaling pathway | MAPK12 | mitogen-activated protein kinase 12 | 6300 | down | 1,37 | 3,32E-03 |
| hsa04068 | FoxO signaling pathway | MAPK8 | mitogen-activated protein kinase 8 | 5599 | up | 1,38 | 3,05E-02 |
| hsa04068 | FoxO signaling pathway | SMAD2 | SMAD family member 2 | 4087 | up | 1,29 | 2,19E-02 |
| hsa04068 | FoxO signaling pathway | STK11 | serine/threonine kinase 11 | 6794 | down | 1,25 | 3,05E-02 |
| hsa04068 | FoxO signaling pathway | SKP2 | S-phase kinase associated protein 2 | 6502 | down | 2,21 | 3,69E-02 |
| hsa04068 | FoxO signaling pathway | PIK3CA | phosphatidylinositol-4,5-bisphosphate 3-kinase catalyti | 5290 | up | 1,33 | 3,17E-02 |
| hsa04068 | FoxO signaling pathway | AKT2 | AKT serine/threonine kinase 2 | 208 | down | 1,28 | 6,33E-03 |
| hsa04068 | FoxO signaling pathway | BRAF | B-Raf proto-oncogene, serine/threonine kinase | 673 | up | 1,52 | 4,53E-02 |
| hsa04068 | FoxO signaling pathway | PRKAB2 | protein kinase AMP-activated non-catalytic subunit beta | 5565 | up | 1,83 | 1,27E-02 |
| hsa04068 | FoxO signaling pathway | PLK3 | polo like kinase 3 | 1263 | up | 2,42 | 3,60E-03 |
| hsa04068 | FoxO signaling pathway | GADD45B | growth arrest and DNA damage inducible beta | 4616 | up | 3,21 | 3,92E-04 |
| hsa04068 | FoxO signaling pathway | CDKN1A | cyclin dependent kinase inhibitor 1A | 1026 | up | 3,62 | 5,43E-03 |
| hsa04068 | FoxO signaling pathway | CDKN2B | cyclin dependent kinase inhibitor 2B | 1030 | up | 2,40 | 3,90E-02 |
| hsa04068 | FoxO signaling pathway | PRKAG2 | protein kinase AMP-activated non-catalytic subunit gan | 51422 | up | 1,28 | 4,57E-02 |
| hsa04068 | FoxO signaling pathway | AKT1 | AKT serine/threonine kinase 1 | 207 | down | 1,37 | 1,42E-03 |
| hsa04068 | FoxO signaling pathway | GADD45A | growth arrest and DNA damage inducible alpha | 1647 | up | 3,05 | 4,75E-04 |
| hsa04068 | FoxO signaling pathway | IRS2 | insulin receptor substrate 2 | 8660 | up | 2,21 | 2,22E-02 |
| hsa04068 | FoxO signaling pathway | PTEN | phosphatase and tensin homolog | 5728 | up | 1,33 | 4,72E-02 |
| hsa04068 | FoxO signaling pathway | KLF2 | Kruppel like factor 2 | 10365 | up | 2,02 | 4,16E-02 |
| hsa04068 | FoxO signaling pathway | ATG12 | autophagy related 12 | 9140 | up | 1,35 | 4,10E-02 |
| hsa04068 | FoxO signaling pathway | IRS1 | insulin receptor substrate 1 | 3667 | down | 1,89 | 6,25E-03 |
| hsa04068 | FoxO signaling pathway | SIRT1 | sirtuin 1 | 23411 | up | 1,60 | 1,00E-02 |
| hsa04068 | FoxO signaling pathway | GABARAPL1 | GABA type A receptor associated protein like 1 | 23710 | up | 2,62 | 4,12E-03 |
| hsa04722 | Neurotrophin signaling pathway | CALM3 | calmodulin 3 | 808 | down | 1,43 | 2,23E-02 |
| hsa04722 | Neurotrophin signaling pathway | BDNF | brain derived neurotrophic factor | 627 | down | 2,98 | 1,15E-02 |
| hsa04722 | Neurotrophin signaling pathway | TP73 | tumor protein p73 | 7161 | down | 2,86 | 3,40E-02 |
| hsa04722 | Neurotrophin signaling pathway | RIPK2 | receptor interacting serine/threonine kinase 2 | 8767 | up | 1,72 | 8,31E-03 |
| hsa04722 | Neurotrophin signaling pathway | PRDM4 | PR/SET domain 4 | 11108 | up | 1,52 | 2,92E-02 |
| hsa04722 | Neurotrophin signaling pathway | AKT1 | AKT serine/threonine kinase 1 | 207 | down | 1,37 | 1,42E-03 |
| hsa04722 | Neurotrophin signaling pathway | CALM2 | calmodulin 2 | 805 | down | 1,43 | 2,23E-02 |
| hsa04722 | Neurotrophin signaling pathway | BAX | BCL2 associated X, apoptosis regulator | 581 | down | 1,22 | 2,61E-02 |
| hsa04722 | Neurotrophin signaling pathway | NGFR | nerve growth factor receptor | 4804 | up | 12,35 | 2,75E-03 |
| hsa04722 | Neurotrophin signaling pathway | IRS1 | insulin receptor substrate 1 | 3667 | down | 1,89 | 6,25E-03 |
| hsa04722 | Neurotrophin signaling pathway | SH2B2 | SH2B adaptor protein 2 | 10603 | down | 2,07 | 1,95E-02 |
| hsa04722 | Neurotrophin signaling pathway | CAMK2D | calcium/calmodulin dependent protein kinase II delta | 817 | up | 1,29 | 1,49E-02 |

| | | | | | | | | |
|---|---|---|---|---|---|---|---|---|
| hsa04722 | Neurotrophin signaling pathway | CALM1 | calmodulin 1 | 801 | down | 1,43 | 2,23E-02 | |
| hsa04722 | Neurotrophin signaling pathway | HRAS | HRas proto-oncogene, GTPase | 3265 | down | 1,31 | 2,19E-03 | |
| hsa04722 | Neurotrophin signaling pathway | PIK3R1 | phosphoinositide-3-kinase regulatory subunit 1 | 5295 | down | 2,04 | 8,40E-03 | |
| hsa04722 | Neurotrophin signaling pathway | ZNF274 | zinc finger protein 274 | 10782 | up | 11,88 | 2,26E-03 | |
| hsa04722 | Neurotrophin signaling pathway | MAPK8 | mitogen-activated protein kinase 8 | 5599 | up | 1,38 | 3,05E-02 | |
| hsa04722 | Neurotrophin signaling pathway | RPS6KA5 | ribosomal protein S6 kinase A5 | 9252 | up | 2,11 | 5,94E-03 | |
| hsa04722 | Neurotrophin signaling pathway | MAPK12 | mitogen-activated protein kinase 12 | 6300 | down | 1,37 | 3,32E-03 | |
| hsa04722 | Neurotrophin signaling pathway | SOS2 | SOS Ras/Rho guanine nucleotide exchange factor 2 | 6655 | up | 1,82 | 1,41E-02 | |
| hsa04722 | Neurotrophin signaling pathway | KIDINS220 | kinase D interacting substrate 220 | 57498 | up | 1,34 | 3,94E-02 | |
| hsa04722 | Neurotrophin signaling pathway | AKT2 | AKT serine/threonine kinase 2 | 208 | down | 1,28 | 6,33E-03 | |
| hsa04722 | Neurotrophin signaling pathway | PIK3CA | phosphatidylinositol-4,5-bisphosphate 3-kinase catalyti | 5290 | up | 1,33 | 3,17E-02 | |
| hsa04722 | Neurotrophin signaling pathway | BRAF | B-Raf proto-oncogene, serine/threonine kinase | 673 | up | 1,52 | 4,53E-02 | |
| hsa04722 | Neurotrophin signaling pathway | PLCG1 | phospholipase C gamma 1 | 5335 | down | 1,51 | 2,85E-02 | |
| hsa04722 | Neurotrophin signaling pathway | SH2B1 | SH2B adaptor protein 1 | 25970 | down | 1,31 | 1,81E-02 | |
| hsa04910 | Insulin signaling pathway | AKT2 | AKT serine/threonine kinase 2 | 208 | down | 1,28 | 6,33E-03 | |
| hsa04910 | Insulin signaling pathway | PIK3CA | phosphatidylinositol-4,5-bisphosphate 3-kinase catalyti | 5290 | up | 1,33 | 3,17E-02 | |
| hsa04910 | Insulin signaling pathway | RPTOR | regulatory associated protein of MTOR complex 1 | 57521 | down | 1,20 | 1,16E-02 | |
| hsa04910 | Insulin signaling pathway | PRKAB2 | protein kinase AMP-activated non-catalytic subunit bet | 5565 | up | 1,83 | 1,27E-02 | |
| hsa04910 | Insulin signaling pathway | RPS6KB2 | ribosomal protein S6 kinase B2 | 6199 | down | 1,22 | 1,01E-02 | |
| hsa04910 | Insulin signaling pathway | PPP1CA | protein phosphatase 1 catalytic subunit alpha | 5499 | down | 1,24 | 9,77E-03 | |
| hsa04910 | Insulin signaling pathway | BRAF | B-Raf proto-oncogene, serine/threonine kinase | 673 | up | 1,52 | 4,53E-02 | |
| hsa04910 | Insulin signaling pathway | HRAS | HRas proto-oncogene, GTPase | 3265 | down | 1,31 | 2,19E-03 | |
| hsa04910 | Insulin signaling pathway | CALM1 | calmodulin 1 | 801 | down | 1,43 | 2,23E-02 | |
| hsa04910 | Insulin signaling pathway | PIK3R1 | phosphoinositide-3-kinase regulatory subunit 1 | 5295 | down | 2,04 | 8,40E-03 | |
| hsa04910 | Insulin signaling pathway | SREBF1 | sterol regulatory element binding transcription factor 1 | 6720 | down | 1,42 | 3,67E-02 | |
| hsa04910 | Insulin signaling pathway | PRKAR1B | protein kinase cAMP-dependent type I regulatory subu | 5575 | down | 1,50 | 6,80E-04 | |
| hsa04910 | Insulin signaling pathway | MAPK8 | mitogen-activated protein kinase 8 | 5599 | up | 1,38 | 3,05E-02 | |
| hsa04910 | Insulin signaling pathway | FASN | fatty acid synthase | 2194 | down | 1,96 | 8,31E-03 | |
| hsa04910 | Insulin signaling pathway | SOS2 | SOS Ras/Rho guanine nucleotide exchange factor 2 | 6655 | up | 1,82 | 1,41E-02 | |
| hsa04910 | Insulin signaling pathway | CALM2 | calmodulin 2 | 805 | down | 1,43 | 2,23E-02 | |
| hsa04910 | Insulin signaling pathway | IRS1 | insulin receptor substrate 1 | 3667 | down | 1,89 | 6,25E-03 | |
| hsa04910 | Insulin signaling pathway | HKDC1 | hexokinase domain containing 1 | 80201 | up | 3,37 | 4,73E-03 | |
| hsa04910 | Insulin signaling pathway | ACACA | acetyl-CoA carboxylase alpha | 31 | down | 1,40 | 2,54E-02 | |
| hsa04910 | Insulin signaling pathway | CBLB | Cbl proto-oncogene B | 868 | up | 1,83 | 4,35E-02 | |
| hsa04910 | Insulin signaling pathway | SH2B2 | SH2B adaptor protein 2 | 10603 | down | 2,07 | 1,95E-02 | |
| hsa04910 | Insulin signaling pathway | PPP1R3B | protein phosphatase 1 regulatory subunit 3B | 79660 | up | 1,41 | 4,66E-02 | |
| hsa04910 | Insulin signaling pathway | CALM3 | calmodulin 3 | 808 | down | 1,43 | 2,23E-02 | |
| hsa04910 | Insulin signaling pathway | FLOT2 | flotillin 2 | 2319 | down | 1,45 | 3,40E-02 | |
| hsa04910 | Insulin signaling pathway | GYS1 | glycogen synthase 1 | 2997 | down | 1,32 | 2,63E-02 | |
| hsa04910 | Insulin signaling pathway | IRS2 | insulin receptor substrate 2 | 8660 | up | 2,21 | 2,22E-02 | |
| hsa04910 | Insulin signaling pathway | PRKAG2 | protein kinase AMP-activated non-catalytic subunit gar | 51422 | up | 1,28 | 4,57E-02 | |
| hsa04910 | Insulin signaling pathway | AKT1 | AKT serine/threonine kinase 1 | 207 | down | 1,37 | 1,42E-03 | |
| hsa03410 | Base excision repair | UNG | uracil DNA glycosylase | 7374 | down | 2,13 | 1,95E-02 | |
| hsa03410 | Base excision repair | POLB | DNA polymerase beta | 5423 | up | 1,25 | 3,75E-02 | |
| hsa03410 | Base excision repair | PARP1 | poly(ADP-ribose) polymerase 1 | 142 | down | 1,50 | 3,92E-02 | |
| hsa03410 | Base excision repair | POLD2 | DNA polymerase delta 2, accessory subunit | 5425 | down | 1,30 | 1,92E-02 | |
| hsa03410 | Base excision repair | APEX2 | apurinic/apyrimidinic endodeoxyribonuclease 2 | 27301 | down | 1,25 | 2,34E-02 | |
| hsa03410 | Base excision repair | PCNA | proliferating cell nuclear antigen | 5111 | down | 2,14 | 4,63E-02 | PCNA down |
| hsa03410 | Base excision repair | POLD1 | DNA polymerase delta 1, catalytic subunit | 5424 | down | 1,59 | 4,62E-02 | |
| hsa03410 | Base excision repair | POLE | DNA polymerase epsilon, catalytic subunit | 5426 | down | 1,55 | 2,67E-02 | |
| hsa03410 | Base excision repair | FEN1 | flap structure-specific endonuclease 1 | 2237 | down | 2,04 | 1,60E-02 | |
| hsa04962 | Vasopressin-regulated water reabsorption | RAB5A | RAB5A, member RAS oncogene family | 5868 | up | 1,78 | 3,10E-03 | |
| hsa04962 | Vasopressin-regulated water reabsorption | DYNC2H1 | dynein cytoplasmic 2 heavy chain 1 | 79659 | up | 1,52 | 9,51E-03 | |
| hsa04962 | Vasopressin-regulated water reabsorption | DYNC2LI1 | dynein cytoplasmic 2 light intermediate chain 1 | 51626 | up | 1,40 | 3,09E-02 | |
| hsa04962 | Vasopressin-regulated water reabsorption | CREB3L2 | cAMP responsive element binding protein 3 like 2 | 64764 | up | 2,31 | 8,54E-03 | |
| hsa04962 | Vasopressin-regulated water reabsorption | DCTN4 | dynactin subunit 4 | 51164 | up | 1,51 | 1,31E-02 | |
| hsa04962 | Vasopressin-regulated water reabsorption | VAMP2 | vesicle associated membrane protein 2 | 6844 | up | 1,50 | 4,82E-02 | |
| hsa04962 | Vasopressin-regulated water reabsorption | ADCY3 | adenylate cyclase 3 | 109 | down | 1,45 | 1,03E-02 | |
| hsa04962 | Vasopressin-regulated water reabsorption | DYNC1LI2 | dynein cytoplasmic 1 light intermediate chain 2 | 1783 | up | 1,21 | 2,47E-02 | |
| hsa04962 | Vasopressin-regulated water reabsorption | CREB1 | cAMP responsive element binding protein 1 | 1385 | up | 1,51 | 7,18E-03 | |
| hsa04962 | Vasopressin-regulated water reabsorption | CREB5 | cAMP responsive element binding protein 5 | 9586 | up | 3,23 | 2,63E-03 | |
| hsa04668 | TNF signaling pathway | TRAF2 | TNF receptor associated factor 2 | 7186 | down | 1,47 | 2,65E-02 | |
| hsa04668 | TNF signaling pathway | ATF2 | activating transcription factor 2 | 1386 | up | 1,79 | 1,01E-02 | |
| hsa04668 | TNF signaling pathway | RPS6KA4 | ribosomal protein S6 kinase A4 | 8986 | down | 1,33 | 5,67E-03 | |
| hsa04668 | TNF signaling pathway | PIK3CA | phosphatidylinositol-4,5-bisphosphate 3-kinase catalyti | 5290 | up | 1,33 | 3,17E-02 | |
| hsa04668 | TNF signaling pathway | AKT2 | AKT serine/threonine kinase 2 | 208 | down | 1,28 | 6,33E-03 | |
| hsa04668 | TNF signaling pathway | MAPK8 | mitogen-activated protein kinase 8 | 5599 | up | 1,38 | 3,05E-02 | |
| hsa04668 | TNF signaling pathway | MAPK12 | mitogen-activated protein kinase 12 | 6300 | down | 1,37 | 3,32E-03 | |
| hsa04668 | TNF signaling pathway | RPS6KA5 | ribosomal protein S6 kinase A5 | 9252 | up | 2,11 | 5,94E-03 | |
| hsa04668 | TNF signaling pathway | PGAM5 | PGAM family member 5, mitochondrial serine/threonine | 192111 | down | 1,31 | 3,68E-02 | |
| hsa04668 | TNF signaling pathway | JAG1 | jagged 1 | 182 | up | 1,84 | 3,27E-02 | |
| hsa04668 | TNF signaling pathway | TAB3 | TGF-beta activated kinase 1 and MAP3K7 binding prot | 257397 | up | 1,40 | 4,94E-02 | |
| hsa04668 | TNF signaling pathway | PIK3R1 | phosphoinositide-3-kinase regulatory subunit 1 | 5295 | down | 2,04 | 8,40E-03 | |
| hsa04668 | TNF signaling pathway | BIRC2 | baculoviral IAP repeat containing 2 | 329 | up | 1,38 | 1,86E-02 | |
| hsa04668 | TNF signaling pathway | CEBPB | CCAAT/enhancer binding protein beta | 1051 | up | 2,12 | 2,85E-02 | |
| hsa04668 | TNF signaling pathway | CREB3L2 | cAMP responsive element binding protein 3 like 2 | 64764 | up | 2,31 | 8,54E-03 | |
| hsa04668 | TNF signaling pathway | ITCH | itchy E3 ubiquitin protein ligase | 83737 | up | 1,51 | 6,64E-03 | |
| hsa04668 | TNF signaling pathway | CREB1 | cAMP responsive element binding protein 1 | 1385 | up | 1,51 | 7,18E-03 | |
| hsa04668 | TNF signaling pathway | TAB1 | TGF-beta activated kinase 1 (MAP3K7) binding protein | 10454 | down | 1,29 | 1,94E-02 | |
| hsa04668 | TNF signaling pathway | CREB5 | cAMP responsive element binding protein 5 | 9586 | up | 3,23 | 2,63E-03 | |
| hsa04668 | TNF signaling pathway | AKT1 | AKT serine/threonine kinase 1 | 207 | down | 1,37 | 1,42E-03 | |
| hsa04668 | TNF signaling pathway | TRAF3 | TNF receptor associated factor 3 | 7187 | down | 1,40 | 1,36E-02 | |
| hsa04668 | TNF signaling pathway | FADD | Fas associated via death domain | 8772 | down | 1,31 | 8,90E-03 | |
| hsa04931 | Insulin resistance | PIK3R1 | phosphoinositide-3-kinase regulatory subunit 1 | 5295 | down | 2,04 | 8,40E-03 | |
| hsa04931 | Insulin resistance | GFPT1 | glutamine--fructose-6-phosphate transaminase 1 | 2673 | up | 2,92 | 2,14E-03 | |
| hsa04931 | Insulin resistance | SREBF1 | sterol regulatory element binding transcription factor 1 | 6720 | down | 1,42 | 3,67E-02 | |
| hsa04931 | Insulin resistance | MAPK8 | mitogen-activated protein kinase 8 | 5599 | up | 1,38 | 3,05E-02 | |
| hsa04931 | Insulin resistance | PIK3CA | phosphatidylinositol-4,5-bisphosphate 3-kinase catalyti | 5290 | up | 1,33 | 3,17E-02 | |
| hsa04931 | Insulin resistance | AKT2 | AKT serine/threonine kinase 2 | 208 | down | 1,28 | 6,33E-03 | |
| hsa04931 | Insulin resistance | SLC27A4 | solute carrier family 27 member 4 | 10999 | down | 1,59 | 7,26E-03 | |
| hsa04931 | Insulin resistance | PPP1CA | protein phosphatase 1 catalytic subunit alpha | 5499 | down | 1,24 | 9,77E-03 | |
| hsa04931 | Insulin resistance | RPS6KB2 | ribosomal protein S6 kinase B2 | 6199 | down | 1,22 | 1,01E-02 | |
| hsa04931 | Insulin resistance | PRKAB2 | protein kinase AMP-activated non-catalytic subunit bet | 5565 | up | 1,83 | 1,27E-02 | |
| hsa04931 | Insulin resistance | PPP1R3B | protein phosphatase 1 regulatory subunit 3B | 79660 | up | 1,41 | 4,66E-02 | |
| hsa04931 | Insulin resistance | AKT1 | AKT serine/threonine kinase 1 | 207 | down | 1,37 | 1,42E-03 | |
| hsa04931 | Insulin resistance | CREB5 | cAMP responsive element binding protein 5 | 9586 | up | 3,23 | 2,63E-03 | |
| hsa04931 | Insulin resistance | PRKAG2 | protein kinase AMP-activated non-catalytic subunit gar | 51422 | up | 1,28 | 4,57E-02 | |
| hsa04931 | Insulin resistance | TRIB3 | tribbles pseudokinase 3 | 57761 | up | 3,17 | 8,07E-03 | |
| hsa04931 | Insulin resistance | PTPA | protein phosphatase 2 phosphatase activator | 5524 | down | 1,34 | 2,58E-02 | |
| hsa04931 | Insulin resistance | GYS1 | glycogen synthase 1 | 2997 | down | 1,32 | 2,63E-02 | |
| hsa04931 | Insulin resistance | IRS2 | insulin receptor substrate 2 | 8660 | up | 2,21 | 2,22E-02 | |
| hsa04931 | Insulin resistance | CREB1 | cAMP responsive element binding protein 1 | 1385 | up | 1,51 | 7,18E-03 | |
| hsa04931 | Insulin resistance | CREB3L2 | cAMP responsive element binding protein 3 like 2 | 64764 | up | 2,31 | 8,54E-03 | |
| hsa04931 | Insulin resistance | PTEN | phosphatase and tensin homolog | 5728 | up | 1,33 | 4,72E-02 | |
| hsa04931 | Insulin resistance | IRS1 | insulin receptor substrate 1 | 3667 | down | 1,89 | 6,25E-03 | |
| hsa04015 | Rap1 signaling pathway | PIK3R1 | phosphoinositide-3-kinase regulatory subunit 1 | 5295 | down | 2,04 | 8,40E-03 | |
| hsa04015 | Rap1 signaling pathway | FGF2 | fibroblast growth factor 2 | 2247 | down | 5,35 | 3,09E-02 | |
| hsa04015 | Rap1 signaling pathway | HRAS | HRas proto-oncogene, GTPase | 3265 | down | 1,31 | 2,19E-03 | |
| hsa04015 | Rap1 signaling pathway | CALM1 | calmodulin 1 | 801 | down | 1,43 | 2,23E-02 | |
| hsa04015 | Rap1 signaling pathway | PARD6A | par-6 family cell polarity regulator alpha | 50855 | down | 1,46 | 8,78E-03 | |
| hsa04015 | Rap1 signaling pathway | EFNA4 | ephrin A4 | 1945 | down | 1,43 | 4,92E-02 | |
| hsa04015 | Rap1 signaling pathway | PFN1 | profilin 1 | 5216 | down | 1,20 | 3,98E-02 | |
| hsa04015 | Rap1 signaling pathway | RGS14 | regulator of G protein signaling 14 | 10636 | down | 1,61 | 4,72E-02 | |
| hsa04015 | Rap1 signaling pathway | MAPK12 | mitogen-activated protein kinase 12 | 6300 | down | 1,37 | 3,32E-03 | |
| hsa04015 | Rap1 signaling pathway | RALGDS | ral guanine nucleotide dissociation stimulator | 5900 | down | 1,30 | 3,49E-02 | |
| hsa04015 | Rap1 signaling pathway | EFNA3 | ephrin A3 | 1944 | down | 2,00 | 4,89E-02 | |
| hsa04015 | Rap1 signaling pathway | AKT2 | AKT serine/threonine kinase 2 | 208 | down | 1,28 | 6,33E-03 | |
| hsa04015 | Rap1 signaling pathway | PLCG1 | phospholipase C gamma 1 | 5335 | down | 1,51 | 2,85E-02 | |
| hsa04015 | Rap1 signaling pathway | RASSF5 | Ras association domain family member 5 | 83593 | down | 2,36 | 2,58E-02 | |
| hsa04015 | Rap1 signaling pathway | FGFR3 | fibroblast growth factor receptor 3 | 2261 | down | 1,70 | 4,60E-02 | |
| hsa04015 | Rap1 signaling pathway | ARAP3 | ArfGAP with RhoGAP domain, ankyrin repeat and PH d | 64411 | down | 1,96 | 6,61E-03 | |
| hsa04015 | Rap1 signaling pathway | CALM3 | calmodulin 3 | 808 | down | 1,43 | 2,23E-02 | |
| hsa04015 | Rap1 signaling pathway | GNAI2 | G protein subunit alpha i2 | 2771 | down | 1,22 | 3,65E-02 | |
| hsa04015 | Rap1 signaling pathway | AKT1 | AKT serine/threonine kinase 1 | 207 | down | 1,37 | 1,42E-03 | |
| hsa04015 | Rap1 signaling pathway | F2R | coagulation factor II thrombin receptor | 2149 | down | 1,54 | 7,69E-03 | |
| hsa04015 | Rap1 signaling pathway | CALM2 | calmodulin 2 | 805 | down | 1,43 | 2,23E-02 | |
| hsa04015 | Rap1 signaling pathway | ADCY3 | adenylate cyclase 3 | 109 | down | 1,45 | 1,03E-02 | |
| hsa01130 | Biosynthesis of antibiotics | CTH | cystathionine gamma-lyase | 1491 | up | 3,41 | 4,94E-03 | |
| hsa01130 | Biosynthesis of antibiotics | PGM3 | phosphoglucomutase 3 | 5238 | up | 2,22 | 4,77E-03 | |
| hsa01130 | Biosynthesis of antibiotics | TGDS | TDP-glucose 4,6-dehydratase | 23483 | up | 1,72 | 2,20E-02 | |
| hsa01130 | Biosynthesis of antibiotics | LSS | lanosterol synthase | 4047 | down | 1,49 | 1,65E-03 | |
| hsa01130 | Biosynthesis of antibiotics | GFPT1 | glutamine--fructose-6-phosphate transaminase 1 | 2673 | up | 2,92 | 2,14E-03 | |
| hsa01130 | Biosynthesis of antibiotics | HADHB | hydroxyacyl-CoA dehydrogenase trifunctional multienzy | 3032 | up | 1,20 | 4,66E-02 | |
| hsa01130 | Biosynthesis of antibiotics | ACAT2 | acetyl-CoA acetyltransferase 2 | 39 | down | 1,69 | 2,52E-02 | |
| hsa01130 | Biosynthesis of antibiotics | MVK | mevalonate kinase | 4598 | down | 1,99 | 3,36E-03 | |
| hsa01130 | Biosynthesis of antibiotics | FNTA | farnesyltransferase, CAAX box, alpha | 2339 | up | 1,33 | 3,73E-03 | |
| hsa01130 | Biosynthesis of antibiotics | SQLE | squalene epoxidase | 6713 | down | 2,05 | 1,00E-02 | |
| hsa01130 | Biosynthesis of antibiotics | DLD | dihydrolipoamide dehydrogenase | 1738 | up | 1,16 | 4,53E-02 | |
| hsa01130 | Biosynthesis of antibiotics | GOT1 | glutamic-oxaloacetic transaminase 1 | 2805 | up | 2,30 | 5,67E-03 | |
| hsa01130 | Biosynthesis of antibiotics | PFAS | phosphoribosylformylglycinamidine synthase | 5198 | down | 1,53 | 1,69E-02 | |
| hsa01130 | Biosynthesis of antibiotics | UGP2 | UDP-glucose pyrophosphorylase 2 | 7360 | up | 1,37 | 3,47E-02 | |
| hsa01130 | Biosynthesis of antibiotics | ISYNA1 | inositol-3-phosphate synthase 1 | 51477 | down | 1,54 | 7,64E-03 | |
| hsa01130 | Biosynthesis of antibiotics | TM7SF2 | transmembrane 7 superfamily member 2 | 7108 | down | 1,51 | 4,00E-02 | |
| hsa01130 | Biosynthesis of antibiotics | PGP | phosphoglycolate phosphatase | 283871 | down | 1,50 | 8,04E-03 | |
| hsa01130 | Biosynthesis of antibiotics | PKM | pyruvate kinase M1/2 | 5315 | down | 1,35 | 4,16E-02 | |
| hsa01130 | Biosynthesis of antibiotics | PGAM1 | phosphoglycerate mutase 1 | 5223 | down | 1,62 | 5,67E-03 | |
| hsa01130 | Biosynthesis of antibiotics | NSDHL | NAD(P) dependent steroid dehydrogenase-like | 50814 | down | 1,36 | 1,99E-02 | |
| hsa01130 | Biosynthesis of antibiotics | ARG2 | arginase 2 | 384 | up | 7,09 | 3,92E-04 | |

| | | | | | | | |
|---|---|---|---|---|---|---|---|
| hsa01130 | Biosynthesis of antibiotics | PYCR3 | pyrroline-5-carboxylate reductase 3 | 65263 | down | 1,77 | 1,11E-03 |
| hsa01130 | Biosynthesis of antibiotics | ALDH1B1 | aldehyde dehydrogenase 1 family member B1 | 219 | down | 1,92 | 9,37E-03 |
| hsa01130 | Biosynthesis of antibiotics | ICMT | isoprenylcysteine carboxyl methyltransferase | 23463 | down | 1,63 | 2,39E-02 |
| hsa01130 | Biosynthesis of antibiotics | AK1 | adenylate kinase 1 | 203 | down | 1,57 | 9,91E-03 |
| hsa01130 | Biosynthesis of antibiotics | GPI | glucose-6-phosphate isomerase | 2821 | down | 1,24 | 1,55E-02 |
| hsa01130 | Biosynthesis of antibiotics | TPI1 | triosephosphate isomerase 1 | 7167 | down | 1,28 | 3,58E-02 |
| hsa01130 | Biosynthesis of antibiotics | ACLY | ATP citrate lyase | 47 | down | 1,41 | 3,09E-02 |
| hsa01130 | Biosynthesis of antibiotics | AASS | aminoadipate-semialdehyde synthase | 10157 | up | 2,01 | 4,95E-02 |
| hsa01130 | Biosynthesis of antibiotics | PCCA | propionyl-CoA carboxylase alpha subunit | 5095 | up | 1,37 | 2,03E-02 |
| hsa01130 | Biosynthesis of antibiotics | BCAT2 | branched chain amino acid transaminase 2 | 587 | down | 1,34 | 1,96E-02 |
| hsa01130 | Biosynthesis of antibiotics | PHGDH | phosphoglycerate dehydrogenase | 26227 | down | 2,34 | 2,54E-02 |
| hsa01130 | Biosynthesis of antibiotics | HKDC1 | hexokinase domain containing 1 | 80201 | up | 3,37 | 4,73E-03 |
| hsa01130 | Biosynthesis of antibiotics | AK4 | adenylate kinase 4 | 205 | down | 1,76 | 4,87E-02 |
| hsa01130 | Biosynthesis of antibiotics | NME4 | NME/NM23 nucleoside diphosphate kinase 4 | 4833 | down | 1,47 | 3,85E-02 |
| hsa00564 | Glycerophospholipid metabolism | ADPRM | ADP-ribose/CDP-alcohol diphosphatase, manganese d | 56985 | up | 2,23 | 2,72E-03 |
| hsa00564 | Glycerophospholipid metabolism | MBOAT7 | membrane bound O-acyltransferase domain containing | 79143 | down | 1,47 | 9,36E-03 |
| hsa00564 | Glycerophospholipid metabolism | PISD | phosphatidylserine decarboxylase | 23761 | down | 1,20 | 2,48E-02 |
| hsa00564 | Glycerophospholipid metabolism | GPCPD1 | glycerophosphocholine phosphodiesterase 1 | 56261 | up | 2,75 | 2,24E-03 |
| hsa00564 | Glycerophospholipid metabolism | LPIN2 | lipin 2 | 9663 | up | 2,16 | 6,88E-03 |
| hsa00564 | Glycerophospholipid metabolism | GPAT3 | glycerol-3-phosphate acyltransferase 3 | 84803 | up | 1,77 | 4,99E-02 |
| hsa00564 | Glycerophospholipid metabolism | LPCAT1 | lysophosphatidylcholine acyltransferase 1 | 79888 | down | 1,76 | 2,60E-02 |
| hsa00564 | Glycerophospholipid metabolism | LYPLA2 | lysophospholipase II | 11313 | down | 1,49 | 2,45E-02 |
| hsa00564 | Glycerophospholipid metabolism | AGPAT2 | 1-acylglycerol-3-phosphate O-acyltransferase 2 | 10555 | down | 1,23 | 2,89E-03 |
| hsa00564 | Glycerophospholipid metabolism | AGPAT5 | 1-acylglycerol-3-phosphate O-acyltransferase 5 | 55326 | down | 1,30 | 4,70E-02 |
| hsa00564 | Glycerophospholipid metabolism | PLD1 | phospholipase D1 | 5337 | up | 1,33 | 1,60E-02 |
| hsa00564 | Glycerophospholipid metabolism | LPCAT4 | lysophosphatidylcholine acyltransferase 4 | 254531 | down | 1,78 | 2,52E-02 |
| hsa00564 | Glycerophospholipid metabolism | PTDSS2 | phosphatidylserine synthase 2 | 81490 | down | 1,30 | 1,46E-02 |
| hsa00564 | Glycerophospholipid metabolism | LPIN3 | lipin 3 | 64900 | down | 1,36 | 4,76E-02 |
| hsa00564 | Glycerophospholipid metabolism | CDIPT | CDP-diacylglycerol--inositol 3-phosphatidyltransferase | 10423 | down | 1,32 | 3,12E-03 |
| hsa00564 | Glycerophospholipid metabolism | AGPAT3 | 1-acylglycerol-3-phosphate O-acyltransferase 3 | 56894 | down | 1,30 | 9,80E-03 |
| hsa00564 | Glycerophospholipid metabolism | GPAT4 | glycerol-3-phosphate acyltransferase 4 | 137964 | down | 1,24 | 4,75E-02 |
| hsa00564 | Glycerophospholipid metabolism | PTDSS1 | phosphatidylserine synthase 1 | 9791 | down | 1,31 | 3,90E-02 |
| hsa03020 | RNA polymerase | POLR1C | RNA polymerase I subunit C | 9533 | down | 1,24 | 4,71E-02 |
| hsa03020 | RNA polymerase | POLR3C | RNA polymerase III subunit C | 10623 | up | 1,43 | 4,98E-02 |
| hsa03020 | RNA polymerase | POLR3D | RNA polymerase III subunit D | 661 | up | 1,52 | 2,72E-02 |
| hsa03020 | RNA polymerase | POLR1A | RNA polymerase I subunit A | 25885 | down | 1,30 | 4,32E-02 |
| hsa03020 | RNA polymerase | POLR3H | RNA polymerase III subunit H | 171568 | down | 1,80 | 7,07E-04 |
| hsa03020 | RNA polymerase | POLR2E | RNA polymerase II subunit E | 5434 | down | 1,31 | 3,18E-03 |
| hsa03020 | RNA polymerase | POLR2B | RNA polymerase II subunit B | 5431 | down | 1,23 | 3,81E-02 |
| hsa03020 | RNA polymerase | POLR2L | RNA polymerase II subunit L | 5441 | down | 1,52 | 3,58E-03 |
| hsa03020 | RNA polymerase | POLR2F | RNA polymerase II subunit F | 5435 | down | 1,41 | 3,22E-02 |
| hsa03020 | RNA polymerase | POLR3F | RNA polymerase III subunit F | 10621 | up | 1,25 | 3,24E-02 |
| hsa03060 | Protein export | SRP19 | signal recognition particle 19 | 6728 | up | 1,68 | 4,13E-02 |
| hsa03060 | Protein export | HSPA5 | heat shock protein family A (Hsp70) member 5 | 3309 | up | 5,02 | 6,98E-04 |
| hsa03060 | Protein export | SRP72 | signal recognition particle 72 | 6731 | up | 1,43 | 9,37E-03 |
| hsa03060 | Protein export | SEC63 | SEC63 homolog, protein translocation regulator | 11231 | up | 1,60 | 8,90E-03 |
| hsa03060 | Protein export | IMMP2L | inner mitochondrial membrane peptidase subunit 2 | 83943 | up | 1,42 | 2,85E-02 |
| hsa03060 | Protein export | SRP68 | signal recognition particle 68 | 6730 | up | 1,40 | 3,02E-02 |
| hsa05214 | Glioma | PIK3R1 | phosphoinositide-3-kinase regulatory subunit 1 | 5295 | down | 2,04 | 8,40E-03 |
| hsa05214 | Glioma | HRAS | HRas proto-oncogene, GTPase | 3265 | down | 1,31 | 2,19E-03 |
| hsa05214 | Glioma | CALM1 | calmodulin 1 | 801 | down | 1,43 | 2,23E-02 |
| hsa05214 | Glioma | CAMK2D | calcium/calmodulin dependent protein kinase II delta | 817 | up | 1,29 | 1,49E-02 |
| hsa05214 | Glioma | CDKN1A | cyclin dependent kinase inhibitor 1A | 1026 | up | 3,62 | 5,43E-03 |
| hsa05214 | Glioma | CALM3 | calmodulin 3 | 808 | down | 1,43 | 2,23E-02 |
| hsa05214 | Glioma | SOS2 | SOS Ras/Rho guanine nucleotide exchange factor 2 | 6655 | up | 1,82 | 1,41E-02 |
| hsa05214 | Glioma | AKT1 | AKT serine/threonine kinase 1 | 207 | down | 1,37 | 1,42E-03 |
| hsa05214 | Glioma | E2F1 | E2F transcription factor 1 | 1869 | down | 1,64 | 3,25E-02 |
| hsa05214 | Glioma | PIK3CA | phosphatidylinositol-4,5-bisphosphate 3-kinase catalyti | 5290 | up | 1,33 | 3,17E-02 |
| hsa05214 | Glioma | AKT2 | AKT serine/threonine kinase 2 | 208 | down | 1,28 | 6,33E-03 |
| hsa05214 | Glioma | PTEN | phosphatase and tensin homolog | 5728 | up | 1,33 | 4,72E-02 |
| hsa05214 | Glioma | CALM2 | calmodulin 2 | 805 | down | 1,43 | 2,23E-02 |
| hsa05214 | Glioma | BRAF | B-Raf proto-oncogene, serine/threonine kinase | 673 | up | 1,52 | 4,53E-02 |
| hsa05214 | Glioma | PLCG1 | phospholipase C gamma 1 | 5335 | down | 1,51 | 2,85E-02 |
| hsa04130 | SNARE interactions in vesicular transport | BNIP1 | BCL2 interacting protein 1 | 662 | up | 1,52 | 2,18E-02 |
| hsa04130 | SNARE interactions in vesicular transport | BET1 | Bet1 golgi vesicular membrane trafficking protein | 10282 | up | 2,34 | 3,79E-03 |
| hsa04130 | SNARE interactions in vesicular transport | VAMP2 | vesicle associated membrane protein 2 | 6844 | up | 1,50 | 4,82E-02 |
| hsa04130 | SNARE interactions in vesicular transport | STX5 | syntaxin 5 | 6811 | up | 1,66 | 2,40E-02 |
| hsa04130 | SNARE interactions in vesicular transport | STX7 | syntaxin 7 | 8417 | up | 1,29 | 3,61E-02 |
| hsa04130 | SNARE interactions in vesicular transport | GOSR2 | golgi SNAP receptor complex member 2 | 9570 | up | 1,55 | 2,21E-02 |
| hsa04130 | SNARE interactions in vesicular transport | GOSR1 | golgi SNAP receptor complex member 1 | 9527 | up | 1,39 | 3,34E-02 |
| hsa05223 | Non-small cell lung cancer | RASSF5 | Ras association domain family member 5 | 83593 | down | 2,36 | 2,58E-02 |
| hsa05223 | Non-small cell lung cancer | BRAF | B-Raf proto-oncogene, serine/threonine kinase | 673 | up | 1,52 | 4,53E-02 |
| hsa05223 | Non-small cell lung cancer | PLCG1 | phospholipase C gamma 1 | 5335 | down | 1,51 | 2,85E-02 |
| hsa05223 | Non-small cell lung cancer | ERBB2 | erb-b2 receptor tyrosine kinase 2 | 2064 | down | 1,47 | 1,93E-02 |
| hsa05223 | Non-small cell lung cancer | RXRA | retinoid X receptor alpha | 6256 | down | 1,34 | 3,67E-03 |
| hsa05223 | Non-small cell lung cancer | AKT2 | AKT serine/threonine kinase 2 | 208 | down | 1,28 | 6,33E-03 |
| hsa05223 | Non-small cell lung cancer | PIK3CA | phosphatidylinositol-4,5-bisphosphate 3-kinase catalyti | 5290 | up | 1,33 | 3,17E-02 |
| hsa05223 | Non-small cell lung cancer | SOS2 | SOS Ras/Rho guanine nucleotide exchange factor 2 | 6655 | up | 1,82 | 1,41E-02 |
| hsa05223 | Non-small cell lung cancer | AKT1 | AKT serine/threonine kinase 1 | 207 | down | 1,37 | 1,42E-03 |
| hsa05223 | Non-small cell lung cancer | E2F1 | E2F transcription factor 1 | 1869 | down | 1,64 | 3,25E-02 |
| hsa05223 | Non-small cell lung cancer | PIK3R1 | phosphoinositide-3-kinase regulatory subunit 1 | 5295 | down | 2,04 | 8,40E-03 |
| hsa05223 | Non-small cell lung cancer | HRAS | HRas proto-oncogene, GTPase | 3265 | down | 1,31 | 2,19E-03 |
| hsa04150 | mTOR signaling pathway | BRAF | B-Raf proto-oncogene, serine/threonine kinase | 673 | up | 1,52 | 4,53E-02 |
| hsa04150 | mTOR signaling pathway | RPS6KB2 | ribosomal protein S6 kinase B2 | 6199 | down | 1,22 | 1,01E-02 |
| hsa04150 | mTOR signaling pathway | RPTOR | regulatory associated protein of MTOR complex 1 | 57521 | down | 1,20 | 1,16E-02 |
| hsa04150 | mTOR signaling pathway | IRS1 | insulin receptor substrate 1 | 3667 | down | 1,89 | 6,25E-03 |
| hsa04150 | mTOR signaling pathway | AKT2 | AKT serine/threonine kinase 2 | 208 | down | 1,28 | 6,33E-03 |
| hsa04150 | mTOR signaling pathway | PIK3CA | phosphatidylinositol-4,5-bisphosphate 3-kinase catalyti | 5290 | up | 1,33 | 3,17E-02 |
| hsa04150 | mTOR signaling pathway | ULK3 | unc-51 like kinase 3 | 25989 | down | 1,22 | 4,13E-02 |
| hsa04150 | mTOR signaling pathway | STK11 | serine/threonine kinase 11 | 6794 | down | 1,25 | 3,05E-02 |
| hsa04150 | mTOR signaling pathway | PTEN | phosphatase and tensin homolog | 5728 | up | 1,33 | 4,72E-02 |
| hsa04150 | mTOR signaling pathway | AKT1 | AKT serine/threonine kinase 1 | 207 | down | 1,37 | 1,42E-03 |
| hsa04150 | mTOR signaling pathway | PIK3R1 | phosphoinositide-3-kinase regulatory subunit 1 | 5295 | down | 2,04 | 8,40E-03 |
| hsa04150 | mTOR signaling pathway | AKT1S1 | AKT1 substrate 1 | 84335 | down | 1,21 | 4,33E-02 |
| hsa04150 | mTOR signaling pathway | RRAGC | Ras related GTP binding C | 64121 | up | 1,78 | 6,10E-03 |
| hsa04360 | Axon guidance | L1CAM | L1 cell adhesion molecule | 3897 | down | 2,86 | 3,09E-02 |
| hsa04360 | Axon guidance | UNC5B | unc-5 netrin receptor B | 219699 | up | 4,39 | 5,27E-03 |
| hsa04360 | Axon guidance | NCK1 | NCK adaptor protein 1 | 4690 | up | 1,48 | 1,42E-02 |
| hsa04360 | Axon guidance | PPP3CC | protein phosphatase 3 catalytic subunit gamma | 5533 | up | 1,52 | 1,78E-02 |
| hsa04360 | Axon guidance | EFNA3 | ephrin A3 | 1944 | down | 2,00 | 4,89E-02 |
| hsa04360 | Axon guidance | SEMA7A | semaphorin 7A (John Milton Hagen blood group) | 8482 | up | 2,44 | 8,66E-03 |
| hsa04360 | Axon guidance | PLXNA1 | plexin A1 | 5361 | down | 1,28 | 9,70E-04 |
| hsa04360 | Axon guidance | NTN4 | netrin 4 | 59277 | up | 2,22 | 5,93E-03 |
| hsa04360 | Axon guidance | EFNA4 | ephrin A4 | 1945 | down | 1,43 | 4,92E-02 |
| hsa04360 | Axon guidance | SLIT1 | slit guidance ligand 1 | 6585 | up | 7,59 | 2,13E-03 |
| hsa04360 | Axon guidance | HRAS | HRas proto-oncogene, GTPase | 3265 | down | 1,31 | 2,19E-03 |
| hsa04360 | Axon guidance | EPHA5 | EPH receptor A5 | 2044 | down | 18,87 | 1,96E-02 |
| hsa04360 | Axon guidance | LIMK1 | LIM domain kinase 1 | 3984 | down | 1,44 | 6,49E-03 |
| hsa04360 | Axon guidance | EPHB2 | EPH receptor B2 | 2048 | down | 1,66 | 2,52E-02 |
| hsa04360 | Axon guidance | PLXNB1 | plexin B1 | 5364 | down | 1,32 | 3,64E-02 |
| hsa04360 | Axon guidance | SEMA3C | semaphorin 3C | 10512 | up | 1,42 | 3,05E-02 |
| hsa04360 | Axon guidance | SEMA4C | semaphorin 4C | 54910 | down | 1,72 | 8,31E-03 |
| hsa04360 | Axon guidance | RND1 | Rho family GTPase 1 | 27289 | up | 4,86 | 3,18E-03 |
| hsa04360 | Axon guidance | EPHA2 | EPH receptor A2 | 1969 | up | 1,89 | 2,51E-02 |
| hsa04360 | Axon guidance | RGS3 | regulator of G protein signaling 3 | 5998 | up | 1,77 | 1,19E-02 |
| hsa04360 | Axon guidance | DPYSL2 | dihydropyrimidinase like 2 | 1808 | down | 1,77 | 2,19E-02 |
| hsa04360 | Axon guidance | GNAI2 | G protein subunit alpha i2 | 2771 | down | 1,22 | 3,65E-02 |
| hsa04360 | Axon guidance | CFL1 | cofilin 1 | 1072 | down | 1,29 | 3,02E-02 |
| hsa01212 | Fatty acid metabolism | ACACA | acetyl-CoA carboxylase alpha | 31 | down | 1,40 | 2,54E-02 |
| hsa01212 | Fatty acid metabolism | FADS1 | fatty acid desaturase 1 | 3992 | down | 1,54 | 8,39E-03 |
| hsa01212 | Fatty acid metabolism | FASN | fatty acid synthase | 2194 | down | 1,96 | 8,31E-03 |
| hsa01212 | Fatty acid metabolism | ACAT2 | acetyl-CoA acetyltransferase 2 | 39 | down | 1,69 | 2,52E-02 |
| hsa01212 | Fatty acid metabolism | MCAT | malonyl-CoA-acyl carrier protein transacylase | 27349 | down | 1,52 | 3,36E-03 |
| hsa01212 | Fatty acid metabolism | ACSL1 | acyl-CoA synthetase long chain family member 1 | 2180 | down | 1,54 | 3,01E-02 |
| hsa01212 | Fatty acid metabolism | SCD | stearoyl-CoA desaturase | 6319 | down | 1,88 | 3,59E-02 |
| hsa01212 | Fatty acid metabolism | PPT2 | palmitoyl-protein thioesterase 2 | 9374 | down | 1,51 | 1,22E-02 |
| hsa04922 | Glucagon signaling pathway | ATF2 | activating transcription factor 2 | 1386 | up | 1,79 | 1,01E-02 |
| hsa04922 | Glucagon signaling pathway | AKT2 | AKT serine/threonine kinase 2 | 208 | down | 1,28 | 6,33E-03 |
| hsa04922 | Glucagon signaling pathway | PPP3CC | protein phosphatase 3 catalytic subunit gamma | 5533 | up | 1,52 | 1,78E-02 |
| hsa04922 | Glucagon signaling pathway | PRKAB2 | protein kinase AMP-activated non-catalytic subunit bet | 5565 | up | 1,83 | 1,27E-02 |
| hsa04922 | Glucagon signaling pathway | PPP4R3B | protein phosphatase 4 regulatory subunit 3B | 57223 | up | 1,33 | 4,10E-02 |
| hsa04922 | Glucagon signaling pathway | CAMK2D | calcium/calmodulin dependent protein kinase II delta | 817 | up | 1,29 | 1,49E-02 |
| hsa04922 | Glucagon signaling pathway | CALM1 | calmodulin 1 | 801 | down | 1,43 | 2,23E-02 |
| hsa04922 | Glucagon signaling pathway | CALM2 | calmodulin 2 | 805 | down | 1,43 | 2,23E-02 |
| hsa04922 | Glucagon signaling pathway | CREB3L2 | cAMP responsive element binding protein 3 like 2 | 64764 | up | 2,31 | 8,54E-03 |
| hsa04922 | Glucagon signaling pathway | ACACA | acetyl-CoA carboxylase alpha | 31 | down | 1,40 | 2,54E-02 |
| hsa04922 | Glucagon signaling pathway | SIRT1 | sirtuin 1 | 23411 | up | 1,60 | 1,00E-02 |
| hsa04922 | Glucagon signaling pathway | CALM3 | calmodulin 3 | 808 | down | 1,43 | 2,23E-02 |
| hsa04922 | Glucagon signaling pathway | PGAM1 | phosphoglycerate mutase 1 | 5223 | down | 1,62 | 5,67E-03 |
| hsa04922 | Glucagon signaling pathway | GYS1 | glycogen synthase 1 | 2997 | down | 1,32 | 2,63E-02 |
| hsa04922 | Glucagon signaling pathway | PKM | pyruvate kinase M1/2 | 5315 | down | 1,35 | 4,16E-02 |
| hsa04922 | Glucagon signaling pathway | CREB1 | cAMP responsive element binding protein 1 | 1385 | up | 1,51 | 7,18E-03 |
| hsa04922 | Glucagon signaling pathway | AKT1 | AKT serine/threonine kinase 1 | 207 | down | 1,37 | 1,42E-03 |
| hsa04922 | Glucagon signaling pathway | PRKAG2 | protein kinase AMP-activated non-catalytic subunit gan | 51422 | up | 1,28 | 4,57E-02 |
| hsa04922 | Glucagon signaling pathway | CREB5 | cAMP responsive element binding protein 5 | 9586 | up | 3,23 | 2,63E-03 |
| hsa04666 | Fc gamma R-mediated phagocytosis | AKT1 | AKT serine/threonine kinase 1 | 207 | down | 1,37 | 1,42E-03 |
| hsa04666 | Fc gamma R-mediated phagocytosis | SPHK1 | sphingosine kinase 1 | 8877 | down | 2,00 | 2,34E-02 |
| hsa04666 | Fc gamma R-mediated phagocytosis | CFL1 | cofilin 1 | 1072 | down | 1,29 | 3,02E-02 |

| | | | | | | | |
|---|---|---|---|---|---|---|---|
| hsa04666 | Fc gamma R-mediated phagocytosis | MARCKSL1 | MARCKS like 1 | 65108 | down | 1,63 | 1,61E-02 |
| hsa04666 | Fc gamma R-mediated phagocytosis | LYN | LYN proto-oncogene, Src family tyrosine kinase | 4067 | down | 1,36 | 2,34E-02 |
| hsa04666 | Fc gamma R-mediated phagocytosis | PIK3R1 | phosphoinositide-3-kinase regulatory subunit 1 | 5295 | down | 2,04 | 8,40E-03 |
| hsa04666 | Fc gamma R-mediated phagocytosis | PLD1 | phospholipase D1 | 5337 | up | 1,33 | 1,60E-02 |
| hsa04666 | Fc gamma R-mediated phagocytosis | PIP5K1C | phosphatidylinositol-4-phosphate 5-kinase type 1 gamm | 23396 | down | 1,39 | 1,49E-03 |
| hsa04666 | Fc gamma R-mediated phagocytosis | RPS6KB2 | ribosomal protein S6 kinase B2 | 6199 | down | 1,22 | 1,01E-02 |
| hsa04666 | Fc gamma R-mediated phagocytosis | LIMK1 | LIM domain kinase 1 | 3984 | down | 1,44 | 6,49E-03 |
| hsa04666 | Fc gamma R-mediated phagocytosis | ASAP1 | ArfGAP with SH3 domain, ankyrin repeat and PH doma | 50807 | up | 1,27 | 4,95E-02 |
| hsa04666 | Fc gamma R-mediated phagocytosis | PLCG1 | phospholipase C gamma 1 | 5335 | down | 1,51 | 2,85E-02 |
| hsa04666 | Fc gamma R-mediated phagocytosis | PIP5K1B | phosphatidylinositol-4-phosphate 5-kinase type 1 beta | 8395 | up | 3,40 | 1,84E-02 |
| hsa04666 | Fc gamma R-mediated phagocytosis | PIK3CA | phosphatidylinositol-4,5-bisphosphate 3-kinase catalyti | 5290 | up | 1,33 | 3,17E-02 |
| hsa04666 | Fc gamma R-mediated phagocytosis | AKT2 | AKT serine/threonine kinase 2 | 208 | down | 1,28 | 6,33E-03 |
| hsa05230 | Central carbon metabolism in cancer | FGFR3 | fibroblast growth factor receptor 3 | 2261 | down | 1,70 | 4,60E-02 |
| hsa05230 | Central carbon metabolism in cancer | PDK1 | pyruvate dehydrogenase kinase 1 | 5163 | up | 1,31 | 2,69E-03 |
| hsa05230 | Central carbon metabolism in cancer | ERBB2 | erb-b2 receptor tyrosine kinase 2 | 2064 | down | 1,47 | 1,93E-02 |
| hsa05230 | Central carbon metabolism in cancer | HKDC1 | hexokinase domain containing 1 | 80201 | up | 3,37 | 4,73E-03 |
| hsa05230 | Central carbon metabolism in cancer | SIRT3 | sirtuin 3 | 23410 | down | 1,27 | 4,49E-02 |
| hsa05230 | Central carbon metabolism in cancer | PIK3CA | phosphatidylinositol-4,5-bisphosphate 3-kinase catalyti | 5290 | up | 1,33 | 3,17E-02 |
| hsa05230 | Central carbon metabolism in cancer | AKT2 | AKT serine/threonine kinase 2 | 208 | down | 1,28 | 6,33E-03 |
| hsa05230 | Central carbon metabolism in cancer | PTEN | phosphatase and tensin homolog | 5728 | up | 1,33 | 4,72E-02 |
| hsa05230 | Central carbon metabolism in cancer | AKT1 | AKT serine/threonine kinase 1 | 207 | down | 1,37 | 1,42E-03 |
| hsa05230 | Central carbon metabolism in cancer | PKM | pyruvate kinase M1/2 | 5315 | down | 1,35 | 4,16E-02 |
| hsa05230 | Central carbon metabolism in cancer | PGAM1 | phosphoglycerate mutase 1 | 5223 | down | 1,62 | 5,67E-03 |
| hsa05230 | Central carbon metabolism in cancer | HIF1A | hypoxia inducible factor 1 alpha subunit | 3091 | up | 1,23 | 4,66E-02 |
| hsa05230 | Central carbon metabolism in cancer | PIK3R1 | phosphoinositide-3-kinase regulatory subunit 1 | 5295 | down | 2,04 | 8,40E-03 |
| hsa05230 | Central carbon metabolism in cancer | HRAS | HRas proto-oncogene, GTPase | 3265 | down | 1,31 | 2,19E-03 |
| hsa00562 | Inositol phosphate metabolism | PIK3C2A | phosphatidylinositol-4-phosphate 3-kinase catalytic sub | 5286 | up | 1,30 | 1,79E-02 |
| hsa00562 | Inositol phosphate metabolism | ITPKC | inositol-trisphosphate 3-kinase C | 80271 | up | 1,63 | 3,32E-02 |
| hsa00562 | Inositol phosphate metabolism | PIP4K2B | phosphatidylinositol-5-phosphate 4-kinase type 2 beta | 8396 | down | 1,53 | 1,01E-02 |
| hsa00562 | Inositol phosphate metabolism | ISYNA1 | inositol-3-phosphate synthase 1 | 51477 | down | 1,54 | 7,64E-03 |
| hsa00562 | Inositol phosphate metabolism | MTMR6 | myotubularin related protein 6 | 9107 | up | 1,33 | 2,92E-02 |
| hsa00562 | Inositol phosphate metabolism | ALDH6A1 | aldehyde dehydrogenase 6 family member A1 | 4329 | up | 1,60 | 3,50E-02 |
| hsa00562 | Inositol phosphate metabolism | ITPKB | inositol-trisphosphate 3-kinase B | 3707 | down | 1,71 | 1,97E-02 |
| hsa00562 | Inositol phosphate metabolism | PTEN | phosphatase and tensin homolog | 5728 | up | 1,33 | 4,72E-02 |
| hsa00562 | Inositol phosphate metabolism | CDIPT | CDP-diacylglycerol--inositol 3-phosphatidyltransferase | 10423 | down | 1,32 | 3,12E-03 |
| hsa00562 | Inositol phosphate metabolism | PIK3CA | phosphatidylinositol-4,5-bisphosphate 3-kinase catalyti | 5290 | up | 1,33 | 3,17E-02 |
| hsa00562 | Inositol phosphate metabolism | PIP5K1B | phosphatidylinositol-4-phosphate 5-kinase type 1 beta | 8395 | up | 3,40 | 1,84E-02 |
| hsa00562 | Inositol phosphate metabolism | TPI1 | triosephosphate isomerase 1 | 7167 | down | 1,28 | 3,58E-02 |
| hsa00562 | Inositol phosphate metabolism | MTMR2 | myotubularin related protein 2 | 8898 | up | 1,23 | 2,61E-02 |
| hsa00562 | Inositol phosphate metabolism | PLCG1 | phospholipase C gamma 1 | 5335 | down | 1,51 | 2,85E-02 |
| hsa00562 | Inositol phosphate metabolism | PIP5K1C | phosphatidylinositol-4-phosphate 5-kinase type 1 gamm | 23396 | down | 1,39 | 1,49E-03 |
| hsa04210 | Apoptosis | AKT2 | AKT serine/threonine kinase 2 | 208 | down | 1,28 | 6,33E-03 |
| hsa04210 | Apoptosis | BAX | BCL2 associated X, apoptosis regulator | 581 | down | 1,22 | 2,61E-02 |
| hsa04210 | Apoptosis | DFFB | DNA fragmentation factor subunit beta | 1677 | down | 2,29 | 3,17E-02 |
| hsa04210 | Apoptosis | BID | BH3 interacting domain death agonist | 637 | down | 1,40 | 2,42E-02 |
| hsa04210 | Apoptosis | TRAF2 | TNF receptor associated factor 2 | 7186 | down | 1,47 | 2,65E-02 |
| hsa04210 | Apoptosis | PIK3R1 | phosphoinositide-3-kinase regulatory subunit 1 | 5295 | down | 2,04 | 8,40E-03 |
| hsa04210 | Apoptosis | FADD | Fas associated via death domain | 8772 | down | 1,31 | 8,90E-03 |
| hsa04210 | Apoptosis | AKT1 | AKT serine/threonine kinase 1 | 207 | down | 1,37 | 1,42E-03 |
| hsa04210 | Apoptosis | CAPN1 | calpain 1 | 823 | down | 1,27 | 2,65E-02 |
| hsa04012 | ErbB signaling pathway | CAMK2D | calcium/calmodulin dependent protein kinase II delta | 817 | up | 1,29 | 1,49E-02 |
| hsa04012 | ErbB signaling pathway | HRAS | HRas proto-oncogene, GTPase | 3265 | down | 1,31 | 2,19E-03 |
| hsa04012 | ErbB signaling pathway | AREG | amphiregulin | 374 | up | 3,33 | 2,37E-02 |
| hsa04012 | ErbB signaling pathway | PIK3R1 | phosphoinositide-3-kinase regulatory subunit 1 | 5295 | down | 2,04 | 8,40E-03 |
| hsa04012 | ErbB signaling pathway | MAPK8 | mitogen-activated protein kinase 8 | 5599 | up | 1,38 | 3,05E-02 |
| hsa04012 | ErbB signaling pathway | SOS2 | SOS Ras/Rho guanine nucleotide exchange factor 2 | 6655 | up | 1,82 | 1,41E-02 |
| hsa04012 | ErbB signaling pathway | NCK1 | NCK adaptor protein 1 | 4690 | up | 1,48 | 1,42E-02 |
| hsa04012 | ErbB signaling pathway | PIK3CA | phosphatidylinositol-4,5-bisphosphate 3-kinase catalyti | 5290 | up | 1,33 | 3,17E-02 |
| hsa04012 | ErbB signaling pathway | AKT2 | AKT serine/threonine kinase 2 | 208 | down | 1,28 | 6,33E-03 |
| hsa04012 | ErbB signaling pathway | PLCG1 | phospholipase C gamma 1 | 5335 | down | 1,51 | 2,85E-02 |
| hsa04012 | ErbB signaling pathway | BRAF | B-Raf proto-oncogene, serine/threonine kinase | 673 | up | 1,52 | 4,53E-02 |
| hsa04012 | ErbB signaling pathway | RPS6KB2 | ribosomal protein S6 kinase B2 | 6199 | down | 1,22 | 1,01E-02 |
| hsa04012 | ErbB signaling pathway | CDKN1A | cyclin dependent kinase inhibitor 1A | 1026 | up | 3,62 | 5,43E-03 |
| hsa04012 | ErbB signaling pathway | HBEGF | heparin binding EGF like growth factor | 1839 | up | 3,67 | 5,67E-03 |
| hsa04012 | ErbB signaling pathway | AKT1 | AKT serine/threonine kinase 1 | 207 | down | 1,37 | 1,42E-03 |
| hsa04012 | ErbB signaling pathway | ERBB2 | erb-b2 receptor tyrosine kinase 2 | 2064 | down | 1,47 | 1,93E-02 |
| hsa04012 | ErbB signaling pathway | CBLB | Cbl proto-oncogene B | 868 | up | 1,83 | 4,35E-02 |
| hsa00970 | Aminoacyl-tRNA biosynthesis | SARS | seryl-tRNA synthetase | 6301 | up | 1,82 | 1,31E-02 |
| hsa00970 | Aminoacyl-tRNA biosynthesis | WARS | tryptophanyl-tRNA synthetase | 7453 | up | 1,88 | 3,69E-02 |
| hsa00970 | Aminoacyl-tRNA biosynthesis | NARS | asparaginyl-tRNA synthetase | 4677 | up | 1,29 | 2,58E-02 |
| hsa00970 | Aminoacyl-tRNA biosynthesis | CARS | cysteinyl-tRNA synthetase | 833 | up | 1,55 | 4,38E-02 |
| hsa00970 | Aminoacyl-tRNA biosynthesis | AARS | alanyl-tRNA synthetase | 16 | up | 1,82 | 1,84E-02 |
| hsa00970 | Aminoacyl-tRNA biosynthesis | TARS | threonyl-tRNA synthetase | 6897 | up | 1,45 | 1,96E-02 |
| hsa00970 | Aminoacyl-tRNA biosynthesis | WARS2 | tryptophanyl tRNA synthetase 2, mitochondrial | 10352 | up | 1,54 | 9,54E-03 |
| hsa00970 | Aminoacyl-tRNA biosynthesis | EPRS | glutamyl-prolyl-tRNA synthetase | 2058 | up | 1,42 | 1,95E-02 |
| hsa00970 | Aminoacyl-tRNA biosynthesis | GARS | glycyl-tRNA synthetase | 2617 | up | 1,65 | 1,79E-02 |
| hsa04710 | Circadian rhythm | PRKAB2 | protein kinase AMP-activated non-catalytic subunit bet | 5565 | up | 1,83 | 1,27E-02 |
| hsa04710 | Circadian rhythm | CREB1 | cAMP responsive element binding protein 1 | 1385 | up | 1,51 | 7,18E-03 |
| hsa04710 | Circadian rhythm | BHLHE40 | basic helix-loop-helix family member e40 | 8553 | up | 1,80 | 3,56E-02 |
| hsa04710 | Circadian rhythm | PRKAG2 | protein kinase AMP-activated non-catalytic subunit gar | 51422 | up | 1,28 | 4,57E-02 |
| hsa04710 | Circadian rhythm | PER1 | period circadian regulator 1 | 5187 | up | 2,35 | 1,38E-02 |
| hsa04710 | Circadian rhythm | ARNTL | aryl hydrocarbon receptor nuclear translocator like | 406 | up | 1,67 | 5,94E-03 |
| hsa04115 | p53 signaling pathway | BID | BH3 interacting domain death agonist | 637 | down | 1,40 | 2,42E-02 |
| hsa04115 | p53 signaling pathway | CD82 | CD82 molecule | 3732 | down | 1,36 | 1,80E-02 |
| hsa04115 | p53 signaling pathway | EI24 | EI24, autophagy associated transmembrane protein | 9538 | down | 1,27 | 2,31E-02 |
| hsa04115 | p53 signaling pathway | BAX | BCL2 associated X, apoptosis regulator | 581 | down | 1,22 | 2,61E-02 |
| hsa04115 | p53 signaling pathway | CCND3 | cyclin D3 | 896 | down | 1,60 | 4,17E-02 |
| hsa04115 | p53 signaling pathway | PTEN | phosphatase and tensin homolog | 5728 | up | 1,33 | 4,72E-02 |
| hsa04115 | p53 signaling pathway | BBC3 | BCL2 binding component 3 | 27113 | up | 2,09 | 3,17E-02 |
| hsa04115 | p53 signaling pathway | SESN2 | sestrin 2 | 83667 | up | 2,90 | 1,62E-02 |
| hsa04115 | p53 signaling pathway | GADD45A | growth arrest and DNA damage inducible alpha | 1647 | up | 3,05 | 4,75E-04 |
| hsa04115 | p53 signaling pathway | SIAH1 | siah E3 ubiquitin protein ligase 1 | 6477 | up | 1,74 | 1,66E-02 |
| hsa04115 | p53 signaling pathway | PMAIP1 | phorbol-12-myristate-13-acetate-induced protein 1 | 5366 | up | 2,01 | 4,52E-02 |
| hsa04115 | p53 signaling pathway | CDKN1A | cyclin dependent kinase inhibitor 1A | 1026 | up | 3,62 | 5,43E-03 |
| hsa04115 | p53 signaling pathway | GADD45B | growth arrest and DNA damage inducible beta | 4616 | up | 3,21 | 3,92E-04 |
| hsa04115 | p53 signaling pathway | TP73 | tumor protein p73 | 7161 | down | 2,86 | 3,40E-02 |
| hsa03420 | Nucleotide excision repair | RAD23B | RAD23 homolog B, nucleotide excision repair protein | 5887 | up | 1,23 | 3,34E-02 |
| hsa03420 | Nucleotide excision repair | GTF2H1 | general transcription factor IIH subunit 1 | 2965 | up | 1,33 | 1,85E-02 |
| hsa03420 | Nucleotide excision repair | POLD1 | DNA polymerase delta 1, catalytic subunit | 5424 | down | 1,59 | 4,62E-02 |
| hsa03420 | Nucleotide excision repair | PCNA | proliferating cell nuclear antigen | 5111 | down | 2,14 | 4,63E-02 |
| hsa03420 | Nucleotide excision repair | ERCC2 | ERCC excision repair 2, TFIIH core complex helicase su | 2068 | down | 1,29 | 2,20E-02 |
| hsa03420 | Nucleotide excision repair | POLE | DNA polymerase epsilon, catalytic subunit | 5426 | down | 1,55 | 2,67E-02 |
| hsa03420 | Nucleotide excision repair | ERCC3 | ERCC excision repair 3, TFIIH core complex helicase su | 2071 | down | 1,20 | 2,37E-02 |
| hsa03420 | Nucleotide excision repair | RFC1 | replication factor C subunit 1 | 5981 | up | 1,15 | 2,18E-02 |
| hsa03420 | Nucleotide excision repair | POLD2 | DNA polymerase delta 2, accessory subunit | 5425 | down | 1,30 | 1,92E-02 |
| hsa03420 | Nucleotide excision repair | RPA1 | replication protein A1 | 6117 | down | 1,41 | 4,93E-02 |
| hsa03420 | Nucleotide excision repair | CDK7 | cyclin dependent kinase 7 | 1022 | up | 1,37 | 4,38E-02 |
| hsa04920 | Adipocytokine signaling pathway | ACSL1 | acyl-CoA synthetase long chain family member 1 | 2180 | down | 1,54 | 3,01E-02 |
| hsa04920 | Adipocytokine signaling pathway | CAMKK2 | calcium/calmodulin dependent protein kinase kinase 2 | 10645 | down | 1,63 | 2,18E-02 |
| hsa04920 | Adipocytokine signaling pathway | AKT1 | AKT serine/threonine kinase 1 | 207 | down | 1,37 | 1,42E-03 |
| hsa04920 | Adipocytokine signaling pathway | PRKAG2 | protein kinase AMP-activated non-catalytic subunit gar | 51422 | up | 1,28 | 4,57E-02 |
| hsa04920 | Adipocytokine signaling pathway | IRS2 | insulin receptor substrate 2 | 8660 | up | 2,21 | 2,22E-02 |
| hsa04920 | Adipocytokine signaling pathway | MAPK8 | mitogen-activated protein kinase 8 | 5599 | up | 1,38 | 3,05E-02 |
| hsa04920 | Adipocytokine signaling pathway | LEPR | leptin receptor | 3953 | up | 1,33 | 4,94E-03 |
| hsa04920 | Adipocytokine signaling pathway | AKT2 | AKT serine/threonine kinase 2 | 208 | down | 1,28 | 6,33E-03 |
| hsa04920 | Adipocytokine signaling pathway | RXRA | retinoid X receptor alpha | 6256 | down | 1,34 | 3,67E-03 |
| hsa04920 | Adipocytokine signaling pathway | STK11 | serine/threonine kinase 11 | 6794 | down | 1,25 | 3,05E-02 |
| hsa04920 | Adipocytokine signaling pathway | CAMKK1 | calcium/calmodulin dependent protein kinase kinase 1 | 84254 | down | 1,83 | 2,57E-02 |
| hsa04920 | Adipocytokine signaling pathway | PRKAB2 | protein kinase AMP-activated non-catalytic subunit bet | 5565 | up | 1,83 | 1,27E-02 |
| hsa04920 | Adipocytokine signaling pathway | TRAF2 | TNF receptor associated factor 2 | 7186 | down | 1,47 | 2,65E-02 |
| hsa04920 | Adipocytokine signaling pathway | IRS1 | insulin receptor substrate 1 | 3667 | down | 1,89 | 6,25E-03 |
| hsa05200 | Pathways in cancer | RALGDS | ral guanine nucleotide dissociation stimulator | 5900 | down | 1,30 | 3,49E-02 |
| hsa05200 | Pathways in cancer | GNA11 | G protein subunit alpha 11 | 2767 | down | 1,18 | 4,37E-02 |
| hsa05200 | Pathways in cancer | FZD4 | frizzled class receptor 4 | 8322 | down | 2,17 | 3,82E-02 |
| hsa05200 | Pathways in cancer | E2F1 | E2F transcription factor 1 | 1869 | down | 1,64 | 3,25E-02 |
| hsa05200 | Pathways in cancer | TGFBR2 | transforming growth factor beta receptor 2 | 7048 | down | 1,34 | 2,74E-02 |
| hsa05200 | Pathways in cancer | BMP4 | bone morphogenetic protein 4 | 652 | down | 2,60 | 1,26E-02 |
| hsa05200 | Pathways in cancer | HRAS | HRas proto-oncogene, GTPase | 3265 | down | 1,31 | 2,19E-03 |
| hsa05200 | Pathways in cancer | FGF2 | fibroblast growth factor 2 | 2247 | down | 5,35 | 3,09E-02 |
| hsa05200 | Pathways in cancer | CTBP1 | C-terminal binding protein 1 | 1487 | down | 1,16 | 3,77E-02 |
| hsa05200 | Pathways in cancer | PIK3R1 | phosphoinositide-3-kinase regulatory subunit 1 | 5295 | down | 2,04 | 8,40E-03 |
| hsa05200 | Pathways in cancer | TRAF2 | TNF receptor associated factor 2 | 7186 | down | 1,47 | 2,65E-02 |
| hsa05200 | Pathways in cancer | FGFR3 | fibroblast growth factor receptor 3 | 2261 | down | 1,70 | 4,60E-02 |
| hsa05200 | Pathways in cancer | PLCG1 | phospholipase C gamma 1 | 5335 | down | 1,51 | 2,85E-02 |
| hsa05200 | Pathways in cancer | WNT10B | Wnt family member 10B | 7480 | down | 1,76 | 4,78E-02 |
| hsa05200 | Pathways in cancer | RASSF5 | Ras association domain family member 5 | 83593 | down | 2,36 | 2,58E-02 |
| hsa05200 | Pathways in cancer | FZD1 | frizzled class receptor 1 | 8321 | down | 1,75 | 2,84E-02 |
| hsa05200 | Pathways in cancer | SUFU | SUFU negative regulator of hedgehog signaling | 51684 | down | 1,41 | 3,37E-02 |
| hsa05200 | Pathways in cancer | AKT2 | AKT serine/threonine kinase 2 | 208 | down | 1,28 | 6,33E-03 |
| hsa05200 | Pathways in cancer | SKP2 | S-phase kinase associated protein 2 | 6502 | down | 2,21 | 3,69E-02 |
| hsa05200 | Pathways in cancer | F2R | coagulation factor II thrombin receptor | 2149 | down | 1,54 | 7,69E-03 |
| hsa05200 | Pathways in cancer | AKT1 | AKT serine/threonine kinase 1 | 207 | down | 1,37 | 1,42E-03 |
| hsa05200 | Pathways in cancer | GNAI2 | G protein subunit alpha i2 | 2771 | down | 1,22 | 3,65E-02 |
| hsa05200 | Pathways in cancer | TRAF3 | TNF receptor associated factor 3 | 7187 | down | 1,40 | 1,36E-02 |
| hsa05200 | Pathways in cancer | FADD | Fas associated via death domain | 8772 | down | 1,31 | 8,90E-03 |
| hsa05200 | Pathways in cancer | AXIN2 | axin 2 | 8313 | down | 2,79 | 3,67E-03 |
| hsa05200 | Pathways in cancer | ERBB2 | erb-b2 receptor tyrosine kinase 2 | 2064 | down | 1,47 | 1,93E-02 |
| hsa05200 | Pathways in cancer | ADCY3 | adenylate cyclase 3 | 109 | down | 1,45 | 1,03E-02 |

| | | | | | | | |
|---|---|---|---|---|---|---|---|
| hsa05200 | Pathways in cancer | BID | BH3 interacting domain death agonist | 637 | down | 1,40 | 2,42E-02 |
| hsa05200 | Pathways in cancer | TCF7 | transcription factor 7 | 6932 | down | 1,47 | 1,07E-02 |
| hsa05200 | Pathways in cancer | BAX | BCL2 associated X, apoptosis regulator | 581 | down | 1,22 | 2,61E-02 |
| hsa05200 | Pathways in cancer | RXRA | retinoid X receptor alpha | 6256 | down | 1,34 | 3,67E-03 |
| hsa00061 | Fatty acid biosynthesis | ACACA | acetyl-CoA carboxylase alpha | 31 | down | 1,40 | 2,54E-02 |
| hsa00061 | Fatty acid biosynthesis | FASN | fatty acid synthase | 2194 | down | 1,96 | 8,31E-03 |
| hsa00061 | Fatty acid biosynthesis | MCAT | malonyl-CoA-acyl carrier protein transacylase | 27349 | down | 1,39 | 3,36E-03 |
| hsa00061 | Fatty acid biosynthesis | ACSL1 | acyl-CoA synthetase long chain family member 1 | 2180 | down | 1,54 | 3,01E-02 |
| hsa04010 | MAPK signaling pathway | TAB1 | TGF-beta activated kinase 1 (MAP3K7) binding protein | 10454 | down | 1,29 | 1,94E-02 |
| hsa04010 | MAPK signaling pathway | GADD45A | growth arrest and DNA damage inducible alpha | 1647 | up | 3,05 | 4,75E-04 |
| hsa04010 | MAPK signaling pathway | AKT1 | AKT serine/threonine kinase 1 | 207 | down | 1,37 | 1,42E-03 |
| hsa04010 | MAPK signaling pathway | GADD45B | growth arrest and DNA damage inducible beta | 4616 | up | 3,21 | 3,92E-04 |
| hsa04010 | MAPK signaling pathway | JUND | JunD proto-oncogene, AP-1 transcription factor subuni | 3727 | up | 1,56 | 4,42E-02 |
| hsa04010 | MAPK signaling pathway | DUSP10 | dual specificity phosphatase 10 | 11221 | up | 2,34 | 2,20E-03 |
| hsa04010 | MAPK signaling pathway | RPS6KA5 | ribosomal protein S6 kinase A5 | 9252 | up | 2,11 | 5,94E-03 |
| hsa04010 | MAPK signaling pathway | MAPK8 | mitogen-activated protein kinase 8 | 5599 | up | 1,38 | 3,05E-02 |
| hsa04010 | MAPK signaling pathway | MAPK12 | mitogen-activated protein kinase 12 | 6300 | down | 1,37 | 3,32E-03 |
| hsa04010 | MAPK signaling pathway | DUSP5 | dual specificity phosphatase 5 | 1847 | up | 3,63 | 9,77E-03 |
| hsa04010 | MAPK signaling pathway | HRAS | HRas proto-oncogene, GTPase | 3265 | down | 1,31 | 2,19E-03 |
| hsa04010 | MAPK signaling pathway | FGF2 | fibroblast growth factor 2 | 2247 | down | 5,35 | 3,09E-02 |
| hsa04010 | MAPK signaling pathway | SRF | serum response factor | 6722 | up | 1,48 | 3,97E-02 |
| hsa04010 | MAPK signaling pathway | CACNB3 | calcium voltage-gated channel auxiliary subunit beta 3 | 784 | down | 1,19 | 3,30E-02 |
| hsa04010 | MAPK signaling pathway | TRAF2 | TNF receptor associated factor 2 | 7186 | down | 1,47 | 2,65E-02 |
| hsa04010 | MAPK signaling pathway | BRAF | B-Raf proto-oncogene, serine/threonine kinase | 673 | up | 1,52 | 4,53E-02 |
| hsa04010 | MAPK signaling pathway | MAP3K2 | mitogen-activated protein kinase kinase kinase 2 | 10746 | up | 1,27 | 8,96E-03 |
| hsa04010 | MAPK signaling pathway | PPM1B | protein phosphatase, Mg2+/Mn2+ dependent 1B | 5495 | up | 1,53 | 2,84E-02 |
| hsa04010 | MAPK signaling pathway | IL1A | interleukin 1 alpha | 3552 | up | 8,56 | 1,05E-02 |
| hsa04010 | MAPK signaling pathway | DUSP1 | dual specificity phosphatase 1 | 1843 | up | 1,99 | 2,51E-02 |
| hsa04010 | MAPK signaling pathway | MAP3K6 | mitogen-activated protein kinase kinase kinase 6 | 9064 | down | 1,68 | 3,57E-02 |
| hsa04010 | MAPK signaling pathway | MAP4K3 | mitogen-activated protein kinase kinase kinase kinase | 8491 | up | 1,23 | 4,82E-02 |
| hsa04010 | MAPK signaling pathway | BDNF | brain derived neurotrophic factor | 627 | down | 2,98 | 1,15E-02 |
| hsa04010 | MAPK signaling pathway | DUSP8 | dual specificity phosphatase 8 | 1850 | up | 2,42 | 1,40E-02 |
| hsa04010 | MAPK signaling pathway | RASGRP3 | RAS guanyl releasing protein 3 | 25780 | up | 2,59 | 1,78E-02 |
| hsa04010 | MAPK signaling pathway | STK3 | serine/threonine kinase 3 | 6788 | up | 1,61 | 2,08E-03 |
| hsa04010 | MAPK signaling pathway | TGFBR2 | transforming growth factor beta receptor 2 | 7048 | down | 1,34 | 2,74E-02 |
| hsa04010 | MAPK signaling pathway | DUSP7 | dual specificity phosphatase 7 | 1849 | down | 1,42 | 1,78E-02 |
| hsa04010 | MAPK signaling pathway | SOS2 | SOS Ras/Rho guanine nucleotide exchange factor 2 | 6655 | up | 1,82 | 1,41E-02 |
| hsa04010 | MAPK signaling pathway | ARRB1 | arrestin beta 1 | 408 | down | 1,79 | 2,42E-02 |
| hsa04010 | MAPK signaling pathway | PPP5C | protein phosphatase 5 catalytic subunit | 5536 | down | 1,29 | 4,15E-02 |
| hsa04010 | MAPK signaling pathway | FGFR3 | fibroblast growth factor receptor 3 | 2261 | down | 1,70 | 4,60E-02 |
| hsa04010 | MAPK signaling pathway | DDIT3 | DNA damage inducible transcript 3 | 1649 | up | 5,65 | 3,67E-03 |
| hsa04010 | MAPK signaling pathway | RPS6KA4 | ribosomal protein S6 kinase A4 | 8986 | down | 1,33 | 5,67E-03 |
| hsa04010 | MAPK signaling pathway | ATF2 | activating transcription factor 2 | 1386 | up | 1,79 | 1,01E-02 |
| hsa04010 | MAPK signaling pathway | AKT2 | AKT serine/threonine kinase 2 | 208 | down | 1,28 | 6,33E-03 |
| hsa04010 | MAPK signaling pathway | PPP3CC | protein phosphatase 3 catalytic subunit gamma | 5533 | up | 1,52 | 1,78E-02 |
| hsa04110 | Cell cycle | PCNA | proliferating cell nuclear antigen | 5111 | down | 2,14 | 4,63E-02 |
| hsa04110 | Cell cycle | MCM2 | minichromosome maintenance complex component 2 | 4171 | down | 2,01 | 1,42E-02 |
| hsa04110 | Cell cycle | E2F1 | E2F transcription factor 1 | 1869 | down | 1,64 | 3,25E-02 |
| hsa04110 | Cell cycle | ANAPC2 | anaphase promoting complex subunit 2 | 29882 | down | 1,26 | 1,99E-02 |
| hsa04110 | Cell cycle | MCM7 | minichromosome maintenance complex component 7 | 4176 | down | 1,67 | 3,35E-02 |
| hsa04110 | Cell cycle | TFDP1 | transcription factor Dp-1 | 7027 | down | 1,77 | 1,08E-02 |
| hsa04110 | Cell cycle | ESPL1 | extra spindle pole bodies like 1, separase | 9700 | down | 1,88 | 3,17E-02 |
| hsa04110 | Cell cycle | MCM5 | minichromosome maintenance complex component 5 | 4174 | down | 1,83 | 2,22E-02 |
| hsa04110 | Cell cycle | CDC25A | cell division cycle 25A | 993 | down | 1,92 | 1,48E-02 |
| hsa04110 | Cell cycle | CCND3 | cyclin D3 | 896 | down | 1,60 | 4,17E-02 |
| hsa04110 | Cell cycle | YWHAH | tyrosine 3-monooxygenase/tryptophan 5-monooxygena | 7533 | down | 1,33 | 4,79E-02 |
| hsa04110 | Cell cycle | SKP2 | S-phase kinase associated protein 2 | 6502 | down | 2,21 | 3,69E-02 |
| hsa04110 | Cell cycle | MCM3 | minichromosome maintenance complex component 3 | 4172 | down | 2,07 | 4,13E-02 |
| hsa05160 | Hepatitis C | PPP2CB | protein phosphatase 2 catalytic subunit beta | 5516 | up | 1,30 | 4,79E-02 |
| hsa05160 | Hepatitis C | CDKN1A | cyclin dependent kinase inhibitor 1A | 1026 | up | 3,62 | 5,43E-03 |
| hsa05160 | Hepatitis C | EIF2AK3 | eukaryotic translation initiation factor 2 alpha kinase 3 | 9451 | up | 2,82 | 1,11E-03 |
| hsa05160 | Hepatitis C | TRAF3 | TNF receptor associated factor 3 | 7187 | down | 1,40 | 1,36E-02 |
| hsa05160 | Hepatitis C | AKT1 | AKT serine/threonine kinase 1 | 207 | down | 1,37 | 1,42E-03 |
| hsa05160 | Hepatitis C | JAK1 | Janus kinase 1 | 3716 | up | 1,53 | 2,13E-02 |
| hsa05160 | Hepatitis C | RXRA | retinoid X receptor alpha | 6256 | down | 1,34 | 3,67E-03 |
| hsa05160 | Hepatitis C | PIK3R1 | phosphoinositide-3-kinase regulatory subunit 1 | 5295 | down | 2,04 | 8,40E-03 |
| hsa05160 | Hepatitis C | HRAS | HRas proto-oncogene, GTPase | 3265 | down | 1,31 | 2,19E-03 |
| hsa05160 | Hepatitis C | CLDN1 | claudin 1 | 9076 | up | 2,13 | 6,96E-03 |
| hsa05160 | Hepatitis C | IRF3 | interferon regulatory factor 3 | 3661 | down | 1,33 | 2,58E-03 |
| hsa05160 | Hepatitis C | SOS2 | SOS Ras/Rho guanine nucleotide exchange factor 2 | 6655 | up | 1,82 | 1,41E-02 |
| hsa05160 | Hepatitis C | MAPK12 | mitogen-activated protein kinase 12 | 6300 | down | 1,37 | 3,32E-03 |
| hsa05160 | Hepatitis C | IKBKE | inhibitor of nuclear factor kappa B kinase subunit epsilo | 9641 | down | 1,67 | 1,00E-02 |
| hsa05160 | Hepatitis C | MAPK8 | mitogen-activated protein kinase 8 | 5599 | up | 1,38 | 3,05E-02 |
| hsa05160 | Hepatitis C | AKT2 | AKT serine/threonine kinase 2 | 208 | down | 1,28 | 6,33E-03 |
| hsa05160 | Hepatitis C | PIK3CA | phosphatidylinositol-4,5-bisphosphate 3-kinase catalyti | 5290 | up | 1,33 | 3,17E-02 |
| hsa05160 | Hepatitis C | CD81 | CD81 molecule | 975 | down | 1,31 | 1,15E-02 |
| hsa05160 | Hepatitis C | BRAF | B-Raf proto-oncogene, serine/threonine kinase | 673 | up | 1,52 | 4,53E-02 |
| hsa05160 | Hepatitis C | SCARB1 | scavenger receptor class B member 1 | 949 | down | 1,45 | 4,26E-02 |
| hsa05160 | Hepatitis C | PPP2R2A | protein phosphatase 2 regulatory subunit Balpha | 5520 | up | 1,26 | 3,61E-02 |
| hsa05160 | Hepatitis C | TRAF2 | TNF receptor associated factor 2 | 7186 | down | 1,47 | 2,65E-02 |
| hsa05212 | Pancreatic cancer | MAPK8 | mitogen-activated protein kinase 8 | 5599 | up | 1,38 | 3,05E-02 |
| hsa05212 | Pancreatic cancer | TGFBR2 | transforming growth factor beta receptor 2 | 7048 | down | 1,34 | 2,74E-02 |
| hsa05212 | Pancreatic cancer | RALGDS | ral guanine nucleotide dissociation stimulator | 5900 | down | 1,30 | 3,49E-02 |
| hsa05212 | Pancreatic cancer | SMAD2 | SMAD family member 2 | 4087 | up | 1,29 | 2,19E-02 |
| hsa05212 | Pancreatic cancer | E2F1 | E2F transcription factor 1 | 1869 | down | 1,64 | 3,25E-02 |
| hsa05212 | Pancreatic cancer | VEGFA | vascular endothelial growth factor A | 7422 | up | 2,38 | 1,68E-02 |
| hsa05212 | Pancreatic cancer | AKT1 | AKT serine/threonine kinase 1 | 207 | down | 1,37 | 1,42E-03 |
| hsa05212 | Pancreatic cancer | JAK1 | Janus kinase 1 | 3716 | up | 1,53 | 2,13E-02 |
| hsa05212 | Pancreatic cancer | PIK3R1 | phosphoinositide-3-kinase regulatory subunit 1 | 5295 | down | 2,04 | 8,40E-03 |
| hsa05212 | Pancreatic cancer | ERBB2 | erb-b2 receptor tyrosine kinase 2 | 2064 | down | 1,47 | 1,93E-02 |
| hsa05212 | Pancreatic cancer | BRAF | B-Raf proto-oncogene, serine/threonine kinase | 673 | up | 1,52 | 4,53E-02 |
| hsa05212 | Pancreatic cancer | AKT2 | AKT serine/threonine kinase 2 | 208 | down | 1,28 | 6,33E-03 |
| hsa05212 | Pancreatic cancer | PIK3CA | phosphatidylinositol-4,5-bisphosphate 3-kinase catalyti | 5290 | up | 1,33 | 3,17E-02 |
| hsa01230 | Biosynthesis of amino acids | BCAT2 | branched chain amino acid transaminase 2 | 587 | down | 1,34 | 1,96E-02 |
| hsa01230 | Biosynthesis of amino acids | PHGDH | phosphoglycerate dehydrogenase | 26227 | down | 2,34 | 2,54E-02 |
| hsa01230 | Biosynthesis of amino acids | MAT2A | methionine adenosyltransferase 2A | 4144 | down | 1,79 | 2,93E-02 |
| hsa01230 | Biosynthesis of amino acids | PYCR3 | pyrroline-5-carboxylate reductase 3 | 65263 | down | 1,77 | 1,11E-03 |
| hsa01230 | Biosynthesis of amino acids | TPI1 | triosephosphate isomerase 1 | 7167 | down | 1,28 | 3,58E-02 |
| hsa01230 | Biosynthesis of amino acids | NAGS | N-acetylglutamate synthase | 162417 | down | 1,62 | 4,90E-02 |
| hsa01230 | Biosynthesis of amino acids | GLUL | glutamate-ammonia ligase | 2752 | down | 1,47 | 6,91E-03 |
| hsa01230 | Biosynthesis of amino acids | PKM | pyruvate kinase M1/2 | 5315 | down | 1,35 | 4,16E-02 |
| hsa01230 | Biosynthesis of amino acids | PGAM1 | phosphoglycerate mutase 1 | 5223 | down | 1,62 | 5,67E-03 |
| hsa04022 | cGMP-PKG signaling pathway | ATF2 | activating transcription factor 2 | 1386 | up | 1,79 | 1,01E-02 |
| hsa04022 | cGMP-PKG signaling pathway | PPP3CC | protein phosphatase 3 catalytic subunit gamma | 5533 | up | 1,52 | 1,78E-02 |
| hsa04022 | cGMP-PKG signaling pathway | AKT2 | AKT serine/threonine kinase 2 | 208 | down | 1,28 | 6,33E-03 |
| hsa04022 | cGMP-PKG signaling pathway | ADRA2A | adrenoceptor alpha 2A | 150 | down | 3,92 | 2,45E-02 |
| hsa04022 | cGMP-PKG signaling pathway | PPP1CA | protein phosphatase 1 catalytic subunit alpha | 5499 | down | 1,24 | 9,77E-03 |
| hsa04022 | cGMP-PKG signaling pathway | CALM1 | calmodulin 1 | 801 | down | 1,43 | 2,23E-02 |
| hsa04022 | cGMP-PKG signaling pathway | GNA13 | G protein subunit alpha 13 | 10672 | up | 1,68 | 2,41E-03 |
| hsa04022 | cGMP-PKG signaling pathway | ATP2B1 | ATPase plasma membrane Ca2+ transporting 1 | 490 | up | 1,35 | 4,17E-02 |
| hsa04022 | cGMP-PKG signaling pathway | SRF | serum response factor | 6722 | up | 1,48 | 3,97E-02 |
| hsa04022 | cGMP-PKG signaling pathway | PPP1R12A | protein phosphatase 1 regulatory subunit 12A | 4659 | up | 1,25 | 1,03E-02 |
| hsa04022 | cGMP-PKG signaling pathway | GNA11 | G protein subunit alpha 11 | 2767 | down | 1,18 | 4,37E-02 |
| hsa04022 | cGMP-PKG signaling pathway | CALM2 | calmodulin 2 | 805 | down | 1,43 | 2,23E-02 |
| hsa04022 | cGMP-PKG signaling pathway | CREB3L2 | cAMP responsive element binding protein 3 like 2 | 64764 | up | 2,31 | 8,54E-03 |
| hsa04022 | cGMP-PKG signaling pathway | MEF2A | myocyte enhancer factor 2A | 4205 | up | 1,55 | 1,41E-02 |
| hsa04022 | cGMP-PKG signaling pathway | ATP2A3 | ATPase sarcoplasmic/endoplasmic reticulum Ca2+ tran | 489 | down | 1,62 | 4,34E-02 |
| hsa04022 | cGMP-PKG signaling pathway | IRS1 | insulin receptor substrate 1 | 3667 | down | 1,89 | 6,25E-03 |
| hsa04022 | cGMP-PKG signaling pathway | ADCY3 | adenylate cyclase 3 | 109 | down | 1,45 | 1,03E-02 |
| hsa04022 | cGMP-PKG signaling pathway | CALM3 | calmodulin 3 | 808 | down | 1,43 | 2,23E-02 |
| hsa04022 | cGMP-PKG signaling pathway | MEF2D | myocyte enhancer factor 2D | 4209 | up | 1,90 | 4,04E-02 |
| hsa04022 | cGMP-PKG signaling pathway | PPIF | peptidylprolyl isomerase F | 10105 | down | 1,66 | 7,16E-03 |
| hsa04022 | cGMP-PKG signaling pathway | CREB1 | cAMP responsive element binding protein 1 | 1385 | up | 1,51 | 7,18E-03 |
| hsa04022 | cGMP-PKG signaling pathway | IRS2 | insulin receptor substrate 2 | 8660 | up | 2,21 | 2,22E-02 |
| hsa04022 | cGMP-PKG signaling pathway | GNAI2 | G protein subunit alpha i2 | 2771 | down | 1,22 | 3,65E-02 |
| hsa04022 | cGMP-PKG signaling pathway | CREB5 | cAMP responsive element binding protein 5 | 9586 | up | 3,23 | 2,63E-03 |
| hsa04022 | cGMP-PKG signaling pathway | AKT1 | AKT serine/threonine kinase 1 | 207 | down | 1,37 | 1,42E-03 |
| hsa05215 | Prostate cancer | CREB1 | cAMP responsive element binding protein 1 | 1385 | up | 1,51 | 7,18E-03 |
| hsa05215 | Prostate cancer | CREB5 | cAMP responsive element binding protein 5 | 9586 | up | 3,23 | 2,63E-03 |
| hsa05215 | Prostate cancer | AKT1 | AKT serine/threonine kinase 1 | 207 | down | 1,37 | 1,42E-03 |
| hsa05215 | Prostate cancer | CDKN1A | cyclin dependent kinase inhibitor 1A | 1026 | up | 3,62 | 5,43E-03 |
| hsa05215 | Prostate cancer | ERBB2 | erb-b2 receptor tyrosine kinase 2 | 2064 | down | 1,47 | 1,93E-02 |
| hsa05215 | Prostate cancer | TCF7 | transcription factor 7 | 6932 | down | 1,47 | 1,07E-02 |
| hsa05215 | Prostate cancer | PTEN | phosphatase and tensin homolog | 5728 | up | 1,33 | 4,72E-02 |
| hsa05215 | Prostate cancer | HSP90B1 | heat shock protein 90 beta family member 1 | 7184 | up | 1,71 | 2,12E-02 |
| hsa05215 | Prostate cancer | CREB3L2 | cAMP responsive element binding protein 3 like 2 | 64764 | up | 2,31 | 8,54E-03 |
| hsa05215 | Prostate cancer | E2F1 | E2F transcription factor 1 | 1869 | down | 1,64 | 3,25E-02 |
| hsa05215 | Prostate cancer | SOS2 | SOS Ras/Rho guanine nucleotide exchange factor 2 | 6655 | up | 1,82 | 1,41E-02 |
| hsa05215 | Prostate cancer | HRAS | HRas proto-oncogene, GTPase | 3265 | down | 1,31 | 2,19E-03 |
| hsa05215 | Prostate cancer | PIK3R1 | phosphoinositide-3-kinase regulatory subunit 1 | 5295 | down | 2,04 | 8,40E-03 |
| hsa05215 | Prostate cancer | BRAF | B-Raf proto-oncogene, serine/threonine kinase | 673 | up | 1,52 | 4,53E-02 |
| hsa05215 | Prostate cancer | AKT2 | AKT serine/threonine kinase 2 | 208 | down | 1,28 | 6,33E-03 |
| hsa05215 | Prostate cancer | PIK3CA | phosphatidylinositol-4,5-bisphosphate 3-kinase catalyti | 5290 | up | 1,33 | 3,17E-02 |
| hsa05213 | Endometrial cancer | ERBB2 | erb-b2 receptor tyrosine kinase 2 | 2064 | down | 1,47 | 1,93E-02 |
| hsa05213 | Endometrial cancer | BRAF | B-Raf proto-oncogene, serine/threonine kinase | 673 | up | 1,52 | 4,53E-02 |
| hsa05213 | Endometrial cancer | TCF7 | transcription factor 7 | 6932 | down | 1,47 | 1,07E-02 |
| hsa05213 | Endometrial cancer | PTEN | phosphatase and tensin homolog | 5728 | up | 1,33 | 4,72E-02 |
| hsa05213 | Endometrial cancer | AKT2 | AKT serine/threonine kinase 2 | 208 | down | 1,28 | 6,33E-03 |

| | | | | | | | |
|---|---|---|---|---|---|---|---|
| hsa05213 | Endometrial cancer | PIK3CA | phosphatidylinositol-4,5-bisphosphate 3-kinase catalyti | 5290 | up | 1,33 | 3,17E-02 |
| hsa05213 | Endometrial cancer | AKT1 | AKT serine/threonine kinase 1 | 207 | down | 1,37 | 1,42E-03 |
| hsa05213 | Endometrial cancer | SOS2 | SOS Ras/Rho guanine nucleotide exchange factor 2 | 6655 | up | 1,82 | 1,41E-02 |
| hsa05213 | Endometrial cancer | HRAS | HRas proto-oncogene, GTPase | 3265 | down | 1,31 | 2,19E-03 |
| hsa05213 | Endometrial cancer | PIK3R1 | phosphoinositide-3-kinase regulatory subunit 1 | 5295 | down | 2,04 | 8,40E-03 |
| hsa05213 | Endometrial cancer | AXIN2 | axin 2 | 8313 | down | 2,79 | 3,67E-03 |
| hsa04071 | Sphingolipid signaling pathway | PTEN | phosphatase and tensin homolog | 5728 | up | 1,33 | 4,72E-02 |
| hsa04071 | Sphingolipid signaling pathway | BAX | BCL2 associated X, apoptosis regulator | 581 | down | 1,22 | 2,61E-02 |
| hsa04071 | Sphingolipid signaling pathway | BID | BH3 interacting domain death agonist | 637 | down | 1,40 | 2,42E-02 |
| hsa04071 | Sphingolipid signaling pathway | PPP2CB | protein phosphatase 2 catalytic subunit beta | 5516 | up | 1,30 | 4,79E-02 |
| hsa04071 | Sphingolipid signaling pathway | SMPD2 | sphingomyelin phosphodiesterase 2 | 6610 | down | 1,22 | 4,95E-02 |
| hsa04071 | Sphingolipid signaling pathway | GNAI2 | G protein subunit alpha i2 | 2771 | down | 1,22 | 3,65E-02 |
| hsa04071 | Sphingolipid signaling pathway | AKT1 | AKT serine/threonine kinase 1 | 207 | down | 1,37 | 1,42E-03 |
| hsa04071 | Sphingolipid signaling pathway | SPHK1 | sphingosine kinase 1 | 8877 | down | 2,00 | 2,34E-02 |
| hsa04071 | Sphingolipid signaling pathway | AKT2 | AKT serine/threonine kinase 2 | 208 | down | 1,28 | 6,33E-03 |
| hsa04071 | Sphingolipid signaling pathway | PIK3CA | phosphatidylinositol-4,5-bisphosphate 3-kinase catalyti | 5290 | up | 1,33 | 3,17E-02 |
| hsa04071 | Sphingolipid signaling pathway | TRAF2 | TNF receptor associated factor 2 | 7186 | down | 1,47 | 2,65E-02 |
| hsa04071 | Sphingolipid signaling pathway | S1PR2 | sphingosine-1-phosphate receptor 2 | 9294 | down | 1,91 | 1,41E-02 |
| hsa04071 | Sphingolipid signaling pathway | SGMS2 | sphingomyelin synthase 2 | 166929 | up | 1,39 | 4,00E-02 |
| hsa04071 | Sphingolipid signaling pathway | PPP2R2A | protein phosphatase 2 regulatory subunit Balpha | 5520 | up | 1,26 | 3,61E-02 |
| hsa04071 | Sphingolipid signaling pathway | PLD1 | phospholipase D1 | 5337 | up | 1,33 | 1,60E-02 |
| hsa04071 | Sphingolipid signaling pathway | HRAS | HRas proto-oncogene, GTPase | 3265 | down | 1,31 | 2,19E-03 |
| hsa04071 | Sphingolipid signaling pathway | GNA13 | G protein subunit alpha 13 | 10672 | up | 1,68 | 2,41E-03 |
| hsa04071 | Sphingolipid signaling pathway | PIK3R1 | phosphoinositide-3-kinase regulatory subunit 1 | 5295 | down | 2,04 | 8,40E-03 |
| hsa04071 | Sphingolipid signaling pathway | MAPK12 | mitogen-activated protein kinase 12 | 6300 | down | 1,37 | 3,32E-03 |
| hsa04071 | Sphingolipid signaling pathway | MAPK8 | mitogen-activated protein kinase 8 | 5599 | up | 1,38 | 3,05E-02 |
| hsa04919 | Thyroid hormone signaling pathway | RCAN1 | regulator of calcineurin 1 | 1827 | up | 1,47 | 4,96E-02 |
| hsa04919 | Thyroid hormone signaling pathway | AKT1 | AKT serine/threonine kinase 1 | 207 | down | 1,37 | 1,42E-03 |
| hsa04919 | Thyroid hormone signaling pathway | MED24 | mediator complex subunit 24 | 9862 | down | 1,33 | 2,93E-02 |
| hsa04919 | Thyroid hormone signaling pathway | NOTCH1 | notch 1 | 4851 | down | 1,62 | 2,46E-02 |
| hsa04919 | Thyroid hormone signaling pathway | RXRA | retinoid X receptor alpha | 6256 | down | 1,34 | 3,67E-03 |
| hsa04919 | Thyroid hormone signaling pathway | SLC9A1 | solute carrier family 9 member A1 | 6548 | up | 2,29 | 9,81E-03 |
| hsa04919 | Thyroid hormone signaling pathway | NCOA3 | nuclear receptor coactivator 3 | 8202 | up | 1,53 | 4,13E-02 |
| hsa04919 | Thyroid hormone signaling pathway | THRA | thyroid hormone receptor, alpha | 7067 | down | 1,45 | 2,63E-02 |
| hsa04919 | Thyroid hormone signaling pathway | BMP4 | bone morphogenetic protein 4 | 652 | down | 2,60 | 1,26E-02 |
| hsa04919 | Thyroid hormone signaling pathway | HRAS | HRas proto-oncogene, GTPase | 3265 | down | 1,31 | 2,19E-03 |
| hsa04919 | Thyroid hormone signaling pathway | SLC16A2 | solute carrier family 16 member 2 | 6567 | down | 3,17 | 2,47E-02 |
| hsa04919 | Thyroid hormone signaling pathway | PIK3R1 | phosphoinositide-3-kinase regulatory subunit 1 | 5295 | down | 2,04 | 8,40E-03 |
| hsa04919 | Thyroid hormone signaling pathway | HIF1A | hypoxia inducible factor 1 alpha subunit | 3091 | up | 1,23 | 4,66E-02 |
| hsa04919 | Thyroid hormone signaling pathway | ACTG1 | actin gamma 1 | 71 | up | 1,58 | 1,66E-02 |
| hsa04919 | Thyroid hormone signaling pathway | KAT2A | lysine acetyltransferase 2A | 2648 | down | 1,39 | 1,20E-02 |
| hsa04919 | Thyroid hormone signaling pathway | PLCG1 | phospholipase C gamma 1 | 5335 | down | 1,51 | 2,85E-02 |
| hsa04919 | Thyroid hormone signaling pathway | PFKFB2 | 6-phosphofructo-2-kinase/fructose-2,6-biphosphatase | 5208 | up | 1,39 | 4,32E-02 |
| hsa04919 | Thyroid hormone signaling pathway | PIK3CA | phosphatidylinositol-4,5-bisphosphate 3-kinase catalyti | 5290 | up | 1,33 | 3,17E-02 |
| hsa04919 | Thyroid hormone signaling pathway | AKT2 | AKT serine/threonine kinase 2 | 208 | down | 1,28 | 6,33E-03 |
| hsa00533 | Glycosaminoglycan biosynthesis - keratan sulfate | ST3GAL2 | ST3 beta-galactoside alpha-2,3-sialyltransferase 2 | 6483 | down | 1,66 | 3,84E-03 |
| hsa00533 | Glycosaminoglycan biosynthesis - keratan sulfate | B3GNT7 | UDP-GlcNAc:betaGal beta-1,3-N-acetylglucosaminyltra | 93010 | down | 2,40 | 3,48E-02 |
| hsa00533 | Glycosaminoglycan biosynthesis - keratan sulfate | B4GAT1 | beta-1,4-glucuronyltransferase 1 | 11041 | down | 2,42 | 1,49E-02 |
| hsa00533 | Glycosaminoglycan biosynthesis - keratan sulfate | B4GALT2 | beta-1,4-galactosyltransferase 2 | 8704 | down | 1,45 | 2,13E-03 |
| hsa04070 | Phosphatidylinositol signaling system | PIK3R1 | phosphoinositide-3-kinase regulatory subunit 1 | 5295 | down | 2,04 | 8,40E-03 |
| hsa04070 | Phosphatidylinositol signaling system | PIP4K2B | phosphatidylinositol-5-phosphate 4-kinase type 2 beta | 8396 | down | 1,53 | 1,01E-02 |
| hsa04070 | Phosphatidylinositol signaling system | ITPKC | inositol-trisphosphate 3-kinase C | 80271 | up | 1,63 | 3,32E-02 |
| hsa04070 | Phosphatidylinositol signaling system | CALM1 | calmodulin 1 | 801 | down | 1,43 | 2,23E-02 |
| hsa04070 | Phosphatidylinositol signaling system | CDIPT | CDP-diacylglycerol--inositol 3-phosphatidyltransferase | 10423 | down | 1,32 | 3,12E-03 |
| hsa04070 | Phosphatidylinositol signaling system | PIK3CA | phosphatidylinositol-4,5-bisphosphate 3-kinase catalyti | 5290 | up | 1,33 | 3,17E-02 |
| hsa04070 | Phosphatidylinositol signaling system | PIP5K1C | phosphatidylinositol-4-phosphate 5-kinase type 1 gam | 23396 | down | 1,39 | 1,49E-03 |
| hsa04070 | Phosphatidylinositol signaling system | PLCG1 | phospholipase C gamma 1 | 5335 | down | 1,51 | 2,85E-02 |
| hsa04070 | Phosphatidylinositol signaling system | PIP4P2 | phosphatidylinositol-4,5-bisphosphate 4-phosphatase | 55529 | up | 1,48 | 2,19E-02 |
| hsa04070 | Phosphatidylinositol signaling system | CALM3 | calmodulin 3 | 808 | down | 1,43 | 2,23E-02 |
| hsa04070 | Phosphatidylinositol signaling system | PIK3C2A | phosphatidylinositol-4-phosphate 3-kinase catalytic sub | 5286 | up | 1,30 | 1,79E-02 |
| hsa04070 | Phosphatidylinositol signaling system | ITPKB | inositol-trisphosphate 3-kinase B | 3707 | down | 1,71 | 1,97E-02 |
| hsa04070 | Phosphatidylinositol signaling system | MTMR6 | myotubularin related protein 6 | 9107 | up | 1,33 | 2,92E-02 |
| hsa04070 | Phosphatidylinositol signaling system | PIP5K1B | phosphatidylinositol-4-phosphate 5-kinase type 1 beta | 8395 | up | 3,40 | 1,84E-02 |
| hsa04070 | Phosphatidylinositol signaling system | PTEN | phosphatase and tensin homolog | 5728 | up | 1,33 | 4,72E-02 |
| hsa04070 | Phosphatidylinositol signaling system | CALM2 | calmodulin 2 | 805 | down | 1,43 | 2,23E-02 |
| hsa04070 | Phosphatidylinositol signaling system | MTMR2 | myotubularin related protein 2 | 8898 | up | 1,23 | 2,61E-02 |
| **hsa05210** | **Colorectal cancer** | **TGFBR2** | **transforming growth factor beta receptor 2** | **7048** | **down** | **1,34** | **2,74E-02** |
| **hsa05210** | **Colorectal cancer** | **RALGDS** | **ral guanine nucleotide dissociation stimulator** | **5900** | **down** | **1,30** | **3,49E-02** |
| **hsa05210** | **Colorectal cancer** | **AKT1** | **AKT serine/threonine kinase 1** | **207** | **down** | **1,37** | **1,42E-03** |
| **hsa05210** | **Colorectal cancer** | **AXIN2** | **axin 2** | **8313** | **down** | **2,79** | **3,67E-03** |
| **hsa05210** | **Colorectal cancer** | **PIK3R1** | **phosphoinositide-3-kinase regulatory subunit 1** | **5295** | **down** | **2,04** | **8,40E-03** |
| **hsa05210** | **Colorectal cancer** | **TCF7** | **transcription factor 7** | **6932** | **down** | **1,47** | **1,07E-02** |
| **hsa05210** | **Colorectal cancer** | **BAX** | **BCL2 associated X, apoptosis regulator** | **581** | **down** | **1,22** | **2,61E-02** |
| **hsa05210** | **Colorectal cancer** | **AKT2** | **AKT serine/threonine kinase 2** | **208** | **down** | **1,28** | **6,33E-03** |
| hsa04320 | Dorso-ventral axis formation | CPEB4 | cytoplasmic polyadenylation element binding protein 4 | 80315 | up | 2,70 | 6,54E-03 |
| hsa04320 | Dorso-ventral axis formation | SPIRE1 | spire type actin nucleation factor 1 | 56907 | up | 2,93 | 2,13E-03 |
| hsa04320 | Dorso-ventral axis formation | ETS1 | ETS proto-oncogene 1, transcription factor | 2113 | up | 2,07 | 1,22E-02 |
| hsa04320 | Dorso-ventral axis formation | CPEB3 | cytoplasmic polyadenylation element binding protein 3 | 22849 | up | 1,80 | 2,54E-02 |
| hsa04320 | Dorso-ventral axis formation | SOS2 | SOS Ras/Rho guanine nucleotide exchange factor 2 | 6655 | up | 1,82 | 1,41E-02 |
| hsa04915 | Estrogen signaling pathway | ADCY3 | adenylate cyclase 3 | 109 | down | 1,45 | 1,03E-02 |
| hsa04915 | Estrogen signaling pathway | CALM2 | calmodulin 2 | 805 | down | 1,43 | 2,23E-02 |
| hsa04915 | Estrogen signaling pathway | CREB3L2 | cAMP responsive element binding protein 3 like 2 | 64764 | up | 2,31 | 8,54E-03 |
| hsa04915 | Estrogen signaling pathway | HSP90B1 | heat shock protein 90 beta family member 1 | 7184 | up | 1,71 | 2,12E-02 |
| hsa04915 | Estrogen signaling pathway | CREB1 | cAMP responsive element binding protein 1 | 1385 | up | 1,51 | 7,18E-03 |
| hsa04915 | Estrogen signaling pathway | GNAI2 | G protein subunit alpha i2 | 2771 | down | 1,22 | 3,65E-02 |
| hsa04915 | Estrogen signaling pathway | CREB5 | cAMP responsive element binding protein 5 | 9586 | up | 3,23 | 2,63E-03 |
| hsa04915 | Estrogen signaling pathway | AKT1 | AKT serine/threonine kinase 1 | 207 | down | 1,37 | 1,42E-03 |
| hsa04915 | Estrogen signaling pathway | CALM3 | calmodulin 3 | 808 | down | 1,43 | 2,23E-02 |
| hsa04915 | Estrogen signaling pathway | HBEGF | heparin binding EGF like growth factor | 1839 | up | 3,67 | 5,67E-03 |
| hsa04915 | Estrogen signaling pathway | ATF2 | activating transcription factor 2 | 1386 | up | 1,79 | 1,01E-02 |
| hsa04915 | Estrogen signaling pathway | AKT2 | AKT serine/threonine kinase 2 | 208 | down | 1,28 | 6,33E-03 |
| hsa04915 | Estrogen signaling pathway | PIK3CA | phosphatidylinositol-4,5-bisphosphate 3-kinase catalyti | 5290 | up | 1,33 | 3,17E-02 |
| hsa04915 | Estrogen signaling pathway | SOS2 | SOS Ras/Rho guanine nucleotide exchange factor 2 | 6655 | up | 1,82 | 1,41E-02 |
| hsa04915 | Estrogen signaling pathway | HRAS | HRas proto-oncogene, GTPase | 3265 | down | 1,31 | 2,19E-03 |
| hsa04915 | Estrogen signaling pathway | CALM1 | calmodulin 1 | 801 | down | 1,43 | 2,23E-02 |
| hsa04915 | Estrogen signaling pathway | PIK3R1 | phosphoinositide-3-kinase regulatory subunit 1 | 5295 | down | 2,04 | 8,40E-03 |
