## Supplemental Table 6 for "Transcriptome profiling of human colonic cells exposed to the gut pathobiont *Streptococcus gallolyticus* subsp. *gallolyticus*"

**Supplemental Table S6. REACTOME**: List of 88 significantly up- and down-regulated pathways found by Reactome pathway analysis in HT29 cells after *SGG* UCN34 vs *SGM* during 24 h.

| Link to REACTOME Pathway | Pathway Description (REACTOME) | Nb Genes in Pathway | Nb Regulated Genes (Up / Down) | P-Value (All) | P-Value (Up) | P-Value (Down) | Min P-Value | Min BH P-Value |
|---|---|---|---|---|---|---|---|---|
| **R-HSA-381038** | **XBP1(S) activates chaperone genes** | **56** | **22 (18/4)** | **1,32E-07** | **3,04E-09** | **NA** | **3,04E-09** | **1,89E-06** |
| **R-HSA-6807878** | **COPI-mediated anterograde transport** | **98** | **28 (23/5)** | **2,92E-06** | **8,05E-09** | **NA** | **8,05E-09** | **2,49E-06** |
| **R-HSA-204005** | **COPII-mediated vesicle transport** | **68** | **20 (16/4)** | **6,87E-05** | **2,50E-06** | **NA** | **2,50E-06** | **5,17E-04** |
| R-HSA-73762 | RNA Polymerase I Transcription Initiation | 47 | 18 (5/13) | 3,73E-06 | NA | 4,64E-06 | 3,73E-06 | 1,20E-03 |
| R-HSA-73772 | RNA Polymerase I Promoter Escape | 30 | 14 (5/9) | 5,11E-06 | NA | 1,33E-04 | 5,11E-06 | 1,23E-03 |
| R-HSA-73863 | RNA Polymerase I Transcription Termination | 31 | 14 (4/10) | 7,95E-06 | NA | 2,39E-05 | 7,95E-06 | 1,53E-03 |
| R-HSA-2426168 | Activation of gene expression by SREBF (SREBP) | 42 | 16 (4/12) | 1,66E-05 | NA | 8,90E-06 | 8,90E-06 | 2,67E-03 |
| R-HSA-5694530 | **Cargo concentration in the ER** | 33 | 11 (10/1) | 1,85E-03 | 4,43E-05 | NA | 4,43E-05 | 6,85E-03 |
| R-HSA-749476 | RNA Polymerase III Abortive And Retractive Initiation | 42 | 15 (5/10) | 7,63E-05 | NA | 3,12E-04 | 7,63E-05 | 9,18E-03 |
| R-HSA-76066 | RNA Polymerase III Transcription Initiation From Type | 28 | 12 (4/8) | 8,24E-05 | NA | 5,42E-04 | 8,24E-05 | 8,81E-03 |
| R-HSA-381183 | ATF6 (ATF6-alpha) activates chaperone genes | 10 | 6 (6/0) | 2,28E-03 | 9,12E-05 | NA | 9,12E-05 | 1,12E-02 |
| R-HSA-76061 | RNA Polymerase III Transcription Initiation From Type | 29 | 12 (4/8) | 1,20E-04 | NA | 6,81E-04 | 1,20E-04 | 1,15E-02 |
| R-HSA-379716 | Cytosolic tRNA aminoacylation | 24 | 8 (8/0) | 1,10E-02 | 2,00E-04 | NA | 2,00E-04 | 2,05E-02 |
| R-HSA-176974 | Unwinding of DNA | 12 | 6 (0/6) | 5,98E-03 | NA | 2,52E-04 | 2,52E-04 | 3,88E-02 |
| R-HSA-191273 | Cholesterol biosynthesis | 20 | 7 (0/7) | 1,59E-02 | NA | 4,70E-04 | 4,70E-04 | 5,14E-02 |
| R-HSA-69166 | Removal of the Flap Intermediate | 14 | 6 (0/6) | 1,25E-02 | NA | 5,82E-04 | 5,82E-04 | 4,95E-02 |
| R-HSA-6782210 | Gap-filling DNA repair synthesis and ligation in TC-NE | 65 | 16 (4/12) | 3,25E-03 | NA | 6,08E-04 | 6,08E-04 | 4,66E-02 |
| R-HSA-6782135 | Dual incision in TC-NER | 66 | 16 (4/12) | 3,80E-03 | NA | 6,95E-04 | 6,95E-04 | 4,44E-02 |
| R-HSA-174437 | Removal of the Flap Intermediate from the C-strand | 10 | 5 (0/5) | 1,65E-02 | NA | 1,32E-03 | 1,32E-03 | 7,70E-02 |
| R-HSA-1368108 | BMAL1:CLOCK,NPAS2 activates circadian gene exp | 43 | 13 (9/4) | 1,48E-03 | 1,87E-03 | NA | 1,48E-03 | 1,22E-01 |
| R-HSA-442717 | No details found for 442717 | 6 | 5 (1/4) | 1,68E-03 | NA | 2,71E-03 | 1,68E-03 | 1,27E-01 |
| R-HSA-75109 | Triglyceride biosynthesis | 26 | 9 (2/7) | 4,71E-03 | NA | 2,12E-03 | 2,12E-03 | 1,12E-01 |
| R-HSA-76071 | RNA Polymerase III Transcription Initiation From Type | 29 | 10 (4/6) | 2,58E-03 | NA | 1,77E-02 | 2,58E-03 | 1,53E-01 |
| R-HSA-73780 | RNA Polymerase III Chain Elongation | 19 | 8 (3/5) | 2,60E-03 | NA | 1,66E-02 | 2,60E-03 | 1,45E-01 |
| R-HSA-73980 | RNA Polymerase III Transcription Termination | 24 | 9 (3/6) | 2,69E-03 | NA | 7,92E-03 | 2,69E-03 | 1,42E-01 |
| R-HSA-69052 | Switching of origins to a post-replicative state | 6 | 4 (0/4) | 1,94E-02 | NA | 2,71E-03 | 2,71E-03 | 1,32E-01 |
| R-HSA-1474151 | Tetrahydrobiopterin (BH4) synthesis, recycling, salvag | 12 | 5 (0/5) | 3,28E-02 | NA | 2,87E-03 | 2,87E-03 | 1,31E-01 |
| R-HSA-2151201 | Transcriptional activation of mitochondrial biogenesis | 41 | 12 (6/6) | 3,26E-03 | NA | NA | 3,26E-03 | 1,53E-01 |
| R-HSA-5675221 | **Negative regulation of MAPK pathway** | 38 | 11 (8/3) | 5,74E-03 | 3,79E-03 | NA | 3,79E-03 | 2,07E-01 |
| R-HSA-5218921 | VEGFR2 mediated cell proliferation | 21 | 7 (1/6) | 2,03E-02 | NA | 4,33E-03 | 4,33E-03 | 1,81E-01 |
| R-HSA-5651801 | PCNA-Dependent Long Patch Base Excision Repair | 21 | 8 (2/6) | 4,96E-03 | NA | 4,33E-03 | 4,33E-03 | 1,81E-01 |
| R-HSA-400253 | Circadian Clock | 49 | 13 (9/4) | 4,88E-03 | 4,39E-03 | NA | 4,39E-03 | 1,93E-01 |
| R-HSA-5696400 | Dual Incision in GG-NER | 41 | 10 (2/8) | 2,79E-02 | NA | 5,62E-03 | 5,62E-03 | 2,18E-01 |
| R-HSA-72165 | mRNA Splicing - Minor Pathway | 52 | 9 (0/9) | NA | NA | 6,07E-03 | 6,07E-03 | 2,22E-01 |
| R-HSA-68962 | Activation of the pre-replicative complex | 32 | 7 (0/7) | NA | NA | 6,36E-03 | 6,36E-03 | 2,21E-01 |
| R-HSA-203927 | MicroRNA (miRNA) biogenesis | 23 | 7 (1/6) | 3,13E-02 | NA | 6,55E-03 | 6,55E-03 | 2,18E-01 |
| R-HSA-111997 | CaM pathway | 3 | 3 (0/3) | 3,23E-02 | NA | 8,31E-03 | 8,31E-03 | 2,47E-01 |
| R-HSA-381070 | IRE1alpha activates chaperones | 3 | 3 (3/0) | 3,23E-02 | 8,45E-03 | NA | 8,45E-03 | 4,09E-01 |
| R-HSA-1834949 | Cytosolic sensors of pathogen-associated DNA | 23 | 8 (3/5) | 8,63E-03 | NA | 3,21E-02 | 8,63E-03 | 2,75E-01 |
| R-HSA-380994 | ATF4 activates genes | 25 | 8 (6/2) | 1,39E-02 | 9,80E-03 | NA | 9,80E-03 | 3,64E-01 |
| R-HSA-111932 | CaMK IV-mediated phosphorylation of CREB | 5 | 4 (1/3) | 1,05E-02 | NA | 2,57E-02 | 1,05E-02 | 3,16E-01 |
| R-HSA-69205 | G1/S-Specific Transcription | 17 | 5 (0/5) | NA | NA | 1,11E-02 | 1,11E-02 | 3,06E-01 |
| R-HSA-113510 | E2F mediated regulation of DNA replication | 17 | 5 (0/5) | NA | NA | 1,11E-02 | 1,11E-02 | 3,06E-01 |
| R-HSA-180024 | DARPP-32 events | 26 | 8 (2/6) | 1,74E-02 | NA | 1,12E-02 | 1,12E-02 | 2,98E-01 |
| R-HSA-983170 | Antigen Presentation: Folding, assembly and peptide | 26 | 6 (6/0) | NA | 1,16E-02 | NA | 1,16E-02 | 4,53E-01 |
| R-HSA-77075 | RNA Pol II CTD phosphorylation and interaction with | 27 | 8 (2/6) | 2,13E-02 | NA | 1,31E-02 | 1,31E-02 | 3,30E-01 |
| R-HSA-167160 | RNA Pol II CTD phosphorylation and interaction with | 27 | 8 (2/6) | 2,13E-02 | NA | 1,31E-02 | 1,31E-02 | 3,30E-01 |
| R-HSA-69183 | Processive synthesis on the lagging strand | 10 | 4 (0/4) | NA | NA | 1,38E-02 | 1,38E-02 | 3,34E-01 |
| R-HSA-983168 | Antigen processing: Ubiquitination & Proteasome deg | 308 | 27 (27/0) | NA | 1,58E-02 | NA | 1,58E-02 | 5,34E-01 |
| R-HSA-6807062 | Cholesterol biosynthesis via lathosterol | 4 | 3 (0/3) | NA | NA | 1,60E-02 | 1,60E-02 | 3,64E-01 |
| R-HSA-442745 | No details found for 442745 | 4 | 3 (0/3) | NA | NA | 1,60E-02 | 1,60E-02 | 3,64E-01 |
| R-HSA-6807047 | Cholesterol biosynthesis via desmosterol | 4 | 3 (0/3) | NA | NA | 1,60E-02 | 1,60E-02 | 3,64E-01 |
| R-HSA-5218920 | VEGFR2 mediated vascular permeability | 29 | 6 (0/6) | NA | NA | 1,77E-02 | 1,77E-02 | 3,74E-01 |
| R-HSA-72086 | mRNA Capping | 29 | 8 (2/6) | 3,11E-02 | NA | 1,77E-02 | 1,77E-02 | 3,74E-01 |
| R-HSA-1368082 | RORA activates gene expression | 27 | 8 (4/4) | 2,13E-02 | NA | NA | 2,13E-02 | 4,40E-01 |
| R-HSA-203615 | eNOS activation | 12 | 4 (0/4) | NA | NA | 2,35E-02 | 2,35E-02 | 4,52E-01 |
| R-HSA-174417 | Telomere C-strand (Lagging Strand) Synthesis | 5 | 3 (0/3) | NA | NA | 2,57E-02 | 2,57E-02 | 4,73E-01 |
| R-HSA-1483166 | Synthesis of PA | 32 | 6 (0/6) | NA | NA | 2,64E-02 | 2,64E-02 | 4,71E-01 |
| R-HSA-450513 | Tristetraprolin (TTP, ZFP36) binds and destabilizes m | 17 | 6 (4/2) | 2,96E-02 | NA | NA | 2,96E-02 | 5,34E-01 |
| R-HSA-442729 | CREB1 phosphorylation through the activation of Ca | 17 | 6 (3/3) | 2,96E-02 | NA | NA | 2,96E-02 | 5,34E-01 |
| R-HSA-113418 | Formation of the Early Elongation Complex | 33 | 6 (0/6) | NA | NA | 2,98E-02 | 2,98E-02 | 5,03E-01 |
| R-HSA-167158 | Formation of the HIV-1 Early Elongation Complex | 33 | 6 (0/6) | NA | NA | 2,98E-02 | 2,98E-02 | 5,03E-01 |
| R-HSA-112409 | RAF-independent MAPK1/3 activation | 13 | 5 (4/1) | 4,34E-02 | 2,99E-02 | NA | 2,99E-02 | 6,23E-01 |
| R-HSA-6781827 | Transcription-Coupled Nucleotide Excision Repair (TC | 45 | 10 (3/7) | 4,80E-02 | NA | 3,19E-02 | 3,19E-02 | 5,16E-01 |
| R-HSA-1989781 | PPARA activates gene expression | 113 | 20 (11/9) | 3,27E-02 | NA | NA | 3,27E-02 | 5,34E-01 |
| R-HSA-5626467 | RHO GTPases activate IQGAPs | 34 | 6 (0/6) | NA | NA | 3,34E-02 | 3,34E-02 | 5,14E-01 |
| R-HSA-174411 | Polymerase switching on the C-strand of the telomere | 14 | 4 (0/4) | NA | NA | 3,59E-02 | 3,59E-02 | 5,30E-01 |
| R-HSA-156581 | Methylation | 14 | 4 (0/4) | NA | NA | 3,59E-02 | 3,59E-02 | 5,30E-01 |
| R-HSA-418359 | Reduction of cytosolic Ca++ levels | 14 | 4 (0/4) | NA | NA | 3,59E-02 | 3,59E-02 | 5,30E-01 |
| R-HSA-69091 | Polymerase switching | 14 | 4 (0/4) | NA | NA | 3,59E-02 | 3,59E-02 | 5,30E-01 |
| R-HSA-174414 | Processive synthesis on the C-strand of the telomere | 6 | 3 (0/3) | NA | NA | 3,73E-02 | 3,73E-02 | 5,34E-01 |
| R-HSA-111957 | Cam-PDE 1 activation | 6 | 3 (0/3) | NA | NA | 3,73E-02 | 3,73E-02 | 5,34E-01 |
| R-HSA-171319 | Telomere Extension By Telomerase | 6 | 3 (0/3) | NA | NA | 3,73E-02 | 3,73E-02 | 5,34E-01 |
| R-HSA-5674400 | Constitutive Signaling by AKT1 E17K in Cancer | 25 | 7 (2/5) | 4,57E-02 | NA | 4,23E-02 | 4,23E-02 | 5,71E-01 |
| R-HSA-5696397 | Gap-filling DNA repair synthesis and ligation in GG-NE | 25 | 5 (0/5) | NA | NA | 4,23E-02 | 4,23E-02 | 5,71E-01 |
| R-HSA-75105 | Fatty acyl-CoA biosynthesis | 25 | 5 (0/5) | NA | NA | 4,23E-02 | 4,23E-02 | 5,71E-01 |
| R-HSA-1169408 | ISG15 antiviral mechanism | 73 | 9 (9/0) | NA | 4,27E-02 | NA | 4,27E-02 | 8,35E-01 |
| R-HSA-68867 | Assembly of the pre-replicative complex | 15 | 4 (0/4) | NA | NA | 4,31E-02 | 4,31E-02 | 5,70E-01 |
| R-HSA-5358565 | Mismatch repair (MMR) directed by MSH2:MSH6 (Mut | 15 | 4 (0/4) | NA | NA | 4,31E-02 | 4,31E-02 | 5,70E-01 |
| R-HSA-111447 | Activation of BAD and translocation to mitochondria | 15 | 4 (0/4) | NA | NA | 4,31E-02 | 4,31E-02 | 5,70E-01 |
| R-HSA-5358606 | Mismatch repair (MMR) directed by MSH2:MSH3 (Mut | 15 | 4 (0/4) | NA | NA | 4,31E-02 | 4,31E-02 | 5,70E-01 |
| R-HSA-1358803 | Downregulation of ERBB2:ERBB3 signaling | 13 | 5 (2/3) | 4,34E-02 | NA | NA | 4,34E-02 | 6,23E-01 |
| R-HSA-176187 | Activation of ATR in response to replication stress | 37 | 6 (0/6) | NA | NA | 4,60E-02 | 4,60E-02 | 5,85E-01 |
| R-HSA-139915 | Activation of PUMA and translocation to mitochondria | 8 | 4 (1/3) | 4,62E-02 | NA | NA | 4,62E-02 | 6,30E-01 |
| R-HSA-199418 | Negative regulation of the PI3K/AKT network | 8 | 4 (2/2) | 4,62E-02 | NA | NA | 4,62E-02 | 6,30E-01 |
| R-HSA-69231 | Cyclin D associated events in G1 | 38 | 9 (5/4) | 4,64E-02 | NA | NA | 4,64E-02 | 6,24E-01 |
| R-HSA-442982 | Ras activation upon Ca2+ influx through NMDA recep | 19 | 6 (2/4) | 4,65E-02 | NA | NA | 4,65E-02 | 6,16E-01 |
| R-HSA-2682334 | EPH-Ephrin signaling | 26 | 5 (0/5) | NA | NA | 4,79E-02 | 4,79E-02 | 5,92E-01 |

***Regulated Genes in Pathways***

| Link to REACTOME Pathway | Pathway Description (REACTOME) | Gene Symbol | Gene Name | Link to Entrez Gene | Regulation | Fold-Change | Adjusted P-Value |
|---|---|---|---|---|---|---|---|
| R-HSA-381038 | XBP1(S) activates chaperone genes | HYOU1 | hypoxia up-regulated 1 | 10525 | up | 2,96 | 5,94E-03 |
| R-HSA-381038 | XBP1(S) activates chaperone genes | SEC31A | SEC31 homolog A, COPII coat complex | 22872 | up | 1,53 | 3,02E-02 |
| R-HSA-381038 | XBP1(S) activates chaperone genes | GFPT1 | glutamine--fructose-6-phosphate transa | 2673 | up | 2,92 | 2,14E-03 |
| R-HSA-381038 | XBP1(S) activates chaperone genes | TSPYL2 | TSPY like 2 | 64061 | up | 2,24 | 3,18E-02 |
| R-HSA-381038 | XBP1(S) activates chaperone genes | DNAJB9 | DnaJ heat shock protein family (Hsp40) | 4189 | up | 7,76 | 1,30E-04 |
| R-HSA-381038 | XBP1(S) activates chaperone genes | DNAJC3 | DnaJ heat shock protein family (Hsp40) | 5611 | up | 2,75 | 2,24E-03 |
| R-HSA-381038 | XBP1(S) activates chaperone genes | SEC63 | SEC63 homolog, protein translocation r | 11231 | up | 1,60 | 8,90E-03 |
| R-HSA-381038 | XBP1(S) activates chaperone genes | KLHDC3 | kelch domain containing 3 | 116138 | down | 1,49 | 2,19E-03 |
| R-HSA-381038 | XBP1(S) activates chaperone genes | WIPI1 | WD repeat domain, phosphoinositide in | 55062 | up | 3,38 | 2,86E-03 |
| R-HSA-381038 | XBP1(S) activates chaperone genes | CXXC1 | CXXC finger protein 1 | 30827 | down | 1,21 | 2,87E-02 |
| R-HSA-381038 | XBP1(S) activates chaperone genes | CUL7 | cullin 7 | 9820 | up | 1,34 | 4,05E-02 |
| R-HSA-381038 | XBP1(S) activates chaperone genes | XBP1 | X-box binding protein 1 | 7494 | up | 2,00 | 5,67E-03 |
| R-HSA-381038 | XBP1(S) activates chaperone genes | TATDN2 | TatD DNase domain containing 2 | 9797 | down | 1,27 | 4,26E-02 |
| R-HSA-381038 | XBP1(S) activates chaperone genes | DNAJB11 | DnaJ heat shock protein family (Hsp40) | 51726 | up | 2,08 | 1,17E-02 |
| R-HSA-381038 | XBP1(S) activates chaperone genes | SSR1 | signal sequence receptor subunit 1 | 6745 | up | 1,73 | 2,06E-02 |
| R-HSA-381038 | XBP1(S) activates chaperone genes | EDEM1 | ER degradation enhancing alpha-mann | 9695 | up | 2,16 | 1,32E-02 |
| R-HSA-381038 | XBP1(S) activates chaperone genes | GOSR2 | golgi SNAP receptor complex member 2 | 9570 | up | 1,55 | 2,21E-02 |
| R-HSA-381038 | XBP1(S) activates chaperone genes | DDX11 | DEAD/H-box helicase 11 | 1663 | down | 1,61 | 1,78E-02 |
| R-HSA-381038 | XBP1(S) activates chaperone genes | SERP1 | stress associated endoplasmic reticulum | 27230 | up | 1,95 | 1,27E-02 |
| R-HSA-381038 | XBP1(S) activates chaperone genes | FKBP14 | FK506 binding protein 14 | 55033 | up | 3,26 | 1,30E-03 |
| R-HSA-381038 | XBP1(S) activates chaperone genes | SYVN1 | synoviolin 1 | 84447 | up | 2,33 | 1,93E-02 |
| R-HSA-381038 | XBP1(S) activates chaperone genes | PREB | prolactin regulatory element binding | 10113 | up | 1,76 | 3,49E-02 |
| R-HSA-6807878 | COPI-mediated anterograde transport | COG5 | component of oligomeric golgi complex | 10466 | up | 1,40 | 3,36E-03 |
| R-HSA-6807878 | COPI-mediated anterograde transport | GOLGB1 | golgin B1 | 2804 | up | 1,62 | 4,76E-04 |
| R-HSA-6807878 | COPI-mediated anterograde transport | RAB1B | RAB1B, member RAS oncogene family | 81876 | down | 1,27 | 4,09E-04 |
| R-HSA-6807878 | COPI-mediated anterograde transport | COPB1 | coatomer protein complex subunit beta | 1315 | up | 1,50 | 1,68E-02 |
| R-HSA-6807878 | COPI-mediated anterograde transport | TMED7 | transmembrane p24 trafficking protein 7 | 51014 | up | 2,23 | 4,98E-02 |
| R-HSA-6807878 | COPI-mediated anterograde transport | DCTN4 | dynactin subunit 4 | 51164 | up | 1,51 | 1,31E-02 |
| R-HSA-6807878 | COPI-mediated anterograde transport | ARFGAP3 | ADP ribosylation factor GTPase activati | 26286 | up | 2,76 | 2,05E-03 |
| R-HSA-6807878 | COPI-mediated anterograde transport | TUBA4A | tubulin alpha 4a | 7277 | down | 1,63 | 2,97E-02 |
| R-HSA-6807878 | COPI-mediated anterograde transport | DYNC1LI2 | dynein cytoplasmic 1 light intermediate | 1783 | up | 1,21 | 2,47E-02 |
| R-HSA-6807878 | COPI-mediated anterograde transport | DYNC2H1 | dynein cytoplasmic 2 heavy chain 1 | 79659 | up | 1,52 | 9,51E-03 |
| R-HSA-6807878 | COPI-mediated anterograde transport | COG3 | component of oligomeric golgi complex | 83548 | up | 1,82 | 2,63E-03 |
| R-HSA-6807878 | COPI-mediated anterograde transport | COPG1 | coatomer protein complex subunit gamm | 22820 | up | 1,50 | 4,74E-02 |
| R-HSA-6807878 | COPI-mediated anterograde transport | TMEM115 | transmembrane protein 115 | 11070 | down | 1,26 | 8,31E-03 |
| R-HSA-6807878 | COPI-mediated anterograde transport | GOSR1 | golgi SNAP receptor complex member 1 | 9527 | up | 1,39 | 3,34E-02 |
| R-HSA-6807878 | COPI-mediated anterograde transport | BET1 | Bet1 golgi vesicular membrane traffickin | 10282 | up | 2,34 | 3,79E-03 |
| R-HSA-6807878 | COPI-mediated anterograde transport | ARF4 | ADP ribosylation factor 4 | 378 | up | 1,69 | 2,56E-02 |
| R-HSA-6807878 | COPI-mediated anterograde transport | CD55 | CD55 molecule (Cromer blood group) | 1604 | up | 3,43 | 2,24E-03 |
| R-HSA-6807878 | COPI-mediated anterograde transport | GOSR2 | golgi SNAP receptor complex member 2 | 9570 | up | 1,55 | 2,21E-02 |
| R-HSA-6807878 | COPI-mediated anterograde transport | GOLGA2 | golgin A2 | 2801 | up | 1,85 | 1,27E-02 |
| R-HSA-6807878 | COPI-mediated anterograde transport | ARCN1 | archain 1 | 372 | up | 1,45 | 2,54E-02 |
| R-HSA-6807878 | COPI-mediated anterograde transport | NAPG | NSF attachment protein gamma | 8774 | up | 1,33 | 1,69E-02 |
| R-HSA-6807878 | COPI-mediated anterograde transport | STX5 | syntaxin 5 | 6811 | up | 1,66 | 2,40E-02 |
| R-HSA-6807878 | COPI-mediated anterograde transport | SPTBN1 | spectrin beta, non-erythrocytic 1 | 6711 | up | 1,42 | 1,66E-02 |
| R-HSA-6807878 | COPI-mediated anterograde transport | ARF3 | ADP ribosylation factor 3 | 377 | down | 1,30 | 4,57E-02 |
| R-HSA-6807878 | COPI-mediated anterograde transport | TUBA1B | tubulin alpha 1b | 10376 | down | 1,78 | 2,30E-02 |
| R-HSA-6807878 | COPI-mediated anterograde transport | DYNC2LI1 | dynein cytoplasmic 2 light intermediate | 51626 | up | 1,40 | 3,09E-02 |
| R-HSA-6807878 | COPI-mediated anterograde transport | USO1 | USO1 vesicle transport factor | 8615 | up | 1,77 | 6,10E-03 |
| R-HSA-6807878 | COPI-mediated anterograde transport | COG1 | component of oligomeric golgi complex | 9382 | up | 1,38 | 4,79E-02 |
| R-HSA-204005 | COPII-mediated vesicle transport | USO1 | USO1 vesicle transport factor | 8615 | up | 1,77 | 6,10E-03 |
| R-HSA-204005 | COPII-mediated vesicle transport | LMAN1 | lectin, mannose binding 1 | 3998 | up | 1,54 | 4,14E-02 |
| R-HSA-204005 | COPII-mediated vesicle transport | NAPG | NSF attachment protein gamma | 8774 | up | 1,33 | 1,69E-02 |
| R-HSA-204005 | COPII-mediated vesicle transport | TFG | TRK-fused gene | 10342 | up | 1,49 | 1,96E-02 |
| R-HSA-204005 | COPII-mediated vesicle transport | PREB | prolactin regulatory element binding | 10113 | up | 1,76 | 3,49E-02 |
| R-HSA-204005 | COPII-mediated vesicle transport | SCFD1 | sec1 family domain containing 1 | 23256 | up | 1,54 | 1,05E-02 |
| R-HSA-204005 | COPII-mediated vesicle transport | STX5 | syntaxin 5 | 6811 | up | 1,66 | 2,40E-02 |
| R-HSA-204005 | COPII-mediated vesicle transport | GOSR2 | golgi SNAP receptor complex member 2 | 9570 | up | 1,55 | 2,21E-02 |
| R-HSA-204005 | COPII-mediated vesicle transport | GOLGA2 | golgin A2 | 2801 | up | 1,85 | 1,27E-02 |
| R-HSA-204005 | COPII-mediated vesicle transport | SEC24D | SEC24 homolog D, COPII coat complex | 9871 | up | 4,45 | 6,80E-04 |
| R-HSA-204005 | COPII-mediated vesicle transport | AREG | amphiregulin | 374 | up | 3,33 | 2,37E-02 |
| R-HSA-204005 | COPII-mediated vesicle transport | BET1 | Bet1 golgi vesicular membrane traffickin | 10282 | up | 2,34 | 3,79E-03 |
| R-HSA-204005 | COPII-mediated vesicle transport | TRAPPC1 | trafficking protein particle complex 1 | 58485 | down | 1,31 | 2,30E-02 |
| R-HSA-204005 | COPII-mediated vesicle transport | TRAPPC6B | trafficking protein particle complex 6B | 122553 | up | 1,70 | 3,07E-02 |
| R-HSA-204005 | COPII-mediated vesicle transport | CNIH2 | cornichon family AMPA receptor auxiliar | 254263 | down | 2,64 | 1,05E-02 |
| R-HSA-204005 | COPII-mediated vesicle transport | SEC24A | SEC24 homolog A, COPII coat complex | 10802 | up | 1,71 | 3,95E-03 |
| R-HSA-204005 | COPII-mediated vesicle transport | SEC23A | Sec23 homolog A, coat complex II comp | 10484 | up | 1,46 | 2,47E-02 |
| R-HSA-204005 | COPII-mediated vesicle transport | TRAPPC2L | trafficking protein particle complex 2 like | 51693 | down | 1,32 | 2,18E-02 |
| R-HSA-204005 | COPII-mediated vesicle transport | SEC31A | SEC31 homolog A, COPII coat complex | 22872 | up | 1,53 | 3,02E-02 |
| R-HSA-204005 | COPII-mediated vesicle transport | RAB1B | RAB1B, member RAS oncogene family | 81876 | down | 1,27 | 4,09E-04 |
| R-HSA-73762 | RNA Polymerase I Transcription Initiation | UBTF | upstream binding transcription factor, RN | 7343 | down | 1,66 | 1,20E-02 |
| R-HSA-73762 | RNA Polymerase I Transcription Initiation | CDK7 | cyclin dependent kinase 7 | 1022 | up | 1,37 | 4,38E-02 |
| R-HSA-73762 | RNA Polymerase I Transcription Initiation | CHD4 | chromodomain helicase DNA binding pr | 1108 | down | 1,44 | 9,37E-03 |
| R-HSA-73762 | RNA Polymerase I Transcription Initiation | KAT2A | lysine acetyltransferase 2A | 2648 | down | 1,39 | 1,20E-02 |
| R-HSA-73762 | RNA Polymerase I Transcription Initiation | POLR1A | RNA polymerase I subunit A | 25885 | down | 1,30 | 4,32E-02 |
| R-HSA-73762 | RNA Polymerase I Transcription Initiation | POLR2L | RNA polymerase II subunit L | 5441 | down | 1,52 | 3,58E-03 |
| R-HSA-73762 | RNA Polymerase I Transcription Initiation | GTF2H1 | general transcription factor IIH subunit 1 | 2965 | up | 1,33 | 1,85E-02 |
| R-HSA-73762 | RNA Polymerase I Transcription Initiation | TAF1D | TATA-box binding protein associated fa | 79101 | up | 1,87 | 1,00E-02 |
| R-HSA-73762 | RNA Polymerase I Transcription Initiation | MBD3 | methyl-CpG binding domain protein 3 | 53615 | down | 1,52 | 4,26E-04 |
| R-HSA-73762 | RNA Polymerase I Transcription Initiation | MTA2 | metastasis associated 1 family member | 9219 | down | 1,27 | 2,45E-02 |
| R-HSA-73762 | RNA Polymerase I Transcription Initiation | ERCC2 | ERCC excision repair 2, TFIIH core com | 2068 | down | 1,29 | 2,20E-02 |
| R-HSA-73762 | RNA Polymerase I Transcription Initiation | ERCC3 | ERCC excision repair 3, TFIIH core com | 2071 | down | 1,20 | 2,37E-02 |
| R-HSA-73762 | RNA Polymerase I Transcription Initiation | TAF1A | TATA-box binding protein associated fa | 9015 | up | 2,20 | 2,24E-03 |
| R-HSA-73762 | RNA Polymerase I Transcription Initiation | CD3EAP | CD3e molecule associated protein | 10849 | down | 1,57 | 3,12E-03 |
| R-HSA-73762 | RNA Polymerase I Transcription Initiation | RRN3 | RRN3 homolog, RNA polymerase I trans | 54700 | up | 1,41 | 4,24E-02 |
| R-HSA-73762 | RNA Polymerase I Transcription Initiation | POLR2F | RNA polymerase II subunit F | 5435 | down | 1,41 | 3,22E-02 |
| R-HSA-73762 | RNA Polymerase I Transcription Initiation | POLR2E | RNA polymerase II subunit E | 5434 | down | 1,31 | 3,18E-03 |
| R-HSA-73762 | RNA Polymerase I Transcription Initiation | POLR1C | RNA polymerase I subunit C | 9533 | down | 1,24 | 4,71E-02 |
| R-HSA-73772 | RNA Polymerase I Promoter Escape | CDK7 | cyclin dependent kinase 7 | 1022 | up | 1,37 | 4,38E-02 |
| R-HSA-73772 | RNA Polymerase I Promoter Escape | UBTF | upstream binding transcription factor, RN | 7343 | down | 1,66 | 1,20E-02 |
| R-HSA-73772 | RNA Polymerase I Promoter Escape | ERCC3 | ERCC excision repair 3, TFIIH core com | 2071 | down | 1,20 | 2,37E-02 |
| R-HSA-73772 | RNA Polymerase I Promoter Escape | ERCC2 | ERCC excision repair 2, TFIIH core com | 2068 | down | 1,29 | 2,20E-02 |
| R-HSA-73772 | RNA Polymerase I Promoter Escape | CD3EAP | CD3e molecule associated protein | 10849 | down | 1,57 | 3,12E-03 |
| R-HSA-73772 | RNA Polymerase I Promoter Escape | TAF1A | TATA-box binding protein associated fa | 9015 | up | 2,20 | 2,24E-03 |
| R-HSA-73772 | RNA Polymerase I Promoter Escape | RRN3 | RRN3 homolog, RNA polymerase I trans | 54700 | up | 1,41 | 4,24E-02 |
| R-HSA-73772 | RNA Polymerase I Promoter Escape | POLR2L | RNA polymerase II subunit L | 5441 | down | 1,52 | 3,58E-03 |
| R-HSA-73772 | RNA Polymerase I Promoter Escape | POLR1A | RNA polymerase I subunit A | 25885 | down | 1,30 | 4,32E-02 |
| R-HSA-73772 | RNA Polymerase I Promoter Escape | POLR2F | RNA polymerase II subunit F | 5435 | down | 1,41 | 3,22E-02 |
| R-HSA-73772 | RNA Polymerase I Promoter Escape | GTF2H1 | general transcription factor IIH subunit 1 | 2965 | up | 1,33 | 1,85E-02 |
| R-HSA-73772 | RNA Polymerase I Promoter Escape | TAF1D | TATA-box binding protein associated fa | 79101 | up | 1,87 | 1,00E-02 |
| R-HSA-73772 | RNA Polymerase I Promoter Escape | POLR2E | RNA polymerase II subunit E | 5434 | down | 1,31 | 3,18E-03 |
| R-HSA-73772 | RNA Polymerase I Promoter Escape | POLR1C | RNA polymerase I subunit C | 9533 | down | 1,24 | 4,71E-02 |
| R-HSA-73863 | RNA Polymerase I Transcription Termination | CD3EAP | CD3e molecule associated protein | 10849 | down | 1,57 | 3,12E-03 |
| R-HSA-73863 | RNA Polymerase I Transcription Termination | TAF1A | TATA-box binding protein associated fa | 9015 | up | 2,20 | 2,24E-03 |
| R-HSA-73863 | RNA Polymerase I Transcription Termination | POLR2F | RNA polymerase II subunit F | 5435 | down | 1,41 | 3,22E-02 |
| R-HSA-73863 | RNA Polymerase I Transcription Termination | GTF2H1 | general transcription factor IIH subunit 1 | 2965 | up | 1,33 | 1,85E-02 |
| R-HSA-73863 | RNA Polymerase I Transcription Termination | POLR2L | RNA polymerase II subunit L | 5441 | down | 1,52 | 3,58E-03 |
| R-HSA-73863 | RNA Polymerase I Transcription Termination | POLR1A | RNA polymerase I subunit A | 25885 | down | 1,30 | 4,32E-02 |
| R-HSA-73863 | RNA Polymerase I Transcription Termination | POLR2E | RNA polymerase II subunit E | 5434 | down | 1,31 | 3,18E-03 |
| R-HSA-73863 | RNA Polymerase I Transcription Termination | TAF1D | TATA-box binding protein associated fa | 79101 | up | 1,87 | 1,00E-02 |
| R-HSA-73863 | RNA Polymerase I Transcription Termination | CAVIN1 | caveolae associated protein 1 | 284119 | down | 1,37 | 3,95E-02 |
| R-HSA-73863 | RNA Polymerase I Transcription Termination | POLR1C | RNA polymerase I subunit C | 9533 | down | 1,24 | 4,71E-02 |
| R-HSA-73863 | RNA Polymerase I Transcription Termination | CDK7 | cyclin dependent kinase 7 | 1022 | up | 1,37 | 4,38E-02 |
| R-HSA-73863 | RNA Polymerase I Transcription Termination | UBTF | upstream binding transcription factor, RN | 7343 | down | 1,66 | 1,20E-02 |
| R-HSA-73863 | RNA Polymerase I Transcription Termination | ERCC3 | ERCC excision repair 3, TFIIH core com | 2071 | down | 1,20 | 2,37E-02 |
| R-HSA-73863 | RNA Polymerase I Transcription Termination | ERCC2 | ERCC excision repair 2, TFIIH core com | 2068 | down | 1,29 | 2,20E-02 |
| R-HSA-2426168 | Activation of gene expression by SREBF (SREBP) | DHCR7 | 7-dehydrocholesterol reductase | 1717 | down | 2,00 | 2,13E-03 |
| R-HSA-2426168 | Activation of gene expression by SREBF (SREBP) | SCD | stearoyl-CoA desaturase | 6319 | down | 1,88 | 3,59E-02 |
| R-HSA-2426168 | Activation of gene expression by SREBF (SREBP) | TGS1 | trimethylguanosine synthase 1 | 96764 | up | 1,63 | 9,37E-03 |

| | | | | | | | |
|---|---|---|---|---|---|---|---|
| R-HSA-2426168 | Activation of gene expression by SREBF (SREBP) | TBL1X | transducin beta like 1 X-linked | 6907 | down | 1,20 | 2,84E-02 |
| R-HSA-2426168 | Activation of gene expression by SREBF (SREBP) | CARM1 | coactivator associated arginine methyltr | 10498 | down | 1,36 | 1,60E-02 |
| R-HSA-2426168 | Activation of gene expression by SREBF (SREBP) | NFYA | nuclear transcription factor Y subunit alp | 4800 | up | 1,35 | 3,30E-02 |
| R-HSA-2426168 | Activation of gene expression by SREBF (SREBP) | FASN | fatty acid synthase | 2194 | down | 1,96 | 8,31E-03 |
| R-HSA-2426168 | Activation of gene expression by SREBF (SREBP) | SREBF1 | sterol regulatory element binding transc | 6720 | down | 1,42 | 3,67E-02 |
| R-HSA-2426168 | Activation of gene expression by SREBF (SREBP) | ACACA | acetyl-CoA carboxylase alpha | 31 | down | 1,40 | 2,54E-02 |
| R-HSA-2426168 | Activation of gene expression by SREBF (SREBP) | MVK | mevalonate kinase | 4598 | down | 1,99 | 3,36E-03 |
| R-HSA-2426168 | Activation of gene expression by SREBF (SREBP) | MTF1 | metal regulatory transcription factor 1 | 4520 | up | 1,42 | 3,66E-02 |
| R-HSA-2426168 | Activation of gene expression by SREBF (SREBP) | TM7SF2 | transmembrane 7 superfamily member 2 | 7108 | down | 1,51 | 4,00E-02 |
| R-HSA-2426168 | Activation of gene expression by SREBF (SREBP) | LSS | lanosterol synthase | 4047 | down | 1,49 | 1,65E-03 |
| R-HSA-2426168 | Activation of gene expression by SREBF (SREBP) | SQLE | squalene epoxidase | 6713 | down | 2,05 | 1,00E-02 |
| R-HSA-2426168 | Activation of gene expression by SREBF (SREBP) | RXRA | retinoid X receptor alpha | 6256 | down | 1,34 | 3,67E-03 |
| R-HSA-2426168 | Activation of gene expression by SREBF (SREBP) | CHD9 | chromodomain helicase DNA binding pr | 80205 | up | 1,32 | 1,42E-03 |
| R-HSA-5694530 | Cargo concentration in the ER | SEC24D | SEC24 homolog D, COPII coat complex | 9871 | up | 4,45 | 6,80E-04 |
| R-HSA-5694530 | Cargo concentration in the ER | MIA2 | melanoma inhibitory activity 2 | 4253 | up | 2,33 | 4,64E-03 |
| R-HSA-5694530 | Cargo concentration in the ER | AREG | amphiregulin | 374 | up | 3,33 | 2,37E-02 |
| R-HSA-5694530 | Cargo concentration in the ER | GOSR2 | golgi SNAP receptor complex member 2 | 9570 | up | 1,55 | 2,21E-02 |
| R-HSA-5694530 | Cargo concentration in the ER | SEC23A | Sec23 homolog A, coat complex II comp | 10484 | up | 1,46 | 2,47E-02 |
| R-HSA-5694530 | Cargo concentration in the ER | PREB | prolactin regulatory element binding | 10113 | up | 1,76 | 3,49E-02 |
| R-HSA-5694530 | Cargo concentration in the ER | MIA3 | MIA family member 3, ER export factor | 375056 | up | 1,93 | 2,25E-03 |
| R-HSA-5694530 | Cargo concentration in the ER | STX5 | syntaxin 5 | 6811 | up | 1,66 | 2,40E-02 |
| R-HSA-5694530 | Cargo concentration in the ER | LMAN1 | lectin, mannose binding 1 | 3998 | up | 1,54 | 4,14E-02 |
| R-HSA-5694530 | Cargo concentration in the ER | SEC24A | SEC24 homolog A, COPII coat complex | 10802 | up | 1,71 | 3,95E-03 |
| R-HSA-5694530 | Cargo concentration in the ER | CNIH2 | cornichon family AMPA receptor auxiliar | 254263 | down | 2,64 | 1,05E-02 |
| R-HSA-76066 | RNA Polymerase III Transcription Initiation From Type | POLR3F | RNA polymerase III subunit F | 10621 | up | 1,25 | 3,24E-02 |
| R-HSA-76066 | RNA Polymerase III Transcription Initiation From Type | POLR2L | RNA polymerase II subunit L | 5441 | down | 1,52 | 3,58E-03 |
| R-HSA-76066 | RNA Polymerase III Transcription Initiation From Type | GTF3C5 | general transcription factor IIIC subunit | 9328 | down | 1,24 | 3,90E-02 |
| R-HSA-76066 | RNA Polymerase III Transcription Initiation From Type | POLR2F | RNA polymerase II subunit F | 5435 | down | 1,41 | 3,22E-02 |
| R-HSA-76066 | RNA Polymerase III Transcription Initiation From Type | POLR1C | RNA polymerase I subunit C | 9533 | down | 1,24 | 4,71E-02 |
| R-HSA-76066 | RNA Polymerase III Transcription Initiation From Type | GTF3C2 | general transcription factor IIIC subunit | 2976 | down | 1,30 | 1,09E-02 |
| R-HSA-76066 | RNA Polymerase III Transcription Initiation From Type | POLR3C | RNA polymerase III subunit C | 10623 | up | 1,43 | 4,98E-02 |
| R-HSA-76066 | RNA Polymerase III Transcription Initiation From Type | POLR3D | RNA polymerase III subunit D | 661 | up | 1,52 | 2,72E-02 |
| R-HSA-76066 | RNA Polymerase III Transcription Initiation From Type | POLR3H | RNA polymerase III subunit H | 171568 | down | 1,80 | 7,07E-04 |
| R-HSA-76066 | RNA Polymerase III Transcription Initiation From Type | POLR2E | RNA polymerase II subunit E | 5434 | down | 1,31 | 3,18E-03 |
| R-HSA-76066 | RNA Polymerase III Transcription Initiation From Type | GTF3C3 | general transcription factor IIIC subunit | 9330 | up | 1,28 | 2,81E-03 |
| R-HSA-76066 | RNA Polymerase III Transcription Initiation From Type | BRF1 | BRF1, RNA polymerase III transcription | 2972 | down | 1,42 | 3,92E-04 |
| R-HSA-749476 | RNA Polymerase III Abortive And Retractive Initiation | GTF3C3 | general transcription factor IIIC subunit | 9330 | up | 1,28 | 2,81E-03 |
| R-HSA-749476 | RNA Polymerase III Abortive And Retractive Initiation | BRF1 | BRF1, RNA polymerase III transcription | 2972 | down | 1,42 | 3,92E-04 |
| R-HSA-749476 | RNA Polymerase III Abortive And Retractive Initiation | SNAPC3 | small nuclear RNA activating complex po | 6619 | up | 1,57 | 2,07E-03 |
| R-HSA-749476 | RNA Polymerase III Abortive And Retractive Initiation | NFIC | nuclear factor I C | 4782 | down | 1,53 | 1,48E-02 |
| R-HSA-749476 | RNA Polymerase III Abortive And Retractive Initiation | POLR2F | RNA polymerase II subunit F | 5435 | down | 1,41 | 3,22E-02 |
| R-HSA-749476 | RNA Polymerase III Abortive And Retractive Initiation | POLR3F | RNA polymerase III subunit F | 10621 | up | 1,25 | 3,24E-02 |
| R-HSA-749476 | RNA Polymerase III Abortive And Retractive Initiation | POLR2L | RNA polymerase II subunit L | 5441 | down | 1,52 | 3,58E-03 |
| R-HSA-749476 | RNA Polymerase III Abortive And Retractive Initiation | GTF3C5 | general transcription factor IIIC subunit | 9328 | down | 1,24 | 3,90E-02 |
| R-HSA-749476 | RNA Polymerase III Abortive And Retractive Initiation | POLR3H | RNA polymerase III subunit H | 171568 | down | 1,80 | 7,07E-04 |
| R-HSA-749476 | RNA Polymerase III Abortive And Retractive Initiation | POLR2E | RNA polymerase II subunit E | 5434 | down | 1,31 | 3,18E-03 |
| R-HSA-749476 | RNA Polymerase III Abortive And Retractive Initiation | SNAPC4 | small nuclear RNA activating complex po | 6621 | down | 1,26 | 4,38E-02 |
| R-HSA-749476 | RNA Polymerase III Abortive And Retractive Initiation | POLR3D | RNA polymerase III subunit D | 661 | up | 1,52 | 2,72E-02 |
| R-HSA-749476 | RNA Polymerase III Abortive And Retractive Initiation | POLR1C | RNA polymerase I subunit C | 9533 | down | 1,24 | 4,71E-02 |
| R-HSA-749476 | RNA Polymerase III Abortive And Retractive Initiation | POLR3C | RNA polymerase III subunit C | 10623 | up | 1,43 | 4,98E-02 |
| R-HSA-749476 | RNA Polymerase III Abortive And Retractive Initiation | GTF3C2 | general transcription factor IIIC subunit | 2976 | down | 1,30 | 1,09E-02 |
| R-HSA-381183 | ATF6 (ATF6-alpha) activates chaperone genes | ATF6 | activating transcription factor 6 | 22926 | up | 1,68 | 1,14E-02 |
| R-HSA-381183 | ATF6 (ATF6-alpha) activates chaperone genes | HSPA5 | heat shock protein family A (Hsp70) mer | 3309 | up | 5,02 | 6,98E-04 |
| R-HSA-381183 | ATF6 (ATF6-alpha) activates chaperone genes | DDIT3 | DNA damage inducible transcript 3 | 1649 | up | 5,65 | 3,67E-03 |
| R-HSA-381183 | ATF6 (ATF6-alpha) activates chaperone genes | HSP90B1 | heat shock protein 90 beta family memb | 7184 | up | 1,71 | 2,12E-02 |
| R-HSA-381183 | ATF6 (ATF6-alpha) activates chaperone genes | NFYA | nuclear transcription factor Y subunit alp | 4800 | up | 1,35 | 3,30E-02 |
| R-HSA-381183 | ATF6 (ATF6-alpha) activates chaperone genes | XBP1 | X-box binding protein 1 | 7494 | up | 2,00 | 5,67E-03 |
| R-HSA-76061 | RNA Polymerase III Transcription Initiation From Type | GTF3C2 | general transcription factor IIIC subunit | 2976 | down | 1,30 | 1,09E-02 |
| R-HSA-76061 | RNA Polymerase III Transcription Initiation From Type | POLR1C | RNA polymerase I subunit C | 9533 | down | 1,24 | 4,71E-02 |
| R-HSA-76061 | RNA Polymerase III Transcription Initiation From Type | POLR3C | RNA polymerase III subunit C | 10623 | up | 1,43 | 4,98E-02 |
| R-HSA-76061 | RNA Polymerase III Transcription Initiation From Type | POLR2E | RNA polymerase II subunit E | 5434 | down | 1,31 | 3,18E-03 |
| R-HSA-76061 | RNA Polymerase III Transcription Initiation From Type | POLR3H | RNA polymerase III subunit H | 171568 | down | 1,80 | 7,07E-04 |
| R-HSA-76061 | RNA Polymerase III Transcription Initiation From Type | POLR3D | RNA polymerase III subunit D | 661 | up | 1,52 | 2,72E-02 |
| R-HSA-76061 | RNA Polymerase III Transcription Initiation From Type | POLR2F | RNA polymerase II subunit F | 5435 | down | 1,41 | 3,22E-02 |
| R-HSA-76061 | RNA Polymerase III Transcription Initiation From Type | POLR2L | RNA polymerase II subunit L | 5441 | down | 1,52 | 3,58E-03 |
| R-HSA-76061 | RNA Polymerase III Transcription Initiation From Type | POLR3F | RNA polymerase III subunit F | 10621 | up | 1,25 | 3,24E-02 |
| R-HSA-76061 | RNA Polymerase III Transcription Initiation From Type | GTF3C5 | general transcription factor IIIC subunit | 9328 | down | 1,24 | 3,90E-02 |
| R-HSA-76061 | RNA Polymerase III Transcription Initiation From Type | GTF3C3 | general transcription factor IIIC subunit | 9330 | up | 1,28 | 2,81E-03 |
| R-HSA-76061 | RNA Polymerase III Transcription Initiation From Type | BRF1 | BRF1, RNA polymerase III transcription | 2972 | down | 1,42 | 3,92E-04 |
| R-HSA-379716 | Cytosolic tRNA aminoacylation | SARS | seryl-tRNA synthetase | 6301 | up | 1,82 | 1,31E-02 |
| R-HSA-379716 | Cytosolic tRNA aminoacylation | CARS | cysteinyl-tRNA synthetase | 833 | up | 1,55 | 4,38E-02 |
| R-HSA-379716 | Cytosolic tRNA aminoacylation | TARS | threonyl-tRNA synthetase | 6897 | up | 1,45 | 1,96E-02 |
| R-HSA-379716 | Cytosolic tRNA aminoacylation | NARS | asparaginyl-tRNA synthetase | 4677 | up | 1,29 | 2,58E-02 |
| R-HSA-379716 | Cytosolic tRNA aminoacylation | WARS | tryptophanyl-tRNA synthetase | 7453 | up | 1,88 | 3,69E-02 |
| R-HSA-379716 | Cytosolic tRNA aminoacylation | AARS | alanyl-tRNA synthetase | 16 | up | 1,82 | 1,84E-02 |
| R-HSA-379716 | Cytosolic tRNA aminoacylation | GARS | glycyl-tRNA synthetase | 2617 | up | 1,65 | 1,79E-02 |
| R-HSA-379716 | Cytosolic tRNA aminoacylation | EPRS | glutamyl-prolyl-tRNA synthetase | 2058 | up | 1,42 | 1,95E-02 |
| R-HSA-176974 | Unwinding of DNA | MCM2 | minichromosome maintenance complex | 4171 | down | 2,01 | 1,42E-02 |
| R-HSA-176974 | Unwinding of DNA | GINS3 | GINS complex subunit 3 | 64785 | down | 1,95 | 1,67E-02 |
| R-HSA-176974 | Unwinding of DNA | GINS1 | GINS complex subunit 1 | 9837 | down | 2,17 | 3,09E-02 |
| R-HSA-176974 | Unwinding of DNA | MCM3 | minichromosome maintenance complex | 4172 | down | 2,07 | 4,13E-02 |
| R-HSA-176974 | Unwinding of DNA | MCM5 | minichromosome maintenance complex | 4174 | down | 1,83 | 2,22E-02 |
| R-HSA-176974 | Unwinding of DNA | MCM7 | minichromosome maintenance complex | 4176 | down | 1,67 | 3,35E-02 |
| R-HSA-6782135 | Dual incision in TC-NER | POLD1 | DNA polymerase delta 1, catalytic subur | 5424 | down | 1,59 | 4,62E-02 |
| R-HSA-6782135 | Dual incision in TC-NER | CDK7 | cyclin dependent kinase 7 | 1022 | up | 1,37 | 4,38E-02 |
| R-HSA-6782135 | Dual incision in TC-NER | POLD2 | DNA polymerase delta 2, accessory sub | 5425 | down | 1,30 | 1,92E-02 |
| R-HSA-6782135 | Dual incision in TC-NER | POLE | DNA polymerase epsilon, catalytic subu | 5426 | down | 1,55 | 2,67E-02 |
| R-HSA-6782135 | Dual incision in TC-NER | GTF2H1 | general transcription factor IIH subunit 1 | 2965 | up | 1,33 | 1,85E-02 |
| R-HSA-6782135 | Dual incision in TC-NER | POLR2L | RNA polymerase II subunit L | 5441 | down | 1,52 | 3,58E-03 |
| R-HSA-6782135 | Dual incision in TC-NER | POLR2B | RNA polymerase II subunit B | 5431 | down | 1,23 | 3,81E-02 |
| R-HSA-6782135 | Dual incision in TC-NER | PRPF19 | pre-mRNA processing factor 19 | 27339 | down | 1,24 | 4,41E-02 |
| R-HSA-6782135 | Dual incision in TC-NER | RFC1 | replication factor C subunit 1 | 5981 | up | 1,15 | 2,18E-02 |
| R-HSA-6782135 | Dual incision in TC-NER | ERCC3 | ERCC excision repair 3, TFIIH core com | 2071 | down | 1,20 | 2,37E-02 |
| R-HSA-6782135 | Dual incision in TC-NER | ERCC2 | ERCC excision repair 2, TFIIH core com | 2068 | down | 1,29 | 2,20E-02 |
| R-HSA-6782135 | Dual incision in TC-NER | POLR2F | RNA polymerase II subunit F | 5435 | down | 1,41 | 3,22E-02 |
| R-HSA-6782135 | Dual incision in TC-NER | TCEA1 | transcription elongation factor A1 | 6917 | up | 1,76 | 1,81E-02 |
| R-HSA-6782135 | Dual incision in TC-NER | PCNA | proliferating cell nuclear antigen | 5111 | down | 2,14 | 4,63E-02 |
| R-HSA-6782135 | Dual incision in TC-NER | POLR2E | RNA polymerase II subunit E | 5434 | down | 1,31 | 3,18E-03 |
| R-HSA-6782135 | Dual incision in TC-NER | RPA1 | replication protein A1 | 6117 | down | 1,41 | 4,93E-02 |
| R-HSA-6782210 | Gap-filling DNA repair synthesis and ligation in TC-NE | ERCC2 | ERCC excision repair 2, TFIIH core com | 2068 | down | 1,29 | 2,20E-02 |
| R-HSA-6782210 | Gap-filling DNA repair synthesis and ligation in TC-NE | ERCC3 | ERCC excision repair 3, TFIIH core com | 2071 | down | 1,20 | 2,37E-02 |
| R-HSA-6782210 | Gap-filling DNA repair synthesis and ligation in TC-NE | RPA1 | replication protein A1 | 6117 | down | 1,41 | 4,93E-02 |
| R-HSA-6782210 | Gap-filling DNA repair synthesis and ligation in TC-NE | POLR2E | RNA polymerase II subunit E | 5434 | down | 1,31 | 3,18E-03 |
| R-HSA-6782210 | Gap-filling DNA repair synthesis and ligation in TC-NE | PCNA | proliferating cell nuclear antigen | 5111 | down | 2,14 | 4,63E-02 |
| R-HSA-6782210 | Gap-filling DNA repair synthesis and ligation in TC-NE | POLR2F | RNA polymerase II subunit F | 5435 | down | 1,41 | 3,22E-02 |
| R-HSA-6782210 | Gap-filling DNA repair synthesis and ligation in TC-NE | TCEA1 | transcription elongation factor A1 | 6917 | up | 1,76 | 1,81E-02 |
| R-HSA-6782210 | Gap-filling DNA repair synthesis and ligation in TC-NE | POLD2 | DNA polymerase delta 2, accessory sub | 5425 | down | 1,30 | 1,92E-02 |
| R-HSA-6782210 | Gap-filling DNA repair synthesis and ligation in TC-NE | CDK7 | cyclin dependent kinase 7 | 1022 | up | 1,37 | 4,38E-02 |
| R-HSA-6782210 | Gap-filling DNA repair synthesis and ligation in TC-NE | POLD1 | DNA polymerase delta 1, catalytic subur | 5424 | down | 1,59 | 4,62E-02 |
| R-HSA-6782210 | Gap-filling DNA repair synthesis and ligation in TC-NE | RFC1 | replication factor C subunit 1 | 5981 | up | 1,15 | 2,18E-02 |
| R-HSA-6782210 | Gap-filling DNA repair synthesis and ligation in TC-NE | POLR2B | RNA polymerase II subunit B | 5431 | down | 1,23 | 3,81E-02 |
| R-HSA-6782210 | Gap-filling DNA repair synthesis and ligation in TC-NE | PRPF19 | pre-mRNA processing factor 19 | 27339 | down | 1,24 | 4,41E-02 |
| R-HSA-6782210 | Gap-filling DNA repair synthesis and ligation in TC-NE | POLR2L | RNA polymerase II subunit L | 5441 | down | 1,52 | 3,58E-03 |
| R-HSA-6782210 | Gap-filling DNA repair synthesis and ligation in TC-NE | GTF2H1 | general transcription factor IIH subunit 1 | 2965 | up | 1,33 | 1,85E-02 |
| R-HSA-6782210 | Gap-filling DNA repair synthesis and ligation in TC-NE | POLE | DNA polymerase epsilon, catalytic subu | 5426 | down | 1,55 | 2,67E-02 |
| R-HSA-69166 | Removal of the Flap Intermediate | PCNA | proliferating cell nuclear antigen | 5111 | down | 2,14 | 4,63E-02 |
| R-HSA-69166 | Removal of the Flap Intermediate | POLD2 | DNA polymerase delta 2, accessory sub | 5425 | down | 1,30 | 1,92E-02 |
| R-HSA-69166 | Removal of the Flap Intermediate | POLA2 | DNA polymerase alpha 2, accessory sub | 23649 | down | 1,72 | 2,46E-02 |
| R-HSA-69166 | Removal of the Flap Intermediate | FEN1 | flap structure-specific endonuclease 1 | 2237 | down | 2,04 | 1,60E-02 |
| R-HSA-69166 | Removal of the Flap Intermediate | RPA1 | replication protein A1 | 6117 | down | 1,41 | 4,93E-02 |
| R-HSA-69166 | Removal of the Flap Intermediate | POLD1 | DNA polymerase delta 1, catalytic subur | 5424 | down | 1,59 | 4,62E-02 |
| R-HSA-191273 | Cholesterol biosynthesis | MVK | mevalonate kinase | 4598 | down | 1,99 | 3,36E-03 |
| R-HSA-191273 | Cholesterol biosynthesis | NSDHL | NAD(P) dependent steroid dehydrogena | 50814 | down | 1,36 | 1,99E-02 |
| R-HSA-191273 | Cholesterol biosynthesis | ACAT2 | acetyl-CoA acetyltransferase 2 | 39 | down | 1,69 | 2,52E-02 |
| R-HSA-191273 | Cholesterol biosynthesis | LSS | lanosterol synthase | 4047 | down | 1,49 | 1,65E-03 |
| R-HSA-191273 | Cholesterol biosynthesis | TM7SF2 | transmembrane 7 superfamily member 2 | 7108 | down | 1,51 | 4,00E-02 |
| R-HSA-191273 | Cholesterol biosynthesis | LBR | lamin B receptor | 3930 | down | 1,48 | 4,92E-02 |
| R-HSA-191273 | Cholesterol biosynthesis | SQLE | squalene epoxidase | 6713 | down | 2,05 | 1,00E-02 |
| R-HSA-174437 | Removal of the Flap Intermediate from the C-strand | FEN1 | flap structure-specific endonuclease 1 | 2237 | down | 2,04 | 1,60E-02 |
| R-HSA-174437 | Removal of the Flap Intermediate from the C-strand | PCNA | proliferating cell nuclear antigen | 5111 | down | 2,14 | 4,63E-02 |
| R-HSA-174437 | Removal of the Flap Intermediate from the C-strand | POLD2 | DNA polymerase delta 2, accessory sub | 5425 | down | 1,30 | 1,92E-02 |
| R-HSA-174437 | Removal of the Flap Intermediate from the C-strand | RPA1 | replication protein A1 | 6117 | down | 1,41 | 4,93E-02 |
| R-HSA-174437 | Removal of the Flap Intermediate from the C-strand | POLD1 | DNA polymerase delta 1, catalytic subur | 5424 | down | 1,59 | 4,62E-02 |
| R-HSA-75109 | Triglyceride biosynthesis | AGPAT2 | 1-acylglycerol-3-phosphate O-acyltransf | 10555 | down | 1,23 | 2,89E-03 |
| R-HSA-75109 | Triglyceride biosynthesis | GPAT4 | glycerol-3-phosphate acyltransferase 4 | 137964 | down | 1,24 | 4,75E-02 |
| R-HSA-75109 | Triglyceride biosynthesis | LPCAT1 | lysophosphatidylcholine acyltransferase | 79888 | down | 1,76 | 2,60E-02 |
| R-HSA-75109 | Triglyceride biosynthesis | GPAT3 | glycerol-3-phosphate acyltransferase 3 | 84803 | up | 1,77 | 4,99E-02 |
| R-HSA-75109 | Triglyceride biosynthesis | LPIN2 | lipin 2 | 9663 | up | 2,16 | 6,88E-03 |
| R-HSA-75109 | Triglyceride biosynthesis | AGPAT3 | 1-acylglycerol-3-phosphate O-acyltransf | 56894 | down | 1,30 | 9,80E-03 |
| R-HSA-75109 | Triglyceride biosynthesis | LPCAT4 | lysophosphatidylcholine acyltransferase | 254531 | down | 1,78 | 2,52E-02 |
| R-HSA-75109 | Triglyceride biosynthesis | LPIN3 | lipin 3 | 64900 | down | 1,36 | 4,76E-02 |
| R-HSA-75109 | Triglyceride biosynthesis | AGPAT5 | 1-acylglycerol-3-phosphate O-acyltransf | 55326 | down | 1,30 | 4,70E-02 |
| R-HSA-1368108 | BMAL1:CLOCK,NPAS2 activates circadian gene exp | CREB1 | cAMP responsive element binding prote | 1385 | up | 1,51 | 7,18E-03 |
| R-HSA-1368108 | BMAL1:CLOCK,NPAS2 activates circadian gene exp | CHD9 | chromodomain helicase DNA binding pr | 80205 | up | 1,32 | 1,42E-03 |
| R-HSA-1368108 | BMAL1:CLOCK,NPAS2 activates circadian gene exp | HIF1A | hypoxia inducible factor 1 alpha subunit | 3091 | up | 1,23 | 4,66E-02 |
| R-HSA-1368108 | BMAL1:CLOCK,NPAS2 activates circadian gene exp | NAMPT | nicotinamide phosphoribosyltransferase | 10135 | up | 1,49 | 4,90E-02 |
| R-HSA-1368108 | BMAL1:CLOCK,NPAS2 activates circadian gene exp | TBL1X | transducin beta like 1 X-linked | 6907 | down | 1,20 | 2,84E-02 |
| R-HSA-1368108 | BMAL1:CLOCK,NPAS2 activates circadian gene exp | RXRA | retinoid X receptor alpha | 6256 | down | 1,34 | 3,67E-03 |
| R-HSA-1368108 | BMAL1:CLOCK,NPAS2 activates circadian gene exp | CARM1 | coactivator associated arginine methyltr | 10498 | down | 1,36 | 1,60E-02 |
| R-HSA-1368108 | BMAL1:CLOCK,NPAS2 activates circadian gene exp | NOCT | nocturnin | 25819 | up | 1,79 | 6,10E-03 |
| R-HSA-1368108 | BMAL1:CLOCK,NPAS2 activates circadian gene exp | TGS1 | trimethylguanosine synthase 1 | 96764 | up | 1,63 | 9,37E-03 |
| R-HSA-1368108 | BMAL1:CLOCK,NPAS2 activates circadian gene exp | BHLHE40 | basic helix-loop-helix family member e40 | 8553 | up | 1,80 | 3,56E-02 |
| R-HSA-1368108 | BMAL1:CLOCK,NPAS2 activates circadian gene exp | ARNTL | aryl hydrocarbon receptor nuclear transl | 406 | up | 1,67 | 5,94E-03 |
| R-HSA-1368108 | BMAL1:CLOCK,NPAS2 activates circadian gene exp | PER1 | period circadian regulator 1 | 5187 | up | 2,35 | 1,38E-02 |
| R-HSA-1368108 | BMAL1:CLOCK,NPAS2 activates circadian gene exp | DBP | D-box binding PAR bZIP transcription fa | 1628 | down | 1,59 | 2,85E-02 |
| R-HSA-442717 | No details found for 442717 | CALM1 | calmodulin 1 | 801 | down | 1,43 | 2,23E-02 |
| R-HSA-442717 | No details found for 442717 | CALM3 | calmodulin 3 | 808 | down | 1,43 | 2,23E-02 |
| R-HSA-442717 | No details found for 442717 | CREB1 | cAMP responsive element binding prote | 1385 | up | 1,51 | 7,18E-03 |
| R-HSA-442717 | No details found for 442717 | CALM2 | calmodulin 2 | 805 | down | 1,43 | 2,23E-02 |
| R-HSA-442717 | No details found for 442717 | CAMKK1 | calcium/calmodulin dependent protein k | 84254 | down | 1,83 | 2,57E-02 |
| R-HSA-1474151 | Tetrahydrobiopterin (BH4) synthesis, recycling, salvag | AKT1 | AKT serine/threonine kinase 1 | 207 | down | 1,37 | 1,42E-03 |
| R-HSA-1474151 | Tetrahydrobiopterin (BH4) synthesis, recycling, salvag | CALM1 | calmodulin 1 | 801 | down | 1,43 | 2,23E-02 |
| R-HSA-1474151 | Tetrahydrobiopterin (BH4) synthesis, recycling, salvag | CALM3 | calmodulin 3 | 808 | down | 1,43 | 2,23E-02 |
| R-HSA-1474151 | Tetrahydrobiopterin (BH4) synthesis, recycling, salvag | CALM2 | calmodulin 2 | 805 | down | 1,43 | 2,23E-02 |
| R-HSA-1474151 | Tetrahydrobiopterin (BH4) synthesis, recycling, salvag | SPR | sepiapterin reductase | 6697 | down | 1,54 | 1,91E-02 |
| R-HSA-69052 | Switching of origins to a post-replicative state | MCM7 | minichromosome maintenance complex | 4176 | down | 1,67 | 3,35E-02 |
| R-HSA-69052 | Switching of origins to a post-replicative state | MCM3 | minichromosome maintenance complex | 4172 | down | 2,07 | 4,13E-02 |
| R-HSA-69052 | Switching of origins to a post-replicative state | MCM5 | minichromosome maintenance complex | 4174 | down | 1,83 | 2,22E-02 |
| R-HSA-69052 | Switching of origins to a post-replicative state | MCM2 | minichromosome maintenance complex | 4171 | down | 2,01 | 1,42E-02 |
| R-HSA-73980 | RNA Polymerase III Transcription Termination | POLR1C | RNA polymerase I subunit C | 9533 | down | 1,24 | 4,71E-02 |
| R-HSA-73980 | RNA Polymerase III Transcription Termination | POLR3C | RNA polymerase III subunit C | 10623 | up | 1,43 | 4,98E-02 |
| R-HSA-73980 | RNA Polymerase III Transcription Termination | POLR2E | RNA polymerase II subunit E | 5434 | down | 1,31 | 3,18E-03 |
| R-HSA-73980 | RNA Polymerase III Transcription Termination | POLR3H | RNA polymerase III subunit H | 171568 | down | 1,80 | 7,07E-04 |
| R-HSA-73980 | RNA Polymerase III Transcription Termination | POLR3D | RNA polymerase III subunit D | 661 | up | 1,52 | 2,72E-02 |
| R-HSA-73980 | RNA Polymerase III Transcription Termination | POLR2F | RNA polymerase II subunit F | 5435 | down | 1,41 | 3,22E-02 |
| R-HSA-73980 | RNA Polymerase III Transcription Termination | POLR2L | RNA polymerase II subunit L | 5441 | down | 1,52 | 3,58E-03 |
| R-HSA-73980 | RNA Polymerase III Transcription Termination | POLR3F | RNA polymerase III subunit F | 10621 | up | 1,25 | 3,24E-02 |
| R-HSA-73980 | RNA Polymerase III Transcription Termination | NFIC | nuclear factor I C | 4782 | down | 1,53 | 1,48E-02 |
| R-HSA-73780 | RNA Polymerase III Chain Elongation | POLR2L | RNA polymerase II subunit L | 5441 | down | 1,52 | 3,58E-03 |
| R-HSA-73780 | RNA Polymerase III Chain Elongation | POLR3F | RNA polymerase III subunit F | 10621 | up | 1,25 | 3,24E-02 |
| R-HSA-73780 | RNA Polymerase III Chain Elongation | POLR2F | RNA polymerase II subunit F | 5435 | down | 1,41 | 3,22E-02 |
| R-HSA-73780 | RNA Polymerase III Chain Elongation | POLR3D | RNA polymerase III subunit D | 661 | up | 1,52 | 2,72E-02 |
| R-HSA-73780 | RNA Polymerase III Chain Elongation | POLR2E | RNA polymerase II subunit E | 5434 | down | 1,31 | 3,18E-03 |
| R-HSA-73780 | RNA Polymerase III Chain Elongation | POLR3H | RNA polymerase III subunit H | 171568 | down | 1,80 | 7,07E-04 |
| R-HSA-73780 | RNA Polymerase III Chain Elongation | POLR3C | RNA polymerase III subunit C | 10623 | up | 1,43 | 4,98E-02 |
| R-HSA-73780 | RNA Polymerase III Chain Elongation | POLR1C | RNA polymerase I subunit C | 9533 | down | 1,24 | 4,71E-02 |
| R-HSA-2151201 | Transcriptional activation of mitochondrial biogenesis | TWNK | twinkle mtDNA helicase | 56652 | down | 1,34 | 4,18E-02 |
| R-HSA-2151201 | Transcriptional activation of mitochondrial biogenesis | SIRT3 | sirtuin 3 | 23410 | down | 1,27 | 4,49E-02 |
| R-HSA-2151201 | Transcriptional activation of mitochondrial biogenesis | CHD9 | chromodomain helicase DNA binding pr | 80205 | up | 1,32 | 1,42E-03 |
| R-HSA-2151201 | Transcriptional activation of mitochondrial biogenesis | CREB1 | cAMP responsive element binding prote | 1385 | up | 1,51 | 7,18E-03 |
| R-HSA-2151201 | Transcriptional activation of mitochondrial biogenesis | MEF2D | myocyte enhancer factor 2D | 4209 | up | 1,90 | 4,04E-02 |
| R-HSA-2151201 | Transcriptional activation of mitochondrial biogenesis | ATF2 | activating transcription factor 2 | 1386 | up | 1,79 | 1,01E-02 |
| R-HSA-2151201 | Transcriptional activation of mitochondrial biogenesis | TGS1 | trimethylguanosine synthase 1 | 96764 | up | 1,63 | 9,37E-03 |
| R-HSA-2151201 | Transcriptional activation of mitochondrial biogenesis | HCFC1 | host cell factor C1 | 3054 | down | 1,33 | 4,82E-04 |
| R-HSA-2151201 | Transcriptional activation of mitochondrial biogenesis | GABPB1 | GA binding protein transcription factor b | 2553 | up | 1,31 | 1,57E-02 |
| R-HSA-2151201 | Transcriptional activation of mitochondrial biogenesis | TBL1X | transducin beta like 1 X-linked | 6907 | down | 1,20 | 2,84E-02 |
| R-HSA-2151201 | Transcriptional activation of mitochondrial biogenesis | RXRA | retinoid X receptor alpha | 6256 | down | 1,34 | 3,67E-03 |
| R-HSA-2151201 | Transcriptional activation of mitochondrial biogenesis | CARM1 | coactivator associated arginine methyltr | 10498 | down | 1,36 | 1,60E-02 |
| R-HSA-76071 | RNA Polymerase III Transcription Initiation From Type | SNAPC3 | small nuclear RNA activating complex po | 6619 | up | 1,57 | 2,07E-03 |
| R-HSA-76071 | RNA Polymerase III Transcription Initiation From Type | POLR2F | RNA polymerase II subunit F | 5435 | down | 1,41 | 3,22E-02 |
| R-HSA-76071 | RNA Polymerase III Transcription Initiation From Type | POLR2L | RNA polymerase II subunit L | 5441 | down | 1,52 | 3,58E-03 |
| R-HSA-76071 | RNA Polymerase III Transcription Initiation From Type | POLR3F | RNA polymerase III subunit F | 10621 | up | 1,25 | 3,24E-02 |
| R-HSA-76071 | RNA Polymerase III Transcription Initiation From Type | POLR3H | RNA polymerase III subunit H | 171568 | down | 1,80 | 7,07E-04 |
| R-HSA-76071 | RNA Polymerase III Transcription Initiation From Type | POLR2E | RNA polymerase II subunit E | 5434 | down | 1,31 | 3,18E-03 |
| R-HSA-76071 | RNA Polymerase III Transcription Initiation From Type | POLR3D | RNA polymerase III subunit D | 661 | up | 1,52 | 2,72E-02 |
| R-HSA-76071 | RNA Polymerase III Transcription Initiation From Type | SNAPC4 | small nuclear RNA activating complex po | 6621 | down | 1,26 | 4,38E-02 |
| R-HSA-76071 | RNA Polymerase III Transcription Initiation From Type | POLR3C | RNA polymerase III subunit C | 10623 | up | 1,43 | 4,98E-02 |
| R-HSA-76071 | RNA Polymerase III Transcription Initiation From Type | POLR1C | RNA polymerase I subunit C | 9533 | down | 1,24 | 4,71E-02 |
| R-HSA-5218921 | VEGFR2 mediated cell proliferation | SPHK1 | sphingosine kinase 1 | 8877 | down | 2,00 | 2,34E-02 |
| R-HSA-5218921 | VEGFR2 mediated cell proliferation | PLCG1 | phospholipase C gamma 1 | 5335 | down | 1,51 | 2,85E-02 |
| R-HSA-5218921 | VEGFR2 mediated cell proliferation | CALM1 | calmodulin 1 | 801 | down | 1,43 | 2,23E-02 |
| R-HSA-5218921 | VEGFR2 mediated cell proliferation | HRAS | HRas proto-oncogene, GTPase | 3265 | down | 1,31 | 2,19E-03 |
| R-HSA-5218921 | VEGFR2 mediated cell proliferation | CALM3 | calmodulin 3 | 808 | down | 1,43 | 2,23E-02 |
| R-HSA-5218921 | VEGFR2 mediated cell proliferation | CALM2 | calmodulin 2 | 805 | down | 1,43 | 2,23E-02 |
| R-HSA-5218921 | VEGFR2 mediated cell proliferation | VEGFA | vascular endothelial growth factor A | 7422 | up | 2,38 | 1,68E-02 |
| R-HSA-5651801 | PCNA-Dependent Long Patch Base Excision Repair | POLE | DNA polymerase epsilon, catalytic subu | 5426 | down | 1,55 | 2,67E-02 |
| R-HSA-5651801 | PCNA-Dependent Long Patch Base Excision Repair | RFC1 | replication factor C subunit 1 | 5981 | up | 1,15 | 2,18E-02 |

| | | | | | | | |
|---|---|---|---|---|---|---|---|
| R-HSA-5651801 | PCNA-Dependent Long Patch Base Excision Repair | PCNA | proliferating cell nuclear antigen | 5111 | down | 2,14 | 4,63E-02 |
| R-HSA-5651801 | PCNA-Dependent Long Patch Base Excision Repair | POLB | DNA polymerase beta | 5423 | up | 1,25 | 3,75E-02 |
| R-HSA-5651801 | PCNA-Dependent Long Patch Base Excision Repair | RPA1 | replication protein A1 | 6117 | down | 1,41 | 4,93E-02 |
| R-HSA-5651801 | PCNA-Dependent Long Patch Base Excision Repair | POLD1 | DNA polymerase delta 1, catalytic subur | 5424 | down | 1,59 | 4,62E-02 |
| R-HSA-5651801 | PCNA-Dependent Long Patch Base Excision Repair | POLD2 | DNA polymerase delta 2, accessory sub | 5425 | down | 1,30 | 1,92E-02 |
| R-HSA-5651801 | PCNA-Dependent Long Patch Base Excision Repair | FEN1 | flap structure-specific endonuclease 1 | 2237 | down | 2,04 | 1,60E-02 |
| R-HSA-400253 | Circadian Clock | PPP1CA | protein phosphatase 1 catalytic subunit | 5499 | down | 1,24 | 9,77E-03 |
| R-HSA-400253 | Circadian Clock | MEF2D | myocyte enhancer factor 2D | 4209 | up | 1,90 | 4,04E-02 |
| R-HSA-400253 | Circadian Clock | CHD9 | chromodomain helicase DNA binding pr | 80205 | up | 1,32 | 1,42E-03 |
| R-HSA-400253 | Circadian Clock | HIF1A | hypoxia inducible factor 1 alpha subunit | 3091 | up | 1,23 | 4,66E-02 |
| R-HSA-400253 | Circadian Clock | CREB1 | cAMP responsive element binding prote | 1385 | up | 1,51 | 7,18E-03 |
| R-HSA-400253 | Circadian Clock | TGS1 | trimethylguanosine synthase 1 | 96764 | up | 1,63 | 9,37E-03 |
| R-HSA-400253 | Circadian Clock | ATF2 | activating transcription factor 2 | 1386 | up | 1,79 | 1,01E-02 |
| R-HSA-400253 | Circadian Clock | CARM1 | coactivator associated arginine methyltr | 10498 | down | 1,36 | 1,60E-02 |
| R-HSA-400253 | Circadian Clock | RXRA | retinoid X receptor alpha | 6256 | down | 1,34 | 3,67E-03 |
| R-HSA-400253 | Circadian Clock | TBL1X | transducin beta like 1 X-linked | 6907 | down | 1,20 | 2,84E-02 |
| R-HSA-400253 | Circadian Clock | PER1 | period circadian regulator 1 | 5187 | up | 2,35 | 1,38E-02 |
| R-HSA-400253 | Circadian Clock | ARNTL | aryl hydrocarbon receptor nuclear transl | 406 | up | 1,67 | 5,94E-03 |
| R-HSA-400253 | Circadian Clock | SIRT1 | sirtuin 1 | 23411 | up | 1,60 | 1,00E-02 |
| R-HSA-5675221 | Negative regulation of MAPK pathway | DUSP5 | dual specificity phosphatase 5 | 1847 | up | 3,63 | 9,77E-03 |
| R-HSA-5675221 | Negative regulation of MAPK pathway | DUSP8 | dual specificity phosphatase 8 | 1850 | up | 2,42 | 1,40E-02 |
| R-HSA-5675221 | Negative regulation of MAPK pathway | HRAS | HRas proto-oncogene, GTPase | 3265 | down | 1,31 | 2,19E-03 |
| R-HSA-5675221 | Negative regulation of MAPK pathway | DUSP10 | dual specificity phosphatase 10 | 11221 | up | 2,34 | 2,20E-03 |
| R-HSA-5675221 | Negative regulation of MAPK pathway | DUSP1 | dual specificity phosphatase 1 | 1843 | up | 1,99 | 2,51E-02 |
| R-HSA-5675221 | Negative regulation of MAPK pathway | PPP5C | protein phosphatase 5 catalytic subunit | 5536 | down | 1,29 | 4,15E-02 |
| R-HSA-5675221 | Negative regulation of MAPK pathway | BRAF | B-Raf proto-oncogene, serine/threonine | 673 | up | 1,52 | 4,53E-02 |
| R-HSA-5675221 | Negative regulation of MAPK pathway | BRAP | BRCA1 associated protein | 8315 | up | 1,34 | 1,14E-02 |
| R-HSA-5675221 | Negative regulation of MAPK pathway | PPP2CB | protein phosphatase 2 catalytic subunit | 5516 | up | 1,30 | 4,79E-02 |
| R-HSA-5675221 | Negative regulation of MAPK pathway | DUSP7 | dual specificity phosphatase 7 | 1849 | down | 1,42 | 1,78E-02 |
| R-HSA-5675221 | Negative regulation of MAPK pathway | MARK3 | microtubule affinity regulating kinase 3 | 4140 | up | 1,40 | 1,07E-02 |
| R-HSA-203927 | MicroRNA (miRNA) biogenesis | AGO3 | argonaute 3, RISC catalytic component | 192669 | up | 1,36 | 1,80E-02 |
| R-HSA-203927 | MicroRNA (miRNA) biogenesis | POLR2E | RNA polymerase II subunit E | 5434 | down | 1,31 | 3,18E-03 |
| R-HSA-203927 | MicroRNA (miRNA) biogenesis | TARBP2 | TARBP2, RISC loading complex RNA bi | 6895 | down | 1,36 | 3,80E-03 |
| R-HSA-203927 | MicroRNA (miRNA) biogenesis | POLR2B | RNA polymerase II subunit B | 5431 | down | 1,23 | 3,81E-02 |
| R-HSA-203927 | MicroRNA (miRNA) biogenesis | DGCR8 | DGCR8, microprocessor complex subun | 54487 | down | 1,84 | 1,01E-02 |
| R-HSA-203927 | MicroRNA (miRNA) biogenesis | POLR2L | RNA polymerase II subunit L | 5441 | down | 1,52 | 3,58E-03 |
| R-HSA-203927 | MicroRNA (miRNA) biogenesis | POLR2F | RNA polymerase II subunit F | 5435 | down | 1,41 | 3,22E-02 |
| R-HSA-5696400 | Dual Incision in GG-NER | ERCC3 | ERCC excision repair 3, TFIIH core com | 2071 | down | 1,20 | 2,37E-02 |
| R-HSA-5696400 | Dual Incision in GG-NER | ERCC2 | ERCC excision repair 2, TFIIH core com | 2068 | down | 1,29 | 2,20E-02 |
| R-HSA-5696400 | Dual Incision in GG-NER | POLD2 | DNA polymerase delta 2, accessory sub | 5425 | down | 1,30 | 1,92E-02 |
| R-HSA-5696400 | Dual Incision in GG-NER | POLD1 | DNA polymerase delta 1, catalytic subur | 5424 | down | 1,59 | 4,62E-02 |
| R-HSA-5696400 | Dual Incision in GG-NER | RFC1 | replication factor C subunit 1 | 5981 | up | 1,15 | 2,18E-02 |
| R-HSA-5696400 | Dual Incision in GG-NER | PCNA | proliferating cell nuclear antigen | 5111 | down | 2,14 | 4,63E-02 |
| R-HSA-5696400 | Dual Incision in GG-NER | RPA1 | replication protein A1 | 6117 | down | 1,41 | 4,93E-02 |
| R-HSA-5696400 | Dual Incision in GG-NER | POLE | DNA polymerase epsilon, catalytic subu | 5426 | down | 1,55 | 2,67E-02 |
| R-HSA-5696400 | Dual Incision in GG-NER | GTF2H1 | general transcription factor IIH subunit 1 | 2965 | up | 1,33 | 1,85E-02 |
| R-HSA-5696400 | Dual Incision in GG-NER | PARP1 | poly(ADP-ribose) polymerase 1 | 142 | down | 1,50 | 3,92E-02 |
| R-HSA-68962 | Activation of the pre-replicative complex | MCM7 | minichromosome maintenance complex | 4176 | down | 1,67 | 3,35E-02 |
| R-HSA-68962 | Activation of the pre-replicative complex | MCM5 | minichromosome maintenance complex | 4174 | down | 1,83 | 2,22E-02 |
| R-HSA-68962 | Activation of the pre-replicative complex | POLA2 | DNA polymerase alpha 2, accessory sub | 23649 | down | 1,72 | 2,46E-02 |
| R-HSA-68962 | Activation of the pre-replicative complex | MCM3 | minichromosome maintenance complex | 4172 | down | 2,07 | 4,13E-02 |
| R-HSA-68962 | Activation of the pre-replicative complex | RPA1 | replication protein A1 | 6117 | down | 1,41 | 4,93E-02 |
| R-HSA-68962 | Activation of the pre-replicative complex | POLE | DNA polymerase epsilon, catalytic subu | 5426 | down | 1,55 | 2,67E-02 |
| R-HSA-68962 | Activation of the pre-replicative complex | MCM2 | minichromosome maintenance complex | 4171 | down | 2,01 | 1,42E-02 |
| R-HSA-72165 | mRNA Splicing - Minor Pathway | POLR2B | RNA polymerase II subunit B | 5431 | down | 1,23 | 3,81E-02 |
| R-HSA-72165 | mRNA Splicing - Minor Pathway | ZRSR2 | zinc finger CCCH-type, RNA binding mo | 8233 | up | 1,53 | 1,67E-02 |
| R-HSA-72165 | mRNA Splicing - Minor Pathway | POLR2F | RNA polymerase II subunit F | 5435 | down | 1,41 | 3,22E-02 |
| R-HSA-72165 | mRNA Splicing - Minor Pathway | POLR2L | RNA polymerase II subunit L | 5441 | down | 1,52 | 3,58E-03 |
| R-HSA-72165 | mRNA Splicing - Minor Pathway | EFTUD2 | elongation factor Tu GTP binding doma | 9343 | down | 1,24 | 3,99E-02 |
| R-HSA-72165 | mRNA Splicing - Minor Pathway | SNRNP35 | small nuclear ribonucleoprotein U11/U1 | 11066 | down | 1,38 | 4,95E-02 |
| R-HSA-72165 | mRNA Splicing - Minor Pathway | POLR2E | RNA polymerase II subunit E | 5434 | down | 1,31 | 3,18E-03 |
| R-HSA-72165 | mRNA Splicing - Minor Pathway | SRSF2 | serine and arginine rich splicing factor 2 | 6427 | down | 1,69 | 1,79E-02 |
| R-HSA-72165 | mRNA Splicing - Minor Pathway | DDX23 | DEAD-box helicase 23 | 9416 | down | 1,32 | 1,25E-02 |
| R-HSA-72165 | mRNA Splicing - Minor Pathway | SNRNP25 | small nuclear ribonucleoprotein U11/U1 | 79622 | down | 1,54 | 4,79E-02 |
| R-HSA-111997 | CaM pathway | CALM1 | calmodulin 1 | 801 | down | 1,43 | 2,23E-02 |
| R-HSA-111997 | CaM pathway | CALM2 | calmodulin 2 | 805 | down | 1,43 | 2,23E-02 |
| R-HSA-111997 | CaM pathway | CALM3 | calmodulin 3 | 808 | down | 1,43 | 2,23E-02 |
| R-HSA-1834949 | Cytosolic sensors of pathogen-associated DNA | POLR2F | RNA polymerase II subunit F | 5435 | down | 1,41 | 3,22E-02 |
| R-HSA-1834949 | Cytosolic sensors of pathogen-associated DNA | POLR2L | RNA polymerase II subunit L | 5441 | down | 1,52 | 3,58E-03 |
| R-HSA-1834949 | Cytosolic sensors of pathogen-associated DNA | POLR3F | RNA polymerase III subunit F | 10621 | up | 1,25 | 3,24E-02 |
| R-HSA-1834949 | Cytosolic sensors of pathogen-associated DNA | POLR2E | RNA polymerase II subunit E | 5434 | down | 1,31 | 3,18E-03 |
| R-HSA-1834949 | Cytosolic sensors of pathogen-associated DNA | POLR3H | RNA polymerase III subunit H | 171568 | down | 1,80 | 7,07E-04 |
| R-HSA-1834949 | Cytosolic sensors of pathogen-associated DNA | POLR3D | RNA polymerase III subunit D | 661 | up | 1,52 | 2,72E-02 |
| R-HSA-1834949 | Cytosolic sensors of pathogen-associated DNA | POLR3C | RNA polymerase III subunit C | 10623 | up | 1,43 | 4,98E-02 |
| R-HSA-1834949 | Cytosolic sensors of pathogen-associated DNA | POLR1C | RNA polymerase I subunit C | 9533 | down | 1,24 | 4,71E-02 |
| R-HSA-180024 | DARPP-32 events | PPP1CA | protein phosphatase 1 catalytic subunit | 5499 | down | 1,24 | 9,77E-03 |
| R-HSA-180024 | DARPP-32 events | PPP2CB | protein phosphatase 2 catalytic subunit | 5516 | up | 1,30 | 4,79E-02 |
| R-HSA-180024 | DARPP-32 events | PPP3CC | protein phosphatase 3 catalytic subunit | 5533 | up | 1,52 | 1,78E-02 |
| R-HSA-180024 | DARPP-32 events | CALM3 | calmodulin 3 | 808 | down | 1,43 | 2,23E-02 |
| R-HSA-180024 | DARPP-32 events | CALM2 | calmodulin 2 | 805 | down | 1,43 | 2,23E-02 |
| R-HSA-180024 | DARPP-32 events | CALM1 | calmodulin 1 | 801 | down | 1,43 | 2,23E-02 |
| R-HSA-180024 | DARPP-32 events | PDE4B | phosphodiesterase 4B | 5142 | down | 2,82 | 4,90E-02 |
| R-HSA-180024 | DARPP-32 events | PRKAR1B | protein kinase cAMP-dependent type I r | 5575 | down | 1,50 | 6,80E-04 |
| R-HSA-69205 | G1/S-Specific Transcription | CDC25A | cell division cycle 25A | 993 | down | 1,92 | 1,48E-02 |
| R-HSA-69205 | G1/S-Specific Transcription | TFDP1 | transcription factor Dp-1 | 7027 | down | 1,77 | 1,08E-02 |
| R-HSA-69205 | G1/S-Specific Transcription | PCNA | proliferating cell nuclear antigen | 5111 | down | 2,14 | 4,63E-02 |
| R-HSA-69205 | G1/S-Specific Transcription | FBXO5 | F-box protein 5 | 26271 | down | 1,57 | 4,56E-02 |
| R-HSA-69205 | G1/S-Specific Transcription | E2F1 | E2F transcription factor 1 | 1869 | down | 1,64 | 3,25E-02 |
| R-HSA-113510 | E2F mediated regulation of DNA replication | E2F1 | E2F transcription factor 1 | 1869 | down | 1,64 | 3,25E-02 |
| R-HSA-113510 | E2F mediated regulation of DNA replication | FBXO5 | F-box protein 5 | 26271 | down | 1,57 | 4,56E-02 |
| R-HSA-113510 | E2F mediated regulation of DNA replication | PCNA | proliferating cell nuclear antigen | 5111 | down | 2,14 | 4,63E-02 |
| R-HSA-113510 | E2F mediated regulation of DNA replication | TFDP1 | transcription factor Dp-1 | 7027 | down | 1,77 | 1,08E-02 |
| R-HSA-113510 | E2F mediated regulation of DNA replication | CDC25A | cell division cycle 25A | 993 | down | 1,92 | 1,48E-02 |
| R-HSA-111932 | CaMK IV-mediated phosphorylation of CREB | CALM1 | calmodulin 1 | 801 | down | 1,43 | 2,23E-02 |
| R-HSA-111932 | CaMK IV-mediated phosphorylation of CREB | CREB1 | cAMP responsive element binding prote | 1385 | up | 1,51 | 7,18E-03 |
| R-HSA-111932 | CaMK IV-mediated phosphorylation of CREB | CALM3 | calmodulin 3 | 808 | down | 1,43 | 2,23E-02 |
| R-HSA-111932 | CaMK IV-mediated phosphorylation of CREB | CALM2 | calmodulin 2 | 805 | down | 1,43 | 2,23E-02 |
| R-HSA-77075 | RNA Pol II CTD phosphorylation and interaction with | ERCC3 | ERCC excision repair 3, TFIIH core com | 2071 | down | 1,20 | 2,37E-02 |
| R-HSA-77075 | RNA Pol II CTD phosphorylation and interaction with | ERCC2 | ERCC excision repair 2, TFIIH core com | 2068 | down | 1,29 | 2,20E-02 |
| R-HSA-77075 | RNA Pol II CTD phosphorylation and interaction with | CDK7 | cyclin dependent kinase 7 | 1022 | up | 1,37 | 4,38E-02 |
| R-HSA-77075 | RNA Pol II CTD phosphorylation and interaction with | POLR2E | RNA polymerase II subunit E | 5434 | down | 1,31 | 3,18E-03 |
| R-HSA-77075 | RNA Pol II CTD phosphorylation and interaction with | POLR2L | RNA polymerase II subunit L | 5441 | down | 1,52 | 3,58E-03 |
| R-HSA-77075 | RNA Pol II CTD phosphorylation and interaction with | POLR2F | RNA polymerase II subunit F | 5435 | down | 1,41 | 3,22E-02 |
| R-HSA-77075 | RNA Pol II CTD phosphorylation and interaction with | GTF2H1 | general transcription factor IIH subunit 1 | 2965 | up | 1,33 | 1,85E-02 |
| R-HSA-77075 | RNA Pol II CTD phosphorylation and interaction with | POLR2B | RNA polymerase II subunit B | 5431 | down | 1,23 | 3,81E-02 |
| R-HSA-167160 | RNA Pol II CTD phosphorylation and interaction with | POLR2E | RNA polymerase II subunit E | 5434 | down | 1,31 | 3,18E-03 |
| R-HSA-167160 | RNA Pol II CTD phosphorylation and interaction with | POLR2L | RNA polymerase II subunit L | 5441 | down | 1,52 | 3,58E-03 |
| R-HSA-167160 | RNA Pol II CTD phosphorylation and interaction with | POLR2F | RNA polymerase II subunit F | 5435 | down | 1,41 | 3,22E-02 |
| R-HSA-167160 | RNA Pol II CTD phosphorylation and interaction with | GTF2H1 | general transcription factor IIH subunit 1 | 2965 | up | 1,33 | 1,85E-02 |
| R-HSA-167160 | RNA Pol II CTD phosphorylation and interaction with | POLR2B | RNA polymerase II subunit B | 5431 | down | 1,23 | 3,81E-02 |
| R-HSA-167160 | RNA Pol II CTD phosphorylation and interaction with | ERCC2 | ERCC excision repair 2, TFIIH core com | 2068 | down | 1,29 | 2,20E-02 |
| R-HSA-167160 | RNA Pol II CTD phosphorylation and interaction with | ERCC3 | ERCC excision repair 3, TFIIH core com | 2071 | down | 1,20 | 2,37E-02 |
| R-HSA-167160 | RNA Pol II CTD phosphorylation and interaction with | CDK7 | cyclin dependent kinase 7 | 1022 | up | 1,37 | 4,38E-02 |
| R-HSA-69183 | Processive synthesis on the lagging strand | PCNA | proliferating cell nuclear antigen | 5111 | down | 2,14 | 4,63E-02 |
| R-HSA-69183 | Processive synthesis on the lagging strand | POLD2 | DNA polymerase delta 2, accessory sub | 5425 | down | 1,30 | 1,92E-02 |
| R-HSA-69183 | Processive synthesis on the lagging strand | POLA2 | DNA polymerase alpha 2, accessory sub | 23649 | down | 1,72 | 2,46E-02 |
| R-HSA-69183 | Processive synthesis on the lagging strand | POLD1 | DNA polymerase delta 1, catalytic subur | 5424 | down | 1,59 | 4,62E-02 |
| R-HSA-442745 | No details found for 442745 | CALM1 | calmodulin 1 | 801 | down | 1,43 | 2,23E-02 |
| R-HSA-442745 | No details found for 442745 | CALM2 | calmodulin 2 | 805 | down | 1,43 | 2,23E-02 |
| R-HSA-442745 | No details found for 442745 | CALM3 | calmodulin 3 | 808 | down | 1,43 | 2,23E-02 |
| R-HSA-6807047 | Cholesterol biosynthesis via desmosterol | DHCR24 | 24-dehydrocholesterol reductase | 1718 | down | 2,17 | 4,89E-02 |
| R-HSA-6807047 | Cholesterol biosynthesis via desmosterol | DHCR7 | 7-dehydrocholesterol reductase | 1717 | down | 2,00 | 2,13E-03 |
| R-HSA-6807047 | Cholesterol biosynthesis via desmosterol | EBP | emopamil binding protein (sterol isomera | 10682 | down | 1,71 | 1,99E-02 |
| R-HSA-6807062 | Cholesterol biosynthesis via lathosterol | DHCR24 | 24-dehydrocholesterol reductase | 1718 | down | 2,17 | 4,89E-02 |
| R-HSA-6807062 | Cholesterol biosynthesis via lathosterol | EBP | emopamil binding protein (sterol isomera | 10682 | down | 1,71 | 1,99E-02 |
| R-HSA-6807062 | Cholesterol biosynthesis via lathosterol | DHCR7 | 7-dehydrocholesterol reductase | 1717 | down | 2,00 | 2,13E-03 |
| R-HSA-380994 | ATF4 activates genes | DDIT3 | DNA damage inducible transcript 3 | 1649 | up | 5,65 | 3,67E-03 |
| R-HSA-380994 | ATF4 activates genes | ATF3 | activating transcription factor 3 | 467 | up | 9,22 | 4,75E-04 |
| R-HSA-380994 | ATF4 activates genes | ATF6 | activating transcription factor 6 | 22926 | up | 1,68 | 1,14E-02 |
| R-HSA-380994 | ATF4 activates genes | EXOSC4 | exosome component 4 | 54512 | down | 1,51 | 2,76E-03 |
| R-HSA-380994 | ATF4 activates genes | ASNS | asparagine synthetase (glutamine-hydro | 440 | up | 2,47 | 2,91E-02 |
| R-HSA-380994 | ATF4 activates genes | HERPUD1 | homocysteine inducible ER protein with | 9709 | up | 4,55 | 1,11E-03 |
| R-HSA-380994 | ATF4 activates genes | EXOSC6 | exosome component 6 | 118460 | down | 1,50 | 1,04E-02 |
| R-HSA-380994 | ATF4 activates genes | NFYA | nuclear transcription factor Y subunit alp | 4800 | up | 1,35 | 3,30E-02 |
| R-HSA-72086 | mRNA Capping | POLR2E | RNA polymerase II subunit E | 5434 | down | 1,31 | 3,18E-03 |
| R-HSA-72086 | mRNA Capping | POLR2B | RNA polymerase II subunit B | 5431 | down | 1,23 | 3,81E-02 |
| R-HSA-72086 | mRNA Capping | POLR2F | RNA polymerase II subunit F | 5435 | down | 1,41 | 3,22E-02 |
| R-HSA-72086 | mRNA Capping | GTF2H1 | general transcription factor IIH subunit 1 | 2965 | up | 1,33 | 1,85E-02 |
| R-HSA-72086 | mRNA Capping | POLR2L | RNA polymerase II subunit L | 5441 | down | 1,52 | 3,58E-03 |
| R-HSA-72086 | mRNA Capping | ERCC2 | ERCC excision repair 2, TFIIH core com | 2068 | down | 1,29 | 2,20E-02 |
| R-HSA-72086 | mRNA Capping | ERCC3 | ERCC excision repair 3, TFIIH core com | 2071 | down | 1,20 | 2,37E-02 |
| R-HSA-72086 | mRNA Capping | CDK7 | cyclin dependent kinase 7 | 1022 | up | 1,37 | 4,38E-02 |
| R-HSA-5218920 | VEGFR2 mediated vascular permeability | AKT1 | AKT serine/threonine kinase 1 | 207 | down | 1,37 | 1,42E-03 |
| R-HSA-5218920 | VEGFR2 mediated vascular permeability | CALM1 | calmodulin 1 | 801 | down | 1,43 | 2,23E-02 |
| R-HSA-5218920 | VEGFR2 mediated vascular permeability | AKT2 | AKT serine/threonine kinase 2 | 208 | down | 1,28 | 6,33E-03 |
| R-HSA-5218920 | VEGFR2 mediated vascular permeability | PRR5 | proline rich 5 | 55615 | down | 1,41 | 1,79E-03 |
| R-HSA-5218920 | VEGFR2 mediated vascular permeability | TRIB3 | tribbles pseudokinase 3 | 57761 | up | 3,17 | 8,07E-03 |
| R-HSA-5218920 | VEGFR2 mediated vascular permeability | CALM3 | calmodulin 3 | 808 | down | 1,43 | 2,23E-02 |
| R-HSA-5218920 | VEGFR2 mediated vascular permeability | CALM2 | calmodulin 2 | 805 | down | 1,43 | 2,23E-02 |
| R-HSA-381070 | IRE1alpha activates chaperones | HSPA5 | heat shock protein family A (Hsp70) mem | 3309 | up | 5,02 | 6,98E-04 |
| R-HSA-381070 | IRE1alpha activates chaperones | ERN1 | endoplasmic reticulum to nucleus signal | 2081 | up | 3,27 | 7,84E-03 |
| R-HSA-381070 | IRE1alpha activates chaperones | XBP1 | X-box binding protein 1 | 7494 | up | 2,00 | 5,67E-03 |
| R-HSA-1368082 | RORA activates gene expression | CHD9 | chromodomain helicase DNA binding pr | 80205 | up | 1,32 | 1,42E-03 |
| R-HSA-1368082 | RORA activates gene expression | SREBF1 | sterol regulatory element binding transc | 6720 | down | 1,42 | 3,67E-02 |
| R-HSA-1368082 | RORA activates gene expression | TBL1X | transducin beta like 1 X-linked | 6907 | down | 1,20 | 2,84E-02 |
| R-HSA-1368082 | RORA activates gene expression | RXRA | retinoid X receptor alpha | 6256 | down | 1,34 | 3,67E-03 |
| R-HSA-1368082 | RORA activates gene expression | CARM1 | coactivator associated arginine methyltr | 10498 | down | 1,36 | 1,60E-02 |
| R-HSA-1368082 | RORA activates gene expression | TGS1 | trimethylguanosine synthase 1 | 96764 | up | 1,63 | 9,37E-03 |
| R-HSA-1368082 | RORA activates gene expression | SIRT1 | sirtuin 1 | 23411 | up | 1,60 | 1,00E-02 |
| R-HSA-1368082 | RORA activates gene expression | ARNTL | aryl hydrocarbon receptor nuclear transl | 406 | up | 1,67 | 5,94E-03 |
| R-HSA-203615 | eNOS activation | CALM1 | calmodulin 1 | 801 | down | 1,43 | 2,23E-02 |
| R-HSA-203615 | eNOS activation | AKT1 | AKT serine/threonine kinase 1 | 207 | down | 1,37 | 1,42E-03 |
| R-HSA-203615 | eNOS activation | CALM2 | calmodulin 2 | 805 | down | 1,43 | 2,23E-02 |
| R-HSA-203615 | eNOS activation | CALM3 | calmodulin 3 | 808 | down | 1,43 | 2,23E-02 |
| R-HSA-983170 | Antigen Presentation: Folding, assembly and peptide | SEC31A | SEC31 homolog A, COPII coat complex | 22872 | up | 1,53 | 3,02E-02 |
| R-HSA-983170 | Antigen Presentation: Folding, assembly and peptide | HSPA5 | heat shock protein family A (Hsp70) mem | 3309 | up | 5,02 | 6,98E-04 |
| R-HSA-983170 | Antigen Presentation: Folding, assembly and peptide | SEC24A | SEC24 homolog A, COPII coat complex | 10802 | up | 1,71 | 3,95E-03 |
| R-HSA-983170 | Antigen Presentation: Folding, assembly and peptide | SEC24D | SEC24 homolog D, COPII coat complex | 9871 | up | 4,45 | 6,80E-04 |
| R-HSA-983170 | Antigen Presentation: Folding, assembly and peptide | SEC23A | Sec23 homolog A, coat complex II com | 10484 | up | 1,46 | 2,47E-02 |
| R-HSA-983170 | Antigen Presentation: Folding, assembly and peptide | CANX | calnexin | 821 | up | 1,36 | 3,98E-02 |
| R-HSA-1483166 | Synthesis of PA | LPCAT4 | lysophosphatidylcholine acyltransferase | 254531 | down | 1,78 | 2,52E-02 |
| R-HSA-1483166 | Synthesis of PA | AGPAT3 | 1-acylglycerol-3-phosphate O-acyltransf | 56894 | down | 1,30 | 9,80E-03 |
| R-HSA-1483166 | Synthesis of PA | AGPAT5 | 1-acylglycerol-3-phosphate O-acyltransf | 55326 | down | 1,30 | 4,70E-02 |
| R-HSA-1483166 | Synthesis of PA | LPCAT1 | lysophosphatidylcholine acyltransferase | 79888 | down | 1,76 | 2,60E-02 |
| R-HSA-1483166 | Synthesis of PA | GPAT4 | glycerol-3-phosphate acyltransferase 4 | 137964 | down | 1,24 | 4,75E-02 |
| R-HSA-1483166 | Synthesis of PA | AGPAT2 | 1-acylglycerol-3-phosphate O-acyltransf | 10555 | down | 1,23 | 2,89E-03 |
| R-HSA-1483166 | Synthesis of PA | PLD1 | phospholipase D1 | 5337 | up | 1,33 | 1,60E-02 |
| R-HSA-1483166 | Synthesis of PA | GPAT3 | glycerol-3-phosphate acyltransferase 3 | 84803 | up | 1,77 | 4,99E-02 |
| R-HSA-174417 | Telomere C-strand (Lagging Strand) Synthesis | PCNA | proliferating cell nuclear antigen | 5111 | down | 2,14 | 4,63E-02 |
| R-HSA-174417 | Telomere C-strand (Lagging Strand) Synthesis | POLD2 | DNA polymerase delta 2, accessory sub | 5425 | down | 1,30 | 1,92E-02 |
| R-HSA-174417 | Telomere C-strand (Lagging Strand) Synthesis | POLD1 | DNA polymerase delta 1, catalytic subur | 5424 | down | 1,59 | 4,62E-02 |
| R-HSA-167158 | Formation of the HIV-1 Early Elongation Complex | POLR2F | RNA polymerase II subunit F | 5435 | down | 1,41 | 3,22E-02 |
| R-HSA-167158 | Formation of the HIV-1 Early Elongation Complex | GTF2H1 | general transcription factor IIH subunit 1 | 2965 | up | 1,33 | 1,85E-02 |
| R-HSA-167158 | Formation of the HIV-1 Early Elongation Complex | POLR2L | RNA polymerase II subunit L | 5441 | down | 1,52 | 3,58E-03 |
| R-HSA-167158 | Formation of the HIV-1 Early Elongation Complex | POLR2B | RNA polymerase II subunit B | 5431 | down | 1,23 | 3,81E-02 |
| R-HSA-167158 | Formation of the HIV-1 Early Elongation Complex | POLR2E | RNA polymerase II subunit E | 5434 | down | 1,31 | 3,18E-03 |
| R-HSA-167158 | Formation of the HIV-1 Early Elongation Complex | CDK7 | cyclin dependent kinase 7 | 1022 | up | 1,37 | 4,38E-02 |
| R-HSA-167158 | Formation of the HIV-1 Early Elongation Complex | ERCC2 | ERCC excision repair 2, TFIIH core com | 2068 | down | 1,29 | 2,20E-02 |
| R-HSA-167158 | Formation of the HIV-1 Early Elongation Complex | ERCC3 | ERCC excision repair 3, TFIIH core com | 2071 | down | 1,20 | 2,37E-02 |
| R-HSA-113418 | Formation of the Early Elongation Complex | POLR2B | RNA polymerase II subunit B | 5431 | down | 1,23 | 3,81E-02 |
| R-HSA-113418 | Formation of the Early Elongation Complex | POLR2F | RNA polymerase II subunit F | 5435 | down | 1,41 | 3,22E-02 |
| R-HSA-113418 | Formation of the Early Elongation Complex | GTF2H1 | general transcription factor IIH subunit 1 | 2965 | up | 1,33 | 1,85E-02 |
| R-HSA-113418 | Formation of the Early Elongation Complex | POLR2L | RNA polymerase II subunit L | 5441 | down | 1,52 | 3,58E-03 |
| R-HSA-113418 | Formation of the Early Elongation Complex | POLR2E | RNA polymerase II subunit E | 5434 | down | 1,31 | 3,18E-03 |
| R-HSA-113418 | Formation of the Early Elongation Complex | CDK7 | cyclin dependent kinase 7 | 1022 | up | 1,37 | 4,38E-02 |
| R-HSA-113418 | Formation of the Early Elongation Complex | ERCC2 | ERCC excision repair 2, TFIIH core com | 2068 | down | 1,29 | 2,20E-02 |
| R-HSA-113418 | Formation of the Early Elongation Complex | ERCC3 | ERCC excision repair 3, TFIIH core com | 2071 | down | 1,20 | 2,37E-02 |
| R-HSA-5626467 | RHO GTPases activate IQGAPs | CALM2 | calmodulin 2 | 805 | down | 1,43 | 2,23E-02 |
| R-HSA-5626467 | RHO GTPases activate IQGAPs | CALM3 | calmodulin 3 | 808 | down | 1,43 | 2,23E-02 |
| R-HSA-5626467 | RHO GTPases activate IQGAPs | CLIP1 | CAP-Gly domain containing linker protei | 6249 | up | 1,72 | 1,42E-03 |
| R-HSA-5626467 | RHO GTPases activate IQGAPs | CALM1 | calmodulin 1 | 801 | down | 1,43 | 2,23E-02 |
| R-HSA-5626467 | RHO GTPases activate IQGAPs | TUBA4A | tubulin alpha 4a | 7277 | down | 1,63 | 2,97E-02 |
| R-HSA-5626467 | RHO GTPases activate IQGAPs | TUBA1B | tubulin alpha 1b | 10376 | down | 1,78 | 2,30E-02 |
| R-HSA-5626467 | RHO GTPases activate IQGAPs | ACTG1 | actin gamma 1 | 71 | up | 1,58 | 1,66E-02 |
| R-HSA-5626467 | RHO GTPases activate IQGAPs | MEN1 | menin 1 | 4221 | down | 1,49 | 4,75E-04 |
| R-HSA-6781827 | Transcription-Coupled Nucleotide Excision Repair (TC | POLR2E | RNA polymerase II subunit E | 5434 | down | 1,31 | 3,18E-03 |
| R-HSA-6781827 | Transcription-Coupled Nucleotide Excision Repair (TC | POLR2L | RNA polymerase II subunit L | 5441 | down | 1,52 | 3,58E-03 |
| R-HSA-6781827 | Transcription-Coupled Nucleotide Excision Repair (TC | POLR2F | RNA polymerase II subunit F | 5435 | down | 1,41 | 3,22E-02 |
| R-HSA-6781827 | Transcription-Coupled Nucleotide Excision Repair (TC | GTF2H1 | general transcription factor IIH subunit 1 | 2965 | up | 1,33 | 1,85E-02 |
| R-HSA-6781827 | Transcription-Coupled Nucleotide Excision Repair (TC | TCEA1 | transcription elongation factor A1 | 6917 | up | 1,76 | 1,81E-02 |
| R-HSA-6781827 | Transcription-Coupled Nucleotide Excision Repair (TC | PRPF19 | pre-mRNA processing factor 19 | 27339 | down | 1,24 | 4,41E-02 |
| R-HSA-6781827 | Transcription-Coupled Nucleotide Excision Repair (TC | POLR2B | RNA polymerase II subunit B | 5431 | down | 1,23 | 3,81E-02 |
| R-HSA-6781827 | Transcription-Coupled Nucleotide Excision Repair (TC | ERCC3 | ERCC excision repair 3, TFIIH core com | 2071 | down | 1,20 | 2,37E-02 |
| R-HSA-6781827 | Transcription-Coupled Nucleotide Excision Repair (TC | ERCC2 | ERCC excision repair 2, TFIIH core com | 2068 | down | 1,29 | 2,20E-02 |
| R-HSA-6781827 | Transcription-Coupled Nucleotide Excision Repair (TC | CDK7 | cyclin dependent kinase 7 | 1022 | up | 1,37 | 4,38E-02 |
| R-HSA-418359 | Reduction of cytosolic Ca++ levels | CALM1 | calmodulin 1 | 801 | down | 1,43 | 2,23E-02 |
| R-HSA-418359 | Reduction of cytosolic Ca++ levels | ATP2B1 | ATPase plasma membrane Ca2+ transp | 490 | up | 1,35 | 4,17E-02 |

| | | | | | | | |
|---|---|---|---|---|---|---|---|
| R-HSA-418359 | Reduction of cytosolic Ca++ levels | CALM2 | calmodulin 2 | 805 | down | 1,43 | 2,23E-02 |
| R-HSA-418359 | Reduction of cytosolic Ca++ levels | ATP2A3 | ATPase sarcoplasmic/endoplasmic retic | 489 | down | 1,62 | 4,34E-02 |
| R-HSA-418359 | Reduction of cytosolic Ca++ levels | CALM3 | calmodulin 3 | 808 | down | 1,43 | 2,23E-02 |
| R-HSA-156581 | Methylation | NNMT | nicotinamide N-methyltransferase | 4837 | down | 2,26 | 3,05E-02 |
| R-HSA-156581 | Methylation | MAT2A | methionine adenosyltransferase 2A | 4144 | down | 1,79 | 2,93E-02 |
| R-HSA-156581 | Methylation | COMT | catechol-O-methyltransferase | 1312 | down | 1,30 | 1,09E-02 |
| R-HSA-156581 | Methylation | AHCY | adenosylhomocysteinase | 191 | down | 1,14 | 2,15E-02 |
| R-HSA-174411 | Polymerase switching on the C-strand of the telomere | POLA2 | DNA polymerase alpha 2, accessory sub | 23649 | down | 1,72 | 2,46E-02 |
| R-HSA-174411 | Polymerase switching on the C-strand of the telomere | RFC1 | replication factor C subunit 1 | 5981 | up | 1,15 | 2,18E-02 |
| R-HSA-174411 | Polymerase switching on the C-strand of the telomere | PCNA | proliferating cell nuclear antigen | 5111 | down | 2,14 | 4,63E-02 |
| R-HSA-174411 | Polymerase switching on the C-strand of the telomere | POLD2 | DNA polymerase delta 2, accessory sub | 5425 | down | 1,30 | 1,92E-02 |
| R-HSA-174411 | Polymerase switching on the C-strand of the telomere | POLD1 | DNA polymerase delta 1, catalytic subur | 5424 | down | 1,59 | 4,62E-02 |
| R-HSA-69091 | Polymerase switching | POLA2 | DNA polymerase alpha 2, accessory sub | 23649 | down | 1,72 | 2,46E-02 |
| R-HSA-69091 | Polymerase switching | POLD2 | DNA polymerase delta 2, accessory sub | 5425 | down | 1,30 | 1,92E-02 |
| R-HSA-69091 | Polymerase switching | PCNA | proliferating cell nuclear antigen | 5111 | down | 2,14 | 4,63E-02 |
| R-HSA-69091 | Polymerase switching | RFC1 | replication factor C subunit 1 | 5981 | up | 1,15 | 2,18E-02 |
| R-HSA-69091 | Polymerase switching | POLD1 | DNA polymerase delta 1, catalytic subur | 5424 | down | 1,59 | 4,62E-02 |
| R-HSA-442729 | CREB1 phosphorylation through the activation of Cal | AKAP9 | A-kinase anchoring protein 9 | 10142 | up | 1,47 | 1,80E-02 |
| R-HSA-442729 | CREB1 phosphorylation through the activation of Cal | CAMK2D | calcium/calmodulin dependent protein k | 817 | up | 1,29 | 1,49E-02 |
| R-HSA-442729 | CREB1 phosphorylation through the activation of Cal | CREB1 | cAMP responsive element binding prote | 1385 | up | 1,51 | 7,18E-03 |
| R-HSA-442729 | CREB1 phosphorylation through the activation of Cal | CALM3 | calmodulin 3 | 808 | down | 1,43 | 2,23E-02 |
| R-HSA-442729 | CREB1 phosphorylation through the activation of Cal | CALM2 | calmodulin 2 | 805 | down | 1,43 | 2,23E-02 |
| R-HSA-442729 | CREB1 phosphorylation through the activation of Cal | CALM1 | calmodulin 1 | 801 | down | 1,43 | 2,23E-02 |
| R-HSA-450513 | Tristetraprolin (TTP, ZFP36) binds and destabilizes m | XRN1 | 5'-3' exoribonuclease 1 | 54464 | up | 1,30 | 1,96E-02 |
| R-HSA-450513 | Tristetraprolin (TTP, ZFP36) binds and destabilizes m | ZFP36 | ZFP36 ring finger protein | 7538 | up | 1,90 | 4,84E-02 |
| R-HSA-450513 | Tristetraprolin (TTP, ZFP36) binds and destabilizes m | EXOSC4 | exosome component 4 | 54512 | down | 1,51 | 2,76E-03 |
| R-HSA-450513 | Tristetraprolin (TTP, ZFP36) binds and destabilizes m | TNPO1 | transportin 1 | 3842 | up | 1,28 | 4,94E-02 |
| R-HSA-450513 | Tristetraprolin (TTP, ZFP36) binds and destabilizes m | DCP1A | decapping mRNA 1A | 55802 | up | 1,33 | 4,54E-02 |
| R-HSA-450513 | Tristetraprolin (TTP, ZFP36) binds and destabilizes m | EXOSC6 | exosome component 6 | 118460 | down | 1,50 | 1,04E-02 |
| R-HSA-983168 | Antigen processing: Ubiquitination & Proteasome deg | ASB1 | ankyrin repeat and SOCS box containin | 51665 | up | 1,81 | 3,60E-02 |
| R-HSA-983168 | Antigen processing: Ubiquitination & Proteasome deg | FBXL19 | F-box and leucine rich repeat protein 19 | 54620 | down | 1,40 | 9,51E-03 |
| R-HSA-983168 | Antigen processing: Ubiquitination & Proteasome deg | UBE2G2 | ubiquitin conjugating enzyme E2 G2 | 7327 | down | 1,41 | 4,12E-03 |
| R-HSA-983168 | Antigen processing: Ubiquitination & Proteasome deg | UBR1 | ubiquitin protein ligase E3 component n | 197131 | up | 1,19 | 2,29E-02 |
| R-HSA-983168 | Antigen processing: Ubiquitination & Proteasome deg | RLIM | ring finger protein, LIM domain interactir | 51132 | up | 1,34 | 9,04E-03 |
| R-HSA-983168 | Antigen processing: Ubiquitination & Proteasome deg | STUB1 | STIP1 homology and U-box containing | 10273 | down | 1,24 | 4,82E-03 |
| R-HSA-983168 | Antigen processing: Ubiquitination & Proteasome deg | ASB13 | ankyrin repeat and SOCS box containin | 79754 | down | 1,87 | 4,56E-03 |
| R-HSA-983168 | Antigen processing: Ubiquitination & Proteasome deg | WSB1 | WD repeat and SOCS box containing 1 | 26118 | up | 1,56 | 2,10E-02 |
| R-HSA-983168 | Antigen processing: Ubiquitination & Proteasome deg | RNF6 | ring finger protein 6 | 6049 | up | 1,45 | 1,98E-02 |
| R-HSA-983168 | Antigen processing: Ubiquitination & Proteasome deg | SH3RF1 | SH3 domain containing ring finger 1 | 57630 | up | 1,79 | 1,11E-02 |
| R-HSA-983168 | Antigen processing: Ubiquitination & Proteasome deg | SIAH1 | siah E3 ubiquitin protein ligase 1 | 6477 | up | 1,74 | 1,66E-02 |
| R-HSA-983168 | Antigen processing: Ubiquitination & Proteasome deg | UFL1 | UFM1 specific ligase 1 | 23376 | up | 1,51 | 1,67E-02 |
| R-HSA-983168 | Antigen processing: Ubiquitination & Proteasome deg | KCTD7 | potassium channel tetramerization doma | 154881 | up | 1,94 | 1,85E-03 |
| R-HSA-983168 | Antigen processing: Ubiquitination & Proteasome deg | CUL7 | cullin 7 | 9820 | up | 1,34 | 4,05E-02 |
| R-HSA-983168 | Antigen processing: Ubiquitination & Proteasome deg | UBA6 | ubiquitin like modifier activating enzyme | 55236 | up | 1,24 | 1,11E-02 |
| R-HSA-983168 | Antigen processing: Ubiquitination & Proteasome deg | UBAC1 | UBA domain containing 1 | 10422 | down | 1,29 | 2,18E-02 |
| R-HSA-983168 | Antigen processing: Ubiquitination & Proteasome deg | CCNF | cyclin F | 899 | down | 2,16 | 6,61E-03 |
| R-HSA-983168 | Antigen processing: Ubiquitination & Proteasome deg | ZNRF1 | zinc and ring finger 1 | 84937 | down | 1,18 | 1,73E-02 |
| R-HSA-983168 | Antigen processing: Ubiquitination & Proteasome deg | FBXL16 | F-box and leucine rich repeat protein 16 | 146330 | down | 2,11 | 4,03E-03 |
| R-HSA-983168 | Antigen processing: Ubiquitination & Proteasome deg | RNF19A | ring finger protein 19A, RBR E3 ubiquiti | 25897 | up | 1,56 | 8,54E-03 |
| R-HSA-983168 | Antigen processing: Ubiquitination & Proteasome deg | UBE2J1 | ubiquitin conjugating enzyme E2 J1 | 51465 | up | 1,86 | 8,97E-03 |
| R-HSA-983168 | Antigen processing: Ubiquitination & Proteasome deg | TRIM4 | tripartite motif containing 4 | 89122 | up | 1,48 | 3,48E-02 |
| R-HSA-983168 | Antigen processing: Ubiquitination & Proteasome deg | TRIM36 | tripartite motif containing 36 | 55521 | up | 2,29 | 6,64E-03 |
| R-HSA-983168 | Antigen processing: Ubiquitination & Proteasome deg | SMURF2 | SMAD specific E3 ubiquitin protein ligas | 64750 | up | 1,49 | 4,16E-02 |
| R-HSA-983168 | Antigen processing: Ubiquitination & Proteasome deg | NEDD4 | neural precursor cell expressed, develop | 4734 | up | 1,99 | 3,68E-03 |
| R-HSA-983168 | Antigen processing: Ubiquitination & Proteasome deg | CBLB | Cbl proto-oncogene B | 868 | up | 1,83 | 4,35E-02 |
| R-HSA-983168 | Antigen processing: Ubiquitination & Proteasome deg | UBE2B | ubiquitin conjugating enzyme E2 B | 7320 | up | 1,65 | 4,38E-02 |
| R-HSA-983168 | Antigen processing: Ubiquitination & Proteasome deg | FBXO9 | F-box protein 9 | 26268 | down | 1,37 | 4,01E-02 |
| R-HSA-983168 | Antigen processing: Ubiquitination & Proteasome deg | RNF41 | ring finger protein 41 | 10193 | up | 1,59 | 2,87E-02 |
| R-HSA-983168 | Antigen processing: Ubiquitination & Proteasome deg | ANAPC2 | anaphase promoting complex subunit 2 | 29882 | down | 1,26 | 1,99E-02 |
| R-HSA-983168 | Antigen processing: Ubiquitination & Proteasome deg | RNF14 | ring finger protein 14 | 9604 | up | 1,52 | 2,07E-02 |
| R-HSA-983168 | Antigen processing: Ubiquitination & Proteasome deg | KLHL2 | kelch like family member 2 | 11275 | up | 1,33 | 3,06E-02 |
| R-HSA-983168 | Antigen processing: Ubiquitination & Proteasome deg | PJA2 | praja ring finger ubiquitin ligase 2 | 9867 | up | 2,21 | 3,10E-03 |
| R-HSA-983168 | Antigen processing: Ubiquitination & Proteasome deg | SKP2 | S-phase kinase associated protein 2 | 6502 | down | 2,21 | 3,69E-02 |
| R-HSA-983168 | Antigen processing: Ubiquitination & Proteasome deg | ITCH | itchy E3 ubiquitin protein ligase | 83737 | up | 1,51 | 6,64E-03 |
| R-HSA-983168 | Antigen processing: Ubiquitination & Proteasome deg | TPP2 | tripeptidyl peptidase 2 | 7174 | up | 1,25 | 4,04E-02 |
| R-HSA-983168 | Antigen processing: Ubiquitination & Proteasome deg | TRIM32 | tripartite motif containing 32 | 22954 | down | 1,22 | 3,95E-02 |
| R-HSA-983168 | Antigen processing: Ubiquitination & Proteasome deg | RNF19B | ring finger protein 19B | 127544 | up | 2,61 | 4,56E-03 |
| R-HSA-983168 | Antigen processing: Ubiquitination & Proteasome deg | HECTD1 | HECT domain E3 ubiquitin protein ligase | 25831 | up | 1,43 | 1,57E-02 |
| R-HSA-1989781 | PPARA activates gene expression | CHD9 | chromodomain helicase DNA binding pr | 80205 | up | 1,32 | 1,42E-03 |
| R-HSA-1989781 | PPARA activates gene expression | CTGF | connective tissue growth factor | 1490 | up | 2,49 | 1,32E-02 |
| R-HSA-1989781 | PPARA activates gene expression | RXRA | retinoid X receptor alpha | 6256 | down | 1,34 | 3,67E-03 |
| R-HSA-1989781 | PPARA activates gene expression | ARNTL | aryl hydrocarbon receptor nuclear transl | 406 | up | 1,67 | 5,94E-03 |
| R-HSA-1989781 | PPARA activates gene expression | TRIB3 | tribbles pseudokinase 3 | 57761 | up | 3,17 | 8,07E-03 |
| R-HSA-1989781 | PPARA activates gene expression | WWTR1 | WW domain containing transcription reg | 25937 | down | 1,99 | 1,00E-02 |
| R-HSA-1989781 | PPARA activates gene expression | MED25 | mediator complex subunit 25 | 81857 | down | 1,23 | 4,79E-02 |
| R-HSA-1989781 | PPARA activates gene expression | MTF1 | metal regulatory transcription factor 1 | 4520 | up | 1,42 | 3,66E-02 |
| R-HSA-1989781 | PPARA activates gene expression | SREBF1 | sterol regulatory element binding transc | 6720 | down | 1,42 | 3,67E-02 |
| R-HSA-1989781 | PPARA activates gene expression | TGS1 | trimethylguanosine synthase 1 | 96764 | up | 1,63 | 9,37E-03 |
| R-HSA-1989781 | PPARA activates gene expression | CYP1A1 | cytochrome P450 family 1 subfamily A m | 1543 | up | 2,69 | 1,95E-02 |
| R-HSA-1989781 | PPARA activates gene expression | CARM1 | coactivator associated arginine methyltr | 10498 | down | 1,36 | 1,60E-02 |
| R-HSA-1989781 | PPARA activates gene expression | ANKRD1 | ankyrin repeat domain 1 | 27063 | up | 3,48 | 3,01E-02 |
| R-HSA-1989781 | PPARA activates gene expression | TBL1X | transducin beta like 1 X-linked | 6907 | down | 1,20 | 2,84E-02 |
| R-HSA-1989781 | PPARA activates gene expression | NFYA | nuclear transcription factor Y subunit alp | 4800 | up | 1,35 | 3,30E-02 |
| R-HSA-1989781 | PPARA activates gene expression | MED24 | mediator complex subunit 24 | 9862 | down | 1,33 | 2,93E-02 |
| R-HSA-1989781 | PPARA activates gene expression | FHL2 | four and a half LIM domains 2 | 2274 | up | 2,10 | 1,13E-02 |
| R-HSA-1989781 | PPARA activates gene expression | NCOA3 | nuclear receptor coactivator 3 | 8202 | up | 1,53 | 4,13E-02 |
| R-HSA-1989781 | PPARA activates gene expression | ACSL1 | acyl-CoA synthetase long chain family m | 2180 | down | 1,54 | 3,01E-02 |
| R-HSA-1989781 | PPARA activates gene expression | FADS1 | fatty acid desaturase 1 | 3992 | down | 1,54 | 8,39E-03 |
| R-HSA-111957 | Cam-PDE 1 activation | CALM1 | calmodulin 1 | 801 | down | 1,43 | 2,23E-02 |
| R-HSA-111957 | Cam-PDE 1 activation | CALM3 | calmodulin 3 | 808 | down | 1,43 | 2,23E-02 |
| R-HSA-111957 | Cam-PDE 1 activation | CALM2 | calmodulin 2 | 805 | down | 1,43 | 2,23E-02 |
| R-HSA-174414 | Processive synthesis on the C-strand of the telomere | POLD1 | DNA polymerase delta 1, catalytic subur | 5424 | down | 1,59 | 4,62E-02 |
| R-HSA-174414 | Processive synthesis on the C-strand of the telomere | PCNA | proliferating cell nuclear antigen | 5111 | down | 2,14 | 4,63E-02 |
| R-HSA-174414 | Processive synthesis on the C-strand of the telomere | POLD2 | DNA polymerase delta 2, accessory sub | 5425 | down | 1,30 | 1,92E-02 |
| R-HSA-171319 | Telomere Extension By Telomerase | NHP2 | NHP2 ribonucleoprotein | 55651 | down | 1,23 | 3,15E-02 |
| R-HSA-171319 | Telomere Extension By Telomerase | WRAP53 | WD repeat containing antisense to TP53 | 55135 | down | 1,52 | 1,69E-02 |
| R-HSA-171319 | Telomere Extension By Telomerase | DKC1 | dyskerin pseudouridine synthase 1 | 1736 | down | 1,25 | 4,47E-02 |
| R-HSA-5358606 | Mismatch repair (MMR) directed by MSH2:MSH3 (MutS | RPA1 | replication protein A1 | 6117 | down | 1,41 | 4,93E-02 |
| R-HSA-5358606 | Mismatch repair (MMR) directed by MSH2:MSH3 (MutS | PCNA | proliferating cell nuclear antigen | 5111 | down | 2,14 | 4,63E-02 |
| R-HSA-5358606 | Mismatch repair (MMR) directed by MSH2:MSH3 (MutS | POLD2 | DNA polymerase delta 2, accessory sub | 5425 | down | 1,30 | 1,92E-02 |
| R-HSA-5358606 | Mismatch repair (MMR) directed by MSH2:MSH3 (MutS | POLD1 | DNA polymerase delta 1, catalytic subur | 5424 | down | 1,59 | 4,62E-02 |
| R-HSA-111447 | Activation of BAD and translocation to mitochondria | AKT1 | AKT serine/threonine kinase 1 | 207 | down | 1,37 | 1,42E-03 |
| R-HSA-111447 | Activation of BAD and translocation to mitochondria | AKT2 | AKT serine/threonine kinase 2 | 208 | down | 1,28 | 6,33E-03 |
| R-HSA-111447 | Activation of BAD and translocation to mitochondria | YWHAH | tyrosine 3-monooxygenase/tryptophan 5 | 7533 | down | 1,33 | 4,79E-02 |
| R-HSA-111447 | Activation of BAD and translocation to mitochondria | BID | BH3 interacting domain death agonist | 637 | down | 1,40 | 2,42E-02 |
| R-HSA-111447 | Activation of BAD and translocation to mitochondria | PPP3CC | protein phosphatase 3 catalytic subunit | 5533 | up | 1,52 | 1,78E-02 |
| R-HSA-5358565 | Mismatch repair (MMR) directed by MSH2:MSH6 (MutS | POLD1 | DNA polymerase delta 1, catalytic subur | 5424 | down | 1,59 | 4,62E-02 |
| R-HSA-5358565 | Mismatch repair (MMR) directed by MSH2:MSH6 (MutS | PCNA | proliferating cell nuclear antigen | 5111 | down | 2,14 | 4,63E-02 |
| R-HSA-5358565 | Mismatch repair (MMR) directed by MSH2:MSH6 (MutS | POLD2 | DNA polymerase delta 2, accessory sub | 5425 | down | 1,30 | 1,92E-02 |
| R-HSA-5358565 | Mismatch repair (MMR) directed by MSH2:MSH6 (MutS | RPA1 | replication protein A1 | 6117 | down | 1,41 | 4,93E-02 |
| R-HSA-68867 | Assembly of the pre-replicative complex | MCM7 | minichromosome maintenance complex | 4176 | down | 1,67 | 3,35E-02 |
| R-HSA-68867 | Assembly of the pre-replicative complex | MCM5 | minichromosome maintenance complex | 4174 | down | 1,83 | 2,22E-02 |
| R-HSA-68867 | Assembly of the pre-replicative complex | MCM3 | minichromosome maintenance complex | 4172 | down | 2,07 | 4,13E-02 |
| R-HSA-68867 | Assembly of the pre-replicative complex | MCM2 | minichromosome maintenance complex | 4171 | down | 2,01 | 1,42E-02 |
| R-HSA-5696397 | Gap-filling DNA repair synthesis and ligation in GG-NE | RFC1 | replication factor C subunit 1 | 5981 | up | 1,15 | 2,18E-02 |
| R-HSA-5696397 | Gap-filling DNA repair synthesis and ligation in GG-NE | PCNA | proliferating cell nuclear antigen | 5111 | down | 2,14 | 4,63E-02 |
| R-HSA-5696397 | Gap-filling DNA repair synthesis and ligation in GG-NE | POLD2 | DNA polymerase delta 2, accessory sub | 5425 | down | 1,30 | 1,92E-02 |
| R-HSA-5696397 | Gap-filling DNA repair synthesis and ligation in GG-NE | RPA1 | replication protein A1 | 6117 | down | 1,41 | 4,93E-02 |
| R-HSA-5696397 | Gap-filling DNA repair synthesis and ligation in GG-NE | POLE | DNA polymerase epsilon, catalytic subu | 5426 | down | 1,55 | 2,67E-02 |
| R-HSA-5696397 | Gap-filling DNA repair synthesis and ligation in GG-NE | POLD1 | DNA polymerase delta 1, catalytic subur | 5424 | down | 1,59 | 4,62E-02 |
| R-HSA-5674400 | Constitutive Signaling by AKT1 E17K in Cancer | AKT1 | AKT serine/threonine kinase 1 | 207 | down | 1,37 | 1,42E-03 |
| R-HSA-5674400 | Constitutive Signaling by AKT1 E17K in Cancer | AKT2 | AKT serine/threonine kinase 2 | 208 | down | 1,28 | 6,33E-03 |
| R-HSA-5674400 | Constitutive Signaling by AKT1 E17K in Cancer | RPS6KB2 | ribosomal protein S6 kinase B2 | 6199 | down | 1,22 | 1,01E-02 |
| R-HSA-5674400 | Constitutive Signaling by AKT1 E17K in Cancer | CDKN1A | cyclin dependent kinase inhibitor 1A | 1026 | up | 3,62 | 5,43E-03 |
| R-HSA-5674400 | Constitutive Signaling by AKT1 E17K in Cancer | AKT1S1 | AKT1 substrate 1 | 84335 | down | 1,21 | 4,33E-02 |
| R-HSA-5674400 | Constitutive Signaling by AKT1 E17K in Cancer | PRR5 | proline rich 5 | 55615 | down | 1,41 | 1,79E-03 |
| R-HSA-5674400 | Constitutive Signaling by AKT1 E17K in Cancer | CREB1 | cAMP responsive element binding prote | 1385 | up | 1,51 | 7,18E-03 |
| R-HSA-75105 | Fatty acyl-CoA biosynthesis | ACACA | acetyl-CoA carboxylase alpha | 31 | down | 1,40 | 2,54E-02 |
| R-HSA-75105 | Fatty acyl-CoA biosynthesis | SCD | stearoyl-CoA desaturase | 6319 | down | 1,88 | 3,59E-02 |
| R-HSA-75105 | Fatty acyl-CoA biosynthesis | FASN | fatty acid synthase | 2194 | down | 1,96 | 8,31E-03 |
| R-HSA-75105 | Fatty acyl-CoA biosynthesis | ACLY | ATP citrate lyase | 47 | down | 1,41 | 3,09E-02 |
| R-HSA-75105 | Fatty acyl-CoA biosynthesis | SLC25A1 | solute carrier family 25 member 1 | 6576 | down | 1,48 | 5,54E-04 |
| R-HSA-176187 | Activation of ATR in response to replication stress | HUS1 | HUS1 checkpoint clamp component | 3364 | up | 1,36 | 4,96E-02 |
| R-HSA-176187 | Activation of ATR in response to replication stress | MCM2 | minichromosome maintenance complex | 4171 | down | 2,01 | 1,42E-02 |
| R-HSA-176187 | Activation of ATR in response to replication stress | MCM7 | minichromosome maintenance complex | 4176 | down | 1,67 | 3,35E-02 |
| R-HSA-176187 | Activation of ATR in response to replication stress | MCM3 | minichromosome maintenance complex | 4172 | down | 2,07 | 4,13E-02 |
| R-HSA-176187 | Activation of ATR in response to replication stress | MCM5 | minichromosome maintenance complex | 4174 | down | 1,83 | 2,22E-02 |
| R-HSA-176187 | Activation of ATR in response to replication stress | RPA1 | replication protein A1 | 6117 | down | 1,41 | 4,93E-02 |
| R-HSA-176187 | Activation of ATR in response to replication stress | CDC25A | cell division cycle 25A | 993 | down | 1,92 | 1,48E-02 |
| R-HSA-2682334 | EPH-Ephrin signaling | YES1 | YES proto-oncogene 1, Src family tyrosi | 7525 | up | 1,30 | 2,08E-02 |
| R-HSA-2682334 | EPH-Ephrin signaling | LYN | LYN proto-oncogene, Src family tyrosine | 4067 | down | 1,36 | 2,34E-02 |
| R-HSA-2682334 | EPH-Ephrin signaling | EPHB2 | EPH receptor B2 | 2048 | down | 1,66 | 2,52E-02 |
| R-HSA-2682334 | EPH-Ephrin signaling | EPHA2 | EPH receptor A2 | 1969 | up | 1,89 | 2,51E-02 |
| R-HSA-2682334 | EPH-Ephrin signaling | EFNA4 | ephrin A4 | 1945 | down | 1,43 | 4,92E-02 |
| R-HSA-2682334 | EPH-Ephrin signaling | EFNA3 | ephrin A3 | 1944 | down | 2,00 | 4,89E-02 |
| R-HSA-2682334 | EPH-Ephrin signaling | EPHA5 | EPH receptor A5 | 2044 | down | 18,87 | 1,96E-02 |
| R-HSA-73777 | RNA Polymerase I Chain Elongation | POLR2E | RNA polymerase II subunit E | 5434 | down | 1,31 | 3,18E-03 |
| R-HSA-73777 | RNA Polymerase I Chain Elongation | TAF1D | TATA-box binding protein associated fa | 79101 | up | 1,87 | 1,00E-02 |
| R-HSA-73777 | RNA Polymerase I Chain Elongation | POLR1C | RNA polymerase I subunit C | 9533 | down | 1,24 | 4,71E-02 |
| R-HSA-73777 | RNA Polymerase I Chain Elongation | TAF1A | TATA-box binding protein associated fa | 9015 | up | 2,20 | 2,24E-03 |
| R-HSA-73777 | RNA Polymerase I Chain Elongation | CD3EAP | CD3e molecule associated protein | 10849 | down | 1,57 | 3,12E-03 |
| R-HSA-73777 | RNA Polymerase I Chain Elongation | POLR2F | RNA polymerase II subunit F | 5435 | down | 1,41 | 3,22E-02 |
| R-HSA-73777 | RNA Polymerase I Chain Elongation | GTF2H1 | general transcription factor IIH subunit 1 | 2965 | up | 1,33 | 1,85E-02 |
| R-HSA-73777 | RNA Polymerase I Chain Elongation | POLR1A | RNA polymerase I subunit A | 25885 | down | 1,30 | 4,32E-02 |
| R-HSA-73777 | RNA Polymerase I Chain Elongation | POLR2L | RNA polymerase II subunit L | 5441 | down | 1,52 | 3,58E-03 |
| R-HSA-73777 | RNA Polymerase I Chain Elongation | ERCC3 | ERCC excision repair 3, TFIIH core com | 2071 | down | 1,20 | 2,37E-02 |
| R-HSA-73777 | RNA Polymerase I Chain Elongation | ERCC2 | ERCC excision repair 2, TFIIH core com | 2068 | down | 1,29 | 2,20E-02 |
| R-HSA-73777 | RNA Polymerase I Chain Elongation | H2AFX | H2A histone family member X | 3014 | down | 2,11 | 1,30E-03 |
| R-HSA-73777 | RNA Polymerase I Chain Elongation | CDK7 | cyclin dependent kinase 7 | 1022 | up | 1,37 | 4,38E-02 |
| R-HSA-73777 | RNA Polymerase I Chain Elongation | UBTF | upstream binding transcription factor, RN | 7343 | down | 1,66 | 1,20E-02 |
| R-HSA-72163 | mRNA Splicing - Major Pathway | SRSF2 | serine and arginine rich splicing factor 2 | 6427 | down | 1,69 | 1,79E-02 |
| R-HSA-72163 | mRNA Splicing - Major Pathway | BCAS2 | BCAS2, pre-mRNA processing factor | 10286 | up | 1,57 | 2,14E-02 |
| R-HSA-72163 | mRNA Splicing - Major Pathway | POLR2E | RNA polymerase II subunit E | 5434 | down | 1,31 | 3,18E-03 |
| R-HSA-72163 | mRNA Splicing - Major Pathway | CLP1 | cleavage and polyadenylation factor I s | 10978 | down | 1,40 | 1,04E-02 |
| R-HSA-72163 | mRNA Splicing - Major Pathway | POLR2F | RNA polymerase II subunit F | 5435 | down | 1,41 | 3,22E-02 |
| R-HSA-72163 | mRNA Splicing - Major Pathway | SLU7 | SLU7 homolog, splicing factor | 10569 | up | 1,24 | 3,63E-02 |
| R-HSA-72163 | mRNA Splicing - Major Pathway | CHERP | calcium homeostasis endoplasmic reticu | 10523 | down | 1,20 | 2,06E-02 |
| R-HSA-72163 | mRNA Splicing - Major Pathway | PTBP1 | polypyrimidine tract binding protein 1 | 5725 | down | 1,16 | 4,21E-02 |
| R-HSA-72163 | mRNA Splicing - Major Pathway | EFTUD2 | elongation factor Tu GTP binding doma | 9343 | down | 1,24 | 3,99E-02 |
| R-HSA-72163 | mRNA Splicing - Major Pathway | POLR2L | RNA polymerase II subunit L | 5441 | down | 1,52 | 3,58E-03 |
| R-HSA-72163 | mRNA Splicing - Major Pathway | PRPF19 | pre-mRNA processing factor 19 | 27339 | down | 1,24 | 4,41E-02 |
| R-HSA-72163 | mRNA Splicing - Major Pathway | POLR2B | RNA polymerase II subunit B | 5431 | down | 1,23 | 3,81E-02 |
| R-HSA-72163 | mRNA Splicing - Major Pathway | PRPF40A | pre-mRNA processing factor 40 homolog | 55660 | up | 1,18 | 3,23E-02 |
| R-HSA-72163 | mRNA Splicing - Major Pathway | CSTF2 | cleavage stimulation factor subunit 2 | 1478 | down | 1,45 | 4,38E-02 |
| R-HSA-72163 | mRNA Splicing - Major Pathway | U2AF2 | U2 small nuclear RNA auxiliary factor 2 | 11338 | down | 1,27 | 8,21E-03 |
| R-HSA-72163 | mRNA Splicing - Major Pathway | LSM4 | LSM4 homolog, U6 small nuclear RNA a | 25804 | down | 1,40 | 2,18E-02 |
| R-HSA-72163 | mRNA Splicing - Major Pathway | PLRG1 | pleiotropic regulator 1 | 5356 | up | 1,28 | 3,18E-02 |
| R-HSA-72163 | mRNA Splicing - Major Pathway | DDX23 | DEAD-box helicase 23 | 9416 | down | 1,32 | 1,25E-02 |
| R-HSA-72163 | mRNA Splicing - Major Pathway | HNRNPUL1 | heterogeneous nuclear ribonucleoprote | 11100 | down | 1,17 | 2,87E-02 |
| R-HSA-72163 | mRNA Splicing - Major Pathway | PRPF31 | pre-mRNA processing factor 31 | 26121 | down | 1,21 | 2,74E-02 |
| R-HSA-72163 | mRNA Splicing - Major Pathway | SYF2 | SYF2 pre-mRNA splicing factor | 25949 | up | 1,27 | 4,77E-02 |
| R-HSA-72163 | mRNA Splicing - Major Pathway | CWC22 | CWC22 spliceosome associated protein | 57703 | up | 1,41 | 8,63E-03 |
| R-HSA-70221 | Glycogen breakdown (glycogenolysis) | CALM3 | calmodulin 3 | 808 | down | 1,43 | 2,23E-02 |
| R-HSA-70221 | Glycogen breakdown (glycogenolysis) | CALM2 | calmodulin 2 | 805 | down | 1,43 | 2,23E-02 |
| R-HSA-70221 | Glycogen breakdown (glycogenolysis) | CALM1 | calmodulin 1 | 801 | down | 1,43 | 2,23E-02 |
| R-HSA-70221 | Glycogen breakdown (glycogenolysis) | GYG2 | glycogenin 2 | 8908 | down | 1,52 | 4,53E-02 |
| R-HSA-111933 | Calmodulin induced events | CALM3 | calmodulin 3 | 808 | down | 1,43 | 2,23E-02 |
| R-HSA-111933 | Calmodulin induced events | CALM2 | calmodulin 2 | 805 | down | 1,43 | 2,23E-02 |
| R-HSA-111933 | Calmodulin induced events | CALM1 | calmodulin 1 | 801 | down | 1,43 | 2,23E-02 |
| R-HSA-442982 | Ras activation upon Ca2+ influx through NMDA recep | CALM2 | calmodulin 2 | 805 | down | 1,43 | 2,23E-02 |
| R-HSA-442982 | Ras activation upon Ca2+ influx through NMDA recep | HRAS | HRas proto-oncogene, GTPase | 3265 | down | 1,31 | 2,19E-03 |
| R-HSA-442982 | Ras activation upon Ca2+ influx through NMDA recep | CALM3 | calmodulin 3 | 808 | down | 1,43 | 2,23E-02 |
| R-HSA-442982 | Ras activation upon Ca2+ influx through NMDA recep | AKAP9 | A-kinase anchoring protein 9 | 10142 | up | 1,47 | 1,80E-02 |
| R-HSA-442982 | Ras activation upon Ca2+ influx through NMDA recep | CAMK2D | calcium/calmodulin dependent protein k | 817 | up | 1,29 | 1,49E-02 |
| R-HSA-442982 | Ras activation upon Ca2+ influx through NMDA recep | CALM1 | calmodulin 1 | 801 | down | 1,43 | 2,23E-02 |
| R-HSA-381340 | Transcriptional regulation of white adipocyte different | CEBPB | CCAAT/enhancer binding protein beta | 1051 | up | 2,12 | 2,85E-02 |
| R-HSA-381340 | Transcriptional regulation of white adipocyte different | CHD9 | chromodomain helicase DNA binding pr | 80205 | up | 1,32 | 1,42E-03 |
| R-HSA-381340 | Transcriptional regulation of white adipocyte different | SREBF1 | sterol regulatory element binding transc | 6720 | down | 1,42 | 3,67E-02 |
| R-HSA-381340 | Transcriptional regulation of white adipocyte different | TGS1 | trimethylguanosine synthase 1 | 96764 | up | 1,63 | 9,37E-03 |
| R-HSA-381340 | Transcriptional regulation of white adipocyte different | NR2F2 | nuclear receptor subfamily 2 group F me | 7026 | down | 1,46 | 8,31E-03 |
| R-HSA-381340 | Transcriptional regulation of white adipocyte different | CARM1 | coactivator associated arginine methyltr | 10498 | down | 1,36 | 1,60E-02 |
| R-HSA-381340 | Transcriptional regulation of white adipocyte different | TBL1X | transducin beta like 1 X-linked | 6907 | down | 1,20 | 2,84E-02 |
| R-HSA-381340 | Transcriptional regulation of white adipocyte different | RXRA | retinoid X receptor alpha | 6256 | down | 1,34 | 3,67E-03 |
| R-HSA-381340 | Transcriptional regulation of white adipocyte different | KLF4 | Kruppel like factor 4 | 9314 | up | 1,96 | 3,94E-02 |
| R-HSA-381340 | Transcriptional regulation of white adipocyte different | MED24 | mediator complex subunit 24 | 9862 | down | 1,33 | 2,93E-02 |
| R-HSA-381340 | Transcriptional regulation of white adipocyte different | NCOA3 | nuclear receptor coactivator 3 | 8202 | up | 1,53 | 4,13E-02 |
| R-HSA-381340 | Transcriptional regulation of white adipocyte different | WNT10B | Wnt family member 10B | 7480 | down | 1,76 | 4,78E-02 |
| R-HSA-381340 | Transcriptional regulation of white adipocyte different | CCND3 | cyclin D3 | 896 | down | 1,60 | 4,17E-02 |
| R-HSA-381340 | Transcriptional regulation of white adipocyte different | MED25 | mediator complex subunit 25 | 81857 | down | 1,23 | 4,79E-02 |
| R-HSA-427413 | NoRC negatively regulates rRNA expression | TAF1A | TATA-box binding protein associated fa | 9015 | up | 2,20 | 2,24E-03 |
| R-HSA-427413 | NoRC negatively regulates rRNA expression | CD3EAP | CD3e molecule associated protein | 10849 | down | 1,57 | 3,12E-03 |
| R-HSA-427413 | NoRC negatively regulates rRNA expression | POLR1A | RNA polymerase I subunit A | 25885 | down | 1,30 | 4,32E-02 |
| R-HSA-427413 | NoRC negatively regulates rRNA expression | POLR2L | RNA polymerase II subunit L | 5441 | down | 1,52 | 3,58E-03 |
| R-HSA-427413 | NoRC negatively regulates rRNA expression | POLR2F | RNA polymerase II subunit F | 5435 | down | 1,41 | 3,22E-02 |
| R-HSA-427413 | NoRC negatively regulates rRNA expression | GTF2H1 | general transcription factor IIH subunit 1 | 2965 | up | 1,33 | 1,85E-02 |
| R-HSA-427413 | NoRC negatively regulates rRNA expression | TAF1D | TATA-box binding protein associated fa | 79101 | up | 1,87 | 1,00E-02 |
| R-HSA-427413 | NoRC negatively regulates rRNA expression | POLR2E | RNA polymerase II subunit E | 5434 | down | 1,31 | 3,18E-03 |

| | | | | | | | |
|---|---|---|---|---|---|---|---|
| R-HSA-427413 | NoRC negatively regulates rRNA expression | POLR1C | RNA polymerase I subunit C | 9533 | down | 1,24 | 4,71E-02 |
| R-HSA-427413 | NoRC negatively regulates rRNA expression | CDK7 | cyclin dependent kinase 7 | 1022 | up | 1,37 | 4,38E-02 |
| R-HSA-427413 | NoRC negatively regulates rRNA expression | UBTF | upstream binding transcription factor, RI | 7343 | down | 1,66 | 1,20E-02 |
| R-HSA-427413 | NoRC negatively regulates rRNA expression | H2AFX | H2A histone family member X | 3014 | down | 2,11 | 1,30E-03 |
| R-HSA-427413 | NoRC negatively regulates rRNA expression | DNMT3B | DNA methyltransferase 3 beta | 1789 | down | 1,70 | 3,62E-02 |
| R-HSA-427413 | NoRC negatively regulates rRNA expression | ERCC2 | ERCC excision repair 2, TFIIH core com | 2068 | down | 1,29 | 2,20E-02 |
| R-HSA-427413 | NoRC negatively regulates rRNA expression | ERCC3 | ERCC excision repair 3, TFIIH core com | 2071 | down | 1,20 | 2,37E-02 |
| R-HSA-1358803 | Downregulation of ERBB2:ERBB3 signaling | AKT2 | AKT serine/threonine kinase 2 | 208 | down | 1,28 | 6,33E-03 |
| R-HSA-1358803 | Downregulation of ERBB2:ERBB3 signaling | AKT1 | AKT serine/threonine kinase 1 | 207 | down | 1,37 | 1,42E-03 |
| R-HSA-1358803 | Downregulation of ERBB2:ERBB3 signaling | ERBB2 | erb-b2 receptor tyrosine kinase 2 | 2064 | down | 1,47 | 1,93E-02 |
| R-HSA-1358803 | Downregulation of ERBB2:ERBB3 signaling | USP8 | ubiquitin specific peptidase 8 | 9101 | up | 1,31 | 7,82E-03 |
| R-HSA-1358803 | Downregulation of ERBB2:ERBB3 signaling | RNF41 | ring finger protein 41 | 10193 | up | 1,59 | 2,87E-02 |
| R-HSA-112409 | RAF-independent MAPK1/3 activation | DUSP10 | dual specificity phosphatase 10 | 11221 | up | 2,34 | 2,20E-03 |
| R-HSA-112409 | RAF-independent MAPK1/3 activation | DUSP8 | dual specificity phosphatase 8 | 1850 | up | 2,42 | 1,40E-02 |
| R-HSA-112409 | RAF-independent MAPK1/3 activation | DUSP5 | dual specificity phosphatase 5 | 1847 | up | 3,63 | 9,77E-03 |
| R-HSA-112409 | RAF-independent MAPK1/3 activation | DUSP1 | dual specificity phosphatase 1 | 1843 | up | 1,99 | 2,51E-02 |
| R-HSA-112409 | RAF-independent MAPK1/3 activation | DUSP7 | dual specificity phosphatase 7 | 1849 | down | 1,42 | 1,78E-02 |
| R-HSA-69231 | Cyclin D associated events in G1 | CDKN1A | cyclin dependent kinase inhibitor 1A | 1026 | up | 3,62 | 5,43E-03 |
| R-HSA-69231 | Cyclin D associated events in G1 | CDK7 | cyclin dependent kinase 7 | 1022 | up | 1,37 | 4,38E-02 |
| R-HSA-69231 | Cyclin D associated events in G1 | TFDP1 | transcription factor Dp-1 | 7027 | down | 1,77 | 1,08E-02 |
| R-HSA-69231 | Cyclin D associated events in G1 | PPP2R2A | protein phosphatase 2 regulatory subur | 5520 | up | 1,26 | 3,61E-02 |
| R-HSA-69231 | Cyclin D associated events in G1 | PPP2CB | protein phosphatase 2 catalytic subunit | 5516 | up | 1,30 | 4,79E-02 |
| R-HSA-69231 | Cyclin D associated events in G1 | CDKN2B | cyclin dependent kinase inhibitor 2B | 1030 | up | 2,40 | 3,90E-02 |
| R-HSA-69231 | Cyclin D associated events in G1 | E2F1 | E2F transcription factor 1 | 1869 | down | 1,64 | 3,25E-02 |
| R-HSA-69231 | Cyclin D associated events in G1 | SKP2 | S-phase kinase associated protein 2 | 6502 | down | 2,21 | 3,69E-02 |
| R-HSA-69231 | Cyclin D associated events in G1 | CCND3 | cyclin D3 | 896 | down | 1,60 | 4,17E-02 |
| R-HSA-1855204 | Synthesis of IP3 and IP4 in the cytosol | CALM1 | calmodulin 1 | 801 | down | 1,43 | 2,23E-02 |
| R-HSA-1855204 | Synthesis of IP3 and IP4 in the cytosol | PLCG1 | phospholipase C gamma 1 | 5335 | down | 1,51 | 2,85E-02 |
| R-HSA-1855204 | Synthesis of IP3 and IP4 in the cytosol | CALM3 | calmodulin 3 | 808 | down | 1,43 | 2,23E-02 |
| R-HSA-1855204 | Synthesis of IP3 and IP4 in the cytosol | ITPKC | inositol-trisphosphate 3-kinase C | 80271 | up | 1,63 | 3,32E-02 |
| R-HSA-1855204 | Synthesis of IP3 and IP4 in the cytosol | ITPKB | inositol-trisphosphate 3-kinase B | 3707 | down | 1,71 | 1,97E-02 |
| R-HSA-1855204 | Synthesis of IP3 and IP4 in the cytosol | CALM2 | calmodulin 2 | 805 | down | 1,43 | 2,23E-02 |
| R-HSA-1855204 | Synthesis of IP3 and IP4 in the cytosol | PTEN | phosphatase and tensin homolog | 5728 | up | 1,33 | 4,72E-02 |
| R-HSA-69300 | No details found for 69300 | MCM2 | minichromosome maintenance complex | 4171 | down | 2,01 | 1,42E-02 |
| R-HSA-69300 | No details found for 69300 | MCM7 | minichromosome maintenance complex | 4176 | down | 1,67 | 3,35E-02 |
| R-HSA-69300 | No details found for 69300 | MCM5 | minichromosome maintenance complex | 4174 | down | 1,83 | 2,22E-02 |
| R-HSA-69300 | No details found for 69300 | MCM3 | minichromosome maintenance complex | 4172 | down | 2,07 | 4,13E-02 |
| R-HSA-199418 | Negative regulation of the PI3K/AKT network | AKT1 | AKT serine/threonine kinase 1 | 207 | down | 1,37 | 1,42E-03 |
| R-HSA-199418 | Negative regulation of the PI3K/AKT network | AKT2 | AKT serine/threonine kinase 2 | 208 | down | 1,28 | 6,33E-03 |
| R-HSA-199418 | Negative regulation of the PI3K/AKT network | TRIB3 | tribbles pseudokinase 3 | 57761 | up | 3,17 | 8,07E-03 |
| R-HSA-199418 | Negative regulation of the PI3K/AKT network | PTEN | phosphatase and tensin homolog | 5728 | up | 1,33 | 4,72E-02 |
| R-HSA-139915 | Activation of PUMA and translocation to mitochondria | TP73 | tumor protein p73 | 7161 | down | 2,86 | 3,40E-02 |
| R-HSA-139915 | Activation of PUMA and translocation to mitochondria | TFDP1 | transcription factor Dp-1 | 7027 | down | 1,77 | 1,08E-02 |
| R-HSA-139915 | Activation of PUMA and translocation to mitochondria | BBC3 | BCL2 binding component 3 | 27113 | up | 2,09 | 3,17E-02 |
| R-HSA-139915 | Activation of PUMA and translocation to mitochondria | E2F1 | E2F transcription factor 1 | 1869 | down | 1,64 | 3,25E-02 |
| R-HSA-6781823 | Formation of TC-NER Pre-Incision Complex | ERCC3 | ERCC excision repair 3, TFIIH core com | 2071 | down | 1,20 | 2,37E-02 |
| R-HSA-6781823 | Formation of TC-NER Pre-Incision Complex | ERCC2 | ERCC excision repair 2, TFIIH core com | 2068 | down | 1,29 | 2,20E-02 |
| R-HSA-6781823 | Formation of TC-NER Pre-Incision Complex | CDK7 | cyclin dependent kinase 7 | 1022 | up | 1,37 | 4,38E-02 |
| R-HSA-6781823 | Formation of TC-NER Pre-Incision Complex | POLR2E | RNA polymerase II subunit E | 5434 | down | 1,31 | 3,18E-03 |
| R-HSA-6781823 | Formation of TC-NER Pre-Incision Complex | PRPF19 | pre-mRNA processing factor 19 | 27339 | down | 1,24 | 4,41E-02 |
| R-HSA-6781823 | Formation of TC-NER Pre-Incision Complex | POLR2B | RNA polymerase II subunit B | 5431 | down | 1,23 | 3,81E-02 |
| R-HSA-6781823 | Formation of TC-NER Pre-Incision Complex | TCEA1 | transcription elongation factor A1 | 6917 | up | 1,76 | 1,81E-02 |
| R-HSA-6781823 | Formation of TC-NER Pre-Incision Complex | GTF2H1 | general transcription factor IIH subunit 1 | 2965 | up | 1,33 | 1,85E-02 |
| R-HSA-6781823 | Formation of TC-NER Pre-Incision Complex | POLR2F | RNA polymerase II subunit F | 5435 | down | 1,41 | 3,22E-02 |
| R-HSA-6781823 | Formation of TC-NER Pre-Incision Complex | POLR2L | RNA polymerase II subunit L | 5441 | down | 1,52 | 3,58E-03 |
| R-HSA-1912408 | Pre-NOTCH Transcription and Translation | E2F1 | E2F transcription factor 1 | 1869 | down | 1,64 | 3,25E-02 |
| R-HSA-1912408 | Pre-NOTCH Transcription and Translation | NOTCH1 | notch 1 | 4851 | down | 1,62 | 2,46E-02 |
| R-HSA-1912408 | Pre-NOTCH Transcription and Translation | KAT2A | lysine acetyltransferase 2A | 2648 | down | 1,39 | 1,20E-02 |
| R-HSA-1912408 | Pre-NOTCH Transcription and Translation | MAMLD1 | mastermind like domain containing 1 | 10046 | down | 3,36 | 4,56E-02 |
| R-HSA-1912408 | Pre-NOTCH Transcription and Translation | TFDP1 | transcription factor Dp-1 | 7027 | down | 1,77 | 1,08E-02 |
| R-HSA-1912408 | Pre-NOTCH Transcription and Translation | AGO3 | argonaute 3, RISC catalytic component | 192669 | up | 1,36 | 1,80E-02 |
| R-HSA-5601884 | PIWI-interacting RNA (piRNA) biogenesis | POLR2F | RNA polymerase II subunit F | 5435 | down | 1,41 | 3,22E-02 |
| R-HSA-5601884 | PIWI-interacting RNA (piRNA) biogenesis | HENMT1 | HEN methyltransferase 1 | 113802 | down | 1,43 | 3,76E-02 |
| R-HSA-5601884 | PIWI-interacting RNA (piRNA) biogenesis | POLR2L | RNA polymerase II subunit L | 5441 | down | 1,52 | 3,58E-03 |
| R-HSA-5601884 | PIWI-interacting RNA (piRNA) biogenesis | POLR2B | RNA polymerase II subunit B | 5431 | down | 1,23 | 3,81E-02 |
| R-HSA-5601884 | PIWI-interacting RNA (piRNA) biogenesis | POLR2E | RNA polymerase II subunit E | 5434 | down | 1,31 | 3,18E-03 |
| R-HSA-163765 | ChREBP activates metabolic gene expression | ACACA | acetyl-CoA carboxylase alpha | 31 | down | 1,40 | 2,54E-02 |
| R-HSA-163765 | ChREBP activates metabolic gene expression | ACLY | ATP citrate lyase | 47 | down | 1,41 | 3,09E-02 |
| R-HSA-163765 | ChREBP activates metabolic gene expression | FASN | fatty acid synthase | 2194 | down | 1,96 | 8,31E-03 |
| R-HSA-2173788 | Downregulation of TGF-beta receptor signaling | TGFBR2 | transforming growth factor beta recepto | 7048 | down | 1,34 | 2,74E-02 |
| R-HSA-2173788 | Downregulation of TGF-beta receptor signaling | STUB1 | STIP1 homology and U-box containing | 10273 | down | 1,24 | 4,82E-03 |
| R-HSA-2173788 | Downregulation of TGF-beta receptor signaling | PPP1R15A | protein phosphatase 1 regulatory subur | 23645 | up | 4,33 | 1,06E-03 |
| R-HSA-2173788 | Downregulation of TGF-beta receptor signaling | SMAD2 | SMAD family member 2 | 4087 | up | 1,29 | 2,19E-02 |
| R-HSA-2173788 | Downregulation of TGF-beta receptor signaling | PPP1CA | protein phosphatase 1 catalytic subunit | 5499 | down | 1,24 | 9,77E-03 |
| R-HSA-2173788 | Downregulation of TGF-beta receptor signaling | SMURF2 | SMAD specific E3 ubiquitin protein ligas | 64750 | up | 1,49 | 4,16E-02 |
| R-HSA-2173788 | Downregulation of TGF-beta receptor signaling | USP15 | ubiquitin specific peptidase 15 | 9958 | up | 1,30 | 2,19E-02 |
| R-HSA-204174 | Regulation of pyruvate dehydrogenase (PDH) comple | PDK1 | pyruvate dehydrogenase kinase 1 | 5163 | up | 1,31 | 2,69E-03 |
| R-HSA-204174 | Regulation of pyruvate dehydrogenase (PDH) comple | PDK2 | pyruvate dehydrogenase kinase 2 | 5164 | down | 1,42 | 1,03E-02 |
| R-HSA-204174 | Regulation of pyruvate dehydrogenase (PDH) comple | DLD | dihydrolipoamide dehydrogenase | 1738 | up | 1,16 | 4,53E-02 |
| R-HSA-204174 | Regulation of pyruvate dehydrogenase (PDH) comple | PDP1 | pyruvate dehyrogenase phosphatase c | 54704 | up | 1,89 | 4,12E-03 |
| R-HSA-204174 | Regulation of pyruvate dehydrogenase (PDH) comple | RXRA | retinoid X receptor alpha | 6256 | down | 1,34 | 3,67E-03 |
| R-HSA-3769402 | Deactivation of the beta-catenin transactivating comp | TCF7 | transcription factor 7 | 6932 | down | 1,47 | 1,07E-02 |
| R-HSA-3769402 | Deactivation of the beta-catenin transactivating comp | CTBP1 | C-terminal binding protein 1 | 1487 | down | 1,16 | 3,77E-02 |
| R-HSA-3769402 | Deactivation of the beta-catenin transactivating comp | AKT1 | AKT serine/threonine kinase 1 | 207 | down | 1,37 | 1,42E-03 |
| R-HSA-3769402 | Deactivation of the beta-catenin transactivating comp | PYGO2 | pygopus family PHD finger 2 | 90780 | down | 1,37 | 1,36E-02 |
| R-HSA-3769402 | Deactivation of the beta-catenin transactivating comp | MEN1 | menin 1 | 4221 | down | 1,49 | 4,75E-04 |
| R-HSA-3769402 | Deactivation of the beta-catenin transactivating comp | AKT2 | AKT serine/threonine kinase 2 | 208 | down | 1,28 | 6,33E-03 |
| R-HSA-167246 | Tat-mediated elongation of the HIV-1 transcript | POLR2B | RNA polymerase II subunit B | 5431 | down | 1,23 | 3,81E-02 |
| R-HSA-167246 | Tat-mediated elongation of the HIV-1 transcript | POLR2L | RNA polymerase II subunit L | 5441 | down | 1,52 | 3,58E-03 |
| R-HSA-167246 | Tat-mediated elongation of the HIV-1 transcript | POLR2F | RNA polymerase II subunit F | 5435 | down | 1,41 | 3,22E-02 |
| R-HSA-167246 | Tat-mediated elongation of the HIV-1 transcript | TCEA1 | transcription elongation factor A1 | 6917 | up | 1,76 | 1,81E-02 |
| R-HSA-167246 | Tat-mediated elongation of the HIV-1 transcript | GTF2H1 | general transcription factor IIH subunit 1 | 2965 | up | 1,33 | 1,85E-02 |
| R-HSA-167246 | Tat-mediated elongation of the HIV-1 transcript | POLR2E | RNA polymerase II subunit E | 5434 | down | 1,31 | 3,18E-03 |
| R-HSA-167246 | Tat-mediated elongation of the HIV-1 transcript | CCNT1 | cyclin T1 | 904 | up | 1,34 | 3,05E-02 |
| R-HSA-167246 | Tat-mediated elongation of the HIV-1 transcript | CDK7 | cyclin dependent kinase 7 | 1022 | up | 1,37 | 4,38E-02 |
| R-HSA-167246 | Tat-mediated elongation of the HIV-1 transcript | ERCC2 | ERCC excision repair 2, TFIIH core com | 2068 | down | 1,29 | 2,20E-02 |
| R-HSA-167246 | Tat-mediated elongation of the HIV-1 transcript | ERCC3 | ERCC excision repair 3, TFIIH core com | 2071 | down | 1,20 | 2,37E-02 |
| R-HSA-110314 | Recognition of DNA damage by PCNA-containing rep | POLE | DNA polymerase epsilon, catalytic subu | 5426 | down | 1,55 | 2,67E-02 |
| R-HSA-110314 | Recognition of DNA damage by PCNA-containing rep | UBE2B | ubiquitin conjugating enzyme E2 B | 7320 | up | 1,65 | 4,38E-02 |
| R-HSA-110314 | Recognition of DNA damage by PCNA-containing rep | POLD1 | DNA polymerase delta 1, catalytic subur | 5424 | down | 1,59 | 4,62E-02 |
| R-HSA-110314 | Recognition of DNA damage by PCNA-containing rep | POLD2 | DNA polymerase delta 2, accessory sub | 5425 | down | 1,30 | 1,92E-02 |
| R-HSA-110314 | Recognition of DNA damage by PCNA-containing rep | PCNA | proliferating cell nuclear antigen | 5111 | down | 2,14 | 4,63E-02 |
| R-HSA-110314 | Recognition of DNA damage by PCNA-containing rep | RFC1 | replication factor C subunit 1 | 5981 | up | 1,15 | 2,18E-02 |
| R-HSA-110314 | Recognition of DNA damage by PCNA-containing rep | RPA1 | replication protein A1 | 6117 | down | 1,41 | 4,93E-02 |
| R-HSA-75955 | RNA Polymerase II Transcription Elongation | POLR2L | RNA polymerase II subunit L | 5441 | down | 1,52 | 3,58E-03 |
| R-HSA-75955 | RNA Polymerase II Transcription Elongation | GTF2H1 | general transcription factor IIH subunit 1 | 2965 | up | 1,33 | 1,85E-02 |
| R-HSA-75955 | RNA Polymerase II Transcription Elongation | POLR2F | RNA polymerase II subunit F | 5435 | down | 1,41 | 3,22E-02 |
| R-HSA-75955 | RNA Polymerase II Transcription Elongation | TCEA1 | transcription elongation factor A1 | 6917 | up | 1,76 | 1,81E-02 |
| R-HSA-75955 | RNA Polymerase II Transcription Elongation | POLR2B | RNA polymerase II subunit B | 5431 | down | 1,23 | 3,81E-02 |
| R-HSA-75955 | RNA Polymerase II Transcription Elongation | CCNT1 | cyclin T1 | 904 | up | 1,34 | 3,05E-02 |
| R-HSA-75955 | RNA Polymerase II Transcription Elongation | POLR2E | RNA polymerase II subunit E | 5434 | down | 1,31 | 3,18E-03 |
| R-HSA-75955 | RNA Polymerase II Transcription Elongation | CDK7 | cyclin dependent kinase 7 | 1022 | up | 1,37 | 4,38E-02 |
| R-HSA-75955 | RNA Polymerase II Transcription Elongation | ERCC2 | ERCC excision repair 2, TFIIH core com | 2068 | down | 1,29 | 2,20E-02 |
| R-HSA-75955 | RNA Polymerase II Transcription Elongation | ERCC3 | ERCC excision repair 3, TFIIH core com | 2071 | down | 1,20 | 2,37E-02 |
| R-HSA-75955 | RNA Polymerase II Transcription Elongation | CDC73 | cell division cycle 73 | 79577 | up | 1,37 | 4,41E-02 |
| R-HSA-111448 | Activation of NOXA and translocation to mitochondria | PMAIP1 | phorbol-12-myristate-13-acetate-induced | 5366 | up | 2,01 | 4,52E-02 |
| R-HSA-111448 | Activation of NOXA and translocation to mitochondria | E2F1 | E2F transcription factor 1 | 1869 | down | 1,64 | 3,25E-02 |
| R-HSA-111448 | Activation of NOXA and translocation to mitochondria | TFDP1 | transcription factor Dp-1 | 7027 | down | 1,77 | 1,08E-02 |
| R-HSA-727802 | Transport of nucleotide sugars | SLC35B2 | solute carrier family 35 member B2 | 347734 | down | 1,57 | 6,39E-03 |
| R-HSA-727802 | Transport of nucleotide sugars | SLC35C1 | solute carrier family 35 member C1 | 55343 | down | 1,25 | 2,93E-02 |
| R-HSA-727802 | Transport of nucleotide sugars | SLC35B4 | solute carrier family 35 member B4 | 84912 | down | 1,57 | 3,98E-02 |
| R-HSA-198693 | AKT phosphorylates targets in the nucleus | AKT2 | AKT serine/threonine kinase 2 | 208 | down | 1,28 | 6,33E-03 |
| R-HSA-198693 | AKT phosphorylates targets in the nucleus | RPS6KB2 | ribosomal protein S6 kinase B2 | 6199 | down | 1,22 | 1,01E-02 |
| R-HSA-198693 | AKT phosphorylates targets in the nucleus | AKT1 | AKT serine/threonine kinase 1 | 207 | down | 1,37 | 1,42E-03 |
| R-HSA-198693 | AKT phosphorylates targets in the nucleus | CREB1 | cAMP responsive element binding prote | 1385 | up | 1,51 | 7,18E-03 |
| R-HSA-4641262 | Disassembly of the destruction complex and recruitme | PPP2CB | protein phosphatase 2 catalytic subunit | 5516 | up | 1,30 | 4,79E-02 |
| R-HSA-4641262 | Disassembly of the destruction complex and recruitme | AMER1 | APC membrane recruitment protein 1 | 139285 | down | 1,78 | 4,15E-02 |
| R-HSA-4641262 | Disassembly of the destruction complex and recruitme | FZD1 | frizzled class receptor 1 | 8321 | down | 1,75 | 2,84E-02 |
| R-HSA-4641262 | Disassembly of the destruction complex and recruitme | CSNK1A1 | casein kinase 1 alpha 1 | 1452 | up | 1,35 | 2,84E-02 |
| R-HSA-4641262 | Disassembly of the destruction complex and recruitme | CSNK1G2 | casein kinase 1 gamma 2 | 1455 | down | 1,34 | 2,24E-03 |
| R-HSA-4641262 | Disassembly of the destruction complex and recruitme | FRAT1 | FRAT1, WNT signaling pathway regulato | 10023 | down | 1,48 | 3,14E-02 |
| R-HSA-4641262 | Disassembly of the destruction complex and recruitme | FRAT2 | FRAT2, WNT signaling pathway regulato | 23401 | down | 1,71 | 2,52E-04 |
| R-HSA-198203 | PI3K/AKT activation | PIK3R1 | phosphoinositide-3-kinase regulatory su | 5295 | down | 2,04 | 8,40E-03 |
| R-HSA-198203 | PI3K/AKT activation | PIK3CA | phosphatidylinositol-4,5-bisphosphate 3 | 5290 | up | 1,33 | 3,17E-02 |
| R-HSA-198203 | PI3K/AKT activation | IRS1 | insulin receptor substrate 1 | 3667 | down | 1,89 | 6,25E-03 |
| R-HSA-198203 | PI3K/AKT activation | IRS2 | insulin receptor substrate 2 | 8660 | up | 2,21 | 2,22E-02 |
| R-HSA-6784531 | tRNA processing in the nucleus | POP1 | POP1 homolog, ribonuclease P/MRP su | 10940 | down | 1,65 | 3,98E-02 |
| R-HSA-6784531 | tRNA processing in the nucleus | XPOT | exportin for tRNA | 11260 | up | 1,58 | 1,20E-02 |
| R-HSA-6784531 | tRNA processing in the nucleus | CLP1 | cleavage and polyadenylation factor I s | 10978 | down | 1,40 | 1,04E-02 |
| R-HSA-6784531 | tRNA processing in the nucleus | NUP58 | nucleoporin 58 | 9818 | up | 1,61 | 1,28E-02 |
| R-HSA-6784531 | tRNA processing in the nucleus | CSTF2 | cleavage stimulation factor subunit 2 | 1478 | down | 1,45 | 4,38E-02 |
| R-HSA-6784531 | tRNA processing in the nucleus | NUP62 | nucleoporin 62 | 23636 | down | 1,52 | 4,39E-03 |
| R-HSA-6784531 | tRNA processing in the nucleus | TSEN54 | tRNA splicing endonuclease subunit 54 | 283989 | down | 1,59 | 9,51E-03 |
| R-HSA-6784531 | tRNA processing in the nucleus | RPP25 | ribonuclease P and MRP subunit p25 | 54913 | down | 1,71 | 4,35E-03 |
| R-HSA-6784531 | tRNA processing in the nucleus | ELAC2 | elaC ribonuclease Z 2 | 60528 | down | 1,30 | 2,77E-02 |
| R-HSA-167200 | Formation of HIV-1 elongation complex containing HI | ERCC3 | ERCC excision repair 3, TFIIH core com | 2071 | down | 1,20 | 2,37E-02 |
| R-HSA-167200 | Formation of HIV-1 elongation complex containing HI | ERCC2 | ERCC excision repair 2, TFIIH core com | 2068 | down | 1,29 | 2,20E-02 |
| R-HSA-167200 | Formation of HIV-1 elongation complex containing HI | CDK7 | cyclin dependent kinase 7 | 1022 | up | 1,37 | 4,38E-02 |
| R-HSA-167200 | Formation of HIV-1 elongation complex containing HI | CCNT1 | cyclin T1 | 904 | up | 1,34 | 3,05E-02 |
| R-HSA-167200 | Formation of HIV-1 elongation complex containing HI | POLR2E | RNA polymerase II subunit E | 5434 | down | 1,31 | 3,18E-03 |
| R-HSA-167200 | Formation of HIV-1 elongation complex containing HI | POLR2L | RNA polymerase II subunit L | 5441 | down | 1,52 | 3,58E-03 |
| R-HSA-167200 | Formation of HIV-1 elongation complex containing HI | POLR2F | RNA polymerase II subunit F | 5435 | down | 1,41 | 3,22E-02 |
| R-HSA-167200 | Formation of HIV-1 elongation complex containing HI | GTF2H1 | general transcription factor IIH subunit 1 | 2965 | up | 1,33 | 1,85E-02 |
| R-HSA-167200 | Formation of HIV-1 elongation complex containing HI | TCEA1 | transcription elongation factor A1 | 6917 | up | 1,76 | 1,81E-02 |
| R-HSA-167200 | Formation of HIV-1 elongation complex containing HI | POLR2B | RNA polymerase II subunit B | 5431 | down | 1,23 | 3,81E-02 |
| R-HSA-936440 | Negative regulators of DDX58/IFIH1 signaling | TRAF3 | TNF receptor associated factor 3 | 7187 | down | 1,40 | 1,36E-02 |
| R-HSA-936440 | Negative regulators of DDX58/IFIH1 signaling | CYLD | CYLD lysine 63 deubiquitinase | 1540 | up | 1,34 | 4,86E-02 |
| R-HSA-936440 | Negative regulators of DDX58/IFIH1 signaling | ATG12 | autophagy related 12 | 9140 | up | 1,35 | 4,10E-02 |
| R-HSA-936440 | Negative regulators of DDX58/IFIH1 signaling | IKBKE | inhibitor of nuclear factor kappa B kinas | 9641 | down | 1,67 | 1,00E-02 |
| R-HSA-936440 | Negative regulators of DDX58/IFIH1 signaling | IRF3 | interferon regulatory factor 3 | 3661 | down | 1,33 | 2,58E-03 |
| R-HSA-936440 | Negative regulators of DDX58/IFIH1 signaling | ITCH | itchy E3 ubiquitin protein ligase | 83737 | up | 1,51 | 6,64E-03 |
| R-HSA-936440 | Negative regulators of DDX58/IFIH1 signaling | NLRX1 | NLR family member X1 | 79671 | down | 1,38 | 1,23E-02 |
| R-HSA-936440 | Negative regulators of DDX58/IFIH1 signaling | TAX1BP1 | Tax1 binding protein 1 | 8887 | up | 1,45 | 1,32E-03 |
| R-HSA-1655829 | Regulation of cholesterol biosynthesis by SREBP (SR | SEC23A | Sec23 homolog A, coat complex II com | 10484 | up | 1,46 | 2,47E-02 |
| R-HSA-1655829 | Regulation of cholesterol biosynthesis by SREBP (SR | SEC24D | SEC24 homolog D, COPII coat complex | 9871 | up | 4,45 | 6,80E-04 |
| R-HSA-1655829 | Regulation of cholesterol biosynthesis by SREBP (SR | INSIG1 | insulin induced gene 1 | 3638 | down | 1,94 | 1,60E-02 |
| R-HSA-1655829 | Regulation of cholesterol biosynthesis by SREBP (SR | SEC24A | SEC24 homolog A, COPII coat complex | 10802 | up | 1,71 | 3,95E-03 |
| R-HSA-1655829 | Regulation of cholesterol biosynthesis by SREBP (SR | SREBF1 | sterol regulatory element binding transc | 6720 | down | 1,42 | 3,67E-02 |
| R-HSA-5656169 | Termination of translesion DNA synthesis | POLE | DNA polymerase epsilon, catalytic subu | 5426 | down | 1,55 | 2,67E-02 |
| R-HSA-5656169 | Termination of translesion DNA synthesis | POLD1 | DNA polymerase delta 1, catalytic subur | 5424 | down | 1,59 | 4,62E-02 |
| R-HSA-5656169 | Termination of translesion DNA synthesis | RFC1 | replication factor C subunit 1 | 5981 | up | 1,15 | 2,18E-02 |
| R-HSA-5656169 | Termination of translesion DNA synthesis | PCNA | proliferating cell nuclear antigen | 5111 | down | 2,14 | 4,63E-02 |
| R-HSA-5656169 | Termination of translesion DNA synthesis | POLD2 | DNA polymerase delta 2, accessory sub | 5425 | down | 1,30 | 1,92E-02 |
| R-HSA-5656169 | Termination of translesion DNA synthesis | RPA1 | replication protein A1 | 6117 | down | 1,41 | 4,93E-02 |
| R-HSA-70263 | Gluconeogenesis | PGAM1 | phosphoglycerate mutase 1 | 5223 | down | 1,62 | 5,67E-03 |
| R-HSA-70263 | Gluconeogenesis | GOT1 | glutamic-oxaloacetic transaminase 1 | 2805 | up | 2,30 | 5,67E-03 |
| R-HSA-70263 | Gluconeogenesis | TPI1 | triosephosphate isomerase 1 | 7167 | down | 1,28 | 3,58E-02 |
| R-HSA-70263 | Gluconeogenesis | SLC25A1 | solute carrier family 25 member 1 | 6576 | down | 1,48 | 5,54E-04 |
| R-HSA-70263 | Gluconeogenesis | GPI | glucose-6-phosphate isomerase | 2821 | down | 1,24 | 1,55E-02 |
| R-HSA-70263 | Gluconeogenesis | SLC25A10 | solute carrier family 25 member 10 | 1468 | down | 1,83 | 5,12E-03 |
| R-HSA-112382 | Formation of RNA Pol II elongation complex | ERCC3 | ERCC excision repair 3, TFIIH core com | 2071 | down | 1,20 | 2,37E-02 |
| R-HSA-112382 | Formation of RNA Pol II elongation complex | ERCC2 | ERCC excision repair 2, TFIIH core com | 2068 | down | 1,29 | 2,20E-02 |
| R-HSA-112382 | Formation of RNA Pol II elongation complex | CDC73 | cell division cycle 73 | 79577 | up | 1,37 | 4,41E-02 |
| R-HSA-112382 | Formation of RNA Pol II elongation complex | CDK7 | cyclin dependent kinase 7 | 1022 | up | 1,37 | 4,38E-02 |
| R-HSA-112382 | Formation of RNA Pol II elongation complex | CCNT1 | cyclin T1 | 904 | up | 1,34 | 3,05E-02 |
| R-HSA-112382 | Formation of RNA Pol II elongation complex | POLR2E | RNA polymerase II subunit E | 5434 | down | 1,31 | 3,18E-03 |
| R-HSA-112382 | Formation of RNA Pol II elongation complex | POLR2L | RNA polymerase II subunit L | 5441 | down | 1,52 | 3,58E-03 |
| R-HSA-112382 | Formation of RNA Pol II elongation complex | TCEA1 | transcription elongation factor A1 | 6917 | up | 1,76 | 1,81E-02 |
| R-HSA-112382 | Formation of RNA Pol II elongation complex | POLR2F | RNA polymerase II subunit F | 5435 | down | 1,41 | 3,22E-02 |
| R-HSA-112382 | Formation of RNA Pol II elongation complex | GTF2H1 | general transcription factor IIH subunit 1 | 2965 | up | 1,33 | 1,85E-02 |
| R-HSA-112382 | Formation of RNA Pol II elongation complex | POLR2B | RNA polymerase II subunit B | 5431 | down | 1,23 | 3,81E-02 |
| R-HSA-1660499 | Synthesis of PIPs at the plasma membrane | PIP5K1C | phosphatidylinositol-4-phosphate 5-kina | 23396 | down | 1,39 | 1,49E-03 |
| R-HSA-1660499 | Synthesis of PIPs at the plasma membrane | PIK3R1 | phosphoinositide-3-kinase regulatory su | 5295 | down | 2,04 | 8,40E-03 |
| R-HSA-1660499 | Synthesis of PIPs at the plasma membrane | MTMR6 | myotubularin related protein 6 | 9107 | up | 1,33 | 2,92E-02 |
| R-HSA-1660499 | Synthesis of PIPs at the plasma membrane | PIK3CA | phosphatidylinositol-4,5-bisphosphate 3 | 5290 | up | 1,33 | 3,17E-02 |
| R-HSA-1660499 | Synthesis of PIPs at the plasma membrane | PIP5K1B | phosphatidylinositol-4-phosphate 5-kina | 8395 | up | 3,40 | 1,84E-02 |
| R-HSA-1660499 | Synthesis of PIPs at the plasma membrane | PIK3C2A | phosphatidylinositol-4-phosphate 3-kina | 5286 | up | 1,30 | 1,79E-02 |
| R-HSA-1660499 | Synthesis of PIPs at the plasma membrane | PTEN | phosphatase and tensin homolog | 5728 | up | 1,33 | 4,72E-02 |
| R-HSA-1660499 | Synthesis of PIPs at the plasma membrane | PIP4K2B | phosphatidylinositol-5-phosphate 4-kina | 8396 | down | 1,53 | 1,01E-02 |
| R-HSA-389357 | CD28 dependent PI3K/Akt signaling | PIK3R1 | phosphoinositide-3-kinase regulatory su | 5295 | down | 2,04 | 8,40E-03 |
| R-HSA-389357 | CD28 dependent PI3K/Akt signaling | AKT2 | AKT serine/threonine kinase 2 | 208 | down | 1,28 | 6,33E-03 |
| R-HSA-389357 | CD28 dependent PI3K/Akt signaling | AKT1 | AKT serine/threonine kinase 1 | 207 | down | 1,37 | 1,42E-03 |
| R-HSA-389357 | CD28 dependent PI3K/Akt signaling | PIK3CA | phosphatidylinositol-4,5-bisphosphate 3 | 5290 | up | 1,33 | 3,17E-02 |
| R-HSA-389357 | CD28 dependent PI3K/Akt signaling | TRIB3 | tribbles pseudokinase 3 | 57761 | up | 3,17 | 8,07E-03 |
| R-HSA-389357 | CD28 dependent PI3K/Akt signaling | PRR5 | proline rich 5 | 55615 | down | 1,41 | 1,79E-03 |
| R-HSA-2032785 | YAP1- and WWTR1 (TAZ)-stimulated gene expressio | CTGF | connective tissue growth factor | 1490 | up | 2,49 | 1,32E-02 |
| R-HSA-2032785 | YAP1- and WWTR1 (TAZ)-stimulated gene expressio | TGS1 | trimethylguanosine synthase 1 | 96764 | up | 1,63 | 9,37E-03 |
| R-HSA-2032785 | YAP1- and WWTR1 (TAZ)-stimulated gene expressio | WWTR1 | WW domain containing transcription reg | 25937 | down | 1,99 | 1,00E-02 |
| R-HSA-2032785 | YAP1- and WWTR1 (TAZ)-stimulated gene expressio | RXRA | retinoid X receptor alpha | 6256 | down | 1,34 | 3,67E-03 |
| R-HSA-2032785 | YAP1- and WWTR1 (TAZ)-stimulated gene expressio | CARM1 | coactivator associated arginine methyltr | 10498 | down | 1,36 | 1,60E-02 |
| R-HSA-2032785 | YAP1- and WWTR1 (TAZ)-stimulated gene expressio | TBL1X | transducin beta like 1 X-linked | 6907 | down | 1,20 | 2,84E-02 |
| R-HSA-2032785 | YAP1- and WWTR1 (TAZ)-stimulated gene expressio | CHD9 | chromodomain helicase DNA binding pr | 80205 | up | 1,32 | 1,42E-03 |
| R-HSA-446652 | Interleukin-1 family signaling | PELI3 | pellino E3 ubiquitin protein ligase family | 246330 | down | 1,56 | 1,44E-02 |
| R-HSA-446652 | Interleukin-1 family signaling | IL1RAP | interleukin 1 receptor accessory protein | 3556 | up | 2,09 | 9,72E-03 |
| R-HSA-446652 | Interleukin-1 family signaling | MYD88 | myeloid differentiation primary response | 4615 | down | 1,55 | 1,85E-02 |
| R-HSA-446652 | Interleukin-1 family signaling | TAB3 | TGF-beta activated kinase 1 and MAP3 | 257397 | up | 1,40 | 4,94E-02 |
| R-HSA-446652 | Interleukin-1 family signaling | RIPK2 | receptor interacting serine/threonine kin | 8767 | up | 1,72 | 8,31E-03 |

| | | | | | | | |
|---|---|---|---|---|---|---|---|
| R-HSA-446652 | Interleukin-1 family signaling | IL1A | interleukin 1 alpha | 3552 | up | 8,56 | 1,05E-02 |
| R-HSA-446652 | Interleukin-1 family signaling | SQSTM1 | sequestosome 1 | 8878 | up | 1,88 | 2,42E-02 |
| R-HSA-446652 | Interleukin-1 family signaling | TAB1 | TGF-beta activated kinase 1 (MAP3K7) | 10454 | down | 1,29 | 1,94E-02 |
| R-HSA-167152 | Formation of HIV elongation complex in the absence | GTF2H1 | general transcription factor IIH subunit 1 | 2965 | up | 1,33 | 1,85E-02 |
| R-HSA-167152 | Formation of HIV elongation complex in the absence | POLR2F | RNA polymerase II subunit F | 5435 | down | 1,41 | 3,22E-02 |
| R-HSA-167152 | Formation of HIV elongation complex in the absence | TCEA1 | transcription elongation factor A1 | 6917 | up | 1,76 | 1,81E-02 |
| R-HSA-167152 | Formation of HIV elongation complex in the absence | POLR2L | RNA polymerase II subunit L | 5441 | down | 1,52 | 3,58E-03 |
| R-HSA-167152 | Formation of HIV elongation complex in the absence | POLR2B | RNA polymerase II subunit B | 5431 | down | 1,23 | 3,81E-02 |
| R-HSA-167152 | Formation of HIV elongation complex in the absence | CCNT1 | cyclin T1 | 904 | up | 1,34 | 3,05E-02 |
| R-HSA-167152 | Formation of HIV elongation complex in the absence | POLR2E | RNA polymerase II subunit E | 5434 | down | 1,31 | 3,18E-03 |
| R-HSA-167152 | Formation of HIV elongation complex in the absence | CDK7 | cyclin dependent kinase 7 | 1022 | up | 1,37 | 4,38E-02 |
| R-HSA-167152 | Formation of HIV elongation complex in the absence | ERCC2 | ERCC excision repair 2, TFIIH core com | 2068 | down | 1,29 | 2,20E-02 |
| R-HSA-167152 | Formation of HIV elongation complex in the absence | ERCC3 | ERCC excision repair 3, TFIIH core com | 2071 | down | 1,20 | 2,37E-02 |
| R-HSA-76042 | RNA Polymerase II Transcription Initiation And Promo | TAF2 | TATA-box binding protein associated fa | 6873 | up | 1,29 | 5,94E-03 |
| R-HSA-76042 | RNA Polymerase II Transcription Initiation And Promo | POLR2E | RNA polymerase II subunit E | 5434 | down | 1,31 | 3,18E-03 |
| R-HSA-76042 | RNA Polymerase II Transcription Initiation And Promo | POLR2L | RNA polymerase II subunit L | 5441 | down | 1,52 | 3,58E-03 |
| R-HSA-76042 | RNA Polymerase II Transcription Initiation And Promo | GTF2H1 | general transcription factor IIH subunit 1 | 2965 | up | 1,33 | 1,85E-02 |
| R-HSA-76042 | RNA Polymerase II Transcription Initiation And Promo | POLR2F | RNA polymerase II subunit F | 5435 | down | 1,41 | 3,22E-02 |
| R-HSA-76042 | RNA Polymerase II Transcription Initiation And Promo | POLR2B | RNA polymerase II subunit B | 5431 | down | 1,23 | 3,81E-02 |
| R-HSA-76042 | RNA Polymerase II Transcription Initiation And Promo | ERCC3 | ERCC excision repair 3, TFIIH core com | 2071 | down | 1,20 | 2,37E-02 |
| R-HSA-76042 | RNA Polymerase II Transcription Initiation And Promo | ERCC2 | ERCC excision repair 2, TFIIH core com | 2068 | down | 1,29 | 2,20E-02 |
| R-HSA-76042 | RNA Polymerase II Transcription Initiation And Promo | CDK7 | cyclin dependent kinase 7 | 1022 | up | 1,37 | 4,38E-02 |
| R-HSA-73776 | RNA Polymerase II Promoter Escape | CDK7 | cyclin dependent kinase 7 | 1022 | up | 1,37 | 4,38E-02 |
| R-HSA-73776 | RNA Polymerase II Promoter Escape | ERCC3 | ERCC excision repair 3, TFIIH core com | 2071 | down | 1,20 | 2,37E-02 |
| R-HSA-73776 | RNA Polymerase II Promoter Escape | ERCC2 | ERCC excision repair 2, TFIIH core com | 2068 | down | 1,29 | 2,20E-02 |
| R-HSA-73776 | RNA Polymerase II Promoter Escape | POLR2B | RNA polymerase II subunit B | 5431 | down | 1,23 | 3,81E-02 |
| R-HSA-73776 | RNA Polymerase II Promoter Escape | POLR2L | RNA polymerase II subunit L | 5441 | down | 1,52 | 3,58E-03 |
| R-HSA-73776 | RNA Polymerase II Promoter Escape | POLR2F | RNA polymerase II subunit F | 5435 | down | 1,41 | 3,22E-02 |
| R-HSA-73776 | RNA Polymerase II Promoter Escape | GTF2H1 | general transcription factor IIH subunit 1 | 2965 | up | 1,33 | 1,85E-02 |
| R-HSA-73776 | RNA Polymerase II Promoter Escape | POLR2E | RNA polymerase II subunit E | 5434 | down | 1,31 | 3,18E-03 |
| R-HSA-73776 | RNA Polymerase II Promoter Escape | TAF2 | TATA-box binding protein associated fa | 6873 | up | 1,29 | 5,94E-03 |
| R-HSA-73779 | RNA Polymerase II Transcription Pre-Initiation And Pr | ERCC3 | ERCC excision repair 3, TFIIH core com | 2071 | down | 1,20 | 2,37E-02 |
| R-HSA-73779 | RNA Polymerase II Transcription Pre-Initiation And Pr | ERCC2 | ERCC excision repair 2, TFIIH core com | 2068 | down | 1,29 | 2,20E-02 |
| R-HSA-73779 | RNA Polymerase II Transcription Pre-Initiation And Pr | CDK7 | cyclin dependent kinase 7 | 1022 | up | 1,37 | 4,38E-02 |
| R-HSA-73779 | RNA Polymerase II Transcription Pre-Initiation And Pr | POLR2E | RNA polymerase II subunit E | 5434 | down | 1,31 | 3,18E-03 |
| R-HSA-73779 | RNA Polymerase II Transcription Pre-Initiation And Pr | TAF2 | TATA-box binding protein associated fa | 6873 | up | 1,29 | 5,94E-03 |
| R-HSA-73779 | RNA Polymerase II Transcription Pre-Initiation And Pr | POLR2B | RNA polymerase II subunit B | 5431 | down | 1,23 | 3,81E-02 |
| R-HSA-73779 | RNA Polymerase II Transcription Pre-Initiation And Pr | GTF2H1 | general transcription factor IIH subunit 1 | 2965 | up | 1,33 | 1,85E-02 |
| R-HSA-73779 | RNA Polymerase II Transcription Pre-Initiation And Pr | POLR2F | RNA polymerase II subunit F | 5435 | down | 1,41 | 3,22E-02 |
| R-HSA-73779 | RNA Polymerase II Transcription Pre-Initiation And Pr | POLR2L | RNA polymerase II subunit L | 5441 | down | 1,52 | 3,58E-03 |
| R-HSA-167161 | HIV Transcription Initiation | ERCC3 | ERCC excision repair 3, TFIIH core com | 2071 | down | 1,20 | 2,37E-02 |
| R-HSA-167161 | HIV Transcription Initiation | ERCC2 | ERCC excision repair 2, TFIIH core com | 2068 | down | 1,29 | 2,20E-02 |
| R-HSA-167161 | HIV Transcription Initiation | CDK7 | cyclin dependent kinase 7 | 1022 | up | 1,37 | 4,38E-02 |
| R-HSA-167161 | HIV Transcription Initiation | POLR2E | RNA polymerase II subunit E | 5434 | down | 1,31 | 3,18E-03 |
| R-HSA-167161 | HIV Transcription Initiation | TAF2 | TATA-box binding protein associated fa | 6873 | up | 1,29 | 5,94E-03 |
| R-HSA-167161 | HIV Transcription Initiation | POLR2B | RNA polymerase II subunit B | 5431 | down | 1,23 | 3,81E-02 |
| R-HSA-167161 | HIV Transcription Initiation | GTF2H1 | general transcription factor IIH subunit 1 | 2965 | up | 1,33 | 1,85E-02 |
| R-HSA-167161 | HIV Transcription Initiation | POLR2F | RNA polymerase II subunit F | 5435 | down | 1,41 | 3,22E-02 |
| R-HSA-167161 | HIV Transcription Initiation | POLR2L | RNA polymerase II subunit L | 5441 | down | 1,52 | 3,58E-03 |
| R-HSA-167162 | RNA Polymerase II HIV Promoter Escape | POLR2B | RNA polymerase II subunit B | 5431 | down | 1,23 | 3,81E-02 |
| R-HSA-167162 | RNA Polymerase II HIV Promoter Escape | POLR2L | RNA polymerase II subunit L | 5441 | down | 1,52 | 3,58E-03 |
| R-HSA-167162 | RNA Polymerase II HIV Promoter Escape | GTF2H1 | general transcription factor IIH subunit 1 | 2965 | up | 1,33 | 1,85E-02 |
| R-HSA-167162 | RNA Polymerase II HIV Promoter Escape | POLR2F | RNA polymerase II subunit F | 5435 | down | 1,41 | 3,22E-02 |
| R-HSA-167162 | RNA Polymerase II HIV Promoter Escape | POLR2E | RNA polymerase II subunit E | 5434 | down | 1,31 | 3,18E-03 |
| R-HSA-167162 | RNA Polymerase II HIV Promoter Escape | TAF2 | TATA-box binding protein associated fa | 6873 | up | 1,29 | 5,94E-03 |
| R-HSA-167162 | RNA Polymerase II HIV Promoter Escape | CDK7 | cyclin dependent kinase 7 | 1022 | up | 1,37 | 4,38E-02 |
| R-HSA-167162 | RNA Polymerase II HIV Promoter Escape | ERCC2 | ERCC excision repair 2, TFIIH core com | 2068 | down | 1,29 | 2,20E-02 |
| R-HSA-167162 | RNA Polymerase II HIV Promoter Escape | ERCC3 | ERCC excision repair 3, TFIIH core com | 2071 | down | 1,20 | 2,37E-02 |
| R-HSA-167172 | Transcription of the HIV genome | POLR2B | RNA polymerase II subunit B | 5431 | down | 1,23 | 3,81E-02 |
| R-HSA-167172 | Transcription of the HIV genome | GTF2H1 | general transcription factor IIH subunit 1 | 2965 | up | 1,33 | 1,85E-02 |
| R-HSA-167172 | Transcription of the HIV genome | POLR2F | RNA polymerase II subunit F | 5435 | down | 1,41 | 3,22E-02 |
| R-HSA-167172 | Transcription of the HIV genome | POLR2L | RNA polymerase II subunit L | 5441 | down | 1,52 | 3,58E-03 |
| R-HSA-167172 | Transcription of the HIV genome | POLR2E | RNA polymerase II subunit E | 5434 | down | 1,31 | 3,18E-03 |
| R-HSA-167172 | Transcription of the HIV genome | TAF2 | TATA-box binding protein associated fa | 6873 | up | 1,29 | 5,94E-03 |
| R-HSA-167172 | Transcription of the HIV genome | CDK7 | cyclin dependent kinase 7 | 1022 | up | 1,37 | 4,38E-02 |
| R-HSA-167172 | Transcription of the HIV genome | ERCC3 | ERCC excision repair 3, TFIIH core com | 2071 | down | 1,20 | 2,37E-02 |
| R-HSA-167172 | Transcription of the HIV genome | ERCC2 | ERCC excision repair 2, TFIIH core com | 2068 | down | 1,29 | 2,20E-02 |
| R-HSA-75953 | RNA Polymerase II Transcription Initiation | ERCC3 | ERCC excision repair 3, TFIIH core com | 2071 | down | 1,20 | 2,37E-02 |
| R-HSA-75953 | RNA Polymerase II Transcription Initiation | ERCC2 | ERCC excision repair 2, TFIIH core com | 2068 | down | 1,29 | 2,20E-02 |
| R-HSA-75953 | RNA Polymerase II Transcription Initiation | CDK7 | cyclin dependent kinase 7 | 1022 | up | 1,37 | 4,38E-02 |
| R-HSA-75953 | RNA Polymerase II Transcription Initiation | TAF2 | TATA-box binding protein associated fa | 6873 | up | 1,29 | 5,94E-03 |
| R-HSA-75953 | RNA Polymerase II Transcription Initiation | POLR2E | RNA polymerase II subunit E | 5434 | down | 1,31 | 3,18E-03 |
| R-HSA-75953 | RNA Polymerase II Transcription Initiation | POLR2L | RNA polymerase II subunit L | 5441 | down | 1,52 | 3,58E-03 |
| R-HSA-75953 | RNA Polymerase II Transcription Initiation | GTF2H1 | general transcription factor IIH subunit 1 | 2965 | up | 1,33 | 1,85E-02 |
| R-HSA-75953 | RNA Polymerase II Transcription Initiation | POLR2F | RNA polymerase II subunit F | 5435 | down | 1,41 | 3,22E-02 |
| R-HSA-75953 | RNA Polymerase II Transcription Initiation | POLR2B | RNA polymerase II subunit B | 5431 | down | 1,23 | 3,81E-02 |
| R-HSA-74713 | IRS activation | GRB10 | growth factor receptor bound protein 10 | 2887 | up | 1,86 | 2,07E-02 |
| R-HSA-74713 | IRS activation | IRS2 | insulin receptor substrate 2 | 8660 | up | 2,21 | 2,22E-02 |
| R-HSA-74713 | IRS activation | IRS1 | insulin receptor substrate 1 | 3667 | down | 1,89 | 6,25E-03 |
| R-HSA-1257604 | PIP3 activates AKT signaling | PIK3R1 | phosphoinositide-3-kinase regulatory su | 5295 | down | 2,04 | 8,40E-03 |
| R-HSA-1257604 | PIP3 activates AKT signaling | HBEGF | heparin binding EGF like growth factor | 1839 | up | 3,67 | 5,67E-03 |
| R-HSA-1257604 | PIP3 activates AKT signaling | FGFR3 | fibroblast growth factor receptor 3 | 2261 | down | 1,70 | 4,60E-02 |
| R-HSA-1257604 | PIP3 activates AKT signaling | PIK3CA | phosphatidylinositol-4,5-bisphosphate 3 | 5290 | up | 1,33 | 3,17E-02 |
| R-HSA-1257604 | PIP3 activates AKT signaling | IRS2 | insulin receptor substrate 2 | 8660 | up | 2,21 | 2,22E-02 |
| R-HSA-1257604 | PIP3 activates AKT signaling | TRIB3 | tribbles pseudokinase 3 | 57761 | up | 3,17 | 8,07E-03 |
| R-HSA-1257604 | PIP3 activates AKT signaling | PRR5 | proline rich 5 | 55615 | down | 1,41 | 1,79E-03 |
| R-HSA-1257604 | PIP3 activates AKT signaling | AKT2 | AKT serine/threonine kinase 2 | 208 | down | 1,28 | 6,33E-03 |
| R-HSA-1257604 | PIP3 activates AKT signaling | FGF2 | fibroblast growth factor 2 | 2247 | down | 5,35 | 3,09E-02 |
| R-HSA-1257604 | PIP3 activates AKT signaling | AKT1 | AKT serine/threonine kinase 1 | 207 | down | 1,37 | 1,42E-03 |
| R-HSA-1257604 | PIP3 activates AKT signaling | IRS1 | insulin receptor substrate 1 | 3667 | down | 1,89 | 6,25E-03 |
| R-HSA-1257604 | PIP3 activates AKT signaling | ERBB2 | erb-b2 receptor tyrosine kinase 2 | 2064 | down | 1,47 | 1,93E-02 |
| R-HSA-1257604 | PIP3 activates AKT signaling | PTEN | phosphatase and tensin homolog | 5728 | up | 1,33 | 4,72E-02 |
| R-HSA-1257604 | PIP3 activates AKT signaling | AGO3 | argonaute 3, RISC catalytic component | 192669 | up | 1,36 | 1,80E-02 |
| R-HSA-111452 | Activation and oligomerization of BAK protein | BAK1 | BCL2 antagonist/killer 1 | 578 | down | 1,27 | 1,41E-02 |
| R-HSA-111452 | Activation and oligomerization of BAK protein | BID | BH3 interacting domain death agonist | 637 | down | 1,40 | 2,42E-02 |
| R-HSA-114294 | Activation, translocation and oligomerization of BAX | BID | BH3 interacting domain death agonist | 637 | down | 1,40 | 2,42E-02 |
| R-HSA-114294 | Activation, translocation and oligomerization of BAX | BAX | BCL2 associated X, apoptosis regulator | 581 | down | 1,22 | 2,61E-02 |
| R-HSA-1483101 | Synthesis of PS | PTDSS1 | phosphatidylserine synthase 1 | 9791 | down | 1,31 | 3,90E-02 |
| R-HSA-1483101 | Synthesis of PS | PTDSS2 | phosphatidylserine synthase 2 | 81490 | down | 1,30 | 1,46E-02 |
| R-HSA-3785653 | Myoclonic epilepsy of Lafora | GYS1 | glycogen synthase 1 | 2997 | down | 1,32 | 2,63E-02 |
| R-HSA-3785653 | Myoclonic epilepsy of Lafora | GYG2 | glycogenin 2 | 8908 | down | 1,52 | 4,53E-02 |
| R-HSA-3785653 | Myoclonic epilepsy of Lafora | NHLRC1 | NHL repeat containing E3 ubiquitin prot | 378884 | down | 1,81 | 1,84E-03 |
| R-HSA-5655291 | Signaling by FGFR4 in disease | HRAS | HRas proto-oncogene, GTPase | 3265 | down | 1,31 | 2,19E-03 |
| R-HSA-5655291 | Signaling by FGFR4 in disease | PIK3CA | phosphatidylinositol-4,5-bisphosphate 3 | 5290 | up | 1,33 | 3,17E-02 |
| R-HSA-5655291 | Signaling by FGFR4 in disease | PIK3R1 | phosphoinositide-3-kinase regulatory su | 5295 | down | 2,04 | 8,40E-03 |
| R-HSA-5655291 | Signaling by FGFR4 in disease | PLCG1 | phospholipase C gamma 1 | 5335 | down | 1,51 | 2,85E-02 |
| R-HSA-196299 | Beta-catenin phosphorylation cascade | CSNK1A1 | casein kinase 1 alpha 1 | 1452 | up | 1,35 | 2,84E-02 |
| R-HSA-196299 | Beta-catenin phosphorylation cascade | FRAT2 | FRAT2, WNT signaling pathway regulato | 23401 | down | 1,71 | 2,52E-04 |
| R-HSA-196299 | Beta-catenin phosphorylation cascade | FRAT1 | FRAT1, WNT signaling pathway regulato | 10023 | down | 1,48 | 3,14E-02 |
| R-HSA-196299 | Beta-catenin phosphorylation cascade | AMER1 | APC membrane recruitment protein 1 | 139285 | down | 1,78 | 4,15E-02 |
| R-HSA-196299 | Beta-catenin phosphorylation cascade | PPP2CB | protein phosphatase 2 catalytic subunit | 5516 | up | 1,30 | 4,79E-02 |
| R-HSA-450385 | Butyrate Response Factor 1 (BRF1) binds and destal | XRN1 | 5'-3' exoribonuclease 1 | 54464 | up | 1,30 | 1,96E-02 |
| R-HSA-450385 | Butyrate Response Factor 1 (BRF1) binds and destal | EXOSC4 | exosome component 4 | 54512 | down | 1,51 | 2,76E-03 |
| R-HSA-450385 | Butyrate Response Factor 1 (BRF1) binds and destal | AKT1 | AKT serine/threonine kinase 1 | 207 | down | 1,37 | 1,42E-03 |
| R-HSA-450385 | Butyrate Response Factor 1 (BRF1) binds and destal | DCP1A | decapping mRNA 1A | 55802 | up | 1,33 | 4,54E-02 |
| R-HSA-450385 | Butyrate Response Factor 1 (BRF1) binds and destal | EXOSC6 | exosome component 6 | 118460 | down | 1,50 | 1,04E-02 |
| R-HSA-5250924 | B-WICH complex positively regulates rRNA expressior | KAT2A | lysine acetyltransferase 2A | 2648 | down | 1,39 | 1,20E-02 |
| R-HSA-5250924 | B-WICH complex positively regulates rRNA expressior | CD3EAP | CD3e molecule associated protein | 10849 | down | 1,57 | 3,12E-03 |
| R-HSA-5250924 | B-WICH complex positively regulates rRNA expressior | TAF1A | TATA-box binding protein associated fa | 9015 | up | 2,20 | 2,24E-03 |
| R-HSA-5250924 | B-WICH complex positively regulates rRNA expressior | POLR2F | RNA polymerase II subunit F | 5435 | down | 1,41 | 3,22E-02 |
| R-HSA-5250924 | B-WICH complex positively regulates rRNA expressior | POLR2L | RNA polymerase II subunit L | 5441 | down | 1,52 | 3,58E-03 |
| R-HSA-5250924 | B-WICH complex positively regulates rRNA expressior | POLR1A | RNA polymerase I subunit A | 25885 | down | 1,30 | 4,32E-02 |
| R-HSA-5250924 | B-WICH complex positively regulates rRNA expressior | POLR2E | RNA polymerase II subunit E | 5434 | down | 1,31 | 3,18E-03 |
| R-HSA-5250924 | B-WICH complex positively regulates rRNA expressior | TAF1D | TATA-box binding protein associated fa | 79101 | up | 1,87 | 1,00E-02 |
| R-HSA-5250924 | B-WICH complex positively regulates rRNA expressior | POLR1C | RNA polymerase I subunit C | 9533 | down | 1,24 | 4,71E-02 |
| R-HSA-5250924 | B-WICH complex positively regulates rRNA expressior | H2AFX | H2A histone family member X | 3014 | down | 2,11 | 1,30E-03 |
| R-HSA-5250924 | B-WICH complex positively regulates rRNA expressior | MYBBP1A | MYB binding protein 1a | 10514 | down | 1,34 | 6,88E-03 |
| R-HSA-2586552 | Signaling by Leptin | IRS1 | insulin receptor substrate 1 | 3667 | down | 1,89 | 6,25E-03 |
| R-HSA-2586552 | Signaling by Leptin | LEPR | leptin receptor | 3953 | up | 1,33 | 4,94E-03 |
| R-HSA-2586552 | Signaling by Leptin | SH2B1 | SH2B adaptor protein 1 | 25970 | down | 1,31 | 1,81E-02 |
| R-HSA-2586552 | Signaling by Leptin | IRS2 | insulin receptor substrate 2 | 8660 | up | 2,21 | 2,22E-02 |
| R-HSA-196780 | Biotin transport and metabolism | PCCA | propionyl-CoA carboxylase alpha subun | 5095 | up | 1,37 | 2,03E-02 |
| R-HSA-196780 | Biotin transport and metabolism | MCCC1 | methylcrotonoyl-CoA carboxylase 1 | 56922 | up | 1,28 | 4,04E-02 |
| R-HSA-196780 | Biotin transport and metabolism | ACACA | acetyl-CoA carboxylase alpha | 31 | down | 1,40 | 2,54E-02 |
| R-HSA-196780 | Biotin transport and metabolism | SLC5A6 | solute carrier family 5 member 6 | 8884 | down | 1,35 | 8,34E-03 |
| R-HSA-200425 | Import of palmitoyl-CoA into the mitochondrial matrix | PRKAG2 | protein kinase AMP-activated non-catal | 51422 | up | 1,28 | 4,57E-02 |
| R-HSA-200425 | Import of palmitoyl-CoA into the mitochondrial matrix | RXRA | retinoid X receptor alpha | 6256 | down | 1,34 | 3,67E-03 |
| R-HSA-200425 | Import of palmitoyl-CoA into the mitochondrial matrix | PRKAB2 | protein kinase AMP-activated non-catal | 5565 | up | 1,83 | 1,27E-02 |
| R-HSA-200425 | Import of palmitoyl-CoA into the mitochondrial matrix | ACACA | acetyl-CoA carboxylase alpha | 31 | down | 1,40 | 2,54E-02 |
| R-HSA-167242 | Abortive elongation of HIV-1 transcript in the absence | POLR2B | RNA polymerase II subunit B | 5431 | down | 1,23 | 3,81E-02 |
| R-HSA-167242 | Abortive elongation of HIV-1 transcript in the absence | POLR2F | RNA polymerase II subunit F | 5435 | down | 1,41 | 3,22E-02 |
| R-HSA-167242 | Abortive elongation of HIV-1 transcript in the absence | POLR2L | RNA polymerase II subunit L | 5441 | down | 1,52 | 3,58E-03 |
| R-HSA-167242 | Abortive elongation of HIV-1 transcript in the absence | POLR2E | RNA polymerase II subunit E | 5434 | down | 1,31 | 3,18E-03 |
| R-HSA-5627123 | RHO GTPases activate PAKs | CALM2 | calmodulin 2 | 805 | down | 1,43 | 2,23E-02 |
| R-HSA-5627123 | RHO GTPases activate PAKs | CALM3 | calmodulin 3 | 808 | down | 1,43 | 2,23E-02 |
| R-HSA-5627123 | RHO GTPases activate PAKs | LIMK1 | LIM domain kinase 1 | 3984 | down | 1,44 | 6,49E-03 |
| R-HSA-5627123 | RHO GTPases activate PAKs | CALM1 | calmodulin 1 | 801 | down | 1,43 | 2,23E-02 |
| R-HSA-5627123 | RHO GTPases activate PAKs | PPP1R12A | protein phosphatase 1 regulatory subur | 4659 | up | 1,25 | 1,03E-02 |
| R-HSA-451308 | Activation of Ca-permeable Kainate Receptor | CALM2 | calmodulin 2 | 805 | down | 1,43 | 2,23E-02 |
| R-HSA-451308 | Activation of Ca-permeable Kainate Receptor | CALM3 | calmodulin 3 | 808 | down | 1,43 | 2,23E-02 |
| R-HSA-451308 | Activation of Ca-permeable Kainate Receptor | CALM1 | calmodulin 1 | 801 | down | 1,43 | 2,23E-02 |
| R-HSA-429958 | mRNA decay by 3' to 5' exoribonuclease | EXOSC6 | exosome component 6 | 118460 | down | 1,50 | 1,04E-02 |
| R-HSA-429958 | mRNA decay by 3' to 5' exoribonuclease | NT5C3B | 5'-nucleotidase, cytosolic IIIB | 115024 | down | 1,24 | 3,10E-03 |
| R-HSA-429958 | mRNA decay by 3' to 5' exoribonuclease | EXOSC4 | exosome component 4 | 54512 | down | 1,51 | 2,76E-03 |
| R-HSA-113501 | Inhibition of replication initiation of damaged DNA by | E2F1 | E2F transcription factor 1 | 1869 | down | 1,64 | 3,25E-02 |
| R-HSA-113501 | Inhibition of replication initiation of damaged DNA by | PPP2CB | protein phosphatase 2 catalytic subunit | 5516 | up | 1,30 | 4,79E-02 |
| R-HSA-113501 | Inhibition of replication initiation of damaged DNA by | TFDP1 | transcription factor Dp-1 | 7027 | down | 1,77 | 1,08E-02 |
| R-HSA-113501 | Inhibition of replication initiation of damaged DNA by | POLA2 | DNA polymerase alpha 2, accessory sub | 23649 | down | 1,72 | 2,46E-02 |
| R-HSA-72203 | Processing of Capped Intron-Containing Pre-mRNA | PTBP1 | polypyrimidine tract binding protein 1 | 5725 | down | 1,16 | 4,21E-02 |
| R-HSA-72203 | Processing of Capped Intron-Containing Pre-mRNA | POLR2B | RNA polymerase II subunit B | 5431 | down | 1,23 | 3,81E-02 |
| R-HSA-72203 | Processing of Capped Intron-Containing Pre-mRNA | POLR2F | RNA polymerase II subunit F | 5435 | down | 1,41 | 3,22E-02 |
| R-HSA-72203 | Processing of Capped Intron-Containing Pre-mRNA | POLR2L | RNA polymerase II subunit L | 5441 | down | 1,52 | 3,58E-03 |
| R-HSA-72203 | Processing of Capped Intron-Containing Pre-mRNA | POLR2E | RNA polymerase II subunit E | 5434 | down | 1,31 | 3,18E-03 |
| R-HSA-5673000 | RAF activation | PPP2CB | protein phosphatase 2 catalytic subunit | 5516 | up | 1,30 | 4,79E-02 |
| R-HSA-5673000 | RAF activation | HRAS | HRas proto-oncogene, GTPase | 3265 | down | 1,31 | 2,19E-03 |
| R-HSA-5673000 | RAF activation | BRAP | BRCA1 associated protein | 8315 | up | 1,34 | 1,14E-02 |
| R-HSA-5673000 | RAF activation | PHB | prohibitin | 5245 | down | 1,29 | 3,98E-02 |
| R-HSA-5673000 | RAF activation | BRAF | B-Raf proto-oncogene, serine/threonine | 673 | up | 1,52 | 4,53E-02 |
| R-HSA-5673000 | RAF activation | MARK3 | microtubule affinity regulating kinase 3 | 4140 | up | 1,40 | 1,07E-02 |
| R-HSA-5620924 | Intraflagellar transport | DYNC2LI1 | dynein cytoplasmic 2 light intermediate c | 51626 | up | 1,40 | 3,09E-02 |
| R-HSA-5620924 | Intraflagellar transport | TUBA1B | tubulin alpha 1b | 10376 | down | 1,78 | 2,30E-02 |
| R-HSA-5620924 | Intraflagellar transport | DYNC2H1 | dynein cytoplasmic 2 heavy chain 1 | 79659 | up | 1,52 | 9,51E-03 |
| R-HSA-5620924 | Intraflagellar transport | TUBA4A | tubulin alpha 4a | 7277 | down | 1,63 | 2,97E-02 |
| R-HSA-5620924 | Intraflagellar transport | KIF3C | kinesin family member 3C | 3797 | down | 1,58 | 4,57E-02 |
| R-HSA-5620924 | Intraflagellar transport | TTC30B | tetratricopeptide repeat domain 30B | 150737 | down | 1,89 | 3,05E-02 |
| R-HSA-5620924 | Intraflagellar transport | TNPO1 | transportin 1 | 3842 | up | 1,28 | 4,94E-02 |
| R-HSA-5620924 | Intraflagellar transport | IFT88 | intraflagellar transport 88 | 8100 | up | 1,17 | 4,84E-02 |
| R-HSA-5620924 | Intraflagellar transport | WDR19 | WD repeat domain 19 | 57728 | up | 1,32 | 4,95E-02 |
| R-HSA-5620924 | Intraflagellar transport | TRIP11 | thyroid hormone receptor interactor 11 | 9321 | up | 1,33 | 2,24E-03 |
| R-HSA-2173796 | SMAD2/SMAD3:SMAD4 heterotrimer regulates transc | TFDP1 | transcription factor Dp-1 | 7027 | down | 1,77 | 1,08E-02 |
| R-HSA-2173796 | SMAD2/SMAD3:SMAD4 heterotrimer regulates transc | CCNT1 | cyclin T1 | 904 | up | 1,34 | 3,05E-02 |
| R-HSA-2173796 | SMAD2/SMAD3:SMAD4 heterotrimer regulates transc | SMAD2 | SMAD family member 2 | 4087 | up | 1,29 | 2,19E-02 |
| R-HSA-2173796 | SMAD2/SMAD3:SMAD4 heterotrimer regulates transc | WWTR1 | WW domain containing transcription reg | 25937 | down | 1,99 | 1,00E-02 |
| R-HSA-2173796 | SMAD2/SMAD3:SMAD4 heterotrimer regulates transc | CDKN2B | cyclin dependent kinase inhibitor 2B | 1030 | up | 2,40 | 3,90E-02 |
| R-HSA-2173796 | SMAD2/SMAD3:SMAD4 heterotrimer regulates transc | TGIF2 | TGFB induced factor homeobox 2 | 60436 | down | 1,76 | 2,54E-02 |
| R-HSA-2173796 | SMAD2/SMAD3:SMAD4 heterotrimer regulates transc | MEN1 | menin 1 | 4221 | down | 1,49 | 4,75E-04 |
| R-HSA-3371453 | Regulation of HSF1-mediated heat shock response | NUP62 | nucleoporin 62 | 23636 | down | 1,52 | 4,39E-03 |
| R-HSA-3371453 | Regulation of HSF1-mediated heat shock response | HSPA5 | heat shock protein family A (Hsp70) mer | 3309 | up | 5,02 | 6,98E-04 |
| R-HSA-3371453 | Regulation of HSF1-mediated heat shock response | CCAR2 | cell cycle and apoptosis regulator 2 | 57805 | down | 1,23 | 2,13E-02 |
| R-HSA-3371453 | Regulation of HSF1-mediated heat shock response | DNAJB1 | DnaJ heat shock protein family (Hsp40) | 3337 | up | 2,02 | 1,41E-02 |
| R-HSA-3371453 | Regulation of HSF1-mediated heat shock response | HSPA9 | heat shock protein family A (Hsp70) mer | 3313 | up | 1,59 | 5,68E-03 |
| R-HSA-3371453 | Regulation of HSF1-mediated heat shock response | DNAJB6 | DnaJ heat shock protein family (Hsp40) | 10049 | down | 1,20 | 4,90E-02 |
| R-HSA-3371453 | Regulation of HSF1-mediated heat shock response | RPA1 | replication protein A1 | 6117 | down | 1,41 | 4,93E-02 |
| R-HSA-3371453 | Regulation of HSF1-mediated heat shock response | SIRT1 | sirtuin 1 | 23411 | up | 1,60 | 1,00E-02 |
| R-HSA-3371453 | Regulation of HSF1-mediated heat shock response | DNAJC2 | DnaJ heat shock protein family (Hsp40) | 27000 | up | 1,50 | 1,50E-02 |
| R-HSA-3371453 | Regulation of HSF1-mediated heat shock response | NUP58 | nucleoporin 58 | 9818 | up | 1,61 | 1,28E-02 |
| R-HSA-3371453 | Regulation of HSF1-mediated heat shock response | BAG3 | BCL2 associated athanogene 3 | 9531 | up | 1,52 | 4,07E-02 |
| R-HSA-3371453 | Regulation of HSF1-mediated heat shock response | HSPA13 | heat shock protein family A (Hsp70) mer | 6782 | up | 3,69 | 7,84E-04 |
| R-HSA-1538133 | G0 and Early G1 | TFDP1 | transcription factor Dp-1 | 7027 | down | 1,77 | 1,08E-02 |
| R-HSA-1538133 | G0 and Early G1 | PCNA | proliferating cell nuclear antigen | 5111 | down | 2,14 | 4,63E-02 |
| R-HSA-1538133 | G0 and Early G1 | CDC25A | cell division cycle 25A | 993 | down | 1,92 | 1,48E-02 |
| R-HSA-1538133 | G0 and Early G1 | E2F1 | E2F transcription factor 1 | 1869 | down | 1,64 | 3,25E-02 |
| R-HSA-163841 | Gamma carboxylation, hypusine formation and arylsu | ICMT | isoprenylcysteine carboxyl methyltransfe | 23463 | down | 1,63 | 2,39E-02 |
| R-HSA-163841 | Gamma carboxylation, hypusine formation and arylsu | FN3KRP | fructosamine 3 kinase related protein | 79672 | down | 1,37 | 1,69E-02 |
| R-HSA-5654227 | Phospholipase C-mediated cascade; FGFR3 | FGF2 | fibroblast growth factor 2 | 2247 | down | 5,35 | 3,09E-02 |
| R-HSA-5654227 | Phospholipase C-mediated cascade; FGFR3 | PLCG1 | phospholipase C gamma 1 | 5335 | down | 1,51 | 2,85E-02 |
| R-HSA-5654227 | Phospholipase C-mediated cascade; FGFR3 | FGFR3 | fibroblast growth factor receptor 3 | 2261 | down | 1,70 | 4,60E-02 |
| R-HSA-198323 | AKT phosphorylates targets in the cytosol | AKT1 | AKT serine/threonine kinase 1 | 207 | down | 1,37 | 1,42E-03 |
| R-HSA-198323 | AKT phosphorylates targets in the cytosol | AKT2 | AKT serine/threonine kinase 2 | 208 | down | 1,28 | 6,33E-03 |
| R-HSA-198323 | AKT phosphorylates targets in the cytosol | CDKN1A | cyclin dependent kinase inhibitor 1A | 1026 | up | 3,62 | 5,43E-03 |
| R-HSA-198323 | AKT phosphorylates targets in the cytosol | AKT1S1 | AKT1 substrate 1 | 84335 | down | 1,21 | 4,33E-02 |
| R-HSA-171007 | p38MAPK events | RALGDS | ral guanine nucleotide dissociation stimu | 5900 | down | 1,30 | 3,49E-02 |
| R-HSA-171007 | p38MAPK events | MAPK12 | mitogen-activated protein kinase 12 | 6300 | down | 1,37 | 3,32E-03 |
| R-HSA-171007 | p38MAPK events | HRAS | HRas proto-oncogene, GTPase | 3265 | down | 1,31 | 2,19E-03 |
| R-HSA-2564830 | Cytosolic iron-sulfur cluster assembly | POLD1 | DNA polymerase delta 1, catalytic subur | 5424 | down | 1,59 | 4,62E-02 |
| R-HSA-2564830 | Cytosolic iron-sulfur cluster assembly | ERCC2 | ERCC excision repair 2, TFIIH core com | 2068 | down | 1,29 | 2,20E-02 |
| R-HSA-2564830 | Cytosolic iron-sulfur cluster assembly | CIAO1 | cytosolic iron-sulfur assembly componer | 9391 | down | 1,24 | 1,25E-02 |

| | | | | | | | |
|---|---|---|---|---|---|---|---|
| R-HSA-4641265 | Repression of WNT target genes | TCF7 | transcription factor 7 | 6932 | down | 1,47 | 1,07E-02 |
| R-HSA-4641265 | Repression of WNT target genes | CTBP1 | C-terminal binding protein 1 | 1487 | down | 1,16 | 3,77E-02 |
| R-HSA-4641265 | Repression of WNT target genes | AES | amino-terminal enhancer of split | 166 | down | 1,18 | 2,84E-02 |
| R-HSA-5607763 | CLEC7A (Dectin-1) induces NFAT activation | CALM1 | calmodulin 1 | 801 | down | 1,43 | 2,23E-02 |
| R-HSA-5607763 | CLEC7A (Dectin-1) induces NFAT activation | CALM3 | calmodulin 3 | 808 | down | 1,43 | 2,23E-02 |
| R-HSA-5607763 | CLEC7A (Dectin-1) induces NFAT activation | CALM2 | calmodulin 2 | 805 | down | 1,43 | 2,23E-02 |
| R-HSA-425561 | Sodium/Calcium exchangers | CALM3 | calmodulin 3 | 808 | down | 1,43 | 2,23E-02 |
| R-HSA-425561 | Sodium/Calcium exchangers | CALM2 | calmodulin 2 | 805 | down | 1,43 | 2,23E-02 |
| R-HSA-425561 | Sodium/Calcium exchangers | CALM1 | calmodulin 1 | 801 | down | 1,43 | 2,23E-02 |
| R-HSA-350054 | Notch-HLH transcription pathway | MAMLD1 | mastermind like domain containing 1 | 10046 | down | 3,36 | 4,56E-02 |
| R-HSA-350054 | Notch-HLH transcription pathway | KAT2A | lysine acetyltransferase 2A | 2648 | down | 1,39 | 1,20E-02 |
| R-HSA-350054 | Notch-HLH transcription pathway | NOTCH1 | notch 1 | 4851 | down | 1,62 | 2,46E-02 |
| R-HSA-5655302 | Signaling by FGFR1 in disease | HRAS | HRas proto-oncogene, GTPase | 3265 | down | 1,31 | 2,19E-03 |
| R-HSA-5655302 | Signaling by FGFR1 in disease | LRRFIP1 | LRR binding FLII interacting protein 1 | 9208 | up | 1,28 | 2,23E-02 |
| R-HSA-5655302 | Signaling by FGFR1 in disease | FGFR1OP2 | FGFR1 oncogene partner 2 | 26127 | up | 1,39 | 2,09E-02 |
| R-HSA-5655302 | Signaling by FGFR1 in disease | FGF2 | fibroblast growth factor 2 | 2247 | down | 5,35 | 3,09E-02 |
| R-HSA-5655302 | Signaling by FGFR1 in disease | PLCG1 | phospholipase C gamma 1 | 5335 | down | 1,51 | 2,85E-02 |
| R-HSA-5655302 | Signaling by FGFR1 in disease | PIK3R1 | phosphoinositide-3-kinase regulatory su | 5295 | down | 2,04 | 8,40E-03 |
| R-HSA-5655302 | Signaling by FGFR1 in disease | PIK3CA | phosphatidylinositol-4,5-bisphosphate 3 | 5290 | up | 1,33 | 3,17E-02 |
| R-HSA-1445148 | Translocation of SLC2A4 (GLUT4) to the plasma mem | ACTG1 | actin gamma 1 | 71 | up | 1,58 | 1,66E-02 |
| R-HSA-1445148 | Translocation of SLC2A4 (GLUT4) to the plasma mem | PRKAG2 | protein kinase AMP-activated non-catal | 51422 | up | 1,28 | 4,57E-02 |
| R-HSA-1445148 | Translocation of SLC2A4 (GLUT4) to the plasma mem | TUBA1B | tubulin alpha 1b | 10376 | down | 1,78 | 2,30E-02 |
| R-HSA-1445148 | Translocation of SLC2A4 (GLUT4) to the plasma mem | TUBA4A | tubulin alpha 4a | 7277 | down | 1,63 | 2,97E-02 |
| R-HSA-1445148 | Translocation of SLC2A4 (GLUT4) to the plasma mem | CALM1 | calmodulin 1 | 801 | down | 1,43 | 2,23E-02 |
| R-HSA-1445148 | Translocation of SLC2A4 (GLUT4) to the plasma mem | AKT1 | AKT serine/threonine kinase 1 | 207 | down | 1,37 | 1,42E-03 |
| R-HSA-1445148 | Translocation of SLC2A4 (GLUT4) to the plasma mem | PRKAB2 | protein kinase AMP-activated non-catal | 5565 | up | 1,83 | 1,27E-02 |
| R-HSA-1445148 | Translocation of SLC2A4 (GLUT4) to the plasma mem | VAMP2 | vesicle associated membrane protein 2 | 6844 | up | 1,50 | 4,82E-02 |
| R-HSA-1445148 | Translocation of SLC2A4 (GLUT4) to the plasma mem | AKT2 | AKT serine/threonine kinase 2 | 208 | down | 1,28 | 6,33E-03 |
| R-HSA-1445148 | Translocation of SLC2A4 (GLUT4) to the plasma mem | YWHAH | tyrosine 3-monooxygenase/tryptophan 5 | 7533 | down | 1,33 | 4,79E-02 |
| R-HSA-1445148 | Translocation of SLC2A4 (GLUT4) to the plasma mem | CALM2 | calmodulin 2 | 805 | down | 1,43 | 2,23E-02 |
| R-HSA-1445148 | Translocation of SLC2A4 (GLUT4) to the plasma mem | CALM3 | calmodulin 3 | 808 | down | 1,43 | 2,23E-02 |
| R-HSA-3371571 | HSF1-dependent transactivation | CAMK2D | calcium/calmodulin dependent protein k | 817 | up | 1,29 | 1,49E-02 |
| R-HSA-3371571 | HSF1-dependent transactivation | AKT1S1 | AKT1 substrate 1 | 84335 | down | 1,21 | 4,33E-02 |
| R-HSA-3371571 | HSF1-dependent transactivation | DNAJB1 | DnaJ heat shock protein family (Hsp40) | 3337 | up | 2,02 | 1,41E-02 |
| R-HSA-3371571 | HSF1-dependent transactivation | HSPB8 | heat shock protein family B (small) mem | 26353 | up | 2,27 | 4,99E-03 |
| R-HSA-3371571 | HSF1-dependent transactivation | RPTOR | regulatory associated protein of MTOR c | 57521 | down | 1,20 | 1,16E-02 |
| R-HSA-70895 | Branched-chain amino acid catabolism | MCCC1 | methylcrotonoyl-CoA carboxylase 1 | 56922 | up | 1,28 | 4,04E-02 |
| R-HSA-70895 | Branched-chain amino acid catabolism | BCAT2 | branched chain amino acid transaminas | 587 | down | 1,34 | 1,96E-02 |
| R-HSA-70895 | Branched-chain amino acid catabolism | PPM1K | protein phosphatase, Mg2+/Mn2+ depe | 152926 | up | 2,16 | 3,86E-02 |
| R-HSA-70895 | Branched-chain amino acid catabolism | DLD | dihydrolipoamide dehydrogenase | 1738 | up | 1,16 | 4,53E-02 |
| R-HSA-70895 | Branched-chain amino acid catabolism | ALDH6A1 | aldehyde dehydrogenase 6 family mem | 4329 | up | 1,60 | 3,50E-02 |
| R-HSA-373752 | Netrin-1 signaling | UNC5B | unc-5 netrin receptor B | 219699 | up | 4,39 | 5,27E-03 |
| R-HSA-373752 | Netrin-1 signaling | SLIT1 | slit guidance ligand 1 | 6585 | up | 7,59 | 2,13E-03 |
| R-HSA-373752 | Netrin-1 signaling | SIAH1 | siah E3 ubiquitin protein ligase 1 | 6477 | up | 1,74 | 1,66E-02 |
| R-HSA-373752 | Netrin-1 signaling | NTN4 | netrin 4 | 59277 | up | 2,22 | 5,93E-03 |
| R-HSA-373752 | Netrin-1 signaling | AGAP2 | ArfGAP with GTPase domain, ankyrin re | 116986 | down | 2,02 | 3,75E-02 |
| R-HSA-5610787 | Hedgehog 'off' state | GPR161 | G protein-coupled receptor 161 | 23432 | down | 2,43 | 1,49E-03 |
| R-HSA-5610787 | Hedgehog 'off' state | PRKAR1B | protein kinase cAMP-dependent type I r | 5575 | down | 1,50 | 6,80E-04 |
| R-HSA-5610787 | Hedgehog 'off' state | TULP3 | tubby like protein 3 | 7289 | up | 2,16 | 7,18E-03 |
| R-HSA-5610787 | Hedgehog 'off' state | SUFU | SUFU negative regulator of hedgehog s | 51684 | down | 1,41 | 3,37E-02 |
| R-HSA-5610787 | Hedgehog 'off' state | TUBA4A | tubulin alpha 4a | 7277 | down | 1,63 | 2,97E-02 |
| R-HSA-5610787 | Hedgehog 'off' state | ADCY3 | adenylate cyclase 3 | 109 | down | 1,45 | 1,03E-02 |
| R-HSA-5610787 | Hedgehog 'off' state | TUBA1B | tubulin alpha 1b | 10376 | down | 1,78 | 2,30E-02 |
| R-HSA-5610787 | Hedgehog 'off' state | DYNC2H1 | dynein cytoplasmic 2 heavy chain 1 | 79659 | up | 1,52 | 9,51E-03 |
| R-HSA-5610787 | Hedgehog 'off' state | IFT88 | intraflagellar transport 88 | 8100 | up | 1,17 | 4,84E-02 |
| R-HSA-5610787 | Hedgehog 'off' state | WDR19 | WD repeat domain 19 | 57728 | up | 1,32 | 4,95E-02 |
| R-HSA-499943 | Interconversion of nucleotide di- and triphosphates | NME4 | NME/NM23 nucleoside diphosphate kin | 4833 | down | 1,47 | 3,85E-02 |
| R-HSA-499943 | Interconversion of nucleotide di- and triphosphates | AK1 | adenylate kinase 1 | 203 | down | 1,57 | 9,91E-03 |
| R-HSA-499943 | Interconversion of nucleotide di- and triphosphates | AK4 | adenylate kinase 4 | 205 | down | 1,76 | 4,87E-02 |
| R-HSA-499943 | Interconversion of nucleotide di- and triphosphates | DCTPP1 | dCTP pyrophosphatase 1 | 79077 | down | 1,69 | 1,60E-02 |
| R-HSA-1971475 | A tetrasaccharide linker sequence is required for GAG | SDC1 | syndecan 1 | 6382 | down | 2,17 | 1,61E-02 |
| R-HSA-1971475 | A tetrasaccharide linker sequence is required for GAG | SDC3 | syndecan 3 | 9672 | down | 1,68 | 1,97E-02 |
| R-HSA-1971475 | A tetrasaccharide linker sequence is required for GAG | CSPG4 | chondroitin sulfate proteoglycan 4 | 1464 | down | 2,22 | 3,99E-02 |
| R-HSA-1971475 | A tetrasaccharide linker sequence is required for GAG | XYLT2 | xylosyltransferase 2 | 64132 | down | 1,98 | 2,03E-02 |
| R-HSA-1169408 | ISG15 antiviral mechanism | NUP58 | nucleoporin 58 | 9818 | up | 1,61 | 1,28E-02 |
| R-HSA-1169408 | ISG15 antiviral mechanism | JAK1 | Janus kinase 1 | 3716 | up | 1,53 | 2,13E-02 |
| R-HSA-1169408 | ISG15 antiviral mechanism | PPM1B | protein phosphatase, Mg2+/Mn2+ depe | 5495 | up | 1,53 | 2,84E-02 |
| R-HSA-1169408 | ISG15 antiviral mechanism | EIF4G3 | eukaryotic translation initiation factor 4 g | 8672 | up | 1,19 | 4,60E-02 |
| R-HSA-1169408 | ISG15 antiviral mechanism | NEDD4 | neural precursor cell expressed, develop | 4734 | up | 1,99 | 3,68E-03 |
| R-HSA-1169408 | ISG15 antiviral mechanism | KPNA5 | karyopherin subunit alpha 5 | 3841 | up | 1,61 | 1,30E-02 |
| R-HSA-1169408 | ISG15 antiviral mechanism | PLCG1 | phospholipase C gamma 1 | 5335 | down | 1,51 | 2,85E-02 |
| R-HSA-1169408 | ISG15 antiviral mechanism | KPNA4 | karyopherin subunit alpha 4 | 3840 | up | 1,61 | 4,03E-03 |
| R-HSA-1169408 | ISG15 antiviral mechanism | IRF3 | interferon regulatory factor 3 | 3661 | down | 1,33 | 2,58E-03 |
| R-HSA-1169408 | ISG15 antiviral mechanism | NUP62 | nucleoporin 62 | 23636 | down | 1,52 | 4,39E-03 |
| R-HSA-1169408 | ISG15 antiviral mechanism | ARIH1 | ariadne RBR E3 ubiquitin protein ligase | 25820 | up | 1,36 | 8,38E-03 |
| R-HSA-1169408 | ISG15 antiviral mechanism | EIF4A2 | eukaryotic translation initiation factor 4A | 1974 | up | 1,68 | 4,70E-03 |
| R-HSA-3134975 | Regulation of innate immune responses to cytosolic D | TRIM32 | tripartite motif containing 32 | 22954 | down | 1,22 | 3,95E-02 |
| R-HSA-3134975 | Regulation of innate immune responses to cytosolic D | IRF3 | interferon regulatory factor 3 | 3661 | down | 1,33 | 2,58E-03 |
| R-HSA-3134975 | Regulation of innate immune responses to cytosolic D | DTX4 | deltex E3 ubiquitin ligase 4 | 23220 | down | 1,52 | 4,63E-02 |
| R-HSA-918233 | TRAF3-dependent IRF activation pathway | IKBKE | inhibitor of nuclear factor kappa B kinas | 9641 | down | 1,67 | 1,00E-02 |
| R-HSA-918233 | TRAF3-dependent IRF activation pathway | IRF3 | interferon regulatory factor 3 | 3661 | down | 1,33 | 2,58E-03 |
| R-HSA-918233 | TRAF3-dependent IRF activation pathway | TRAF3 | TNF receptor associated factor 3 | 7187 | down | 1,40 | 1,36E-02 |
| R-HSA-5655253 | Signaling by FGFR2 in disease | PIK3R1 | phosphoinositide-3-kinase regulatory su | 5295 | down | 2,04 | 8,40E-03 |
| R-HSA-5655253 | Signaling by FGFR2 in disease | PLCG1 | phospholipase C gamma 1 | 5335 | down | 1,51 | 2,85E-02 |
| R-HSA-5655253 | Signaling by FGFR2 in disease | FGF2 | fibroblast growth factor 2 | 2247 | down | 5,35 | 3,09E-02 |
| R-HSA-5655253 | Signaling by FGFR2 in disease | PIK3CA | phosphatidylinositol-4,5-bisphosphate 3 | 5290 | up | 1,33 | 3,17E-02 |
| R-HSA-5655253 | Signaling by FGFR2 in disease | HRAS | HRas proto-oncogene, GTPase | 3265 | down | 1,31 | 2,19E-03 |
| R-HSA-5357786 | TNFR1-induced proapoptotic signaling | CYLD | CYLD lysine 63 deubiquitinase | 1540 | up | 1,34 | 4,86E-02 |
| R-HSA-5357786 | TNFR1-induced proapoptotic signaling | FADD | Fas associated via death domain | 8772 | down | 1,31 | 8,90E-03 |
| R-HSA-5357786 | TNFR1-induced proapoptotic signaling | OTUD7B | OTU deubiquitinase 7B | 56957 | up | 1,27 | 3,81E-02 |
| R-HSA-5357786 | TNFR1-induced proapoptotic signaling | TRAF2 | TNF receptor associated factor 2 | 7186 | down | 1,47 | 2,65E-02 |
| R-HSA-209543 | p75NTR recruits signalling complexes | RIPK2 | receptor interacting serine/threonine kin | 8767 | up | 1,72 | 8,31E-03 |
| R-HSA-209543 | p75NTR recruits signalling complexes | MYD88 | myeloid differentiation primary response | 4615 | down | 1,55 | 1,85E-02 |
| R-HSA-209543 | p75NTR recruits signalling complexes | SQSTM1 | sequestosome 1 | 8878 | up | 1,88 | 2,42E-02 |
| R-HSA-209543 | p75NTR recruits signalling complexes | NGFR | nerve growth factor receptor | 4804 | up | 12,35 | 2,75E-03 |
| R-HSA-400206 | Regulation of lipid metabolism by Peroxisome prolifera | CHD9 | chromodomain helicase DNA binding pr | 80205 | up | 1,32 | 1,42E-03 |
| R-HSA-400206 | Regulation of lipid metabolism by Peroxisome prolifera | RXRA | retinoid X receptor alpha | 6256 | down | 1,34 | 3,67E-03 |
| R-HSA-400206 | Regulation of lipid metabolism by Peroxisome prolifera | CARM1 | coactivator associated arginine methyltr | 10498 | down | 1,36 | 1,60E-02 |
| R-HSA-400206 | Regulation of lipid metabolism by Peroxisome prolifera | TBL1X | transducin beta like 1 X-linked | 6907 | down | 1,20 | 2,84E-02 |
| R-HSA-400206 | Regulation of lipid metabolism by Peroxisome prolifera | TGS1 | trimethylguanosine synthase 1 | 96764 | up | 1,63 | 9,37E-03 |
| R-HSA-452723 | Transcriptional regulation of pluripotent stem cells | POLR2E | RNA polymerase II subunit E | 5434 | down | 1,31 | 3,18E-03 |
| R-HSA-452723 | Transcriptional regulation of pluripotent stem cells | SMAD2 | SMAD family member 2 | 4087 | up | 1,29 | 2,19E-02 |
| R-HSA-452723 | Transcriptional regulation of pluripotent stem cells | KLF4 | Kruppel like factor 4 | 9314 | up | 1,96 | 3,94E-02 |
| R-HSA-452723 | Transcriptional regulation of pluripotent stem cells | POLR2B | RNA polymerase II subunit B | 5431 | down | 1,23 | 3,81E-02 |
| R-HSA-452723 | Transcriptional regulation of pluripotent stem cells | POLR2F | RNA polymerase II subunit F | 5435 | down | 1,41 | 3,22E-02 |
| R-HSA-452723 | Transcriptional regulation of pluripotent stem cells | POLR2L | RNA polymerase II subunit L | 5441 | down | 1,52 | 3,58E-03 |
| R-HSA-2022854 | Keratan sulfate biosynthesis | ST3GAL2 | ST3 beta-galactoside alpha-2,3-sialyltra | 6483 | down | 1,66 | 3,84E-03 |
| R-HSA-2022854 | Keratan sulfate biosynthesis | B4GALT2 | beta-1,4-galactosyltransferase 2 | 8704 | down | 1,45 | 2,13E-03 |
| R-HSA-2022854 | Keratan sulfate biosynthesis | B3GNT7 | UDP-GlcNAc:betaGal beta-1,3-N-acetylg | 93010 | down | 2,40 | 3,48E-02 |
| R-HSA-2022854 | Keratan sulfate biosynthesis | B4GAT1 | beta-1,4-glucuronyltransferase 1 | 11041 | down | 2,42 | 1,49E-02 |
| R-HSA-3928662 | EPHB-mediated forward signaling | YES1 | YES proto-oncogene 1, Src family tyros | 7525 | up | 1,30 | 2,08E-02 |
| R-HSA-3928662 | EPHB-mediated forward signaling | LYN | LYN proto-oncogene, Src family tyrosine | 4067 | down | 1,36 | 2,34E-02 |
| R-HSA-3928662 | EPHB-mediated forward signaling | CFL1 | cofilin 1 | 1072 | down | 1,29 | 3,02E-02 |
| R-HSA-3928662 | EPHB-mediated forward signaling | HRAS | HRas proto-oncogene, GTPase | 3265 | down | 1,31 | 2,19E-03 |
| R-HSA-3928662 | EPHB-mediated forward signaling | LIMK1 | LIM domain kinase 1 | 3984 | down | 1,44 | 6,49E-03 |
| R-HSA-3928662 | EPHB-mediated forward signaling | EPHB2 | EPH receptor B2 | 2048 | down | 1,66 | 2,52E-02 |
| R-HSA-3928662 | EPHB-mediated forward signaling | ACTG1 | actin gamma 1 | 71 | up | 1,58 | 1,66E-02 |
| R-HSA-5637810 | Constitutive Signaling by EGFRvIII | HRAS | HRas proto-oncogene, GTPase | 3265 | down | 1,31 | 2,19E-03 |
| R-HSA-5637810 | Constitutive Signaling by EGFRvIII | PIK3CA | phosphatidylinositol-4,5-bisphosphate 3 | 5290 | up | 1,33 | 3,17E-02 |
| R-HSA-5637810 | Constitutive Signaling by EGFRvIII | PIK3R1 | phosphoinositide-3-kinase regulatory su | 5295 | down | 2,04 | 8,40E-03 |
| R-HSA-5637810 | Constitutive Signaling by EGFRvIII | PLCG1 | phospholipase C gamma 1 | 5335 | down | 1,51 | 2,85E-02 |
| R-HSA-917937 | Iron uptake and transport | SLC46A1 | solute carrier family 46 member 1 | 113235 | down | 1,66 | 4,58E-02 |
| R-HSA-917937 | Iron uptake and transport | SLC40A1 | solute carrier family 40 member 1 | 30061 | down | 1,80 | 2,30E-02 |
| R-HSA-917937 | Iron uptake and transport | HMOX2 | heme oxygenase 2 | 3163 | down | 1,29 | 4,87E-02 |
| R-HSA-3322077 | Glycogen synthesis | GYG2 | glycogenin 2 | 8908 | down | 1,52 | 4,53E-02 |
| R-HSA-3322077 | Glycogen synthesis | NHLRC1 | NHL repeat containing E3 ubiquitin prot | 378884 | down | 1,81 | 1,84E-03 |
| R-HSA-3322077 | Glycogen synthesis | GYS1 | glycogen synthase 1 | 2997 | down | 1,32 | 2,63E-02 |
| R-HSA-3322077 | Glycogen synthesis | UGP2 | UDP-glucose pyrophosphorylase 2 | 7360 | up | 1,37 | 3,47E-02 |
| R-HSA-3928663 | EPHA-mediated growth cone collapse | EPHA5 | EPH receptor A5 | 2044 | down | 18,87 | 1,96E-02 |
| R-HSA-3928663 | EPHA-mediated growth cone collapse | EFNA3 | ephrin A3 | 1944 | down | 2,00 | 4,89E-02 |
| R-HSA-3928663 | EPHA-mediated growth cone collapse | EPHA2 | EPH receptor A2 | 1969 | up | 1,89 | 2,51E-02 |
| R-HSA-3928663 | EPHA-mediated growth cone collapse | EFNA4 | ephrin A4 | 1945 | down | 1,43 | 4,92E-02 |
| R-HSA-3928663 | EPHA-mediated growth cone collapse | YES1 | YES proto-oncogene 1, Src family tyros | 7525 | up | 1,30 | 2,08E-02 |
| R-HSA-3928663 | EPHA-mediated growth cone collapse | VEGFA | vascular endothelial growth factor A | 7422 | up | 2,38 | 1,68E-02 |
| R-HSA-3928663 | EPHA-mediated growth cone collapse | LYN | LYN proto-oncogene, Src family tyrosine | 4067 | down | 1,36 | 2,34E-02 |
| R-HSA-112412 | SOS-mediated signalling | IRS1 | insulin receptor substrate 1 | 3667 | down | 1,89 | 6,25E-03 |
| R-HSA-112412 | SOS-mediated signalling | HRAS | HRas proto-oncogene, GTPase | 3265 | down | 1,31 | 2,19E-03 |
| R-HSA-112412 | SOS-mediated signalling | IRS2 | insulin receptor substrate 2 | 8660 | up | 2,21 | 2,22E-02 |
| R-HSA-2660826 | Constitutive Signaling by NOTCH1 t(7;9)(NOTCH1:M1 | ADAM17 | ADAM metallopeptidase domain 17 | 6868 | up | 2,59 | 1,60E-03 |
| R-HSA-2660826 | Constitutive Signaling by NOTCH1 t(7;9)(NOTCH1:M1 | JAG1 | jagged 1 | 182 | up | 1,84 | 3,27E-02 |
| R-HSA-2660826 | Constitutive Signaling by NOTCH1 t(7;9)(NOTCH1:M1 | NOTCH1 | notch 1 | 4851 | down | 1,62 | 2,46E-02 |
| R-HSA-1251932 | PLCG1 events in ERBB2 signaling | PLCG1 | phospholipase C gamma 1 | 5335 | down | 1,51 | 2,85E-02 |
| R-HSA-1251932 | PLCG1 events in ERBB2 signaling | ERBB2 | erb-b2 receptor tyrosine kinase 2 | 2064 | down | 1,47 | 1,93E-02 |
| R-HSA-199220 | Vitamin B5 (pantothenate) metabolism | SLC5A6 | solute carrier family 5 member 6 | 8884 | down | 1,35 | 8,34E-03 |
| R-HSA-199220 | Vitamin B5 (pantothenate) metabolism | FASN | fatty acid synthase | 2194 | down | 1,96 | 8,31E-03 |
| R-HSA-211163 | AKT-mediated inactivation of FOXO1A | AKT1 | AKT serine/threonine kinase 1 | 207 | down | 1,37 | 1,42E-03 |
| R-HSA-211163 | AKT-mediated inactivation of FOXO1A | AKT2 | AKT serine/threonine kinase 2 | 208 | down | 1,28 | 6,33E-03 |
| R-HSA-5696395 | Formation of Incision Complex in GG-NER | RAD23B | RAD23 homolog B, nucleotide excision | 5887 | up | 1,23 | 3,34E-02 |
| R-HSA-5696395 | Formation of Incision Complex in GG-NER | ERCC2 | ERCC excision repair 2, TFIIH core com | 2068 | down | 1,29 | 2,20E-02 |
| R-HSA-5696395 | Formation of Incision Complex in GG-NER | ERCC3 | ERCC excision repair 3, TFIIH core com | 2071 | down | 1,20 | 2,37E-02 |
| R-HSA-5696395 | Formation of Incision Complex in GG-NER | CDK7 | cyclin dependent kinase 7 | 1022 | up | 1,37 | 4,38E-02 |
| R-HSA-5696395 | Formation of Incision Complex in GG-NER | SUMO3 | small ubiquitin-like modifier 3 | 6612 | down | 1,24 | 4,67E-02 |
| R-HSA-5696395 | Formation of Incision Complex in GG-NER | RPA1 | replication protein A1 | 6117 | down | 1,41 | 4,93E-02 |
| R-HSA-5696395 | Formation of Incision Complex in GG-NER | PARP1 | poly(ADP-ribose) polymerase 1 | 142 | down | 1,50 | 3,92E-02 |
| R-HSA-5696395 | Formation of Incision Complex in GG-NER | GTF2H1 | general transcription factor IIH subunit 1 | 2965 | up | 1,33 | 1,85E-02 |
| R-HSA-168325 | Viral Messenger RNA Synthesis | POLR2E | RNA polymerase II subunit E | 5434 | down | 1,31 | 3,18E-03 |
| R-HSA-168325 | Viral Messenger RNA Synthesis | NUP62 | nucleoporin 62 | 23636 | down | 1,52 | 4,39E-03 |
| R-HSA-168325 | Viral Messenger RNA Synthesis | POLR2F | RNA polymerase II subunit F | 5435 | down | 1,41 | 3,22E-02 |
| R-HSA-168325 | Viral Messenger RNA Synthesis | POLR2L | RNA polymerase II subunit L | 5441 | down | 1,52 | 3,58E-03 |
| R-HSA-168325 | Viral Messenger RNA Synthesis | NUP58 | nucleoporin 58 | 9818 | up | 1,61 | 1,28E-02 |
| R-HSA-168325 | Viral Messenger RNA Synthesis | POLR2B | RNA polymerase II subunit B | 5431 | down | 1,23 | 3,81E-02 |
| R-HSA-168638 | NOD1/2 Signaling Pathway | MAPK12 | mitogen-activated protein kinase 12 | 6300 | down | 1,37 | 3,32E-03 |
| R-HSA-168638 | NOD1/2 Signaling Pathway | ITCH | itchy E3 ubiquitin protein ligase | 83737 | up | 1,51 | 6,64E-03 |
| R-HSA-168638 | NOD1/2 Signaling Pathway | CYLD | CYLD lysine 63 deubiquitinase | 1540 | up | 1,34 | 4,86E-02 |
| R-HSA-168638 | NOD1/2 Signaling Pathway | TAB1 | TGF-beta activated kinase 1 (MAP3K7) | 10454 | down | 1,29 | 1,94E-02 |
| R-HSA-168638 | NOD1/2 Signaling Pathway | RIPK2 | receptor interacting serine/threonine kin | 8767 | up | 1,72 | 8,31E-03 |
| R-HSA-168638 | NOD1/2 Signaling Pathway | TAB3 | TGF-beta activated kinase 1 and MAP3 | 257397 | up | 1,40 | 4,94E-02 |
| R-HSA-168638 | NOD1/2 Signaling Pathway | BIRC2 | baculoviral IAP repeat containing 2 | 329 | up | 1,38 | 1,86E-02 |
| R-HSA-165159 | mTOR signalling | AKT1S1 | AKT1 substrate 1 | 84335 | down | 1,21 | 4,33E-02 |
| R-HSA-165159 | mTOR signalling | RRAGC | Ras related GTP binding C | 64121 | up | 1,78 | 6,10E-03 |
| R-HSA-165159 | mTOR signalling | AKT1 | AKT serine/threonine kinase 1 | 207 | down | 1,37 | 1,42E-03 |
| R-HSA-165159 | mTOR signalling | RPTOR | regulatory associated protein of MTOR c | 57521 | down | 1,20 | 1,16E-02 |
| R-HSA-399954 | Sema3A PAK dependent Axon repulsion | PLXNA1 | plexin A1 | 5361 | down | 1,28 | 9,70E-04 |
| R-HSA-399954 | Sema3A PAK dependent Axon repulsion | CFL1 | cofilin 1 | 1072 | down | 1,29 | 3,02E-02 |
| R-HSA-399954 | Sema3A PAK dependent Axon repulsion | LIMK1 | LIM domain kinase 1 | 3984 | down | 1,44 | 6,49E-03 |
| R-HSA-162658 | Golgi Cisternae Pericentriolar Stack Reorganization | RAB1B | RAB1B, member RAS oncogene family | 81876 | down | 1,27 | 4,09E-04 |
| R-HSA-162658 | Golgi Cisternae Pericentriolar Stack Reorganization | BLZF1 | basic leucine zipper nuclear factor 1 | 8548 | up | 1,43 | 2,30E-02 |
| R-HSA-162658 | Golgi Cisternae Pericentriolar Stack Reorganization | GOLGA2 | golgin A2 | 2801 | up | 1,85 | 1,27E-02 |
| R-HSA-162658 | Golgi Cisternae Pericentriolar Stack Reorganization | USO1 | USO1 vesicle transport factor | 8615 | up | 1,77 | 6,10E-03 |
| R-HSA-5213460 | RIPK1-mediated regulated necrosis | FADD | Fas associated via death domain | 8772 | down | 1,31 | 8,90E-03 |
| R-HSA-5213460 | RIPK1-mediated regulated necrosis | TNFRSF10B | TNF receptor superfamily member 10b | 8795 | up | 2,77 | 1,99E-03 |
| R-HSA-5213460 | RIPK1-mediated regulated necrosis | TRAF2 | TNF receptor associated factor 2 | 7186 | down | 1,47 | 2,65E-02 |
| R-HSA-5213460 | RIPK1-mediated regulated necrosis | BIRC2 | baculoviral IAP repeat containing 2 | 329 | up | 1,38 | 1,86E-02 |
| R-HSA-4615885 | SUMOylation of DNA replication proteins | TOP1 | DNA topoisomerase I | 7150 | up | 1,51 | 1,94E-02 |
| R-HSA-4615885 | SUMOylation of DNA replication proteins | SUMO3 | small ubiquitin-like modifier 3 | 6612 | down | 1,24 | 4,67E-02 |
| R-HSA-4615885 | SUMOylation of DNA replication proteins | INCENP | inner centromere protein | 3619 | down | 1,78 | 2,08E-02 |
| R-HSA-4615885 | SUMOylation of DNA replication proteins | NUP58 | nucleoporin 58 | 9818 | up | 1,61 | 1,28E-02 |
| R-HSA-4615885 | SUMOylation of DNA replication proteins | RANGAP1 | Ran GTPase activating protein 1 | 5905 | down | 1,48 | 1,41E-02 |
| R-HSA-4615885 | SUMOylation of DNA replication proteins | NUP62 | nucleoporin 62 | 23636 | down | 1,52 | 4,39E-03 |
| R-HSA-4615885 | SUMOylation of DNA replication proteins | PCNA | proliferating cell nuclear antigen | 5111 | down | 2,14 | 4,63E-02 |
| R-HSA-5545619 | XAV939 inhibits tankyrase, stabilizing AXIN | TNKS2 | tankyrase 2 | 80351 | up | 1,68 | 3,02E-03 |
| R-HSA-5545619 | XAV939 inhibits tankyrase, stabilizing AXIN | TNKS | tankyrase | 8658 | up | 1,42 | 8,31E-03 |
| R-HSA-111453 | BH3-only proteins associate with and inactivate anti-a | BBC3 | BCL2 binding component 3 | 27113 | up | 2,09 | 3,17E-02 |
| R-HSA-111453 | BH3-only proteins associate with and inactivate anti-a | BID | BH3 interacting domain death agonist | 637 | down | 1,40 | 2,42E-02 |
| R-HSA-111453 | BH3-only proteins associate with and inactivate anti-a | PMAIP1 | phorbol-12-myristate-13-acetate-induced | 5366 | up | 2,01 | 4,52E-02 |
| R-HSA-163680 | AMPK inhibits chREBP transcriptional activation activi | PRKAG2 | protein kinase AMP-activated non-catal | 51422 | up | 1,28 | 4,57E-02 |
| R-HSA-163680 | AMPK inhibits chREBP transcriptional activation activi | STK11 | serine/threonine kinase 11 | 6794 | down | 1,25 | 3,05E-02 |
| R-HSA-163680 | AMPK inhibits chREBP transcriptional activation activi | PRKAB2 | protein kinase AMP-activated non-catal | 5565 | up | 1,83 | 1,27E-02 |
| R-HSA-110362 | POLB-Dependent Long Patch Base Excision Repair | POLB | DNA polymerase beta | 5423 | up | 1,25 | 3,75E-02 |
| R-HSA-110362 | POLB-Dependent Long Patch Base Excision Repair | FEN1 | flap structure-specific endonuclease 1 | 2237 | down | 2,04 | 1,60E-02 |
| R-HSA-110362 | POLB-Dependent Long Patch Base Excision Repair | PARP1 | poly(ADP-ribose) polymerase 1 | 142 | down | 1,50 | 3,92E-02 |
| R-HSA-6787450 | tRNA modification in the mitochondrion | MTO1 | mitochondrial tRNA translation optimizat | 25821 | up | 1,47 | 2,12E-02 |
| R-HSA-6787450 | tRNA modification in the mitochondrion | GTPBP3 | GTP binding protein 3, mitochondrial | 84705 | down | 1,48 | 4,00E-02 |
| R-HSA-6787450 | tRNA modification in the mitochondrion | TRMU | tRNA 5-methylaminomethyl-2-thiouridyla | 55687 | down | 1,54 | 1,56E-02 |
| R-HSA-3371599 | Defective HLCS causes multiple carboxylase deficienc | MCCC1 | methylcrotonoyl-CoA carboxylase 1 | 56922 | up | 1,28 | 4,04E-02 |
| R-HSA-3371599 | Defective HLCS causes multiple carboxylase deficienc | PCCA | propionyl-CoA carboxylase alpha subun | 5095 | up | 1,37 | 2,03E-02 |
| R-HSA-3371599 | Defective HLCS causes multiple carboxylase deficienc | ACACA | acetyl-CoA carboxylase alpha | 31 | down | 1,40 | 2,54E-02 |
| R-HSA-1433557 | Signaling by SCF-KIT | FER | FER tyrosine kinase | 2241 | up | 1,40 | 6,10E-03 |
| R-HSA-1433557 | Signaling by SCF-KIT | GRB10 | growth factor receptor bound protein 10 | 2887 | up | 1,86 | 2,07E-02 |
| R-HSA-1433557 | Signaling by SCF-KIT | HRAS | HRas proto-oncogene, GTPase | 3265 | down | 1,31 | 2,19E-03 |
| R-HSA-1433557 | Signaling by SCF-KIT | LYN | LYN proto-oncogene, Src family tyrosine | 4067 | down | 1,36 | 2,34E-02 |
| R-HSA-1433557 | Signaling by SCF-KIT | YES1 | YES proto-oncogene 1, Src family tyros | 7525 | up | 1,30 | 2,08E-02 |
| R-HSA-1433557 | Signaling by SCF-KIT | PIK3R1 | phosphoinositide-3-kinase regulatory su | 5295 | down | 2,04 | 8,40E-03 |
| R-HSA-1433557 | Signaling by SCF-KIT | PIK3CA | phosphatidylinositol-4,5-bisphosphate 3 | 5290 | up | 1,33 | 3,17E-02 |
| R-HSA-5628897 | TP53 Regulates Metabolic Genes | RPTOR | regulatory associated protein of MTOR c | 57521 | down | 1,20 | 1,16E-02 |
| R-HSA-5628897 | TP53 Regulates Metabolic Genes | LRPPRC | leucine rich pentatricopeptide repeat co | 10128 | up | 1,30 | 4,03E-03 |
| R-HSA-5628897 | TP53 Regulates Metabolic Genes | RRAGC | Ras related GTP binding C | 64121 | up | 1,78 | 6,10E-03 |

| | | | | | | | |
|---|---|---|---|---|---|---|---|
| R-HSA-5628897 | TP53 Regulates Metabolic Genes | PRKAG2 | protein kinase AMP-activated non-cataly | 51422 | up | 1,28 | 4,57E-02 |
| R-HSA-5628897 | TP53 Regulates Metabolic Genes | AKT2 | AKT serine/threonine kinase 2 | 208 | down | 1,28 | 6,33E-03 |
| R-HSA-5628897 | TP53 Regulates Metabolic Genes | AKT1 | AKT serine/threonine kinase 1 | 207 | down | 1,37 | 1,42E-03 |
| R-HSA-5628897 | TP53 Regulates Metabolic Genes | PRKAB2 | protein kinase AMP-activated non-cataly | 5565 | up | 1,83 | 1,27E-02 |
| R-HSA-5628897 | TP53 Regulates Metabolic Genes | PTEN | phosphatase and tensin homolog | 5728 | up | 1,33 | 4,72E-02 |
| R-HSA-5628897 | TP53 Regulates Metabolic Genes | AGO3 | argonaute 3, RISC catalytic component | 192669 | up | 1,36 | 1,80E-02 |
| R-HSA-5628897 | TP53 Regulates Metabolic Genes | PRDX2 | peroxiredoxin 2 | 7001 | down | 1,26 | 3,14E-02 |
| R-HSA-5628897 | TP53 Regulates Metabolic Genes | SESN2 | sestrin 2 | 83667 | up | 2,90 | 1,62E-02 |
| R-HSA-5628897 | TP53 Regulates Metabolic Genes | GPI | glucose-6-phosphate isomerase | 2821 | down | 1,24 | 1,55E-02 |
| R-HSA-5628897 | TP53 Regulates Metabolic Genes | YWHAH | tyrosine 3-monooxygenase/tryptophan 5 | 7533 | down | 1,33 | 4,79E-02 |
| R-HSA-5205685 | Pink/Parkin Mediated Mitophagy | ATG12 | autophagy related 12 | 9140 | up | 1,35 | 4,10E-02 |
| R-HSA-5205685 | Pink/Parkin Mediated Mitophagy | MAP1LC3B | microtubule associated protein 1 light ch | 81631 | up | 2,92 | 2,24E-03 |
| R-HSA-5205685 | Pink/Parkin Mediated Mitophagy | TOMM40 | translocase of outer mitochondrial memb | 10452 | down | 1,29 | 4,09E-04 |
| R-HSA-5205685 | Pink/Parkin Mediated Mitophagy | TOMM70 | translocase of outer mitochondrial memb | 9868 | up | 1,24 | 3,74E-02 |
| R-HSA-5205685 | Pink/Parkin Mediated Mitophagy | SQSTM1 | sequestosome 1 | 8878 | up | 1,88 | 2,42E-02 |
| R-HSA-389513 | CTLA4 inhibitory signaling | AKT1 | AKT serine/threonine kinase 1 | 207 | down | 1,37 | 1,42E-03 |
| R-HSA-389513 | CTLA4 inhibitory signaling | AKT2 | AKT serine/threonine kinase 2 | 208 | down | 1,28 | 6,33E-03 |
| R-HSA-389513 | CTLA4 inhibitory signaling | YES1 | YES proto-oncogene 1, Src family tyros | 7525 | up | 1,30 | 2,08E-02 |
| R-HSA-389513 | CTLA4 inhibitory signaling | LYN | LYN proto-oncogene, Src family tyrosine | 4067 | down | 1,36 | 2,34E-02 |
| R-HSA-389513 | CTLA4 inhibitory signaling | PPP2CB | protein phosphatase 2 catalytic subunit | 5516 | up | 1,30 | 4,79E-02 |
| R-HSA-2122948 | Activated NOTCH1 Transmits Signal to the Nucleus | ITCH | itchy E3 ubiquitin protein ligase | 83737 | up | 1,51 | 6,64E-03 |
| R-HSA-2122948 | Activated NOTCH1 Transmits Signal to the Nucleus | ARRB1 | arrestin beta 1 | 408 | down | 1,79 | 2,42E-02 |
| R-HSA-2122948 | Activated NOTCH1 Transmits Signal to the Nucleus | ADAM17 | ADAM metallopeptidase domain 17 | 6868 | up | 2,59 | 1,60E-03 |
| R-HSA-2122948 | Activated NOTCH1 Transmits Signal to the Nucleus | JAG1 | jagged 1 | 182 | up | 1,84 | 3,27E-02 |
| R-HSA-2122948 | Activated NOTCH1 Transmits Signal to the Nucleus | NOTCH1 | notch 1 | 4851 | down | 1,62 | 2,46E-02 |
| R-HSA-2122948 | Activated NOTCH1 Transmits Signal to the Nucleus | DTX4 | deltex E3 ubiquitin ligase 4 | 23220 | down | 1,52 | 4,63E-02 |
| R-HSA-375170 | CDO in myogenesis | TCF12 | transcription factor 12 | 6938 | up | 1,34 | 4,98E-02 |
| R-HSA-375170 | CDO in myogenesis | MEF2D | myocyte enhancer factor 2D | 4209 | up | 1,90 | 4,04E-02 |
| R-HSA-375170 | CDO in myogenesis | MAPK12 | mitogen-activated protein kinase 12 | 6300 | down | 1,37 | 3,32E-03 |
| R-HSA-375170 | CDO in myogenesis | SPAG9 | sperm associated antigen 9 | 9043 | up | 1,62 | 6,49E-03 |
| R-HSA-375170 | CDO in myogenesis | BNIP2 | BCL2 interacting protein 2 | 663 | up | 1,31 | 3,45E-02 |
| R-HSA-375170 | CDO in myogenesis | MEF2A | myocyte enhancer factor 2A | 4205 | up | 1,55 | 1,41E-02 |
| R-HSA-5357956 | TNFR1-induced NFkappaB signaling pathway | TAB3 | TGF-beta activated kinase 1 and MAP3 | 257397 | up | 1,40 | 4,94E-02 |
| R-HSA-5357956 | TNFR1-induced NFkappaB signaling pathway | BIRC2 | baculoviral IAP repeat containing 2 | 329 | up | 1,38 | 1,86E-02 |
| R-HSA-5357956 | TNFR1-induced NFkappaB signaling pathway | TRAF2 | TNF receptor associated factor 2 | 7186 | down | 1,47 | 2,65E-02 |
| R-HSA-5357956 | TNFR1-induced NFkappaB signaling pathway | TAB1 | TGF-beta activated kinase 1 (MAP3K7) | 10454 | down | 1,29 | 1,94E-02 |
| R-HSA-5357956 | TNFR1-induced NFkappaB signaling pathway | CYLD | CYLD lysine 63 deubiquitinase | 1540 | up | 1,34 | 4,86E-02 |
| R-HSA-5357956 | TNFR1-induced NFkappaB signaling pathway | OTUD7B | OTU deubiquitinase 7B | 56957 | up | 1,27 | 3,81E-02 |
| R-HSA-2559585 | Oncogene Induced Senescence | ETS1 | ETS proto-oncogene 1, transcription fac | 2113 | up | 2,07 | 1,22E-02 |
| R-HSA-2559585 | Oncogene Induced Senescence | CDKN2B | cyclin dependent kinase inhibitor 2B | 1030 | up | 2,40 | 3,90E-02 |
| R-HSA-2559585 | Oncogene Induced Senescence | E2F1 | E2F transcription factor 1 | 1869 | down | 1,64 | 3,25E-02 |
| R-HSA-2559585 | Oncogene Induced Senescence | TFDP1 | transcription factor Dp-1 | 7027 | down | 1,77 | 1,08E-02 |
| R-HSA-2559585 | Oncogene Induced Senescence | ETS2 | ETS proto-oncogene 2, transcription fac | 2114 | down | 1,49 | 4,28E-02 |
| R-HSA-2559585 | Oncogene Induced Senescence | AGO3 | argonaute 3, RISC catalytic component | 192669 | up | 1,36 | 1,80E-02 |
| R-HSA-674695 | RNA Polymerase II Pre-transcription Events | CDK7 | cyclin dependent kinase 7 | 1022 | up | 1,37 | 4,38E-02 |
| R-HSA-674695 | RNA Polymerase II Pre-transcription Events | CDC73 | cell division cycle 73 | 79577 | up | 1,37 | 4,41E-02 |
| R-HSA-674695 | RNA Polymerase II Pre-transcription Events | ERCC2 | ERCC excision repair 2, TFIIH core com | 2068 | down | 1,29 | 2,20E-02 |
| R-HSA-674695 | RNA Polymerase II Pre-transcription Events | ERCC3 | ERCC excision repair 3, TFIIH core com | 2071 | down | 1,20 | 2,37E-02 |
| R-HSA-674695 | RNA Polymerase II Pre-transcription Events | POLR2B | RNA polymerase II subunit B | 5431 | down | 1,23 | 3,81E-02 |
| R-HSA-674695 | RNA Polymerase II Pre-transcription Events | POLR2L | RNA polymerase II subunit L | 5441 | down | 1,52 | 3,58E-03 |
| R-HSA-674695 | RNA Polymerase II Pre-transcription Events | POLR2F | RNA polymerase II subunit F | 5435 | down | 1,41 | 3,22E-02 |
| R-HSA-674695 | RNA Polymerase II Pre-transcription Events | GTF2H1 | general transcription factor IIH subunit 1 | 2965 | up | 1,33 | 1,85E-02 |
| R-HSA-674695 | RNA Polymerase II Pre-transcription Events | TCEA1 | transcription elongation factor A1 | 6917 | up | 1,76 | 1,81E-02 |
| R-HSA-674695 | RNA Polymerase II Pre-transcription Events | POLR2E | RNA polymerase II subunit E | 5434 | down | 1,31 | 3,18E-03 |
| R-HSA-674695 | RNA Polymerase II Pre-transcription Events | TAF2 | TATA-box binding protein associated fa | 6873 | up | 1,29 | 5,94E-03 |
| R-HSA-674695 | RNA Polymerase II Pre-transcription Events | CCNT1 | cyclin T1 | 904 | up | 1,34 | 3,05E-02 |
| R-HSA-5576892 | Phase 0 - rapid depolarisation | CALM3 | calmodulin 3 | 808 | down | 1,43 | 2,23E-02 |
| R-HSA-5576892 | Phase 0 - rapid depolarisation | CALM2 | calmodulin 2 | 805 | down | 1,43 | 2,23E-02 |
| R-HSA-5576892 | Phase 0 - rapid depolarisation | SCN5A | sodium voltage-gated channel alpha su | 6331 | down | 15,38 | 3,09E-02 |
| R-HSA-5576892 | Phase 0 - rapid depolarisation | CACNB3 | calcium voltage-gated channel auxiliary | 784 | down | 1,19 | 3,30E-02 |
| R-HSA-5576892 | Phase 0 - rapid depolarisation | CAMK2D | calcium/calmodulin dependent protein k | 817 | up | 1,29 | 1,49E-02 |
| R-HSA-5576892 | Phase 0 - rapid depolarisation | CALM1 | calmodulin 1 | 801 | down | 1,43 | 2,23E-02 |
| R-HSA-450604 | KSRP (KHSRP) binds and destabilizes mRNA | EXOSC6 | exosome component 6 | 118460 | down | 1,50 | 1,04E-02 |
| R-HSA-450604 | KSRP (KHSRP) binds and destabilizes mRNA | AKT1 | AKT serine/threonine kinase 1 | 207 | down | 1,37 | 1,42E-03 |
| R-HSA-450604 | KSRP (KHSRP) binds and destabilizes mRNA | EXOSC4 | exosome component 4 | 54512 | down | 1,51 | 2,76E-03 |
| R-HSA-432722 | Golgi Associated Vesicle Biogenesis | SNX9 | sorting nexin 9 | 51429 | up | 1,42 | 4,22E-02 |
| R-HSA-432722 | Golgi Associated Vesicle Biogenesis | DNAJC6 | DnaJ heat shock protein family (Hsp40) | 9829 | up | 1,87 | 1,56E-02 |
| R-HSA-432722 | Golgi Associated Vesicle Biogenesis | YIPF6 | Yip1 domain family member 6 | 286451 | up | 1,44 | 2,29E-02 |
| R-HSA-432722 | Golgi Associated Vesicle Biogenesis | PIK3C2A | phosphatidylinositol-4-phosphate 3-kina | 5286 | up | 1,30 | 1,79E-02 |
| R-HSA-432722 | Golgi Associated Vesicle Biogenesis | ARRB1 | arrestin beta 1 | 408 | down | 1,79 | 2,42E-02 |
| R-HSA-432722 | Golgi Associated Vesicle Biogenesis | VAMP2 | vesicle associated membrane protein 2 | 6844 | up | 1,50 | 4,82E-02 |
| R-HSA-432722 | Golgi Associated Vesicle Biogenesis | NECAP1 | NECAP endocytosis associated 1 | 25977 | up | 1,50 | 1,94E-02 |
| R-HSA-432722 | Golgi Associated Vesicle Biogenesis | AP1M1 | adaptor related protein complex 1 mu 1 | 8907 | down | 1,24 | 1,30E-02 |
| R-HSA-432722 | Golgi Associated Vesicle Biogenesis | PICALM | phosphatidylinositol binding clathrin ass | 8301 | up | 1,34 | 3,22E-02 |
| R-HSA-70171 | Glycolysis | TPI1 | triosephosphate isomerase 1 | 7167 | down | 1,28 | 3,58E-02 |
| R-HSA-70171 | Glycolysis | PGAM1 | phosphoglycerate mutase 1 | 5223 | down | 1,62 | 5,67E-03 |
| R-HSA-70171 | Glycolysis | GPI | glucose-6-phosphate isomerase | 2821 | down | 1,24 | 1,55E-02 |
| R-HSA-70171 | Glycolysis | PFKFB2 | 6-phosphofructo-2-kinase/fructose-2,6-b | 5208 | up | 1,39 | 4,32E-02 |
| R-HSA-70171 | Glycolysis | PKM | pyruvate kinase M1/2 | 5315 | down | 1,35 | 4,16E-02 |
| R-HSA-70171 | Glycolysis | PPP2CB | protein phosphatase 2 catalytic subunit | 5516 | up | 1,30 | 4,79E-02 |
| R-HSA-112411 | MAPK1 (ERK2) activation | IL6R | interleukin 6 receptor | 3570 | up | 2,48 | 3,34E-02 |
| R-HSA-112411 | MAPK1 (ERK2) activation | IL6ST | interleukin 6 signal transducer | 3572 | up | 1,52 | 1,96E-02 |
| R-HSA-112411 | MAPK1 (ERK2) activation | JAK1 | Janus kinase 1 | 3716 | up | 1,53 | 2,13E-02 |
| R-HSA-3878781 | Glycogen storage disease type IV (GBE1) | GYS1 | glycogen synthase 1 | 2997 | down | 1,32 | 2,63E-02 |
| R-HSA-3878781 | Glycogen storage disease type IV (GBE1) | GYG2 | glycogenin 2 | 8908 | down | 1,52 | 4,53E-02 |
| R-HSA-1614517 | Sulfide oxidation to sulfate | SLC25A10 | solute carrier family 25 member 10 | 1468 | down | 1,83 | 5,12E-03 |
| R-HSA-1614517 | Sulfide oxidation to sulfate | SUOX | sulfite oxidase | 6821 | down | 1,44 | 3,35E-02 |
| R-HSA-426048 | Arachidonate production from DAG | DAGLA | diacylglycerol lipase alpha | 747 | down | 1,27 | 4,46E-02 |
| R-HSA-426048 | Arachidonate production from DAG | ABHD6 | abhydrolase domain containing 6 | 57406 | down | 1,49 | 2,16E-02 |
| R-HSA-2122947 | NOTCH1 Intracellular Domain Regulates Transcription | TBL1X | transducin beta like 1 X-linked | 6907 | down | 1,20 | 2,84E-02 |
| R-HSA-2122947 | NOTCH1 Intracellular Domain Regulates Transcription | NOTCH1 | notch 1 | 4851 | down | 1,62 | 2,46E-02 |
| R-HSA-2122947 | NOTCH1 Intracellular Domain Regulates Transcription | MAMLD1 | mastermind like domain containing 1 | 10046 | down | 3,36 | 4,56E-02 |
| R-HSA-2122947 | NOTCH1 Intracellular Domain Regulates Transcription | HIF1A | hypoxia inducible factor 1 alpha subunit | 3091 | up | 1,23 | 4,66E-02 |
| R-HSA-2122947 | NOTCH1 Intracellular Domain Regulates Transcription | HDAC4 | histone deacetylase 4 | 9759 | down | 1,38 | 1,08E-02 |
| R-HSA-2122947 | NOTCH1 Intracellular Domain Regulates Transcription | KAT2A | lysine acetyltransferase 2A | 2648 | down | 1,39 | 1,20E-02 |
| R-HSA-2122947 | NOTCH1 Intracellular Domain Regulates Transcription | HEY1 | hes related family bHLH transcription fac | 23462 | up | 3,28 | 1,40E-02 |
| R-HSA-2122947 | NOTCH1 Intracellular Domain Regulates Transcription | HDAC9 | histone deacetylase 9 | 9734 | up | 2,67 | 1,19E-02 |
| R-HSA-983189 | Kinesins | KIF5B | kinesin family member 5B | 3799 | up | 1,40 | 7,18E-03 |
| R-HSA-983189 | Kinesins | KIF1C | kinesin family member 1C | 10749 | down | 1,20 | 2,45E-02 |
| R-HSA-983189 | Kinesins | TUBA1B | tubulin alpha 1b | 10376 | down | 1,78 | 2,30E-02 |
| R-HSA-983189 | Kinesins | KIF3C | kinesin family member 3C | 3797 | down | 1,58 | 4,57E-02 |
| R-HSA-983189 | Kinesins | KIFC1 | kinesin family member C1 | 3833 | down | 1,95 | 2,60E-02 |
| R-HSA-983189 | Kinesins | TUBA4A | tubulin alpha 4a | 7277 | down | 1,63 | 2,97E-02 |
| R-HSA-983189 | Kinesins | KIF18B | kinesin family member 18B | 146909 | down | 1,94 | 3,01E-02 |
| R-HSA-983189 | Kinesins | KIF16B | kinesin family member 16B | 55614 | up | 1,64 | 5,27E-03 |
| R-HSA-5654704 | SHC-mediated cascade:FGFR3 | HRAS | HRas proto-oncogene, GTPase | 3265 | down | 1,31 | 2,19E-03 |
| R-HSA-5654704 | SHC-mediated cascade:FGFR3 | FGF2 | fibroblast growth factor 2 | 2247 | down | 5,35 | 3,09E-02 |
| R-HSA-5654704 | SHC-mediated cascade:FGFR3 | FGFR3 | fibroblast growth factor receptor 3 | 2261 | down | 1,70 | 4,60E-02 |
| R-HSA-5654710 | PI-3K cascade:FGFR3 | FGF2 | fibroblast growth factor 2 | 2247 | down | 5,35 | 3,09E-02 |
| R-HSA-5654710 | PI-3K cascade:FGFR3 | PIK3R1 | phosphoinositide-3-kinase regulatory su | 5295 | down | 2,04 | 8,40E-03 |
| R-HSA-5654710 | PI-3K cascade:FGFR3 | FGFR3 | fibroblast growth factor receptor 3 | 2261 | down | 1,70 | 4,60E-02 |
| R-HSA-5654710 | PI-3K cascade:FGFR3 | PIK3CA | phosphatidylinositol-4,5-bisphosphate 3 | 5290 | up | 1,33 | 3,17E-02 |
| R-HSA-1912420 | Pre-NOTCH Processing in Golgi | SEL1L | SEL1L ERAD E3 ligase adaptor subunit | 6400 | up | 2,76 | 4,03E-03 |
| R-HSA-1912420 | Pre-NOTCH Processing in Golgi | LFNG | LFNG O-fucosylpeptide 3-beta-N-acetylg | 3955 | down | 2,26 | 1,16E-02 |
| R-HSA-1912420 | Pre-NOTCH Processing in Golgi | ATP2A3 | ATPase sarcoplasmic/endoplasmic retic | 489 | down | 1,62 | 4,34E-02 |
| R-HSA-1912420 | Pre-NOTCH Processing in Golgi | NOTCH1 | notch 1 | 4851 | down | 1,62 | 2,46E-02 |
| R-HSA-373753 | Nephrin family interactions | PIK3CA | phosphatidylinositol-4,5-bisphosphate 3 | 5290 | up | 1,33 | 3,17E-02 |
| R-HSA-373753 | Nephrin family interactions | PIK3R1 | phosphoinositide-3-kinase regulatory su | 5295 | down | 2,04 | 8,40E-03 |
| R-HSA-373753 | Nephrin family interactions | NCK1 | NCK adaptor protein 1 | 4690 | up | 1,48 | 1,42E-02 |
| R-HSA-373753 | Nephrin family interactions | SPTBN1 | spectrin beta, non-erythrocytic 1 | 6711 | up | 1,42 | 1,66E-02 |
| R-HSA-373753 | Nephrin family interactions | CD2AP | CD2 associated protein | 23607 | up | 1,26 | 4,32E-02 |
| R-HSA-2173795 | Downregulation of SMAD2/3:SMAD4 transcriptional a | SMAD2 | SMAD family member 2 | 4087 | up | 1,29 | 2,19E-02 |
| R-HSA-2173795 | Downregulation of SMAD2/3:SMAD4 transcriptional a | SMURF2 | SMAD specific E3 ubiquitin protein ligas | 64750 | up | 1,49 | 4,16E-02 |
| R-HSA-2173795 | Downregulation of SMAD2/3:SMAD4 transcriptional a | PARP1 | poly(ADP-ribose) polymerase 1 | 142 | down | 1,50 | 3,92E-02 |
| R-HSA-2173795 | Downregulation of SMAD2/3:SMAD4 transcriptional a | SKIL | SKI like proto-oncogene | 6498 | up | 1,70 | 1,91E-02 |
| R-HSA-2173795 | Downregulation of SMAD2/3:SMAD4 transcriptional a | TGIF2 | TGFB induced factor homeobox 2 | 60436 | down | 1,76 | 2,54E-02 |
| R-HSA-5685942 | HDR through Homologous Recombination (HRR) | POLD1 | DNA polymerase delta 1, catalytic subur | 5424 | down | 1,59 | 4,62E-02 |
| R-HSA-5685942 | HDR through Homologous Recombination (HRR) | NBN | nibrin | 4683 | up | 1,25 | 3,15E-02 |
| R-HSA-5685942 | HDR through Homologous Recombination (HRR) | POLE | DNA polymerase epsilon, catalytic subu | 5426 | down | 1,55 | 2,67E-02 |
| R-HSA-5685942 | HDR through Homologous Recombination (HRR) | RPA1 | replication protein A1 | 6117 | down | 1,41 | 4,93E-02 |
| R-HSA-5685942 | HDR through Homologous Recombination (HRR) | RFC1 | replication factor C subunit 1 | 5981 | up | 1,15 | 2,18E-02 |
| R-HSA-5685942 | HDR through Homologous Recombination (HRR) | PCNA | proliferating cell nuclear antigen | 5111 | down | 2,14 | 4,63E-02 |
| R-HSA-5685942 | HDR through Homologous Recombination (HRR) | POLD2 | DNA polymerase delta 2, accessory sub | 5425 | down | 1,30 | 1,92E-02 |
| R-HSA-933541 | TRAF6 mediated IRF7 activation | MYD88 | myeloid differentiation primary response | 4615 | down | 1,55 | 1,85E-02 |
| R-HSA-933541 | TRAF6 mediated IRF7 activation | IRF3 | interferon regulatory factor 3 | 3661 | down | 1,33 | 2,58E-03 |
| R-HSA-933541 | TRAF6 mediated IRF7 activation | IKBKE | inhibitor of nuclear factor kappa B kinas | 9641 | down | 1,67 | 1,00E-02 |
| R-HSA-933541 | TRAF6 mediated IRF7 activation | TRAF2 | TNF receptor associated factor 2 | 7186 | down | 1,47 | 2,65E-02 |
| R-HSA-1632852 | Macroautophagy | PRKAB2 | protein kinase AMP-activated non-cataly | 5565 | up | 1,83 | 1,27E-02 |
| R-HSA-1632852 | Macroautophagy | ATG12 | autophagy related 12 | 9140 | up | 1,35 | 4,10E-02 |
| R-HSA-1632852 | Macroautophagy | GABARAPL1 | GABA type A receptor associated prote | 23710 | up | 2,62 | 4,12E-03 |
| R-HSA-1632852 | Macroautophagy | RRAGC | Ras related GTP binding C | 64121 | up | 1,78 | 6,10E-03 |
| R-HSA-1632852 | Macroautophagy | RPTOR | regulatory associated protein of MTOR c | 57521 | down | 1,20 | 1,16E-02 |
| R-HSA-1632852 | Macroautophagy | WIPI1 | WD repeat domain, phosphoinositide int | 55062 | up | 3,38 | 2,86E-03 |
| R-HSA-1632852 | Macroautophagy | MAP1LC3B | microtubule associated protein 1 light ch | 81631 | up | 2,92 | 2,24E-03 |
| R-HSA-1632852 | Macroautophagy | PRKAG2 | protein kinase AMP-activated non-cataly | 51422 | up | 1,28 | 4,57E-02 |
| R-HSA-1632852 | Macroautophagy | RB1CC1 | RB1 inducible coiled-coil 1 | 9821 | up | 1,55 | 2,96E-02 |
| R-HSA-437239 | Recycling pathway of L1 | L1CAM | L1 cell adhesion molecule | 3897 | down | 2,86 | 3,09E-02 |
| R-HSA-437239 | Recycling pathway of L1 | ACTG1 | actin gamma 1 | 71 | up | 1,58 | 1,66E-02 |
| R-HSA-437239 | Recycling pathway of L1 | RPS6KA5 | ribosomal protein S6 kinase A5 | 9252 | up | 2,11 | 5,94E-03 |
| R-HSA-437239 | Recycling pathway of L1 | TUBA1B | tubulin alpha 1b | 10376 | down | 1,78 | 2,30E-02 |
| R-HSA-437239 | Recycling pathway of L1 | DPYSL2 | dihydropyrimidinase like 2 | 1808 | down | 1,77 | 2,19E-02 |
| R-HSA-437239 | Recycling pathway of L1 | RPS6KA4 | ribosomal protein S6 kinase A4 | 8986 | down | 1,33 | 5,67E-03 |
| R-HSA-437239 | Recycling pathway of L1 | TUBA4A | tubulin alpha 4a | 7277 | down | 1,63 | 2,97E-02 |
| R-HSA-1236382 | Constitutive Signaling by Ligand-Responsive EGFR C | PIK3CA | phosphatidylinositol-4,5-bisphosphate 3 | 5290 | up | 1,33 | 3,17E-02 |
| R-HSA-1236382 | Constitutive Signaling by Ligand-Responsive EGFR C | PLCG1 | phospholipase C gamma 1 | 5335 | down | 1,51 | 2,85E-02 |
| R-HSA-1236382 | Constitutive Signaling by Ligand-Responsive EGFR C | PIK3R1 | phosphoinositide-3-kinase regulatory su | 5295 | down | 2,04 | 8,40E-03 |
| R-HSA-1236382 | Constitutive Signaling by Ligand-Responsive EGFR C | HRAS | HRas proto-oncogene, GTPase | 3265 | down | 1,31 | 2,19E-03 |
| R-HSA-936964 | Activation of IRF3/IRF7 mediated by TBK1/IKK epsilo | IKBKE | inhibitor of nuclear factor kappa B kinas | 9641 | down | 1,67 | 1,00E-02 |
| R-HSA-936964 | Activation of IRF3/IRF7 mediated by TBK1/IKK epsilo | IRF3 | interferon regulatory factor 3 | 3661 | down | 1,33 | 2,58E-03 |
| R-HSA-936964 | Activation of IRF3/IRF7 mediated by TBK1/IKK epsilo | TRAF3 | TNF receptor associated factor 3 | 7187 | down | 1,40 | 1,36E-02 |
| R-HSA-1963642 | PI3K events in ERBB2 signaling | HBEGF | heparin binding EGF like growth factor | 1839 | up | 3,67 | 5,67E-03 |
| R-HSA-1963642 | PI3K events in ERBB2 signaling | PIK3R1 | phosphoinositide-3-kinase regulatory su | 5295 | down | 2,04 | 8,40E-03 |
| R-HSA-1963642 | PI3K events in ERBB2 signaling | PIK3CA | phosphatidylinositol-4,5-bisphosphate 3 | 5290 | up | 1,33 | 3,17E-02 |
| R-HSA-1963642 | PI3K events in ERBB2 signaling | ERBB2 | erb-b2 receptor tyrosine kinase 2 | 2064 | down | 1,47 | 1,93E-02 |
| R-HSA-196757 | Metabolism of folate and pterines | SLC46A1 | solute carrier family 46 member 1 | 113235 | down | 1,66 | 4,58E-02 |
| R-HSA-196757 | Metabolism of folate and pterines | MTHFD2 | methylenetetrahydrofolate dehydrogena | 10797 | up | 2,09 | 9,37E-03 |
| R-HSA-196757 | Metabolism of folate and pterines | MTHFD1L | methylenetetrahydrofolate dehydrogena | 25902 | up | 1,36 | 4,51E-02 |
| R-HSA-196757 | Metabolism of folate and pterines | SLC19A1 | solute carrier family 19 member 1 | 6573 | down | 1,93 | 4,76E-03 |
| R-HSA-6787639 | GDP-fucose biosynthesis | FUOM | fucose mutarotase | 282969 | down | 1,30 | 1,85E-02 |
| R-HSA-6787639 | GDP-fucose biosynthesis | SLC35C1 | solute carrier family 35 member C1 | 55343 | down | 1,25 | 2,93E-02 |
| R-HSA-1606341 | IRF3 mediated activation of type 1 IFN | DTX4 | deltex E3 ubiquitin ligase 4 | 23220 | down | 1,52 | 4,63E-02 |
| R-HSA-1606341 | IRF3 mediated activation of type 1 IFN | IRF3 | interferon regulatory factor 3 | 3661 | down | 1,33 | 2,58E-03 |
| R-HSA-500753 | Pyrimidine biosynthesis | UMPS | uridine monophosphate synthetase | 7372 | down | 1,34 | 4,41E-02 |
| R-HSA-500753 | Pyrimidine biosynthesis | CAD | carbamoyl-phosphate synthetase 2, asp | 790 | down | 1,43 | 7,14E-03 |
| R-HSA-5173105 | O-linked glycosylation | LARGE2 | LARGE xylosyl- and glucuronyltransfera | 120071 | down | 2,23 | 3,90E-02 |
| R-HSA-5173105 | O-linked glycosylation | DAG1 | dystroglycan 1 | 1605 | down | 1,65 | 1,04E-02 |
| R-HSA-3928665 | EPH-ephrin mediated repulsion of cells | LYN | LYN proto-oncogene, Src family tyrosine | 4067 | down | 1,36 | 2,34E-02 |
| R-HSA-3928665 | EPH-ephrin mediated repulsion of cells | EFNA4 | ephrin A4 | 1945 | down | 1,43 | 4,92E-02 |
| R-HSA-3928665 | EPH-ephrin mediated repulsion of cells | EPHA2 | EPH receptor A2 | 1969 | up | 1,89 | 2,51E-02 |
| R-HSA-3928665 | EPH-ephrin mediated repulsion of cells | EPHB2 | EPH receptor B2 | 2048 | down | 1,66 | 2,52E-02 |
| R-HSA-3928665 | EPH-ephrin mediated repulsion of cells | EPHA5 | EPH receptor A5 | 2044 | down | 18,87 | 1,96E-02 |
| R-HSA-3928665 | EPH-ephrin mediated repulsion of cells | YES1 | YES proto-oncogene 1, Src family tyros | 7525 | up | 1,30 | 2,08E-02 |
| R-HSA-3928665 | EPH-ephrin mediated repulsion of cells | ACTG1 | actin gamma 1 | 71 | up | 1,58 | 1,66E-02 |
| R-HSA-3928665 | EPH-ephrin mediated repulsion of cells | EFNA3 | ephrin A3 | 1944 | down | 2,00 | 4,89E-02 |
| R-HSA-917729 | Endosomal Sorting Complex Required For Transport ( | CHMP5 | charged multivesicular body protein 5 | 51510 | up | 1,32 | 3,37E-02 |
| R-HSA-917729 | Endosomal Sorting Complex Required For Transport ( | STAM | signal transducing adaptor molecule | 8027 | up | 1,23 | 3,52E-02 |
| R-HSA-917729 | Endosomal Sorting Complex Required For Transport ( | VPS37A | VPS37A, ESCRT-I subunit | 137492 | up | 1,48 | 1,14E-02 |
| R-HSA-917729 | Endosomal Sorting Complex Required For Transport ( | STAM2 | signal transducing adaptor molecule 2 | 10254 | up | 1,41 | 6,99E-03 |
| R-HSA-917729 | Endosomal Sorting Complex Required For Transport ( | VPS37D | VPS37D, ESCRT-I subunit | 155382 | up | 2,22 | 5,83E-03 |
| R-HSA-901032 | ER Quality Control Compartment (ERQC) | SYVN1 | synoviolin 1 | 84447 | up | 2,33 | 1,93E-02 |
| R-HSA-901032 | ER Quality Control Compartment (ERQC) | EDEM1 | ER degradation enhancing alpha-mann | 9695 | up | 2,16 | 1,32E-02 |
| R-HSA-901032 | ER Quality Control Compartment (ERQC) | SEL1L | SEL1L ERAD E3 ligase adaptor subunit | 6400 | up | 2,76 | 4,03E-03 |
| R-HSA-901032 | ER Quality Control Compartment (ERQC) | DERL2 | derlin 2 | 51009 | up | 1,68 | 4,88E-02 |
| R-HSA-110056 | MAPK3 (ERK1) activation | IL6ST | interleukin 6 signal transducer | 3572 | up | 1,52 | 1,96E-02 |
| R-HSA-110056 | MAPK3 (ERK1) activation | JAK1 | Janus kinase 1 | 3716 | up | 1,53 | 2,13E-02 |
| R-HSA-110056 | MAPK3 (ERK1) activation | IL6R | interleukin 6 receptor | 3570 | up | 2,48 | 3,34E-02 |
| R-HSA-1614635 | Sulfur amino acid metabolism | AHCY | adenosylhomocysteinase | 191 | down | 1,14 | 2,15E-02 |
| R-HSA-1614635 | Sulfur amino acid metabolism | GCLC | glutamate-cysteine ligase catalytic subu | 2729 | up | 2,09 | 2,27E-02 |
| R-HSA-1614635 | Sulfur amino acid metabolism | CNDP2 | carnosine dipeptidase 2 | 55748 | down | 1,20 | 4,78E-02 |
| R-HSA-3371378 | Regulation by c-FLIP | TRAF2 | TNF receptor associated factor 2 | 7186 | down | 1,47 | 2,65E-02 |
| R-HSA-3371378 | Regulation by c-FLIP | TNFRSF10B | TNF receptor superfamily member 10b | 8795 | up | 2,77 | 1,99E-03 |
| R-HSA-3371378 | Regulation by c-FLIP | FADD | Fas associated via death domain | 8772 | down | 1,31 | 8,90E-03 |
| R-HSA-5218900 | CASP8 activity is inhibited | TRAF2 | TNF receptor associated factor 2 | 7186 | down | 1,47 | 2,65E-02 |
| R-HSA-5218900 | CASP8 activity is inhibited | FADD | Fas associated via death domain | 8772 | down | 1,31 | 8,90E-03 |
| R-HSA-5218900 | CASP8 activity is inhibited | TNFRSF10B | TNF receptor superfamily member 10b | 8795 | up | 2,77 | 1,99E-03 |
| R-HSA-2408508 | Metabolism of ingested SeMet, Sec, MeSec into H2S | CTH | cystathionine gamma-lyase | 1491 | up | 3,41 | 4,94E-03 |
| R-HSA-2408508 | Metabolism of ingested SeMet, Sec, MeSec into H2S | AHCY | adenosylhomocysteinase | 191 | down | 1,14 | 2,15E-02 |
| R-HSA-2408508 | Metabolism of ingested SeMet, Sec, MeSec into H2S | NNMT | nicotinamide N-methyltransferase | 4837 | down | 2,26 | 3,05E-02 |
| R-HSA-69416 | Dimerization of procaspase-8 | TRAF2 | TNF receptor associated factor 2 | 7186 | down | 1,47 | 2,65E-02 |
| R-HSA-69416 | Dimerization of procaspase-8 | TNFRSF10B | TNF receptor superfamily member 10b | 8795 | up | 2,77 | 1,99E-03 |
| R-HSA-69416 | Dimerization of procaspase-8 | FADD | Fas associated via death domain | 8772 | down | 1,31 | 8,90E-03 |
| R-HSA-982772 | Growth hormone receptor signaling | ADAM17 | ADAM metallopeptidase domain 17 | 6868 | up | 2,59 | 1,60E-03 |
| R-HSA-982772 | Growth hormone receptor signaling | IRS2 | insulin receptor substrate 2 | 8660 | up | 2,21 | 2,22E-02 |
| R-HSA-982772 | Growth hormone receptor signaling | LYN | LYN proto-oncogene, Src family tyrosine | 4067 | down | 1,36 | 2,34E-02 |
| R-HSA-982772 | Growth hormone receptor signaling | SH2B1 | SH2B adaptor protein 1 | 25970 | down | 1,31 | 1,81E-02 |
| R-HSA-982772 | Growth hormone receptor signaling | IRS1 | insulin receptor substrate 1 | 3667 | down | 1,89 | 6,25E-03 |
| R-HSA-3560783 | Defective B4GALT7 causes EDS, progeroid type | CSPG4 | chondroitin sulfate proteoglycan 4 | 1464 | down | 2,22 | 3,99E-02 |

| | | | | | | | |
|---|---|---|---|---|---|---|---|
| R-HSA-3560783 | Defective B4GALT7 causes EDS, progeroid type | SDC1 | syndecan 1 | 6382 | down | 2,17 | 1,61E-02 |
| R-HSA-3560783 | Defective B4GALT7 causes EDS, progeroid type | SDC3 | syndecan 3 | 9672 | down | 1,68 | 1,97E-02 |
| R-HSA-3560801 | Defective B3GAT3 causes JDSSDHD | CSPG4 | chondroitin sulfate proteoglycan 4 | 1464 | down | 2,22 | 3,99E-02 |
| R-HSA-3560801 | Defective B3GAT3 causes JDSSDHD | SDC3 | syndecan 3 | 9672 | down | 1,68 | 1,97E-02 |
| R-HSA-3560801 | Defective B3GAT3 causes JDSSDHD | SDC1 | syndecan 1 | 6382 | down | 2,17 | 1,61E-02 |
| R-HSA-4420332 | Defective B3GALT6 causes EDSP2 and SEMDJL1 | SDC3 | syndecan 3 | 9672 | down | 1,68 | 1,97E-02 |
| R-HSA-4420332 | Defective B3GALT6 causes EDSP2 and SEMDJL1 | SDC1 | syndecan 1 | 6382 | down | 2,17 | 1,61E-02 |
| R-HSA-4420332 | Defective B3GALT6 causes EDSP2 and SEMDJL1 | CSPG4 | chondroitin sulfate proteoglycan 4 | 1464 | down | 2,22 | 3,99E-02 |
| R-HSA-5654706 | FRS-mediated FGFR3 signaling | FGFR3 | fibroblast growth factor receptor 3 | 2261 | down | 1,70 | 4,60E-02 |
| R-HSA-5654706 | FRS-mediated FGFR3 signaling | FGF2 | fibroblast growth factor 2 | 2247 | down | 5,35 | 3,09E-02 |
| R-HSA-5654706 | FRS-mediated FGFR3 signaling | HRAS | HRas proto-oncogene, GTPase | 3265 | down | 1,31 | 2,19E-03 |
| R-HSA-5620922 | BBSome-mediated cargo-targeting to cilium | BBS2 | Bardet-Biedl syndrome 2 | 583 | up | 1,21 | 3,77E-02 |
| R-HSA-5620922 | BBSome-mediated cargo-targeting to cilium | LZTFL1 | leucine zipper transcription factor like 1 | 54585 | up | 1,44 | 4,05E-02 |
| R-HSA-5620922 | BBSome-mediated cargo-targeting to cilium | BBIP1 | BBSome interacting protein 1 | 92482 | up | 1,33 | 3,49E-02 |
| R-HSA-5620922 | BBSome-mediated cargo-targeting to cilium | BBS12 | Bardet-Biedl syndrome 12 | 166379 | up | 1,27 | 4,93E-02 |
| R-HSA-1059683 | Interleukin-6 signaling | IL6ST | interleukin 6 signal transducer | 3572 | up | 1,52 | 1,96E-02 |
| R-HSA-1059683 | Interleukin-6 signaling | JAK1 | Janus kinase 1 | 3716 | up | 1,53 | 2,13E-02 |
| R-HSA-1059683 | Interleukin-6 signaling | IL6R | interleukin 6 receptor | 3570 | up | 2,48 | 3,34E-02 |
| R-HSA-2132295 | MHC class II antigen presentation | CANX | calnexin | 821 | up | 1,36 | 3,98E-02 |
| R-HSA-2132295 | MHC class II antigen presentation | AP1M1 | adaptor related protein complex 1 mu 1 | 8907 | down | 1,24 | 1,30E-02 |
| R-HSA-2132295 | MHC class II antigen presentation | SEC24D | SEC24 homolog D, COPII coat complex | 9871 | up | 4,45 | 6,80E-04 |
| R-HSA-2132295 | MHC class II antigen presentation | SEC31A | SEC31 homolog A, COPII coat complex | 22872 | up | 1,53 | 3,02E-02 |
| R-HSA-2132295 | MHC class II antigen presentation | DCTN4 | dynactin subunit 4 | 51164 | up | 1,51 | 1,31E-02 |
| R-HSA-2132295 | MHC class II antigen presentation | SEC23A | Sec23 homolog A, coat complex II comp | 10484 | up | 1,46 | 2,47E-02 |
| R-HSA-2132295 | MHC class II antigen presentation | KIF5B | kinesin family member 5B | 3799 | up | 1,40 | 7,18E-03 |
| R-HSA-2132295 | MHC class II antigen presentation | OSBPL1A | oxysterol binding protein like 1A | 114876 | up | 1,40 | 2,14E-03 |
| R-HSA-2132295 | MHC class II antigen presentation | SEC24A | SEC24 homolog A, COPII coat complex | 10802 | up | 1,71 | 3,95E-03 |
| R-HSA-2132295 | MHC class II antigen presentation | DYNC1LI2 | dynein cytoplasmic 1 light intermediate c | 1783 | up | 1,21 | 2,47E-02 |
| R-HSA-2132295 | MHC class II antigen presentation | DYNC2H1 | dynein cytoplasmic 2 heavy chain 1 | 79659 | up | 1,52 | 9,51E-03 |
| R-HSA-2132295 | MHC class II antigen presentation | DYNC2LI1 | dynein cytoplasmic 2 light intermediate c | 51626 | up | 1,40 | 3,09E-02 |
| R-HSA-2132295 | MHC class II antigen presentation | TUBA1B | tubulin alpha 1b | 10376 | down | 1,78 | 2,30E-02 |
| R-HSA-2132295 | MHC class II antigen presentation | KIF3C | kinesin family member 3C | 3797 | down | 1,58 | 4,57E-02 |
| R-HSA-2132295 | MHC class II antigen presentation | TUBA4A | tubulin alpha 4a | 7277 | down | 1,63 | 2,97E-02 |
| R-HSA-2894862 | Constitutive Signaling by NOTCH1 HD+PEST Domain | ADAM17 | ADAM metallopeptidase domain 17 | 6868 | up | 2,59 | 1,60E-03 |
| R-HSA-2894862 | Constitutive Signaling by NOTCH1 HD+PEST Domain | MAMLD1 | mastermind like domain containing 1 | 10046 | down | 3,36 | 4,56E-02 |
| R-HSA-2894862 | Constitutive Signaling by NOTCH1 HD+PEST Domain | TBL1X | transducin beta like 1 X-linked | 6907 | down | 1,20 | 2,84E-02 |
| R-HSA-2894862 | Constitutive Signaling by NOTCH1 HD+PEST Domain | NOTCH1 | notch 1 | 4851 | down | 1,62 | 2,46E-02 |
| R-HSA-2894862 | Constitutive Signaling by NOTCH1 HD+PEST Domain | HDAC9 | histone deacetylase 9 | 9734 | up | 2,67 | 1,19E-02 |
| R-HSA-2894862 | Constitutive Signaling by NOTCH1 HD+PEST Domain | HEY1 | hes related family bHLH transcription fac | 23462 | up | 3,28 | 1,40E-02 |
| R-HSA-2894862 | Constitutive Signaling by NOTCH1 HD+PEST Domain | KAT2A | lysine acetyltransferase 2A | 2648 | down | 1,39 | 1,20E-02 |
| R-HSA-2894862 | Constitutive Signaling by NOTCH1 HD+PEST Domain | HDAC4 | histone deacetylase 4 | 9759 | down | 1,38 | 1,08E-02 |
| R-HSA-2894862 | Constitutive Signaling by NOTCH1 HD+PEST Domain | JAG1 | jagged 1 | 182 | up | 1,84 | 3,27E-02 |
| R-HSA-2644606 | Constitutive Signaling by NOTCH1 PEST Domain Mut | MAMLD1 | mastermind like domain containing 1 | 10046 | down | 3,36 | 4,56E-02 |
| R-HSA-2644606 | Constitutive Signaling by NOTCH1 PEST Domain Mut | ADAM17 | ADAM metallopeptidase domain 17 | 6868 | up | 2,59 | 1,60E-03 |
| R-HSA-2644606 | Constitutive Signaling by NOTCH1 PEST Domain Mut | NOTCH1 | notch 1 | 4851 | down | 1,62 | 2,46E-02 |
| R-HSA-2644606 | Constitutive Signaling by NOTCH1 PEST Domain Mut | TBL1X | transducin beta like 1 X-linked | 6907 | down | 1,20 | 2,84E-02 |
| R-HSA-2644606 | Constitutive Signaling by NOTCH1 PEST Domain Mut | HEY1 | hes related family bHLH transcription fac | 23462 | up | 3,28 | 1,40E-02 |
| R-HSA-2644606 | Constitutive Signaling by NOTCH1 PEST Domain Mut | HDAC9 | histone deacetylase 9 | 9734 | up | 2,67 | 1,19E-02 |
| R-HSA-2644606 | Constitutive Signaling by NOTCH1 PEST Domain Mut | JAG1 | jagged 1 | 182 | up | 1,84 | 3,27E-02 |
| R-HSA-2644606 | Constitutive Signaling by NOTCH1 PEST Domain Mut | HDAC4 | histone deacetylase 4 | 9759 | down | 1,38 | 1,08E-02 |
| R-HSA-2644606 | Constitutive Signaling by NOTCH1 PEST Domain Mut | KAT2A | lysine acetyltransferase 2A | 2648 | down | 1,39 | 1,20E-02 |
| R-HSA-167238 | Pausing and recovery of Tat-mediated HIV elongatio | POLR2B | RNA polymerase II subunit B | 5431 | down | 1,23 | 3,81E-02 |
| R-HSA-167238 | Pausing and recovery of Tat-mediated HIV elongatio | TCEA1 | transcription elongation factor A1 | 6917 | up | 1,76 | 1,81E-02 |
| R-HSA-167238 | Pausing and recovery of Tat-mediated HIV elongatio | POLR2F | RNA polymerase II subunit F | 5435 | down | 1,41 | 3,22E-02 |
| R-HSA-167238 | Pausing and recovery of Tat-mediated HIV elongatio | POLR2L | RNA polymerase II subunit L | 5441 | down | 1,52 | 3,58E-03 |
| R-HSA-167238 | Pausing and recovery of Tat-mediated HIV elongatio | POLR2E | RNA polymerase II subunit E | 5434 | down | 1,31 | 3,18E-03 |
| R-HSA-167238 | Pausing and recovery of Tat-mediated HIV elongatio | CCNT1 | cyclin T1 | 904 | up | 1,34 | 3,05E-02 |
| R-HSA-109704 | PI3K Cascade | IRS2 | insulin receptor substrate 2 | 8660 | up | 2,21 | 2,22E-02 |
| R-HSA-109704 | PI3K Cascade | IRS1 | insulin receptor substrate 1 | 3667 | down | 1,89 | 6,25E-03 |
| R-HSA-109704 | PI3K Cascade | PIK3CA | phosphatidylinositol-4,5-bisphosphate 3 | 5290 | up | 1,33 | 3,17E-02 |
| R-HSA-109704 | PI3K Cascade | FGFR3 | fibroblast growth factor receptor 3 | 2261 | down | 1,70 | 4,60E-02 |
| R-HSA-109704 | PI3K Cascade | PIK3R1 | phosphoinositide-3-kinase regulatory su | 5295 | down | 2,04 | 8,40E-03 |
| R-HSA-109704 | PI3K Cascade | FGF2 | fibroblast growth factor 2 | 2247 | down | 5,35 | 3,09E-02 |
| R-HSA-167243 | Tat-mediated HIV elongation arrest and recovery | CCNT1 | cyclin T1 | 904 | up | 1,34 | 3,05E-02 |
| R-HSA-167243 | Tat-mediated HIV elongation arrest and recovery | POLR2E | RNA polymerase II subunit E | 5434 | down | 1,31 | 3,18E-03 |
| R-HSA-167243 | Tat-mediated HIV elongation arrest and recovery | TCEA1 | transcription elongation factor A1 | 6917 | up | 1,76 | 1,81E-02 |
| R-HSA-167243 | Tat-mediated HIV elongation arrest and recovery | POLR2F | RNA polymerase II subunit F | 5435 | down | 1,41 | 3,22E-02 |
| R-HSA-167243 | Tat-mediated HIV elongation arrest and recovery | POLR2L | RNA polymerase II subunit L | 5441 | down | 1,52 | 3,58E-03 |
| R-HSA-167243 | Tat-mediated HIV elongation arrest and recovery | POLR2B | RNA polymerase II subunit B | 5431 | down | 1,23 | 3,81E-02 |
| R-HSA-4641263 | Regulation of FZD by ubiquitination | USP8 | ubiquitin specific peptidase 8 | 9101 | up | 1,31 | 7,82E-03 |
| R-HSA-4641263 | Regulation of FZD by ubiquitination | FZD4 | frizzled class receptor 4 | 8322 | down | 2,17 | 3,82E-02 |
| R-HSA-4641263 | Regulation of FZD by ubiquitination | LGR5 | leucine rich repeat containing G protein- | 8549 | down | 2,70 | 3,08E-02 |
| R-HSA-4641263 | Regulation of FZD by ubiquitination | ZNRF3 | zinc and ring finger 3 | 84133 | down | 1,62 | 2,51E-02 |
| R-HSA-5621575 | CD209 (DC-SIGN) signaling | RPS6KA5 | ribosomal protein S6 kinase A5 | 9252 | up | 2,11 | 5,94E-03 |
| R-HSA-5621575 | CD209 (DC-SIGN) signaling | ICAM3 | intercellular adhesion molecule 3 | 3385 | down | 1,36 | 1,58E-02 |
| R-HSA-5621575 | CD209 (DC-SIGN) signaling | HRAS | HRas proto-oncogene, GTPase | 3265 | down | 1,31 | 2,19E-03 |
| R-HSA-5621575 | CD209 (DC-SIGN) signaling | LYN | LYN proto-oncogene, Src family tyrosine | 4067 | down | 1,36 | 2,34E-02 |
| R-HSA-1227986 | Signaling by ERBB2 | HBEGF | heparin binding EGF like growth factor | 1839 | up | 3,67 | 5,67E-03 |
| R-HSA-1227986 | Signaling by ERBB2 | STUB1 | STIP1 homology and U-box containing | 10273 | down | 1,24 | 4,82E-03 |
| R-HSA-1227986 | Signaling by ERBB2 | ERBB2 | erb-b2 receptor tyrosine kinase 2 | 2064 | down | 1,47 | 1,93E-02 |
| R-HSA-1227986 | Signaling by ERBB2 | ERBIN | erbb2 interacting protein | 55914 | up | 1,44 | 6,49E-03 |
| R-HSA-1227986 | Signaling by ERBB2 | YES1 | YES proto-oncogene 1, Src family tyrosi | 7525 | up | 1,30 | 2,08E-02 |
| R-HSA-416572 | Sema4D induced cell migration and growth-cone colla | ERBB2 | erb-b2 receptor tyrosine kinase 2 | 2064 | down | 1,47 | 1,93E-02 |
| R-HSA-416572 | Sema4D induced cell migration and growth-cone colla | LIMK1 | LIM domain kinase 1 | 3984 | down | 1,44 | 6,49E-03 |
| R-HSA-416572 | Sema4D induced cell migration and growth-cone colla | PLXNB1 | plexin B1 | 5364 | down | 1,32 | 3,64E-02 |
| R-HSA-416572 | Sema4D induced cell migration and growth-cone colla | RND1 | Rho family GTPase 1 | 27289 | up | 4,86 | 3,18E-03 |
| R-HSA-416572 | Sema4D induced cell migration and growth-cone colla | RHOB | ras homolog family member B | 388 | up | 1,65 | 2,44E-02 |
| R-HSA-5663220 | RHO GTPases Activate Formins | PPP2CB | protein phosphatase 2 catalytic subunit | 5516 | up | 1,30 | 4,79E-02 |
| R-HSA-5663220 | RHO GTPases Activate Formins | TUBA4A | tubulin alpha 4a | 7277 | down | 1,63 | 2,97E-02 |
| R-HSA-5663220 | RHO GTPases Activate Formins | PFN1 | profilin 1 | 5216 | down | 1,20 | 3,98E-02 |
| R-HSA-5663220 | RHO GTPases Activate Formins | RHOB | ras homolog family member B | 388 | up | 1,65 | 2,44E-02 |
| R-HSA-5663220 | RHO GTPases Activate Formins | CENPO | centromere protein O | 79172 | down | 1,38 | 4,86E-02 |
| R-HSA-5663220 | RHO GTPases Activate Formins | NDEL1 | nudE neurodevelopment protein 1 like 1 | 81565 | up | 1,97 | 6,96E-03 |
| R-HSA-5663220 | RHO GTPases Activate Formins | ERCC6L | ERCC excision repair 6 like, spindle ass | 54821 | down | 2,03 | 2,60E-02 |
| R-HSA-5663220 | RHO GTPases Activate Formins | SRF | serum response factor | 6722 | up | 1,48 | 3,97E-02 |
| R-HSA-5663220 | RHO GTPases Activate Formins | CLIP1 | CAP-Gly domain containing linker protei | 6249 | up | 1,72 | 1,42E-03 |
| R-HSA-5663220 | RHO GTPases Activate Formins | SPC24 | SPC24, NDC80 kinetochore complex co | 147841 | down | 2,07 | 3,62E-02 |
| R-HSA-5663220 | RHO GTPases Activate Formins | TUBA1B | tubulin alpha 1b | 10376 | down | 1,78 | 2,30E-02 |
| R-HSA-5663220 | RHO GTPases Activate Formins | DAAM1 | dishevelled associated activator of morp | 23002 | up | 1,56 | 1,63E-02 |
| R-HSA-5663220 | RHO GTPases Activate Formins | B9D2 | B9 domain containing 2 | 80776 | down | 1,43 | 1,56E-02 |
| R-HSA-5663220 | RHO GTPases Activate Formins | ACTG1 | actin gamma 1 | 71 | up | 1,58 | 1,66E-02 |
| R-HSA-5663220 | RHO GTPases Activate Formins | INCENP | inner centromere protein | 3619 | down | 1,78 | 2,08E-02 |
| R-HSA-5663220 | RHO GTPases Activate Formins | DIAPH2 | diaphanous related formin 2 | 1730 | up | 1,25 | 2,60E-02 |
| R-HSA-5663220 | RHO GTPases Activate Formins | RANGAP1 | Ran GTPase activating protein 1 | 5905 | down | 1,48 | 1,41E-02 |
| R-HSA-5663220 | RHO GTPases Activate Formins | MIS12 | MIS12, kinetochore complex componen | 79003 | up | 1,78 | 1,57E-02 |
| R-HSA-442742 | CREB1 phosphorylation through NMDA receptor-med | CREB1 | cAMP responsive element binding prote | 1385 | up | 1,51 | 7,18E-03 |
| R-HSA-442742 | CREB1 phosphorylation through NMDA receptor-med | BRAF | B-Raf proto-oncogene, serine/threonine | 673 | up | 1,52 | 4,53E-02 |
| R-HSA-442742 | CREB1 phosphorylation through NMDA receptor-med | CAMK2D | calcium/calmodulin dependent protein k | 817 | up | 1,29 | 1,49E-02 |
| R-HSA-4411364 | Binding of TCF/LEF:CTNNB1 to target gene promote | AXIN2 | axin 2 | 8313 | down | 2,79 | 3,67E-03 |
| R-HSA-4411364 | Binding of TCF/LEF:CTNNB1 to target gene promote | TCF7 | transcription factor 7 | 6932 | down | 1,47 | 1,07E-02 |
| R-HSA-351200 | Interconversion of polyamines | SMOX | spermine oxidase | 54498 | up | 2,90 | 6,10E-03 |
| R-HSA-351200 | Interconversion of polyamines | SAT1 | spermidine/spermine N1-acetyltransfera | 6303 | up | 2,30 | 3,55E-02 |
| R-HSA-193681 | Ceramide signalling | SMPD2 | sphingomyelin phosphodiesterase 2 | 6610 | down | 1,22 | 4,95E-02 |
| R-HSA-193681 | Ceramide signalling | NGFR | nerve growth factor receptor | 4804 | up | 12,35 | 2,75E-03 |
| R-HSA-425397 | Transport of vitamins, nucleosides, and related molec | SLC33A1 | solute carrier family 33 member 1 | 9197 | up | 1,72 | 2,23E-02 |
| R-HSA-425397 | Transport of vitamins, nucleosides, and related molec | SLC5A6 | solute carrier family 5 member 6 | 8884 | down | 1,35 | 8,34E-03 |
| R-HSA-173599 | Formation of the active cofactor, UDP-glucuronate | UGDH | UDP-glucose 6-dehydrogenase | 7358 | up | 1,53 | 3,49E-02 |
| R-HSA-173599 | Formation of the active cofactor, UDP-glucuronate | UGP2 | UDP-glucose pyrophosphorylase 2 | 7360 | up | 1,37 | 3,47E-02 |
| R-HSA-2151209 | Activation of PPARGC1A (PGC-1alpha) by phosphory | PRKAB2 | protein kinase AMP-activated non-cataly | 5565 | up | 1,83 | 1,27E-02 |
| R-HSA-2151209 | Activation of PPARGC1A (PGC-1alpha) by phosphory | PRKAG2 | protein kinase AMP-activated non-cataly | 51422 | up | 1,28 | 4,57E-02 |
| R-HSA-2151209 | Activation of PPARGC1A (PGC-1alpha) by phosphory | MAPK12 | mitogen-activated protein kinase 12 | 6300 | down | 1,37 | 3,32E-03 |
| R-HSA-74749 | Signal attenuation | GRB10 | growth factor receptor bound protein 10 | 2887 | up | 1,86 | 2,07E-02 |
| R-HSA-74749 | Signal attenuation | IRS2 | insulin receptor substrate 2 | 8660 | up | 2,21 | 2,22E-02 |
| R-HSA-74749 | Signal attenuation | IRS1 | insulin receptor substrate 1 | 3667 | down | 1,89 | 6,25E-03 |
| R-HSA-70635 | Urea cycle | NMRAL1 | NmrA like redox sensor 1 | 57407 | down | 1,28 | 3,98E-02 |
| R-HSA-70635 | Urea cycle | ARG2 | arginase 2 | 384 | up | 7,09 | 3,92E-04 |
| R-HSA-70635 | Urea cycle | NAGS | N-acetylglutamate synthase | 162417 | down | 1,62 | 4,90E-02 |
| R-HSA-1250342 | PI3K events in ERBB4 signaling | PIK3CA | phosphatidylinositol-4,5-bisphosphate 3 | 5290 | up | 1,33 | 3,17E-02 |
| R-HSA-1250342 | PI3K events in ERBB4 signaling | PIK3R1 | phosphoinositide-3-kinase regulatory su | 5295 | down | 2,04 | 8,40E-03 |
| R-HSA-1250342 | PI3K events in ERBB4 signaling | HBEGF | heparin binding EGF like growth factor | 1839 | up | 3,67 | 5,67E-03 |
| R-HSA-5685939 | HDR through MMEJ (alt-NHEJ) | FEN1 | flap structure-specific endonuclease 1 | 2237 | down | 2,04 | 1,60E-02 |
| R-HSA-5685939 | HDR through MMEJ (alt-NHEJ) | NBN | nibrin | 4683 | up | 1,25 | 3,15E-02 |
| R-HSA-5685939 | HDR through MMEJ (alt-NHEJ) | PARP1 | poly(ADP-ribose) polymerase 1 | 142 | down | 1,50 | 3,92E-02 |
| R-HSA-167290 | Pausing and recovery of HIV elongation | POLR2B | RNA polymerase II subunit B | 5431 | down | 1,23 | 3,81E-02 |
| R-HSA-167290 | Pausing and recovery of HIV elongation | POLR2L | RNA polymerase II subunit L | 5441 | down | 1,52 | 3,58E-03 |
| R-HSA-167290 | Pausing and recovery of HIV elongation | POLR2F | RNA polymerase II subunit F | 5435 | down | 1,41 | 3,22E-02 |
| R-HSA-167290 | Pausing and recovery of HIV elongation | TCEA1 | transcription elongation factor A1 | 6917 | up | 1,76 | 1,81E-02 |
| R-HSA-167290 | Pausing and recovery of HIV elongation | POLR2E | RNA polymerase II subunit E | 5434 | down | 1,31 | 3,18E-03 |
| R-HSA-167290 | Pausing and recovery of HIV elongation | CCNT1 | cyclin T1 | 904 | up | 1,34 | 3,05E-02 |
| R-HSA-167287 | HIV elongation arrest and recovery | CCNT1 | cyclin T1 | 904 | up | 1,34 | 3,05E-02 |
| R-HSA-167287 | HIV elongation arrest and recovery | POLR2E | RNA polymerase II subunit E | 5434 | down | 1,31 | 3,18E-03 |
| R-HSA-167287 | HIV elongation arrest and recovery | POLR2L | RNA polymerase II subunit L | 5441 | down | 1,52 | 3,58E-03 |
| R-HSA-167287 | HIV elongation arrest and recovery | TCEA1 | transcription elongation factor A1 | 6917 | up | 1,76 | 1,81E-02 |
| R-HSA-167287 | HIV elongation arrest and recovery | POLR2F | RNA polymerase II subunit F | 5435 | down | 1,41 | 3,22E-02 |
| R-HSA-167287 | HIV elongation arrest and recovery | POLR2B | RNA polymerase II subunit B | 5431 | down | 1,23 | 3,81E-02 |
| R-HSA-2129379 | Molecules associated with elastic fibres | BMP4 | bone morphogenetic protein 4 | 652 | down | 2,60 | 1,26E-02 |
| R-HSA-2129379 | Molecules associated with elastic fibres | EMILIN2 | elastin microfibril interfacer 2 | 84034 | down | 1,40 | 4,61E-02 |
| R-HSA-2129379 | Molecules associated with elastic fibres | EFEMP1 | EGF containing fibulin extracellular matr | 2202 | down | 3,55 | 4,54E-02 |
| R-HSA-2129379 | Molecules associated with elastic fibres | EMILIN1 | elastin microfibril interfacer 1 | 11117 | down | 2,02 | 1,49E-02 |
| R-HSA-5617472 | Activation of anterior HOX genes in hindbrain develop | RXRA | retinoid X receptor alpha | 6256 | down | 1,34 | 3,67E-03 |
| R-HSA-5617472 | Activation of anterior HOX genes in hindbrain develop | H2AFX | H2A histone family member X | 3014 | down | 2,11 | 1,30E-03 |
| R-HSA-5617472 | Activation of anterior HOX genes in hindbrain develop | MEIS1 | Meis homeobox 1 | 4211 | down | 1,47 | 3,49E-02 |
| R-HSA-5617472 | Activation of anterior HOX genes in hindbrain develop | CNOT9 | CCR4-NOT transcription complex subun | 9125 | down | 1,30 | 4,90E-02 |
| R-HSA-5617472 | Activation of anterior HOX genes in hindbrain develop | POLR2B | RNA polymerase II subunit B | 5431 | down | 1,23 | 3,81E-02 |
| R-HSA-5617472 | Activation of anterior HOX genes in hindbrain develop | WDR5 | WD repeat domain 5 | 11091 | down | 1,28 | 3,81E-03 |
| R-HSA-5617472 | Activation of anterior HOX genes in hindbrain develop | POLR2L | RNA polymerase II subunit L | 5441 | down | 1,52 | 3,58E-03 |
| R-HSA-5617472 | Activation of anterior HOX genes in hindbrain develop | POLR2F | RNA polymerase II subunit F | 5435 | down | 1,41 | 3,22E-02 |
| R-HSA-5617472 | Activation of anterior HOX genes in hindbrain develop | POLR2E | RNA polymerase II subunit E | 5434 | down | 1,31 | 3,18E-03 |
| R-HSA-5617472 | Activation of anterior HOX genes in hindbrain develop | NCOA3 | nuclear receptor coactivator 3 | 8202 | up | 1,53 | 4,13E-02 |
| R-HSA-1839117 | Signaling by cytosolic FGFR1 fusion mutants | PIK3CA | phosphatidylinositol-4,5-bisphosphate 3 | 5290 | up | 1,33 | 3,17E-02 |
| R-HSA-1839117 | Signaling by cytosolic FGFR1 fusion mutants | PIK3R1 | phosphoinositide-3-kinase regulatory su | 5295 | down | 2,04 | 8,40E-03 |
| R-HSA-1839117 | Signaling by cytosolic FGFR1 fusion mutants | FGFR1OP2 | FGFR1 oncogene partner 2 | 26127 | up | 1,39 | 2,09E-02 |
| R-HSA-1839117 | Signaling by cytosolic FGFR1 fusion mutants | LRRFIP1 | LRR binding FLII interacting protein 1 | 9208 | up | 1,28 | 2,23E-02 |
| R-HSA-264876 | Insulin processing | VAMP2 | vesicle associated membrane protein 2 | 6844 | up | 1,50 | 4,82E-02 |
| R-HSA-264876 | Insulin processing | KIF5B | kinesin family member 5B | 3799 | up | 1,40 | 7,18E-03 |
| R-HSA-264876 | Insulin processing | SLC30A7 | solute carrier family 30 member 7 | 148867 | up | 1,55 | 2,39E-02 |
| R-HSA-264876 | Insulin processing | ERO1B | endoplasmic reticulum oxidoreductase 1 | 56605 | up | 8,94 | 1,01E-02 |
| R-HSA-381033 | ATF6 (ATF6-alpha) activates chaperones | ATF6 | activating transcription factor 6 | 22926 | up | 1,68 | 1,14E-02 |
| R-HSA-381033 | ATF6 (ATF6-alpha) activates chaperones | HSPA5 | heat shock protein family A (Hsp70) mem | 3309 | up | 5,02 | 6,98E-04 |
| R-HSA-381042 | PERK regulates gene expression | HSPA5 | heat shock protein family A (Hsp70) mem | 3309 | up | 5,02 | 6,98E-04 |
| R-HSA-381042 | PERK regulates gene expression | EIF2AK3 | eukaryotic translation initiation factor 2 a | 9451 | up | 2,82 | 1,11E-03 |
| R-HSA-205043 | NRIF signals cell death from the nucleus | NGFR | nerve growth factor receptor | 4804 | up | 12,35 | 2,75E-03 |
| R-HSA-205043 | NRIF signals cell death from the nucleus | SQSTM1 | sequestosome 1 | 8878 | up | 1,88 | 2,42E-02 |
| R-HSA-205043 | NRIF signals cell death from the nucleus | MAPK8 | mitogen-activated protein kinase 8 | 5599 | up | 1,38 | 3,05E-02 |
| R-HSA-2565942 | Regulation of PLK1 Activity at G2/M Transition | TUBA4A | tubulin alpha 4a | 7277 | down | 1,63 | 2,97E-02 |
| R-HSA-2565942 | Regulation of PLK1 Activity at G2/M Transition | CEP290 | centrosomal protein 290 | 80184 | up | 1,63 | 1,81E-02 |
| R-HSA-2565942 | Regulation of PLK1 Activity at G2/M Transition | OPTN | optineurin | 10133 | up | 1,48 | 2,60E-02 |
| R-HSA-2565942 | Regulation of PLK1 Activity at G2/M Transition | CEP164 | centrosomal protein 164 | 22897 | up | 1,46 | 2,91E-02 |
| R-HSA-2565942 | Regulation of PLK1 Activity at G2/M Transition | PCM1 | pericentriolar material 1 | 5108 | up | 1,23 | 5,27E-03 |
| R-HSA-2565942 | Regulation of PLK1 Activity at G2/M Transition | PPP1R12A | protein phosphatase 1 regulatory subur | 4659 | up | 1,25 | 1,03E-02 |
| R-HSA-2565942 | Regulation of PLK1 Activity at G2/M Transition | TUBG1 | tubulin gamma 1 | 7283 | down | 1,50 | 1,42E-02 |
| R-HSA-2565942 | Regulation of PLK1 Activity at G2/M Transition | CEP63 | centrosomal protein 63 | 80254 | up | 1,42 | 2,18E-02 |
| R-HSA-2565942 | Regulation of PLK1 Activity at G2/M Transition | CEP57 | centrosomal protein 57 | 9702 | up | 1,57 | 1,19E-02 |
| R-HSA-2565942 | Regulation of PLK1 Activity at G2/M Transition | TUBB | tubulin beta class I | 203068 | down | 1,56 | 2,67E-02 |
| R-HSA-2565942 | Regulation of PLK1 Activity at G2/M Transition | AKAP9 | A-kinase anchoring protein 9 | 10142 | up | 1,47 | 1,80E-02 |
| R-HSA-2046105 | Linoleic acid (LA) metabolism | FADS1 | fatty acid desaturase 1 | 3992 | down | 1,54 | 8,39E-03 |
| R-HSA-2046105 | Linoleic acid (LA) metabolism | ACSL1 | acyl-CoA synthetase long chain family m | 2180 | down | 1,54 | 3,01E-02 |
| R-HSA-174430 | Telomere C-strand synthesis initiation | POLE | DNA polymerase epsilon, catalytic subu | 5426 | down | 1,55 | 2,67E-02 |
| R-HSA-174430 | Telomere C-strand synthesis initiation | POLA2 | DNA polymerase alpha 2, accessory sub | 23649 | down | 1,72 | 2,46E-02 |
| R-HSA-68952 | DNA replication initiation | POLA2 | DNA polymerase alpha 2, accessory sub | 23649 | down | 1,72 | 2,46E-02 |
| R-HSA-68952 | DNA replication initiation | POLE | DNA polymerase epsilon, catalytic subu | 5426 | down | 1,55 | 2,67E-02 |
| R-HSA-3595174 | Defective CHST14 causes EDS, musculocontractural | CHST14 | carbohydrate sulfotransferase 14 | 113189 | down | 1,92 | 4,01E-03 |
| R-HSA-3595174 | Defective CHST14 causes EDS, musculocontractural | CSPG4 | chondroitin sulfate proteoglycan 4 | 1464 | down | 2,22 | 3,99E-02 |
| R-HSA-68877 | Mitotic Prometaphase | SPC24 | SPC24, NDC80 kinetochore complex co | 147841 | down | 2,07 | 3,62E-02 |
| R-HSA-68877 | Mitotic Prometaphase | CLIP1 | CAP-Gly domain containing linker protei | 6249 | up | 1,72 | 1,42E-03 |
| R-HSA-68877 | Mitotic Prometaphase | NDEL1 | nudE neurodevelopment protein 1 like 1 | 81565 | up | 1,97 | 6,96E-03 |
| R-HSA-68877 | Mitotic Prometaphase | ERCC6L | ERCC excision repair 6 like, spindle ass | 54821 | down | 2,03 | 2,60E-02 |
| R-HSA-68877 | Mitotic Prometaphase | PPP2CB | protein phosphatase 2 catalytic subunit | 5516 | up | 1,30 | 4,79E-02 |
| R-HSA-68877 | Mitotic Prometaphase | CENPO | centromere protein O | 79172 | down | 1,38 | 4,86E-02 |
| R-HSA-68877 | Mitotic Prometaphase | RANGAP1 | Ran GTPase activating protein 1 | 5905 | down | 1,48 | 1,41E-02 |
| R-HSA-68877 | Mitotic Prometaphase | MIS12 | MIS12, kinetochore complex componen | 79003 | up | 1,78 | 1,57E-02 |
| R-HSA-68877 | Mitotic Prometaphase | INCENP | inner centromere protein | 3619 | down | 1,78 | 2,08E-02 |
| R-HSA-68877 | Mitotic Prometaphase | B9D2 | B9 domain containing 2 | 80776 | down | 1,43 | 1,56E-02 |
| R-HSA-68877 | Mitotic Prometaphase | TUBA1B | tubulin alpha 1b | 10376 | down | 1,78 | 2,30E-02 |
| R-HSA-68877 | Mitotic Prometaphase | TUBA4A | tubulin alpha 4a | 7277 | down | 1,63 | 2,97E-02 |
| R-HSA-936837 | Ion transport by P-type ATPases | ATP11A | ATPase phospholipid transporting 11A | 23250 | down | 1,28 | 4,20E-02 |
| R-HSA-936837 | Ion transport by P-type ATPases | CALM2 | calmodulin 2 | 805 | down | 1,43 | 2,23E-02 |
| R-HSA-936837 | Ion transport by P-type ATPases | ATP2A3 | ATPase sarcoplasmic/endoplasmic retic | 489 | down | 1,62 | 4,34E-02 |
| R-HSA-936837 | Ion transport by P-type ATPases | CALM3 | calmodulin 3 | 808 | down | 1,43 | 2,23E-02 |
| R-HSA-936837 | Ion transport by P-type ATPases | ATP11B | ATPase phospholipid transporting 11B | 23200 | up | 1,40 | 9,63E-03 |
| R-HSA-936837 | Ion transport by P-type ATPases | CALM1 | calmodulin 1 | 801 | down | 1,43 | 2,23E-02 |
| R-HSA-936837 | Ion transport by P-type ATPases | CAMK2D | calcium/calmodulin dependent protein k | 817 | up | 1,29 | 1,49E-02 |
| R-HSA-936837 | Ion transport by P-type ATPases | ATP2B1 | ATPase plasma membrane Ca2+ transp | 490 | up | 1,35 | 4,17E-02 |
| R-HSA-6783310 | Fanconi Anemia Pathway | RPA1 | replication protein A1 | 6117 | down | 1,41 | 4,93E-02 |
| R-HSA-6783310 | Fanconi Anemia Pathway | CENPX | centromere protein X | 201254 | down | 1,50 | 2,52E-02 |
| R-HSA-6783310 | Fanconi Anemia Pathway | DCLRE1B | DNA cross-link repair 1B | 64858 | down | 1,70 | 1,16E-02 |
| R-HSA-6783310 | Fanconi Anemia Pathway | FAAP20 | Fanconi anemia core complex associate | 199990 | down | 1,22 | 1,49E-02 |

| | | | | | | | |
|---|---|---|---|---|---|---|---|
| R-HSA-373760 | L1CAM interactions | L1CAM | L1 cell adhesion molecule | 3897 | down | 2,86 | 3,09E-02 |
| R-HSA-373760 | L1CAM interactions | EPHB2 | EPH receptor B2 | 2048 | down | 1,66 | 2,52E-02 |
| R-HSA-373760 | L1CAM interactions | RANBP9 | RAN binding protein 9 | 10048 | up | 1,43 | 1,66E-02 |
| R-HSA-180292 | GAB1 signalosome | PIK3R1 | phosphoinositide-3-kinase regulatory su | 5295 | down | 2,04 | 8,40E-03 |
| R-HSA-180292 | GAB1 signalosome | CSK | C-terminal Src kinase | 1445 | down | 1,24 | 5,67E-03 |
| R-HSA-180292 | GAB1 signalosome | PIK3CA | phosphatidylinositol-4,5-bisphosphate 3 | 5290 | up | 1,33 | 3,17E-02 |
| R-HSA-428790 | No details found for 428790 | SLC2A6 | solute carrier family 2 member 6 | 11182 | down | 1,81 | 1,03E-02 |
| R-HSA-428790 | No details found for 428790 | SLC2A8 | solute carrier family 2 member 8 | 29988 | down | 1,22 | 3,58E-03 |
| R-HSA-380972 | Energy dependent regulation of mTOR by LKB1-AMF | RPTOR | regulatory associated protein of MTOR c | 57521 | down | 1,20 | 1,16E-02 |
| R-HSA-380972 | Energy dependent regulation of mTOR by LKB1-AMF | RRAGC | Ras related GTP binding C | 64121 | up | 1,78 | 6,10E-03 |
| R-HSA-380972 | Energy dependent regulation of mTOR by LKB1-AMF | PRKAB2 | protein kinase AMP-activated non-catal | 5565 | up | 1,83 | 1,27E-02 |
| R-HSA-380972 | Energy dependent regulation of mTOR by LKB1-AMF | STK11 | serine/threonine kinase 11 | 6794 | down | 1,25 | 3,05E-02 |
| R-HSA-380972 | Energy dependent regulation of mTOR by LKB1-AMF | PRKAG2 | protein kinase AMP-activated non-catal | 51422 | up | 1,28 | 4,57E-02 |
| R-HSA-210991 | Basigin interactions | SLC3A2 | solute carrier family 3 member 2 | 6520 | up | 2,41 | 7,95E-03 |
| R-HSA-210991 | Basigin interactions | PPIL2 | peptidylprolyl isomerase like 2 | 23759 | down | 1,39 | 6,52E-03 |
| R-HSA-210991 | Basigin interactions | L1CAM | L1 cell adhesion molecule | 3897 | down | 2,86 | 3,09E-02 |
| R-HSA-210991 | Basigin interactions | SLC7A6 | solute carrier family 7 member 6 | 9057 | down | 1,49 | 1,37E-02 |
| R-HSA-3108214 | SUMOylation of DNA damage response and repair pr | CBX4 | chromobox 4 | 8535 | up | 2,70 | 4,00E-03 |
| R-HSA-3108214 | SUMOylation of DNA damage response and repair pr | SP100 | SP100 nuclear antigen | 6672 | up | 1,31 | 2,71E-02 |
| R-HSA-3108214 | SUMOylation of DNA damage response and repair pr | NUP58 | nucleoporin 58 | 9818 | up | 1,61 | 1,28E-02 |
| R-HSA-3108214 | SUMOylation of DNA damage response and repair pr | PARP1 | poly(ADP-ribose) polymerase 1 | 142 | down | 1,50 | 3,92E-02 |
| R-HSA-3108214 | SUMOylation of DNA damage response and repair pr | CBX2 | chromobox 2 | 84733 | down | 2,70 | 8,09E-03 |
| R-HSA-3108214 | SUMOylation of DNA damage response and repair pr | STAG2 | stromal antigen 2 | 10735 | up | 1,26 | 3,69E-02 |
| R-HSA-3108214 | SUMOylation of DNA damage response and repair pr | RPA1 | replication protein A1 | 6117 | down | 1,41 | 4,93E-02 |
| R-HSA-3108214 | SUMOylation of DNA damage response and repair pr | CBX8 | chromobox 8 | 57332 | down | 1,30 | 2,19E-02 |
| R-HSA-3108214 | SUMOylation of DNA damage response and repair pr | SUMO3 | small ubiquitin-like modifier 3 | 6612 | down | 1,24 | 4,67E-02 |
| R-HSA-3108214 | SUMOylation of DNA damage response and repair pr | NUP62 | nucleoporin 62 | 23636 | down | 1,52 | 4,39E-03 |
| R-HSA-5673001 | RAF/MAP kinase cascade | CALM2 | calmodulin 2 | 805 | down | 1,43 | 2,23E-02 |
| R-HSA-5673001 | RAF/MAP kinase cascade | CALM3 | calmodulin 3 | 808 | down | 1,43 | 2,23E-02 |
| R-HSA-5673001 | RAF/MAP kinase cascade | AKAP9 | A-kinase anchoring protein 9 | 10142 | up | 1,47 | 1,80E-02 |
| R-HSA-5673001 | RAF/MAP kinase cascade | FGF2 | fibroblast growth factor 2 | 2247 | down | 5,35 | 3,09E-02 |
| R-HSA-5673001 | RAF/MAP kinase cascade | CALM1 | calmodulin 1 | 801 | down | 1,43 | 2,23E-02 |
| R-HSA-5673001 | RAF/MAP kinase cascade | IRS1 | insulin receptor substrate 1 | 3667 | down | 1,89 | 6,25E-03 |
| R-HSA-5673001 | RAF/MAP kinase cascade | ERBB2 | erb-b2 receptor tyrosine kinase 2 | 2064 | down | 1,47 | 1,93E-02 |
| R-HSA-5673001 | RAF/MAP kinase cascade | IRS2 | insulin receptor substrate 2 | 8660 | up | 2,21 | 2,22E-02 |
| R-HSA-5673001 | RAF/MAP kinase cascade | HRAS | HRas proto-oncogene, GTPase | 3265 | down | 1,31 | 2,19E-03 |
| R-HSA-5673001 | RAF/MAP kinase cascade | CAMK2D | calcium/calmodulin dependent protein k | 817 | up | 1,29 | 1,49E-02 |
| R-HSA-5673001 | RAF/MAP kinase cascade | SPTBN1 | spectrin beta, non-erythrocytic 1 | 6711 | up | 1,42 | 1,66E-02 |
| R-HSA-5673001 | RAF/MAP kinase cascade | JAK1 | Janus kinase 1 | 3716 | up | 1,53 | 2,13E-02 |
| R-HSA-5673001 | RAF/MAP kinase cascade | RASGRP3 | RAS guanyl releasing protein 3 | 25780 | up | 2,59 | 1,78E-02 |
| R-HSA-5673001 | RAF/MAP kinase cascade | HBEGF | heparin binding EGF like growth factor | 1839 | up | 3,67 | 5,67E-03 |
| R-HSA-5673001 | RAF/MAP kinase cascade | FGFR3 | fibroblast growth factor receptor 3 | 2261 | down | 1,70 | 4,60E-02 |
| R-HSA-5362517 | Signaling by Retinoic Acid | PDK1 | pyruvate dehydrogenase kinase 1 | 5163 | up | 1,31 | 2,69E-03 |
| R-HSA-5362517 | Signaling by Retinoic Acid | PDK2 | pyruvate dehydrogenase kinase 2 | 5164 | down | 1,42 | 1,03E-02 |
| R-HSA-5362517 | Signaling by Retinoic Acid | DLD | dihydrolipoamide dehydrogenase | 1738 | up | 1,16 | 4,53E-02 |
| R-HSA-5362517 | Signaling by Retinoic Acid | RXRA | retinoid X receptor alpha | 6256 | down | 1,34 | 3,67E-03 |
| R-HSA-450321 | JNK (c-Jun kinases) phosphorylation and activation m | TAB1 | TGF-beta activated kinase 1 (MAP3K7) | 10454 | down | 1,29 | 1,94E-02 |
| R-HSA-450321 | JNK (c-Jun kinases) phosphorylation and activation m | RIPK2 | receptor interacting serine/threonine kin | 8767 | up | 1,72 | 8,31E-03 |
| R-HSA-450321 | JNK (c-Jun kinases) phosphorylation and activation m | MAPK8 | mitogen-activated protein kinase 8 | 5599 | up | 1,38 | 3,05E-02 |
| R-HSA-450321 | JNK (c-Jun kinases) phosphorylation and activation m | TAB3 | TGF-beta activated kinase 1 and MAP3 | 257397 | up | 1,40 | 4,94E-02 |
| R-HSA-69895 | Transcriptional activation of cell cycle inhibitor p21 | CDKN1A | cyclin dependent kinase inhibitor 1A | 1026 | up | 3,62 | 5,43E-03 |
| R-HSA-69895 | Transcriptional activation of cell cycle inhibitor p21 | ZNF385A | zinc finger protein 385A | 25946 | down | 1,64 | 3,03E-02 |
| R-HSA-165158 | Activation of AKT2 | TRIB3 | tribbles pseudokinase 3 | 57761 | up | 3,17 | 8,07E-03 |
| R-HSA-165158 | Activation of AKT2 | AKT2 | AKT serine/threonine kinase 2 | 208 | down | 1,28 | 6,33E-03 |
| R-HSA-198753 | ERK/MAPK targets | MEF2A | myocyte enhancer factor 2A | 4205 | up | 1,55 | 1,41E-02 |
| R-HSA-198753 | ERK/MAPK targets | RPS6KA5 | ribosomal protein S6 kinase A5 | 9252 | up | 2,11 | 5,94E-03 |
| R-HSA-198753 | ERK/MAPK targets | PPP2CB | protein phosphatase 2 catalytic subunit | 5516 | up | 1,30 | 4,79E-02 |
| R-HSA-5620912 | Anchoring of the basal body to the plasma membrane | TCTN2 | tectonic family member 2 | 79867 | down | 1,44 | 2,63E-02 |
| R-HSA-5620912 | Anchoring of the basal body to the plasma membrane | TUBG1 | tubulin gamma 1 | 7283 | down | 1,50 | 1,42E-02 |
| R-HSA-5620912 | Anchoring of the basal body to the plasma membrane | B9D2 | B9 domain containing 2 | 80776 | down | 1,43 | 1,56E-02 |
| R-HSA-5620912 | Anchoring of the basal body to the plasma membrane | PCM1 | pericentriolar material 1 | 5108 | up | 1,23 | 5,27E-03 |
| R-HSA-5620912 | Anchoring of the basal body to the plasma membrane | CEP164 | centrosomal protein 164 | 22897 | up | 1,46 | 2,91E-02 |
| R-HSA-5620912 | Anchoring of the basal body to the plasma membrane | CEP290 | centrosomal protein 290 | 80184 | up | 1,63 | 1,81E-02 |
| R-HSA-5620912 | Anchoring of the basal body to the plasma membrane | TUBA4A | tubulin alpha 4a | 7277 | down | 1,63 | 2,97E-02 |
| R-HSA-5620912 | Anchoring of the basal body to the plasma membrane | IQCB1 | IQ motif containing B1 | 9657 | up | 1,34 | 1,42E-02 |
| R-HSA-5620912 | Anchoring of the basal body to the plasma membrane | AKAP9 | A-kinase anchoring protein 9 | 10142 | up | 1,47 | 1,80E-02 |
| R-HSA-5620912 | Anchoring of the basal body to the plasma membrane | TUBB | tubulin beta class I | 203068 | down | 1,56 | 2,67E-02 |
| R-HSA-5620912 | Anchoring of the basal body to the plasma membrane | AHI1 | Abelson helper integration site 1 | 54806 | up | 1,35 | 2,72E-03 |
| R-HSA-5620912 | Anchoring of the basal body to the plasma membrane | CEP57 | centrosomal protein 57 | 9702 | up | 1,57 | 1,19E-02 |
| R-HSA-5620912 | Anchoring of the basal body to the plasma membrane | CEP63 | centrosomal protein 63 | 80254 | up | 1,42 | 2,18E-02 |
| R-HSA-187706 | Signalling to p38 via RIT and RIN | RIT1 | Ras like without CAAX 1 | 6016 | up | 1,78 | 1,44E-02 |
| R-HSA-187706 | Signalling to p38 via RIT and RIN | BRAF | B-Raf proto-oncogene, serine/threonine | 673 | up | 1,52 | 4,53E-02 |
| R-HSA-71032 | Propionyl-CoA catabolism | PCCA | propionyl-CoA carboxylase alpha subun | 5095 | up | 1,37 | 2,03E-02 |
| R-HSA-71032 | Propionyl-CoA catabolism | MUT | methylmalonyl-CoA mutase | 4594 | up | 1,31 | 4,57E-03 |
| R-HSA-73817 | Purine ribonucleoside monophosphate biosynthesis | IMPDH1 | inosine monophosphate dehydrogenas | 3614 | down | 1,56 | 1,49E-03 |
| R-HSA-73817 | Purine ribonucleoside monophosphate biosynthesis | PFAS | phosphoribosylformylglycinamidine synth | 5198 | down | 1,53 | 1,69E-02 |
| R-HSA-73817 | Purine ribonucleoside monophosphate biosynthesis | ADSS | adenylosuccinate synthase | 159 | up | 1,21 | 2,47E-02 |
| R-HSA-446353 | Cell-extracellular matrix interactions | LIMS2 | LIM zinc finger domain containing 2 | 55679 | down | 1,42 | 1,11E-02 |
| R-HSA-446353 | Cell-extracellular matrix interactions | ACTG1 | actin gamma 1 | 71 | up | 1,58 | 1,66E-02 |
| R-HSA-446353 | Cell-extracellular matrix interactions | FERMT2 | fermitin family member 2 | 10979 | up | 5,34 | 2,64E-03 |
| R-HSA-210990 | PECAM1 interactions | LYN | LYN proto-oncogene, Src family tyrosine | 4067 | down | 1,36 | 2,34E-02 |
| R-HSA-210990 | PECAM1 interactions | YES1 | YES proto-oncogene 1, Src family tyros | 7525 | up | 1,30 | 2,08E-02 |
| R-HSA-210990 | PECAM1 interactions | PLCG1 | phospholipase C gamma 1 | 5335 | down | 1,51 | 2,85E-02 |
| R-HSA-2219530 | Constitutive Signaling by Aberrant PI3K in Cancer | FGF2 | fibroblast growth factor 2 | 2247 | down | 5,35 | 3,09E-02 |
| R-HSA-2219530 | Constitutive Signaling by Aberrant PI3K in Cancer | IRS1 | insulin receptor substrate 1 | 3667 | down | 1,89 | 6,25E-03 |
| R-HSA-2219530 | Constitutive Signaling by Aberrant PI3K in Cancer | ERBB2 | erb-b2 receptor tyrosine kinase 2 | 2064 | down | 1,47 | 1,93E-02 |
| R-HSA-2219530 | Constitutive Signaling by Aberrant PI3K in Cancer | IRS2 | insulin receptor substrate 2 | 8660 | up | 2,21 | 2,22E-02 |
| R-HSA-2219530 | Constitutive Signaling by Aberrant PI3K in Cancer | PIK3R1 | phosphoinositide-3-kinase regulatory su | 5295 | down | 2,04 | 8,40E-03 |
| R-HSA-2219530 | Constitutive Signaling by Aberrant PI3K in Cancer | HBEGF | heparin binding EGF like growth factor | 1839 | up | 3,67 | 5,67E-03 |
| R-HSA-2219530 | Constitutive Signaling by Aberrant PI3K in Cancer | PIK3CA | phosphatidylinositol-4,5-bisphosphate 3 | 5290 | up | 1,33 | 3,17E-02 |
| R-HSA-2219530 | Constitutive Signaling by Aberrant PI3K in Cancer | FGFR3 | fibroblast growth factor receptor 3 | 2261 | down | 1,70 | 4,60E-02 |
| R-HSA-4086398 | Ca2+ pathway | TCF7 | transcription factor 7 | 6932 | down | 1,47 | 1,07E-02 |
| R-HSA-4086398 | Ca2+ pathway | FZD4 | frizzled class receptor 4 | 8322 | down | 2,17 | 3,82E-02 |
| R-HSA-4086398 | Ca2+ pathway | CALM3 | calmodulin 3 | 808 | down | 1,43 | 2,23E-02 |
| R-HSA-4086398 | Ca2+ pathway | AGO3 | argonaute 3, RISC catalytic component | 192669 | up | 1,36 | 1,80E-02 |
| R-HSA-4086398 | Ca2+ pathway | CALM2 | calmodulin 2 | 805 | down | 1,43 | 2,23E-02 |
| R-HSA-4086398 | Ca2+ pathway | CALM1 | calmodulin 1 | 801 | down | 1,43 | 2,23E-02 |
| R-HSA-193648 | NRAGE signals death through JNK | FGD4 | FYVE, RhoGEF and PH domain contain | 121512 | up | 1,60 | 1,34E-02 |
| R-HSA-193648 | NRAGE signals death through JNK | NGFR | nerve growth factor receptor | 4804 | up | 12,35 | 2,75E-03 |
| R-HSA-193648 | NRAGE signals death through JNK | ARHGEF2 | Rho/Rac guanine nucleotide exchange | 9181 | up | 2,09 | 1,10E-02 |
| R-HSA-193648 | NRAGE signals death through JNK | ARHGEF19 | Rho guanine nucleotide exchange facto | 128272 | down | 1,55 | 1,82E-02 |
| R-HSA-193648 | NRAGE signals death through JNK | MAPK8 | mitogen-activated protein kinase 8 | 5599 | up | 1,38 | 3,05E-02 |
| R-HSA-193648 | NRAGE signals death through JNK | SOS2 | SOS Ras/Rho guanine nucleotide exch | 6655 | up | 1,82 | 1,41E-02 |
| R-HSA-186763 | Downstream signal transduction | PIK3CA | phosphatidylinositol-4,5-bisphosphate 3 | 5290 | up | 1,33 | 3,17E-02 |
| R-HSA-186763 | Downstream signal transduction | PLCG1 | phospholipase C gamma 1 | 5335 | down | 1,51 | 2,85E-02 |
| R-HSA-186763 | Downstream signal transduction | NCK1 | NCK adaptor protein 1 | 4690 | up | 1,48 | 1,42E-02 |
| R-HSA-186763 | Downstream signal transduction | PIK3R1 | phosphoinositide-3-kinase regulatory su | 5295 | down | 2,04 | 8,40E-03 |
| R-HSA-186763 | Downstream signal transduction | HRAS | HRas proto-oncogene, GTPase | 3265 | down | 1,31 | 2,19E-03 |
| R-HSA-442380 | Zinc influx into cells by the SLC39 gene family | SLC39A3 | solute carrier family 39 member 3 | 29985 | down | 1,40 | 3,92E-04 |
| R-HSA-442380 | Zinc influx into cells by the SLC39 gene family | SLC39A4 | solute carrier family 39 member 4 | 55630 | down | 1,33 | 4,55E-03 |
| R-HSA-190377 | FGFR2b ligand binding and activation | FGF2 | fibroblast growth factor 2 | 2247 | down | 5,35 | 3,09E-02 |
| R-HSA-190377 | FGFR2b ligand binding and activation | FGFBP1 | fibroblast growth factor binding protein 1 | 9982 | down | 2,40 | 2,73E-02 |
| R-HSA-3000170 | Syndecan interactions | FGF2 | fibroblast growth factor 2 | 2247 | down | 5,35 | 3,09E-02 |
| R-HSA-3000170 | Syndecan interactions | SDC3 | syndecan 3 | 9672 | down | 1,68 | 1,97E-02 |
| R-HSA-3000170 | Syndecan interactions | SDC1 | syndecan 1 | 6382 | down | 2,17 | 1,61E-02 |
| R-HSA-5357905 | Regulation of TNFR1 signaling | TRAF2 | TNF receptor associated factor 2 | 7186 | down | 1,47 | 2,65E-02 |
| R-HSA-5357905 | Regulation of TNFR1 signaling | BIRC2 | baculoviral IAP repeat containing 2 | 329 | up | 1,38 | 1,86E-02 |
| R-HSA-5357905 | Regulation of TNFR1 signaling | TAX1BP1 | Tax1 binding protein 1 | 8887 | up | 1,45 | 1,32E-03 |
| R-HSA-5357905 | Regulation of TNFR1 signaling | OTUD7B | OTU deubiquitinase 7B | 56957 | up | 1,27 | 3,81E-02 |
| R-HSA-5357905 | Regulation of TNFR1 signaling | CYLD | CYLD lysine 63 deubiquitinase | 1540 | up | 1,34 | 4,86E-02 |
| R-HSA-1660662 | Glycosphingolipid metabolism | SUMF1 | sulfatase modifying factor 1 | 285362 | down | 1,29 | 3,58E-02 |
| R-HSA-1660662 | Glycosphingolipid metabolism | SMPD4 | sphingomyelin phosphodiesterase 4 | 55627 | down | 1,40 | 2,24E-03 |
| R-HSA-1660662 | Glycosphingolipid metabolism | CPTP | ceramide-1-phosphate transfer protein | 80772 | down | 1,34 | 1,22E-02 |
| R-HSA-1660662 | Glycosphingolipid metabolism | SMPD2 | sphingomyelin phosphodiesterase 2 | 6610 | down | 1,22 | 4,95E-02 |
| R-HSA-1660662 | Glycosphingolipid metabolism | ARSG | arylsulfatase G | 22901 | up | 1,36 | 3,94E-02 |
| R-HSA-196741 | Cobalamin (Cbl, vitamin B12) transport and metabolis | PRSS1 | protease, serine 1 | 5644 | up | 2,54 | 4,99E-02 |
| R-HSA-196741 | Cobalamin (Cbl, vitamin B12) transport and metabolis | MUT | methylmalonyl-CoA mutase | 4594 | up | 1,31 | 4,57E-03 |
| R-HSA-196741 | Cobalamin (Cbl, vitamin B12) transport and metabolis | CD320 | CD320 molecule | 51293 | down | 1,40 | 1,01E-02 |
| R-HSA-196741 | Cobalamin (Cbl, vitamin B12) transport and metabolis | MMADHC | methylmalonic aciduria and homocystinu | 27249 | up | 1,35 | 3,57E-02 |
| R-HSA-193670 | p75NTR negatively regulates cell cycle via SC1 | PRDM4 | PR/SET domain 4 | 11108 | up | 1,52 | 2,92E-02 |
| R-HSA-193670 | p75NTR negatively regulates cell cycle via SC1 | NGFR | nerve growth factor receptor | 4804 | up | 12,35 | 2,75E-03 |
| R-HSA-400042 | Adrenaline,noradrenaline inhibits insulin secretion | CACNB3 | calcium voltage-gated channel auxiliary | 784 | down | 1,19 | 3,30E-02 |
| R-HSA-400042 | Adrenaline,noradrenaline inhibits insulin secretion | ADRA2A | adrenoceptor alpha 2A | 150 | down | 3,92 | 2,45E-02 |
| R-HSA-400042 | Adrenaline,noradrenaline inhibits insulin secretion | GNAI2 | G protein subunit alpha i2 | 2771 | down | 1,22 | 3,65E-02 |
| R-HSA-2022923 | Dermatan sulfate biosynthesis | CSPG4 | chondroitin sulfate proteoglycan 4 | 1464 | down | 2,22 | 3,99E-02 |
| R-HSA-2022923 | Dermatan sulfate biosynthesis | CHST14 | carbohydrate sulfotransferase 14 | 113189 | down | 1,92 | 4,01E-03 |
| R-HSA-5661231 | Metallothioneins bind metals | MT1A | metallothionein 1A | 4489 | down | 1,67 | 4,76E-02 |
| R-HSA-5661231 | Metallothioneins bind metals | MT2A | metallothionein 2A | 4502 | down | 1,67 | 7,14E-03 |
| R-HSA-418597 | G alpha (z) signalling events | RGS19 | regulator of G protein signaling 19 | 10287 | down | 1,45 | 1,44E-03 |
| R-HSA-418597 | G alpha (z) signalling events | GNAI2 | G protein subunit alpha i2 | 2771 | down | 1,22 | 3,65E-02 |
| R-HSA-418597 | G alpha (z) signalling events | ADRA2A | adrenoceptor alpha 2A | 150 | down | 3,92 | 2,45E-02 |
| R-HSA-418597 | G alpha (z) signalling events | ADCY3 | adenylate cyclase 3 | 109 | down | 1,45 | 1,03E-02 |
| R-HSA-1483213 | Synthesis of PE | LPIN3 | lipin 3 | 64900 | down | 1,36 | 4,76E-02 |
| R-HSA-1483213 | Synthesis of PE | LPIN2 | lipin 2 | 9663 | up | 2,16 | 6,88E-03 |
| R-HSA-1483213 | Synthesis of PE | PISD | phosphatidylserine decarboxylase | 23761 | down | 1,20 | 2,48E-02 |
| R-HSA-2995383 | Initiation of Nuclear Envelope Reformation | VRK2 | vaccinia related kinase 2 | 7444 | up | 1,28 | 9,54E-03 |
| R-HSA-2995383 | Initiation of Nuclear Envelope Reformation | LMNB1 | lamin B1 | 4001 | down | 2,12 | 3,85E-02 |
| R-HSA-2995383 | Initiation of Nuclear Envelope Reformation | PPP2R2A | protein phosphatase 2 regulatory subur | 5520 | up | 1,26 | 3,61E-02 |
| R-HSA-2500257 | Resolution of Sister Chromatid Cohesion | CENPO | centromere protein O | 79172 | down | 1,38 | 4,86E-02 |
| R-HSA-2500257 | Resolution of Sister Chromatid Cohesion | MIS12 | MIS12, kinetochore complex componen | 79003 | up | 1,78 | 1,57E-02 |
| R-HSA-2500257 | Resolution of Sister Chromatid Cohesion | RANGAP1 | Ran GTPase activating protein 1 | 5905 | down | 1,48 | 1,41E-02 |
| R-HSA-2500257 | Resolution of Sister Chromatid Cohesion | INCENP | inner centromere protein | 3619 | down | 1,78 | 2,08E-02 |
| R-HSA-2500257 | Resolution of Sister Chromatid Cohesion | B9D2 | B9 domain containing 2 | 80776 | down | 1,43 | 1,56E-02 |
| R-HSA-2500257 | Resolution of Sister Chromatid Cohesion | TUBA1B | tubulin alpha 1b | 10376 | down | 1,78 | 2,30E-02 |
| R-HSA-2500257 | Resolution of Sister Chromatid Cohesion | STAG2 | stromal antigen 2 | 10735 | up | 1,26 | 3,69E-02 |
| R-HSA-2500257 | Resolution of Sister Chromatid Cohesion | TUBA4A | tubulin alpha 4a | 7277 | down | 1,63 | 2,97E-02 |
| R-HSA-2500257 | Resolution of Sister Chromatid Cohesion | SPC24 | SPC24, NDC80 kinetochore complex co | 147841 | down | 2,07 | 3,62E-02 |
| R-HSA-2500257 | Resolution of Sister Chromatid Cohesion | CLIP1 | CAP-Gly domain containing linker protei | 6249 | up | 1,72 | 1,42E-03 |
| R-HSA-2500257 | Resolution of Sister Chromatid Cohesion | ERCC6L | ERCC excision repair 6 like, spindle ass | 54821 | down | 2,03 | 2,60E-02 |
| R-HSA-2500257 | Resolution of Sister Chromatid Cohesion | NDEL1 | nudE neurodevelopment protein 1 like 1 | 81565 | up | 1,97 | 6,96E-03 |
| R-HSA-2500257 | Resolution of Sister Chromatid Cohesion | PPP2CB | protein phosphatase 2 catalytic subunit | 5516 | up | 1,30 | 4,79E-02 |
| R-HSA-1839130 | Signaling by activated point mutants of FGFR3 | FGFR3 | fibroblast growth factor receptor 3 | 2261 | down | 1,70 | 4,60E-02 |
| R-HSA-1839130 | Signaling by activated point mutants of FGFR3 | FGF2 | fibroblast growth factor 2 | 2247 | down | 5,35 | 3,09E-02 |
| R-HSA-2033514 | FGFR3 mutant receptor activation | FGFR3 | fibroblast growth factor receptor 3 | 2261 | down | 1,70 | 4,60E-02 |
| R-HSA-2033514 | FGFR3 mutant receptor activation | FGF2 | fibroblast growth factor 2 | 2247 | down | 5,35 | 3,09E-02 |
| R-HSA-70614 | Amino acid synthesis and interconversion (transamina | GLUL | glutamate-ammonia ligase | 2752 | down | 1,47 | 6,91E-03 |
| R-HSA-70614 | Amino acid synthesis and interconversion (transamina | GOT1 | glutamic-oxaloacetic transaminase 1 | 2805 | up | 2,30 | 5,67E-03 |
| R-HSA-70614 | Amino acid synthesis and interconversion (transamina | ASNS | asparagine synthetase (glutamine-hydro | 440 | up | 2,47 | 2,91E-02 |
| R-HSA-70614 | Amino acid synthesis and interconversion (transamina | PYCR3 | pyrroline-5-carboxylate reductase 3 | 65263 | down | 1,77 | 1,11E-03 |
| R-HSA-912631 | Regulation of signaling by CBL | LYN | LYN proto-oncogene, Src family tyrosine | 4067 | down | 1,36 | 2,34E-02 |
| R-HSA-912631 | Regulation of signaling by CBL | YES1 | YES proto-oncogene 1, Src family tyros | 7525 | up | 1,30 | 2,08E-02 |
| R-HSA-912631 | Regulation of signaling by CBL | PIK3CA | phosphatidylinositol-4,5-bisphosphate 3 | 5290 | up | 1,33 | 3,17E-02 |
| R-HSA-912631 | Regulation of signaling by CBL | PIK3R1 | phosphoinositide-3-kinase regulatory su | 5295 | down | 2,04 | 8,40E-03 |
| R-HSA-69273 | Cyclin A/B1/B2 associated events during G2/M transi | CDK7 | cyclin dependent kinase 7 | 1022 | up | 1,37 | 4,38E-02 |
| R-HSA-69273 | Cyclin A/B1/B2 associated events during G2/M transi | PPP2CB | protein phosphatase 2 catalytic subunit | 5516 | up | 1,30 | 4,79E-02 |
| R-HSA-69273 | Cyclin A/B1/B2 associated events during G2/M transi | CDC25A | cell division cycle 25A | 993 | down | 1,92 | 1,48E-02 |
| R-HSA-69273 | Cyclin A/B1/B2 associated events during G2/M transi | PPP2R2A | protein phosphatase 2 regulatory subur | 5520 | up | 1,26 | 3,61E-02 |
| R-HSA-182971 | EGFR downregulation | SPRY1 | sprouty RTK signaling antagonist 1 | 10252 | down | 1,88 | 2,51E-02 |
| R-HSA-182971 | EGFR downregulation | STAM2 | signal transducing adaptor molecule 2 | 10254 | up | 1,41 | 6,99E-03 |
| R-HSA-182971 | EGFR downregulation | EPS15 | epidermal growth factor receptor pathwa | 2060 | up | 1,30 | 7,72E-03 |
| R-HSA-182971 | EGFR downregulation | STAM | signal transducing adaptor molecule | 8027 | up | 1,23 | 3,52E-02 |
| R-HSA-375165 | NCAM signaling for neurite out-growth | RPS6KA5 | ribosomal protein S6 kinase A5 | 9252 | up | 2,11 | 5,94E-03 |
| R-HSA-375165 | NCAM signaling for neurite out-growth | CREB1 | cAMP responsive element binding prote | 1385 | up | 1,51 | 7,18E-03 |
| R-HSA-375165 | NCAM signaling for neurite out-growth | HRAS | HRas proto-oncogene, GTPase | 3265 | down | 1,31 | 2,19E-03 |
| R-HSA-375165 | NCAM signaling for neurite out-growth | SPTBN1 | spectrin beta, non-erythrocytic 1 | 6711 | up | 1,42 | 1,66E-02 |
| R-HSA-5674135 | MAP2K and MAPK activation | BRAF | B-Raf proto-oncogene, serine/threonine | 673 | up | 1,52 | 4,53E-02 |
| R-HSA-5674135 | MAP2K and MAPK activation | ARRB1 | arrestin beta 1 | 408 | down | 1,79 | 2,42E-02 |
| R-HSA-5674135 | MAP2K and MAPK activation | HRAS | HRas proto-oncogene, GTPase | 3265 | down | 1,31 | 2,19E-03 |
| R-HSA-5674135 | MAP2K and MAPK activation | CSK | C-terminal Src kinase | 1445 | down | 1,24 | 5,67E-03 |
| R-HSA-5674135 | MAP2K and MAPK activation | ACTG1 | actin gamma 1 | 71 | up | 1,58 | 1,66E-02 |
| R-HSA-5674135 | MAP2K and MAPK activation | MARK3 | microtubule affinity regulating kinase 3 | 4140 | up | 1,40 | 1,07E-02 |
| R-HSA-380320 | Recruitment of NuMA to mitotic centrosomes | TUBA4A | tubulin alpha 4a | 7277 | down | 1,63 | 2,97E-02 |
| R-HSA-380320 | Recruitment of NuMA to mitotic centrosomes | TUBA1B | tubulin alpha 1b | 10376 | down | 1,78 | 2,30E-02 |
| R-HSA-380320 | Recruitment of NuMA to mitotic centrosomes | TUBG1 | tubulin gamma 1 | 7283 | down | 1,50 | 1,42E-02 |
| R-HSA-2424491 | DAP12 signaling | HRAS | HRas proto-oncogene, GTPase | 3265 | down | 1,31 | 2,19E-03 |
| R-HSA-2424491 | DAP12 signaling | PLCG1 | phospholipase C gamma 1 | 5335 | down | 1,51 | 2,85E-02 |
| R-HSA-2424491 | DAP12 signaling | PIK3R1 | phosphoinositide-3-kinase regulatory su | 5295 | down | 2,04 | 8,40E-03 |
| R-HSA-2424491 | DAP12 signaling | PIK3CA | phosphatidylinositol-4,5-bisphosphate 3 | 5290 | up | 1,33 | 3,17E-02 |
| R-HSA-3134973 | LRR FLII-interacting protein 1 (LRRFIP1) activates typ | IRF3 | interferon regulatory factor 3 | 3661 | down | 1,33 | 2,58E-03 |
| R-HSA-3134973 | LRR FLII-interacting protein 1 (LRRFIP1) activates typ | LRRFIP1 | LRR binding FLII interacting protein 1 | 9208 | up | 1,28 | 2,23E-02 |
| R-HSA-1169092 | Activation of RAS in B cells | HRAS | HRas proto-oncogene, GTPase | 3265 | down | 1,31 | 2,19E-03 |
| R-HSA-1169092 | Activation of RAS in B cells | RASGRP3 | RAS guanyl releasing protein 3 | 25780 | up | 2,59 | 1,78E-02 |
| R-HSA-201681 | TCF dependent signaling in response to WNT | RNF146 | ring finger protein 146 | 81847 | up | 1,28 | 3,57E-02 |
| R-HSA-201681 | TCF dependent signaling in response to WNT | FZD1 | frizzled class receptor 1 | 8321 | down | 1,75 | 2,84E-02 |
| R-HSA-201681 | TCF dependent signaling in response to WNT | AXIN2 | axin 2 | 8313 | down | 2,79 | 3,67E-03 |
| R-HSA-201681 | TCF dependent signaling in response to WNT | TNKS | tankyrase | 8658 | up | 1,42 | 8,31E-03 |
| R-HSA-201681 | TCF dependent signaling in response to WNT | TNKS2 | tankyrase 2 | 80351 | up | 1,68 | 3,02E-03 |
| R-HSA-380259 | Loss of Nlp from mitotic centrosomes | CEP164 | centrosomal protein 164 | 22897 | up | 1,46 | 2,91E-02 |
| R-HSA-380259 | Loss of Nlp from mitotic centrosomes | PCM1 | pericentriolar material 1 | 5108 | up | 1,23 | 5,27E-03 |
| R-HSA-380259 | Loss of Nlp from mitotic centrosomes | CEP290 | centrosomal protein 290 | 80184 | up | 1,63 | 1,81E-02 |
| R-HSA-380259 | Loss of Nlp from mitotic centrosomes | TUBA4A | tubulin alpha 4a | 7277 | down | 1,63 | 2,97E-02 |
| R-HSA-380259 | Loss of Nlp from mitotic centrosomes | TUBG1 | tubulin gamma 1 | 7283 | down | 1,50 | 1,42E-02 |
| R-HSA-380259 | Loss of Nlp from mitotic centrosomes | AKAP9 | A-kinase anchoring protein 9 | 10142 | up | 1,47 | 1,80E-02 |
| R-HSA-380259 | Loss of Nlp from mitotic centrosomes | TUBB | tubulin beta class I | 203068 | down | 1,56 | 2,67E-02 |
| R-HSA-380259 | Loss of Nlp from mitotic centrosomes | CEP57 | centrosomal protein 57 | 9702 | up | 1,57 | 1,19E-02 |
| R-HSA-380259 | Loss of Nlp from mitotic centrosomes | CEP63 | centrosomal protein 63 | 80254 | up | 1,42 | 2,18E-02 |
| R-HSA-170984 | ARMS-mediated activation | BRAF | B-Raf proto-oncogene, serine/threonine | 673 | up | 1,52 | 4,53E-02 |

| | | | | | | | |
|---|---|---|---|---|---|---|---|
| R-HSA-170984 | ARMS-mediated activation | KIDINS220 | kinase D interacting substrate 220 | 57498 | up | 1,34 | 3,94E-02 |
| R-HSA-199920 | CREB phosphorylation | RPS6KA5 | ribosomal protein S6 kinase A5 | 9252 | up | 2,11 | 5,94E-03 |
| R-HSA-199920 | CREB phosphorylation | CREB1 | cAMP responsive element binding prote | 1385 | up | 1,51 | 7,18E-03 |
| R-HSA-399955 | SEMA3A-Plexin repulsion signaling by inhibiting Integ | RND1 | Rho family GTPase 1 | 27289 | up | 4,86 | 3,18E-03 |
| R-HSA-399955 | SEMA3A-Plexin repulsion signaling by inhibiting Integ | PLXNA1 | plexin A1 | 5361 | down | 1,28 | 9,70E-04 |
| R-HSA-399955 | SEMA3A-Plexin repulsion signaling by inhibiting Integ | PIP5K1C | phosphatidylinositol-4-phosphate 5-kina | 23396 | down | 1,39 | 1,49E-03 |
| R-HSA-5467348 | Truncations of AMER1 destabilize the destruction cor | AMER1 | APC membrane recruitment protein 1 | 139285 | down | 1,78 | 4,15E-02 |
| R-HSA-5467348 | Truncations of AMER1 destabilize the destruction cor | PPP2CB | protein phosphatase 2 catalytic subunit | 5516 | up | 1,30 | 4,79E-02 |
| R-HSA-5467348 | Truncations of AMER1 destabilize the destruction cor | CSNK1A1 | casein kinase 1 alpha 1 | 1452 | up | 1,35 | 2,84E-02 |
| R-HSA-5467337 | APC truncation mutants have impaired AXIN binding | CSNK1A1 | casein kinase 1 alpha 1 | 1452 | up | 1,35 | 2,84E-02 |
| R-HSA-5467337 | APC truncation mutants have impaired AXIN binding | AMER1 | APC membrane recruitment protein 1 | 139285 | down | 1,78 | 4,15E-02 |
| R-HSA-5467337 | APC truncation mutants have impaired AXIN binding | PPP2CB | protein phosphatase 2 catalytic subunit | 5516 | up | 1,30 | 4,79E-02 |
| R-HSA-5467340 | AXIN missense mutants destabilize the destruction co | AMER1 | APC membrane recruitment protein 1 | 139285 | down | 1,78 | 4,15E-02 |
| R-HSA-5467340 | AXIN missense mutants destabilize the destruction co | PPP2CB | protein phosphatase 2 catalytic subunit | 5516 | up | 1,30 | 4,79E-02 |
| R-HSA-5467340 | AXIN missense mutants destabilize the destruction co | CSNK1A1 | casein kinase 1 alpha 1 | 1452 | up | 1,35 | 2,84E-02 |
| R-HSA-69202 | Cyclin E associated events during G1/S transition | CDK7 | cyclin dependent kinase 7 | 1022 | up | 1,37 | 4,38E-02 |
| R-HSA-69202 | Cyclin E associated events during G1/S transition | CDKN1A | cyclin dependent kinase inhibitor 1A | 1026 | up | 3,62 | 5,43E-03 |
| R-HSA-69202 | Cyclin E associated events during G1/S transition | CDC25A | cell division cycle 25A | 993 | down | 1,92 | 1,48E-02 |
| R-HSA-1660661 | Sphingolipid de novo biosynthesis | ORMDL3 | ORMDL sphingolipid biosynthesis regula | 94103 | down | 1,35 | 1,52E-02 |
| R-HSA-1660661 | Sphingolipid de novo biosynthesis | SPHK1 | sphingosine kinase 1 | 8877 | down | 2,00 | 2,34E-02 |
| R-HSA-1660661 | Sphingolipid de novo biosynthesis | CSNK1G2 | casein kinase 1 gamma 2 | 1455 | down | 1,34 | 2,24E-03 |
| R-HSA-1660661 | Sphingolipid de novo biosynthesis | OSBP | oxysterol binding protein | 5007 | up | 1,53 | 1,85E-02 |
| R-HSA-1660661 | Sphingolipid de novo biosynthesis | SGMS2 | sphingomyelin synthase 2 | 166929 | up | 1,39 | 4,00E-02 |
| R-HSA-1660661 | Sphingolipid de novo biosynthesis | VAPA | VAMP associated protein A | 9218 | up | 1,29 | 4,33E-02 |
| R-HSA-2022928 | HS-GAG biosynthesis | NDST1 | N-deacetylase and N-sulfotransferase 1 | 3340 | down | 1,59 | 2,40E-03 |
| R-HSA-2022928 | HS-GAG biosynthesis | SDC1 | syndecan 1 | 6382 | down | 2,17 | 1,61E-02 |
| R-HSA-2022928 | HS-GAG biosynthesis | SDC3 | syndecan 3 | 9672 | down | 1,68 | 1,97E-02 |
| R-HSA-3656237 | Defective EXT2 causes exostoses 2 | SDC1 | syndecan 1 | 6382 | down | 2,17 | 1,61E-02 |
| R-HSA-3656237 | Defective EXT2 causes exostoses 2 | SDC3 | syndecan 3 | 9672 | down | 1,68 | 1,97E-02 |
| R-HSA-74217 | Purine salvage | GMPR | guanosine monophosphate reductase | 2766 | down | 1,65 | 3,56E-02 |
| R-HSA-74217 | Purine salvage | APRT | adenine phosphoribosyltransferase | 353 | down | 1,25 | 1,64E-03 |
| R-HSA-3656253 | Defective EXT1 causes exostoses 1, TRPS2 and CH | SDC1 | syndecan 1 | 6382 | down | 2,17 | 1,61E-02 |
| R-HSA-3656253 | Defective EXT1 causes exostoses 1, TRPS2 and CH | SDC3 | syndecan 3 | 9672 | down | 1,68 | 1,97E-02 |
| R-HSA-190372 | FGFR3c ligand binding and activation | FGFR3 | fibroblast growth factor receptor 3 | 2261 | down | 1,70 | 4,60E-02 |
| R-HSA-190372 | FGFR3c ligand binding and activation | FGF2 | fibroblast growth factor 2 | 2247 | down | 5,35 | 3,09E-02 |
| R-HSA-2046106 | alpha-linolenic acid (ALA) metabolism | ACSL1 | acyl-CoA synthetase long chain family m | 2180 | down | 1,54 | 3,01E-02 |
| R-HSA-2046106 | alpha-linolenic acid (ALA) metabolism | FADS1 | fatty acid desaturase 1 | 3992 | down | 1,54 | 8,39E-03 |
| R-HSA-1253288 | Downregulation of ERBB4 signaling | ITCH | itchy E3 ubiquitin protein ligase | 83737 | up | 1,51 | 6,64E-03 |
| R-HSA-1253288 | Downregulation of ERBB4 signaling | NEDD4 | neural precursor cell expressed, develop | 4734 | up | 1,99 | 3,68E-03 |
| R-HSA-446210 | Synthesis of UDP-N-acetyl-glucosamine | PGM3 | phosphoglucomutase 3 | 5238 | up | 2,22 | 4,77E-03 |
| R-HSA-446210 | Synthesis of UDP-N-acetyl-glucosamine | GFPT1 | glutamine--fructose-6-phosphate transa | 2673 | up | 2,92 | 2,14E-03 |
| R-HSA-72731 | Recycling of eIF2:GDP | EIF2S2 | eukaryotic translation initiation factor 2 s | 8894 | up | 1,40 | 2,40E-02 |
| R-HSA-72731 | Recycling of eIF2:GDP | EIF2S3 | eukaryotic translation initiation factor 2 s | 1968 | up | 1,19 | 2,93E-02 |
| R-HSA-383280 | Nuclear Receptor transcription pathway | NR1D2 | nuclear receptor subfamily 1 group D me | 9975 | up | 2,02 | 4,50E-03 |
| R-HSA-383280 | Nuclear Receptor transcription pathway | THRA | thyroid hormone receptor, alpha | 7067 | down | 1,45 | 2,63E-02 |
| R-HSA-383280 | Nuclear Receptor transcription pathway | RXRA | retinoid X receptor alpha | 6256 | down | 1,34 | 3,67E-03 |
| R-HSA-383280 | Nuclear Receptor transcription pathway | NR2F1 | nuclear receptor subfamily 2 group F me | 7025 | down | 1,82 | 1,73E-02 |
| R-HSA-383280 | Nuclear Receptor transcription pathway | NR2F6 | nuclear receptor subfamily 2 group F me | 2063 | down | 1,48 | 1,28E-02 |
| R-HSA-383280 | Nuclear Receptor transcription pathway | NRBF2 | nuclear receptor binding factor 2 | 29982 | up | 2,19 | 2,71E-03 |
| R-HSA-456926 | Thrombin signalling through proteinase activated rece | ARRB1 | arrestin beta 1 | 408 | down | 1,79 | 2,42E-02 |
| R-HSA-456926 | Thrombin signalling through proteinase activated rece | GNA13 | G protein subunit alpha 13 | 10672 | up | 1,68 | 2,41E-03 |
| R-HSA-456926 | Thrombin signalling through proteinase activated rece | GNA11 | G protein subunit alpha 11 | 2767 | down | 1,18 | 4,37E-02 |
| R-HSA-456926 | Thrombin signalling through proteinase activated rece | F2R | coagulation factor II thrombin receptor | 2149 | down | 1,54 | 7,69E-03 |
| R-HSA-5358747 | S33 mutants of beta-catenin aren't phosphorylated | PPP2CB | protein phosphatase 2 catalytic subunit | 5516 | up | 1,30 | 4,79E-02 |
| R-HSA-5358747 | S33 mutants of beta-catenin aren't phosphorylated | AMER1 | APC membrane recruitment protein 1 | 139285 | down | 1,78 | 4,15E-02 |
| R-HSA-5358747 | S33 mutants of beta-catenin aren't phosphorylated | CSNK1A1 | casein kinase 1 alpha 1 | 1452 | up | 1,35 | 2,84E-02 |
| R-HSA-5339716 | Misspliced GSK3beta mutants stabilize beta-catenin | CSNK1A1 | casein kinase 1 alpha 1 | 1452 | up | 1,35 | 2,84E-02 |
| R-HSA-5339716 | Misspliced GSK3beta mutants stabilize beta-catenin | AMER1 | APC membrane recruitment protein 1 | 139285 | down | 1,78 | 4,15E-02 |
| R-HSA-5339716 | Misspliced GSK3beta mutants stabilize beta-catenin | PPP2CB | protein phosphatase 2 catalytic subunit | 5516 | up | 1,30 | 4,79E-02 |
| R-HSA-5358752 | T41 mutants of beta-catenin aren't phosphorylated | CSNK1A1 | casein kinase 1 alpha 1 | 1452 | up | 1,35 | 2,84E-02 |
| R-HSA-5358752 | T41 mutants of beta-catenin aren't phosphorylated | PPP2CB | protein phosphatase 2 catalytic subunit | 5516 | up | 1,30 | 4,79E-02 |
| R-HSA-5358752 | T41 mutants of beta-catenin aren't phosphorylated | AMER1 | APC membrane recruitment protein 1 | 139285 | down | 1,78 | 4,15E-02 |
| R-HSA-5358749 | S37 mutants of beta-catenin aren't phosphorylated | AMER1 | APC membrane recruitment protein 1 | 139285 | down | 1,78 | 4,15E-02 |
| R-HSA-5358749 | S37 mutants of beta-catenin aren't phosphorylated | PPP2CB | protein phosphatase 2 catalytic subunit | 5516 | up | 1,30 | 4,79E-02 |
| R-HSA-5358749 | S37 mutants of beta-catenin aren't phosphorylated | CSNK1A1 | casein kinase 1 alpha 1 | 1452 | up | 1,35 | 2,84E-02 |
| R-HSA-430039 | mRNA decay by 5' to 3' exoribonuclease | DCP1A | decapping mRNA 1A | 55802 | up | 1,33 | 4,54E-02 |
| R-HSA-430039 | mRNA decay by 5' to 3' exoribonuclease | XRN1 | 5'-3' exoribonuclease 1 | 54464 | up | 1,30 | 1,96E-02 |
| R-HSA-430039 | mRNA decay by 5' to 3' exoribonuclease | LSM4 | LSM4 homolog, U6 small nuclear RNA a | 25804 | down | 1,40 | 2,18E-02 |
| R-HSA-4419969 | Depolymerisation of the Nuclear Lamina | LMNB1 | lamin B1 | 4001 | down | 2,12 | 3,85E-02 |
| R-HSA-4419969 | Depolymerisation of the Nuclear Lamina | LPIN3 | lipin 3 | 64900 | down | 1,36 | 4,76E-02 |
| R-HSA-4419969 | Depolymerisation of the Nuclear Lamina | LPIN2 | lipin 2 | 9663 | up | 2,16 | 6,88E-03 |
| R-HSA-5358751 | S45 mutants of beta-catenin aren't phosphorylated | CSNK1A1 | casein kinase 1 alpha 1 | 1452 | up | 1,35 | 2,84E-02 |
| R-HSA-5358751 | S45 mutants of beta-catenin aren't phosphorylated | PPP2CB | protein phosphatase 2 catalytic subunit | 5516 | up | 1,30 | 4,79E-02 |
| R-HSA-5358751 | S45 mutants of beta-catenin aren't phosphorylated | AMER1 | APC membrane recruitment protein 1 | 139285 | down | 1,78 | 4,15E-02 |
| R-HSA-2691232 | Constitutive Signaling by NOTCH1 HD Domain Mutan | NOTCH1 | notch 1 | 4851 | down | 1,62 | 2,46E-02 |
| R-HSA-2691232 | Constitutive Signaling by NOTCH1 HD Domain Mutan | JAG1 | jagged 1 | 182 | up | 1,84 | 3,27E-02 |
| R-HSA-2691232 | Constitutive Signaling by NOTCH1 HD Domain Mutan | ADAM17 | ADAM metallopeptidase domain 17 | 6868 | up | 2,59 | 1,60E-03 |
| R-HSA-446193 | Biosynthesis of the N-glycan precursor (dolichol lipid-l | ALG3 | ALG3, alpha-1,3- mannosyltransferase | 10195 | down | 1,33 | 3,98E-02 |
| R-HSA-446193 | Biosynthesis of the N-glycan precursor (dolichol lipid-l | ALG2 | ALG2, alpha-1,3/1,6-mannosyltransfera | 85365 | up | 1,61 | 2,74E-02 |
| R-HSA-446193 | Biosynthesis of the N-glycan precursor (dolichol lipid-l | ALG1 | ALG1, chitobiosyldiphosphodolichol bet | 56052 | down | 1,35 | 3,85E-02 |
| R-HSA-140534 | Caspase activation via Death Receptors in the preser | TNFRSF10B | TNF receptor superfamily member 10b | 8795 | up | 2,77 | 1,99E-03 |
| R-HSA-140534 | Caspase activation via Death Receptors in the preser | FADD | Fas associated via death domain | 8772 | down | 1,31 | 8,90E-03 |
| R-HSA-140534 | Caspase activation via Death Receptors in the preser | TRAF2 | TNF receptor associated factor 2 | 7186 | down | 1,47 | 2,65E-02 |
| R-HSA-69656 | Cyclin A:Cdk2-associated events at S phase entry | CDK7 | cyclin dependent kinase 7 | 1022 | up | 1,37 | 4,38E-02 |
| R-HSA-69656 | Cyclin A:Cdk2-associated events at S phase entry | CDKN1A | cyclin dependent kinase inhibitor 1A | 1026 | up | 3,62 | 5,43E-03 |
| R-HSA-69656 | Cyclin A:Cdk2-associated events at S phase entry | CDC25A | cell division cycle 25A | 993 | down | 1,92 | 1,48E-02 |
| R-HSA-3304356 | SMAD2/3 Phosphorylation Motif Mutants in Cancer | TGFBR2 | transforming growth factor beta recepto | 7048 | down | 1,34 | 2,74E-02 |
| R-HSA-3304356 | SMAD2/3 Phosphorylation Motif Mutants in Cancer | SMAD2 | SMAD family member 2 | 4087 | up | 1,29 | 2,19E-02 |
| R-HSA-947581 | Molybdenum cofactor biosynthesis | MOCS3 | molybdenum cofactor synthesis 3 | 27304 | down | 1,34 | 6,80E-04 |
| R-HSA-947581 | Molybdenum cofactor biosynthesis | MOCOS | molybdenum cofactor sulfurase | 55034 | up | 1,70 | 2,54E-02 |
| R-HSA-6782861 | Synthesis of wybutosine at G37 of tRNA(Phe) | TYW5 | tRNA-yW synthesizing protein 5 | 129450 | up | 1,52 | 3,54E-02 |
| R-HSA-6782861 | Synthesis of wybutosine at G37 of tRNA(Phe) | LCMT2 | leucine carboxyl methyltransferase 2 | 9836 | down | 1,38 | 1,66E-02 |
| R-HSA-446199 | Synthesis of Dolichyl-phosphate | DOLK | dolichol kinase | 22845 | down | 1,73 | 2,74E-03 |
| R-HSA-446199 | Synthesis of Dolichyl-phosphate | NUS1 | NUS1 dehydrodolichyl diphosphate syn | 116150 | up | 1,54 | 4,27E-02 |
| R-HSA-2428928 | IRS-related events triggered by IGF1R | IRS1 | insulin receptor substrate 1 | 3667 | down | 1,89 | 6,25E-03 |
| R-HSA-2428928 | IRS-related events triggered by IGF1R | IRS2 | insulin receptor substrate 2 | 8660 | up | 2,21 | 2,22E-02 |
| R-HSA-210500 | Glutamate Neurotransmitter Release Cycle | PPFIA1 | PTPRF interacting protein alpha 1 | 8500 | up | 1,30 | 3,68E-02 |
| R-HSA-210500 | Glutamate Neurotransmitter Release Cycle | VAMP2 | vesicle associated membrane protein 2 | 6844 | up | 1,50 | 4,82E-02 |
| R-HSA-210500 | Glutamate Neurotransmitter Release Cycle | RAB3A | RAB3A, member RAS oncogene family | 5864 | down | 1,80 | 4,63E-02 |
| R-HSA-210500 | Glutamate Neurotransmitter Release Cycle | SLC38A2 | solute carrier family 38 member 2 | 54407 | up | 1,62 | 4,57E-02 |
| R-HSA-3270619 | IRF3-mediated induction of type I IFN | DTX4 | deltex E3 ubiquitin ligase 4 | 23220 | down | 1,52 | 4,63E-02 |
| R-HSA-3270619 | IRF3-mediated induction of type I IFN | IRF3 | interferon regulatory factor 3 | 3661 | down | 1,33 | 2,58E-03 |
| R-HSA-170670 | Adenylate cyclase inhibitory pathway | ADCY3 | adenylate cyclase 3 | 109 | down | 1,45 | 1,03E-02 |
| R-HSA-170670 | Adenylate cyclase inhibitory pathway | GNAI2 | G protein subunit alpha i2 | 2771 | down | 1,22 | 3,65E-02 |
| R-HSA-114608 | Platelet degranulation | CDC37L1 | cell division cycle 37 like 1 | 55664 | up | 1,53 | 1,69E-02 |
| R-HSA-114608 | Platelet degranulation | TUBA4A | tubulin alpha 4a | 7277 | down | 1,63 | 2,97E-02 |
| R-HSA-114608 | Platelet degranulation | PFN1 | profilin 1 | 5216 | down | 1,20 | 3,98E-02 |
| R-HSA-114608 | Platelet degranulation | VEGFA | vascular endothelial growth factor A | 7422 | up | 2,38 | 1,68E-02 |
| R-HSA-114608 | Platelet degranulation | GTPBP2 | GTP binding protein 2 | 54676 | up | 3,38 | 2,11E-03 |
| R-HSA-114608 | Platelet degranulation | TEX264 | testis expressed 264 | 51368 | down | 1,28 | 3,32E-02 |
| R-HSA-114608 | Platelet degranulation | MANF | mesencephalic astrocyte derived neurot | 7873 | up | 2,27 | 6,96E-03 |
| R-HSA-114608 | Platelet degranulation | CALM1 | calmodulin 1 | 801 | down | 1,43 | 2,23E-02 |
| R-HSA-114608 | Platelet degranulation | HABP4 | hyaluronan binding protein 4 | 22927 | up | 1,24 | 3,62E-02 |
| R-HSA-114608 | Platelet degranulation | CFL1 | cofilin 1 | 1072 | down | 1,29 | 3,02E-02 |
| R-HSA-114608 | Platelet degranulation | CALM2 | calmodulin 2 | 805 | down | 1,43 | 2,23E-02 |
| R-HSA-114608 | Platelet degranulation | LHFPL2 | LHFPL tetraspan subfamily member 2 | 10184 | up | 1,52 | 1,20E-02 |
| R-HSA-114608 | Platelet degranulation | CHID1 | chitinase domain containing 1 | 66005 | down | 1,27 | 2,24E-03 |
| R-HSA-114608 | Platelet degranulation | CALM3 | calmodulin 3 | 808 | down | 1,43 | 2,23E-02 |
| R-HSA-114608 | Platelet degranulation | HSPA5 | heat shock protein family A (Hsp70) mer | 3309 | up | 5,02 | 6,98E-04 |
| R-HSA-416482 | G alpha (12/13) signalling events | SOS2 | SOS Ras/Rho guanine nucleotide exch | 6655 | up | 1,82 | 1,41E-02 |
| R-HSA-416482 | G alpha (12/13) signalling events | ARHGEF2 | Rho/Rac guanine nucleotide exchange | 9181 | up | 2,09 | 1,10E-02 |
| R-HSA-416482 | G alpha (12/13) signalling events | PLXNB1 | plexin B1 | 5364 | down | 1,32 | 3,64E-02 |
| R-HSA-416482 | G alpha (12/13) signalling events | PIK3CA | phosphatidylinositol-4,5-bisphosphate 3 | 5290 | up | 1,33 | 3,17E-02 |
| R-HSA-416482 | G alpha (12/13) signalling events | PIK3R1 | phosphoinositide-3-kinase regulatory su | 5295 | down | 2,04 | 8,40E-03 |
| R-HSA-416482 | G alpha (12/13) signalling events | GNA13 | G protein subunit alpha 13 | 10672 | up | 1,68 | 2,41E-03 |
| R-HSA-416482 | G alpha (12/13) signalling events | FGD4 | FYVE, RhoGEF and PH domain contain | 121512 | up | 1,60 | 1,34E-02 |
| R-HSA-416482 | G alpha (12/13) signalling events | RHOB | ras homolog family member B | 388 | up | 1,65 | 2,44E-02 |
| R-HSA-416482 | G alpha (12/13) signalling events | ARHGEF19 | Rho guanine nucleotide exchange facto | 128272 | down | 1,55 | 1,82E-02 |
| R-HSA-983695 | Antigen activates B Cell Receptor (BCR) leading to ge | NCK1 | NCK adaptor protein 1 | 4690 | up | 1,48 | 1,42E-02 |
| R-HSA-983695 | Antigen activates B Cell Receptor (BCR) leading to ge | PIK3R1 | phosphoinositide-3-kinase regulatory su | 5295 | down | 2,04 | 8,40E-03 |
| R-HSA-983695 | Antigen activates B Cell Receptor (BCR) leading to ge | CBLB | Cbl proto-oncogene B | 868 | up | 1,83 | 4,35E-02 |
| R-HSA-983695 | Antigen activates B Cell Receptor (BCR) leading to ge | PLCG1 | phospholipase C gamma 1 | 5335 | down | 1,51 | 2,85E-02 |
| R-HSA-983695 | Antigen activates B Cell Receptor (BCR) leading to ge | CALM1 | calmodulin 1 | 801 | down | 1,43 | 2,23E-02 |
| R-HSA-983695 | Antigen activates B Cell Receptor (BCR) leading to ge | CALM3 | calmodulin 3 | 808 | down | 1,43 | 2,23E-02 |
| R-HSA-983695 | Antigen activates B Cell Receptor (BCR) leading to ge | CALM2 | calmodulin 2 | 805 | down | 1,43 | 2,23E-02 |
| R-HSA-983695 | Antigen activates B Cell Receptor (BCR) leading to ge | LYN | LYN proto-oncogene, Src family tyrosine | 4067 | down | 1,36 | 2,34E-02 |
| R-HSA-983695 | Antigen activates B Cell Receptor (BCR) leading to ge | DAPP1 | dual adaptor of phosphotyrosine and 3- | 27071 | up | 2,40 | 3,92E-02 |
| R-HSA-165054 | Rev-mediated nuclear export of HIV RNA | NUP58 | nucleoporin 58 | 9818 | up | 1,61 | 1,28E-02 |
| R-HSA-165054 | Rev-mediated nuclear export of HIV RNA | RCC1 | regulator of chromosome condensation | 1104 | down | 1,51 | 6,96E-03 |
| R-HSA-165054 | Rev-mediated nuclear export of HIV RNA | RANGAP1 | Ran GTPase activating protein 1 | 5905 | down | 1,48 | 1,41E-02 |
| R-HSA-165054 | Rev-mediated nuclear export of HIV RNA | NUP62 | nucleoporin 62 | 23636 | down | 1,52 | 4,39E-03 |
| R-HSA-445355 | Smooth Muscle Contraction | CALM1 | calmodulin 1 | 801 | down | 1,43 | 2,23E-02 |
| R-HSA-445355 | Smooth Muscle Contraction | ANXA1 | annexin A1 | 301 | up | 2,22 | 9,88E-03 |
| R-HSA-445355 | Smooth Muscle Contraction | CALM3 | calmodulin 3 | 808 | down | 1,43 | 2,23E-02 |
| R-HSA-445355 | Smooth Muscle Contraction | CALM2 | calmodulin 2 | 805 | down | 1,43 | 2,23E-02 |
| R-HSA-380270 | Recruitment of mitotic centrosome proteins and comp | TUBB | tubulin beta class I | 203068 | down | 1,56 | 2,67E-02 |
| R-HSA-380270 | Recruitment of mitotic centrosome proteins and comp | AKAP9 | A-kinase anchoring protein 9 | 10142 | up | 1,47 | 1,80E-02 |
| R-HSA-380270 | Recruitment of mitotic centrosome proteins and comp | CEP63 | centrosomal protein 63 | 80254 | up | 1,42 | 2,18E-02 |
| R-HSA-380270 | Recruitment of mitotic centrosome proteins and comp | CEP57 | centrosomal protein 57 | 9702 | up | 1,57 | 1,19E-02 |
| R-HSA-380270 | Recruitment of mitotic centrosome proteins and comp | PCM1 | pericentriolar material 1 | 5108 | up | 1,23 | 5,27E-03 |
| R-HSA-380270 | Recruitment of mitotic centrosome proteins and comp | CEP164 | centrosomal protein 164 | 22897 | up | 1,46 | 2,91E-02 |
| R-HSA-380270 | Recruitment of mitotic centrosome proteins and comp | TUBA4A | tubulin alpha 4a | 7277 | down | 1,63 | 2,97E-02 |
| R-HSA-380270 | Recruitment of mitotic centrosome proteins and comp | CEP290 | centrosomal protein 290 | 80184 | up | 1,63 | 1,81E-02 |
| R-HSA-380270 | Recruitment of mitotic centrosome proteins and comp | TUBG1 | tubulin gamma 1 | 7283 | down | 1,50 | 1,42E-02 |
| R-HSA-5654228 | Phospholipase C-mediated cascade; FGFR4 | FGF2 | fibroblast growth factor 2 | 2247 | down | 5,35 | 3,09E-02 |
| R-HSA-5654228 | Phospholipase C-mediated cascade; FGFR4 | PLCG1 | phospholipase C gamma 1 | 5335 | down | 1,51 | 2,85E-02 |
| R-HSA-193807 | Synthesis of bile acids and bile salts via 27-hydroxych | HSD3B7 | hydroxy-delta-5-steroid dehydrogenase, | 80270 | down | 1,64 | 3,72E-02 |
| R-HSA-193807 | Synthesis of bile acids and bile salts via 27-hydroxych | RXRA | retinoid X receptor alpha | 6256 | down | 1,34 | 3,67E-03 |
| R-HSA-162710 | Synthesis of glycosylphosphatidylinositol (GPI) | PIGM | phosphatidylinositol glycan anchor biosy | 93183 | down | 1,39 | 4,66E-02 |
| R-HSA-162710 | Synthesis of glycosylphosphatidylinositol (GPI) | PIGA | phosphatidylinositol glycan anchor biosy | 5277 | up | 2,32 | 6,64E-03 |
| R-HSA-162710 | Synthesis of glycosylphosphatidylinositol (GPI) | PIGW | phosphatidylinositol glycan anchor biosy | 284098 | down | 1,53 | 1,78E-02 |
| R-HSA-1433559 | Regulation of KIT signaling | LYN | LYN proto-oncogene, Src family tyrosine | 4067 | down | 1,36 | 2,34E-02 |
| R-HSA-1433559 | Regulation of KIT signaling | YES1 | YES proto-oncogene 1, Src family tyros | 7525 | up | 1,30 | 2,08E-02 |
| R-HSA-1433559 | Regulation of KIT signaling | SH2B2 | SH2B adaptor protein 2 | 10603 | down | 2,07 | 1,95E-02 |
| R-HSA-1963640 | GRB2 events in ERBB2 signaling | HBEGF | heparin binding EGF like growth factor | 1839 | up | 3,67 | 5,67E-03 |
| R-HSA-1963640 | GRB2 events in ERBB2 signaling | ERBB2 | erb-b2 receptor tyrosine kinase 2 | 2064 | down | 1,47 | 1,93E-02 |
| R-HSA-1963640 | GRB2 events in ERBB2 signaling | HRAS | HRas proto-oncogene, GTPase | 3265 | down | 1,31 | 2,19E-03 |
| R-HSA-532668 | N-glycan trimming in the ER and Calnexin/Calreticulin | ENGASE | endo-beta-N-acetylglucosaminidase | 64772 | down | 1,38 | 3,04E-02 |
| R-HSA-532668 | N-glycan trimming in the ER and Calnexin/Calreticulin | RAD23B | RAD23 homolog B, nucleotide excision | 5887 | up | 1,23 | 3,34E-02 |
| R-HSA-532668 | N-glycan trimming in the ER and Calnexin/Calreticulin | NGLY1 | N-glycanase 1 | 55768 | up | 1,38 | 1,27E-02 |
| R-HSA-110312 | Translesion synthesis by REV1 | RPA1 | replication protein A1 | 6117 | down | 1,41 | 4,93E-02 |
| R-HSA-110312 | Translesion synthesis by REV1 | RFC1 | replication factor C subunit 1 | 5981 | up | 1,15 | 2,18E-02 |
| R-HSA-110312 | Translesion synthesis by REV1 | PCNA | proliferating cell nuclear antigen | 5111 | down | 2,14 | 4,63E-02 |
| R-HSA-983231 | Factors involved in megakaryocyte development and | JMJD1C | jumonji domain containing 1C | 221037 | up | 1,87 | 6,38E-03 |
| R-HSA-983231 | Factors involved in megakaryocyte development and | SH2B2 | SH2B adaptor protein 2 | 10603 | down | 2,07 | 1,95E-02 |
| R-HSA-983231 | Factors involved in megakaryocyte development and | SH2B1 | SH2B adaptor protein 1 | 25970 | down | 1,31 | 1,81E-02 |
| R-HSA-983231 | Factors involved in megakaryocyte development and | RAB5A | RAB5A, member RAS oncogene family | 5868 | up | 1,78 | 3,10E-03 |
| R-HSA-983231 | Factors involved in megakaryocyte development and | MAFG | MAF bZIP transcription factor G | 4097 | up | 1,78 | 3,51E-02 |
| R-HSA-983231 | Factors involved in megakaryocyte development and | PRKAR1B | protein kinase cAMP-dependent type I r | 5575 | down | 1,50 | 6,80E-04 |
| R-HSA-983231 | Factors involved in megakaryocyte development and | IRF3 | interferon regulatory factor 3 | 3661 | down | 1,33 | 2,58E-03 |
| R-HSA-983231 | Factors involved in megakaryocyte development and | GATA2 | GATA binding protein 2 | 2624 | down | 2,16 | 2,07E-02 |
| R-HSA-983231 | Factors involved in megakaryocyte development and | MAFF | MAF bZIP transcription factor F | 23764 | up | 2,78 | 1,03E-02 |
| R-HSA-983231 | Factors involved in megakaryocyte development and | AKAP1 | A-kinase anchoring protein 1 | 8165 | down | 1,38 | 4,54E-02 |
| R-HSA-983231 | Factors involved in megakaryocyte development and | GATA3 | GATA binding protein 3 | 2625 | down | 2,92 | 3,45E-02 |
| R-HSA-1268020 | Mitochondrial protein import | TIMM21 | translocase of inner mitochondrial memb | 29090 | down | 1,56 | 1,69E-02 |
| R-HSA-1268020 | Mitochondrial protein import | TIMM22 | translocase of inner mitochondrial memb | 29928 | down | 1,18 | 2,54E-02 |
| R-HSA-1268020 | Mitochondrial protein import | PMPCA | peptidase, mitochondrial processing alp | 23203 | down | 1,26 | 2,77E-02 |
| R-HSA-1268020 | Mitochondrial protein import | HSPA9 | heat shock protein family A (Hsp70) mer | 3313 | up | 1,59 | 5,68E-03 |
| R-HSA-1268020 | Mitochondrial protein import | TOMM70 | translocase of outer mitochondrial memb | 9868 | up | 1,24 | 3,74E-02 |
| R-HSA-1268020 | Mitochondrial protein import | TIMM9 | translocase of inner mitochondrial memb | 26520 | up | 1,31 | 2,36E-02 |
| R-HSA-1268020 | Mitochondrial protein import | TOMM40 | translocase of outer mitochondrial memb | 10452 | down | 1,29 | 4,09E-04 |
| R-HSA-72187 | mRNA 3'-end processing | SRSF2 | serine and arginine rich splicing factor 2 | 6427 | down | 1,69 | 1,79E-02 |
| R-HSA-72187 | mRNA 3'-end processing | U2AF2 | U2 small nuclear RNA auxiliary factor 2 | 11338 | down | 1,27 | 8,21E-03 |
| R-HSA-72187 | mRNA 3'-end processing | CSTF2 | cleavage stimulation factor subunit 2 | 1478 | down | 1,45 | 4,38E-02 |
| R-HSA-72187 | mRNA 3'-end processing | SARNP | SAP domain containing ribonucleoprote | 84324 | up | 1,48 | 4,35E-02 |
| R-HSA-72187 | mRNA 3'-end processing | CLP1 | cleavage and polyadenylation factor I s | 10978 | down | 1,40 | 1,04E-02 |
| R-HSA-72187 | mRNA 3'-end processing | SLU7 | SLU7 homolog, splicing factor | 10569 | up | 1,24 | 3,63E-02 |
| R-HSA-72649 | Translation initiation complex formation | RPS4Y1 | ribosomal protein S4, Y-linked 1 | 6192 | down | 5,59 | 7,82E-03 |
| R-HSA-72649 | Translation initiation complex formation | EIF4A2 | eukaryotic translation initiation factor 4A | 1974 | up | 1,68 | 4,70E-03 |
| R-HSA-72649 | Translation initiation complex formation | EIF2S3 | eukaryotic translation initiation factor 2 s | 1968 | up | 1,19 | 2,93E-02 |
| R-HSA-72649 | Translation initiation complex formation | EIF3M | eukaryotic translation initiation factor 3 s | 10480 | up | 1,25 | 2,51E-02 |
| R-HSA-72649 | Translation initiation complex formation | EIF2S2 | eukaryotic translation initiation factor 2 s | 8894 | up | 1,40 | 2,40E-02 |
| R-HSA-72649 | Translation initiation complex formation | EIF3B | eukaryotic translation initiation factor 3 s | 8662 | down | 1,27 | 1,76E-02 |
| R-HSA-72649 | Translation initiation complex formation | EIF3H | eukaryotic translation initiation factor 3 s | 8667 | up | 1,22 | 3,84E-02 |
| R-HSA-72702 | Ribosomal scanning and start codon recognition | EIF3H | eukaryotic translation initiation factor 3 s | 8667 | up | 1,22 | 3,84E-02 |
| R-HSA-72702 | Ribosomal scanning and start codon recognition | EIF3B | eukaryotic translation initiation factor 3 s | 8662 | down | 1,27 | 1,76E-02 |
| R-HSA-72702 | Ribosomal scanning and start codon recognition | EIF2S2 | eukaryotic translation initiation factor 2 s | 8894 | up | 1,40 | 2,40E-02 |
| R-HSA-72702 | Ribosomal scanning and start codon recognition | EIF3M | eukaryotic translation initiation factor 3 s | 10480 | up | 1,25 | 2,51E-02 |
| R-HSA-72702 | Ribosomal scanning and start codon recognition | EIF2S3 | eukaryotic translation initiation factor 2 s | 1968 | up | 1,19 | 2,93E-02 |
| R-HSA-72702 | Ribosomal scanning and start codon recognition | EIF4A2 | eukaryotic translation initiation factor 4A | 1974 | up | 1,68 | 4,70E-03 |
| R-HSA-72702 | Ribosomal scanning and start codon recognition | RPS4Y1 | ribosomal protein S4, Y-linked 1 | 6192 | down | 5,59 | 7,82E-03 |
| R-HSA-162588 | Budding and maturation of HIV virion | VPS37A | VPS37A, ESCRT-I subunit | 137492 | up | 1,48 | 1,14E-02 |
| R-HSA-162588 | Budding and maturation of HIV virion | VPS37D | VPS37D, ESCRT-I subunit | 155382 | up | 2,22 | 5,83E-03 |
| R-HSA-162588 | Budding and maturation of HIV virion | CHMP5 | charged multivesicular body protein 5 | 51510 | up | 1,32 | 3,37E-02 |
| R-HSA-1660517 | Synthesis of PIPs at the late endosome membrane | MTMR2 | myotubularin related protein 2 | 8898 | up | 1,23 | 2,61E-02 |
| R-HSA-1660517 | Synthesis of PIPs at the late endosome membrane | PIK3C2A | phosphatidylinositol-4-phosphate 3-kina | 5286 | up | 1,30 | 1,79E-02 |
| R-HSA-193692 | Regulated proteolysis of p75NTR | NGFR | nerve growth factor receptor | 4804 | up | 12,35 | 2,75E-03 |

| | | | | | | | |
|---|---|---|---|---|---|---|---|
| R-HSA-193692 | Regulated proteolysis of p75NTR | ADAM17 | ADAM metallopeptidase domain 17 | 6868 | up | 2,59 | 1,60E-03 |
| R-HSA-1234158 | Regulation of gene expression by Hypoxia-inducible F | VEGFA | vascular endothelial growth factor A | 7422 | up | 2,38 | 1,68E-02 |
| R-HSA-1234158 | Regulation of gene expression by Hypoxia-inducible F | HIF1A | hypoxia inducible factor 1 alpha subunit | 3091 | up | 1,23 | 4,66E-02 |
| R-HSA-450341 | Activation of the AP-1 family of transcription factors | ATF2 | activating transcription factor 2 | 1386 | up | 1,79 | 1,01E-02 |
| R-HSA-450341 | Activation of the AP-1 family of transcription factors | MAPK8 | mitogen-activated protein kinase 8 | 5599 | up | 1,38 | 3,05E-02 |
| R-HSA-419812 | Calcitonin-like ligand receptors | ADM2 | adrenomedullin 2 | 79924 | up | 3,79 | 7,16E-03 |
| R-HSA-419812 | Calcitonin-like ligand receptors | ADM | adrenomedullin | 133 | up | 1,84 | 4,75E-04 |
| R-HSA-2470946 | Cohesin Loading onto Chromatin | NIPBL | NIPBL, cohesin loading factor | 25836 | up | 1,30 | 4,13E-02 |
| R-HSA-2470946 | Cohesin Loading onto Chromatin | STAG2 | stromal antigen 2 | 10735 | up | 1,26 | 3,69E-02 |
| R-HSA-163615 | PKA activation | ADCY3 | adenylate cyclase 3 | 109 | down | 1,45 | 1,03E-02 |
| R-HSA-163615 | PKA activation | PRKAR1B | protein kinase cAMP-dependent type I r | 5575 | down | 1,50 | 6,80E-04 |
| R-HSA-2173791 | TGF-beta receptor signaling in EMT (epithelial to mes | PARD6A | par-6 family cell polarity regulator alpha | 50855 | down | 1,46 | 8,78E-03 |
| R-HSA-2173791 | TGF-beta receptor signaling in EMT (epithelial to mes | TGFBR2 | transforming growth factor beta receptor | 7048 | down | 1,34 | 2,74E-02 |
| R-HSA-399956 | CRMPs in Sema3A signaling | PLXNA1 | plexin A1 | 5361 | down | 1,28 | 9,70E-04 |
| R-HSA-399956 | CRMPs in Sema3A signaling | DPYSL2 | dihydropyrimidinase like 2 | 1808 | down | 1,77 | 2,19E-02 |
| R-HSA-5654219 | Phospholipase C-mediated cascade: FGFR1 | FGF2 | fibroblast growth factor 2 | 2247 | down | 5,35 | 3,09E-02 |
| R-HSA-5654219 | Phospholipase C-mediated cascade: FGFR1 | PLCG1 | phospholipase C gamma 1 | 5335 | down | 1,51 | 2,85E-02 |
| R-HSA-77595 | Processing of Intronless Pre-mRNAs | CSTF2 | cleavage stimulation factor subunit 2 | 1478 | down | 1,45 | 4,38E-02 |
| R-HSA-77595 | Processing of Intronless Pre-mRNAs | CLP1 | cleavage and polyadenylation factor I s | 10978 | down | 1,40 | 1,04E-02 |
| R-HSA-114604 | GPVI-mediated activation cascade | PIK3CA | phosphatidylinositol-4,5-bisphosphate 3 | 5290 | up | 1,33 | 3,17E-02 |
| R-HSA-114604 | GPVI-mediated activation cascade | AKT1 | AKT serine/threonine kinase 1 | 207 | down | 1,37 | 1,42E-03 |
| R-HSA-114604 | GPVI-mediated activation cascade | PIK3R1 | phosphoinositide-3-kinase regulatory su | 5295 | down | 2,04 | 8,40E-03 |
| R-HSA-114604 | GPVI-mediated activation cascade | AKT2 | AKT serine/threonine kinase 2 | 208 | down | 1,28 | 6,33E-03 |
| R-HSA-114604 | GPVI-mediated activation cascade | JAK1 | Janus kinase 1 | 3716 | up | 1,53 | 2,13E-02 |
| R-HSA-114604 | GPVI-mediated activation cascade | LYN | LYN proto-oncogene, Src family tyrosine | 4067 | down | 1,36 | 2,34E-02 |
| R-HSA-114604 | GPVI-mediated activation cascade | RHOB | ras homolog family member B | 388 | up | 1,65 | 2,44E-02 |
| R-HSA-5578775 | Ion homeostasis | ATP2A3 | ATPase sarcoplasmic/endoplasmic retic | 489 | down | 1,62 | 4,34E-02 |
| R-HSA-5578775 | Ion homeostasis | CALM3 | calmodulin 3 | 808 | down | 1,43 | 2,23E-02 |
| R-HSA-5578775 | Ion homeostasis | CALM2 | calmodulin 2 | 805 | down | 1,43 | 2,23E-02 |
| R-HSA-5578775 | Ion homeostasis | CAMK2D | calcium/calmodulin dependent protein k | 817 | up | 1,29 | 1,49E-02 |
| R-HSA-5578775 | Ion homeostasis | ATP2B1 | ATPase plasma membrane Ca2+ transp | 490 | up | 1,35 | 4,17E-02 |
| R-HSA-5578775 | Ion homeostasis | CALM1 | calmodulin 1 | 801 | down | 1,43 | 2,23E-02 |
| R-HSA-2514859 | Inactivation, recovery and regulation of the phototran | FNTA | farnesyltransferase, CAAX box, alpha | 2339 | up | 1,33 | 3,73E-03 |
| R-HSA-2514859 | Inactivation, recovery and regulation of the phototran | CALM1 | calmodulin 1 | 801 | down | 1,43 | 2,23E-02 |
| R-HSA-2514859 | Inactivation, recovery and regulation of the phototran | CALM2 | calmodulin 2 | 805 | down | 1,43 | 2,23E-02 |
| R-HSA-2514859 | Inactivation, recovery and regulation of the phototran | CALM3 | calmodulin 3 | 808 | down | 1,43 | 2,23E-02 |
| R-HSA-75893 | TNF signaling | TRAF2 | TNF receptor associated factor 2 | 7186 | down | 1,47 | 2,65E-02 |
| R-HSA-75893 | TNF signaling | ADAM17 | ADAM metallopeptidase domain 17 | 6868 | up | 2,59 | 1,60E-03 |
| R-HSA-3656532 | TGFBR1 KD Mutants in Cancer | TGFBR2 | transforming growth factor beta receptor | 7048 | down | 1,34 | 2,74E-02 |
| R-HSA-3656532 | TGFBR1 KD Mutants in Cancer | SMAD2 | SMAD family member 2 | 4087 | up | 1,29 | 2,19E-02 |
| R-HSA-442720 | CREB1 phosphorylation through the activation of Ade | ADCY3 | adenylate cyclase 3 | 109 | down | 1,45 | 1,03E-02 |
| R-HSA-442720 | CREB1 phosphorylation through the activation of Ade | CREB1 | cAMP responsive element binding prote | 1385 | up | 1,51 | 7,18E-03 |
| R-HSA-75158 | TRAIL signaling | FADD | Fas associated via death domain | 8772 | down | 1,31 | 8,90E-03 |
| R-HSA-75158 | TRAIL signaling | TNFRSF10B | TNF receptor superfamily member 10b | 8795 | up | 2,77 | 1,99E-03 |
| R-HSA-3315487 | SMAD2/3 MH2 Domain Mutants in Cancer | TGFBR2 | transforming growth factor beta receptor | 7048 | down | 1,34 | 2,74E-02 |
| R-HSA-3315487 | SMAD2/3 MH2 Domain Mutants in Cancer | SMAD2 | SMAD family member 2 | 4087 | up | 1,29 | 2,19E-02 |
| R-HSA-420597 | Nectin/Necl trans heterodimerization | NECTIN1 | nectin cell adhesion molecule 1 | 5818 | down | 1,48 | 7,71E-03 |
| R-HSA-420597 | Nectin/Necl trans heterodimerization | PVR | poliovirus receptor | 5817 | up | 1,67 | 4,17E-02 |
| R-HSA-2468052 | Establishment of Sister Chromatid Cohesion | ESCO1 | establishment of sister chromatid cohes | 114799 | up | 1,52 | 2,97E-02 |
| R-HSA-2468052 | Establishment of Sister Chromatid Cohesion | STAG2 | stromal antigen 2 | 10735 | up | 1,26 | 3,69E-02 |
| R-HSA-5625970 | RHO GTPases activate KTN1 | KIF5B | kinesin family member 5B | 3799 | up | 1,40 | 7,18E-03 |
| R-HSA-5625970 | RHO GTPases activate KTN1 | KTN1 | kinectin 1 | 3895 | up | 1,35 | 2,24E-03 |
| R-HSA-351906 | Apoptotic cleavage of cell adhesion proteins | DSG2 | desmoglein 2 | 1829 | up | 1,19 | 2,34E-02 |
| R-HSA-351906 | Apoptotic cleavage of cell adhesion proteins | TJP1 | tight junction protein 1 | 7082 | up | 1,46 | 8,91E-03 |
| R-HSA-174490 | Membrane binding and targetting of GAG proteins | VPS37D | VPS37D, ESCRT-I subunit | 155382 | up | 2,22 | 5,83E-03 |
| R-HSA-174490 | Membrane binding and targetting of GAG proteins | VPS37A | VPS37A, ESCRT-I subunit | 137492 | up | 1,48 | 1,14E-02 |
| R-HSA-3214858 | RMTs methylate histone arginines | H2AFX | H2A histone family member X | 3014 | down | 2,11 | 1,30E-03 |
| R-HSA-3214858 | RMTs methylate histone arginines | PRMT7 | protein arginine methyltransferase 7 | 54496 | down | 1,44 | 1,87E-02 |
| R-HSA-3214858 | RMTs methylate histone arginines | SMARCD1 | SWI/SNF related, matrix associated, act | 6602 | down | 1,23 | 2,52E-02 |
| R-HSA-3214858 | RMTs methylate histone arginines | WDR5 | WD repeat domain 5 | 11091 | down | 1,28 | 3,81E-03 |
| R-HSA-3214858 | RMTs methylate histone arginines | CARM1 | coactivator associated arginine methyltr | 10498 | down | 1,36 | 1,60E-02 |
| R-HSA-5656121 | Translesion synthesis by POLI | PCNA | proliferating cell nuclear antigen | 5111 | down | 2,14 | 4,63E-02 |
| R-HSA-5656121 | Translesion synthesis by POLI | RFC1 | replication factor C subunit 1 | 5981 | up | 1,15 | 2,18E-02 |
| R-HSA-5656121 | Translesion synthesis by POLI | RPA1 | replication protein A1 | 6117 | down | 1,41 | 4,93E-02 |
| R-HSA-264642 | Acetylcholine Neurotransmitter Release Cycle | PPFIA1 | PTPRF interacting protein alpha 1 | 8500 | up | 1,30 | 3,68E-02 |
| R-HSA-264642 | Acetylcholine Neurotransmitter Release Cycle | VAMP2 | vesicle associated membrane protein 2 | 6844 | up | 1,50 | 4,82E-02 |
| R-HSA-264642 | Acetylcholine Neurotransmitter Release Cycle | RAB3A | RAB3A, member RAS oncogene family | 5864 | down | 1,80 | 4,63E-02 |
| R-HSA-5655862 | Translesion synthesis by POLK | PCNA | proliferating cell nuclear antigen | 5111 | down | 2,14 | 4,63E-02 |
| R-HSA-5655862 | Translesion synthesis by POLK | RFC1 | replication factor C subunit 1 | 5981 | up | 1,15 | 2,18E-02 |
| R-HSA-5655862 | Translesion synthesis by POLK | RPA1 | replication protein A1 | 6117 | down | 1,41 | 4,93E-02 |
| R-HSA-164378 | PKA activation in glucagon signalling | ADCY3 | adenylate cyclase 3 | 109 | down | 1,45 | 1,03E-02 |
| R-HSA-164378 | PKA activation in glucagon signalling | PRKAR1B | protein kinase cAMP-dependent type I r | 5575 | down | 1,50 | 6,80E-04 |
| R-HSA-5696394 | DNA Damage Recognition in GG-NER | PARP1 | poly(ADP-ribose) polymerase 1 | 142 | down | 1,50 | 3,92E-02 |
| R-HSA-5696394 | DNA Damage Recognition in GG-NER | ACTR5 | ARP5 actin related protein 5 homolog | 79913 | down | 1,54 | 9,20E-03 |
| R-HSA-5696394 | DNA Damage Recognition in GG-NER | INO80E | INO80 complex subunit E | 283899 | down | 1,19 | 2,67E-02 |
| R-HSA-5696394 | DNA Damage Recognition in GG-NER | RAD23B | RAD23 homolog B, nucleotide excision | 5887 | up | 1,23 | 3,34E-02 |
| R-HSA-5696394 | DNA Damage Recognition in GG-NER | INO80 | INO80 complex subunit | 54617 | up | 1,36 | 1,66E-02 |
| R-HSA-1475029 | Reversible hydration of carbon dioxide | CA13 | carbonic anhydrase 13 | 377677 | up | 1,89 | 7,37E-03 |
| R-HSA-1475029 | Reversible hydration of carbon dioxide | CA5B | carbonic anhydrase 5B | 11238 | up | 1,74 | 2,94E-02 |
| R-HSA-71064 | Lysine catabolism | DLD | dihydrolipoamide dehydrogenase | 1738 | up | 1,16 | 4,53E-02 |
| R-HSA-71064 | Lysine catabolism | AASS | aminoadipate-semialdehyde synthase | 10157 | up | 2,01 | 4,95E-02 |
| R-HSA-73621 | Pyrimidine catabolism | UPP1 | uridine phosphorylase 1 | 7378 | up | 4,41 | 1,62E-03 |
| R-HSA-73621 | Pyrimidine catabolism | NT5C3A | 5'-nucleotidase, cytosolic IIIA | 51251 | up | 1,49 | 1,52E-02 |
| R-HSA-390471 | Association of TriC/CCT with target proteins during bio | NOP56 | NOP56 ribonucleoprotein | 10528 | down | 1,40 | 2,84E-02 |
| R-HSA-390471 | Association of TriC/CCT with target proteins during bio | WRAP53 | WD repeat containing antisense to TP53 | 55135 | down | 1,52 | 1,69E-02 |
| R-HSA-390471 | Association of TriC/CCT with target proteins during bio | SPHK1 | sphingosine kinase 1 | 8877 | down | 2,00 | 2,34E-02 |
| R-HSA-5676594 | TNF receptor superfamily (TNFSF) members mediating | TRAF3 | TNF receptor associated factor 3 | 7187 | down | 1,40 | 1,36E-02 |
| R-HSA-5676594 | TNF receptor superfamily (TNFSF) members mediating | TRAF2 | TNF receptor associated factor 2 | 7186 | down | 1,47 | 2,65E-02 |
| R-HSA-5676594 | TNF receptor superfamily (TNFSF) members mediating | BIRC2 | baculoviral IAP repeat containing 2 | 329 | up | 1,38 | 1,86E-02 |
| R-HSA-5654221 | Phospholipase C-mediated cascade; FGFR2 | FGF2 | fibroblast growth factor 2 | 2247 | down | 5,35 | 3,09E-02 |
| R-HSA-5654221 | Phospholipase C-mediated cascade; FGFR2 | PLCG1 | phospholipase C gamma 1 | 5335 | down | 1,51 | 2,85E-02 |
| R-HSA-210993 | Tie2 Signaling | PIK3R1 | phosphoinositide-3-kinase regulatory su | 5295 | down | 2,04 | 8,40E-03 |
| R-HSA-210993 | Tie2 Signaling | PIK3CA | phosphatidylinositol-4,5-bisphosphate 3 | 5290 | up | 1,33 | 3,17E-02 |
| R-HSA-210993 | Tie2 Signaling | HRAS | HRas proto-oncogene, GTPase | 3265 | down | 1,31 | 2,19E-03 |
| R-HSA-209560 | NF-kB is activated and signals survival | NGFR | nerve growth factor receptor | 4804 | up | 12,35 | 2,75E-03 |
| R-HSA-209560 | NF-kB is activated and signals survival | SQSTM1 | sequestosome 1 | 8878 | up | 1,88 | 2,42E-02 |
| R-HSA-5362798 | Release of Hh-Np from the secreting cell | ADAM17 | ADAM metallopeptidase domain 17 | 6868 | up | 2,59 | 1,60E-03 |
| R-HSA-5362798 | Release of Hh-Np from the secreting cell | DISP2 | dispatched RND transporter family mem | 85455 | down | 1,57 | 3,64E-02 |
| R-HSA-5358493 | Synthesis of diphthamide-EEF2 | DPH6 | diphthamine biosynthesis 6 | 89978 | up | 1,26 | 3,02E-02 |
| R-HSA-5358493 | Synthesis of diphthamide-EEF2 | DPH2 | DPH2 homolog | 1802 | down | 1,33 | 6,98E-04 |
| R-HSA-196783 | Coenzyme A biosynthesis | SLC25A16 | solute carrier family 25 member 16 | 8034 | up | 1,41 | 4,87E-02 |
| R-HSA-196783 | Coenzyme A biosynthesis | PANK4 | pantothenate kinase 4 | 55229 | down | 1,39 | 4,37E-02 |
| R-HSA-425410 | Metal ion SLC transporters | SLC31A1 | solute carrier family 31 member 1 | 1317 | up | 1,86 | 1,00E-02 |
| R-HSA-425410 | Metal ion SLC transporters | SLC40A1 | solute carrier family 40 member 1 | 30061 | down | 1,80 | 2,30E-02 |
| R-HSA-416550 | Sema4D mediated inhibition of cell attachment and m | PLXNB1 | plexin B1 | 5364 | down | 1,32 | 3,64E-02 |
| R-HSA-416550 | Sema4D mediated inhibition of cell attachment and m | RND1 | Rho family GTPase 1 | 27289 | up | 4,86 | 3,18E-03 |
| R-HSA-390918 | Peroxisomal lipid metabolism | MLYCD | malonyl-CoA decarboxylase | 23417 | down | 1,35 | 2,54E-02 |
| R-HSA-390918 | Peroxisomal lipid metabolism | PECR | peroxisomal trans-2-enoyl-CoA reductas | 55825 | up | 1,76 | 4,63E-02 |
| R-HSA-3214842 | HDMs demethylate histones | KDM3A | lysine demethylase 3A | 55818 | up | 1,75 | 2,45E-02 |
| R-HSA-3214842 | HDMs demethylate histones | JMJD6 | arginine demethylase and lysine hydrox | 23210 | up | 1,37 | 4,71E-02 |
| R-HSA-3214842 | HDMs demethylate histones | KDM6B | lysine demethylase 6B | 23135 | up | 2,16 | 2,06E-02 |
| R-HSA-3214842 | HDMs demethylate histones | KDM7A | lysine demethylase 7A | 80853 | up | 1,89 | 3,97E-02 |
| R-HSA-3214842 | HDMs demethylate histones | ARID5B | AT-rich interaction domain 5B | 84159 | down | 2,09 | 3,59E-02 |
| R-HSA-3214842 | HDMs demethylate histones | KDM2B | lysine demethylase 2B | 84678 | down | 1,53 | 2,57E-02 |
| R-HSA-181430 | Norepinephrine Neurotransmitter Release Cycle | VAMP2 | vesicle associated membrane protein 2 | 6844 | up | 1,50 | 4,82E-02 |
| R-HSA-181430 | Norepinephrine Neurotransmitter Release Cycle | PPFIA1 | PTPRF interacting protein alpha 1 | 8500 | up | 1,30 | 3,68E-02 |
| R-HSA-181430 | Norepinephrine Neurotransmitter Release Cycle | RAB3A | RAB3A, member RAS oncogene family | 5864 | down | 1,80 | 4,63E-02 |
| R-HSA-181429 | Serotonin Neurotransmitter Release Cycle | RAB3A | RAB3A, member RAS oncogene family | 5864 | down | 1,80 | 4,63E-02 |
| R-HSA-181429 | Serotonin Neurotransmitter Release Cycle | PPFIA1 | PTPRF interacting protein alpha 1 | 8500 | up | 1,30 | 3,68E-02 |
| R-HSA-181429 | Serotonin Neurotransmitter Release Cycle | VAMP2 | vesicle associated membrane protein 2 | 6844 | up | 1,50 | 4,82E-02 |
| R-HSA-5627117 | RHO GTPases Activate ROCKs | PPP1R12A | protein phosphatase 1 regulatory subur | 4659 | up | 1,25 | 1,03E-02 |
| R-HSA-5627117 | RHO GTPases Activate ROCKs | RHOB | ras homolog family member B | 388 | up | 1,65 | 2,44E-02 |
| R-HSA-5627117 | RHO GTPases Activate ROCKs | LIMK1 | LIM domain kinase 1 | 3984 | down | 1,44 | 6,49E-03 |
| R-HSA-3214847 | HATs acetylate histones | ING5 | inhibitor of growth family member 5 | 84289 | down | 1,31 | 4,38E-02 |
| R-HSA-3214847 | HATs acetylate histones | ATF2 | activating transcription factor 2 | 1386 | up | 1,79 | 1,01E-02 |
| R-HSA-3214847 | HATs acetylate histones | HCFC1 | host cell factor C1 | 3054 | down | 1,33 | 4,82E-04 |
| R-HSA-3214847 | HATs acetylate histones | ING3 | inhibitor of growth family member 3 | 54556 | up | 1,67 | 2,14E-02 |
| R-HSA-3214847 | HATs acetylate histones | JADE2 | jade family PHD finger 2 | 23338 | down | 1,80 | 8,91E-03 |
| R-HSA-3214847 | HATs acetylate histones | EPC1 | enhancer of polycomb homolog 1 | 80314 | up | 1,40 | 2,88E-02 |
| R-HSA-3214847 | HATs acetylate histones | ZZZ3 | zinc finger ZZ-type containing 3 | 26009 | up | 1,44 | 3,36E-03 |
| R-HSA-3214847 | HATs acetylate histones | SUPT7L | SPT7 like, STAGA complex gamma sub | 9913 | up | 1,30 | 3,94E-02 |
| R-HSA-3214847 | HATs acetylate histones | KAT6B | lysine acetyltransferase 6B | 23522 | up | 1,44 | 4,05E-02 |
| R-HSA-3214847 | HATs acetylate histones | ELP2 | elongator acetyltransferase complex sub | 55250 | up | 1,26 | 4,37E-02 |
| R-HSA-3214847 | HATs acetylate histones | SUPT3H | SPT3 homolog, SAGA and STAGA com | 8464 | up | 1,38 | 1,01E-02 |
| R-HSA-3214847 | HATs acetylate histones | SUPT20H | SPT20 homolog, SAGA complex compo | 55578 | up | 1,29 | 4,65E-02 |
| R-HSA-3214847 | HATs acetylate histones | WDR5 | WD repeat domain 5 | 11091 | down | 1,28 | 3,81E-03 |
| R-HSA-3214847 | HATs acetylate histones | KAT2A | lysine acetyltransferase 2A | 2648 | down | 1,39 | 1,20E-02 |
| R-HSA-1482925 | Acyl chain remodelling of PG | LPCAT4 | lysophosphatidylcholine acyltransferase | 254531 | down | 1,78 | 2,52E-02 |
| R-HSA-1482925 | Acyl chain remodelling of PG | LPCAT1 | lysophosphatidylcholine acyltransferase | 79888 | down | 1,76 | 2,60E-02 |
| R-HSA-110320 | Translesion Synthesis by POLH | RFC1 | replication factor C subunit 1 | 5981 | up | 1,15 | 2,18E-02 |
| R-HSA-110320 | Translesion Synthesis by POLH | PCNA | proliferating cell nuclear antigen | 5111 | down | 2,14 | 4,63E-02 |
| R-HSA-110320 | Translesion Synthesis by POLH | RPA1 | replication protein A1 | 6117 | down | 1,41 | 4,93E-02 |
| R-HSA-1250196 | SHC1 events in ERBB2 signaling | HBEGF | heparin binding EGF like growth factor | 1839 | up | 3,67 | 5,67E-03 |
| R-HSA-1250196 | SHC1 events in ERBB2 signaling | ERBB2 | erb-b2 receptor tyrosine kinase 2 | 2064 | down | 1,47 | 1,93E-02 |
| R-HSA-1250196 | SHC1 events in ERBB2 signaling | HRAS | HRas proto-oncogene, GTPase | 3265 | down | 1,31 | 2,19E-03 |
| R-HSA-2672351 | Stimuli-sensing channels | CLCN3 | chloride voltage-gated channel 3 | 1182 | up | 1,84 | 9,81E-03 |
| R-HSA-2672351 | Stimuli-sensing channels | OSTM1 | osteopetrosis associated transmembran | 28962 | up | 1,78 | 9,61E-03 |
| R-HSA-2672351 | Stimuli-sensing channels | CLCN5 | chloride voltage-gated channel 5 | 1184 | up | 1,57 | 9,66E-03 |
| R-HSA-2672351 | Stimuli-sensing channels | CALM3 | calmodulin 3 | 808 | down | 1,43 | 2,23E-02 |
| R-HSA-2672351 | Stimuli-sensing channels | CALM2 | calmodulin 2 | 805 | down | 1,43 | 2,23E-02 |
| R-HSA-2672351 | Stimuli-sensing channels | TTYH3 | tweety family member 3 | 80727 | down | 1,75 | 6,96E-03 |
| R-HSA-2672351 | Stimuli-sensing channels | CALM1 | calmodulin 1 | 801 | down | 1,43 | 2,23E-02 |
| R-HSA-2672351 | Stimuli-sensing channels | ANO7 | anoctamin 7 | 50636 | down | 1,61 | 5,94E-03 |
| R-HSA-2672351 | Stimuli-sensing channels | BEST1 | bestrophin 1 | 7439 | up | 2,40 | 5,90E-03 |
| R-HSA-877312 | Regulation of IFNG signaling | JAK1 | Janus kinase 1 | 3716 | up | 1,53 | 2,13E-02 |
| R-HSA-877312 | Regulation of IFNG signaling | PTPN2 | protein tyrosine phosphatase, non-rece | 5771 | up | 1,40 | 1,72E-02 |
| R-HSA-1295596 | Spry regulation of FGF signaling | PPP2CB | protein phosphatase 2 catalytic subunit | 5516 | up | 1,30 | 4,79E-02 |
| R-HSA-1295596 | Spry regulation of FGF signaling | BRAF | B-Raf proto-oncogene, serine/threonine | 673 | up | 1,52 | 4,53E-02 |
| R-HSA-1660516 | Synthesis of PIPs at the early endosome membrane | PIK3C2A | phosphatidylinositol-4-phosphate 3-kina | 5286 | up | 1,30 | 1,79E-02 |
| R-HSA-1660516 | Synthesis of PIPs at the early endosome membrane | MTMR2 | myotubularin related protein 2 | 8898 | up | 1,23 | 2,61E-02 |
| R-HSA-6782315 | tRNA modification in the nucleus and cytosol | TRMT11 | tRNA methyltransferase 11 homolog | 60487 | up | 1,68 | 6,36E-03 |
| R-HSA-6782315 | tRNA modification in the nucleus and cytosol | THG1L | tRNA-histidine guanylyltransferase 1 like | 54974 | up | 1,29 | 4,59E-02 |
| R-HSA-6782315 | tRNA modification in the nucleus and cytosol | LAGE3 | L antigen family member 3 | 8270 | down | 1,23 | 3,97E-02 |
| R-HSA-6782315 | tRNA modification in the nucleus and cytosol | EPRS | glutamyl-prolyl-tRNA synthetase | 2058 | up | 1,42 | 1,95E-02 |
| R-HSA-190840 | Microtubule-dependent trafficking of connexons from | TUBA4A | tubulin alpha 4a | 7277 | down | 1,63 | 2,97E-02 |
| R-HSA-190840 | Microtubule-dependent trafficking of connexons from | TUBA1B | tubulin alpha 1b | 10376 | down | 1,78 | 2,30E-02 |
| R-HSA-2022870 | Chondroitin sulfate biosynthesis | CSPG4 | chondroitin sulfate proteoglycan 4 | 1464 | down | 2,22 | 3,99E-02 |
| R-HSA-2022870 | Chondroitin sulfate biosynthesis | CSGALNACT2 | chondroitin sulfate N-acetylgalactosamir | 55454 | up | 2,21 | 1,16E-02 |
| R-HSA-2022870 | Chondroitin sulfate biosynthesis | CHST12 | carbohydrate sulfotransferase 12 | 55501 | down | 1,51 | 1,30E-04 |
| R-HSA-5654720 | PI-3K cascade:FGFR4 | PIK3CA | phosphatidylinositol-4,5-bisphosphate 3 | 5290 | up | 1,33 | 3,17E-02 |
| R-HSA-5654720 | PI-3K cascade:FGFR4 | PIK3R1 | phosphoinositide-3-kinase regulatory su | 5295 | down | 2,04 | 8,40E-03 |
| R-HSA-5654720 | PI-3K cascade:FGFR4 | FGF2 | fibroblast growth factor 2 | 2247 | down | 5,35 | 3,09E-02 |
| R-HSA-2467813 | Separation of Sister Chromatids | TUBA1B | tubulin alpha 1b | 10376 | down | 1,78 | 2,30E-02 |
| R-HSA-2467813 | Separation of Sister Chromatids | TUBA4A | tubulin alpha 4a | 7277 | down | 1,63 | 2,97E-02 |
| R-HSA-2467813 | Separation of Sister Chromatids | STAG2 | stromal antigen 2 | 10735 | up | 1,26 | 3,69E-02 |
| R-HSA-2467813 | Separation of Sister Chromatids | MIS12 | MIS12, kinetochore complex componen | 79003 | up | 1,78 | 1,57E-02 |
| R-HSA-2467813 | Separation of Sister Chromatids | RANGAP1 | Ran GTPase activating protein 1 | 5905 | down | 1,48 | 1,41E-02 |
| R-HSA-2467813 | Separation of Sister Chromatids | ANAPC2 | anaphase promoting complex subunit 2 | 29882 | down | 1,26 | 1,99E-02 |
| R-HSA-2467813 | Separation of Sister Chromatids | CENPO | centromere protein O | 79172 | down | 1,38 | 4,86E-02 |
| R-HSA-2467813 | Separation of Sister Chromatids | ESPL1 | extra spindle pole bodies like 1, separas | 9700 | down | 1,88 | 3,17E-02 |
| R-HSA-2467813 | Separation of Sister Chromatids | B9D2 | B9 domain containing 2 | 80776 | down | 1,43 | 1,56E-02 |
| R-HSA-2467813 | Separation of Sister Chromatids | INCENP | inner centromere protein | 3619 | down | 1,78 | 2,08E-02 |
| R-HSA-2467813 | Separation of Sister Chromatids | CLIP1 | CAP-Gly domain containing linker protei | 6249 | up | 1,72 | 1,42E-03 |
| R-HSA-2467813 | Separation of Sister Chromatids | PPP2CB | protein phosphatase 2 catalytic subunit | 5516 | up | 1,30 | 4,79E-02 |
| R-HSA-2467813 | Separation of Sister Chromatids | NDEL1 | nudE neurodevelopment protein 1 like 1 | 81565 | up | 1,97 | 6,96E-03 |
| R-HSA-2467813 | Separation of Sister Chromatids | ERCC6L | ERCC excision repair 6 like, spindle ass | 54821 | down | 2,03 | 2,60E-02 |
| R-HSA-2467813 | Separation of Sister Chromatids | SPC24 | SPC24, NDC80 kinetochore complex co | 147841 | down | 2,07 | 3,62E-02 |
| R-HSA-109688 | Cleavage of Growing Transcript in the Termination Re | SARNP | SAP domain containing ribonucleoprote | 84324 | up | 1,48 | 4,35E-02 |
| R-HSA-109688 | Cleavage of Growing Transcript in the Termination Re | CLP1 | cleavage and polyadenylation factor I s | 10978 | down | 1,40 | 1,04E-02 |
| R-HSA-109688 | Cleavage of Growing Transcript in the Termination Re | SLU7 | SLU7 homolog, splicing factor | 10569 | up | 1,24 | 3,63E-02 |
| R-HSA-109688 | Cleavage of Growing Transcript in the Termination Re | SRSF2 | serine and arginine rich splicing factor 2 | 6427 | down | 1,69 | 1,79E-02 |
| R-HSA-109688 | Cleavage of Growing Transcript in the Termination Re | CSTF2 | cleavage stimulation factor subunit 2 | 1478 | down | 1,45 | 4,38E-02 |
| R-HSA-109688 | Cleavage of Growing Transcript in the Termination Re | U2AF2 | U2 small nuclear RNA auxiliary factor 2 | 11338 | down | 1,27 | 8,21E-03 |
| R-HSA-975634 | Retinoid metabolism and transport | SDC1 | syndecan 1 | 6382 | down | 2,17 | 1,61E-02 |
| R-HSA-975634 | Retinoid metabolism and transport | SDC3 | syndecan 3 | 9672 | down | 1,68 | 1,97E-02 |
| R-HSA-975634 | Retinoid metabolism and transport | LRP8 | LDL receptor related protein 8 | 7804 | down | 1,85 | 4,38E-02 |
| R-HSA-432040 | Vasopressin regulates renal water homeostasis via Aq | ADCY3 | adenylate cyclase 3 | 109 | down | 1,45 | 1,03E-02 |
| R-HSA-432040 | Vasopressin regulates renal water homeostasis via Aq | RAB11FIP2 | RAB11 family interacting protein 2 | 22841 | up | 1,50 | 4,23E-02 |
| R-HSA-432040 | Vasopressin regulates renal water homeostasis via Aq | AQP1 | aquaporin 1 (Colton blood group) | 358 | down | 2,24 | 2,60E-02 |
| R-HSA-432040 | Vasopressin regulates renal water homeostasis via Aq | PRKAR1B | protein kinase cAMP-dependent type I r | 5575 | down | 1,50 | 6,80E-04 |
| R-HSA-5578749 | Transcriptional regulation by small RNAs | POLR2B | RNA polymerase II subunit B | 5431 | down | 1,23 | 3,81E-02 |
| R-HSA-5578749 | Transcriptional regulation by small RNAs | NUP58 | nucleoporin 58 | 9818 | up | 1,61 | 1,28E-02 |
| R-HSA-5578749 | Transcriptional regulation by small RNAs | POLR2L | RNA polymerase II subunit L | 5441 | down | 1,52 | 3,58E-03 |
| R-HSA-5578749 | Transcriptional regulation by small RNAs | POLR2F | RNA polymerase II subunit F | 5435 | down | 1,41 | 3,22E-02 |
| R-HSA-5578749 | Transcriptional regulation by small RNAs | NUP62 | nucleoporin 62 | 23636 | down | 1,52 | 4,39E-03 |
| R-HSA-5578749 | Transcriptional regulation by small RNAs | POLR2E | RNA polymerase II subunit E | 5434 | down | 1,31 | 3,18E-03 |
| R-HSA-5578749 | Transcriptional regulation by small RNAs | H2AFX | H2A histone family member X | 3014 | down | 2,11 | 1,30E-03 |
| R-HSA-5654688 | SHC-mediated cascade:FGFR1 | HRAS | HRas proto-oncogene, GTPase | 3265 | down | 1,31 | 2,19E-03 |
| R-HSA-5654688 | SHC-mediated cascade:FGFR1 | FGF2 | fibroblast growth factor 2 | 2247 | down | 5,35 | 3,09E-02 |
| R-HSA-5654689 | PI-3K cascade:FGFR1 | PIK3R1 | phosphoinositide-3-kinase regulatory su | 5295 | down | 2,04 | 8,40E-03 |
| R-HSA-5654689 | PI-3K cascade:FGFR1 | FGF2 | fibroblast growth factor 2 | 2247 | down | 5,35 | 3,09E-02 |
| R-HSA-5654689 | PI-3K cascade:FGFR1 | PIK3CA | phosphatidylinositol-4,5-bisphosphate 3 | 5290 | up | 1,33 | 3,17E-02 |
| R-HSA-4086400 | PCP/CE pathway | DAAM1 | dishevelled associated activator of morp | 23002 | up | 1,56 | 1,63E-02 |
| R-HSA-4086400 | PCP/CE pathway | FZD1 | frizzled class receptor 1 | 8321 | down | 1,75 | 2,84E-02 |
| R-HSA-4086400 | PCP/CE pathway | PFN1 | profilin 1 | 5216 | down | 1,20 | 3,98E-02 |
| R-HSA-5617833 | Cilium Assembly | TUBA4A | tubulin alpha 4a | 7277 | down | 1,63 | 2,97E-02 |
| R-HSA-5617833 | Cilium Assembly | TUBA1B | tubulin alpha 1b | 10376 | down | 1,78 | 2,30E-02 |

| | | | | | | | |
|---|---|---|---|---|---|---|---|
| R-HSA-5368286 | Mitochondrial translation initiation | MTIF2 | mitochondrial translational initiation fact | 4528 | up | 1,26 | 1,83E-02 |
| R-HSA-5368286 | Mitochondrial translation initiation | MRPL4 | mitochondrial ribosomal protein L4 | 51073 | down | 1,33 | 1,11E-03 |
| R-HSA-5368286 | Mitochondrial translation initiation | MRPS10 | mitochondrial ribosomal protein S10 | 55173 | up | 1,41 | 1,62E-02 |
| R-HSA-5368286 | Mitochondrial translation initiation | MRPS12 | mitochondrial ribosomal protein S12 | 6183 | down | 1,29 | 9,88E-03 |
| R-HSA-5368286 | Mitochondrial translation initiation | MRPS34 | mitochondrial ribosomal protein S34 | 65993 | down | 1,51 | 1,07E-02 |
| R-HSA-5368286 | Mitochondrial translation initiation | MTFMT | mitochondrial methionyl-tRNA formyltran | 123263 | down | 1,40 | 4,59E-02 |
| R-HSA-5368286 | Mitochondrial translation initiation | MRPS26 | mitochondrial ribosomal protein S26 | 64949 | down | 1,49 | 2,65E-02 |
| R-HSA-445989 | TAK1 activates NFkB by phosphorylation and activati | MYD88 | myeloid differentiation primary response | 4615 | down | 1,55 | 1,85E-02 |
| R-HSA-445989 | TAK1 activates NFkB by phosphorylation and activati | TAB1 | TGF-beta activated kinase 1 (MAP3K7) | 10454 | down | 1,29 | 1,94E-02 |
| R-HSA-445989 | TAK1 activates NFkB by phosphorylation and activati | RIPK2 | receptor interacting serine/threonine kin | 8767 | up | 1,72 | 8,31E-03 |
| R-HSA-445989 | TAK1 activates NFkB by phosphorylation and activati | TAB3 | TGF-beta activated kinase 1 and MAP3 | 257397 | up | 1,40 | 4,94E-02 |
| R-HSA-2142816 | Synthesis of (16-20)-hydroxyeicosatetraenoic acids (H | CYP1A1 | cytochrome P450 family 1 subfamily A m | 1543 | up | 2,69 | 1,95E-02 |
| R-HSA-2142816 | Synthesis of (16-20)-hydroxyeicosatetraenoic acids (H | CYP2U1 | cytochrome P450 family 2 subfamily U m | 113612 | down | 1,88 | 2,19E-02 |
| R-HSA-5666185 | RHO GTPases Activate Rhotekin and Rhophilins | RHOB | ras homolog family member B | 388 | up | 1,65 | 2,44E-02 |
| R-HSA-5666185 | RHO GTPases Activate Rhotekin and Rhophilins | RTKN | rhotekin | 6242 | down | 1,39 | 3,82E-02 |
| R-HSA-74751 | Insulin receptor signalling cascade | GRB10 | growth factor receptor bound protein 10 | 2887 | up | 1,86 | 2,07E-02 |
| R-HSA-74751 | Insulin receptor signalling cascade | HRAS | HRas proto-oncogene, GTPase | 3265 | down | 1,31 | 2,19E-03 |
| R-HSA-2179392 | EGFR Transactivation by Gastrin | HRAS | HRas proto-oncogene, GTPase | 3265 | down | 1,31 | 2,19E-03 |
| R-HSA-2179392 | EGFR Transactivation by Gastrin | HBEGF | heparin binding EGF like growth factor | 1839 | up | 3,67 | 5,67E-03 |
| R-HSA-2993913 | Clearance of Nuclear Envelope Membranes from Chro | VRK2 | vaccinia related kinase 2 | 7444 | up | 1,28 | 9,54E-03 |
| R-HSA-2993913 | Clearance of Nuclear Envelope Membranes from Chro | LMNB1 | lamin B1 | 4001 | down | 2,12 | 3,85E-02 |
| R-HSA-426486 | Small interfering RNA (siRNA) biogenesis | TARBP2 | TARBP2, RISC loading complex RNA bi | 6895 | down | 1,36 | 3,80E-03 |
| R-HSA-426486 | Small interfering RNA (siRNA) biogenesis | AGO3 | argonaute 3, RISC catalytic component | 192669 | up | 1,36 | 1,80E-02 |
| R-HSA-389356 | CD28 co-stimulation | LYN | LYN proto-oncogene, Src family tyrosine | 4067 | down | 1,36 | 2,34E-02 |
| R-HSA-389356 | CD28 co-stimulation | YES1 | YES proto-oncogene 1, Src family tyros | 7525 | up | 1,30 | 2,08E-02 |
| R-HSA-170968 | Frs2-mediated activation | BRAF | B-Raf proto-oncogene, serine/threonine | 673 | up | 1,52 | 4,53E-02 |
| R-HSA-170968 | Frs2-mediated activation | PLCG1 | phospholipase C gamma 1 | 5335 | down | 1,51 | 2,85E-02 |
| R-HSA-428540 | Activation of RAC1 | NCK1 | NCK adaptor protein 1 | 4690 | up | 1,48 | 1,42E-02 |
| R-HSA-428540 | Activation of RAC1 | SOS2 | SOS Ras/Rho guanine nucleotide exch | 6655 | up | 1,82 | 1,41E-02 |
| R-HSA-72695 | Formation of the ternary complex, and subsequently, | RPS4Y1 | ribosomal protein S4, Y-linked 1 | 6192 | down | 5,59 | 7,82E-03 |
| R-HSA-72695 | Formation of the ternary complex, and subsequently, | EIF2S3 | eukaryotic translation initiation factor 2 s | 1968 | up | 1,19 | 2,93E-02 |
| R-HSA-72695 | Formation of the ternary complex, and subsequently, | EIF3M | eukaryotic translation initiation factor 3 s | 10480 | up | 1,25 | 2,51E-02 |
| R-HSA-72695 | Formation of the ternary complex, and subsequently, | EIF3H | eukaryotic translation initiation factor 3 s | 8667 | up | 1,22 | 3,84E-02 |
| R-HSA-72695 | Formation of the ternary complex, and subsequently, | EIF3B | eukaryotic translation initiation factor 3 s | 8662 | down | 1,27 | 1,76E-02 |
| R-HSA-72695 | Formation of the ternary complex, and subsequently, | EIF2S2 | eukaryotic translation initiation factor 2 s | 8894 | up | 1,40 | 2,40E-02 |
| R-HSA-4641257 | Degradation of AXIN | SMURF2 | SMAD specific E3 ubiquitin protein ligas | 64750 | up | 1,49 | 4,16E-02 |
| R-HSA-4641257 | Degradation of AXIN | RNF146 | ring finger protein 146 | 81847 | up | 1,28 | 3,57E-02 |
| R-HSA-4641257 | Degradation of AXIN | AXIN2 | axin 2 | 8313 | down | 2,79 | 3,67E-03 |
| R-HSA-4641257 | Degradation of AXIN | TNKS | tankyrase | 8658 | up | 1,42 | 8,31E-03 |
| R-HSA-4641257 | Degradation of AXIN | TNKS2 | tankyrase 2 | 80351 | up | 1,68 | 3,02E-03 |
| R-HSA-2028269 | Signaling by Hippo | STK3 | serine/threonine kinase 3 | 6788 | up | 1,61 | 2,08E-03 |
| R-HSA-2028269 | Signaling by Hippo | TJP1 | tight junction protein 1 | 7082 | up | 1,46 | 8,91E-03 |
| R-HSA-2028269 | Signaling by Hippo | WWTR1 | WW domain containing transcription reg | 25937 | down | 1,99 | 1,00E-02 |
| R-HSA-5654712 | FRS-mediated FGFR4 signaling | FGF2 | fibroblast growth factor 2 | 2247 | down | 5,35 | 3,09E-02 |
| R-HSA-5654712 | FRS-mediated FGFR4 signaling | HRAS | HRas proto-oncogene, GTPase | 3265 | down | 1,31 | 2,19E-03 |
| R-HSA-5654732 | Negative regulation of FGFR3 signaling | FGF2 | fibroblast growth factor 2 | 2247 | down | 5,35 | 3,09E-02 |
| R-HSA-5654732 | Negative regulation of FGFR3 signaling | FGFR3 | fibroblast growth factor receptor 3 | 2261 | down | 1,70 | 4,60E-02 |
| R-HSA-2024096 | HS-GAG degradation | SDC1 | syndecan 1 | 6382 | down | 2,17 | 1,61E-02 |
| R-HSA-2024096 | HS-GAG degradation | SDC3 | syndecan 3 | 9672 | down | 1,68 | 1,97E-02 |
| R-HSA-2871809 | FCERI mediated Ca+2 mobilization | CALM1 | calmodulin 1 | 801 | down | 1,43 | 2,23E-02 |
| R-HSA-2871809 | FCERI mediated Ca+2 mobilization | PLCG1 | phospholipase C gamma 1 | 5335 | down | 1,51 | 2,85E-02 |
| R-HSA-2871809 | FCERI mediated Ca+2 mobilization | LYN | LYN proto-oncogene, Src family tyrosine | 4067 | down | 1,36 | 2,34E-02 |
| R-HSA-2871809 | FCERI mediated Ca+2 mobilization | CALM2 | calmodulin 2 | 805 | down | 1,43 | 2,23E-02 |
| R-HSA-2871809 | FCERI mediated Ca+2 mobilization | CALM3 | calmodulin 3 | 808 | down | 1,43 | 2,23E-02 |
| R-HSA-5362768 | Hh mutants that don't undergo autocatalytic processi | DERL2 | derlin 2 | 51009 | up | 1,68 | 4,88E-02 |
| R-HSA-5362768 | Hh mutants that don't undergo autocatalytic processi | SEL1L | SEL1L ERAD E3 ligase adaptor subuni | 6400 | up | 2,76 | 4,03E-03 |
| R-HSA-5362768 | Hh mutants that don't undergo autocatalytic processi | SYVN1 | synoviolin 1 | 84447 | up | 2,33 | 1,93E-02 |
| R-HSA-5362768 | Hh mutants that don't undergo autocatalytic processi | ERLEC1 | endoplasmic reticulum lectin 1 | 27248 | up | 1,96 | 7,71E-03 |
| R-HSA-352230 | Amino acid transport across the plasma membrane | SLC38A2 | solute carrier family 38 member 2 | 54407 | up | 1,62 | 4,57E-02 |
| R-HSA-352230 | Amino acid transport across the plasma membrane | SLC38A5 | solute carrier family 38 member 5 | 92745 | down | 1,46 | 4,02E-02 |
| R-HSA-352230 | Amino acid transport across the plasma membrane | SLC7A6 | solute carrier family 7 member 6 | 9057 | down | 1,49 | 1,37E-02 |
| R-HSA-352230 | Amino acid transport across the plasma membrane | SLC3A2 | solute carrier family 3 member 2 | 6520 | up | 2,41 | 7,95E-03 |
| R-HSA-445095 | Interaction between L1 and Ankyrins | SCN5A | sodium voltage-gated channel alpha su | 6331 | down | 15,38 | 3,09E-02 |
| R-HSA-445095 | Interaction between L1 and Ankyrins | SPTBN1 | spectrin beta, non-erythrocytic 1 | 6711 | up | 1,42 | 1,66E-02 |
| R-HSA-445095 | Interaction between L1 and Ankyrins | ACTG1 | actin gamma 1 | 71 | up | 1,58 | 1,66E-02 |
| R-HSA-445095 | Interaction between L1 and Ankyrins | L1CAM | L1 cell adhesion molecule | 3897 | down | 2,86 | 3,09E-02 |
| R-HSA-4420097 | VEGFA-VEGFR2 Pathway | PIK3CA | phosphatidylinositol-4,5-bisphosphate 3 | 5290 | up | 1,33 | 3,17E-02 |
| R-HSA-4420097 | VEGFA-VEGFR2 Pathway | ACTG1 | actin gamma 1 | 71 | up | 1,58 | 1,66E-02 |
| R-HSA-4420097 | VEGFA-VEGFR2 Pathway | PIK3R1 | phosphoinositide-3-kinase regulatory su | 5295 | down | 2,04 | 8,40E-03 |
| R-HSA-4420097 | VEGFA-VEGFR2 Pathway | NCK1 | NCK adaptor protein 1 | 4690 | up | 1,48 | 1,42E-02 |
| R-HSA-4420097 | VEGFA-VEGFR2 Pathway | VEGFA | vascular endothelial growth factor A | 7422 | up | 2,38 | 1,68E-02 |
| R-HSA-4420097 | VEGFA-VEGFR2 Pathway | CYBA | cytochrome b-245 alpha chain | 1535 | down | 1,34 | 1,09E-02 |
| R-HSA-4420097 | VEGFA-VEGFR2 Pathway | MAPK12 | mitogen-activated protein kinase 12 | 6300 | down | 1,37 | 3,32E-03 |
| R-HSA-438066 | Unblocking of NMDA receptors, glutamate binding an | CAMK2D | calcium/calmodulin dependent protein k | 817 | up | 1,29 | 1,49E-02 |
| R-HSA-438066 | Unblocking of NMDA receptors, glutamate binding an | AKAP9 | A-kinase anchoring protein 9 | 10142 | up | 1,47 | 1,80E-02 |
| R-HSA-168276 | NS1 Mediated Effects on Host Pathways | KPNA5 | karyopherin subunit alpha 5 | 3841 | up | 1,61 | 1,30E-02 |
| R-HSA-168276 | NS1 Mediated Effects on Host Pathways | NUP58 | nucleoporin 58 | 9818 | up | 1,61 | 1,28E-02 |
| R-HSA-168276 | NS1 Mediated Effects on Host Pathways | KPNA4 | karyopherin subunit alpha 4 | 3840 | up | 1,61 | 4,03E-03 |
| R-HSA-168276 | NS1 Mediated Effects on Host Pathways | NUP62 | nucleoporin 62 | 23636 | down | 1,52 | 4,39E-03 |
| R-HSA-5620916 | VxPx cargo-targeting to cilium | RAB11FIP3 | RAB11 family interacting protein 3 | 9727 | down | 1,28 | 2,24E-02 |
| R-HSA-5620916 | VxPx cargo-targeting to cilium | ASAP1 | ArfGAP with SH3 domain, ankyrin repea | 50807 | up | 1,27 | 4,95E-02 |
| R-HSA-5620916 | VxPx cargo-targeting to cilium | ARF4 | ADP ribosylation factor 4 | 378 | up | 1,69 | 2,56E-02 |
| R-HSA-450302 | activated TAK1 mediates p38 MAPK activation | TAB1 | TGF-beta activated kinase 1 (MAP3K7) | 10454 | down | 1,29 | 1,94E-02 |
| R-HSA-450302 | activated TAK1 mediates p38 MAPK activation | TAB3 | TGF-beta activated kinase 1 and MAP3 | 257397 | up | 1,40 | 4,94E-02 |
| R-HSA-450302 | activated TAK1 mediates p38 MAPK activation | RIPK2 | receptor interacting serine/threonine kin | 8767 | up | 1,72 | 8,31E-03 |
| R-HSA-379726 | Mitochondrial tRNA aminoacylation | GARS | glycyl-tRNA synthetase | 2617 | up | 1,65 | 1,79E-02 |
| R-HSA-379726 | Mitochondrial tRNA aminoacylation | 37316 | methionyl-tRNA synthetase 2, mitochon | 92935 | down | 1,29 | 2,13E-02 |
| R-HSA-379726 | Mitochondrial tRNA aminoacylation | WARS2 | tryptophanyl tRNA synthetase 2, mitoch | 10352 | up | 1,54 | 9,54E-03 |
| R-HSA-5635838 | Activation of SMO | CSNK1A1 | casein kinase 1 alpha 1 | 1452 | up | 1,35 | 2,84E-02 |
| R-HSA-5635838 | Activation of SMO | ARRB1 | arrestin beta 1 | 408 | down | 1,79 | 2,42E-02 |
| R-HSA-937042 | IRAK2 mediated activation of TAK1 complex | TAB1 | TGF-beta activated kinase 1 (MAP3K7) | 10454 | down | 1,29 | 1,94E-02 |
| R-HSA-937042 | IRAK2 mediated activation of TAK1 complex | TAB3 | TGF-beta activated kinase 1 and MAP3 | 257397 | up | 1,40 | 4,94E-02 |
| R-HSA-975163 | IRAK2 mediated activation of TAK1 complex upon TL | TAB3 | TGF-beta activated kinase 1 and MAP3 | 257397 | up | 1,40 | 4,94E-02 |
| R-HSA-975163 | IRAK2 mediated activation of TAK1 complex upon TL | TAB1 | TGF-beta activated kinase 1 (MAP3K7) | 10454 | down | 1,29 | 1,94E-02 |
| R-HSA-73614 | Pyrimidine salvage | UCK2 | uridine-cytidine kinase 2 | 7371 | down | 1,45 | 2,72E-02 |
| R-HSA-73614 | Pyrimidine salvage | UPP1 | uridine phosphorylase 1 | 7378 | up | 4,41 | 1,62E-03 |
| R-HSA-191859 | snRNP Assembly | GEMIN7 | gem nuclear organelle associated prote | 79760 | down | 1,40 | 2,72E-03 |
| R-HSA-191859 | snRNP Assembly | GEMIN4 | gem nuclear organelle associated prote | 50628 | down | 1,89 | 5,67E-03 |
| R-HSA-191859 | snRNP Assembly | TGS1 | trimethylguanosine synthase 1 | 96764 | up | 1,63 | 9,37E-03 |
| R-HSA-5654695 | PI-3K cascade:FGFR2 | PIK3CA | phosphatidylinositol-4,5-bisphosphate 3 | 5290 | up | 1,33 | 3,17E-02 |
| R-HSA-5654695 | PI-3K cascade:FGFR2 | PIK3R1 | phosphoinositide-3-kinase regulatory su | 5295 | down | 2,04 | 8,40E-03 |
| R-HSA-5654695 | PI-3K cascade:FGFR2 | FGF2 | fibroblast growth factor 2 | 2247 | down | 5,35 | 3,09E-02 |
| R-HSA-5654693 | FRS-mediated FGFR1 signaling | FGF2 | fibroblast growth factor 2 | 2247 | down | 5,35 | 3,09E-02 |
| R-HSA-5654693 | FRS-mediated FGFR1 signaling | HRAS | HRas proto-oncogene, GTPase | 3265 | down | 1,31 | 2,19E-03 |
| R-HSA-5654719 | SHC-mediated cascade:FGFR4 | HRAS | HRas proto-oncogene, GTPase | 3265 | down | 1,31 | 2,19E-03 |
| R-HSA-5654719 | SHC-mediated cascade:FGFR4 | FGF2 | fibroblast growth factor 2 | 2247 | down | 5,35 | 3,09E-02 |
| R-HSA-166208 | mTORC1-mediated signalling | RPTOR | regulatory associated protein of MTOR c | 57521 | down | 1,20 | 1,16E-02 |
| R-HSA-166208 | mTORC1-mediated signalling | RRAGC | Ras related GTP binding C | 64121 | up | 1,78 | 6,10E-03 |
| R-HSA-166208 | mTORC1-mediated signalling | AKT1S1 | AKT1 substrate 1 | 84335 | down | 1,21 | 4,33E-02 |
| R-HSA-354192 | Integrin alphaIIb beta3 signaling | CSK | C-terminal Src kinase | 1445 | down | 1,24 | 5,67E-03 |
| R-HSA-354192 | Integrin alphaIIb beta3 signaling | AKT1 | AKT serine/threonine kinase 1 | 207 | down | 1,37 | 1,42E-03 |
| R-HSA-4570464 | SUMOylation of RNA binding proteins | CBX4 | chromobox 4 | 8535 | up | 2,70 | 4,00E-03 |
| R-HSA-4570464 | SUMOylation of RNA binding proteins | NUP58 | nucleoporin 58 | 9818 | up | 1,61 | 1,28E-02 |
| R-HSA-4570464 | SUMOylation of RNA binding proteins | CBX8 | chromobox 8 | 57332 | down | 1,30 | 2,19E-02 |
| R-HSA-4570464 | SUMOylation of RNA binding proteins | CBX2 | chromobox 2 | 84733 | down | 2,70 | 8,09E-03 |
| R-HSA-4570464 | SUMOylation of RNA binding proteins | NUP62 | nucleoporin 62 | 23636 | down | 1,52 | 4,39E-03 |
| R-HSA-193368 | Synthesis of bile acids and bile salts via 7alpha-hydro | RXRA | retinoid X receptor alpha | 6256 | down | 1,34 | 3,67E-03 |
| R-HSA-193368 | Synthesis of bile acids and bile salts via 7alpha-hydro | HSD3B7 | hydroxy-delta-5-steroid dehydrogenase, | 80270 | down | 1,64 | 3,72E-02 |
| R-HSA-933542 | TRAF6 mediated NF-kB activation | MYD88 | myeloid differentiation primary response | 4615 | down | 1,55 | 1,85E-02 |
| R-HSA-933542 | TRAF6 mediated NF-kB activation | TRAF2 | TNF receptor associated factor 2 | 7186 | down | 1,47 | 2,65E-02 |
| R-HSA-176408 | Regulation of APC/C activators between G1/S and ea | ANAPC2 | anaphase promoting complex subunit 2 | 29882 | down | 1,26 | 1,99E-02 |
| R-HSA-176408 | Regulation of APC/C activators between G1/S and ea | FBXO5 | F-box protein 5 | 26271 | down | 1,57 | 4,56E-02 |
| R-HSA-389977 | Post-chaperonin tubulin folding pathway | TUBA1B | tubulin alpha 1b | 10376 | down | 1,78 | 2,30E-02 |
| R-HSA-389977 | Post-chaperonin tubulin folding pathway | TUBA4A | tubulin alpha 4a | 7277 | down | 1,63 | 2,97E-02 |
| R-HSA-392451 | G beta:gamma signalling through PI3Kgamma | AKT2 | AKT serine/threonine kinase 2 | 208 | down | 1,28 | 6,33E-03 |
| R-HSA-392451 | G beta:gamma signalling through PI3Kgamma | JAK1 | Janus kinase 1 | 3716 | up | 1,53 | 2,13E-02 |
| R-HSA-392451 | G beta:gamma signalling through PI3Kgamma | PIK3R1 | phosphoinositide-3-kinase regulatory su | 5295 | down | 2,04 | 8,40E-03 |
| R-HSA-392451 | G beta:gamma signalling through PI3Kgamma | PIK3CA | phosphatidylinositol-4,5-bisphosphate 3 | 5290 | up | 1,33 | 3,17E-02 |
| R-HSA-392451 | G beta:gamma signalling through PI3Kgamma | AKT1 | AKT serine/threonine kinase 1 | 207 | down | 1,37 | 1,42E-03 |
| R-HSA-373080 | Class B/2 (Secretin family receptors) | CD55 | CD55 molecule (Cromer blood group) | 1604 | up | 3,43 | 2,24E-03 |
| R-HSA-373080 | Class B/2 (Secretin family receptors) | FZD4 | frizzled class receptor 4 | 8322 | down | 2,17 | 3,82E-02 |
| R-HSA-373080 | Class B/2 (Secretin family receptors) | WNT10B | Wnt family member 10B | 7480 | down | 1,76 | 4,78E-02 |
| R-HSA-373080 | Class B/2 (Secretin family receptors) | FZD1 | frizzled class receptor 1 | 8321 | down | 1,75 | 2,84E-02 |
| R-HSA-111465 | Apoptotic cleavage of cellular proteins | BIRC2 | baculoviral IAP repeat containing 2 | 329 | up | 1,38 | 1,86E-02 |
| R-HSA-111465 | Apoptotic cleavage of cellular proteins | FNTA | farnesyltransferase, CAAX box, alpha | 2339 | up | 1,33 | 3,73E-03 |
| R-HSA-5358346 | Hedgehog ligand biogenesis | DERL2 | derlin 2 | 51009 | up | 1,68 | 4,88E-02 |
| R-HSA-5358346 | Hedgehog ligand biogenesis | SYVN1 | synoviolin 1 | 84447 | up | 2,33 | 1,93E-02 |
| R-HSA-5358346 | Hedgehog ligand biogenesis | ERLEC1 | endoplasmic reticulum lectin 1 | 27248 | up | 1,96 | 7,71E-03 |
| R-HSA-5358346 | Hedgehog ligand biogenesis | SEL1L | SEL1L ERAD E3 ligase adaptor subuni | 6400 | up | 2,76 | 4,03E-03 |
| R-HSA-68949 | Orc1 removal from chromatin | MCM3 | minichromosome maintenance complex | 4172 | down | 2,07 | 4,13E-02 |
| R-HSA-68949 | Orc1 removal from chromatin | MCM5 | minichromosome maintenance complex | 4174 | down | 1,83 | 2,22E-02 |
| R-HSA-68949 | Orc1 removal from chromatin | MCM7 | minichromosome maintenance complex | 4176 | down | 1,67 | 3,35E-02 |
| R-HSA-68949 | Orc1 removal from chromatin | CDKN1A | cyclin dependent kinase inhibitor 1A | 1026 | up | 3,62 | 5,43E-03 |
| R-HSA-68949 | Orc1 removal from chromatin | MCM2 | minichromosome maintenance complex | 4171 | down | 2,01 | 1,42E-02 |
| R-HSA-5654700 | FRS-mediated FGFR2 signaling | FGF2 | fibroblast growth factor 2 | 2247 | down | 5,35 | 3,09E-02 |
| R-HSA-5654700 | FRS-mediated FGFR2 signaling | HRAS | HRas proto-oncogene, GTPase | 3265 | down | 1,31 | 2,19E-03 |
| R-HSA-75876 | Synthesis of very long-chain fatty acyl-CoAs | PPT2 | palmitoyl-protein thioesterase 2 | 9374 | down | 1,51 | 1,22E-02 |
| R-HSA-75876 | Synthesis of very long-chain fatty acyl-CoAs | ACSL1 | acyl-CoA synthetase long chain family m | 2180 | down | 1,54 | 3,01E-02 |
| R-HSA-212436 | Generic Transcription Pathway | ZNF274 | zinc finger protein 274 | 10782 | up | 11,88 | 2,26E-03 |
| R-HSA-212436 | Generic Transcription Pathway | ZNF75D | zinc finger protein 75D | 7626 | up | 1,95 | 7,64E-03 |
| R-HSA-212436 | Generic Transcription Pathway | ZNF324B | zinc finger protein 324B | 388569 | down | 1,67 | 2,93E-02 |
| R-HSA-212436 | Generic Transcription Pathway | ZFP69B | ZFP69 zinc finger protein B | 65243 | up | 1,57 | 2,17E-02 |
| R-HSA-212436 | Generic Transcription Pathway | MED24 | mediator complex subunit 24 | 9862 | down | 1,33 | 2,93E-02 |
| R-HSA-212436 | Generic Transcription Pathway | ZNF222 | zinc finger protein 222 | 7673 | up | 3,33 | 1,41E-03 |
| R-HSA-212436 | Generic Transcription Pathway | ZNF641 | zinc finger protein 641 | 121274 | up | 1,63 | 1,30E-02 |
| R-HSA-212436 | Generic Transcription Pathway | ZNF333 | zinc finger protein 333 | 84449 | up | 1,80 | 2,50E-03 |
| R-HSA-212436 | Generic Transcription Pathway | ZNF540 | zinc finger protein 540 | 163255 | down | 2,09 | 2,24E-03 |
| R-HSA-212436 | Generic Transcription Pathway | ZNF442 | zinc finger protein 442 | 79973 | up | 1,56 | 2,52E-02 |
| R-HSA-212436 | Generic Transcription Pathway | TRIM28 | tripartite motif containing 28 | 10155 | down | 1,24 | 4,18E-02 |
| R-HSA-212436 | Generic Transcription Pathway | ZNF544 | zinc finger protein 544 | 27300 | down | 1,27 | 4,85E-02 |
| R-HSA-212436 | Generic Transcription Pathway | ZNF793 | zinc finger protein 793 | 390927 | down | 2,21 | 4,11E-02 |
| R-HSA-212436 | Generic Transcription Pathway | ZNF689 | zinc finger protein 689 | 115509 | down | 1,50 | 9,77E-03 |
| R-HSA-212436 | Generic Transcription Pathway | ZNF699 | zinc finger protein 699 | 374879 | up | 1,88 | 2,17E-02 |
| R-HSA-212436 | Generic Transcription Pathway | ZNF792 | zinc finger protein 792 | 126375 | down | 2,40 | 1,00E-02 |
| R-HSA-212436 | Generic Transcription Pathway | ZNF555 | zinc finger protein 555 | 148254 | up | 1,77 | 5,21E-03 |
| R-HSA-212436 | Generic Transcription Pathway | ZNF791 | zinc finger protein 791 | 163049 | up | 1,34 | 2,88E-02 |
| R-HSA-212436 | Generic Transcription Pathway | ZNF668 | zinc finger protein 668 | 79759 | down | 1,27 | 3,49E-02 |
| R-HSA-212436 | Generic Transcription Pathway | ZNF180 | zinc finger protein 180 | 7733 | down | 1,63 | 1,84E-02 |
| R-HSA-212436 | Generic Transcription Pathway | ZNF703 | zinc finger protein 703 | 80139 | down | 1,48 | 4,66E-02 |
| R-HSA-212436 | Generic Transcription Pathway | ZKSCAN4 | zinc finger with KRAB and SCAN domai | 387032 | down | 1,46 | 3,85E-02 |
| R-HSA-212436 | Generic Transcription Pathway | ZNF551 | zinc finger protein 551 | 90233 | down | 1,79 | 1,18E-02 |
| R-HSA-212436 | Generic Transcription Pathway | ZNF697 | zinc finger protein 697 | 90874 | up | 1,82 | 6,32E-03 |
| R-HSA-212436 | Generic Transcription Pathway | ZNF77 | zinc finger protein 77 | 58492 | up | 1,54 | 2,97E-02 |
| R-HSA-212436 | Generic Transcription Pathway | ZNF250 | zinc finger protein 250 | 58500 | down | 1,59 | 2,28E-02 |
| R-HSA-212436 | Generic Transcription Pathway | ZNF530 | zinc finger protein 530 | 348327 | down | 1,72 | 1,40E-02 |
| R-HSA-212436 | Generic Transcription Pathway | ZNF611 | zinc finger protein 611 | 81856 | up | 1,25 | 2,68E-02 |
| R-HSA-212436 | Generic Transcription Pathway | ZNF433 | zinc finger protein 433 | 163059 | up | 1,69 | 5,67E-03 |
| R-HSA-212436 | Generic Transcription Pathway | ZNF552 | zinc finger protein 552 | 79818 | down | 1,71 | 1,41E-02 |
| R-HSA-212436 | Generic Transcription Pathway | ZNF441 | zinc finger protein 441 | 126068 | up | 1,58 | 3,64E-02 |
| R-HSA-212436 | Generic Transcription Pathway | ZNF70 | zinc finger protein 70 | 7621 | up | 1,68 | 4,63E-02 |
| R-HSA-212436 | Generic Transcription Pathway | ZNF562 | zinc finger protein 562 | 54811 | up | 1,33 | 5,21E-03 |
| R-HSA-212436 | Generic Transcription Pathway | ZNF184 | zinc finger protein 184 | 7738 | up | 1,28 | 2,30E-02 |
| R-HSA-212436 | Generic Transcription Pathway | MED25 | mediator complex subunit 25 | 81857 | down | 1,23 | 4,79E-02 |
| R-HSA-212436 | Generic Transcription Pathway | ZNF25 | zinc finger protein 25 | 219749 | up | 1,82 | 5,69E-04 |
| R-HSA-382556 | ABC-family proteins mediated transport | DERL2 | derlin 2 | 51009 | up | 1,68 | 4,88E-02 |
| R-HSA-382556 | ABC-family proteins mediated transport | EIF2S2 | eukaryotic translation initiation factor 2 s | 8894 | up | 1,40 | 2,40E-02 |
| R-HSA-382556 | ABC-family proteins mediated transport | SEL1L | SEL1L ERAD E3 ligase adaptor subuni | 6400 | up | 2,76 | 4,03E-03 |
| R-HSA-382556 | ABC-family proteins mediated transport | ERLEC1 | endoplasmic reticulum lectin 1 | 27248 | up | 1,96 | 7,71E-03 |
| R-HSA-382556 | ABC-family proteins mediated transport | EIF2S3 | eukaryotic translation initiation factor 2 s | 1968 | up | 1,19 | 2,93E-02 |
| R-HSA-6788467 | IL-6-type cytokine receptor ligand interactions | JAK1 | Janus kinase 1 | 3716 | up | 1,53 | 2,13E-02 |
| R-HSA-6788467 | IL-6-type cytokine receptor ligand interactions | IL6ST | interleukin 6 signal transducer | 3572 | up | 1,52 | 1,96E-02 |
| R-HSA-5675482 | Regulation of necroptotic cell death | TRAF2 | TNF receptor associated factor 2 | 7186 | down | 1,47 | 2,65E-02 |
| R-HSA-5675482 | Regulation of necroptotic cell death | BIRC2 | baculoviral IAP repeat containing 2 | 329 | up | 1,38 | 1,86E-02 |
| R-HSA-376172 | DSCAM interactions | MAPK12 | mitogen-activated protein kinase 12 | 6300 | down | 1,37 | 3,32E-03 |
| R-HSA-376172 | DSCAM interactions | MAPK8 | mitogen-activated protein kinase 8 | 5599 | up | 1,38 | 3,05E-02 |
| R-HSA-5654699 | SHC-mediated cascade:FGFR2 | FGF2 | fibroblast growth factor 2 | 2247 | down | 5,35 | 3,09E-02 |
| R-HSA-5654699 | SHC-mediated cascade:FGFR2 | HRAS | HRas proto-oncogene, GTPase | 3265 | down | 1,31 | 2,19E-03 |
| R-HSA-212676 | Dopamine Neurotransmitter Release Cycle | RAB3A | RAB3A, member RAS oncogene family | 5864 | down | 1,80 | 4,63E-02 |
| R-HSA-212676 | Dopamine Neurotransmitter Release Cycle | VAMP2 | vesicle associated membrane protein 2 | 6844 | up | 1,50 | 4,82E-02 |
| R-HSA-212676 | Dopamine Neurotransmitter Release Cycle | PPFIA1 | PTPRF interacting protein alpha 1 | 8500 | up | 1,30 | 3,68E-02 |
| R-HSA-429947 | Deadenylation of mRNA | CNOT2 | CCR4-NOT transcription complex subun | 4848 | up | 1,19 | 4,40E-02 |
| R-HSA-429947 | Deadenylation of mRNA | EIF4A2 | eukaryotic translation initiation factor 4A | 1974 | up | 1,68 | 4,70E-03 |
| R-HSA-429947 | Deadenylation of mRNA | CNOT9 | CCR4-NOT transcription complex subun | 9125 | down | 1,30 | 4,90E-02 |
| R-HSA-877300 | Interferon gamma signaling | TRIM3 | tripartite motif containing 3 | 10612 | down | 1,49 | 2,76E-02 |
| R-HSA-877300 | Interferon gamma signaling | IRF3 | interferon regulatory factor 3 | 3661 | down | 1,33 | 2,58E-03 |
| R-HSA-877300 | Interferon gamma signaling | CAMK2D | calcium/calmodulin dependent protein k | 817 | up | 1,29 | 1,49E-02 |
| R-HSA-877300 | Interferon gamma signaling | MT2A | metallothionein 2A | 4502 | down | 1,67 | 7,14E-03 |
| R-HSA-877300 | Interferon gamma signaling | TRIM38 | tripartite motif containing 38 | 10475 | up | 1,27 | 3,49E-02 |
| R-HSA-877300 | Interferon gamma signaling | SP100 | SP100 nuclear antigen | 6672 | up | 1,31 | 2,71E-02 |
| R-HSA-877300 | Interferon gamma signaling | JAK1 | Janus kinase 1 | 3716 | up | 1,53 | 2,13E-02 |
| R-HSA-877300 | Interferon gamma signaling | TRIM5 | tripartite motif containing 5 | 85363 | up | 1,56 | 4,94E-02 |
| R-HSA-389960 | Formation of tubulin folding intermediates by CCT/TriC | TUBA1B | tubulin alpha 1b | 10376 | down | 1,78 | 2,30E-02 |
| R-HSA-389960 | Formation of tubulin folding intermediates by CCT/TriC | TUBA4A | tubulin alpha 4a | 7277 | down | 1,63 | 2,97E-02 |
| R-HSA-1482788 | Acyl chain remodelling of PC | PNPLA8 | patatin like phospholipase domain conta | 50640 | up | 2,48 | 3,36E-03 |
| R-HSA-1482788 | Acyl chain remodelling of PC | LPCAT1 | lysophosphatidylcholine acyltransferase | 79888 | down | 1,76 | 2,60E-02 |
| R-HSA-1482788 | Acyl chain remodelling of PC | LPCAT4 | lysophosphatidylcholine acyltransferase | 254531 | down | 1,78 | 2,52E-02 |
| R-HSA-389661 | Glyoxylate metabolism and glycine degradation | LIPT1 | lipoyltransferase 1 | 51601 | up | 1,63 | 1,73E-02 |

| | | | | | | | |
|---|---|---|---|---|---|---|---|
| R-HSA-389661 | Glyoxylate metabolism and glycine degradation | DLD | dihydrolipoamide dehydrogenase | 1738 | up | 1,16 | 4,53E-02 |
| R-HSA-432720 | Lysosome Vesicle Biogenesis | ARRB1 | arrestin beta 1 | 408 | down | 1,79 | 2,42E-02 |
| R-HSA-432720 | Lysosome Vesicle Biogenesis | AP1M1 | adaptor related protein complex 1 mu 1 | 8907 | down | 1,24 | 1,30E-02 |
| R-HSA-432720 | Lysosome Vesicle Biogenesis | DNAJC6 | DnaJ heat shock protein family (Hsp40) | 9829 | up | 1,87 | 1,56E-02 |
| R-HSA-432720 | Lysosome Vesicle Biogenesis | VAMP2 | vesicle associated membrane protein 2 | 6844 | up | 1,50 | 4,82E-02 |
| R-HSA-2173789 | TGF-beta receptor signaling activates SMADs | TGFBR2 | transforming growth factor beta recepto | 7048 | down | 1,34 | 2,74E-02 |
| R-HSA-2173789 | TGF-beta receptor signaling activates SMADs | SMAD2 | SMAD family member 2 | 4087 | up | 1,29 | 2,19E-02 |
| R-HSA-202670 | ERKs are inactivated | PPP2CB | protein phosphatase 2 catalytic subunit | 5516 | up | 1,30 | 4,79E-02 |
| R-HSA-202670 | ERKs are inactivated | DUSP7 | dual specificity phosphatase 7 | 1849 | down | 1,42 | 1,78E-02 |
| R-HSA-388841 | Costimulation by the CD28 family | PIK3R1 | phosphoinositide-3-kinase regulatory su | 5295 | down | 2,04 | 8,40E-03 |
| R-HSA-388841 | Costimulation by the CD28 family | PIK3CA | phosphatidylinositol-4,5-bisphosphate 3 | 5290 | up | 1,33 | 3,17E-02 |
| R-HSA-2197563 | NOTCH2 intracellular domain regulates transcription | MAMLD1 | mastermind like domain containing 1 | 10046 | down | 3,36 | 4,56E-02 |
| R-HSA-2197563 | NOTCH2 intracellular domain regulates transcription | CREB1 | cAMP responsive element binding prote | 1385 | up | 1,51 | 7,18E-03 |
| R-HSA-2160916 | Hyaluronan uptake and degradation | HYAL2 | hyaluronoglucosaminidase 2 | 8692 | down | 1,71 | 7,16E-03 |
| R-HSA-2160916 | Hyaluronan uptake and degradation | SLC9A1 | solute carrier family 9 member A1 | 6548 | up | 2,29 | 9,81E-03 |
| R-HSA-1474228 | Degradation of the extracellular matrix | HTRA1 | HtrA serine peptidase 1 | 5654 | down | 3,48 | 2,23E-02 |
| R-HSA-1474228 | Degradation of the extracellular matrix | CAPN7 | calpain 7 | 23473 | up | 1,25 | 4,72E-02 |
| R-HSA-1474228 | Degradation of the extracellular matrix | LAMB3 | laminin subunit beta 3 | 3914 | up | 2,74 | 1,16E-02 |
| R-HSA-1474228 | Degradation of the extracellular matrix | ADAMTS9 | ADAM metallopeptidase with thrombosp | 56999 | down | 1,98 | 4,91E-02 |
| R-HSA-1474228 | Degradation of the extracellular matrix | CAST | calpastatin | 831 | up | 1,22 | 4,60E-02 |
| R-HSA-1474228 | Degradation of the extracellular matrix | CAPN5 | calpain 5 | 726 | up | 1,82 | 1,32E-02 |
| R-HSA-1474228 | Degradation of the extracellular matrix | CAPN1 | calpain 1 | 823 | down | 1,27 | 2,65E-02 |
| R-HSA-1474228 | Degradation of the extracellular matrix | MMP15 | matrix metallopeptidase 15 | 4324 | down | 1,49 | 4,35E-02 |
| R-HSA-1799339 | SRP-dependent cotranslational protein targeting to m | SRP68 | signal recognition particle 68 | 6730 | up | 1,40 | 3,02E-02 |
| R-HSA-1799339 | SRP-dependent cotranslational protein targeting to m | RPS4Y1 | ribosomal protein S4, Y-linked 1 | 6192 | down | 5,59 | 7,82E-03 |
| R-HSA-1799339 | SRP-dependent cotranslational protein targeting to m | SSR3 | signal sequence receptor subunit 3 | 6747 | up | 1,43 | 4,56E-02 |
| R-HSA-1799339 | SRP-dependent cotranslational protein targeting to m | TRAM1 | translocation associated membrane pro | 23471 | up | 1,76 | 4,37E-02 |
| R-HSA-1799339 | SRP-dependent cotranslational protein targeting to m | SRP72 | signal recognition particle 72 | 6731 | up | 1,43 | 9,37E-03 |
| R-HSA-1799339 | SRP-dependent cotranslational protein targeting to m | SSR1 | signal sequence receptor subunit 1 | 6745 | up | 1,73 | 2,06E-02 |
| R-HSA-1799339 | SRP-dependent cotranslational protein targeting to m | SRP19 | signal recognition particle 19 | 6728 | up | 1,68 | 4,13E-02 |
| R-HSA-1251985 | Nuclear signaling by ERBB4 | HBEGF | heparin binding EGF like growth factor | 1839 | up | 3,67 | 5,67E-03 |
| R-HSA-1251985 | Nuclear signaling by ERBB4 | ADAM17 | ADAM metallopeptidase domain 17 | 6868 | up | 2,59 | 1,60E-03 |
| R-HSA-1221632 | Meiotic synapsis | SYNE1 | spectrin repeat containing nuclear enve | 23345 | up | 3,01 | 1,11E-03 |
| R-HSA-1221632 | Meiotic synapsis | LMNB1 | lamin B1 | 4001 | down | 2,12 | 3,85E-02 |
| R-HSA-1221632 | Meiotic synapsis | DIDO1 | death inducer-obliterator 1 | 11083 | down | 1,37 | 4,03E-03 |
| R-HSA-1221632 | Meiotic synapsis | TERF2IP | TERF2 interacting protein | 54386 | up | 1,41 | 1,57E-02 |
| R-HSA-1221632 | Meiotic synapsis | STAG2 | stromal antigen 2 | 10735 | up | 1,26 | 3,69E-02 |
| R-HSA-1221632 | Meiotic synapsis | ACD | ACD, shelterin complex subunit and telo | 65057 | down | 1,25 | 2,08E-02 |
| R-HSA-1221632 | Meiotic synapsis | H2AFX | H2A histone family member X | 3014 | down | 2,11 | 1,30E-03 |
| R-HSA-194840 | Rho GTPase cycle | SYDE1 | synapse defective Rho GTPase homolo | 85360 | down | 1,99 | 3,40E-03 |
| R-HSA-194840 | Rho GTPase cycle | RHOF | ras homolog family member F, filopodia | 54509 | up | 1,79 | 3,66E-02 |
| R-HSA-194840 | Rho GTPase cycle | FGD4 | FYVE, RhoGEF and PH domain contair | 121512 | up | 1,60 | 1,34E-02 |
| R-HSA-194840 | Rho GTPase cycle | ARHGAP29 | Rho GTPase activating protein 29 | 9411 | up | 2,27 | 7,26E-03 |
| R-HSA-194840 | Rho GTPase cycle | ARAP1 | ArfGAP with RhoGAP domain, ankyrin r | 116985 | down | 1,24 | 4,90E-02 |
| R-HSA-194840 | Rho GTPase cycle | RHOB | ras homolog family member B | 388 | up | 1,65 | 2,44E-02 |
| R-HSA-194840 | Rho GTPase cycle | ARHGEF19 | Rho guanine nucleotide exchange facto | 128272 | down | 1,55 | 1,82E-02 |
| R-HSA-194840 | Rho GTPase cycle | SOS2 | SOS Ras/Rho guanine nucleotide exch | 6655 | up | 1,82 | 1,41E-02 |
| R-HSA-194840 | Rho GTPase cycle | RHOT2 | ras homolog family member T2 | 89941 | down | 1,19 | 2,93E-02 |
| R-HSA-194840 | Rho GTPase cycle | ARAP3 | ArfGAP with RhoGAP domain, ankyrin r | 64411 | down | 1,96 | 6,61E-03 |
| R-HSA-194840 | Rho GTPase cycle | RHOBTB2 | Rho related BTB domain containing 2 | 23221 | down | 2,29 | 7,15E-03 |
| R-HSA-194840 | Rho GTPase cycle | ARHGEF2 | Rho/Rac guanine nucleotide exchange | 9181 | up | 2,09 | 1,10E-02 |
| R-HSA-1663150 | The activation of arylsulfatases | ARSG | arylsulfatase G | 22901 | up | 1,36 | 3,94E-02 |
| R-HSA-1663150 | The activation of arylsulfatases | SUMF1 | sulfatase modifying factor 1 | 285362 | down | 1,29 | 3,58E-02 |
| R-HSA-75205 | Dissolution of Fibrin Clot | SERPINB8 | serpin family B member 8 | 5271 | up | 2,74 | 1,05E-02 |
| R-HSA-75205 | Dissolution of Fibrin Clot | PLAT | plasminogen activator, tissue type | 5327 | down | 6,67 | 2,47E-02 |
| R-HSA-888590 | GABA synthesis, release, reuptake and degradation | VAMP2 | vesicle associated membrane protein 2 | 6844 | up | 1,50 | 4,82E-02 |
| R-HSA-888590 | GABA synthesis, release, reuptake and degradation | RAB3A | RAB3A, member RAS oncogene family | 5864 | down | 1,80 | 4,63E-02 |
| R-HSA-3238698 | WNT ligand biogenesis and trafficking | WNT10B | Wnt family member 10B | 7480 | down | 1,76 | 4,78E-02 |
| R-HSA-3238698 | WNT ligand biogenesis and trafficking | TMED5 | transmembrane p24 trafficking protein 5 | 50999 | up | 1,48 | 4,72E-02 |
| R-HSA-3238698 | WNT ligand biogenesis and trafficking | VPS26A | VPS26, retromer complex component A | 9559 | up | 1,24 | 4,07E-02 |
| R-HSA-174403 | Glutathione synthesis and recycling | CNDP2 | carnosine dipeptidase 2 | 55748 | down | 1,20 | 4,78E-02 |
| R-HSA-174403 | Glutathione synthesis and recycling | GCLC | glutamate-cysteine ligase catalytic subu | 2729 | up | 2,09 | 2,27E-02 |
| R-HSA-1250347 | SHC1 events in ERBB4 signaling | HRAS | HRas proto-oncogene, GTPase | 3265 | down | 1,31 | 2,19E-03 |
| R-HSA-1250347 | SHC1 events in ERBB4 signaling | HBEGF | heparin binding EGF like growth factor | 1839 | up | 3,67 | 5,67E-03 |
| R-HSA-912526 | Interleukin receptor SHC signaling | PIK3CA | phosphatidylinositol-4,5-bisphosphate 3 | 5290 | up | 1,33 | 3,17E-02 |
| R-HSA-912526 | Interleukin receptor SHC signaling | PIK3R1 | phosphoinositide-3-kinase regulatory su | 5295 | down | 2,04 | 8,40E-03 |
| R-HSA-912526 | Interleukin receptor SHC signaling | JAK1 | Janus kinase 1 | 3716 | up | 1,53 | 2,13E-02 |
| R-HSA-5625740 | RHO GTPases activate PKNs | PPP1R12A | protein phosphatase 1 regulatory subur | 4659 | up | 1,25 | 1,03E-02 |
| R-HSA-5625740 | RHO GTPases activate PKNs | YWHAH | tyrosine 3-monooxygenase/tryptophan 5 | 7533 | down | 1,33 | 4,79E-02 |
| R-HSA-5625740 | RHO GTPases activate PKNs | RHOB | ras homolog family member B | 388 | up | 1,65 | 2,44E-02 |
| R-HSA-3000171 | Non-integrin membrane-ECM interactions | FGF2 | fibroblast growth factor 2 | 2247 | down | 5,35 | 3,09E-02 |
| R-HSA-3000171 | Non-integrin membrane-ECM interactions | DAG1 | dystroglycan 1 | 1605 | down | 1,65 | 1,04E-02 |
| R-HSA-3000171 | Non-integrin membrane-ECM interactions | LAMB3 | laminin subunit beta 3 | 3914 | up | 2,74 | 1,16E-02 |
| R-HSA-3000171 | Non-integrin membrane-ECM interactions | NTN4 | netrin 4 | 59277 | up | 2,22 | 5,93E-03 |
| R-HSA-180746 | Nuclear import of Rev protein | NUP62 | nucleoporin 62 | 23636 | down | 1,52 | 4,39E-03 |
| R-HSA-180746 | Nuclear import of Rev protein | RCC1 | regulator of chromosome condensation | 1104 | down | 1,51 | 6,96E-03 |
| R-HSA-180746 | Nuclear import of Rev protein | NUP58 | nucleoporin 58 | 9818 | up | 1,61 | 1,28E-02 |
| R-HSA-422356 | Regulation of insulin secretion | CACNB3 | calcium voltage-gated channel auxiliary | 784 | down | 1,19 | 3,30E-02 |
| R-HSA-422356 | Regulation of insulin secretion | GNAI2 | G protein subunit alpha i2 | 2771 | down | 1,22 | 3,65E-02 |
| R-HSA-422356 | Regulation of insulin secretion | VAMP2 | vesicle associated membrane protein 2 | 6844 | up | 1,50 | 4,82E-02 |
| R-HSA-420029 | Tight junction interactions | CLDN12 | claudin 12 | 9069 | up | 1,38 | 3,08E-02 |
| R-HSA-420029 | Tight junction interactions | PARD6A | par-6 family cell polarity regulator alpha | 50855 | down | 1,46 | 8,78E-03 |
| R-HSA-420029 | Tight junction interactions | CLDN1 | claudin 1 | 9076 | up | 2,13 | 6,96E-03 |
| R-HSA-5389840 | Mitochondrial translation elongation | MRPS26 | mitochondrial ribosomal protein S26 | 64949 | down | 1,49 | 2,65E-02 |
| R-HSA-5389840 | Mitochondrial translation elongation | MRPS34 | mitochondrial ribosomal protein S34 | 65993 | down | 1,51 | 1,07E-02 |
| R-HSA-5389840 | Mitochondrial translation elongation | MRPS12 | mitochondrial ribosomal protein S12 | 6183 | down | 1,29 | 9,88E-03 |
| R-HSA-5389840 | Mitochondrial translation elongation | MRPL4 | mitochondrial ribosomal protein L4 | 51073 | down | 1,33 | 1,11E-03 |
| R-HSA-5389840 | Mitochondrial translation elongation | MRPS10 | mitochondrial ribosomal protein S10 | 55173 | up | 1,41 | 1,62E-02 |
| R-HSA-5419276 | Mitochondrial translation termination | MRPS10 | mitochondrial ribosomal protein S10 | 55173 | up | 1,41 | 1,62E-02 |
| R-HSA-5419276 | Mitochondrial translation termination | MRPL4 | mitochondrial ribosomal protein L4 | 51073 | down | 1,33 | 1,11E-03 |
| R-HSA-5419276 | Mitochondrial translation termination | MRPS12 | mitochondrial ribosomal protein S12 | 6183 | down | 1,29 | 9,88E-03 |
| R-HSA-5419276 | Mitochondrial translation termination | MRPS34 | mitochondrial ribosomal protein S34 | 65993 | down | 1,51 | 1,07E-02 |
| R-HSA-5419276 | Mitochondrial translation termination | MRPS26 | mitochondrial ribosomal protein S26 | 64949 | down | 1,49 | 2,65E-02 |
| R-HSA-1222556 | ROS, RNS production in phagocytes | CYBA | cytochrome b-245 alpha chain | 1535 | down | 1,34 | 1,09E-02 |
| R-HSA-1222556 | ROS, RNS production in phagocytes | ATP6V0E2 | ATPase H+ transporting V0 subunit e2 | 155066 | down | 1,61 | 2,71E-03 |
| R-HSA-3299685 | Detoxification of Reactive Oxygen Species | CYBA | cytochrome b-245 alpha chain | 1535 | down | 1,34 | 1,09E-02 |
| R-HSA-3299685 | Detoxification of Reactive Oxygen Species | PRDX2 | peroxiredoxin 2 | 7001 | down | 1,26 | 3,14E-02 |
| R-HSA-201722 | Formation of the beta-catenin:TCF transactivating co | PYGO2 | pygopus family PHD finger 2 | 90780 | down | 1,37 | 1,36E-02 |
| R-HSA-201722 | Formation of the beta-catenin:TCF transactivating co | MEN1 | menin 1 | 4221 | down | 1,49 | 4,75E-04 |
| R-HSA-201722 | Formation of the beta-catenin:TCF transactivating co | CDC73 | cell division cycle 73 | 79577 | up | 1,37 | 4,41E-02 |
| R-HSA-201722 | Formation of the beta-catenin:TCF transactivating co | TCF7 | transcription factor 7 | 6932 | down | 1,47 | 1,07E-02 |
| R-HSA-201722 | Formation of the beta-catenin:TCF transactivating co | H2AFX | H2A histone family member X | 3014 | down | 2,11 | 1,30E-03 |
| R-HSA-391160 | Signal regulatory protein family interactions | SKAP2 | src kinase associated phosphoprotein 2 | 8935 | up | 1,19 | 3,74E-02 |
| R-HSA-391160 | Signal regulatory protein family interactions | SIRPA | signal regulatory protein alpha | 140885 | down | 2,17 | 1,00E-02 |
| R-HSA-4608870 | Asymmetric localization of PCP proteins | FZD4 | frizzled class receptor 4 | 8322 | down | 2,17 | 3,82E-02 |
| R-HSA-4608870 | Asymmetric localization of PCP proteins | SMURF2 | SMAD specific E3 ubiquitin protein ligas | 64750 | up | 1,49 | 4,16E-02 |
| R-HSA-4608870 | Asymmetric localization of PCP proteins | FZD1 | frizzled class receptor 1 | 8321 | down | 1,75 | 2,84E-02 |
| R-HSA-4608870 | Asymmetric localization of PCP proteins | PARD6A | par-6 family cell polarity regulator alpha | 50855 | down | 1,46 | 8,78E-03 |
| R-HSA-5334118 | DNA methylation | DNMT3B | DNA methyltransferase 3 beta | 1789 | down | 1,70 | 3,62E-02 |
| R-HSA-5334118 | DNA methylation | H2AFX | H2A histone family member X | 3014 | down | 2,11 | 1,30E-03 |
| R-HSA-5334118 | DNA methylation | UHRF1 | ubiquitin like with PHD and ring finger do | 29128 | down | 2,85 | 1,96E-02 |
| R-HSA-5678895 | Defective CFTR causes cystic fibrosis | DERL2 | derlin 2 | 51009 | up | 1,68 | 4,88E-02 |
| R-HSA-5678895 | Defective CFTR causes cystic fibrosis | SEL1L | SEL1L ERAD E3 ligase adaptor subunit | 6400 | up | 2,76 | 4,03E-03 |
| R-HSA-5678895 | Defective CFTR causes cystic fibrosis | ERLEC1 | endoplasmic reticulum lectin 1 | 27248 | up | 1,96 | 7,71E-03 |
| R-HSA-5632684 | Hedgehog 'on' state | SUFU | SUFU negative regulator of hedgehog s | 51684 | down | 1,41 | 3,37E-02 |
| R-HSA-5632684 | Hedgehog 'on' state | CDC73 | cell division cycle 73 | 79577 | up | 1,37 | 4,41E-02 |
| R-HSA-5632684 | Hedgehog 'on' state | ITCH | itchy E3 ubiquitin protein ligase | 83737 | up | 1,51 | 6,64E-03 |
| R-HSA-5632684 | Hedgehog 'on' state | ULK3 | unc-51 like kinase 3 | 25989 | down | 1,22 | 4,13E-02 |
| R-HSA-5632684 | Hedgehog 'on' state | SMURF2 | SMAD specific E3 ubiquitin protein ligas | 64750 | up | 1,49 | 4,16E-02 |
| R-HSA-5632684 | Hedgehog 'on' state | GPR161 | G protein-coupled receptor 161 | 23432 | down | 2,43 | 1,49E-03 |
| R-HSA-190861 | Gap junction assembly | TUBA1B | tubulin alpha 1b | 10376 | down | 1,78 | 2,30E-02 |
| R-HSA-190861 | Gap junction assembly | TUBA4A | tubulin alpha 4a | 7277 | down | 1,63 | 2,97E-02 |
| R-HSA-937072 | TRAF6-mediated induction of TAK1 complex within TI | TAB1 | TGF-beta activated kinase 1 (MAP3K7) | 10454 | down | 1,29 | 1,94E-02 |
| R-HSA-937072 | TRAF6-mediated induction of TAK1 complex within TI | TAB3 | TGF-beta activated kinase 1 and MAP3 | 257397 | up | 1,40 | 4,94E-02 |
| R-HSA-432142 | Platelet sensitization by LDL | LRP8 | LDL receptor related protein 8 | 7804 | down | 1,85 | 4,38E-02 |
| R-HSA-432142 | Platelet sensitization by LDL | PPP2CB | protein phosphatase 2 catalytic subunit | 5516 | up | 1,30 | 4,79E-02 |
| R-HSA-3301854 | Nuclear Pore Complex (NPC) Disassembly | NUP62 | nucleoporin 62 | 23636 | down | 1,52 | 4,39E-03 |
| R-HSA-3301854 | Nuclear Pore Complex (NPC) Disassembly | NEK7 | NIMA related kinase 7 | 140609 | up | 1,48 | 2,18E-02 |
| R-HSA-3301854 | Nuclear Pore Complex (NPC) Disassembly | NUP58 | nucleoporin 58 | 9818 | up | 1,61 | 1,28E-02 |
| R-HSA-5687128 | MAPK6/MAPK4 signaling | DNAJB1 | DnaJ heat shock protein family (Hsp40) | 3337 | up | 2,02 | 1,41E-02 |
| R-HSA-5687128 | MAPK6/MAPK4 signaling | MAPK6 | mitogen-activated protein kinase 6 | 5597 | up | 2,04 | 5,54E-03 |
| R-HSA-5687128 | MAPK6/MAPK4 signaling | AGO3 | argonaute 3, RISC catalytic component | 192669 | up | 1,36 | 1,80E-02 |
| R-HSA-5687128 | MAPK6/MAPK4 signaling | CCND3 | cyclin D3 | 896 | down | 1,60 | 4,17E-02 |
| R-HSA-5687128 | MAPK6/MAPK4 signaling | NCOA3 | nuclear receptor coactivator 3 | 8202 | up | 1,53 | 4,13E-02 |
| R-HSA-5663213 | RHO GTPases Activate WASPs and WAVEs | NCK1 | NCK adaptor protein 1 | 4690 | up | 1,48 | 1,42E-02 |
| R-HSA-5663213 | RHO GTPases Activate WASPs and WAVEs | ACTG1 | actin gamma 1 | 71 | up | 1,58 | 1,66E-02 |
| R-HSA-416476 | G alpha (q) signalling events | GNA11 | G protein subunit alpha 11 | 2767 | down | 1,18 | 4,37E-02 |
| R-HSA-416476 | G alpha (q) signalling events | PIK3CA | phosphatidylinositol-4,5-bisphosphate 3 | 5290 | up | 1,33 | 3,17E-02 |
| R-HSA-416476 | G alpha (q) signalling events | HRH1 | histamine receptor H1 | 3269 | up | 1,51 | 2,03E-02 |
| R-HSA-416476 | G alpha (q) signalling events | NPFFR1 | neuropeptide FF receptor 1 | 64106 | down | 2,16 | 3,35E-02 |
| R-HSA-416476 | G alpha (q) signalling events | F2R | coagulation factor II thrombin receptor | 2149 | down | 1,54 | 7,69E-03 |
| R-HSA-416476 | G alpha (q) signalling events | ANXA1 | annexin A1 | 301 | up | 2,22 | 9,88E-03 |
| R-HSA-416476 | G alpha (q) signalling events | RGS19 | regulator of G protein signaling 19 | 10287 | down | 1,45 | 1,44E-03 |
| R-HSA-416476 | G alpha (q) signalling events | PIK3R1 | phosphoinositide-3-kinase regulatory su | 5295 | down | 2,04 | 8,40E-03 |
| R-HSA-416476 | G alpha (q) signalling events | NMB | neuromedin B | 4828 | down | 1,31 | 2,56E-02 |
| R-HSA-416476 | G alpha (q) signalling events | RGS3 | regulator of G protein signaling 3 | 5998 | up | 1,77 | 1,19E-02 |
| R-HSA-416476 | G alpha (q) signalling events | P2RY11 | purinergic receptor P2Y11 | 5032 | down | 1,54 | 2,90E-02 |
| R-HSA-3214815 | HDACs deacetylate histones | TBL1X | transducin beta like 1 X-linked | 6907 | down | 1,20 | 2,84E-02 |
| R-HSA-3214815 | HDACs deacetylate histones | CHD4 | chromodomain helicase DNA binding pr | 1108 | down | 1,44 | 9,37E-03 |
| R-HSA-3214815 | HDACs deacetylate histones | MTA2 | metastasis associated 1 family member | 9219 | down | 1,27 | 2,45E-02 |
| R-HSA-3214815 | HDACs deacetylate histones | MBD3 | methyl-CpG binding domain protein 3 | 53615 | down | 1,52 | 4,26E-04 |
| R-HSA-5685938 | HDR through Single Strand Annealing (SSA) | NBN | nibrin | 4683 | up | 1,25 | 3,15E-02 |
| R-HSA-5685938 | HDR through Single Strand Annealing (SSA) | HUS1 | HUS1 checkpoint clamp component | 3364 | up | 1,36 | 4,96E-02 |
| R-HSA-5685938 | HDR through Single Strand Annealing (SSA) | RPA1 | replication protein A1 | 6117 | down | 1,41 | 4,93E-02 |
| R-HSA-72706 | GTP hydrolysis and joining of the 60S ribosomal subu | EIF3M | eukaryotic translation initiation factor 3 s | 10480 | up | 1,25 | 2,51E-02 |
| R-HSA-72706 | GTP hydrolysis and joining of the 60S ribosomal subu | EIF3B | eukaryotic translation initiation factor 3 s | 8662 | down | 1,27 | 1,76E-02 |
| R-HSA-72706 | GTP hydrolysis and joining of the 60S ribosomal subu | EIF3H | eukaryotic translation initiation factor 3 s | 8667 | up | 1,22 | 3,84E-02 |
| R-HSA-72706 | GTP hydrolysis and joining of the 60S ribosomal subu | EIF2S2 | eukaryotic translation initiation factor 2 s | 8894 | up | 1,40 | 2,40E-02 |
| R-HSA-72706 | GTP hydrolysis and joining of the 60S ribosomal subu | EIF4A2 | eukaryotic translation initiation factor 4A | 1974 | up | 1,68 | 4,70E-03 |
| R-HSA-72706 | GTP hydrolysis and joining of the 60S ribosomal subu | RPS4Y1 | ribosomal protein S4, Y-linked 1 | 6192 | down | 5,59 | 7,82E-03 |
| R-HSA-72706 | GTP hydrolysis and joining of the 60S ribosomal subu | EIF2S3 | eukaryotic translation initiation factor 2 s | 1968 | up | 1,19 | 2,93E-02 |
| R-HSA-1660514 | Synthesis of PIPs at the Golgi membrane | ARF3 | ADP ribosylation factor 3 | 377 | down | 1,30 | 4,57E-02 |
| R-HSA-1660514 | Synthesis of PIPs at the Golgi membrane | PIK3C2A | phosphatidylinositol-4-phosphate 3-kina | 5286 | up | 1,30 | 1,79E-02 |
| R-HSA-156827 | L13a-mediated translational silencing of Ceruloplasmi | EIF2S3 | eukaryotic translation initiation factor 2 s | 1968 | up | 1,19 | 2,93E-02 |
| R-HSA-156827 | L13a-mediated translational silencing of Ceruloplasmi | EIF4A2 | eukaryotic translation initiation factor 4A | 1974 | up | 1,68 | 4,70E-03 |
| R-HSA-156827 | L13a-mediated translational silencing of Ceruloplasmi | RPS4Y1 | ribosomal protein S4, Y-linked 1 | 6192 | down | 5,59 | 7,82E-03 |
| R-HSA-156827 | L13a-mediated translational silencing of Ceruloplasmi | EIF3H | eukaryotic translation initiation factor 3 s | 8667 | up | 1,22 | 3,84E-02 |
| R-HSA-156827 | L13a-mediated translational silencing of Ceruloplasmi | EIF3B | eukaryotic translation initiation factor 3 s | 8662 | down | 1,27 | 1,76E-02 |
| R-HSA-156827 | L13a-mediated translational silencing of Ceruloplasmi | EIF2S2 | eukaryotic translation initiation factor 2 s | 8894 | up | 1,40 | 2,40E-02 |
| R-HSA-156827 | L13a-mediated translational silencing of Ceruloplasmi | EIF3M | eukaryotic translation initiation factor 3 s | 10480 | up | 1,25 | 2,51E-02 |
| R-HSA-2559586 | DNA Damage/Telomere Stress Induced Senescence | CDKN1A | cyclin dependent kinase inhibitor 1A | 1026 | up | 3,62 | 5,43E-03 |
| R-HSA-2559586 | DNA Damage/Telomere Stress Induced Senescence | NBN | nibrin | 4683 | up | 1,25 | 3,15E-02 |
| R-HSA-2559586 | DNA Damage/Telomere Stress Induced Senescence | TERF2IP | TERF2 interacting protein | 54386 | up | 1,41 | 1,57E-02 |
| R-HSA-2559586 | DNA Damage/Telomere Stress Induced Senescence | H2AFX | H2A histone family member X | 3014 | down | 2,11 | 1,30E-03 |
| R-HSA-2559586 | DNA Damage/Telomere Stress Induced Senescence | ACD | ACD, shelterin complex subunit and telo | 65057 | down | 1,25 | 2,08E-02 |
| R-HSA-5693616 | Presynaptic phase of homologous DNA pairing and s | RPA1 | replication protein A1 | 6117 | down | 1,41 | 4,93E-02 |
| R-HSA-5693616 | Presynaptic phase of homologous DNA pairing and s | NBN | nibrin | 4683 | up | 1,25 | 3,15E-02 |
| R-HSA-5693616 | Presynaptic phase of homologous DNA pairing and s | HUS1 | HUS1 checkpoint clamp component | 3364 | up | 1,36 | 4,96E-02 |
| R-HSA-427359 | SIRT1 negatively regulates rRNA expression | H2AFX | H2A histone family member X | 3014 | down | 2,11 | 1,30E-03 |
| R-HSA-427359 | SIRT1 negatively regulates rRNA expression | SIRT1 | sirtuin 1 | 23411 | up | 1,60 | 1,00E-02 |
| R-HSA-427359 | SIRT1 negatively regulates rRNA expression | TAF1D | TATA-box binding protein associated fa | 79101 | up | 1,87 | 1,00E-02 |
| R-HSA-427359 | SIRT1 negatively regulates rRNA expression | TAF1A | TATA-box binding protein associated fa | 9015 | up | 2,20 | 2,24E-03 |
| R-HSA-427359 | SIRT1 negatively regulates rRNA expression | SUV39H1 | suppressor of variegation 3-9 homolog 1 | 6839 | down | 1,55 | 1,62E-02 |
| R-HSA-2559580 | Oxidative Stress Induced Senescence | CBX8 | chromobox 8 | 57332 | down | 1,30 | 2,19E-02 |
| R-HSA-2559580 | Oxidative Stress Induced Senescence | MAPK8 | mitogen-activated protein kinase 8 | 5599 | up | 1,38 | 3,05E-02 |
| R-HSA-2559580 | Oxidative Stress Induced Senescence | AGO3 | argonaute 3, RISC catalytic component | 192669 | up | 1,36 | 1,80E-02 |
| R-HSA-2559580 | Oxidative Stress Induced Senescence | H2AFX | H2A histone family member X | 3014 | down | 2,11 | 1,30E-03 |
| R-HSA-2559580 | Oxidative Stress Induced Senescence | TFDP1 | transcription factor Dp-1 | 7027 | down | 1,77 | 1,08E-02 |
| R-HSA-2559580 | Oxidative Stress Induced Senescence | E2F1 | E2F transcription factor 1 | 1869 | down | 1,64 | 3,25E-02 |
| R-HSA-2559580 | Oxidative Stress Induced Senescence | CDKN2B | cyclin dependent kinase inhibitor 2B | 1030 | up | 2,40 | 3,90E-02 |
| R-HSA-2559580 | Oxidative Stress Induced Senescence | CBX4 | chromobox 4 | 8535 | up | 2,70 | 4,00E-03 |
| R-HSA-2559580 | Oxidative Stress Induced Senescence | CBX2 | chromobox 2 | 84733 | down | 2,70 | 8,09E-03 |
| R-HSA-2559580 | Oxidative Stress Induced Senescence | KDM6B | lysine demethylase 6B | 23135 | up | 2,16 | 2,06E-02 |
| R-HSA-416700 | Other semaphorin interactions | SEMA7A | semaphorin 7A (John Milton Hagen bloo | 8482 | up | 2,44 | 8,66E-03 |
| R-HSA-416700 | Other semaphorin interactions | PLXNA1 | plexin A1 | 5361 | down | 1,28 | 9,70E-04 |
| R-HSA-3928664 | Ephrin signaling | SDCBP | syndecan binding protein | 6386 | up | 1,44 | 8,07E-03 |
| R-HSA-3928664 | Ephrin signaling | EPHB2 | EPH receptor B2 | 2048 | down | 1,66 | 2,52E-02 |
| R-HSA-159236 | Transport of Mature mRNA derived from an Intron-Co | NUP62 | nucleoporin 62 | 23636 | down | 1,52 | 4,39E-03 |
| R-HSA-159236 | Transport of Mature mRNA derived from an Intron-Co | U2AF2 | U2 small nuclear RNA auxiliary factor 2 | 11338 | down | 1,27 | 8,21E-03 |
| R-HSA-159236 | Transport of Mature mRNA derived from an Intron-Co | SRSF2 | serine and arginine rich splicing factor 2 | 6427 | down | 1,69 | 1,79E-02 |
| R-HSA-159236 | Transport of Mature mRNA derived from an Intron-Co | NUP58 | nucleoporin 58 | 9818 | up | 1,61 | 1,28E-02 |
| R-HSA-159236 | Transport of Mature mRNA derived from an Intron-Co | SLU7 | SLU7 homolog, splicing factor | 10569 | up | 1,24 | 3,63E-02 |
| R-HSA-159236 | Transport of Mature mRNA derived from an Intron-Co | SARNP | SAP domain containing ribonucleoprote | 84324 | up | 1,48 | 4,35E-02 |
| R-HSA-606279 | Deposition of new CENPA-containing nucleosomes a | RSF1 | remodeling and spacing factor 1 | 51773 | up | 1,26 | 3,93E-02 |
| R-HSA-606279 | Deposition of new CENPA-containing nucleosomes a | CENPO | centromere protein O | 79172 | down | 1,38 | 4,86E-02 |
| R-HSA-606279 | Deposition of new CENPA-containing nucleosomes a | H2AFX | H2A histone family member X | 3014 | down | 2,11 | 1,30E-03 |
| R-HSA-606279 | Deposition of new CENPA-containing nucleosomes a | CENPX | centromere protein X | 201254 | down | 1,50 | 2,52E-02 |
| R-HSA-5223345 | Miscellaneous transport and binding events | MAGT1 | magnesium transporter 1 | 84061 | up | 1,54 | 4,98E-02 |
| R-HSA-5223345 | Miscellaneous transport and binding events | ANKH | ANKH inorganic pyrophosphate transpo | 56172 | down | 1,24 | 4,14E-02 |
| R-HSA-1483191 | Synthesis of PC | LPIN2 | lipin 2 | 9663 | up | 2,16 | 6,88E-03 |
| R-HSA-1483191 | Synthesis of PC | LPIN3 | lipin 3 | 64900 | down | 1,36 | 4,76E-02 |
| R-HSA-202424 | Downstream TCR signaling | PTEN | phosphatase and tensin homolog | 5728 | up | 1,33 | 4,72E-02 |
| R-HSA-202424 | Downstream TCR signaling | BCL10 | B-cell CLL/lymphoma 10 | 8915 | up | 2,06 | 5,25E-04 |
| R-HSA-202424 | Downstream TCR signaling | RIPK2 | receptor interacting serine/threonine kin | 8767 | up | 1,72 | 8,31E-03 |
| R-HSA-202424 | Downstream TCR signaling | PIK3CA | phosphatidylinositol-4,5-bisphosphate 3 | 5290 | up | 1,33 | 3,17E-02 |
| R-HSA-202424 | Downstream TCR signaling | PIK3R1 | phosphoinositide-3-kinase regulatory su | 5295 | down | 2,04 | 8,40E-03 |
| R-HSA-428930 | Thromboxane signalling through TP receptor | GNA13 | G protein subunit alpha 13 | 10672 | up | 1,68 | 2,41E-03 |

| | | | | | | | |
|---|---|---|---|---|---|---|---|
| R-HSA-428930 | Thromboxane signalling through TP receptor | GNA11 | G protein subunit alpha 11 | 2767 | down | 1,18 | 4,37E-02 |
| R-HSA-2559582 | Senescence-Associated Secretory Phenotype (SASP | CEBPB | CCAAT/enhancer binding protein beta | 1051 | up | 2,12 | 2,85E-02 |
| R-HSA-2559582 | Senescence-Associated Secretory Phenotype (SASP | IGFBP7 | insulin like growth factor binding protein | 3490 | down | 1,86 | 2,80E-02 |
| R-HSA-2559582 | Senescence-Associated Secretory Phenotype (SASP | H2AFX | H2A histone family member X | 3014 | down | 2,11 | 1,30E-03 |
| R-HSA-2559582 | Senescence-Associated Secretory Phenotype (SASP | CDKN1A | cyclin dependent kinase inhibitor 1A | 1026 | up | 3,62 | 5,43E-03 |
| R-HSA-2559582 | Senescence-Associated Secretory Phenotype (SASP | IL1A | interleukin 1 alpha | 3552 | up | 8,56 | 1,05E-02 |
| R-HSA-2559582 | Senescence-Associated Secretory Phenotype (SASP | CDKN2B | cyclin dependent kinase inhibitor 2B | 1030 | up | 2,40 | 3,90E-02 |
| R-HSA-2559582 | Senescence-Associated Secretory Phenotype (SASP | ANAPC2 | anaphase promoting complex subunit 2 | 29882 | down | 1,26 | 1,99E-02 |
| R-HSA-2029485 | Role of phospholipids in phagocytosis | PIK3CA | phosphatidylinositol-4,5-bisphosphate 3 | 5290 | up | 1,33 | 3,17E-02 |
| R-HSA-2029485 | Role of phospholipids in phagocytosis | PLPP5 | phospholipid phosphatase 5 | 84513 | up | 1,85 | 4,15E-02 |
| R-HSA-2029485 | Role of phospholipids in phagocytosis | PLCG1 | phospholipase C gamma 1 | 5335 | down | 1,51 | 2,85E-02 |
| R-HSA-2029485 | Role of phospholipids in phagocytosis | PIK3R1 | phosphoinositide-3-kinase regulatory su | 5295 | down | 2,04 | 8,40E-03 |
| R-HSA-2029485 | Role of phospholipids in phagocytosis | PLD1 | phospholipase D1 | 5337 | up | 1,33 | 1,60E-02 |
| R-HSA-171306 | Packaging Of Telomere Ends | ACD | ACD, shelterin complex subunit and telo | 65057 | down | 1,25 | 2,08E-02 |
| R-HSA-171306 | Packaging Of Telomere Ends | H2AFX | H2A histone family member X | 3014 | down | 2,11 | 1,30E-03 |
| R-HSA-171306 | Packaging Of Telomere Ends | TERF2IP | TERF2 interacting protein | 54386 | up | 1,41 | 1,57E-02 |
| R-HSA-512988 | Interleukin-3, Interleukin-5 and GM-CSF signaling | PIK3CA | phosphatidylinositol-4,5-bisphosphate 3 | 5290 | up | 1,33 | 3,17E-02 |
| R-HSA-512988 | Interleukin-3, Interleukin-5 and GM-CSF signaling | PIK3R1 | phosphoinositide-3-kinase regulatory su | 5295 | down | 2,04 | 8,40E-03 |
| R-HSA-202733 | Cell surface interactions at the vascular wall | SIRPA | signal regulatory protein alpha | 140885 | down | 2,17 | 1,00E-02 |
| R-HSA-202733 | Cell surface interactions at the vascular wall | LYN | LYN proto-oncogene, Src family tyrosine | 4067 | down | 1,36 | 2,34E-02 |
| R-HSA-2871796 | FCERI mediated MAPK activation | MAPK8 | mitogen-activated protein kinase 8 | 5599 | up | 1,38 | 3,05E-02 |
| R-HSA-2871796 | FCERI mediated MAPK activation | PLCG1 | phospholipase C gamma 1 | 5335 | down | 1,51 | 2,85E-02 |
| R-HSA-2871796 | FCERI mediated MAPK activation | LYN | LYN proto-oncogene, Src family tyrosine | 4067 | down | 1,36 | 2,34E-02 |
| R-HSA-2871796 | FCERI mediated MAPK activation | HRAS | HRas proto-oncogene, GTPase | 3265 | down | 1,31 | 2,19E-03 |
| R-HSA-418555 | G alpha (s) signalling events | P2RY11 | purinergic receptor P2Y11 | 5032 | down | 1,54 | 2,90E-02 |
| R-HSA-418555 | G alpha (s) signalling events | GNAI2 | G protein subunit alpha i2 | 2771 | down | 1,22 | 3,65E-02 |
| R-HSA-418555 | G alpha (s) signalling events | PTGER4 | prostaglandin E receptor 4 | 5734 | up | 2,03 | 2,72E-03 |
| R-HSA-418555 | G alpha (s) signalling events | PDE4B | phosphodiesterase 4B | 5142 | down | 2,82 | 4,90E-02 |
| R-HSA-418555 | G alpha (s) signalling events | ADM | adrenomedullin | 133 | up | 1,84 | 4,75E-04 |
| R-HSA-418555 | G alpha (s) signalling events | ARRB1 | arrestin beta 1 | 408 | down | 1,79 | 2,42E-02 |
| R-HSA-418555 | G alpha (s) signalling events | ADCY3 | adenylate cyclase 3 | 109 | down | 1,45 | 1,03E-02 |
| R-HSA-418555 | G alpha (s) signalling events | ADM2 | adrenomedullin 2 | 79924 | up | 3,79 | 7,16E-03 |
| R-HSA-913709 | O-linked glycosylation of mucins | C1GALT1 | core 1 synthase, glycoprotein-N-acetylga | 56913 | up | 1,51 | 1,69E-02 |
| R-HSA-913709 | O-linked glycosylation of mucins | B3GNT7 | UDP-GlcNAc:betaGal beta-1,3-N-acetylg | 93010 | down | 2,40 | 3,48E-02 |
| R-HSA-913709 | O-linked glycosylation of mucins | B3GNT5 | UDP-GlcNAc:betaGal beta-1,3-N-acetylg | 84002 | up | 1,57 | 1,69E-02 |
| R-HSA-5576893 | Phase 2 - plateau phase | AKAP9 | A-kinase anchoring protein 9 | 10142 | up | 1,47 | 1,80E-02 |
| R-HSA-5576893 | Phase 2 - plateau phase | CACNB3 | calcium voltage-gated channel auxiliary | 784 | down | 1,19 | 3,30E-02 |
| R-HSA-389957 | Prefoldin mediated transfer of substrate to CCT/TriC | TUBA4A | tubulin alpha 4a | 7277 | down | 1,63 | 2,97E-02 |
| R-HSA-389957 | Prefoldin mediated transfer of substrate to CCT/TriC | PFDN2 | prefoldin subunit 2 | 5202 | up | 1,66 | 1,44E-02 |
| R-HSA-1266695 | Interleukin-7 signaling | PIK3R1 | phosphoinositide-3-kinase regulatory su | 5295 | down | 2,04 | 8,40E-03 |
| R-HSA-1266695 | Interleukin-7 signaling | JAK1 | Janus kinase 1 | 3716 | up | 1,53 | 2,13E-02 |
| R-HSA-73728 | RNA Polymerase I Promoter Opening | UBTF | upstream binding transcription factor, RN | 7343 | down | 1,66 | 1,20E-02 |
| R-HSA-73728 | RNA Polymerase I Promoter Opening | H2AFX | H2A histone family member X | 3014 | down | 2,11 | 1,30E-03 |
| R-HSA-3214841 | PKMTs methylate histone lysines | WDR5 | WD repeat domain 5 | 11091 | down | 1,28 | 3,81E-03 |
| R-HSA-3214841 | PKMTs methylate histone lysines | SUV39H1 | suppressor of variegation 3-9 homolog 1 | 6839 | down | 1,55 | 1,62E-02 |
| R-HSA-1482839 | Acyl chain remodelling of PE | LPCAT4 | lysophosphatidylcholine acyltransferase | 254531 | down | 1,78 | 2,52E-02 |
| R-HSA-1482839 | Acyl chain remodelling of PE | PNPLA8 | patatin like phospholipase domain conta | 50640 | up | 2,48 | 3,36E-03 |
| R-HSA-195253 | Degradation of beta-catenin by the destruction comp | AMER1 | APC membrane recruitment protein 1 | 139285 | down | 1,78 | 4,15E-02 |
| R-HSA-195253 | Degradation of beta-catenin by the destruction comp | PPP2CB | protein phosphatase 2 catalytic subunit | 5516 | up | 1,30 | 4,79E-02 |
| R-HSA-195253 | Degradation of beta-catenin by the destruction comp | CSNK1A1 | casein kinase 1 alpha 1 | 1452 | up | 1,35 | 2,84E-02 |
| R-HSA-909733 | Interferon alpha/beta signaling | JAK1 | Janus kinase 1 | 3716 | up | 1,53 | 2,13E-02 |
| R-HSA-909733 | Interferon alpha/beta signaling | IRF3 | interferon regulatory factor 3 | 3661 | down | 1,33 | 2,58E-03 |
| R-HSA-909733 | Interferon alpha/beta signaling | SAMHD1 | SAM and HD domain containing deoxyn | 25939 | down | 1,62 | 4,03E-03 |
| R-HSA-170822 | Regulation of Glucokinase by Glucokinase Regulator | NUP58 | nucleoporin 58 | 9818 | up | 1,61 | 1,28E-02 |
| R-HSA-170822 | Regulation of Glucokinase by Glucokinase Regulator | NUP62 | nucleoporin 62 | 23636 | down | 1,52 | 4,39E-03 |
| R-HSA-5658442 | Regulation of RAS by GAPs | HRAS | HRas proto-oncogene, GTPase | 3265 | down | 1,31 | 2,19E-03 |
| R-HSA-5658442 | Regulation of RAS by GAPs | DAB2IP | DAB2 interacting protein | 153090 | down | 1,62 | 1,57E-02 |
| R-HSA-5668541 | TNFR2 non-canonical NF-kB pathway | TRAF3 | TNF receptor associated factor 3 | 7187 | down | 1,40 | 1,36E-02 |
| R-HSA-5668541 | TNFR2 non-canonical NF-kB pathway | TRAF2 | TNF receptor associated factor 2 | 7186 | down | 1,47 | 2,65E-02 |
| R-HSA-5668541 | TNFR2 non-canonical NF-kB pathway | BIRC2 | baculoviral IAP repeat containing 2 | 329 | up | 1,38 | 1,86E-02 |
| R-HSA-375276 | Peptide ligand-binding receptors | NMB | neuromedin B | 4828 | down | 1,31 | 2,56E-02 |
| R-HSA-375276 | Peptide ligand-binding receptors | F2R | coagulation factor II thrombin receptor | 2149 | down | 1,54 | 7,69E-03 |
| R-HSA-375276 | Peptide ligand-binding receptors | APLN | apelin | 8862 | down | 3,89 | 3,02E-02 |
| R-HSA-418990 | Adherens junctions interactions | NECTIN1 | nectin cell adhesion molecule 1 | 5818 | down | 1,48 | 7,71E-03 |
| R-HSA-418990 | Adherens junctions interactions | ACTG1 | actin gamma 1 | 71 | up | 1,58 | 1,66E-02 |
| R-HSA-2730905 | Role of LAT2/NTAL/LAB on calcium mobilization | PIK3R1 | phosphoinositide-3-kinase regulatory su | 5295 | down | 2,04 | 8,40E-03 |
| R-HSA-2730905 | Role of LAT2/NTAL/LAB on calcium mobilization | PIK3CA | phosphatidylinositol-4,5-bisphosphate 3 | 5290 | up | 1,33 | 3,17E-02 |
| R-HSA-2730905 | Role of LAT2/NTAL/LAB on calcium mobilization | LYN | LYN proto-oncogene, Src family tyrosine | 4067 | down | 1,36 | 2,34E-02 |
| R-HSA-418594 | G alpha (i) signalling events | GNAI2 | G protein subunit alpha i2 | 2771 | down | 1,22 | 3,65E-02 |
| R-HSA-418594 | G alpha (i) signalling events | ADRA2A | adrenoceptor alpha 2A | 150 | down | 3,92 | 2,45E-02 |
| R-HSA-418594 | G alpha (i) signalling events | S1PR2 | sphingosine-1-phosphate receptor 2 | 9294 | down | 1,91 | 1,41E-02 |
| R-HSA-418594 | G alpha (i) signalling events | APLN | apelin | 8862 | down | 3,89 | 3,02E-02 |
| R-HSA-418594 | G alpha (i) signalling events | RGS19 | regulator of G protein signaling 19 | 10287 | down | 1,45 | 1,44E-03 |
| R-HSA-418594 | G alpha (i) signalling events | ANXA1 | annexin A1 | 301 | up | 2,22 | 9,88E-03 |
| R-HSA-418594 | G alpha (i) signalling events | RGS16 | regulator of G protein signaling 16 | 6004 | up | 2,69 | 4,57E-03 |
| R-HSA-418594 | G alpha (i) signalling events | RGS14 | regulator of G protein signaling 14 | 10636 | down | 1,61 | 4,72E-02 |
| R-HSA-418594 | G alpha (i) signalling events | RGS12 | regulator of G protein signaling 12 | 6002 | down | 1,37 | 2,75E-02 |
| R-HSA-418594 | G alpha (i) signalling events | ADCY3 | adenylate cyclase 3 | 109 | down | 1,45 | 1,03E-02 |
| R-HSA-174178 | APC/C:Cdh1 mediated degradation of Cdc20 and oth | SKP2 | S-phase kinase associated protein 2 | 6502 | down | 2,21 | 3,69E-02 |
| R-HSA-174178 | APC/C:Cdh1 mediated degradation of Cdc20 and oth | ANAPC2 | anaphase promoting complex subunit 2 | 29882 | down | 1,26 | 1,99E-02 |
| R-HSA-1592389 | Activation of Matrix Metalloproteinases | PRSS1 | protease, serine 1 | 5644 | up | 2,54 | 4,99E-02 |
| R-HSA-1592389 | Activation of Matrix Metalloproteinases | MMP15 | matrix metallopeptidase 15 | 4324 | down | 1,49 | 4,35E-02 |
| R-HSA-180910 | Vpr-mediated nuclear import of PICs | NUP58 | nucleoporin 58 | 9818 | up | 1,61 | 1,28E-02 |
| R-HSA-180910 | Vpr-mediated nuclear import of PICs | NUP62 | nucleoporin 62 | 23636 | down | 1,52 | 4,39E-03 |
| R-HSA-212300 | PRC2 methylates histones and DNA | H2AFX | H2A histone family member X | 3014 | down | 2,11 | 1,30E-03 |
| R-HSA-212300 | PRC2 methylates histones and DNA | DNMT3B | DNA methyltransferase 3 beta | 1789 | down | 1,70 | 3,62E-02 |
| R-HSA-5693565 | Recruitment and ATM-mediated phosphorylation of re | PPP5C | protein phosphatase 5 catalytic subunit | 5536 | down | 1,29 | 4,15E-02 |
| R-HSA-5693565 | Recruitment and ATM-mediated phosphorylation of re | H2AFX | H2A histone family member X | 3014 | down | 2,11 | 1,30E-03 |
| R-HSA-5693565 | Recruitment and ATM-mediated phosphorylation of re | MAPK8 | mitogen-activated protein kinase 8 | 5599 | up | 1,38 | 3,05E-02 |
| R-HSA-5693565 | Recruitment and ATM-mediated phosphorylation of re | NBN | nibrin | 4683 | up | 1,25 | 3,15E-02 |
| R-HSA-202433 | Generation of second messenger molecules | NCK1 | NCK adaptor protein 1 | 4690 | up | 1,48 | 1,42E-02 |
| R-HSA-202433 | Generation of second messenger molecules | PLCG1 | phospholipase C gamma 1 | 5335 | down | 1,51 | 2,85E-02 |
| R-HSA-5607764 | CLEC7A (Dectin-1) signaling | TAB3 | TGF-beta activated kinase 1 and MAP3 | 257397 | up | 1,40 | 4,94E-02 |
| R-HSA-5607764 | CLEC7A (Dectin-1) signaling | TAB1 | TGF-beta activated kinase 1 (MAP3K7) | 10454 | down | 1,29 | 1,94E-02 |
| R-HSA-5607764 | CLEC7A (Dectin-1) signaling | BCL10 | B-cell CLL/lymphoma 10 | 8915 | up | 2,06 | 5,25E-04 |
| R-HSA-69473 | G2/M DNA damage checkpoint | HUS1 | HUS1 checkpoint clamp component | 3364 | up | 1,36 | 4,96E-02 |
| R-HSA-69473 | G2/M DNA damage checkpoint | NBN | nibrin | 4683 | up | 1,25 | 3,15E-02 |
| R-HSA-69473 | G2/M DNA damage checkpoint | H2AFX | H2A histone family member X | 3014 | down | 2,11 | 1,30E-03 |
| R-HSA-69473 | G2/M DNA damage checkpoint | RPA1 | replication protein A1 | 6117 | down | 1,41 | 4,93E-02 |
| R-HSA-216083 | Integrin cell surface interactions | ICAM3 | intercellular adhesion molecule 3 | 3385 | down | 1,36 | 1,58E-02 |
| R-HSA-216083 | Integrin cell surface interactions | DAG1 | dystroglycan 1 | 1605 | down | 1,65 | 1,04E-02 |
| R-HSA-912446 | Meiotic recombination | NBN | nibrin | 4683 | up | 1,25 | 3,15E-02 |
| R-HSA-912446 | Meiotic recombination | H2AFX | H2A histone family member X | 3014 | down | 2,11 | 1,30E-03 |
| R-HSA-912446 | Meiotic recombination | RPA1 | replication protein A1 | 6117 | down | 1,41 | 4,93E-02 |
| R-HSA-2408557 | Selenocysteine synthesis | EEFSEC | eukaryotic elongation factor, selenocyst | 60678 | down | 1,36 | 4,95E-02 |
| R-HSA-2408557 | Selenocysteine synthesis | SECISBP2 | SECIS binding protein 2 | 79048 | up | 1,25 | 3,39E-02 |
| R-HSA-2408557 | Selenocysteine synthesis | SARS | seryl-tRNA synthetase | 6301 | up | 1,82 | 1,31E-02 |
| R-HSA-2408557 | Selenocysteine synthesis | RPS4Y1 | ribosomal protein S4, Y-linked 1 | 6192 | down | 5,59 | 7,82E-03 |
| R-HSA-975956 | Nonsense Mediated Decay (NMD) independent of the | RPS4Y1 | ribosomal protein S4, Y-linked 1 | 6192 | down | 5,59 | 7,82E-03 |
| R-HSA-975956 | Nonsense Mediated Decay (NMD) independent of the | UPF1 | UPF1, RNA helicase and ATPase | 5976 | down | 1,27 | 2,46E-02 |
| R-HSA-5693607 | Processing of DNA double-strand break ends | H2AFX | H2A histone family member X | 3014 | down | 2,11 | 1,30E-03 |
| R-HSA-5693607 | Processing of DNA double-strand break ends | RPA1 | replication protein A1 | 6117 | down | 1,41 | 4,93E-02 |
| R-HSA-5693607 | Processing of DNA double-strand break ends | HUS1 | HUS1 checkpoint clamp component | 3364 | up | 1,36 | 4,96E-02 |
| R-HSA-5693607 | Processing of DNA double-strand break ends | NBN | nibrin | 4683 | up | 1,25 | 3,15E-02 |
| R-HSA-977225 | Amyloid fiber formation | H2AFX | H2A histone family member X | 3014 | down | 2,11 | 1,30E-03 |
| R-HSA-977225 | Amyloid fiber formation | GGA2 | golgi associated, gamma adaptin ear co | 23062 | down | 1,27 | 3,86E-02 |
| R-HSA-977225 | Amyloid fiber formation | SIAH1 | siah E3 ubiquitin protein ligase 1 | 6477 | up | 1,74 | 1,66E-02 |
| R-HSA-72689 | Formation of a pool of free 40S subunits | EIF3B | eukaryotic translation initiation factor 3 s | 8662 | down | 1,27 | 1,76E-02 |
| R-HSA-72689 | Formation of a pool of free 40S subunits | EIF3H | eukaryotic translation initiation factor 3 s | 8667 | up | 1,22 | 3,84E-02 |
| R-HSA-72689 | Formation of a pool of free 40S subunits | EIF3M | eukaryotic translation initiation factor 3 s | 10480 | up | 1,25 | 2,51E-02 |
| R-HSA-72689 | Formation of a pool of free 40S subunits | RPS4Y1 | ribosomal protein S4, Y-linked 1 | 6192 | down | 5,59 | 7,82E-03 |
| R-HSA-975957 | Nonsense Mediated Decay (NMD) enhanced by the E | RPS4Y1 | ribosomal protein S4, Y-linked 1 | 6192 | down | 5,59 | 7,82E-03 |
| R-HSA-975957 | Nonsense Mediated Decay (NMD) enhanced by the E | UPF1 | UPF1, RNA helicase and ATPase | 5976 | down | 1,27 | 2,46E-02 |
| R-HSA-975957 | Nonsense Mediated Decay (NMD) enhanced by the E | PPP2R2A | protein phosphatase 2 regulatory subun | 5520 | up | 1,26 | 3,61E-02 |
| R-HSA-2029482 | Regulation of actin dynamics for phagocytic cup form | CFL1 | cofilin 1 | 1072 | down | 1,29 | 3,02E-02 |
| R-HSA-2029482 | Regulation of actin dynamics for phagocytic cup form | LIMK1 | LIM domain kinase 1 | 3984 | down | 1,44 | 6,49E-03 |
| R-HSA-2029482 | Regulation of actin dynamics for phagocytic cup form | NCK1 | NCK adaptor protein 1 | 4690 | up | 1,48 | 1,42E-02 |
| R-HSA-2029482 | Regulation of actin dynamics for phagocytic cup form | ACTG1 | actin gamma 1 | 71 | up | 1,58 | 1,66E-02 |
| R-HSA-4641258 | Degradation of DVL | KLHL12 | kelch like family member 12 | 59349 | up | 1,31 | 4,38E-02 |
| R-HSA-4641258 | Degradation of DVL | DACT1 | dishevelled binding antagonist of beta c | 51339 | down | 5,29 | 2,23E-02 |
| R-HSA-5610783 | Degradation of GLI2 by the proteasome | SUFU | SUFU negative regulator of hedgehog s | 51684 | down | 1,41 | 3,37E-02 |
| R-HSA-5610783 | Degradation of GLI2 by the proteasome | CSNK1A1 | casein kinase 1 alpha 1 | 1452 | up | 1,35 | 2,84E-02 |
| R-HSA-5610785 | GLI3 is processed to GLI3R by the proteasome | CSNK1A1 | casein kinase 1 alpha 1 | 1452 | up | 1,35 | 2,84E-02 |
| R-HSA-5610785 | GLI3 is processed to GLI3R by the proteasome | SUFU | SUFU negative regulator of hedgehog s | 51684 | down | 1,41 | 3,37E-02 |
| R-HSA-5610780 | Degradation of GLI1 by the proteasome | SUFU | SUFU negative regulator of hedgehog s | 51684 | down | 1,41 | 3,37E-02 |
| R-HSA-5610780 | Degradation of GLI1 by the proteasome | ITCH | itchy E3 ubiquitin protein ligase | 83737 | up | 1,51 | 6,64E-03 |
| R-HSA-187577 | SCF(Skp2)-mediated degradation of p27/p21 | CDKN1A | cyclin dependent kinase inhibitor 1A | 1026 | up | 3,62 | 5,43E-03 |
| R-HSA-187577 | SCF(Skp2)-mediated degradation of p27/p21 | SKP2 | S-phase kinase associated protein 2 | 6502 | down | 2,21 | 3,69E-02 |
| R-HSA-1442490 | Collagen degradation | ADAM17 | ADAM metallopeptidase domain 17 | 6868 | up | 2,59 | 1,60E-03 |
| R-HSA-1442490 | Collagen degradation | MMP15 | matrix metallopeptidase 15 | 4324 | down | 1,49 | 4,35E-02 |
| R-HSA-2871837 | FCERI mediated NF-kB activation | BCL10 | B-cell CLL/lymphoma 10 | 8915 | up | 2,06 | 5,25E-04 |
| R-HSA-2871837 | FCERI mediated NF-kB activation | LYN | LYN proto-oncogene, Src family tyrosine | 4067 | down | 1,36 | 2,34E-02 |
| R-HSA-2871837 | FCERI mediated NF-kB activation | TAB1 | TGF-beta activated kinase 1 (MAP3K7) | 10454 | down | 1,29 | 1,94E-02 |
| R-HSA-2871837 | FCERI mediated NF-kB activation | TAB3 | TGF-beta activated kinase 1 and MAP3 | 257397 | up | 1,40 | 4,94E-02 |
| R-HSA-198933 | Immunoregulatory interactions between a Lymphoid a | PVR | poliovirus receptor | 5817 | up | 1,67 | 4,17E-02 |
| R-HSA-198933 | Immunoregulatory interactions between a Lymphoid a | ULBP1 | UL16 binding protein 1 | 80329 | up | 2,14 | 2,19E-02 |
| R-HSA-198933 | Immunoregulatory interactions between a Lymphoid a | OSCAR | osteoclast associated, immunoglobulin-l | 126014 | up | 4,60 | 2,10E-02 |
| R-HSA-5693571 | Nonhomologous End-Joining (NHEJ) | NBN | nibrin | 4683 | up | 1,25 | 3,15E-02 |
| R-HSA-5693571 | Nonhomologous End-Joining (NHEJ) | H2AFX | H2A histone family member X | 3014 | down | 2,11 | 1,30E-03 |
| R-HSA-2029481 | FCGR activation | YES1 | YES proto-oncogene 1, Src family tyrosi | 7525 | up | 1,30 | 2,08E-02 |
| R-HSA-2029481 | FCGR activation | LYN | LYN proto-oncogene, Src family tyrosine | 4067 | down | 1,36 | 2,34E-02 |
| R-HSA-2299718 | Condensation of Prophase Chromosomes | H2AFX | H2A histone family member X | 3014 | down | 2,11 | 1,30E-03 |
| R-HSA-2299718 | Condensation of Prophase Chromosomes | MCPH1 | microcephalin 1 | 79648 | up | 1,25 | 4,62E-02 |
| R-HSA-192823 | Viral mRNA Translation | DNAJC3 | DnaJ heat shock protein family (Hsp40) | 5611 | up | 2,75 | 2,24E-03 |
| R-HSA-192823 | Viral mRNA Translation | RPS4Y1 | ribosomal protein S4, Y-linked 1 | 6192 | down | 5,59 | 7,82E-03 |
