## Supplemental Table 7 for "Transcriptome profiling of human colonic cells exposed to the gut pathobiont *Streptococcus gallolyticus* subsp. *gallolyticus*"

**Supplemental Table S7.** GSEA: List of 14 gene sets enriched in FHC cells induced by *SGG* UCN34 infection.

| Gene Set | Number of Genes | Total number of genes | PValue | FDR | Direction | Genes names |
|---|---|---|---|---|---|---|
| HALLMARK_OXIDATIVE_PHOSPHORYLATION | 199 | 200 | 7,32E-11 | 3,66E-09 | Up | ABCB7;ACAA1;ACAA2;ACADM;ACADSB;ACADVL;ACAT1;ACO2;AFG3L2;AIFM1;ALAS1;ALDH6A1;ATP1B1;ATP5A1;ATP5B;ATP5C1;ATP5D;ATP5E;ATP5F1;ATP5G1;ATP5G2;ATP5G3;ATP5H;ATP5I;ATP5J;ATP5J2;ATP5L;ATP5O;ATP6AP1;ATP6V0B;ATP6V0E1;ATP6V1C1;ATP6V1D;ATP6V1E1;ATP6V1F;ATP6V1G1;ATP6V1H;BAX;BCKDHA;BDH2;CASP7;COX10;COX11;COX15;COX17;COX4I1;COX5A;COX5B;COX6A1;COX6B1;COX6C;COX7A2;COX7A2L;COX7B;COX7C;COX8A;CPT1A;CS;CYB5A;CYB5R3;CYC1;CYCS;DECR1;DLAT;DLD;DLST;ECH1;ECHS1;ECI1;ETFA;ETFB;ETFDH;FDX1;FH;FXN;GLUD1;GOT2;GPI;GPX4;GRPEL1;HADHA;HADHB;HCCS;HSD17B10;HSPA9;HTRA2;IDH1;IDH2;IDH3A;IDH3B;IDH3G;IMMT;ISCA1;ISCU;LDHA;LDHB;LRPPRC;MAOB;MDH1;MDH2;MFN2;MGST3;MPC1;MRPL11;MRPL15;MRPL34;MRPL35;MRPS11;MRPS12;MRPS15;MRPS22;MRPS30;MTRF1;MTRR;MTX2;NDUFA1;NDUFA2;NDUFA3;NDUFA4;NDUFA5;NDUFA6;NDUFA7;NDUFA8;NDUFA9;NDUFAB1;NDUFB1;NDUFB2;NDUFB3;NDUFB4;NDUFB5;NDUFB6;NDUFB7;NDUFB8;NDUFC1;NDUFC2;NDUFS1;NDUFS2;NDUFS3;NDUFS4;NDUFS6;NDUFS7;NDUFS8;NDUFV1;ND |
| HALLMARK_ADIPOGENESIS | 184 | 200 | 3,90E-05 | 0,000689113 | Up | ABCA1;ABCB8;ACAA2;ACADM;ACADS;ACLY;ACO2;ACOX1;ADCY6;ADIPOR2;AGPAT3;AIFM1;AK2;ALDH2;ALDOA;ANGPT1;ANGPTL4;APLP2;APOE;ARAF;ARL4A;ATL2;ATP1B3;ATP5O;BAZ2A;BCKDHA;BCL2L13;BCL6;C3;CAT;CAVIN1;CAVIN2;CCNG2;CD151;CD302;CD36;CDKN2C;CHCHD10;CHUK;CMBL;CMPK1;COL15A1;COL4A1;COQ3;COQ5;COQ9;COX6A1;COX7B;COX8A;CPT2;CRAT;CS;CYC1;DBT;DECR1;DGAT1;DHCR7;DHRS7;DHRS7B;DLAT;DLD;DNAJB9;DNAJC15;DRAM2;ECH1;ECHS1;ELMOD3;ELOVL6;ENPP2;EPHX2;ESRRA;ESYT1;ETFB;FAH;FZD4;G3BP2;GADD45A;GBE1;GHITM;GPAM;GPAT4;GPD2;GPHN;GPX3;GPX4;GRPEL1;HADH;HIBCH;HSPB8;IDH1;IDH3A;IDH3G;IFNGR1;IMMT;ITGA7;ITSN1;JAGN1;LAMA4;LIFR;LIPE;LPCAT3;LPL;MAP4K3;MARC2;MCCC1;MDH2;ME1;MGLL;MGST3;MIGA2;MRPL15;MTCH2;MYLK;NABP1;NDUFA5;NDUFAB1;NDUFB7;NDUFS3;NKIRAS1;NMT1;PDCD4;PEMT;PEX14;PFKFB3;PFKL;PGM1;PHLDB1;PHYH;PIM3;PLIN2;POR;PPARG;PPM1B;PPP1R15B;PQLC3;PRDX3;PREB;PTCD3;QDPR;RAB34;REEP5;REEP6;RETSAT;RIOK3;RMDN3;RNF11;RREB1;RTN3;SAMM50;SCARB1;SCP2;SDHB;SDHC;SLC19A1;SLC1A5;SLC25 |
| HALLMARK_REACTIVE_OXIGEN_SPECIES_PATHWAY | 46 | 49 | 5,47E-05 | 0,000689113 | Up | ABCC1;ATOX1;CAT;CDKN2D;EGLN2;ERCC2;FES;FTL;G6PD;GCLC;GCLM;GLRX;GLRX2;GPX3;GPX4;GSR;HHEX;HMOX2;JUNB;LAMTOR5;MBP;MGST1;MSRA;NDUFA6;NDUFB4;NDUFS2;NQO1;OXSR1;PDLIM1;PFKP;PRDX1;PRDX2;PRDX4;PRDX6;PRNP;PTPA;SBNO2;SCAF4;SELENOS;SOD1;SOD2;SRXN1;STK25;TXN;TXNRD1;TXNRD2 |
| HALLMARK_CHOLESTEROL_HOMEOSTASIS | 72 | 74 | 5,52E-05 | 0,000689113 | Down | ABCA2;ACAT2;ACSS2;ACTG1;ALCAM;ALDOC;ANTXR2;ANXA13;ANXA5;ATF3;ATF5;ATXN2;CBS;CD9;CHKA;CLU;CPEB2;CTNNB1;CXCL16;CYP51A1;DHCR7;EBP;ECH1;ERRFI1;ETHE1;FABP5;FADS2;FAM129A;FASN;FBXO6;FDFT1;FDPS;GLDC;GNAI1;GPX8;GSTM2;GUSB;HMGCR;HMGCS1;HSD17B7;IDI1;JAG1;LDLR;LGALS3;LGMN;LPL;LSS;MAL2;MVD;MVK;NFIL3;NSDHL;PCYT2;PDK3;PLAUR;PLSCR1;PMVK;PNRC1;PPARG;S100A11;SC5D;SCD;SEMA3B;SQLE;SREBF2;STARD4;STX5;TM7SF2;TMEM97;TNFRSF12A;TP53INP1;TRIB3 |
| HALLMARK_INTERFERON_ALPHA_RESPONSE | 91 | 97 | 6,89E-05 | 0,000689113 | Up | ADAR;B2M;BATF2;BST2;C1S;CASP1;CASP8;CCRL2;CD47;CMPK2;CMTR1;CNP;CSF1;CXCL10;DDX60;DHX58;EIF2AK2;ELF1;EPSTI1;FAM46A;GBP2;GBP4;GMPR;HELZ2;HERC6;HLA-C;IFI27;IFI35;IFI44;IFIH1;IFIT2;IFIT3;IFITM1;IFITM2;IFITM3;IL15;IL4R;IRF1;IRF2;IRF7;ISG15;ISG20;LAMP3;LAP3;LGALS3BP;LPAR6;LY6E;MOV10;MVB12A;MX1;NCOA7;NMI;NUB1;OAS1;OASL;OGFR;PARP12;PARP14;PARP9;PLSCR1;PNPT1;PROCR;PSMA3;PSMB8;PSMB9;PSME1;PSME2;RIPK2;RNF31;RSAD2;RTP4;SAMD9;SAMD9L;SELL;SLC25A28;SP110;STAT2;TAP1;TDRD7;TMEM140;TRAFD1;TRIM14;TRIM21;TRIM25;TRIM26;TRIM5;TXNIP;UBA7;UBE2L6;USP18;WARS |
| HALLMARK_MYC_TARGETS_V1 | 199 | 200 | 0,00015197 | 0,001266389 | Up | ABCE1;ACP1;AIMP2;AP3S1;APEX1;BUB3;C1QBP;CAD;CANX;CBX3;CCNA2;CCT2;CCT3;CCT4;CCT5;CCT7;CDC20;CDC45;CDK2;CDK4;CLNS1A;CNBP;COPS5;COX5A;CSTF2;CTPS1;CUL1;CYC1;DDX18;DDX21;DEK;DHX15;DUT;EEF1B2;EIF1AX;EIF2S1;EIF2S2;EIF3B;EIF3D;EIF3J;EIF4A1;EIF4E;EIF4G2;EIF4H;EPRS;ERH;ETF1;EXOSC7;FAM120A;FBL;G3BP1;GLO1;GNL3;GOT2;GSPT1;H2AFZ;HDAC2;HDDC2;HDGF;HNRNPA1;HNRNPA2B1;HNRNPA3;HNRNPC;HNRNPD;HNRNPR;HNRNPU;HPRT1;HSP90AB1;HSPD1;HSPE1;IARS;IFRD1;ILF2;IMPDH2;KARS;KPNA2;KPNB1;LDHA;LSM2;LSM7;MAD2L1;MCM2;MCM4;MCM5;MCM6;MCM7;MRPL23;MRPL9;MRPS18B;MYC;NAP1L1;NCBP1;NCBP2;NDUFAB1;NHP2;NME1;NOLC1;NOP16;NOP56;NPM1;ODC1;ORC2;PA2G4;PABPC1;PABPC4;PCBP1;PCNA;PGK1;PHB;PHB2;POLD2;POLE3;PPIA;PPM1G;PRDX3;PRDX4;PRPF31;PRPS2;PSMA1;PSMA2;PSMA4;PSMA6;PSMA7;PSMB2;PSMB3;PSMC4;PSMC6;PSMD1;PSMD14;PSMD3;PSMD7;PSMD8;PTGES3;PWP1;RACK1;RAD23B;RAN;RANBP1;RFC4;RNPS1;RPL14;RPL18;RPL22;RPL34;RPL6;RPLP0;RPS10;RPS2;RPS3;RPS5;RPS6;RRM1;RRP9;RSL1D1;RUVBL2;SERBP1;SET;S |
| HALLMARK_PROTEIN_SECRETION | 95 | 96 | 0,00023553 | 0,001617622 | Up | ABCA1;ADAM10;ANP32E;AP1G1;AP2B1;AP2M1;AP2S1;AP3B1;AP3S1;ARCN1;ARF1;ARFGAP3;ARFGEF1;ARFGEF2;ARFIP1;ATP1A1;ATP6V1B1;ATP6V1H;ATP7A;BET1;BNIP3;CAV2;CD63;CLCN3;CLN5;CLTA;CLTC;COG2;COPB1;COPB2;COPE;CTSC;DNM1L;DOPEY1;DST;EGFR;ERGIC3;GALC;GBF1;GLA;GNAS;GOLGA4;GOSR2;ICA1;IGF2R;KIF1B;KRT18;LAMP2;LMAN1;M6PR;MAPK1;MON2;NAPA;NAPG;OCRL;PAM;PPT1;RAB14;RAB22A;RAB2A;RAB5A;RAB9A;RER1;RPS6KA3;SCAMP1;SCAMP3;SCRN1;SEC22B;SEC24D;SEC31A;SGMS1;SNAP23;SNX2;SOD1;SSPN;STAM;STX12;STX16;STX7;TMED10;TMED2;TMX1;TOM1L1;TPD52;TSG101;TSPAN8;USO1;VAMP3;VAMP4;VAMP7;VPS45;VPS4B;YIPF6;YKT6;ZW10 |
| HALLMARK_FATTY_ACID_METABOLISM | 136 | 158 | 0,00025882 | 0,001617622 | Up | AADAT;ACAA1;ACAA2;ACADM;ACADS;ACADVL;ACAT2;ACO2;ACOT2;ACOT8;ACOX1;ACSL1;ACSL4;ACSL5;ACSM3;ACSS1;ADIPOR2;ADSL;ALAD;ALDH1A1;ALDH3A1;ALDH3A2;ALDH9A1;ALDOA;APEX1;AUH;BCKDHB;BLVRA;BPHL;CA2;CBR1;CBR3;CCDC58;CD36;CEL;CPOX;CPT1A;CPT2;CRAT;CRYZ;CYP1A1;D2HGDH;DECR1;DHCR24;DLD;DLST;ECH1;ECHS1;ECI1;ECI2;EHHADH;ELOVL5;ENO2;ENO3;EPHX1;ERP29;ETFDH;FABP1;FASN;FH;G0S2;GABARAPL1;GCDH;GLUL;GPD1;GPD2;GRHPR;GSTZ1;H2AFZ;HADH;HADHB;HCCS;HIBCH;HMGCL;HMGCS1;HMGCS2;HPGD;HSD17B10;HSD17B11;HSD17B4;HSD17B7;HSDL2;HSP90AA1;HSPH1;IDH1;IDH3B;IDH3G;IDI1;INMT;KMT5A;LDHA;LGALS1;MAOA;MCEE;MDH1;MDH2;ME1;METAP1;MGLL;MIF;MLYCD;NBN;NCAPH2;NSDHL;NTHL1;ODC1;OSTC;PCBD1;PDHA1;PDHB;PPARA;PRDX6;PSME1;PTPRG;PTS;RAP1GDS1;RDH11;REEP6;RETSAT;S100A10;SDHA;SDHC;SDHD;SERINC1;SLC22A5;SMS;SUCLA2;SUCLG1;SUCLG2;TP53INP2;UBE2L6;UGDH;UROD;UROS;XIST;YWHAH |
| HALLMARK_EPITHELIAL_MESENCHYMAL_TRANSITION | 183 | 200 | 0,00049137 | 0,002729811 | Up | ACTA2;ADAM12;ANPEP;APLP1;AREG;BASP1;BDNF;BGN;BMP1;CADM1;CALD1;CALU;CAP2;CAPG;CD44;CD59;CDH11;CDH2;CDH6;COL11A1;COL12A1;COL16A1;COL1A1;COL1A2;COL3A1;COL4A1;COL4A2;COL5A1;COL5A2;COL6A2;COL6A3;COL7A1;COL8A2;COLGALT1;COPA;CRLF1;CTGF;CTHRC1;CXCL1;CXCL8;CYR61;DAB2;DCN;DKK1;DPYSL3;DST;ECM1;EDIL3;EFEMP2;ELN;EMP3;ENO2;FAP;FAS;FBLN1;FBLN2;FBLN5;FBN1;FBN2;FERMT2;FGF2;FLNA;FMOD;FN1;FOXC2;FSTL1;FSTL3;FUCA1;FZD8;GADD45A;GADD45B;GAS1;GEM;GJA1;GLIPR1;GPC1;GPX7;GREM1;HTRA1;ID2;IGFBP2;IGFBP3;IGFBP4;IL15;IL32;IL6;INHBA;ITGA2;ITGA5;ITGAV;ITGB1;ITGB3;ITGB5;JUN;LAMA1;LAMA2;LAMA3;LAMC1;LAMC2;LGALS1;LOX;LOXL1;LOXL2;LRP1;LRRC15;LUM;MATN2;MATN3;MCM7;MEST;MMP1;MMP14;MMP2;MMP3;MSX1;MXRA5;MYL9;MYLK;NID2;NNMT;NOTCH2;NT5E;NTM;OXTR;P3H1;PCOLCE;PCOLCE2;PDGFRB;PDLIM4;PFN2;PLAUR;PLOD1;PLOD2;PLOD3;PMEPA1;PMP22;PPIB;PRSS2;PTHLH;PTX3;PVR;QSOX1;RHOB;SAT1;SCG2;SDC1;SDC4;SERPINE1;SERPINE2;SERPINH1;SFRP1;SGCB;SGCG;SLC6A8;SNAI2;SNTB1;SPARC;SPOCK1;SPP1;T |
| HALLMARK_INTERFERON_GAMMA_RESPONSE | 164 | 200 | 0,00127627 | 0,006381333 | Up | ADAR;APOL6;ARID5B;ARL4A;AUTS2;B2M;BANK1;BATF2;BPGM;BST2;BTG1;C1R;C1S;CASP1;CASP3;CASP4;CASP7;CASP8;CD274;CD40;CD69;CDKN1A;CFH;CMPK2;CMTR1;CSF2RB;CXCL10;DDX58;DDX60;DHX58;EIF2AK2;EIF4E3;EPSTI1;FAS;FPR1;GBP4;GCH1;HELZ2;HERC6;HIF1A;HLA-A;HLA-B;HLA-DMA;HLA-DQA1;HLA-G;ICAM1;IFI27;IFI35;IFI44;IFIH1;IFIT1;IFIT2;IFIT3;IFITM2;IFITM3;IFNAR2;IL15;IL15RA;IL18BP;IL4R;IL6;IRF1;IRF2;IRF5;IRF7;IRF8;ISG15;ISG20;ISOC1;ITGB7;JAK2;LAP3;LATS2;LGALS3BP;LY6E;LYSMD2;METTL7B;MT2A;MTHFD2;MVP;MX1;MX2;MYD88;NAMPT;NCOA3;NFKB1;NFKBIA;NLRC5;NMI;NOD1;NUP93;OAS2;OAS3;OASL;OGFR;PARP12;PARP14;PDE4B;PELI1;PFKP;PIM1;PLA2G4A;PLSCR1;PML;PNP;PNPT1;PSMA2;PSMA3;PSMB10;PSMB2;PSMB8;PSMB9;PSME1;PSME2;PTGS2;PTPN1;PTPN2;PTPN6;RAPGEF6;RBCK1;RIPK1;RIPK2;RNF213;RNF31;RSAD2;RTP4;SAMD9L;SAMHD1;SECTM1;SERPING1;SLC25A28;SOCS1;SOCS3;SOD2;SP110;SPPL2A;SRI;SSPN;ST3GAL5;STAT1;STAT2;STAT3;STAT4;TAP1;TAPBP;TDRD7;TNFAIP2;TNFAIP3;TNFSF10;TOR1B;TRAFD1;TRIM14;TRIM21;TRIM25;TRIM26;TXNIP;UBE2L6;UPP1; |
| HALLMARK_MYC_TARGETS_V2 | 58 | 58 | 0,003059 | 0,013904561 | Down | AIMP2;BYSL;CBX3;CDK4;DCTPP1;DDX18;DUSP2;EXOSC5;FARSA;GNL3;GRWD1;HK2;HSPD1;HSPE1;IMP4;IPO4;LAS1L;MAP3K6;MCM4;MCM5;MPHOSPH10;MRTO4;MYBBP1A;MYC;NDUFAF4;NIP7;NOC4L;NOLC1;NOP16;NOP2;NOP56;NPM1;PA2G4;PES1;PHB;PLK1;PLK4;PPAN;PPRC1;PRMT3;PUS1;RABEPK;RCL1;RRP12;RRP9;SLC19A1;SLC29A2;SORD;SRM;SUPV3L1;TBRG4;TCOF1;TFB2M;TMEM97;UNG;UTP20;WDR43;WDR74 |
| HALLMARK_HYPOXIA | 177 | 200 | 0,00579276 | 0,024136484 | Up | ACKR3;ADM;ADORA2B;AK4;AKAP12;ALDOA;ALDOC;AMPD3;ANGPTL4;ANKZF1;ANXA2;ATF3;ATP7A;B3GALT6;B4GALNT2;BCL2;BGN;BHLHE40;BNIP3L;BTG1;CA12;CASP6;CAV1;CAVIN1;CAVIN3;CCNG2;CDKN1A;CDKN1B;CDKN1C;CHST2;CHST3;CITED2;COL5A1;CSRP2;CTGF;CXCR4;CYR61;DCN;DDIT3;DDIT4;DPYSL4;DTNA;DUSP1;EFNA1;EFNA3;EGFR;ENO1;ENO2;ENO3;ERO1A;ERRFI1;ETS1;EXT1;F3;FAM162A;FBP1;FOS;FOSL2;FOXO3;GAA;GALK1;GAPDH;GBE1;GCNT2;GLRX;GPC1;GPC3;GPC4;GPI;GRHPR;GYS1;HDLBP;HEXA;HK1;HK2;HMOX1;HOXB9;HS3ST1;HSPA5;IDS;IER3;IGFBP1;IGFBP3;IL6;ILVBL;IRS2;ISG20;JMJD6;JUN;KDELR3;KDM3A;KLF6;KLF7;KLHL24;LARGE1;LDHA;LOX;LXN;MAFF;MAP3K1;MIF;MT1E;MT2A;MXI1;MYH9;NAGK;NDRG1;NDST1;NDST2;NEDD4L;NFIL3;NOCT;NR3C1;P4HA1;P4HA2;PAM;PDGFB;PDK1;PDK3;PFKFB3;PFKL;PFKP;PGF;PGK1;PGM1;PGM2;PIM1;PKP1;PLAC8;PLAUR;PLIN2;PNRC1;PPARGC1A;PPP1R15A;PPP1R3C;PRDX5;PRKCA;RBPJ;RORA;RRAGD;S100A4;SAP30;SCARB1;SDC2;SDC3;SDC4;SELENBP1;SERPINE1;SIAH2;SLC25A1;SLC2A1;SLC2A3;SLC37A4;SLC6A6;SRPX;STBD1;STC1;STC2;SULT2B1;TES;TGF |
| HALLMARK_ESTROGEN_RESPONSE_EARLY | 185 | 200 | 0,00758651 | 0,0291789 | Down | ABAT;ABCA3;ABHD2;ABLIM1;ADCY1;ADCY9;ADD3;AFF1;AKAP1;ALDH3B1;AMFR;ANXA9;AQP3;AR;AREG;ARL3;ASB13;B4GALT1;BAG1;BCL11B;BCL2;BHLHE40;BLVRB;CA12;CALB2;CANT1;CCND1;CD44;CELSR1;CELSR2;CHPT1;CISH;CLDN7;CLIC3;CYP26B1;DEPTOR;DHCR7;DHRS2;DHRS3;DLC1;DYNLT3;EGR3;ELF1;ELF3;ELOVL5;ENDOD1;ESRP2;FAM102A;FARP1;FASN;FCMR;FDFT1;FHL2;FKBP4;FKBP5;FLNB;FOS;FOXC1;FRK;GAB2;GFRA1;GJA1;GLA;GREB1;HES1;HR;HSPB8;IGF1R;IGFBP4;IL17RB;IL6ST;INHBB;INPP5F;ISG20L2;ITPK1;JAK2;KAZN;KCNK15;KCNK5;KDM4B;KLF10;KLF4;KLK10;KRT13;KRT15;KRT18;KRT19;KRT8;LAD1;LRIG1;MAPT;MAST4;MED13L;MED24;MICB;MINDY1;MLPH;MREG;MUC1;MYB;MYBBP1A;MYBL1;MYC;MYOF;NADSYN1;NAV2;NBL1;NCOR2;NRIP1;NXT1;OLFM1;OLFML3;OPN3;OVOL2;P2RY2;PAPSS2;PDZK1;PEX11A;PLA2G16;PMAIP1;PODXL;PPIF;PRSS23;PTGES;RAB17;RAB31;RAPGEFL1;RARA;RASGRP1;RBBP8;RET;RETREG1;RHOBTB3;RHOD;RPS6KA2;RRP12;SCARB1;SCNN1A;SEC14L2;SEMA3B;SFN;SH3BP5;SIAH2;SLC16A1;SLC19A2;SLC1A1;SLC1A4;SLC22A5;SLC26A2;SLC27A2;SLC2A1;SLC37A1;SLC39A6;SL |
| HALLMARK_DNA_REPAIR | 148 | 150 | 0,01211822 | 0,043279343 | Up | AAAS;ADA;ADCY6;ADRM1;AGO4;AK1;AK3;ALYREF;APRT;ARL6IP1;BCAM;BCAP31;BRF2;CANT1;CCNO;CDA;CETN2;CLP1;CMPK2;COX17;CSTF3;DAD1;DCTN4;DDB1;DDB2;DGCR8;DGUOK;DUT;EDF1;EIF1B;ELL;ELOA;ERCC1;ERCC2;ERCC3;ERCC4;ERCC5;ERCC8;FEN1;GMPR2;GPX4;GSDME;GTF2A2;GTF2B;GTF2F1;GTF2H1;GTF2H3;GTF2H5;GTF3C5;GUK1;HCLS1;HPRT1;IMPDH2;ITPA;LIG1;MPC2;MPG;MRPL40;NCBP2;NELFB;NELFCD;NELFE;NFX1;NME1;NME3;NME4;NPR2;NT5C;NT5C3A;NUDT21;NUDT9;PCNA;PDE4B;PDE6G;PNP;POLA1;POLA2;POLB;POLD1;POLD3;POLD4;POLE4;POLH;POLL;POLR1C;POLR1D;POLR2C;POLR2D;POLR2E;POLR2F;POLR2G;POLR2H;POLR2I;POLR2J;POLR2K;POLR3C;POLR3GL;POM121;PRIM1;RAD51;RAD52;RAE1;RALA;RBX1;REV3L;RFC2;RFC3;RFC4;RFC5;RNMT;RPA2;RPA3;RRM2B;SAC3D1;SDCBP;SEC61A1;SF3A3;SMAD5;SNAPC4;SNAPC5;SRSF6;SSRP1;STX3;SUPT4H1;SUPT5H;SURF1;TAF10;TAF12;TAF13;TAF1C;TAF6;TAF9;TARBP2;TK2;TMED2;TP53;TSG101;TYMS;UMPS;UPF3B;USP11;VPS28;VPS37B;VPS37D;XPC;ZNF707;ZNRD1;ZWINT |
