## Supplemental Table 8 for "Transcriptome profiling of human colonic cells exposed to the gut pathobiont *Streptococcus gallolyticus* subsp. *gallolyticus*"

**Supplemental Table S8. GSEA:** List of 9 gene sets enriched in HT29 cells induced by *SGG* UCN34 infection.

| Gene Set | Number of Genes | Total number of genes | PValue | FDR | Direction | Genes names |
|---|---|---|---|---|---|---|
| HALLMARK_UNFOLDED_PROTEIN_RESPONSE | 111 | 113 | Up | 1,17E-09 | 5,85E-08 | ALDH18A1;ARFGAP1;ASNS;ATF3;ATF4;ATF6;ATP6V0D1;BAG3;BANF1;CALR;CEBPB;CEBPG;CHAC1;CKS1B;CNOT2;CNOT4;CNOT6;CXXC1;DCP1A;DCP2;DCTN1;DDIT4;DDX10;DKC1;DNAJA4;DNAJB9;DNAJC3;EDC4;EDEM1;EEF2;EIF2AK3;EIF2S1;EIF4A1;EIF4A2;EIF4A3;EIF4E;EIF4EBP1;EIF4G1;ERN1;ERO1A;EXOC2;EXOSC1;EXOSC10;EXOSC2;EXOSC4;EXOSC5;EXOSC9;FKBP14;FUS;GEMIN4;GOSR2;H2AFX;HERPUD1;HSP90B1;HSPA5;HSPA9;HYOU1;IARS;IFIT1;IMP3;KDELR3;KHSRP;KIF5B;LSM1;LSM4;MTHFD2;NABP1;NFYA;NFYB;NHP2;NOLC1;NOP14;NOP56;NPM1;PAIP1;PARN;PDIA5;PDIA6;POP4;PREB;PSAT1;RPS14;RRP9;SDAD1;SEC11A;SEC31A;SERP1;SHC1;SKIV2L2;SLC1A4;SLC30A5;SLC7A5;SPCS1;SPCS3;SRPRA;SRPRB;SSR1;STC2;TARS;TATDN2;TSPYL2;TTC37;TUBB2A;VEGFA;WFS1;WIPI1;XBP1;XPOT;YIF1A;YWHAZ;ZBTB17 |
| HALLMARK_E2F_TARGETS | 199 | 200 | Down | 3,74E-08 | 9,35E-07 | AK2;ANP32E;ASF1A;ASF1B;ATAD2;AURKA;AURKB;BARD1;BIRC5;BRCA1;BRCA2;BRMS1L;BUB1B;CBX5;CCNB2;CCNE1;CCP110;CDC20;CDC25A;CDC25B;CDCA3;CDCA8;CDK1;CDK4;CDKN1A;CDKN1B;CDKN2A;CDKN2C;CDKN3;CENPE;CENPM;CHEK1;CHEK2;CIT;CKS1B;CKS2;CNOT9;CSE1L;CTCF;CTPS1;DCK;DCLRE1B;DCTPP1;DDX39A;DEK;DEPDC1;DIAPH3;DLGAP5;DNMT1;DONSON;DSCC1;DUT;E2F8;EED;EIF2S1;ESPL1;EXOSC8;EZH2;GINS1;GINS3;GINS4;GSPT1;H2AFX;H2AFZ;HELLS;HMGA1;HMGB2;HMGB3;HMMR;HNRNPD;HUS1;ILF3;ING3;IPO7;JPT1;KIF18B;KIF22;KIF2C;KIF4A;KPNA2;LBR;LIG1;LMNB1;LSM8;LUC7L3;LYAR;MAD2L1;MCM2;MCM3;MCM4;MCM5;MCM6;MCM7;MELK;MKI67;MLH1;MMS22L;MRE11;MSH2;MTHFD2;MXD3;MYBL2;MYC;NAP1L1;NASP;NBN;NCAPD2;NME1;NOLC1;NOP56;NUDT21;NUP107;NUP153;NUP205;ORC2;ORC6;PA2G4;PAICS;PAN2;PCNA;PDS5B [illegible] |
| HALLMARK_PROTEIN_SECRETION | 94 | 96 | Up | 1,46E-07 | 2,44E-06 | ABCA1;ADAM10;ANP32E;AP1G1;AP2B1;AP2M1;AP2S1;AP3B1;AP3S1;ARCN1;ARF1;ARFGAP3;ARFGEF1;ARFGEF2;ARFIP1;ATP1A1;ATP6V1H;ATP7A;BET1;BNIP3;CAV2;CD63;CLCN3;CLN5;CLTA;CLTC;COG2;COPB1;COPB2;COPE;CTSC;DNM1L;DOPEY1;DST;EGFR;ERGIC3;GALC;GBF1;GLA;GNAS;GOLGA4;GOSR2;ICA1;IGF2R;KIF1B;KRT18;LAMP2;LMAN1;M6PR;MAPK1;MON2;NAPA;NAPG;OCRL;PAM;PPT1;RAB14;RAB22A;RAB2A;RAB5A;RAB9A;RER1;RPS6KA3;SCAMP1;SCAMP3;SCRN1;SEC22B;SEC24D;SEC31A;SGMS1;SNAP23;SNX2;SOD1;SSPN;STAM;STX12;STX16;STX7;TMED10;TMED2;TMX1;TOM1L1;TPD52;TSG101;TSPAN8;USO1;VAMP3;VAMP4;VAMP7;VPS45;VPS4B;YIPF6;YKT6;ZW10 |
| HALLMARK_TNFA_SIGNALING_VIA_NFKB | 178 | 200 | Up | 2,70E-06 | 3,37E-05 | ABCA1;ACKR3;AREG;ATF3;ATP2B1;B4GALT1;B4GALT5;BCL3;BCL6;BHLHE40;BIRC2;BIRC3;BMP2;BTG1;BTG2;BTG3;CCND1;CCNL1;CCRL2;CD44;CD69;CD83;CDKN1A;CEBPB;CEBPD;CFLAR;CLCF1;CSF1;CSF2;CXCL1;CXCL10;CYR61;DDX58;DENND5A;DNAJB4;DRAM1;DUSP1;DUSP2;DUSP4;DUSP5;EDN1;EFNA1;EGR1;EGR2;EHD1;EIF1;ETS2;F2RL1;F3;FJX1;FOS;FOSB;FOSL1;FOSL2;FUT4;G0S2;GADD45A;GADD45B;GCH1;GEM;GFPT2;HBEGF;HES1;ICAM1;ID2;IER2;IER3;IER5;IFIH1;IFIT2;IFNGR2;IL15RA;IL18;IL1A;IL23A;IL6;IL6ST;IL7R;INHBA;IRF1;IRS2;JAG1;JUN;JUNB;KDM6B;KLF10;KLF2;KLF4;KLF6;KLF9;KYNU;LAMB3;LDLR;LIF;LITAF;MAFF;MAP2K3;MAP3K8;MARCKS;MCL1;MSC;MXD1;MYC;NAMPT;NFAT5;NFE2L2;NFIL3;NFKB1;NFKB2;NFKBIA;NFKBIE;NINJ1;NR4A1;NR4A2;NR4A3;OLR1;PANX1;PDE4B;PDLIM5;PER1;PFKFB3;PHLDA1;PHLDA2;PLAU;PLAUR;PLK2;PLPP3;PMEPA1;[illegible] |
| HALLMARK_G2M_CHECKPOINT | 198 | 200 | Down | 5,54E-06 | 5,54E-05 | ABL1;AMD1;ARID4A;ATF5;ATRX;AURKA;AURKB;BARD1;BCL3;BIRC5;BRCA2;BUB1;BUB3;CASP8AP2;CBX1;CCNA2;CCNB2;CCND1;CCNF;CCNT1;CDC20;CDC25A;CDC25B;CDC27;CDC45;CDC6;CDC7;CDK1;CDK4;CDKN1B;CDKN2C;CDKN3;CENPA;CENPE;CENPF;CHAF1A;CHEK1;CHMP1A;CKS1B;CKS2;CTCF;CUL1;CUL3;CUL4A;CUL5;DBF4;DDX39A;DKC1;DMD;DR1;DTYMK;E2F1;E2F2;E2F3;E2F4;EFNA5;EGF;ESPL1;EWSR1;EXO1;EZH2;FANCC;FBXO5;FOXN3;G3BP1;GINS2;GSPT1;H2AFV;H2AFX;H2AFZ;HIF1A;HIRA;HIST1H2BK;HMGA1;HMGB3;HMGN2;HMMR;HNRNPD;HNRNPU;HSPA8;HUS1;ILF3;INCENP;JPT1;KATNA1;KIF11;KIF15;KIF20B;KIF22;KIF23;KIF2C;KIF4A;KIF5B;KMT5A;KNL1;KPNA2;KPNB1;LBR;LIG3;LMNB1;MAD2L1;MAP3K20;MAPK14;MARCKS;MCM2;MCM3;MCM5;MCM6;MEIS1;MEIS2;MKI67;MNAT1;MT2A;MTF2;MYBL2;MYC;NASP;NCL;NDC80;NEK2;NOLC1;NOTCH2;NSD2;[illegible] |
| HALLMARK_KRAS_SIGNALING_DN | 90 | 200 | Down | 0,00216882 | 0,01807351 | AKR1B10;ASB7;BARD1;BTG2;C5;CAMK1D;CAPN9;CCDC106;CDKAL1;CELSR2;CHST2;CLSTN3;COPZ2;COQ8A;CPEB3;CYP39A1;DTNB;EDN1;EFHD1;EGF;ENTPD7;EPHA5;FAM46C;FGFR3;FGGY;GAMT;GDNF;GPR19;GPR3;GPRC5C;GTF3C5;H2AFY2;HNF1A;HSD11B2;HTR1D;IDUA;IGFBP2;ITIH3;KCND1;KMT2D;KRT13;KRT15;LFNG;LYPD3;MAGIX;MAST3;MFSD6;MSH5;MTHFR;MX1;NR4A2;NR6A1;NTF3;NUDT11;P2RX6;PDK2;PKP1;PLAG1;PRODH;PTPRJ;RGS11;RIBC2;RSAD2;SELENOP;SGK1;SIDT1;SKIL;SLC16A7;SLC25A23;SLC29A3;SNCB;SNN;SPHK2;SPRR3;SPTBN2;SYNPO;TCF7L1;TFCP2L1;TG;TGFB2;TGM1;THNSL2;THRB;TNNI3;VPS50;WNT16;YBX2;YPEL1;ZC2HC1C;ZNF112 |
| HALLMARK_MYC_TARGETS_V2 | 58 | 58 | Down | 0,00396268 | 0,02830485 | AIMP2;BYSL;CBX3;CDK4;DCTPP1;DDX18;DUSP2;EXOSC5;FARSA;GNL3;GRWD1;HK2;HSPD1;HSPE1;IMP4;IPO4;LAS1L;MAP3K6;MCM4;MCM5;MPHOSPH10;MRTO4;MYBBP1A;MYC;NDUFAF4;NIP7;NOC4L;NOLC1;NOP16;NOP2;NOP56;NPM1;PA2G4;PES1;PHB;PLK1;PLK4;PPAN;PPRC1;PRMT3;PUS1;RABEPK;RCL1;RRP12;RRP9;SLC19A1;SLC29A2;SORD;SRM;SUPV3L1;TBRG4;TCOF1;TFB2M;TMEM97;UNG;UTP20;WDR43;WDR74 |
| HALLMARK_EPITHELIAL_MESENCHYMAL_TRANSITION | 181 | 200 | Down | 0,00499892 | 0,03124324 | ACTA2;ANPEP;APLP1;AREG;BASP1;BDNF;BGN;BMP1;CADM1;CALD1;CALU;CAP2;CAPG;CD44;CD59;CDH11;CDH2;CDH6;COL11A1;COL12A1;COL16A1;COL1A1;COL1A2;COL3A1;COL4A1;COL4A2;COL5A1;COL5A2;COL6A2;COL6A3;COL7A1;COL8A2;COLGALT1;COPA;CRLF1;CTGF;CTHRC1;CXCL1;CXCL8;CYR61;DAB2;DCN;DKK1;DPYSL3;DST;ECM1;EDIL3;EFEMP2;EMP3;ENO2;FAP;FAS;FBLN1;FBLN2;FBLN5;FBN1;FBN2;FERMT2;FGF2;FLNA;FMOD;FN1;FOXC2;FSTL1;FSTL3;FUCA1;FZD8;GADD45A;GADD45B;GAS1;GEM;GJA1;GLIPR1;GPC1;GPX7;GREM1;HTRA1;ID2;IGFBP2;IGFBP3;IGFBP4;IL15;IL32;IL6;INHBA;ITGA2;ITGA5;ITGAV;ITGB1;ITGB3;ITGB5;JUN;LAMA1;LAMA2;LAMA3;LAMC1;LAMC2;LGALS1;LOX;LOXL1;LOXL2;LRP1;LRRC15;LUM;MATN2;MATN3;MCM7;MEST;MMP1;MMP14;MMP2;MMP3;MSX1;MXRA5;MYL9;MYLK;NID2;NNMT;NOTCH2;NT5E;NTM;OXTR;P3H1;PCOLCE;[illegible] |
| HALLMARK_HEME_METABOLISM | 168 | 200 | Up | 0,00831155 | 0,04617528 | ABCB6;ABCG2;ACP5;ADD1;ADD2;ADIPOR1;AGPAT4;ALAD;ALDH1L1;ALDH6A1;ANK1;AQP3;ARHGEF12;ARL2BP;ASNS;ATG4A;ATP6V0A1;BACH1;BCAM;BLVRA;BLVRB;BMP2K;BNIP3L;BPGM;BSG;BTG2;BTRC;C3;CA2;CAST;CAT;CCDC28A;CCND3;CDC27;CDR2;CIR1;CLCN3;CLIC2;CPOX;CROCCP2;CTNS;CTSB;CTSE;DAAM1;DCAF10;DCAF11;DCUN1D1;DMTN;E2F2;EIF2AK1;ELL2;ENDOD1;EPB41;EPOR;ERMAP;EZH1;FAM46C;FBXO34;FBXO7;FBXO9;FECH;FN3K;FOXJ2;FOXO3;GAPVD1;GCLC;GCLM;GDE1;GLRX5;GMPS;GYPC;H1F0;HAGH;HDGF;HEBP1;HMBS;HTATIP2;HTRA2;ICAM4;IGSF3;ISCA1;KAT2B;KDM7A;KHNYN;KLF3;LAMP2;LMO2;LPIN2;LRP10;MAP2K3;MARCH2;MARCH8;MARK3;MBOAT2;MFHAS1;MGST3;MINPP1;MKRN1;MOCOS;MOSPD1;MPP1;MXI1;NARF;NCOA4;NEK7;NFE2L1;NNT;NR3C1;NUDT4;OPTN;OSBP2;P4HA2;PC;PDZK1IP1;PGLS;PICALM;PIGQ;PPOX;PPP2R5B;[illegible] |

| |
|---|
| ;PHF5A;PLK1;PLK4;PMS2;PNN;POLA2;POLD1;POLD2;POLD3;POLE;POLE4;POP7;PPM1D;PPP1R8;PRDX4;PRIM2;PRKDC;PRPS1;PSIP1;PSMC3IP;PTTG1;RACGAP1;RAD1;RAD21;RAD50;RAD51AP1;RAD51C;RAN;RANBP1;RBBP7;RFC1;RFC2;RFC3;RPA1;RPA2;RPA3;RRM2;SHMT1;SLBP;SMC1A;SMC3;SMC4;SMC6;SNRPB;SPAG5;SPC24;SPC25;SRSF1;SRSF2;SSRP1;STAG1;STMN1;SUV39H1;SYNCRIP; |
| C1;PPP1R15A;PTGER4;PTGS2;PTPRE;PTX3;RCAN1;REL;RELA;RELB;RHOB;RIPK2;RNF19B;SAT1;SDC4;SERPINB8;SERPINE1;SGK1;SLC16A6;SLC2A3;SLC2A6;SMAD3;SNN;SOCS3;SOD2;SPHK1;SPSB1;SQSTM1;STAT5A;TANK;TAP1;TGIF1;TIPARP;TNC;TNFAIP2;TNFAIP3;TNFAIP8;TNFSF9;TNIP1;TNIP2;TRAF1;TRIB1;TRIP10;TSC22D1;TUBB2A;VEGFA;YRDC; |
| NUMA1;NUP50;NUP98;NUSAP1;ODC1;ODF2;ORC5;ORC6;PAFAH1B1;PAPD7;PBK;PDS5B;PLK1;PLK4;PML;POLA2;POLE;POLQ;PRC1;PRIM2;PRMT5;PRPF4B;PTTG1;PURA;RACGAP1;RAD21;RAD23B;RAD54L;RASAL2;RBL1;RBM14;RPA2;RPS6KA5;SAP30;SFPQ;SLC12A2;SLC38A1;SLC7A1;SLC7A5;SMAD3;SMARCC1;SMC1A;SMC2;SMC4;SNRPD1;SQLE;SRSF1;SRSF10;SRSF2;SS18;STAG1;STIL;ST |
| COLCE2;PDGFRB;PDLIM4;PFN2;PLAUR;PLOD1;PLOD2;PLOD3;PMEPA1;PMP22;PPIB;PRSS2;PTHLH;PTX3;PVR;QSOX1;RHOB;SAT1;SCG2;SDC1;SDC4;SERPINE1;SERPINE2;SERPINH1;SFRP1;SGCB;SGCG;SLC6A8;SNAI2;SNTB1;SPARC;SPOCK1;SPP1;TAGLN;TFPI2;TGFB1;TGFBI;TGFBR3;TGM2;THBS1;THBS2;THY1;TIMP1;TIMP3;TNC;TNFAIP3;TNFRSF11B;TNFRSF12A;TPM1;TPM2;TPM4; |
| LC1;PRDX2;PSMD9;RAD23A;RANBP10;RAP1GAP;RBM38;RBM5;RCL1;RIOK3;RNF123;RNF19A;SDCBP;SEC14L1;SELENBP1;SIDT2;SLC10A3;SLC11A2;SLC22A4;SLC25A37;SLC25A38;SLC2A1;SLC30A1;SLC6A8;SLC6A9;SLC7A11;SMOX;SPTB;SYNJ1;TCEA1;TFDP2;TFRC;TMCC2;TMEM9B;TNRC6B;TNS1;TOP1;TRAK2;TRIM10;TSPAN5;UBAC1;UCP2;UROD;UROS;USP15;VEZF1;XK;XPO7;YPEL5 |
